# Spatial proteomics of human diabetic kidney disease, from health to class III

**DOI:** 10.1101/2023.04.12.534028

**Authors:** Ayano Kondo, Monee McGrady, Dhiraj Nallapothula, Hira Ali, Alexandro E. Trevino, Amy Lam, Ryan Preska, H. Blaize D’Angio, Zhenqin Wu, Lauren Lopez, Harshanna Kaur Badhesha, Chenoa Rochel Vargas, Achyuta Ramesh, Nasim Wiegley, Seung Seok Han, Marc Dall’Era, Kuang-Yu Jen, Aaron T. Mayer, Maryam Afkarian

**Affiliations:** Enable Medicine, Menlo Park, CA, 94025; Division of Nephrology, Department of Medicine, University of California, Davis, CA 95618; University of California, San Diego, CA 92093; Department of Internal Medicine, Seoul National University Hospital, Seoul, South Korea; Department of Urology, University of California-Davis Medical Center, Sacramento, CA 95817, USA; Department of Pathology and Laboratory Medicine, University of California, Davis School of Medicine, Sacramento, CA

**Keywords:** Diabetic kidney disease (DKD), tissue proteomics, multiplex immunofluorescence, spatially resolved proteomics, CODEX

## Abstract

**Aims/Hypothesis:** Diabetic kidney disease (DKD) remains a significant cause of morbidity and mortality in people with diabetes. Though animal models have taught us much about the molecular mechanisms of DKD, translating these findings to human disease requires greater knowledge of the molecular changes caused by diabetes in human kidneys. Establishing this knowledge base requires building carefully curated, reliable, and complete repositories of human kidney tissue, as well as tissue proteomics platforms capable of simultaneous, spatially resolved examination of multiple proteins.

**Methods:** We used the multiplexed immunofluorescence platform CO-Detection by indexing (CODEX) to image and analyze the expression of 21 proteins in 23 tissue sections from 12 individuals with diabetes and healthy kidneys (DM, 5 individuals), DKD classes IIA, and IIB (2 individuals per class), IIA-B intermediate (2 individuals), and III (one individual).

**Results:** Analysis of the 21-plex immunofluorescence images revealed 18 cellular clusters, corresponding to 10 known kidney compartments and cell types, including proximal tubules, distal nephron, podocytes, glomerular endothelial and peritubular capillaries, blood vessels, including endothelial cells and vascular smooth muscle cells, macrophages, cells of the myeloid lineage, broad CD45+ inflammatory cells and the basement membrane. DKD progression was associated with co-localized increase in collagen IV deposition and infiltration of inflammatory cells, as well as loss of native proteins of each nephron segment at variable rates. Compartment-specific cellular changes corroborated this general theme, with compartment-specific variations. Cell type frequency and cell-to-cell adjacency highlighted (statistically) significant increase in inflammatory cells and their adjacency to tubular and αSMA+ cells in DKD kidneys. Finally, DKD progression was marked by substantial regional variability within single tissue sections, as well as variability across patients within the same DKD class. The sizable intra-personal variability in DKD severity impacts pathologic classifications, and the attendant clinical decisions, which are usually based on small tissue biopsies.

**Conclusions/Interpretations:** High-plex immunofluorescence images revealed changes in protein expression corresponding to differences in cellular phenotypic composition and microenvironment structure with DKD progression. This initial dataset demonstrates the combined power of curated human kidney tissues, multiplexed immunofluorescence and powerful analysis tools in revealing pathophysiology of human DKD.

## INTRODUCTION

Diabetic kidney disease (DKD) remains a significant cause of morbidity and mortality in people with diabetes worldwide [1]. Current diagnostic tests are limited, especially for the detection of early disease. Despite recent advances, consistently effective tools for management of advanced disease are likewise lacking. Expanding our diagnostic and therapeutic repository for DKD will require a detailed understanding of the molecular mechanisms underlying disease progression in humans. To date, research on the molecular pathobiology of DKD has fallen into two categories: first, examination of human biofluids to identify putative disease markers; and second, dissection of experimental cellular or animal models of DKD. Data bridging the detailed molecular profiling seen in animal studies and human disease progression has been sparse, impeding the translation of insights gleaned from experimental DKD models to clinical improvements. Bulk and single-cell RNA sequencing (RNA-seq) experiments in human kidneys have begun to fill this gap, allowing researchers to define how gene expression patterns between animal models and human patients relate.

More recently, high-parameter *in situ* molecular profiling technologies (reviewed in [1]), such as multiplexed immunofluorescence and spatial transcriptomics, have made it possible to draw connections between disease progression measured by pathologic review, and molecular and cellular states defined by RNA or protein expression. These spatially resolved platforms hold great promise for hypothesis generation directly in the context of human disease.

Here, we describe the expression of a 21-protein panel in 23 regions of interest (ROIs) from 12 individuals with diabetes and healthy kidneys and those with DKD classes IIA through III. We preferentially targeted proteins (over RNA) to address our resources to areas with lower data density (e.g. high-plexed spatial proteomics in human tissues). This manuscript lays out a framework for analysis of tissue proteomics data in human DKD.

## MATERIALS AND METHODS

### Tissue repository

Kidney tissue resection was performed by UC Davis Urologic Surgery. Tissue procurement was supervised by UC Davis Surgical Pathology, following NCI’s Best Practices for procurement of remnant surgical research tissue. Resected tissues were immediately transported to UC Davis Pathology lab. Tissue preservation and processing was conducted within 60 minutes of receipt by the CAP/CLIA-accredited UC Davis Biorepository/Pathology Lab, which performs the subsequent processing steps per standard clinical protocols. The histotechnologists at the UC Davis Biorepository/Pathology Lab are experienced in generating FFPE blocks, tissue sections, microarrays, routine histology, immunohistochemistry and immunofluorescence. (https://health.ucdavis.edu/cancer/research/sharedresources/biorepository/specimen-services.html). Tissue acquisition and use for research is conducted under the GU001 protocol, which is an IRB-approved protocol allowing UC Davis patients to volunteer their biospecimens (surgical tissue, blood and/or urine) for research studies by the UC Davis GenitoUrinary Research Program, consisting of clinicians, basic science researchers, and designated research support personnel. Each patient signs an IRB approved consent form that allows their biospecimens to be collected and stored for research studies. Collection of tissue and associated data is conducted under regulatory processes (IRB, HIPAA or SRC) of the University of California (UC) Davis Institutional Review Board, governing ethical conduct of research by UC Davis investigators. Tissues also undergo pathological consultation by a UC Davis Pathologist. Investigators are provided histopathologic parameters from the final diagnostic report. Specimens are dispersed and tracked via a rapid and standardized approval and monitoring process. Dr. Afkarian is a participating investigator in GU001 and a long-time collaborator with Dr. Dall’Era, the GU001 principal investigator.

### Tissue preparation and processing

#### Kidney tissue storage

Kidney tissues from the 12 individuals for the current study were stored in liquid nitrogen for 3-10 years until use (**Table 1**).

**Table 1.** Characteristics of tissue donors.

| DKD Class | Patient | Age | Sex | Race | DM type | HTN | OR Date | PMH | Medications | BP | Wt (lb) | SCr (mg/dL) | Urine prot |
| --- | --- | --- | --- | --- | --- | --- | --- | --- | --- | --- | --- | --- | --- |
| DM with healthy kidneys | 1 | 69 | M | NHW | 2 | Y | 2012 | Former smoke, Hyperlipidemia, obstructive sleep apnea, chronic lymphocytic leukemia | Cyclobenzaprine, Ezetimibe, Fenofibrate, Metformin, Multivitamin capsule, Omeprazole, Sitagliptin, Vitamin D, hydrocodone/acetaminophen | 160/85 | 270 | 1.3 | Neg (UA) |
|  | 2 | 69 | F | NHW | 2 | Y | 2014 | Endometrial cancer, severe obesity, cardiomyopathy, asthma, pulmonary embolus, arthritis | Acetaminophen, Albuterol, Allopurinol, **Benazepril**, Calcium carbonate/vitamin D3, Carvediol, Dalteparin, Docusate, Gabapentin, Letrozole, Metformin, Hydromorphone, Ondansetron, Letrozole, Simvastatin, Triamterene/ Hydrochlorothiazide, Warfarin | 91//51 | 274 | 1.2 | Trace (UA) |
|  | 3 | 58 | M | NHW | 2 | Y | 2017 | Esophageal motility disorder, adenomatous colonic polyps, rectal bleeding, diarrhea, arthritis of knee, chronic leg and low back pain, hyperlipidemia, insomnia, gout, lower urinary tract symptoms, diaphragmatic hernia | Allopurinol, Aspirin, Atorvastatin, Cholecalciferol, Diazepam, Fentanyl, Gabapentin, Hylan G-F 20, Insulin Lispro, **Lisinopril**-hydrochlorothiazide, Multivitamin capsule, Oxycodone, Pantoprazole, Terazosin, Verapamil | 128/69 | 275 | 0.9-1 | Neg (UA) |
|  | 4 | 66 | M | NHW | 2 | Y | 2010 | Liver problems, nephrolithiasis, heart murmur, spinal cord injury, testicular cancer | Bystolic, Amlodipine, multivitamins | 141/81 |  | 0.87 |  |
|  | 5 | 48 | F | NHW | 2 | N | 2013 | Dyslipidemia, psoriasis, DOE | Citalopram, Klonopin, Lortab, Gemfibrozil, Lamotrigine, Pravastatin, Aleve | 132/80 |  | 0.9 |  |
| DKDIIA | 6 | 73 | F | NHW | 2 | Y | 2011 | Dyslipidemia, smoker, COPD | albuterol, hydrochlorothiazide, hyzaar, isosorbide mononitrate, levothyroxine, nexium, betapace, signulair, zetia, cymbalta, ferrous sulfate, fexofenadine, flonase, **fosinopril**, glyburide, potassium chloride, mag-oxide, klor-con m10, abilify, metformin, pristig (desvenlafaxine), meloxicam welchol, hydrocodone-acetaminophen, phenazopyridine, ciprofloxacin. | 145/59 |  | 1.1 |  |
|  | 7 | 61 | F | NHW | 2 | Y | 2014 | CAD, severe obesity, Obstructive sleep apnea on CPAP, hiatial hernia, gout, anemia, DDD, osteoarthritis | albuterol, Allopurinol, Amlodipine, Ascorbic acid, Aspirin, **Benazepril**, Budesonide/Formoterol, Calcium, Ferrous Sulfate, Glucosamine/Chondroitin sulf A, Glyburide, Hydrochlorothiazide, Metformin, Multivitamin, Niacin, Omega-3 FA Fish Oil, Omeprazole, Oxybutynin, Simvastatin | 126/49 | 251 | 0.9-1.1 | Neg (UA) |
| DKDIIA-B | 8 | 75 | M | UK | 2 | Y | 2016 | CAD, hyperlipidemia, chronic kidney disease, severe obesity, hyperkalemia, hyperlipidemia | Amlodipine, Chlorthalidone, Diclofenac, Docusate, Ferrous sulfate, hydrocodone/acetaminophen, metoprolol tartrate, minoxidil, simvastatin | 182/51 | 217 | 1.73 | 30 (UA) |
|  | 9 | 68 | F | UK | 2 | Y | 2014 | Morbid obesity, CAD, h/o breast cancer | Insulin demetir, Liraglutide, Anastrozole, Atenolol, Calcium, Furosemide, Levothyroxine, **Losartan**, PPI, Rosuvastatin, Iron | 140/68 | 308 | 1.05 |  |
| DKDIIB | 10 | 49 | F | UK | 2 | Y | 2014 | Osteopenia, morbid obesity, gastroparesis | ASA, **Enalapril**, Fluoxetine, Folate, Gaba, Insulin aspart, glargine, **Lisinopril**, Mg, Metformin, Methocarbamol, Metoclopramide, Modafinil, MV, Omega3, Propranolol, Simvastatin, Trazodone | 126/69 | 297 | 0.99 |  |
|  | 11 | 75 | F | NHW | 2 | Y | 2017 | Former smoker 0.25 ppd for 5 years (stopped smoking 17 years ago). Hyperlipidemia, anorexia, thrombectomy | Amlodipine, aspirin 81 mg, metformin, simvastatin, valsartan, docusate, hydrocodone 5mg/ acetaminiphen 325 mg, indomethacin | 145/71 | 174 | 1.2 | Trace (UA) |
| DKDIII | 12 | 50 | M | Hisp | 2 | Y | 2019 | Dyslipidemia, CHF, CKD | Insulin degludec, **spironolactone**, **losartan**, semaglutide, atorvastatin, bumetanide | 122/81 | 183 | 3.1 |  |

Abbreviations- DM: diabetes mellitus; NHW: non-Hispanic white; UK: unknown; UA: urinalysis; DKD: diabetic kidney disease; COPD: chronic obstructive pulmonary disease; CAD: coronary artery disease; DDD: degenerative disc disease; CHF: congestive heart failure; CKD: chronic kidney disease.

#### Fixation and embedding

Kidney tissues were fixed in 20-fold (v:v) 10% neutral buffered formalin (Fisher Scientific, MA) for 48 hours at room temperature. Formalin-fixed tissue was paraffin-embedded and sectioned following UC Davis Anatomic Pathology protocols. Briefly, a processed tissue cassette block was placed into the embedding chamber with orientation confirmed by inspection prior to embedding. The mold was filled with molten paraffin. Using a forceps, the tissue was carefully moved from the cassette to the paraffin in the mold. The tissue was held in place while the paraffin was solidifying in the mold, located on a cold plate. Blocks were removed from the molds when completely cold. Embedding stations, forceps, tampers and molds were kept clean at all times to prevent cross-contamination.

#### Sectioning the blocks

The paraffin blocks were trimmed until the tissue was seen in full-face. Tissue block was then placed in wet ice for 20 minutes at room temperature and then subsequently placed on dry ice for 2-5 minutes, if needed. The tissue block was clamped into the microtome chuck with safety brakes unlocked. The hand wheel was rotated to cut through the tissue, cutting 5-micron sections. Tissue sections were floated onto hot water in a bath and then moved onto the coverslips using forceps, wiping the excess water with a Kim wipe. Coverslips were placed upright into a draining rack and allowed to air dry for 15-20 minutes.

#### PAS staining

The tissue section from the face of the block underwent PAS staining, using the STAT lab (McKinney, TX) staining kit and protocol (https://www.statlab.com/pdfs/ifu/KTPAS.pdf).

#### Generation of the kidney tissue microarray (TMA)

Twenty-three cores from 12 tissue blocks (12 individuals with diabetes with kidneys ranging from healthy to DKDIIB and a DKDIII biopsy core) were used to generate the TMA (**Supplementary Figure 1**). The PAS-stained sections of each block were viewed to mark the areas for inclusion in the TMA. A hollow needle was used to excise tissue cores of 0.5mm diameter from the designated regions in the tissue blocks, as identified by microscopic examination of the PAS-stained section from the face of each paraffin-embedded tissue block. These tissue cores were then inserted in a recipient paraffin block in a precisely spaced, array pattern within a 15 x 15 mm area. The cores were positioned on the TMA such that those from donors with diabetes, DKDIIA, DKDIIA-B, DKDIIB and DKDIII alternated so that DKD classes were dispersed. This was done to prevent segregation of one or other DKD class by location. Two positions (marked ‘Blank’) were kept vacant to establish TMA orientation. The TMA block was sectioned using the protocol described below (‘Sectioning the blocks’). Adjacent tissue sections were used for multiplex immunofluorescence and confirmatory PAS staining.

### Tissue characterization

Kidney tissues were characterized by (a) evaluation of existing donor clinical data by a nephrologist experienced in DKD care (M. A.) and (b) histopathologic examination by an expert renal pathologist (K-U. J.), who assigned the DKD class to each section by microscopic examination of the PAS-stained sections, following the Tervaert classification.[2] The tissue microarray utilized for the current study included 23 tissue sections from 12 individuals with diabetes. Ten kidney tissue sections came from 5 diabetic individuals with healthy kidneys (DM); twelve tissue sections were from two individuals each with DKD IIA, IIA-B and IIB; and one tissue biopsy came from one individual with DKDIII.

### Tissue staining

#### Antibody conjugation

A 21-plex antibody staining panel was developed for this study (**Supplementary Table 1**). To create CODEX-compatible antibodies, DNA barcodes were chemically conjugated to serum-free antibodies. For the panel, five conjugated antibodies were purchased from Akoya Biosciences, MA, and sixteen were conjugated at the staining site (Enable Medicine Laboratory) following a protocol provided by Akoya Biosciences (https://www.akoyabio.com/wp-content/uploads/2021/01/CODEX-User-Manual.pdf). Briefly, unconjugated antibodies were partially reduced and incubated with Barcode Solution for 2 hours at room temperature. After the 2-hour incubation, conjugates were purified by four rounds of centrifugation and buffer exchange with Purification Solution (Akoya) through 50 kDa molecular weight cutoff filters (MilliporeSigma). After conjugation, successful conjugation and purification was verified by denaturing gradient gel electrophoresis (BioRad, TGX Mini-Protean 4-15%) and Coomassie staining (Thermo Fisher). The conjugated antibodies were also validated *in situ* by performing limited CODEX experiments on control tissue sections prior to the experiments described in this manuscript.

#### Antibody Titer Optimization

For antibodies purchased from Akoya Biosciences, titers were optimized on FFPE human tonsil sections, and concentrations adjusted for optimal use at 1:100 or 1:200 dilutions. For antibodies conjugated in the Enable laboratory, a standard dilution of 1:50 was validated on FFPE human tonsil sections. Each antibody was assessed for signal specificity by tissue type, cell type, subcellular localization, as well as qualitatively assessed for high signal to noise ratio relative to background.

#### Staining and Acquisition

CODEX staining and imaging was performed on kidney tissue samples using the 21-plex panel described above. Staining was performed according to manufacturer guidelines (https://www.akoyabio.com/wp-content/uploads/2021/01/CODEX-User-Manual.pdf). Briefly, the tissue sections were heated at 55℃ for 25 minutes, deparaffinized, and re-hydrated. Subsequently, antigen retrieval was performed by heating slides in a pressure cooker at 95° C for 20 minutes in the presence of TRIS-EDTA buffer. pH = 9.0. The tissue sections were then washed and equilibrated to room temperature in Staining Buffer (Akoya Biosciences, MA), and then incubated with the complete antibody cocktail overnight at 4° C. The tissue sections were then fixed and stored at 4℃° C in Storage Buffer (Akoya Biosciences, MA) until image acquisition. Images from the stained tissue sections were acquired on the CODEX instrument (currently called Phenocycler, Akoya Biosciences, MA), attached to a Keyence BZ-X810 fluorescent microscope (Keyence, IL). The CODEX Instrument Manager software was used to specify regions of focus and acquire the images at 20x objective lens resolution with the use of the BZ-X800 Viewer software. The following filter cubes were used to capture the various biomarkers: DAPI, Atto-750, Atto-550, and Cy5. The images taken were stored as ome.tiff image files.

#### Image Processing

The raw fluorescent TIFF images were processed by performing deconvolution across all dimensions, then performing alignment and stitching of image tiles. The images were processed for background subtraction using the extended depth of field algorithm. Finally, staining quality and image processing were assessed on the Enable Medicine platform before proceeding to data analysis.

### CODEX Data Analysis

#### Cell segmentation

The first step, cell segmentation, was performed using the DeepCell deep-learning algorithm, which identified individual nuclei in the images using the DAPI channel. The nuclear masks were then dilated to approximate cell surface. Cell segmentation was performed on pre-processed DAPI images using the DeepCell model [4]. After identifying individual nuclei, nuclear segmentation masks were expanded by constrained stochastic dilation to approximate cell membranes. The dilation algorithm runs as follows:

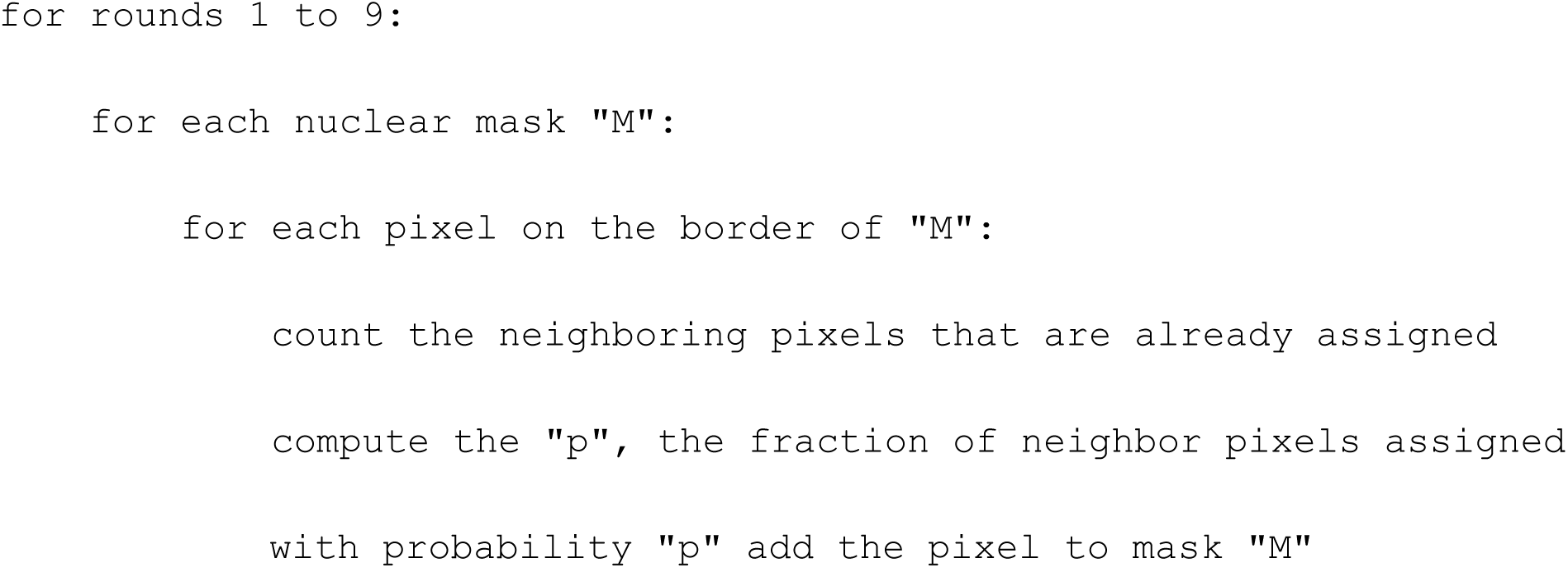

#### Construction of a protein expression matrix

To quantify the protein expression patterns, single cell- and tissue-level information were extracted using both supervised and unsupervised methods. Following cell segmentation, a matrix of single-cell protein expression was generated for each sample. For each image channel, the mean pixel signal intensity inside each segmented cell was calculated, as a surrogate for protein expression levels. The result was unique 21-protein expression profiles for all segmented cells in the study.

#### Identification of distinct cell populations by unsupervised clustering

Data analysis and visualization was performed in R 4.1.2 unless otherwise noted. A custom R package (https://docs.enablemedicine.com/spatialmap) was used to facilitate data normalization, cell phenotyping, and data visualization.

Within each CODEX sample, single cell expression was normalized for each protein by first scaling values by five times the 20th percentile, then applying an inverse hyperbolic sine transformation (“base::asinh” – following the R convention of “<PACKAGE_NAME>::<FUNCTION_NAME>” notation) to the scaled values. Next, normalized values were z-scaled across markers, then across cells.

For cell clustering and cell neighborhood analysis, expression data from the 23 patient samples were combined. To cluster cells, dimensionality reduction was first performed on scaled expression values using principal component analysis with 20 components (“stats::prcomp”). Next, a k-nearest neighbor graph was constructed to build a similarity network between cells in principal component space (“dbscan::kNN”, k = 30). Finally, cells were clustered using the Leiden graph clustering algorithm (“igraph::cluster_leiden”, cluster_resolution = 1.0). To label clusters, a heatmap showing the average normalized marker expression in each cluster was plotted. Clusters were annotated using their average expression to identify cell types, and these annotations were validated by manual inspection of multiplexed images. Rows and columns of the expression heatmap matrix were hierarchically clustered.

Clustered expression data were visualized using Voronoi diagrams, which were constructed on the spatial coordinates of segmented cell centroids. The data were further visualized using the uniform manifold approximation and projection (UMAP) algorithm applied to the similarity network between cells. To confirm a lack of egregious batch effects, UMAP plots were colored by cell cluster, sample, and other covariates.

#### Biomarker Expression Masks for Compartment-Based Analysis

Six kidney compartments were outlined for ROI analysis using the Enable Medicine Visualizer (**Supplementary Figure 2**). These compartments included glomeruli, blood vessels, distal tubules, all-tubules, Collagen IV^+^ areas, and the interstitium. Two compartments (glomeruli and blood vessels) were outlined manually and these annotations were used to create binary masks for these two compartments. Three compartments were isolated by creation of binary masks based on thresholding the expression of relevant proteins (MUC1 for distal nephron, CXCR3 for all-tubules and collagen IV for basement membrane) and morphological fill operations. For example, a binary mask for the distal nephron in each sample was created by feeding in the MUC1 channel from the CODEX image to a custom image processing tool, thresholding pixel values on MUC1 expression, and gap filling and expanding the mask to identify contiguous regions of interest. For downstream analysis, the proximal tubule compartment was determined by subtracting the distal nephron compartment from the all-tubules compartment. The interstitial compartment was marked by a mask that was created by subtracting the masks of the other five compartments from a mask of the entire tissue region. The masks were then used to label the cells in each sample by compartment for further analysis.

#### Cell proportion analysis

Cell proportions were determined across all cell type clusters and summarized by sample, disease stage, and tissue compartment; values were also summarized across disease stage for individual cell types. These data were visualized using stacked barplots and box plots (“ggplot2::geom_bar” with ‘pos =”fill”’ and “ggplot2::geom_boxplot”, respectively).

#### Projection of tissue compartments

To investigate how cell and protein expression dynamics in different tissue compartments contribute to overall kidney pathology across DKD, we projected tissue subregions, specified by a pathologist, as additional samples into a multidimensional scaling plot, along with the full samples. For each sample or sub-region, expression of each marker was summarized by taking the median normalized expression value across cells. The multidimensional scaling was performed on the resulting region/subregion by marker matrix using “limma::plotMDS” with default arguments. In this projection, fibrotic regions localized more closely with diseased samples, as defined by clinical tests, indicating that the presence of fibrotic regions is a mechanistic correlate of clinical stage.

#### Compartment protein expression

Protein *e*xpression in glomeruli was calculated by summing signals over the corresponding regions in the images as follows: For each single channel image, lower and upper intensity thresholds were determined based on the histogram of pixel intensities. Next, the image was min-max normalized according to these thresholds. The normalization process was performed to ensure that images from different regions/acquisitions would have the same dynamic range.

## RESULTS

### Patient characteristics

The majority of the 12 donors were non-Hispanic white (**Table 1**). All groups, except DKDIIB and DKDIII, included kidney tissue from both women and men. All donors had type 2 diabetes; hypertension was present in 4 out of 5 donors with healthy kidneys and all donors with DKD. The mean age of the donors with healthy kidneys (DM), those with DKDIIA, IIA-B and IIB was 62, 71, 72, and 62, respectively; the age of the single donor with DKDIII was 50. The nephrectomies were performed between 2010 and 2019, and the tissues from participants with DM, DKDIIA, IIA-B, IIB, and III were stored in liquid nitrogen for an average of 9, 10, 7, 7, and 3 years, respectively. Information on glomerular filtration rate (GFR) as measured by CKD-EPI[3] and urine protein (urinalysis) was obtained prior to nephrectomy, when possible.

### Using multiplex immunofluorescence to visualize spatial distribution of 21 proteins in human kidneys

We performed multiplexed immunofluorescence to examine tissue expression of 21 proteins (**Supplementary Table 1**) in 23 sections/ROIs from 12 individuals (**Figure 1, Supplementary Figure 1, Supplementary Figures 3 and 4**), using CO-Detection by indexing (CODEX). These proteins were selected because of their expression in human kidney tissue based on prior data (**Supplementary Figure 3**) and relevance to kidney function or DKD pathophysiology. For example, CD45, CD68, and CD11b staining identifies the immune cells infiltrating the kidney (**Figure 1B, 1D** boxes 3 and 6); nestin and CCR6 distinguish glomerular podocytes and endothelial cells (**Figure 1C, 1D** box 1); αSMA, CXCR3, MUC1 and collagen IV identify blood vessels, all tubules, the distal nephron and the basement membrane, respectively (**Figure 1D**, boxes 2, 4 and 5). **Supplementary Figure 5** displays expression of these proteins in all 23 ROIs. **Supplementary Table 1** outlines the utilized clones, commercial antibody sources and dilutions as well as the CODEX barcode and fluorophore used for each monoclonal antibody.

**Figure 1.**
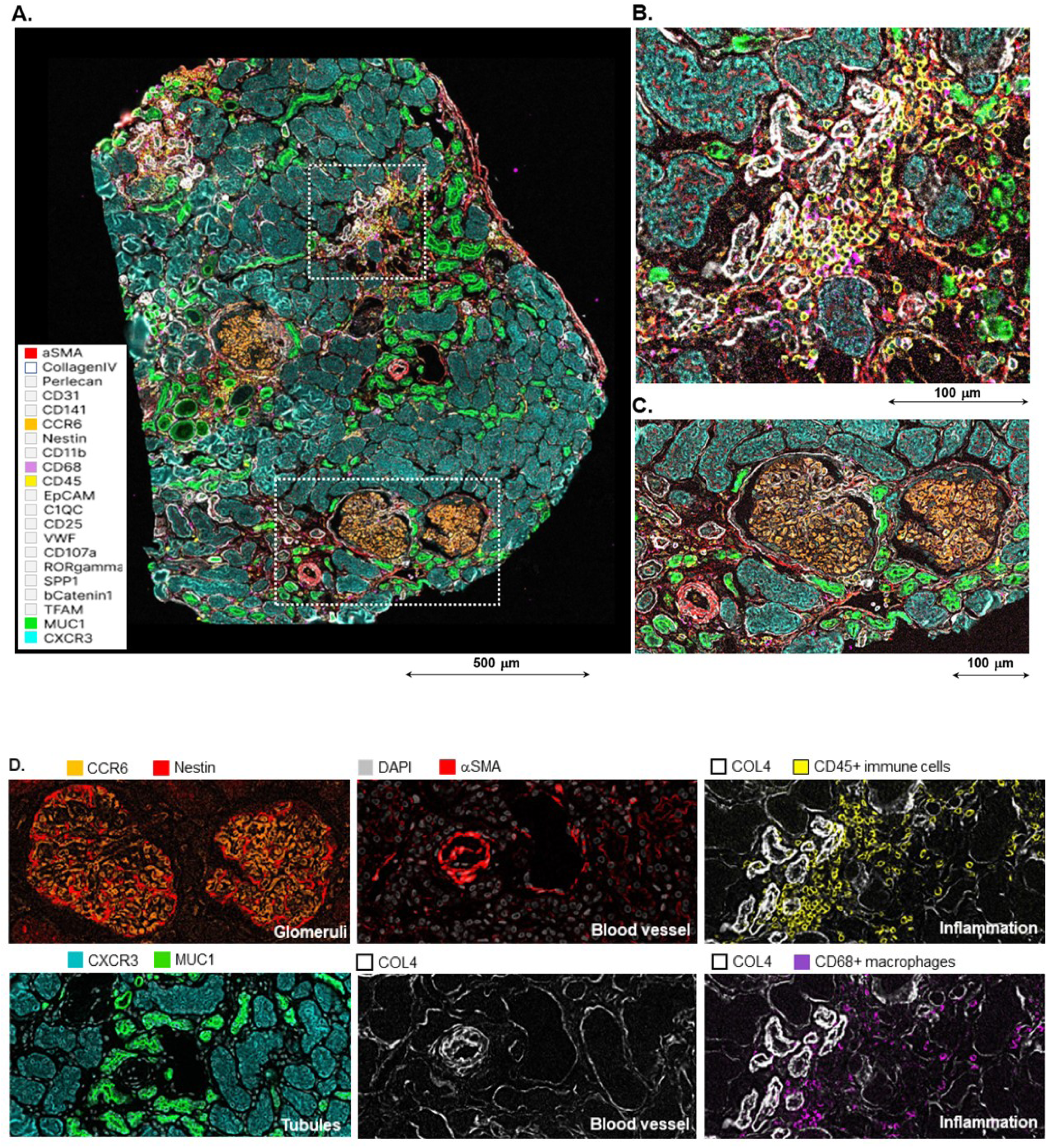
Representative multiplexed immunofluorescence image showing protein expression in a kidney section with DKD. **A.** A representative human kidney cortical tissue section with DKDIIB, displaying the basement membrane (Collagen IV, white), macrophages (CD68, purple), broad immune cells (CD45, yellow), smooth muscle and interstitial cells (αSMA, red), glomerular endothelial and peritubular capillary cells (CCR6, orange), all tubules (CXCR3, teal) and distal nephron (MUC1, bright green). Two zoomed-in regions from this sample show areas of increased immune infiltration and fibrosis (section **B**), a blood vessel and glomerular compartments (section **C**). **D**. Nestin (red), (CCR6) (orange), αSMA (red), CD45 (yellow), CXCR3 (teal), MUC1 (green), Collagen IV (white) and CD68 (purple) highlight podocytes, glomerular endothelial cells, blood vessels, the inflammatory infiltrate, all tubules, the distal nephron, basement membrane, and macrophages, respectively (section **D** panels from top left to bottom right).

### Summary of data on expression of the targeted proteins in human kidneys

Expression of the 21 targeted proteins in our human kidney samples (**Supplementary Figure 4**) was compared with prior protein expression data, including the human protein atlas (https://www.proteinatlas.org/) (**Supplementary Figure 3**, summarized in **Table 2**). For 16 of the 21 proteins, protein expression in our samples was consistent with reported expression in prior literature and/or the Human Protein Atlas. For the other four, including nestin, CXCR3 (CD183) and osteopontin (SPP1), our data diverged from some or all prior reports, as summarized in **Table 2**.

**Table 2.** Summary of data on expression of the 21 targeted proteins in human kidneys.

| Protein | Prior data | Human Protein Atlas | Our data | Conclusion for DM |
| --- | --- | --- | --- | --- |
| **αSMA**<br>**(ACTA2)** | Blood vessels, interstitium.(26808537, 22460095) | 3/3 mAbs interstitial, BVs<br>1/3 some proximal tubular stain | Blood vessels, interstitium | Blood vessels, interstitium |
| **C1QC** | **C1Q, not C1QC**<br>Interstitial/immune cells [13]<br>None.[28-30]<br>Glomeruli, arterioles [14] | All tubules with variations | Proximal tubules >> Distal<br>Nephron 2 | Proximal tubules >> Distal<br>Nephron 2 |
| **CCR6**<br>**(CD196)** | Glom Endothelial Cells (GEC) and PeriTubular<br>Capillary endothelial cells (PTC)[31] | GEC, PTC<br>tubules in some individuals with<br>some of the Abs | GEC, PTC | GEC, PTC |
| **CD11b** | Monocytes and granulocytes[32, 33] | Individual tissue resident cells<br>low level expression in PT | Monocytes and granulocytes | Monocytes and granulocytes |
| **CD31** | Glomeruli, venules, arteries > capillaries[34] | Glomeruli, venules, capillaries | Glomeruli, venules, arteries,<br>capillaries | Glomeruli, venules, arteries<br>capillaries |
| **CD45** | All hematopoietic cells except erythrocytes [35] | All hematopoietic cells except<br>erythrocytes | All hematopoietic cells except<br>erythrocytes | All hematopoietic cells except<br>erythrocytes |
| **CD68** | Monocytes[13] and macrophages [36] | Not detected (individual cells<br>visible, however.) | Monocytes and granulocytes | Monocytes and granulocytes |
| **Collagen**<br>**IV** | Mesangial matrix (4A1), Bowman's capsule (4A1,<br>4A5), GBM (4A3, 4A5), tubular BM (4A1)[37] | (4A1, 2) Mesangial matrix,<br>Bowman's capsule, tubular BM<br>(4A3) Glomerular mesangial<br>cells, proximal tubules | Mesangial matrix, GBM,<br>Bowman's capsule, tubular BM | Mesangial matrix, GBM,<br>Bowman's capsule, tubular<br>BM |
| **CTNBB1** | Proximal tubules, thin + thick ascending limbs, distal<br>convoluted tubules, collecting duct.[38] | Tubular epithelia | All tubular epithelia | All tubular epithelia |
| **CXCR3**<br>**(CD183)** | mRNA from micro-dissected human proximal tubules<br>(and rat IHC) [11], none [5-7], vascular smooth muscle<br>cells,[8, 10] endothelial cells,[8] afferent arteriole, low<br>level glomerular cells,[10] proximal, distal convoluted,<br>low grade in gloms (Thermo Fisher 26756-1-AP)<br>Collecting tubule in medulla (Thermo Fisher PA5-23679) | One mAb used- no expression<br>detected | Proximal and distal convoluted<br>tubules, collecting ducts |  |
| **EpCAM** | Distal convoluted tubule ~ Loop of Henle > Collecting<br>duct.[39]<br>Distal convoluted tubule > Collecting duct.[40]<br>Collecting duct > Distal convoluted tubule,[41]<br>cortical collecting duct[42] | Collecting duct >>> Distal<br>tubules | Collecting duct >>> Distal<br>tubules | Collecting duct >>> Distal<br>tubules |
| **HSPG2**<br>**(perlecan)** | Blood vessels (arterioles,[43] arteries [44]), tubular<br>BM,[43, 44] GBM.[44] | Arteries, arterioles, glomerular<br>vascular pole (3/4)<br>low-medium ubiquitous<br>expression (3/4) | Blood vessels, tubular BM, weak<br>glomerular BM and mesangial<br>matrix | Blood vessels, tubular BM,<br>weak glomerular BM |
| **LAMP1**<br>**(CD107a)** | Ubiquitous [45] | Ubiquitous | Ubiquitous | Ubiquitous |
| **mtTFA** | All tubules, none in gloms [46] | All tubules | All tubules | All tubules |
| **MUC1 (CD227)** | Distal convoluted tubule, collecting duct.[47, 48] | Distal convoluted tubule, collecting duct | Distal convoluted tubule, collecting duct | Distal convoluted tubule, collecting duct |
| **Nestin** | Glomerular podocytes.[25, 26] | Glomerular podocytes, peritubular capillaries, endothelium, low level in proximal tubules (1/4 Abs) | Podocytes, proximal tubules, small medullary vessels | Podocytes, small medullary vessels |
| **RORγ** | RORγ isoform 1 (RORγ1) mRNA is expressed in kidneys.[17-19] This mAb recognizes both isoforms but is commercially defined as identifying RORγt because that is the only isoform in immune cells. No data on protein. | No expression | Ubiquitous nuclear expression | Ubiquitous nuclear expression |
| **SPP1 (OPN)** | Thick ascending LH,[21] distal convoluted tubules,[21-23] collecting duct,[22, 23] some proximal tubules.[22] | Proximal and distal convoluted tubules | Proximal and distal convoluted tubules, collecting ducts |  |
| **TM (CD141)** | Weak & segmental in glomerular vessels, stronger in peritubular capillaries.[49] Vascular pole of glomerulus, reduced in glomeruli of people with DM.[12] Glomerular vascular pole, peritubular capillaries.[50] | Peritubular capillaries, glomerular vascular pole, blood vessels | Peritubular capillaries, blood vessels | Peritubular capillaries, blood vessels. Staining in glom vascular pole reduced due to DM in our samples. |
| **vWF** | Venules, arterioles, capillaries.[34] Argument about whether there's vWF in gloms (yes [51]; no [34]) | Reported as not detected but is visible in venules and arterioles | High intensity staining in venules, arterioles and arteries. Low grade staining in parenchyma | Venules, arterioles, capillaries |

### Identification of the known cell types and tissue compartments in the kidney

Cell types were assigned to all the segmented cells in the dataset using unsupervised clustering from the 21-protein expression profiles, leading to identification of 18 clusters. The clusters were classified into ten distinct cell populations based on the bulk expression profiles of each cluster (**Figure 2A-B**). The ten identified cell populations were proximal tubules (CXCR3^++^/MUC1^-^), distal nephron (CXCR3^+^/MUC1^+^), endothelial cells (CCR6^lo^/CD31^+^), glomerular endothelial cells (GEC) or peritubular capillaries (PTC) (CCR6^+^/CD31^+^), αSMA-expressing cells, including the vascular smooth muscle cells (VSMC) and interstitital αSMA-expressing cells, nestin^+^ cells (primarily podocytes of the glomeruli), macrophages (CD68^+^), myeloid lineage cells (CD11b^+^), and other immune cells (CD45^+^/CD68^-^/CD11b^-^). Cells located in the basement membrane (Collagen IV^+^/HSPG^+^) were categorized as “basement membrane cells”. Cells exhibiting low expression of all biomarkers in the panel were categorized as “low-expressing” cells.

**Figure 2.**
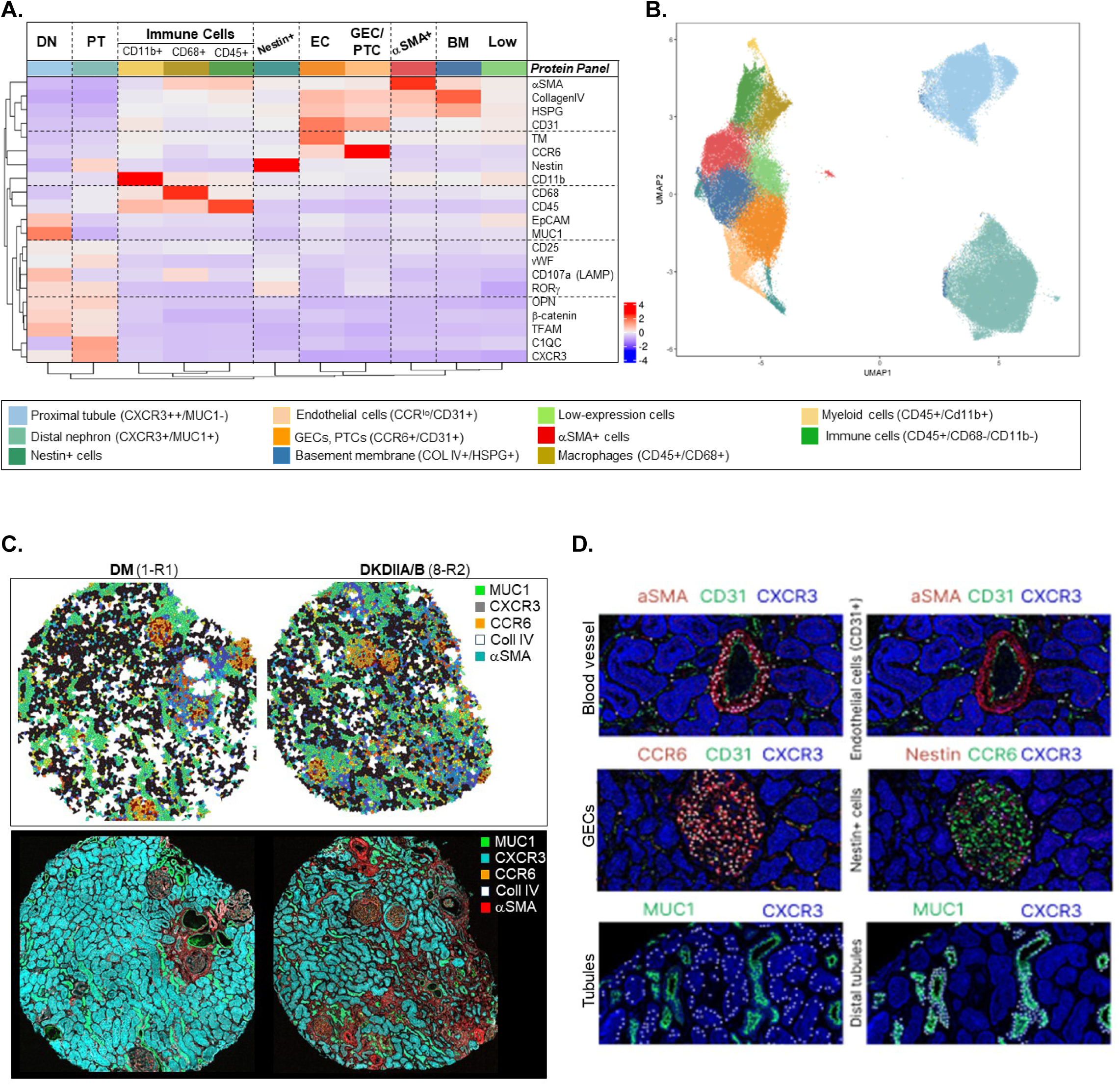
Classification of kidney cell types and tissue compartments. **A.** The heatmap of protein expression by phenotype shows unique expression profiles for the 10 cell populations identified in this study, as well as the low-expressing cells. **B.** A UMAP representations of all cells in the study, colored by cell type. **C.** A Voronoi representation of cortical sections from a healthy kidney sample and one from a donor with DKDIIA-B, colored by cell type (top panels), compared with expression of compartment-identifying proteins in the same tissue sections in the bottom panels. **D.** Cell types identified from unsupervised clustering were validated by overlaying cell annotations (white dots) with cell type-specific marker protein channels.

Because the initial cell classification results were generated algorithmically and without supervision, we validated the cell classification results using several methods. <u>First</u>, we verified that each identified cell population expressed the expected combination of proteins (**Figure 2A-B**). <u>Secondly</u>, we determined that minimal batch effects were observed from unsupervised clustering (**Supplementary Figure 6**). <u>Thirdly</u>, we confirmed that the Voronoi representations of the samples showed the expected spatial localization of the identified cell populations (**Figure 2C**). Finally, we confirmed the results by overlaying cell annotations with CODEX image channels of cell-type specific biomarkers (**Figure 2D**).

### Global and compartment-wise changes in protein expression from diabetes to DKD

After cell classification, we quantitatively compared the cellular composition of the kidney samples across disease progression (**Figure 3A**). The transition from DM to progressive DKD was associated with an increase in the fraction of all inflammatory cells (macrophages, inflammatory cells of the myeloid lineage, and other broad CD45^+^ cells). Furthermore, we observed a decrease in the abundance of proximal tubules from DM to DKD. (**Figure 3A**). In addition to global cell type identification by unsupervised clustering in entire sections, we generated tissue masks for six distinct tissue compartments in the kidney (glomeruli, blood vessels, distal tubules, all-tubules, Collagen IV^+^ areas, and the interstitium), either by manual outlining, using expression of compartment-identifying proteins or both (**Supplementary Figure 2**). Cell frequencies were then examined in these six tissue compartments, defined by the tissue masks (**Figure 3B**). In progression from DM to DKD3, the glomerular compartment showed a decline in podocytes (nestin^+^ cells) and CCR6^+^/CD31^+^ glomerular endothelial cells, and an increase in ɑSMA^+^ cells and cells localized to the Collagen IV^+^/HSPG^+^ basement membrane. The proximal tubule compartment showed a decrease in the actual proximal tubule cells and an increase in CD45^+^ immune cells and macrophages, while the cellular composition of the distal nephron compartment, marked by MUC1 expression, was grossly unchanged. Blood vessels showed a reduction in ɑSMA^+^ cells and a mild reduction in CCR6^lo^/CD31^+^ endothelial cells, and the interstitium had a subtle drop in CCR6^lo^/CD31^+^ endothelial cells and an increase in CD45^+^ immune cells and low-expressing interstitial cells. Finally, the basement membrane showed an increase in Collagen IV^+^/HSPG^+^ regions and inflammatory cells (**Figure 3B**).

**Figure 3.**
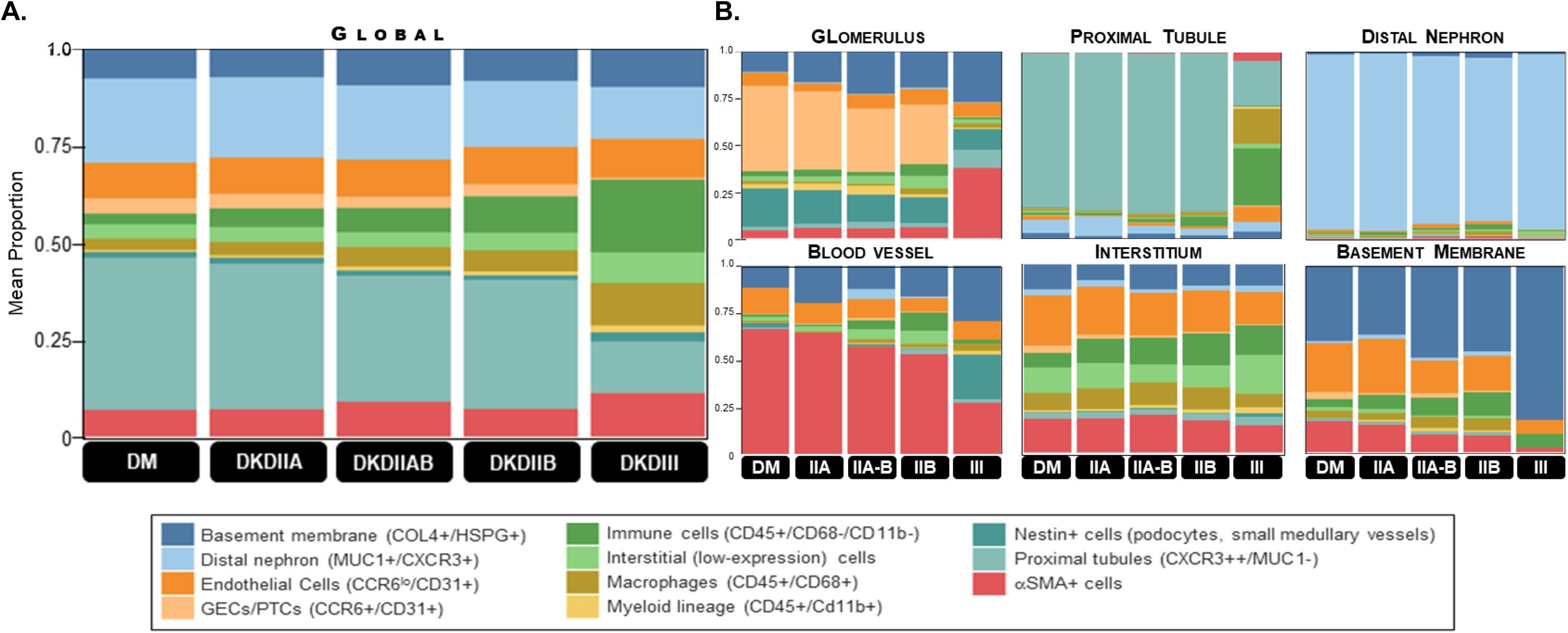
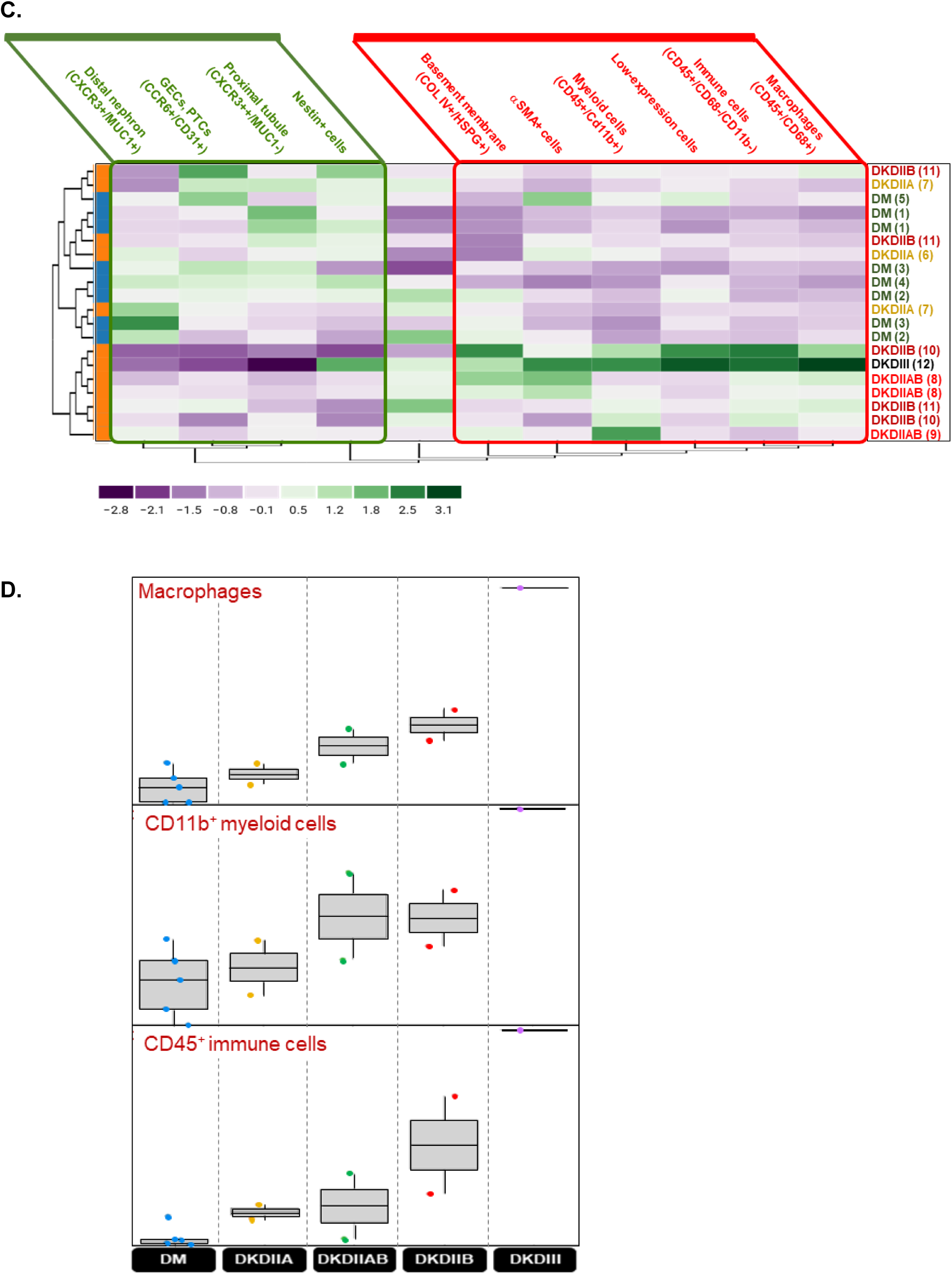
Global and compartment-wise changes in cell type proportions and protein expression from diabetes to DKD. **A.** Bar graph of cell frequency shows global change in fraction of cell types when the data is aggregated by DKD class. **B.** Compartment-wise change in distribution of cellular composition in glomeruli, proximal tubules, distal nephron, blood vessels, interstitium and basement membrane, with DKD progression. **C.** Hierarchical clustering based on cell type frequency in the cortical tissue sections, where the columns represent cell types identified from unsupervised clustering, and the rows represent individual tissue sections. Cell type frequencies are represented as z-scores per column (cell type) and values are normalized by column. Heatmap shows coarse segregation of cortical sections from people with DM and DKD, but with overlap between classes. **D.** Boxplots of cell type frequency, grouped by DKD class, show that increase in immune cells is continuous from DM to DKDIII. Each dot represents a single individual within each DKD class.

Hierarchical clustering based on cell frequencies in the cortical tissue sections reiterated the higher abundance of immune cells, αSMA^+^ cells and collagen IV^+^/HSPG^+^ cells in DKD tissues *vs.* DM (red-bordered box), while sections from healthy kidneys had more cells from proximal tubules and the distal nephron, as well as glomerular nestin^+^ and CCR6^+^ cells (green-bordered box) (**Figure 3C**). The increase in inflammatory cells was continuous from DM to DKDIII (**Figure 3D**).

### Visual examination of protein changes from health to DKDIII in diabetes

Visual examination reiterated the data obtained from bioinformatic analysis in above sections, i.e. the rise in fibrosis and inflammatory infiltrate and a progressive reduction in expression of proteins marking tubular and glomerular compartments with DKD progression. Also notable was a difference in the trajectory of change in expression of the segment marker proteins. For example, MUC1 maintained expression from health to DKDIII class while CCR6 and CXCR3 lost or reduced expression earlier. In addition, co-staining for collagen IV, CD45 and CD68 showed that (a) inflammatory infiltrate and fibrosis occurred in the same areas of the examined tissues and (b) the extent of tissue injury, as shown by increase in collagen IV deposition and CD45^+^ or CD68^+^ cell infiltration, was regional and patchy (**Figure 4**).

**Figure 4.**
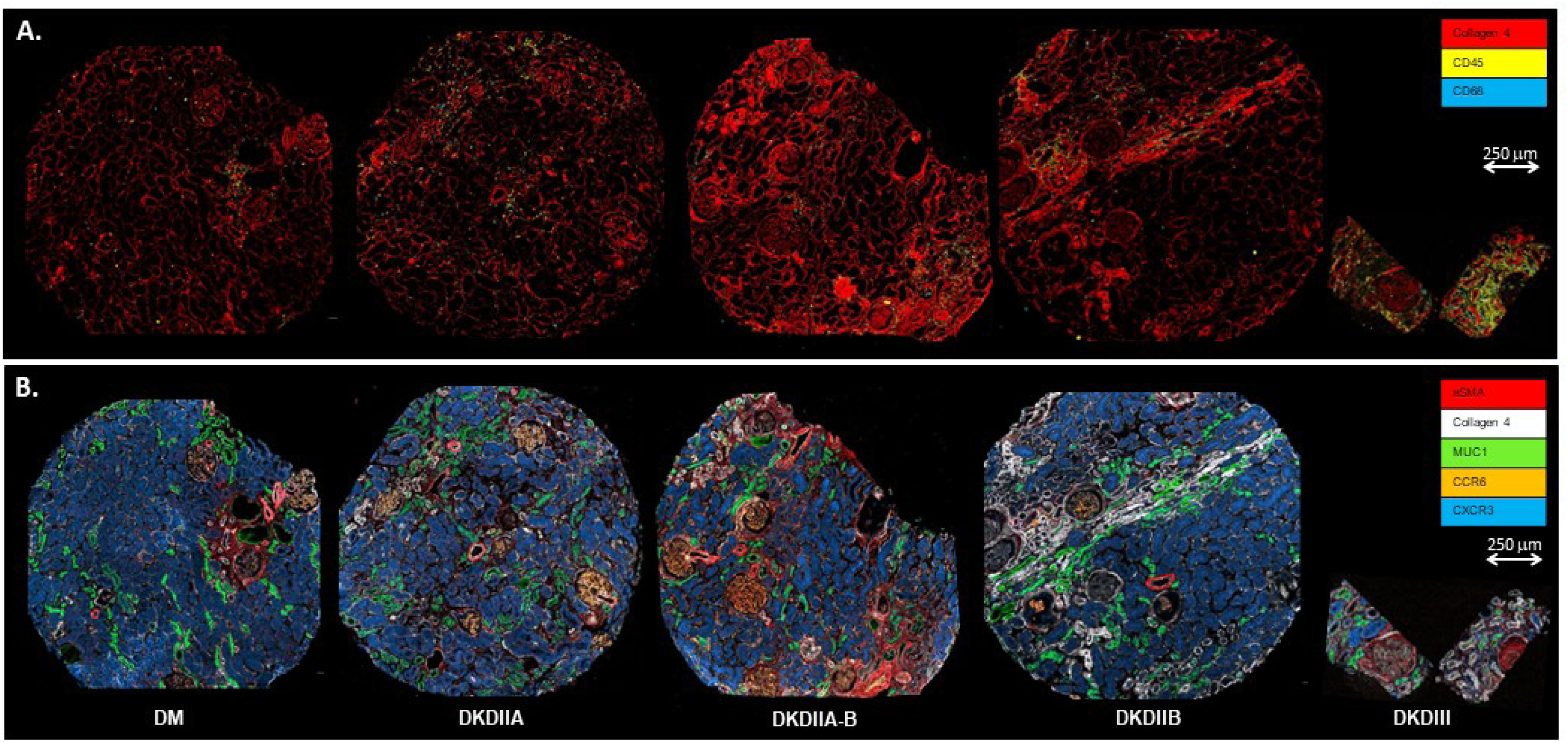
Representative multiplex immunofluorescence images showing protein expression across the spectrum from healthy kidneys to progressive DKD. A. Staining for basement membrane (collagen IV, red), broad inflammatory cells (CD45^+^, yellow) and infiltrating macrophages (CD68^+^, blue) highlights two points: first, disease progression, manifested by basement membrane (collagen IV) thickening is patchy and heterogeneous. Secondly, inflammatory infiltrate, including macrophages, coincide with areas of greater collagen IV deposition. B. Third point emerging is that compartment-identifying proteins (CXCR3-teal, CCR6-orange, MUC1-green, αSMA-red and collagen IV-white) change expression at different DKD classes.

### DKD is patchy: quantifying section- and patient-level heterogeneity in cellular composition and protein expression

Our initial observations suggested that immune infiltration and development of fibrosis are regional and progress in patches (**Figure 1, Figure 4**). Therefore, we aimed to map the correlation between histopathological features/sub-regions of a single tissue to the overall DKD class assigned to the individual. Sub-regions of variable DKD severity were manually outlined in a tissue section from a patient with DKDIIB (Sample ID: 10) by a pathologist. Based on histopathologic features of DKD severity, these areas were labeled as healthy, moderately fibrotic, or severely fibrotic (**Figure 5A**). The sub-regions were then projected as individual specimens onto the principal component space defined by the 23 tissue sections with DKDIIA to III. Healthy, moderately fibrotic and severely fibrotic sub-regions from one individual tissue section localized with tissue sections from healthy kidneys (DM), those with intermediate DKD (IIA to IIAB) and those with severe DKD. Thus, a single tissue section from one individual displayed high pathologic variability, running the gamut from DM to DKDIII (**Figure 4B**). To further assess intra- and inter-individual variability in protein expression, we examined variability in expression of the CCR6 protein, which marks the glomerular compartment. This compartment was selected because (1) glomeruli were manually outlined, and (2) glomerular sclerosis is a known feature of DKD progression. Normalized CCR6 expression in glomerular endothelial cells was calculated in outlined glomeruli from two individuals with DKDIIB (individuals 10 and 11). We observed substantial variability in normalized glomerular CCR6 expression both within an individual participant (e.g. in individual 10, **Figure 5C-D**) and between the two individuals with the DKDIIB (individuals 10 and 11, **Figures 5C-D**).

**Figure 5.**
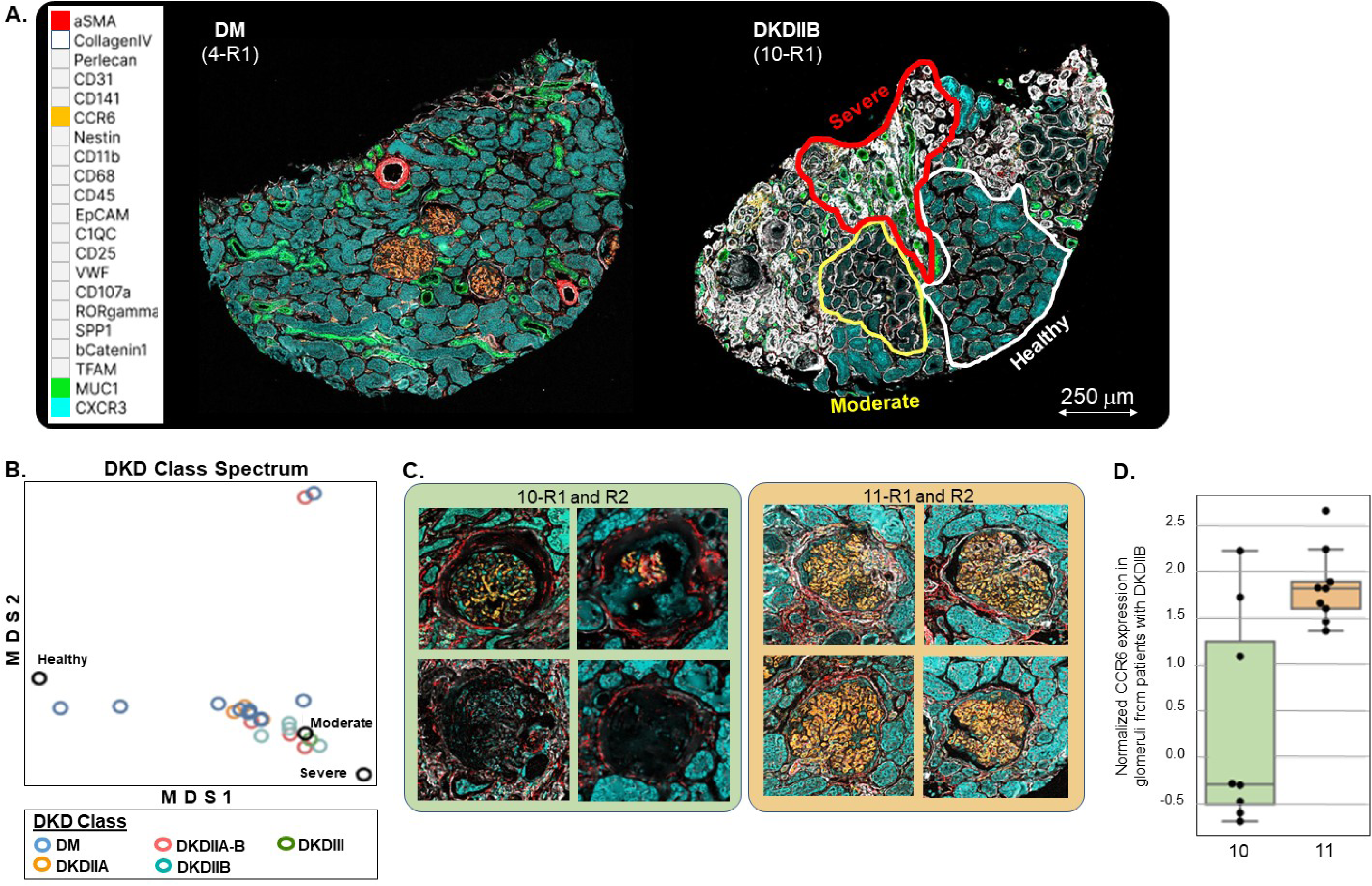
DKD is heterogeneous. **A.** Representative multiplex immunofluorescence images show DKD heterogeneity (sample 10-R1, top right) in alteration of protein expression, compared with normal tissue (sample 4-R1, top left). Regions with variable DKD severity are manually outlined. **B.** Multidimensional scaling (MDS) plot of 20 cortical sections (omitting the three medullary sections), colored by disease class, as well as the three manually outlined sub-regions. **C.** Representative images of manually outlined glomeruli in two DKDIIB sections from patients 10 and 11. **D.** A boxplot comparing normalized CCR6 expression in glomeruli from the two sections. Each dot represents CCR6 expression in a single outlined glomerulus.

### Hierarchical clustering of individual cortical tissue sections based on adjacency between cell types

Heatmap dendrograms of cell-cell adjacency segregated healthy and DKD kidney tissues. Consistent with changes in cell types observed in prior sections, DKD tissues had a greater increase in cell-cell adjacencies involving cells of the inflammatory infiltrate, αSMA^+^ cells and the basement membrane (red-bordered box). Also consistently, healthy tissues showed more frequent cell-cell adjacencies involving cells of the proximal and distal nephron, as well as the nestin^+^ and GEC cells (green-bordered box) (**Figure 6A**). The increase in proximity between CD45^+^ immune cells and cells of the distal nephron, proximal tubules, as well as cells expressing αSMA, was robust to Bonferroni adjustment for multiple testing (**Figure 6B**).

**Figure 6.**
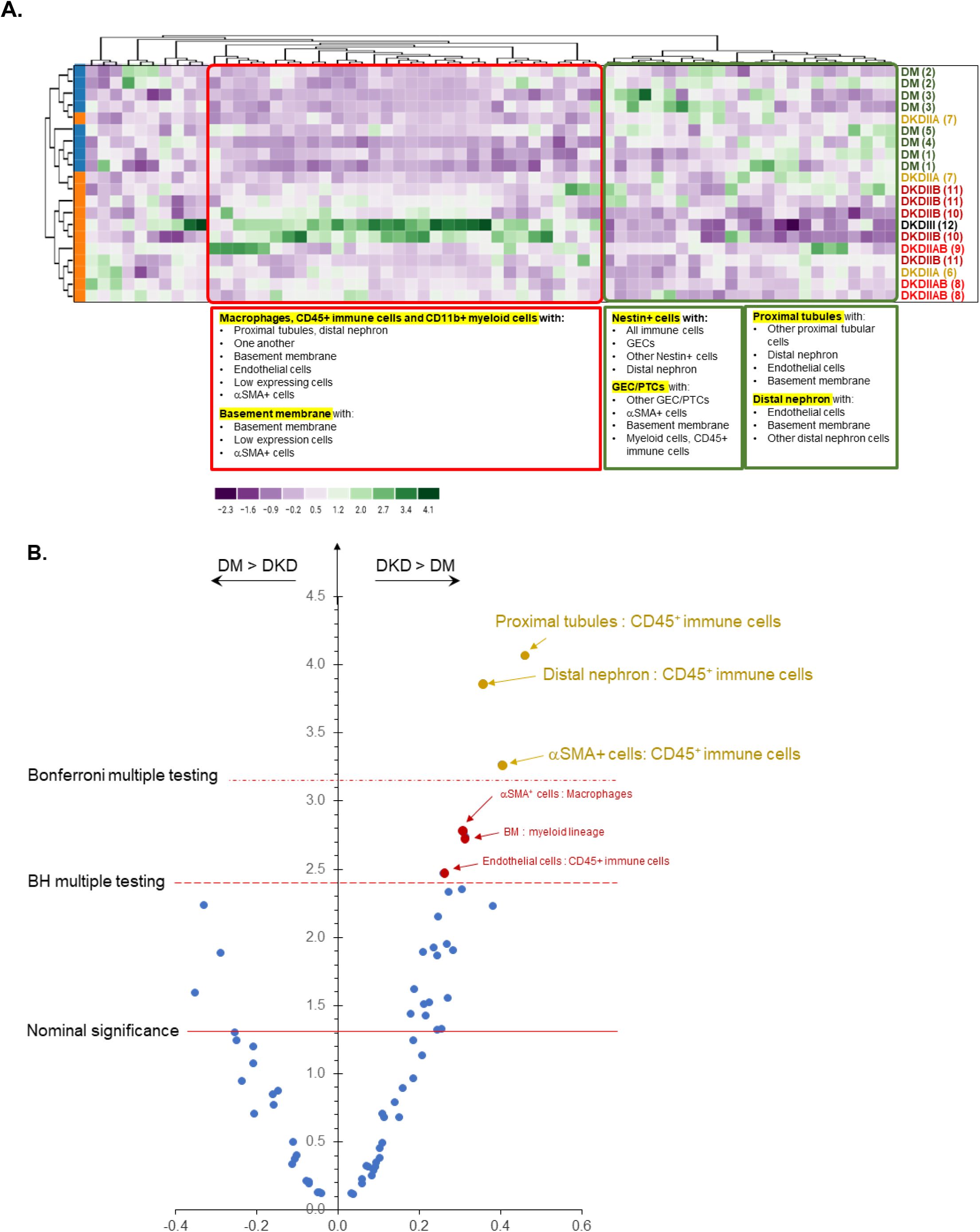
Hierarchical clustering of cortical tissue sections based on cell-cell adjacency between cell types. **A.** Heatmap dendrogram of cell-cell adjacency segregates healthy kidneys from DKD. Cell-cell adjacencies also show significant inter-individual heterogeneity within each DKD class. Values are normalized per column using z-scores; data is clustered by rows. **B.** A Volcano plot of cell adjacencies shows enrichment of proximity between immune cells and tubular (proximal, distal) and αSMA^+^ cells in DKD, compared with healthy kidneys, after Bonferroni adjustment for multiple testing.

## Discussion

We analyze expression of 21 proteins in human kidneys from people with diabetes and healthy kidneys (10 sections from 5 individuals), to those with DKD classes IIA to III (13 sections from 7 individuals). Expression of these 21 proteins, quantified by a multiplex immunofluorescence platform,[4] identified 10 functionally significant kidney compartments or cell types. In people with diabetes, DKD progression was associated with co-localized increase in the inflammatory infiltrate and collagen IV deposition, as well as reduction of the native proteins marking proximal tubules and glomeruli. The expression of proteins marking different nephron segments followed distinct trajectories: CXCR3 and CCR6 reduction occurred by DKDIII, while MUC1 expression persisted through DKDIII. Importantly, DKD severity was patchy, introducing sizable intra- and inter-individual variability in molecular pathology of disease progression in the kidney tissue and highlighting the limitations of kidney biopsies in providing a whole-kidney assessment of the extent of injury. Clustering based on cell type or cell-to-cell proximities confirmed the increases in inflammatory cells in DKD and in addition showed statistically significant proximity between inflammatory cells and proximal and distal tubular cells, as well as those expressing αSMA, despite adjustment for multiple testing.

*<u>First</u>*, this report presents data on expression of 21 proteins in human kidneys, which is compared with prior literature and annotated with pre-analytic variables. To draw firm conclusions on expression of the 21 targeted proteins, our data was compared to existing literature on expression of each protein (including the human protein atlas), and critically analyzed (**Supplementary Figure 3**, summarized in **Table 2**). For 16 of 21 proteins, data reported here was supported by prior literature and allowed determination of protein expression in human kidneys from people with diabetes, with or without DKD. We included all available data on potential sources of pre-analytic variability (theoretical sources outlined in **Supplementary Table 2**) for our samples, including donor characteristics (demographic, clinical, etc.), tissue source and processing and utilized antibodies, to enable similar comparisons by other authors (**Table 1**). Our observed protein expression patterns were consistent with prior data for all but four proteins (CXCR3, RORγ, osteopontin and nestin), where we either varied from prior data or were met with absence of a consensus on expression of the specific protein in human kidneys.

We observed CXCR3 expression in all tubules. In contrast, Human Protein Atlas found no CXCR3 protein in human kidney sections and others reported staining patterns ranging from no expression[5–7] to expression in the inflammatory infiltrate,[8, 9] vascular smooth muscle and endothelial cells,[8, 10] afferent arterioles,[10] or all tubules (data from commercial antibody sources) (**Table 2**). CXCR3 has three known isoforms,[5] (**Supplementary Figure 3**, pages 32-33) whose tissue-specific expression may explain the observed differences. Interestingly, mRNAs for both CXCR3-A and -B isoforms are expressed in micro-dissected human proximal tubules[11] and whole human kidneys.[5] However, the isoform-specificity of the antibodies utilized in the previous studies did not clearly explain the observed discrepancies (**Supplementary Table 3**). One key source of difference between our data and prior reports is that all our samples were from people with diabetes while the donor diabetes status for the other data sources is unknown. Diabetes has been shown to alter kidney protein expression, even without kidney disease. For example, thrombomodulin [12] is lower and C9, CFD,[13] C1Q, C5b-9 and C4d[14] proteins are higher in kidneys of people with diabetes and no DKD. Consistently, CXCR3 protein was expressed in proximal tubules and parietal epithelial cells (PEC) of glomeruli in diabetic db/db mice[15] and upregulated in tubular cells in mouse models of tubular injury.[16] As such, diabetes may induce diffuse CXCR3 protein expression in human kidney tubules, a hypothesis requiring examination in subsequent studies.

We observed ubiquitous RORγ protein expression, localizing to the nuclei in all cells of the kidney, regardless of DKD presence or class. RORγ has two isoforms which differ in their first exon (**Supplementary Figure 3**, pages 80, 82): RORγ1 uses exon 1γ and 2γ, which are replaced by exon 1γt in isoform2 (RORγt). RORγ1 is widely expressed in many tissues, including liver, kidney, lung, muscle, heart, brain and adipose tissue (**Supplementary Figure 3**, pages 81-83); RORγt (RORγ2) is primarily expressed in immature thymocytes and some immune cells.[17] RORγ mRNA is expressed in human kidneys.[18–20] However, to our knowledge, RORγ protein expression was not previously reported in human kidneys.

In our sample set, osteopontin (SPP1) protein was present in proximal tubules as well as the distal nephron. The Human Protein Atlas showed a similar expression, using a polyclonal rabbit antibody verified by orthogonal immunohistochemistry, enhanced immunocytochemical staining and western blotting (**Supplementary Figure 3**, pages 88-89). Prior reports have documented osteopontin expression in thick ascending loop of Henle,[21] distal convoluted tubules,[21–23] collecting duct [22, 23] and some proximal tubules.[22] Interestingly, implantation biopsies also showed perinuclear osteopontin staining in the slightly damaged proximal tubular cells (in addition to pronounced apical expression in the distal tubular cells), suggesting that osteopontin expression in proximal tubules may be induced with injury.[24] However, the effect of variation in pre-analytic sources, e.g. diabetes or hypertension, on osteopontin expression require further study.

Nestin was previously reported to be expressed in podocytes only.[25–27] We also find strong expression in podocytes but in addition observe expression in proximal tubules. Interestingly, the same pattern was observed with one (out of 4) antibodies utilized in the human protein atlas (**Table 2 and Supplementary Figure 3**, page 73, CAB005889). As with the other three proteins discussed above, the effect of pre-analytic variables (e.g. diabetes or hypertension) on nestin expression remains unknown and in need of further studies.

*<u>Second</u>*, this report shows the ability of a relatively small (21-protein) panel to segregate kidneys into 18 clusters, corresponding to several of the known and functionally important kidney compartments and cell types: podocytes, GEC/PTCs, proximal tubules, distal nephron, basement membrane, blood vessels (endothelium, smooth muscle cells) and the inflammatory infiltrate including macrophages, myeloid cells and other inflammatory cells. Each cluster is identifiable by its specific expression profile of the 21-protein panel, including expression of a marker proteins (e.g. CCR6 identifying GECs or PTCs or MUC1 identifying the distal nephron). Within each cluster/compartment, quantification of all 21 proteins allows characterization of disease-associated alterations in these proteins from health to advanced DKD. Comparing cell types in cortical sections from DM to DKD reiterated the themes of concomitant rise in fibrosis and inflammatory infiltrate and reduction in proteins marking proximal tubular and glomerular compartments. Compartment-wise examination of these changes showed reduction in proteins specific to the compartment, but with trajectories distinct to each protein and compartment. For example, CCR6^+^ and nestin^+^ cells were reduced in glomeruli and CXCR3^+^ cells in proximal tubules while distal tubules showed minimal change in MUC1 staining. In addition, examining protein alterations by compartment revealed compartment-specific nuances in DKD-associated global changes in protein expression. For example, with DKD progression, αSMA^+^ cells were increased in glomeruli but reduced in blood vessels, suggesting an alteration not only in quantity but site of αSMA expression, away from its normal cell types in vessel walls and towards abnormal sites such as glomeruli. This examination also highlighted the importance of manual outlining of compartments based on histologic appearances because the compartment- or cell-type marker proteins may no longer be expressed with DKD progression or their pattern of expression may change.

*<u>Third</u>*, we observe striking heterogeneity and patchiness in DKD severity, both within a single tissue section and between individuals with the same DKD class. This marked intra- and inter-individual variability was evaluated by visual examination and quantified by bioinformatic analyses. While visual examination is clearly limiting for the substantial number of sections needed to overcome biologic variability in human tissues, it provided the impetus and direction for quantitative assessment of this variability using the more powerful and far-reaching bioinformatic tools. Importantly, juxtaposition of an average kidney biopsy core with the research tissue sections obtained from partial nephrectomy (**Figure 4**) stresses the scale and impact of this limitation in tissue sampling on our clinical assessment of DKD severity in entire kidneys.

*<u>Fourth</u>*, Cell frequency and cell-cell adjacency clustering studies provide other examples of bioinformatic analyses of spatial proteomics which can shed light on disease mechanisms. As proof of principle, we present an example of these analyses highlighting the increase in inflammatory cells and their proximity to the cells of the nephron in DKD tissues vs. healthy kidneys from people with diabetes. Interestingly, heterogeneity was again in evidence in both cell frequency and cell-cell adjacency analyses, displaying significant variability between tissue sections in the same DKD class. These variations may be due to the difference between the true DKD class in the multiplex immunofluorescence-stained section *vs.* the section used for pathologic staging (which may be a few microns apart) or intra-individual heterogeneity in protein expression within the same tissue section, or the inter-individual variability within each DKD class.

*<u>Finally</u>*, we describe a collaborative human kidney proteomics pipeline, composed of a human kidney biorepository, a multiplex immunofluorescence platform and combined expertise in clinical nephrology, renal pathology, histotechnology/histopathology, epidemiology, biostatistics and bioinformatics required for generation of reliable proteomics data from human kidney. The tissue repository includes the required controls (e.g. kidney tissue from non-diabetic donors with or without kidney disease and donors with diabetes but without kidney disease) as well as tissues with DKDI to III, classified by an expert renal pathologist. Tissues from donors with DKD classes IV and V are not included because severe fibrosis and scarring in these classes significantly reduces their data content for changes in protein expression. DKDI was not included in this initial effort because identifying these tissues required electron microscopy data and preliminary data was necessary to support a search for molecular changes of DKD in this class.

This study adds to the novel and exciting body of work using tissue proteomics to expand our existing molecular companion to human DKD pathologic classification. Not surprisingly, this initial study has generated several new questions, to be addressed in subsequent studies. For example, while tissues from people with diabetes are the appropriate controls for those with DKD, a full understanding of changes in protein expression requires inclusion of kidney tissues from people without diabetes or DKD, as well as those with nondiabetic CKD. For example, diabetes alone (without kidney disease) may cause changes in protein expression which can only be identified using the tissue from donors without diabetes and kidney disease. Addition of samples from a variety of patients with CKD of non-diabetic etiology is also planned in future studies to ascertain if the observed molecular changes are specific to DKD, or general to CKD. In addition, performing protein staining in the same section as pathologic DKD classification helps pair protein expression and tissue pathology more closely by reducing the inherent variation in DKD class from section to section of the same tissue. In addition, it may be worthwhile to include DKD class I tissue samples, which would require electron microscopy-based ascertainment of DKD class to (a) avoid misclassification of DKDI as normal kidney and (b) detect changes in protein expression which occur prior to class IIA. Finally, as with any other study in human participants, strength is in numbers, of individuals, tissue sections per individual and proteins assessed. As such, access to a high-volume pipeline of available tissues and a flexible, expandable multiplex immunofluorescence platform, as presented here, is of paramount importance.

In conclusion, we present data on tissue protein expression of 21 proteins in 23 tissue sections from five individuals with diabetes and healthy kidneys and seven individuals with DKDIIA to III. This work adds to, and augments, current efforts targeting greater understanding of the changes in protein expression and cell composition as human kidneys progress from health to DKD in people with diabetes.

## Acknowledgements

Specimens and services were provided by the GU001 protocol and the UC Davis Pathology Biorepository which is jointly funded by the UC Davis Comprehensive Cancer Center Support Grant (CCSG), awarded by the National Cancer Institute (NCI P30CA093373), and the UC Davis Department of Pathology and Laboratory Medicine. The authors owe a debt of gratitude to Irmgard Feldman, Ramona Clarke, Allison Gail Proffitt and Romina Sacchi for their solid knowledge base, peerless expertise and work ethics, meticulous and productive work and incredible collaborative spirit.

## Contributions

AK performed the bulk of the data analyses and contributed to writing the manuscript and generating figures. MM contributed to data analyses, generating figures, and writing the methods section of the manuscript. RP performed the bulk of the multiplex immunoflourescence experiments. HA, AET, and AL contributed to writing the manuscript. ZW contributed to data analysis and writing the methods section of the manuscript. BD’A contributed to the biomarker panel optimization and design. DN, HKB, CRV and AR contributed to the image analysis. HKB, CRV, LL and AR contributed to the data generation and literature review. AR contributed to the literature review and image analysis. NW and SSH contributed to study idea and image analysis. K-YJ reviewed all tissues, determined DKD classes, contributed to the idea and study design and critically reviewed the data. ATM contributed to the idea and study design, directed the performance of studies and bioinformatic analysis of the data, and generated figures. MA is primarily responsible for the study idea, experimental design, selection of tissues and regions of interest for generation of the tissue microarray, generation of figures, writing of the manuscript and performing some of the data analyses. All authors participated in the critical review of the data and the manuscript.

This work was supported by the Richard A. and Nora Eccles Foundation award (A20-0111) and the Richard A. and Nora Eccles Harrison Endowed Chair in Diabetes Research (M.A.).

