## Supplementary data for "Spatial proteomics of human diabetic kidney disease, from health to class III"

### Supplementary Figures and Tables

**Supplementary Table 1. Antibody clones, vendors, fluorophores and dilutions**

| Protein | Short/alternative Names | Uniprot Accession | Barcode | Fluorophore | Clone | Vendor | Dilution |
|---|---|---|---|---|---|---|---|
| Aortic smooth muscle actin | αSMA (ACTA2) | P62736 | BX030 | Cy5 | EPR5368 | Abcam | 1:50 |
| Beta-Catenin | CTNNB (CTNNB1) | P35222 | BX020 | ATTO550 | 12F7 | Akoya Biosciences | 1:200 |
| C-C chemokine receptor type 6 | CCR6 (CD196) | P51684 | BX043 | AF750 | EPR22259 | Abcam | 1:50 |
| Collagen IV alpha chain | CO4A1, CO4A2, CO4A3, CO4A4, CO4A5, CO4A6 | P02462, P08572, Q01955, P53420, P29400, Q14031 | BX023 | ATTO550 | EPR20966 | Abcam | 1:50 |
| Complement C1q subcomponent subunit C | C1QC | P02747 | BX003 | Cy5 | EPR2984Y | Abcam | 1:50 |
| Cyclin-dependent kinase 11B | CD11b | | | | | | |
| C-X-C chemokine receptor type 3 | CD183 (CXCR3) | P49682 | BX042 | Cy5 | G025H7 | BioLegend | 1:50 |
| Epithelial cell adhesion molecule | Ep-CAM (CD326) | P16422 | BX010 | Cy5 | EPR20532-225 | Abcam | 1:50 |
| Lysosome-associated membrane glycoprotein 1 | LAMP1 (CD107a) | P11279 | BX006 | Cy5 | H4A3 | Akoya Biosciences | 1:200 |
| Macrosialin | Gp110 (CD68) | P34810 | BX015 | Cy5 | KP1 | Akoya Biosciences | 1:200 |
| Mucin-1 | MUC-1 (CD227) | P15941 | BX004 | AF750 | HMPV | BD Biosciences | 1:100 |
| Nestin | NES | P48681 | BX034 | AF750 | SP103 | Abcam | 1:50 |
| Nuclear receptor ROR-gamma | RORγT (RORC) | P51449 | BX027 | Cy5 | 6F3.1 | EMD Millipore | 1:50 |
| Osteopontin (Secreted phosphoprotein 1) | OPN (SPP1) | P10451 | BX052 | ATTO550 | 7C5H12 | Abcam | 1:50 |
| Perlecan (Basement membrane-specific heparan sulfate proteoglycan core protein) | HSPG | P98160 | BX017 | ATTO550 | 5D7-2E4 | BD Biosciences | 1:50 |
| Platelet endothelial cell adhesion molecule | PECAM-1 (CD31) | P16284 | BX001 | AF750 | EP3095 | Akoya Biosciences | 1:100 |
| Receptor-type tyrosine-protein phosphatase C | PTPRC (CD45) | P08575 | BX021 | Cy5 | D9M8I | Cell Signaling Technology | 1:50 |
| Thrombomodulin | TM (CD141) | P07204 | BX013 | AF750 | E7Y9P | Cell Signaling Technology | 1:50 |
| Transcription factor A, mitochondrial | mtTFA (TFAM, TCF6) | Q00059 | BX029 | ATTO550 | 18G102B2E11 | Akoya Biosciences | 1:200 |
| von Willebrand factor | vWF | P04275 | BX045 | Cy5 | C-12 | Santa Cruz Biotechnology | 1:100 |

**Supplementary Table 2. Causes of pre-analytic variation in kidney protein expression**

| Cause | | Control |
|---|---|---|
| **Individual variables** | 1. Demographic- age, race, sex<br>2. Clinical- diabetes, hypertension, immunosuppression, other medications, smoking<br>3. Other- habits (tobacco, alcohol, drug use), medications, etc. | 1. Examining protein expression patterns in a large number of tissues.<br>2. Statistical adjustment of protein quantities for individual variables and/or stratified analysis. |
| **Tissue** | 1. Autopsy vs. biopsy or resection<br>2. Frozen (OCT) vs. FFPE<br>3. Antigen retrieval method | 1. Uniform tissue acquisition, processing, preservation and staining<br>2. Comparing expression to uniform processing protocol when data obtained otherwise |
| **Antibody (monoclonal)** | 1. Epitope present on >1 isoform<br>2. Epitope specific to one isoform but kidney isoform specificity is unknown, varies between animal and human or is extrapolated from expression in other human tissues. | 1. Determining protein isoform expression in human kidneys<br>2. Determining isoform specificity of utilized antibodies |

#### Supplementary Table 3. Potential sources of preclinical variation for CXCR3 kidney protein expression

| Individual variables | | Tissue | | Antibody | | | Isoforms | | | | Reference |
|---|---|---|---|---|---|---|---|---|---|---|---|
| Demographic | Clinical | Source & Preparation | Antigen Retrieval | Clone | | Immunogen | A | B | Alt | Staining | |
| Table 1 (current study) | | Biopsy and nephrectomy, FFPE | HIER pH6 | G025H7 | Monoclonal mouse IgG1k | Human CXCR3 (?isoform) transfectant | ? | ? | ? | Prox tubules, distal tubules | Current report |
| Female (30%)<br>Age 37 + 25<br>Race/ethnicity unknown | Not available | Nephrectomy FFPE | HIER pH6 | HPA045942 | pAb | QVSDHQVLNDAEVAALLENFSSSYDY GENESDSCCTSPPCPQDFSLNFDRAF | Likely all 3 isoforms (first Q only in B) | | | None | HPA (22688270, 25613900) |
| Female (27%)<br>Age 52 + 15<br>Race/ethnicity unknown | Kidney allograft donor (6) and recipients (37) | Allograft biopsy FFPE | HIER pH6 (microwave) | 1C6 (9466968, 9064356) | Monoclonal mouse IgG1 | MVLEVSDHQVLNDAEVAALLENFSSS YDYGENESD | X | ? | X | None | 16421159 |
| Unknown (n=3) | Unknown | Nephrectomy FFPE | HIER pH6 | | | MVLEVSDHQVLNDAEVAALLENFSSS YDYGENESD | X | ? | X | None | 15857922 |
| Unknown (n=18) | Lupus Nephritis (no control) | Biopsy FFPE | HIER pH6 | | | MVLEVSDHQVLNDAEVAALLENFSSS YDYGENESD | X | ? | X | Inflammatory infiltrate (LN) | 19116922 |
| Unknown (n=5) | "Normal" | Nephrectomy PFA-fixed | None | 49801.111* | Monoclonal mouse IgG1 | Human CXCR3-A (aa 1-368) transfected NSO mouse myeloma cells. | X | X | ? | vascular smooth muscle cells, endothelial cells in normal kidneys, inflammatory infiltrate in GN | 11134180 |
| F/M breakdown unknown<br>Age 35-54<br>Race/ethnicity unknown | "Normal" (n=4) | Nephrectomy PFA-fixed | None | | | Human CXCR3-A (aa 1-368) transfected NSO mouse myeloma cells. | X | X | ? | vascular smooth muscle cells, afferent arteriole, low level glomerular cells | 10589690 |
| Unknown | Unknown | Nephrectomy PFA-fixed | None | PL1 | IgG1 | CXCR3-B only (private Ab): MELRKYGPGRLAGTVIGGAAQSKSQT KSDSITKEFLPGLYTAPSSPFPPSQ | | X | | None | 12782716 |
| Unknown | Unknown | Unknown FFPE | Unknown | 26756-1-AP | Polyclonal rabbit IgG | MVLEVSDHQVLNDAEVAALLENFSSS YDYGENESDSCCTSPPCPQDFSLNFDR | X | ?X | X | Proximal, distal convoluted, low grade in gloms | Thermo Fisher |
| Unknown | Unknown | Unknown FFPE | Unknown | PA5-23679 | Polyclonal rabbit IgG | CXCR3-A (140-167): LLACISFDRYLNIVHATQLYRRGP PAR | X | X | X | Collecting tubule in medulla | Thermo Fisher |

*This antibody does not work well in FFPE tissue

**Supplementary Figure 1. Tissue microarray.**

**A.** Layout of the tissue microarray. The healthy and disease cores are denoted in black and red font, respectively.

| | | | | |
|---|---|---|---|---|
| Blank | 19SP-06319 (DKDIII) | Blank | KD-0783 (DM) | KD-0783 (DM) |
| DKDT3 (DKDIIA-B) | DKDT3 (DKDIIA-B) | KD-1557 (DM) | KD-1557 (DM) | DKDT7 (DKDIIB) |
| DKDT7 (DKDIIB) | KD-1984 (DM) | KD-1984 (DM) | KD-1279 (DKDIIA) | KD-1279 (DKDIIA) |
| WD-18360 (DM) | WD-18360 (DM) | KD-1780 (DKDIIA-B) | KD-1780 (DKDIIA-B) | WD-18279 (DM) |
| WD-18279 (DM) | KD-1974 (DKDIIB) | KD-1974 (DKDIIB) | KD-1974 (DKDIIB) | WD-18278 (DKDIIA) |

**B.** TMA block and the PAS-stained TMA section.

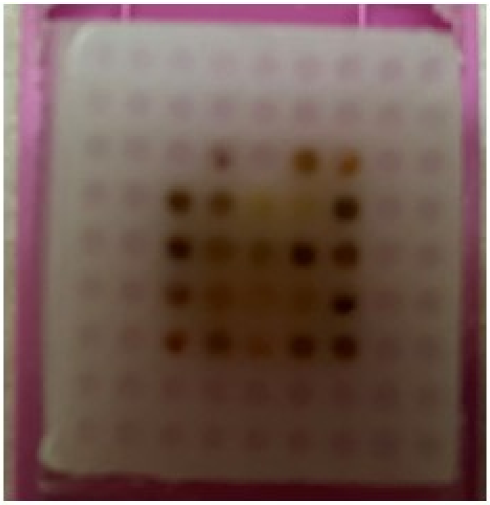

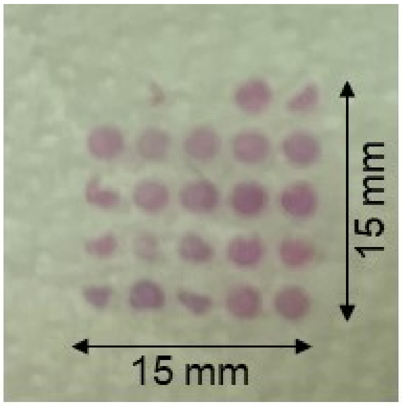

**Supplementary Figure 2.** Six compartments were identified using a combination of histologic appearance and marker protein expression from the multiplex immunofluorescence images and binary masks created. Glomeruli were identified based on histologic appearance; blood vessels were identified based on histologic experience and αSMA expression. Masks for distal nephron, all tubules, and the basement membrane were generated based on the expression of MUC1, CXCR3, and Collagen IV, respectively. The interstitium mask includes all tissue not within the previous five compartments.

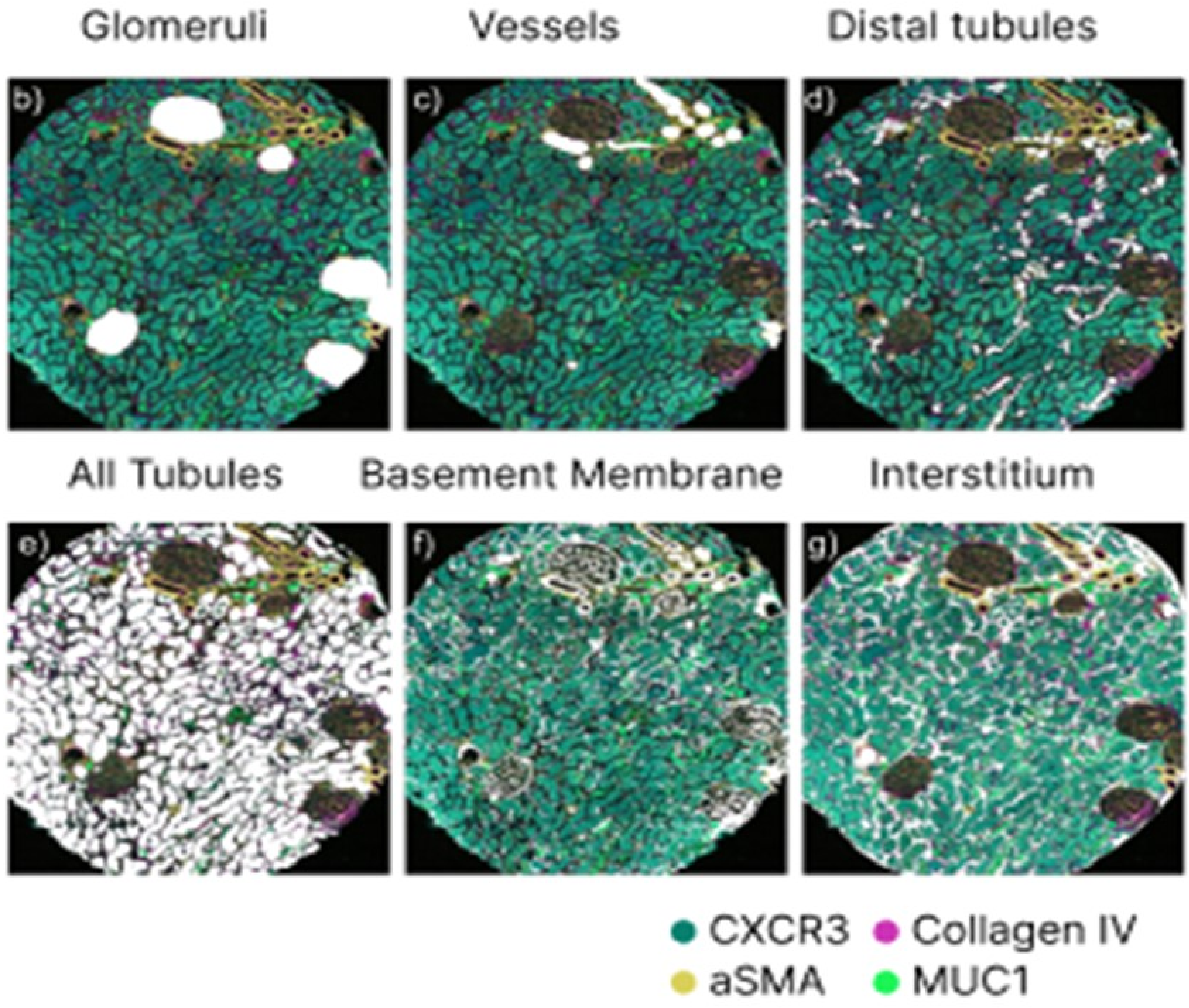

**Supplementary Figure 3. Prior data on kidney expression of the 21 targeted proteins** (110-page pdf)

**Supplementary Figure 4. Tissue expression for all proteins in all sections** (96-page pdf)

**Supplementary Figure 5. Multiplexed immunofluorescence in human kidney tissues from health to DKDIII. A.** Representative multiplexed immunofluorescence images showing protein expression in medullary and cortical sections in healthy kidneys from donors with diabetes.

A.

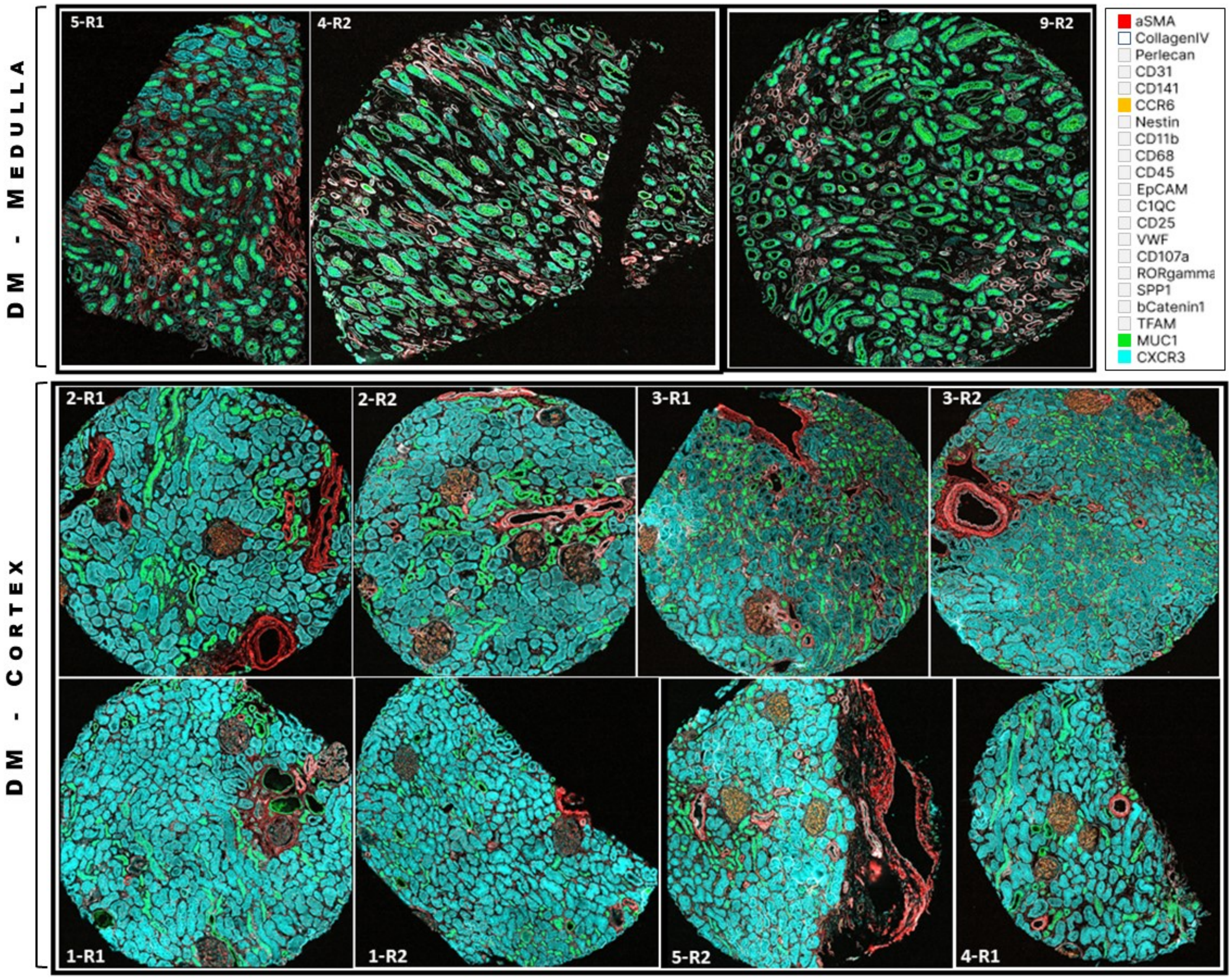

**Supplementary Figure 5. Multiplexed immunofluorescence in human kidney tissues from health to DKDIII. B.** Representative multiplexed immunofluorescence images showing protein expression in cortical sections in kidneys from donors with diabetes and DKDIIA to III.

**B.**

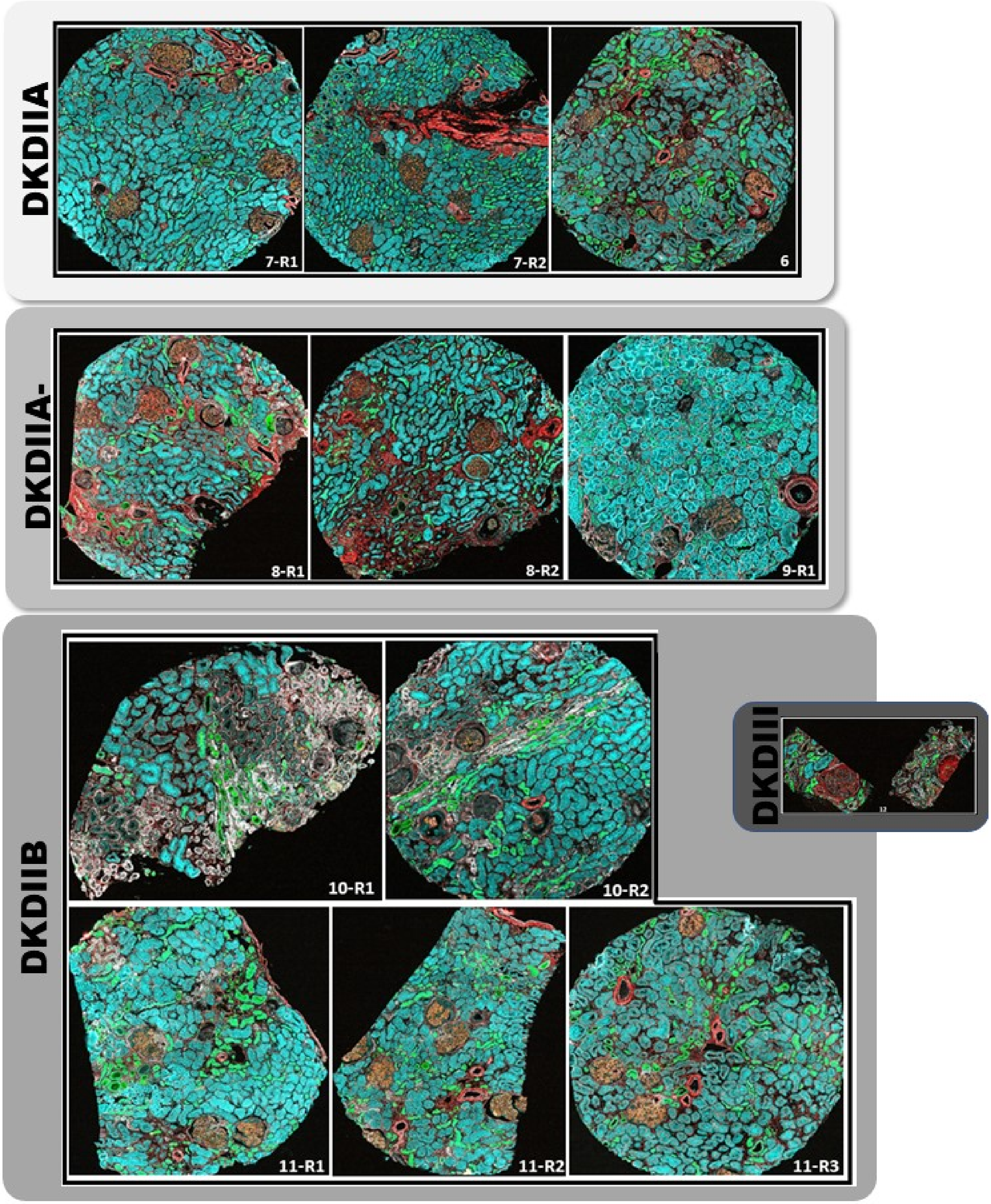

**Supplementary Figure 6. Looking for sections with outlier influence.** Dimension reduction by tissue sections shows all tissues contribute to every cluster. The expected exception is the absence of cortical tissue sections from medullary distal nephron clusters (cluster 9 and tip of cluster 8), which are from the medullary tissue sections only (highlighted in yellow in the legend).

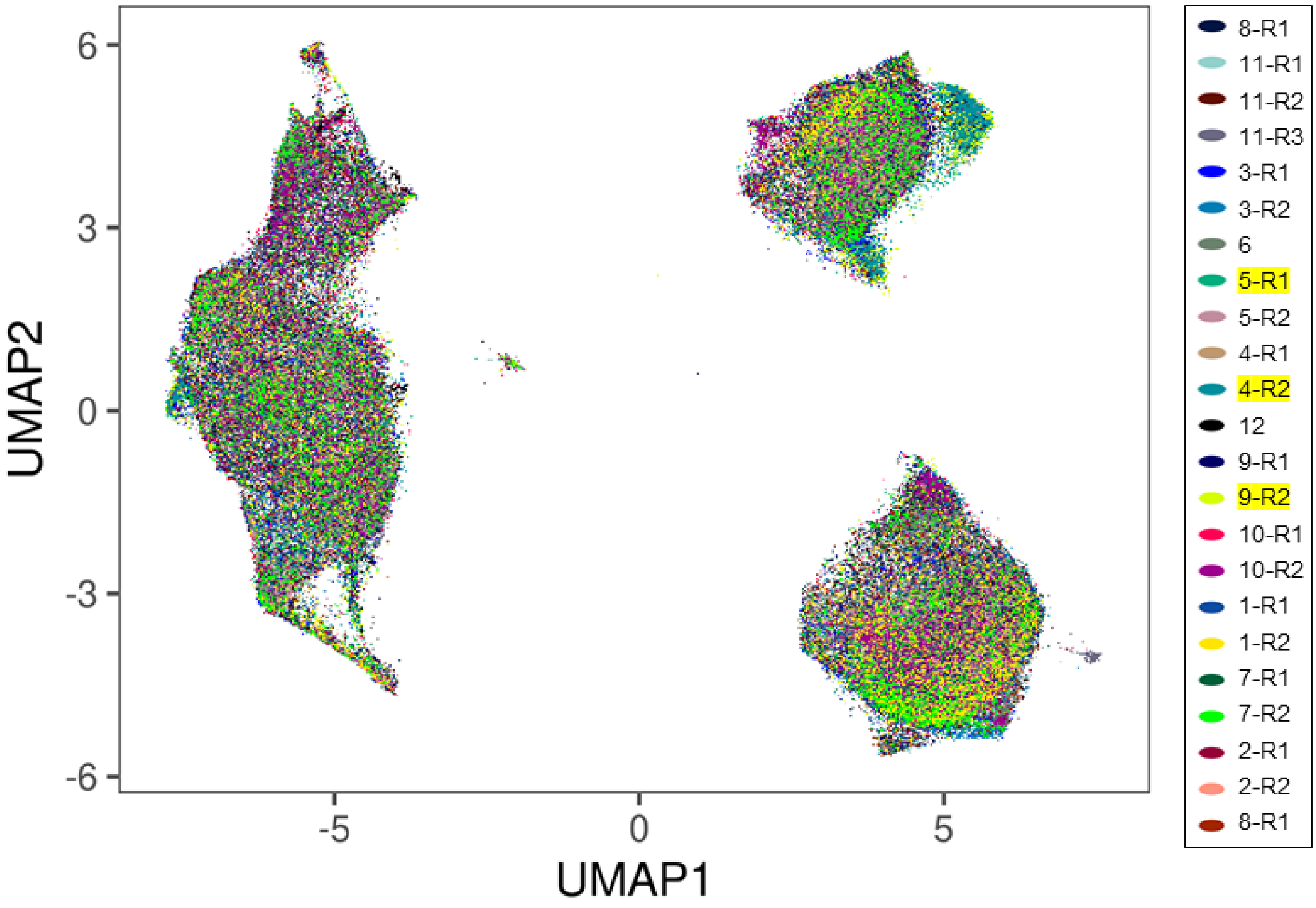
