## Supplementary Figure 3 for "Spatial proteomics of human diabetic kidney disease, from health to class III"

### α-SMA

#### Human Protein Atlas

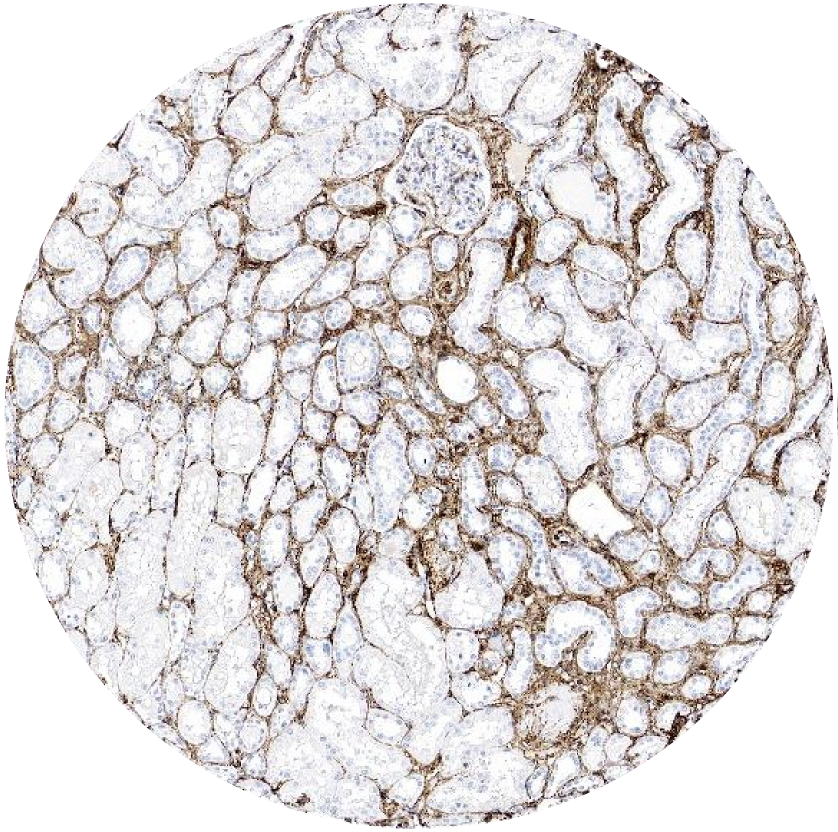

CAB000002
(Agilent M0851, mouse mAb supernatant
one band on western. No Protein array)

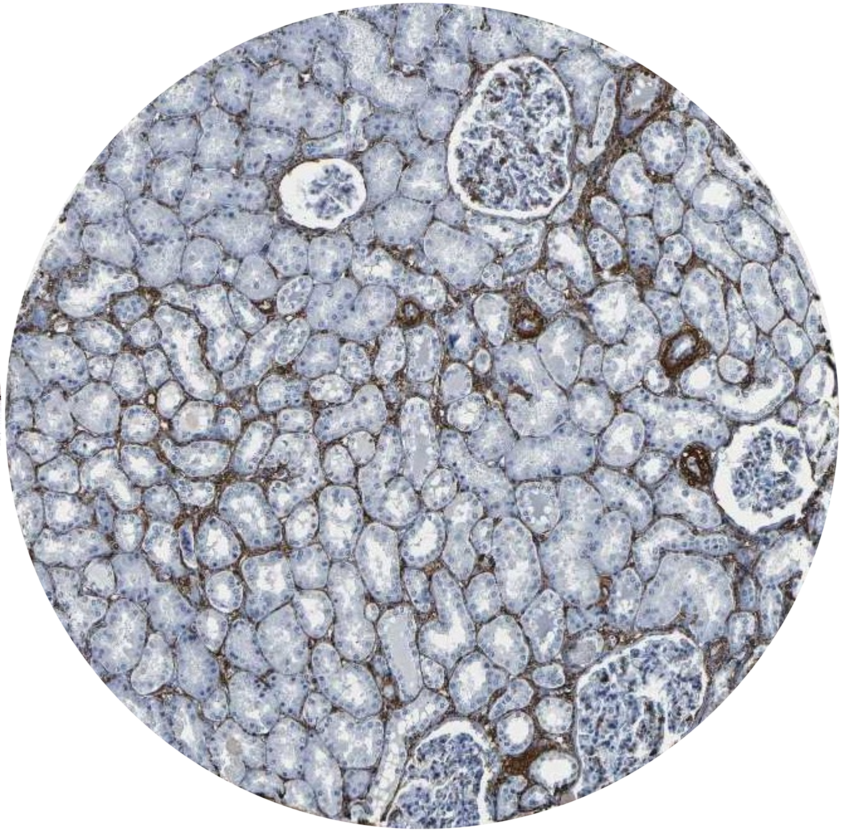

CAB013531
(Abcam 1184-1, Rabbit mAb supernatant
one band on western. No Protein array)

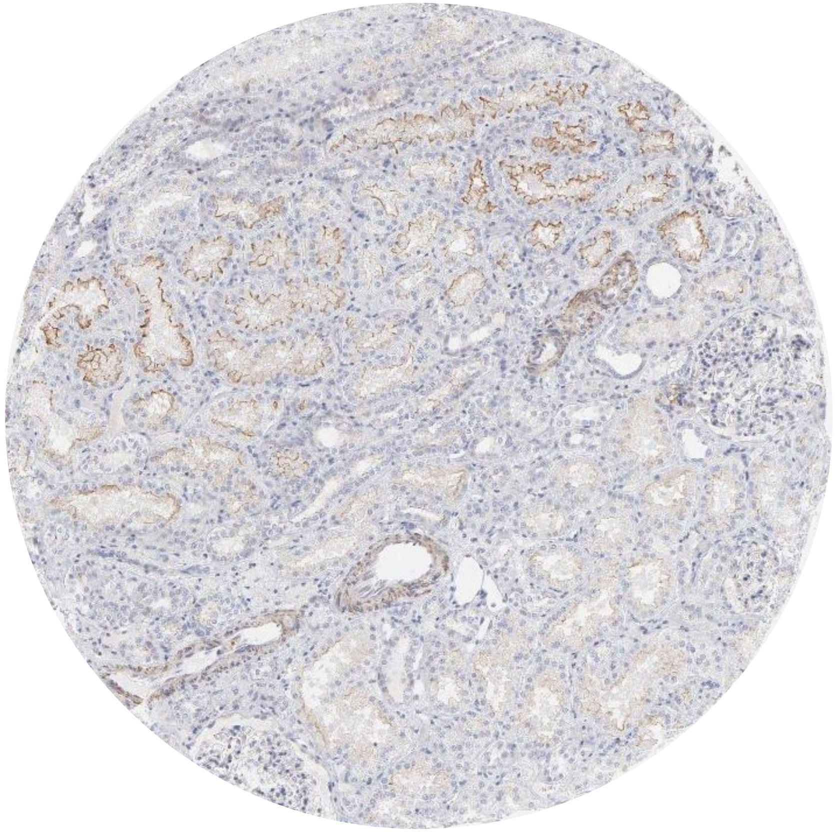

CAB003761
(Thermo Fisher 18-0106, mouse mAb
one band on western. No Protein array)

### α-SMA

**PMID 26808537**

**Figure 8. Expression of ENT1 and A3AR in human kidney.** A. Immunohistochemical detection of ENT1 and A3 receptor proteins were carried out in human kidney sections from non-diabetic normal tissue and biopsies from diabetic nephropathy patients. Selected images denote representative progressive stages of renal injury probed by the content of α-smooth muscle actin (α-SMA) and pathological analysis. Arrows indicate interstitial distribution of the immune signal. Original magnification 200x.

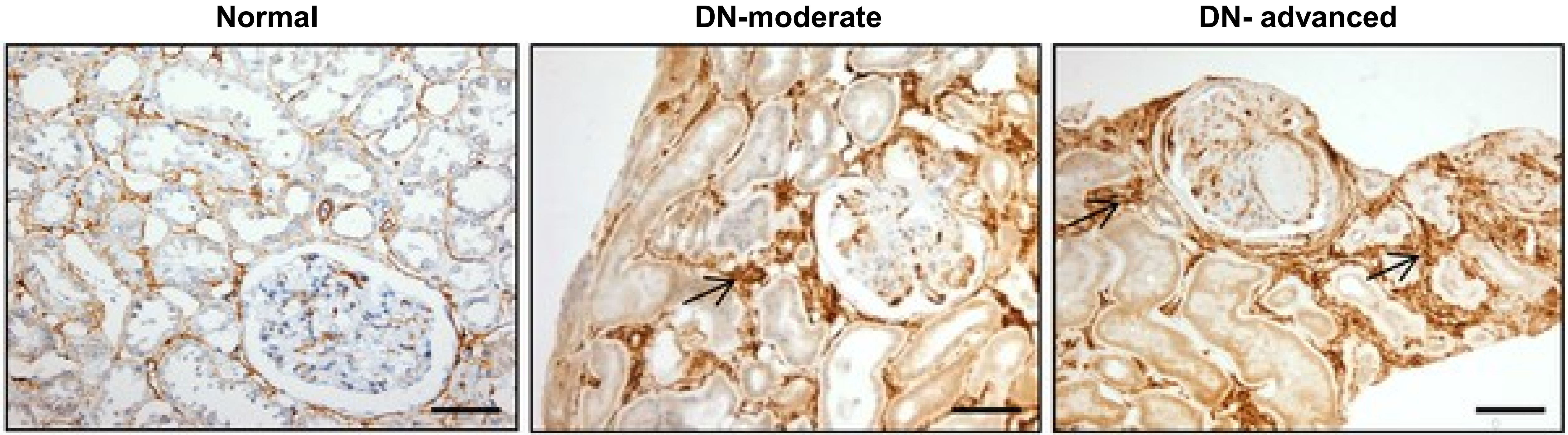

### α-SMA

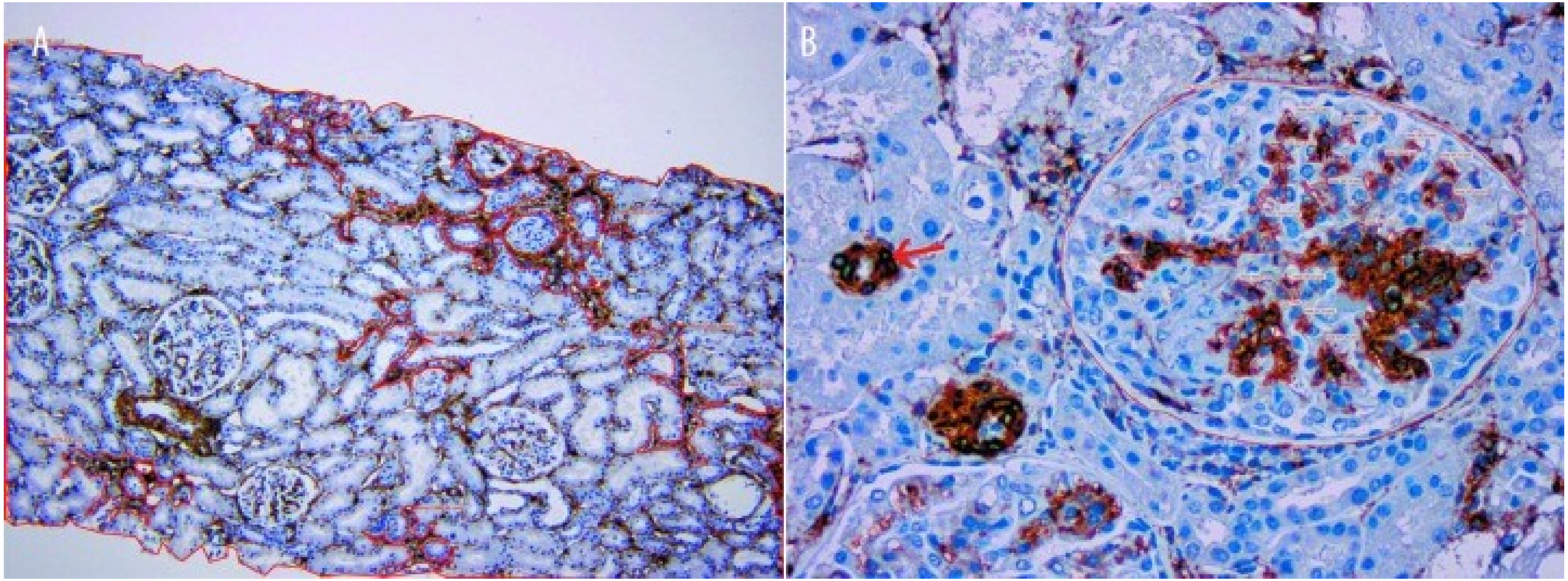


**PMID 22460095**

Figure 1. (A) Morphometric analysis of ASMA expression in interstitium and (B) glomeruli; arteriolar tunica media as positive control (arrow) (ASMA/HRP 100× and 400×).

# C1QC

#### Human Protein Atlas

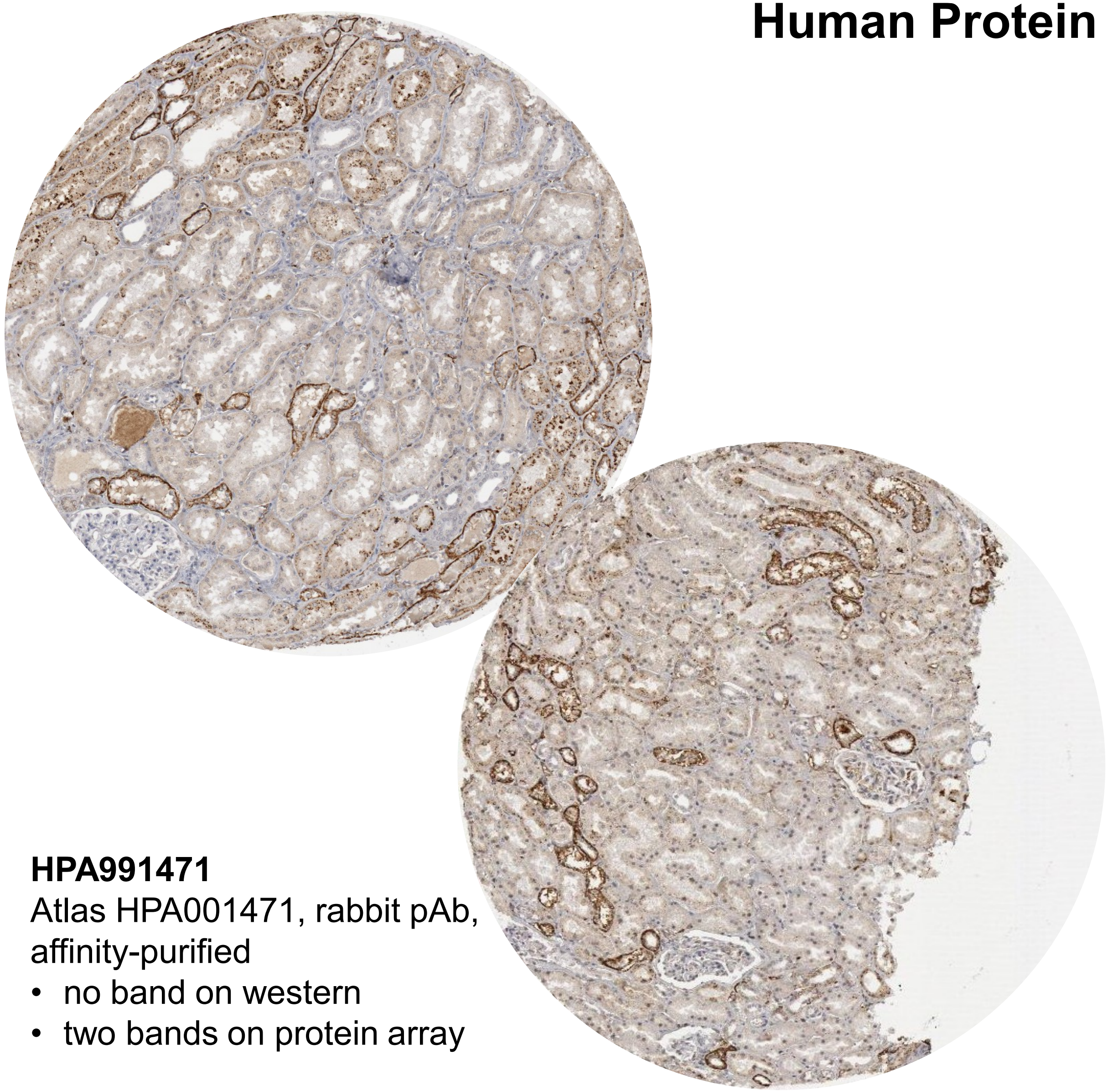

**HPA991471**
Atlas HPA001471, rabbit pAb,
affinity-purified
- no band on western
- two bands on protein array

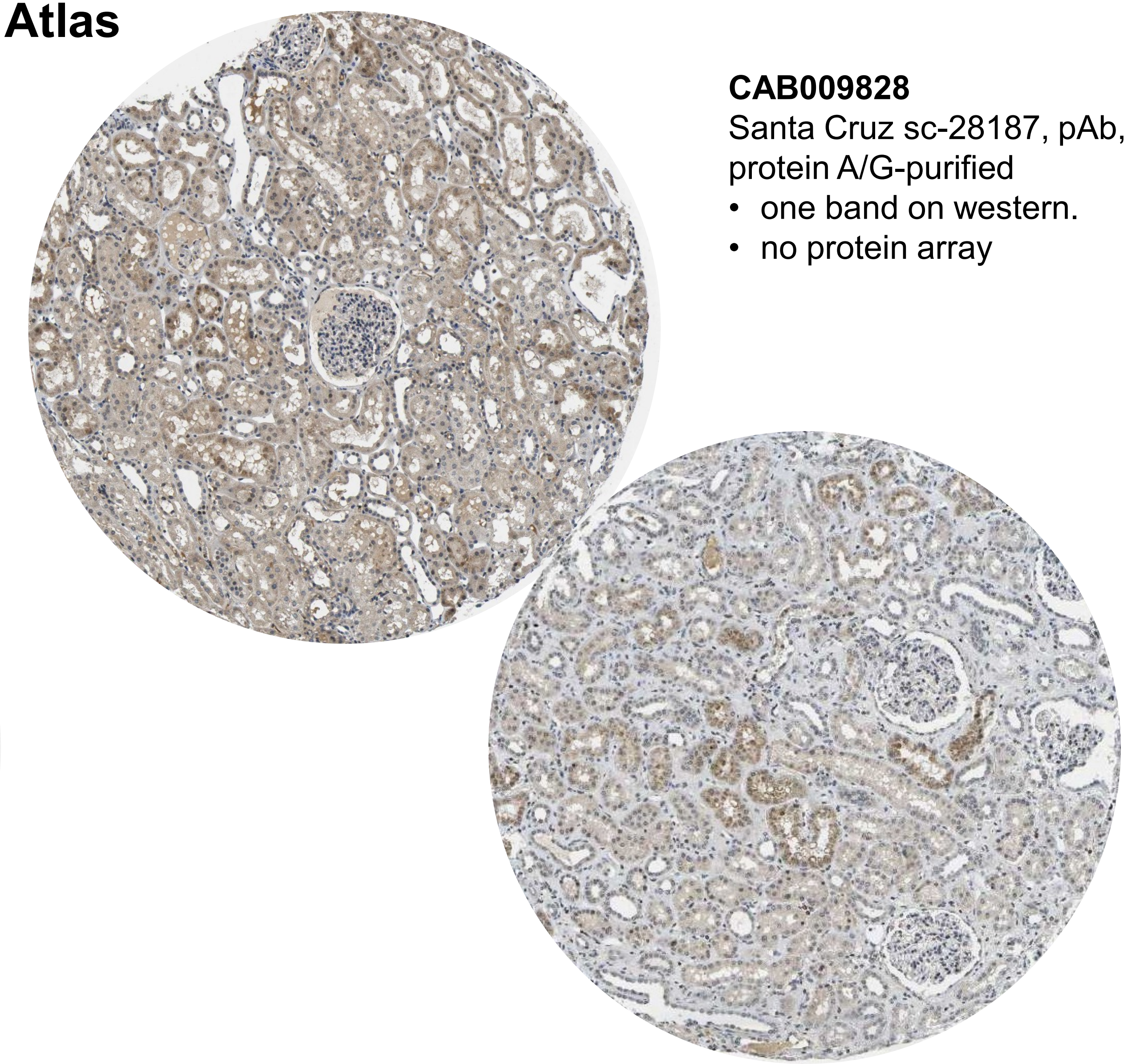

**CAB009828**
Santa Cruz sc-28187, pAb,
protein A/G-purified
- one band on western.
- no protein array

# C1QC

#### PMID 30286468

**Fig 4. C1q, a marker for the classical complement activation pathway, in zero-biopsies and transplant biopsies with DGF, ABMR and TCMR.** Complement factor C1q (C1q) was evaluated in the glomerular (A-C), in peritubular (D-F) and vascular (G-I) compartment using immunohistochemistry and semi-quantitative scoring (C, F, I). Renal transplant biopsies diagnosed for delayed graft function (DGF D), antibody mediated rejection (ABMR D) and T-cell mediated rejection (TCMR D) were compared to 1 year protocol biopsies (ctr 1y) as controls and corresponding zero-biopsies (ctr 0, DGF 0, ABMR 0 and TCMR 0). *p< 0.05, **p< 0.01.

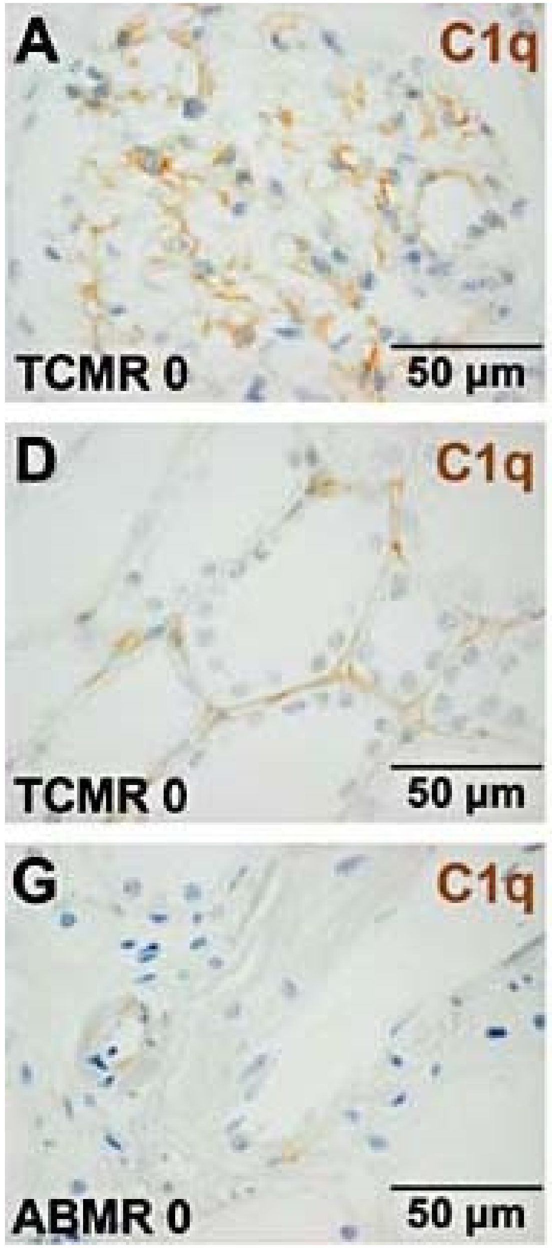


- Tissue: people on immunosuppression (post kidney transplant), some with diabetes
- Preservation: FFPE, Pronase E or HIER pH 6 antigen retrieval
- Rabbit polyclonal against C1q (A0126; DAKO)

# C1QC

**PMID: 34433493**

**Fig 2.** Immunohistochemical analysis of complement components in the glomeruli and renal tubules of LN patients and healthy controls. A Expression of C1q, C3, C5, C3aR, C5aR1, and CR3 in glomeruli (GLO) and renal tubules (TUB). B Immunohistochemistry -paraffin section (IHC-P) scores of C1q (N = 3, $p < 0.01$). C IHC-P scores of C3 (N = 6, $p < 0.001$). D IHC-P scores of C3aR (N = 5, $p < 0.01$). E IHC-P scores of C5 (N = 6, $p < 0.001$). F IHC-P scores of C5aR1 (N = 6, $p < 0.001$). G The IHC-P scores of CR3 (N = 3, $p < 0.01$). All IHC-P scores were determined using ImageJ software

- Tissue: Controls are tissue from healthy kidney donors (~half) or nephrectomy tissue from 'healthy' controls
- Preservation: FFPE, ?unclear retrieval protocol
- Antibodies: Bioss Biotechnology Co, Beijng, China

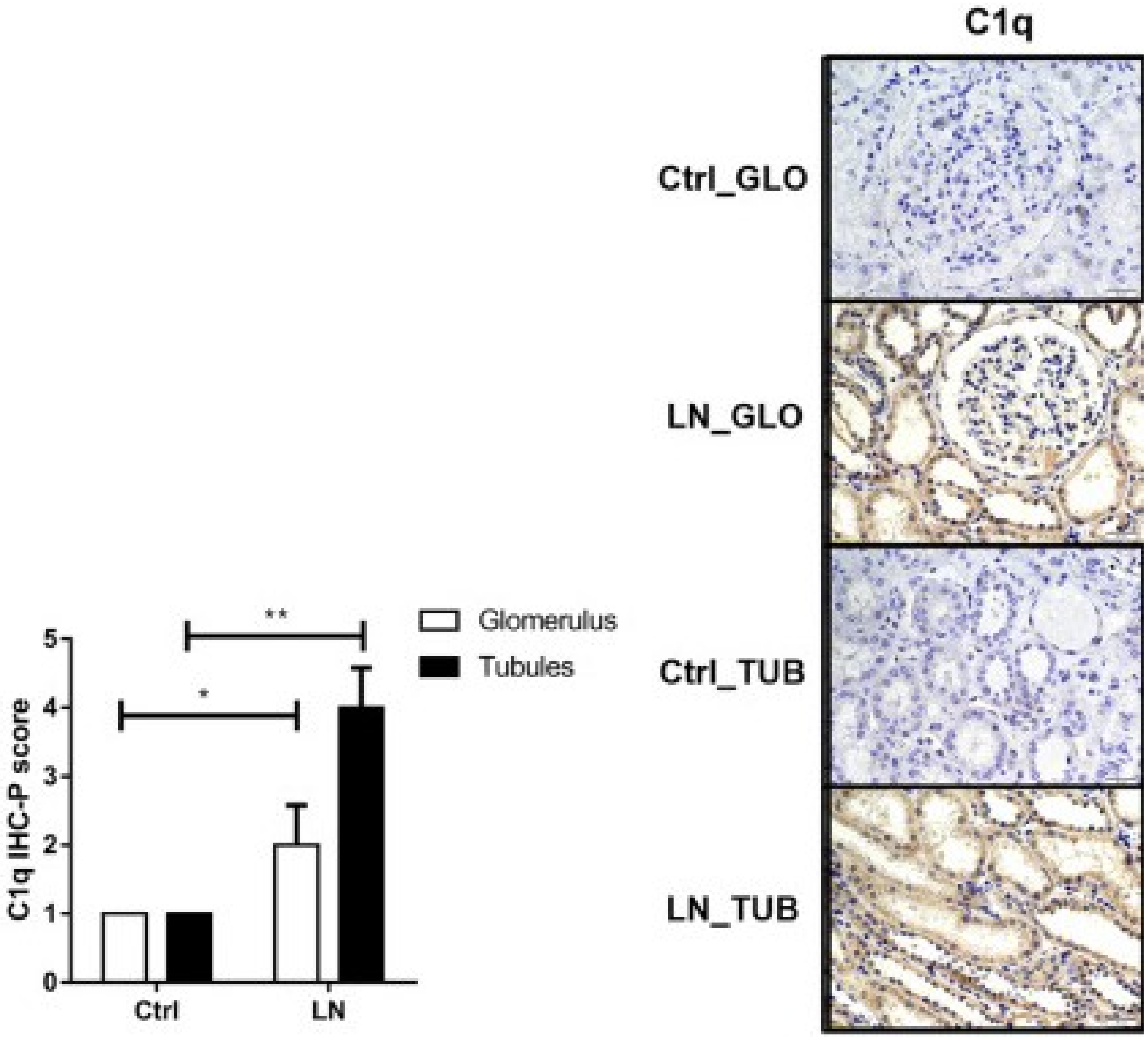

# C1QC

**PMID: 24658070**

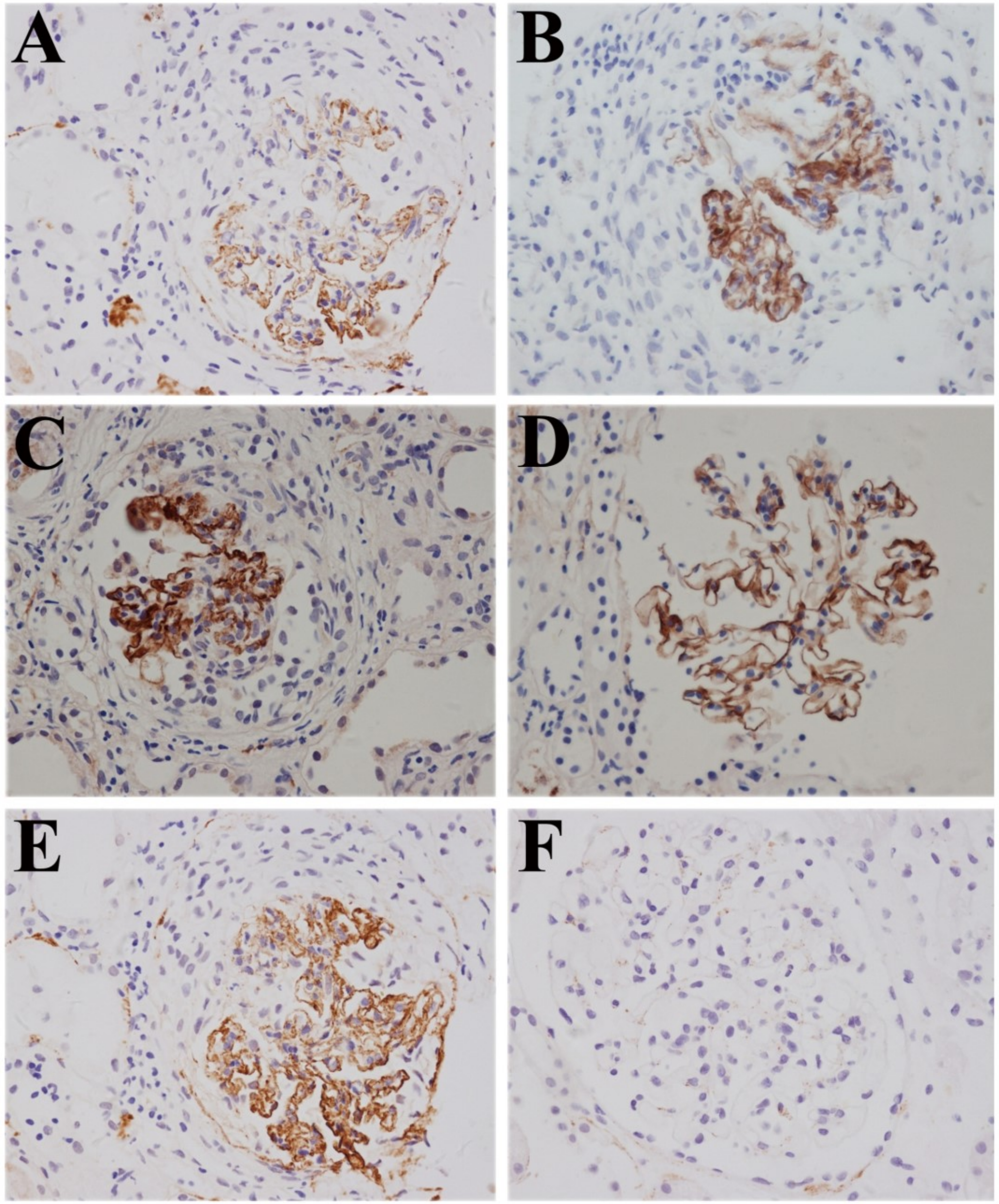


**Figure 1.** Complement components C5b-9, C3d, C4d, C1q and factor B were detected on paraffin sections of renal tissues from patients with anti-GBM disease by immunohistochemistry (magnification ×400). **A.** C5b-9 linear deposition along the glomerular capillary wall (Patient #7); **B.** C3d linear deposition along the capillary wall (Patient #2); **C.** C4d linear deposition along the capillary wall (Patient #5); **D.** C1q linear deposition along the capillary wall (Patient #1); **E.** factor B linear deposition along the capillary wall (Patient #7); **F.** all the complement components above were negative in the renal tissues from patients with minimal change disease

- Negative control was minimal change, not healthy control
- Ab clones not identified (Table 1 lists mouse anti-human mAb from abcam only)

# C1QC

**PMID: 24658070**

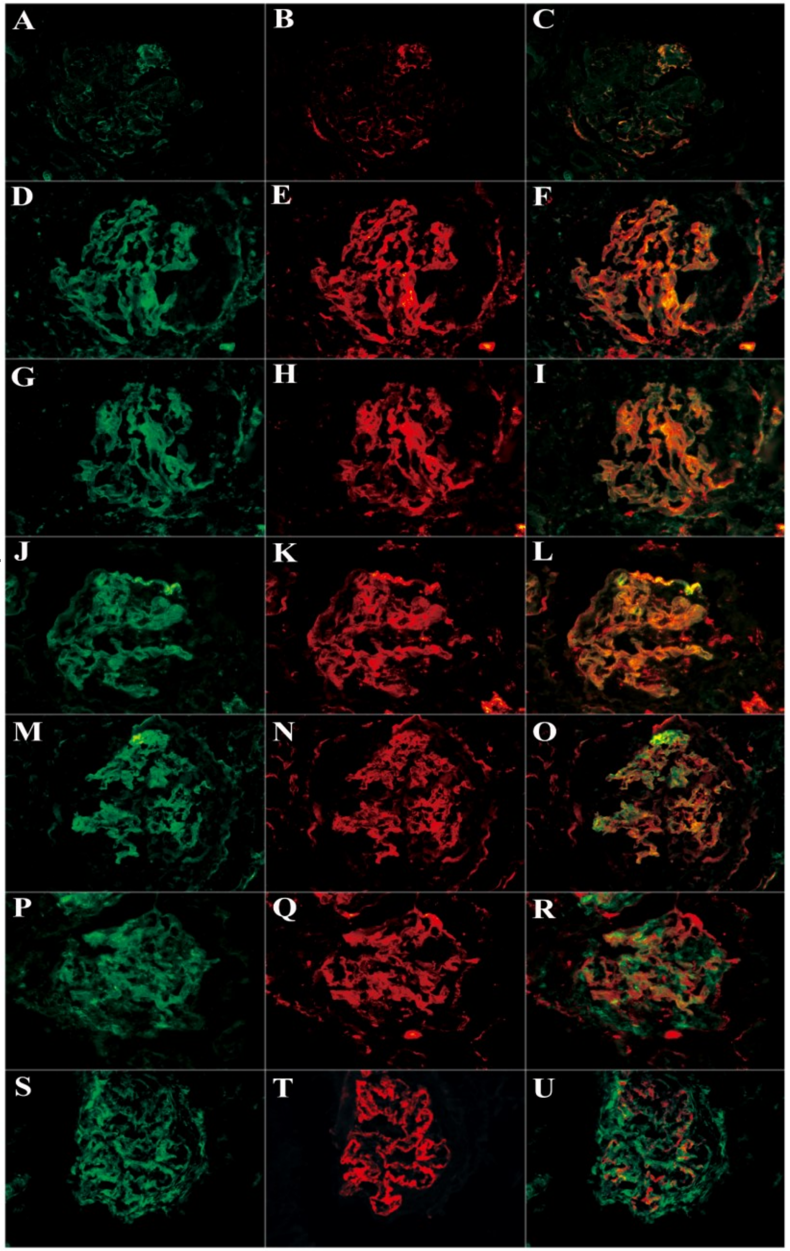


**Figure 3.** Co-localization of various complement components were detected in frozen sections by immunofluorescence using laser confocal microscopy (magnification ×400). A: IgG linear deposition along the glomerular capillary wall; B: C3d granular deposition along the glomerular capillary wall in the same section; C: IgG and C3d co-localized completely; **D: C1q linear deposition along the glomerular capillary wall;** E: C5b-9 granular deposition along the glomerular capillary wall in the same section; F: C1q and C5b-9 co-localized completely; G: factor B linear deposition along the glomerular capillary wall; H: C5b-9 granular deposition along the glomerular capillary wall in the same section; I: factor B and C5b-9 co-localized completely; J: properdin linear deposition along the glomerular capillary wall; K: C5b-9 granular deposition along the glomerular capillary wall in the same section; L: properdin and C5b-9 co-localized completely; M: properdin linear deposition along the glomerular capillary wall; N: C3d granular deposition along the glomerular capillary wall in the same section; O: properdin and C3d co-localized completely; P: MBL diffusive deposition; Q: C5b-9 granular deposition along the glomerular capillary wall in the same section; R: MBL and C5b-9 could not co-localize; S: MBL diffusive deposition; T: C4d granular deposition along the glomerular capillary wall in the same section; U: MBL and C4d partially co-localized along the glomerular capillary wall.

- Negative control was minimal change, not healthy control, and not identified in this figure
- Ab clones not identified (Table 1 lists mouse antihuman mAb from abcam only)

# C1QC

**PMID: 29725633**

**Figure 3.** Representative images of complement staining in patients. Kidney sections were immuno-stained for the indicated proteins, and representative images containing the glomeruli, glomerular hili, arterioles, and arterial branches are shown. Mannose-binding lection (MBL) staining was negative in the glomerular hilus, arterioles, and arterial branches. Bars = 50 µm.

**Text:** Glomerular C1q was present in 36% of the cases with diabetes, and the staining pattern was predominantly focal and global. The prevalence of glomerular C1q was not significantly different between the cases with diabetes and control cases without diabetes (37% vs. 49%, respectively; *P* = 0.150) (Figure 2b and Table 3). In contrast, the prevalence of C1q in the glomerular hili, arterioles, and arterial branches was significantly higher in the cases with diabetes than in the control cases without diabetes ($P \leq 0.001$). Furthermore, among the cases with diabetes, the cases with DN had a significantly higher prevalence of C1q in the glomerular hili and arterioles compared with the cases without DN ($P < 0.05$), whereas the prevalence of C1q in the arterial branches did not differ significantly between these 2 groups (*p* = 0.106). Finally, the presence of C1q deposits was correlated with the presence of C4d deposits in the glomeruli (*P* = 0.006), glomerular hili (*P* = 0.027), and arterioles ($P < 0.001$).

- Tissue: Autopsy, confirmatory data in HC, DKD biopsy
- No tubulointerstitial data
- Ab clone not identified but appears to be from DakoCytomation (?polyclonal, mAb?)

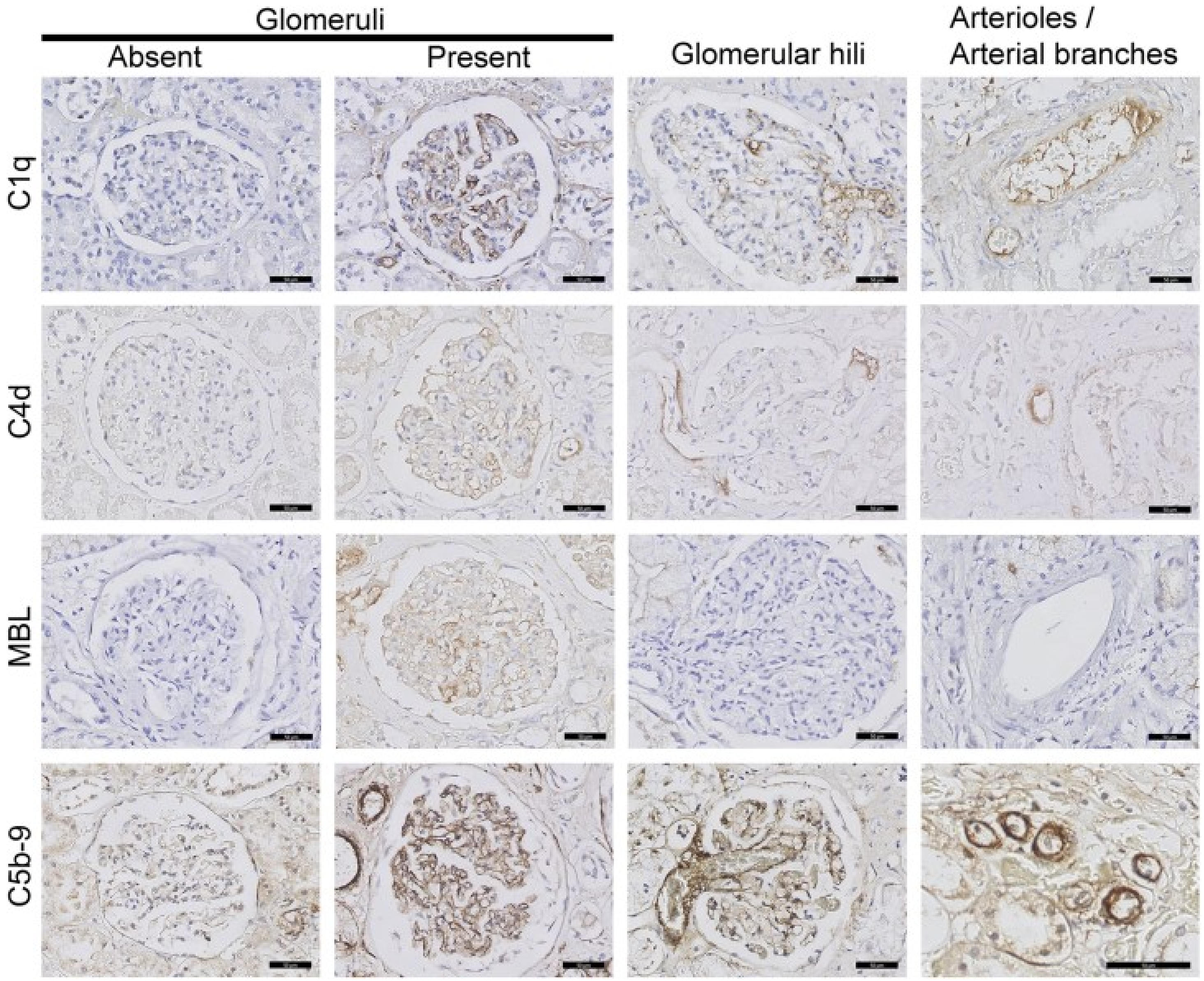

# C1QC

**PMID: 29725633**

**Figure 2.** (a–c) Prevalence of mannose-binding lection (MBL), C1q, and C5b-9 deposits in cases and controls. The percentage of cases and controls with complement factor MBL (a), C1q (b), and/or C5b-9 (c) is shown for the indicated renal structures. The P values shown between the 2 groups represent post hoc analyses. ***P < 0.001, χ2 test between nondiabetic controls, diabetic cases without diabetes nephropathy (DN), and diabetic cases with DN.

**Text:** Glomerular C1q was present in 36% of the cases with diabetes, and the staining pattern was predominantly focal and global. The prevalence of glomerular C1q was not significantly different between the cases with diabetes and control cases without diabetes (37% vs. 49%, respectively; *P* = 0.150) (Figure 2b and Table 3). In contrast, the prevalence of C1q in the glomerular hili, arterioles, and arterial branches was significantly higher in the cases with diabetes than in the control cases without diabetes (*P* ≤ 0.001). Furthermore, among the cases with diabetes, the cases with DN had a significantly higher prevalence of C1q in the glomerular hili and arterioles compared with the cases without DN (*P* < 0.05), whereas the prevalence of C1q in the arterial branches did not differ significantly between these 2 groups (*p* = 0.106). Finally, the presence of C1q deposits was correlated with the presence of C4d deposits in the glomeruli (*P* = 0.006), glomerular hili (*P* = 0.027), and arterioles (*P* < 0.001).

- Autopsy tissue
- Ab clone not identified but appears to be from DakoCytomation (?polyclonal, mAb?)

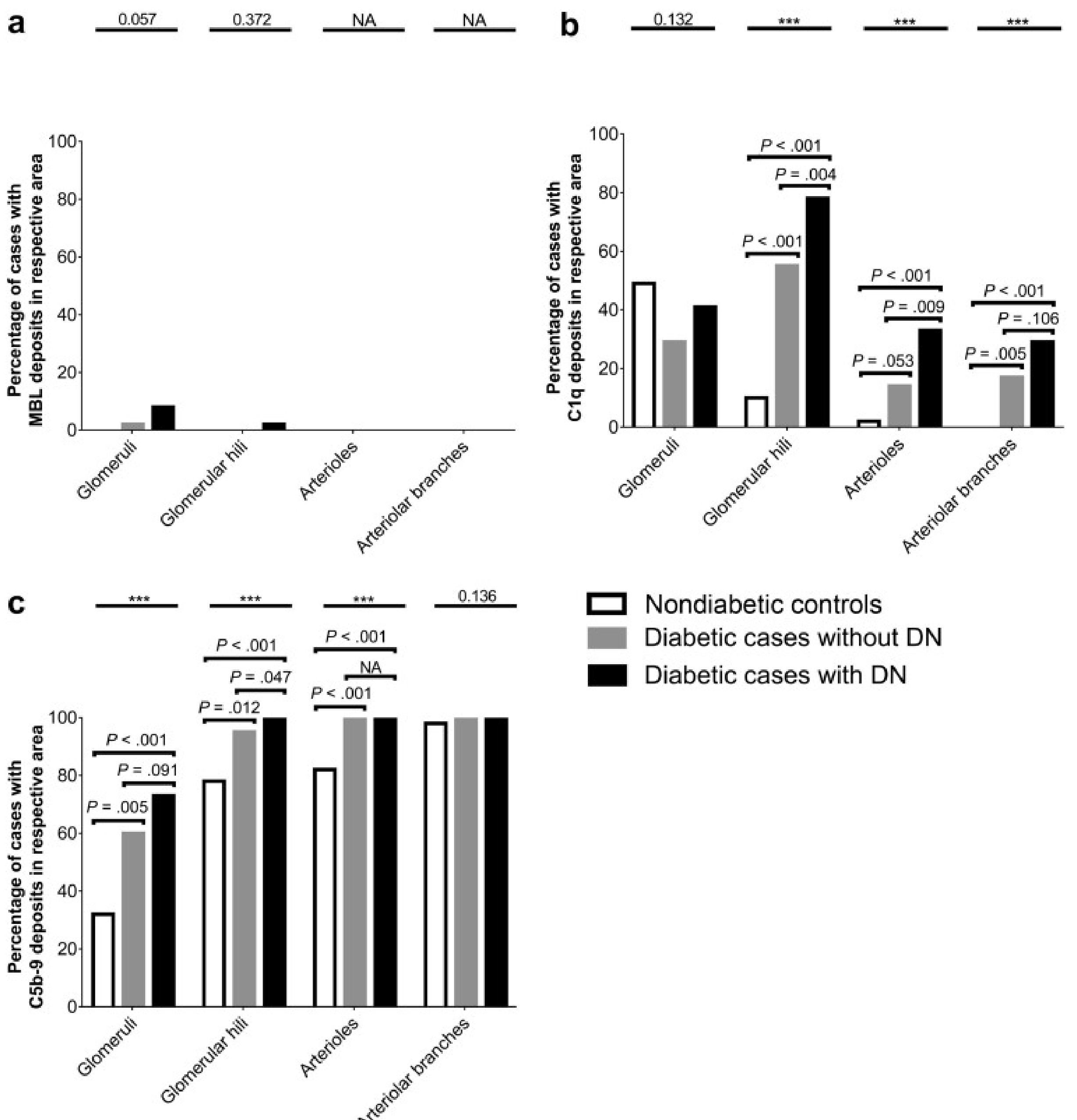

# C1QC

**PMID: 33584662**

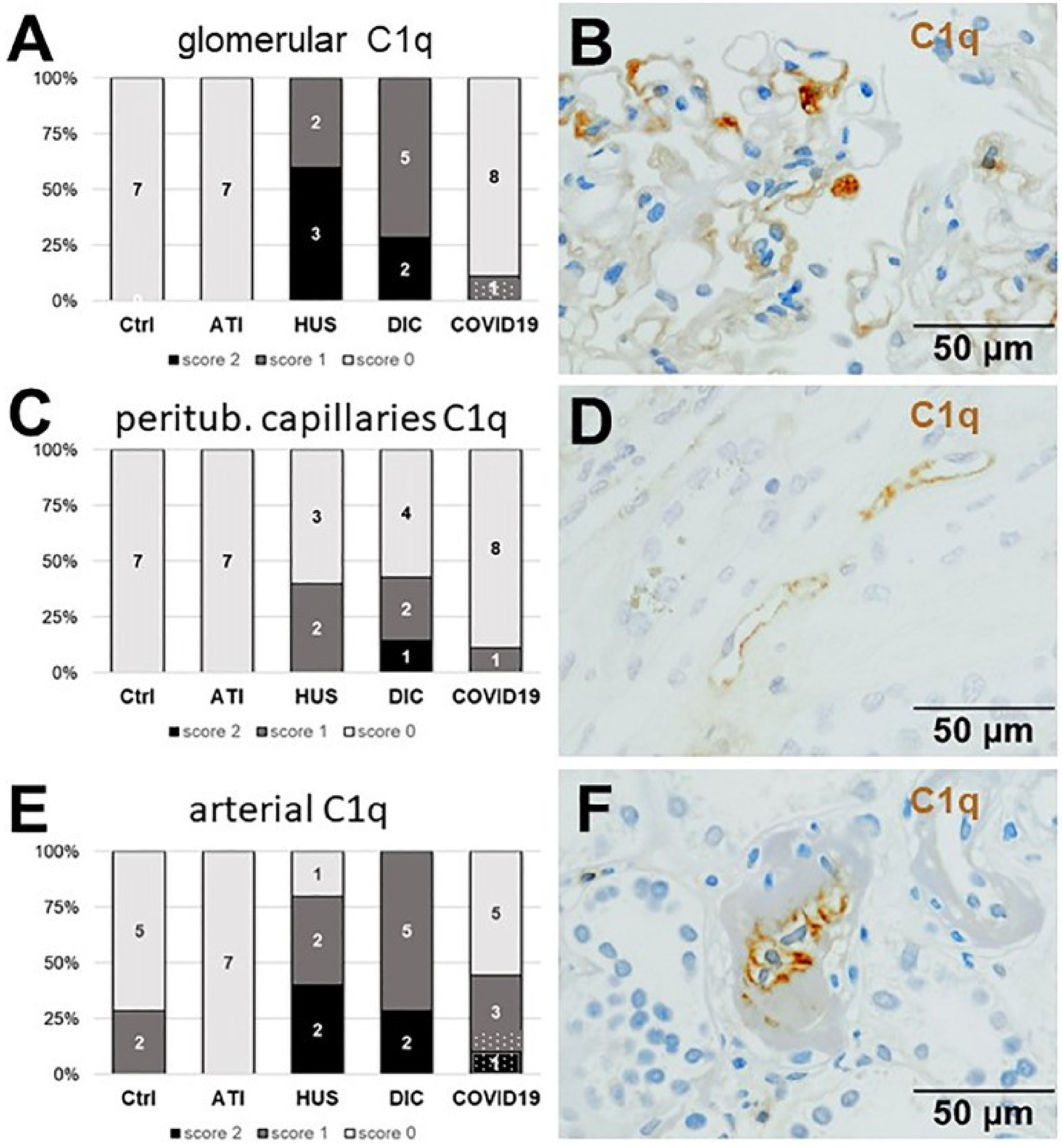


**Figure 6.** Frequency and vascular localization of C1q deposition in renal biopsies with COVID-19 compared to Ctrl, ATI, HUS, and DIC. Frequency and amount of C1q deposition were analyzed in glomerular capillaries (A), peritubular capillaries (C), and renal arteries (E) in renal control biopsies taken 1 year after transplantation (Ctrl, n = 7), biopsies with acute tubular injury (ATI, n = 7), hemolytic uremic syndrome (HUS, n = 5), disseminated intravascular coagulation (DIC, n = 7) and COVID-19 (COVID-19, n = 9) using immunohistochemistry and semi-quantitative scoring. The proportion of C1q-positive stained cases that belongs to cases with comorbidities with known involvement of complement activation is marked by hatched bars representing the COVID-19 cohort. Representative pictures of COVID-19 biopsies positive for C1q were shown for glomerular (B, brown staining), peritubular (D, brown staining) and arterial localization (F, brown staining.

- Controls were 0-time biopsies from stable kidney transplant recipients
- FFPE tissue preservation, pronase-E or heating antigen retrieval
- Rabbit polyclonal antibody against human C1q (A0136; DAKO Deutschland GmbH)

# C1QC

**PMID: 36059556**

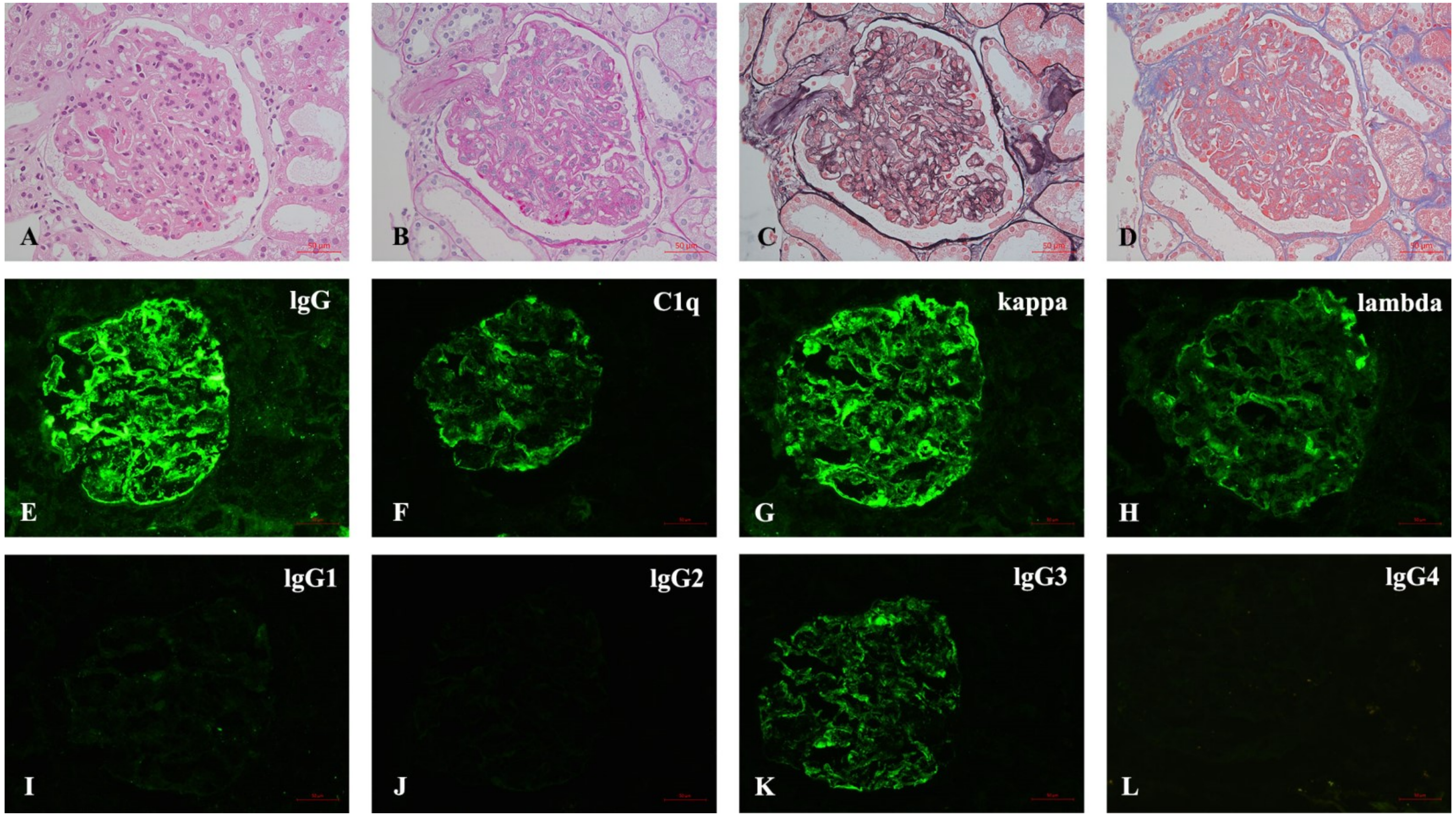


**Figure 1.** Pathological findings of renal biopsy. (A–D) Light microscopy shows a MPGN pattern of glomerular lesion (A), HE; (B), PAS; (C), PASM; (D), MASSON, 400×). (E–L) Immunofluorescence depicts granular mesangial and wall deposits with IgG, C1q, kappa, lambda, IgG3, and IgG2 trace, while IgG1 and IgG4 are negative [(E–L), 400×].

- One patient with MPGN. No negative control (case report)
- Gloms only
- Biopsy but no info on tissue preservation (?FFPE or fresh frozen OCT), antibody clone, etc

# C1QC

**PMID: 34440207**

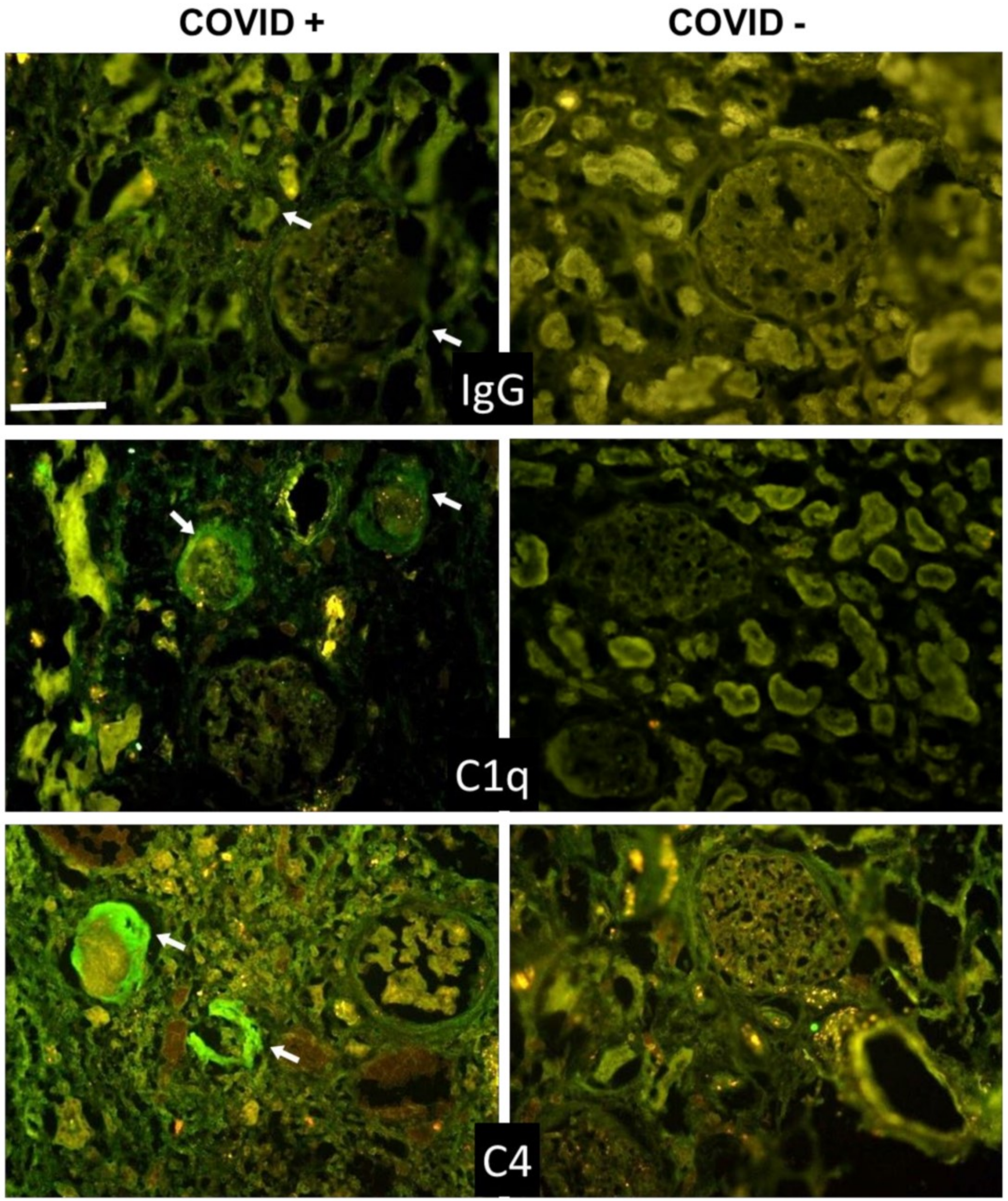


**Figure 3.** Detection of IgG, C1q, and C4 in kidney sections by immunofluorescence analysis. The panel shows representative immunofluorescence images obtained from the analysis of several sections of kidney autopsy samples from 12 SARS-CoV-2-positive and 3 negative cases. See legend to Figure 1 for further details. White arrows show IgG, C1q, and C4 periglomerular deposition. Scale bar = 50 µm.

- Fresh frozen tissue, FFPE
- Goat anti-C1q (The Binding Site, Birmingham, UK)
- Unclear what positive staining is

### CD196 (CCR6)

**Human Protein Atlas**

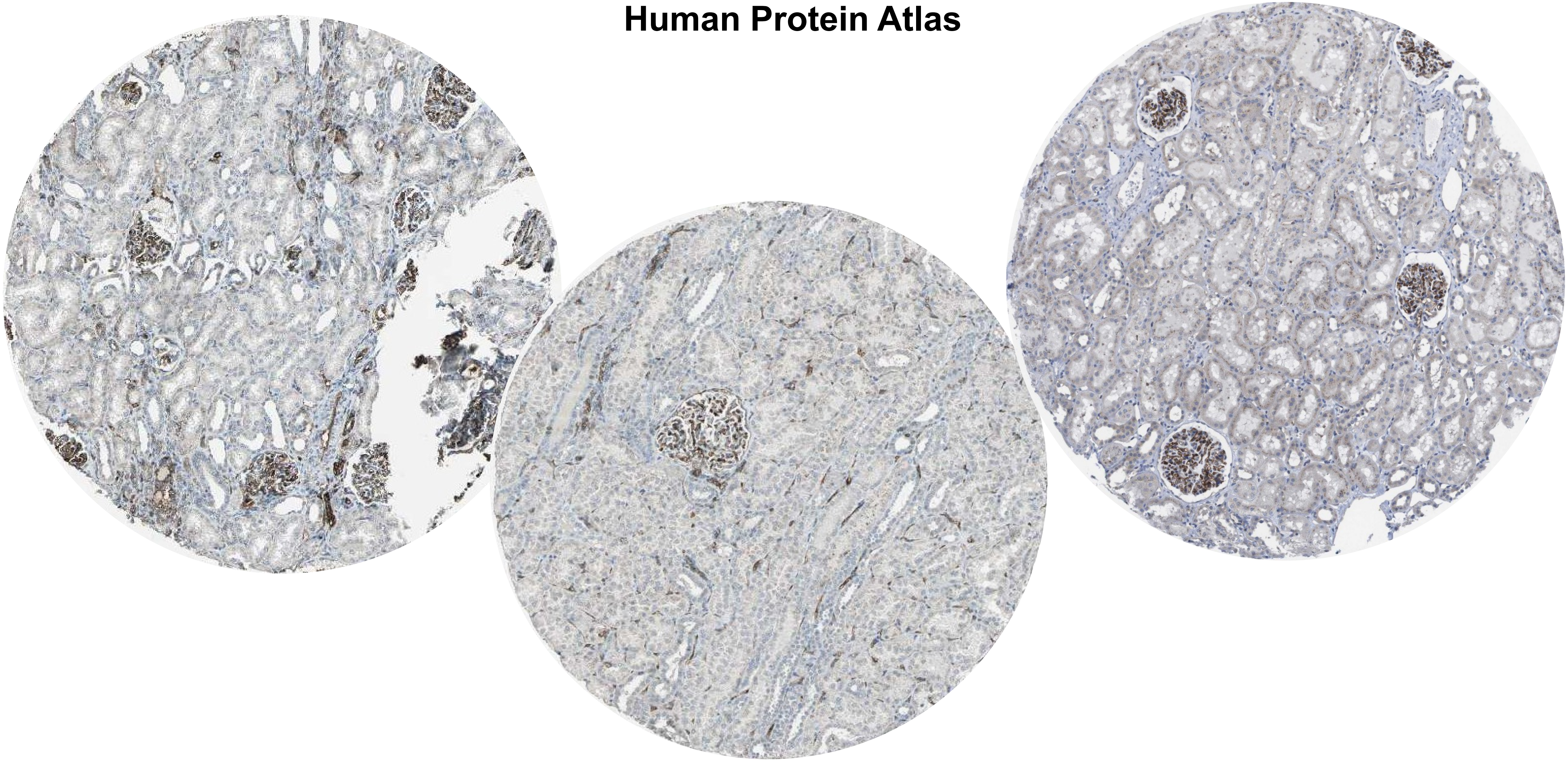

### CD196 (CCR6)

**PMID: 20844183**

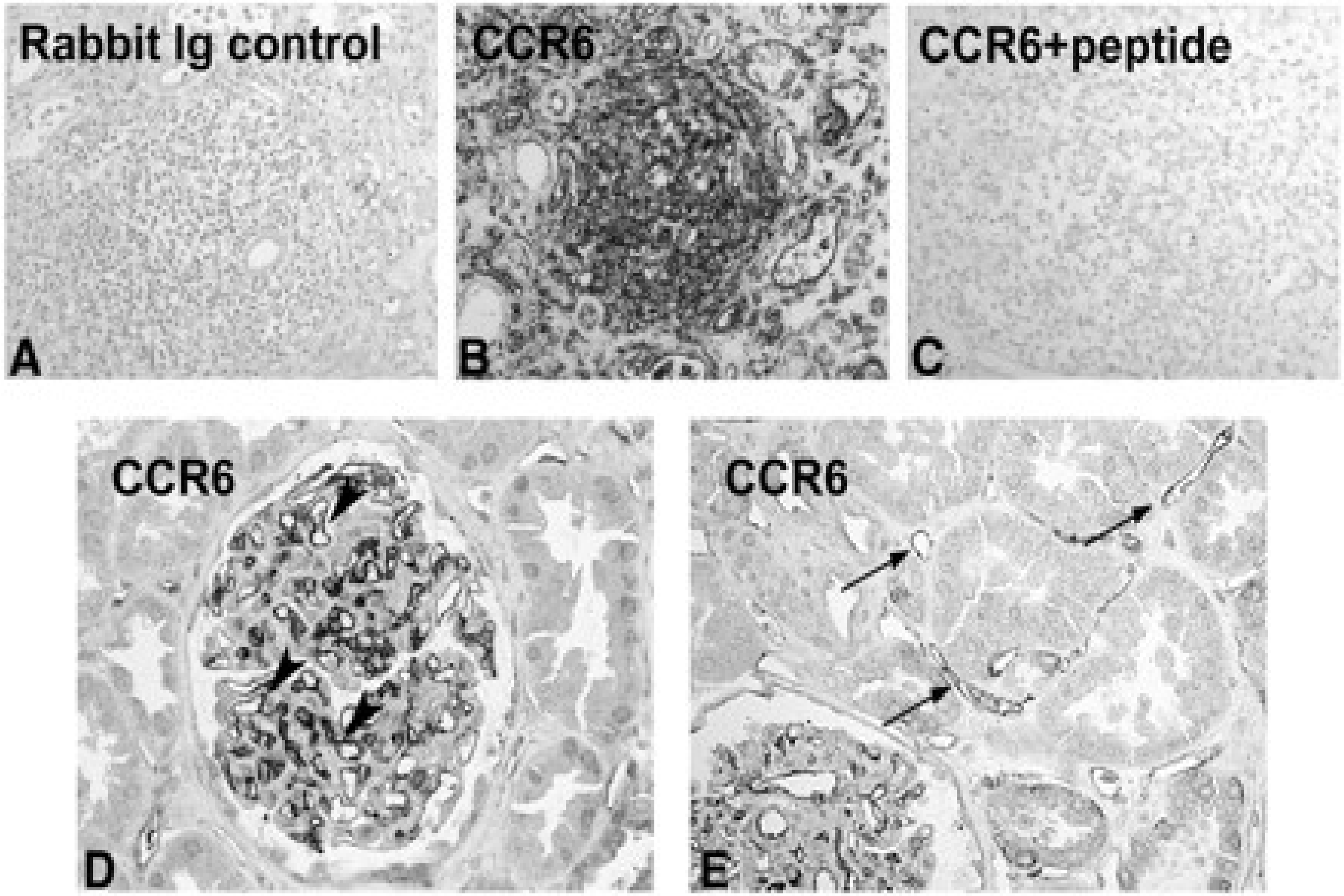


**Fig. 1** Establishment of the anti-CCR6 antibody and expression in normal human kidney. Immunohistochemistry was performed on an allograft nephrectomy (A–C), and a pre-transplant biopsy (D and E) with a polyclonal antibody against CCR6 (B, D and E, Chemicon), a non-immune rabbit serum (A), and after pre-absorption of the antiserum with the peptide (C, original × 250 D, E, original × 200 in A–C). The negative controls performed with non-immune rabbit serum (A) and pre-absorption with the peptide used for the induction of the antiserum did not demonstrate a positive colour product (C). Prominent signal of a nodular infiltrate in the allograft nephrectomy (B). In the well-preserved pre-transplant biopsies, a positive signal was present on endothelial cells of glomerular capillaries (arrowheads, D) and peritubular capillaries (E, arrows).

Two polyclonal antibodies against CCR6 raised in two different rabbits were tested and resulted in the same pattern (anti-human CCR6 by MBL International Corporation, Woburn, MA, USA and Chemicon, Temecula, CA, USA). The primary polyclonal antibody against CCR6 (Chemicon).

# CD196

**PMID: 28552941**

**Fig 7.** CD3$^+$CCR6$^+$IL-17$^+$ cells are detected by immunofluorescence staining for section from IgAN patients or NS (non-IgAN nephrotic syndrome).

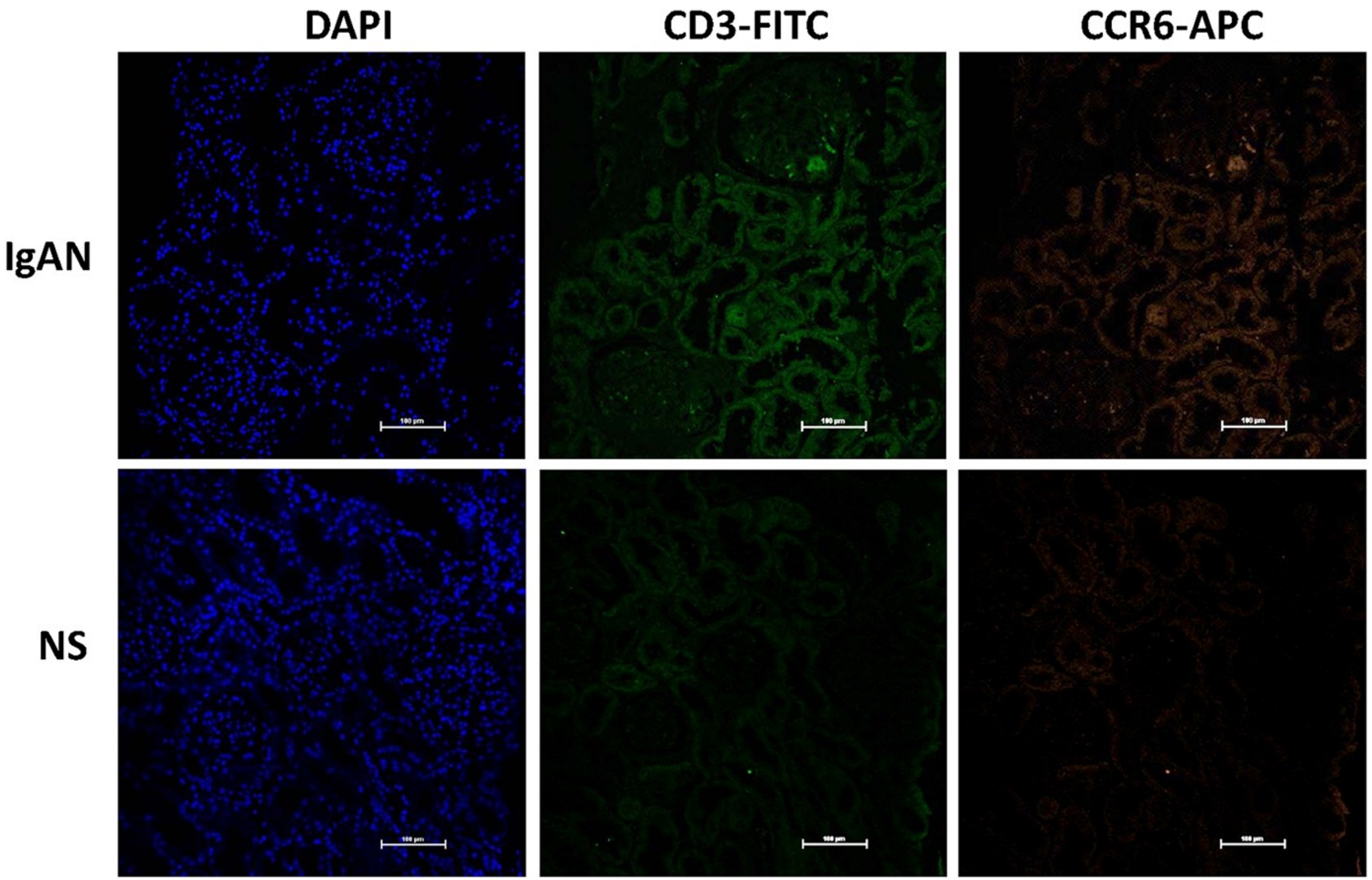


- No control (nephrotic syndrome is used as control)

### CD11b (ITAM, CR3)

**Human Protein Atlas**

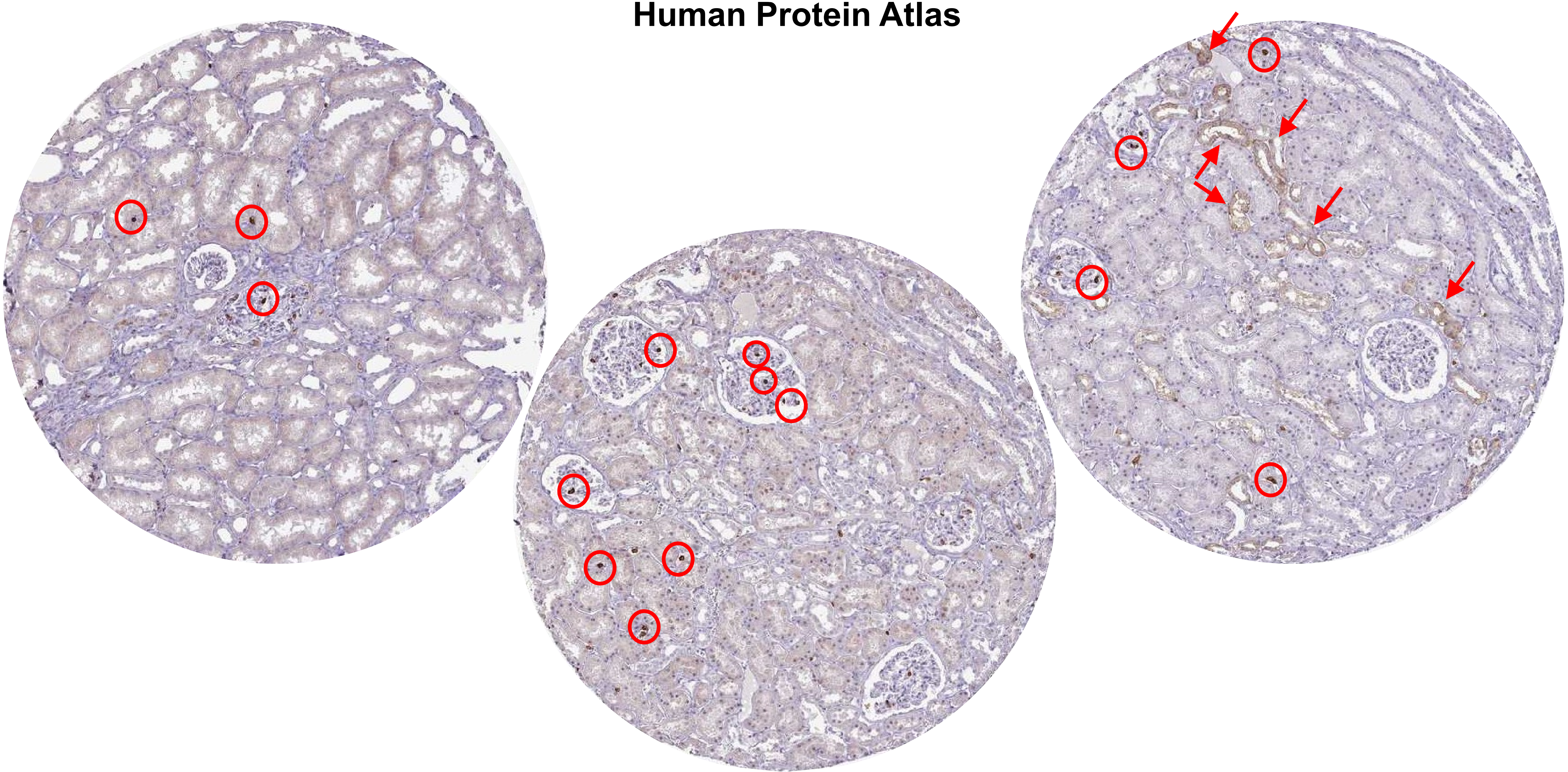

### CD11b (ITAM, CR3A)

**PMID: 1346576-** Predominantly expressed in monocytes and granulocytes

**PMID: 21193407-** Expressed in neutrophils

**Uniprot (https://www.uniprot.org/uniprotkb/P11215/entry)**

- Integrin ITGAM/ITGB2 is implicated in various adhesive interactions of monocytes, macrophages and granulocytes as well as in mediating the uptake of complement-coated particles and pathogens (PubMed:9558116, PubMed:20008295).
- It is identical with CR-3, the receptor for the iC3b fragment of the third complement component. It probably recognizes the R-G-D peptide in C3b.
- Integrin ITGAM/ITGB2 is also a receptor for fibrinogen, factor X and ICAM1. It recognizes P1 and P2 peptides of fibrinogen gamma chain.
- Regulates neutrophil migration (PubMed:28807980).
- In association with beta subunit ITGB2/CD18, required for CD177-PRTN3-mediated activation of TNF primed neutrophils (PubMed:21193407).

### CD31 (PECAM1)

**Human Protein Atlas**

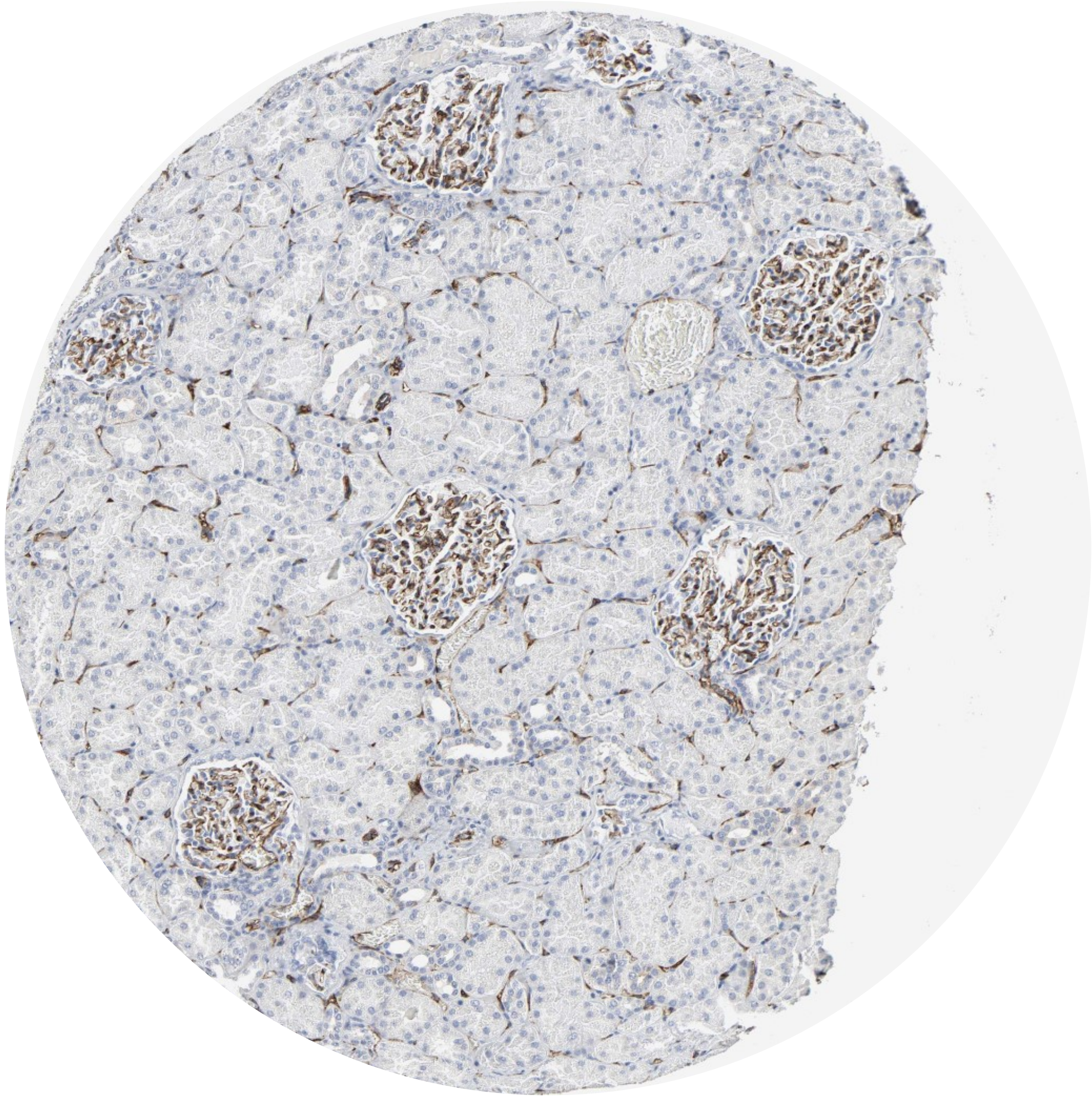

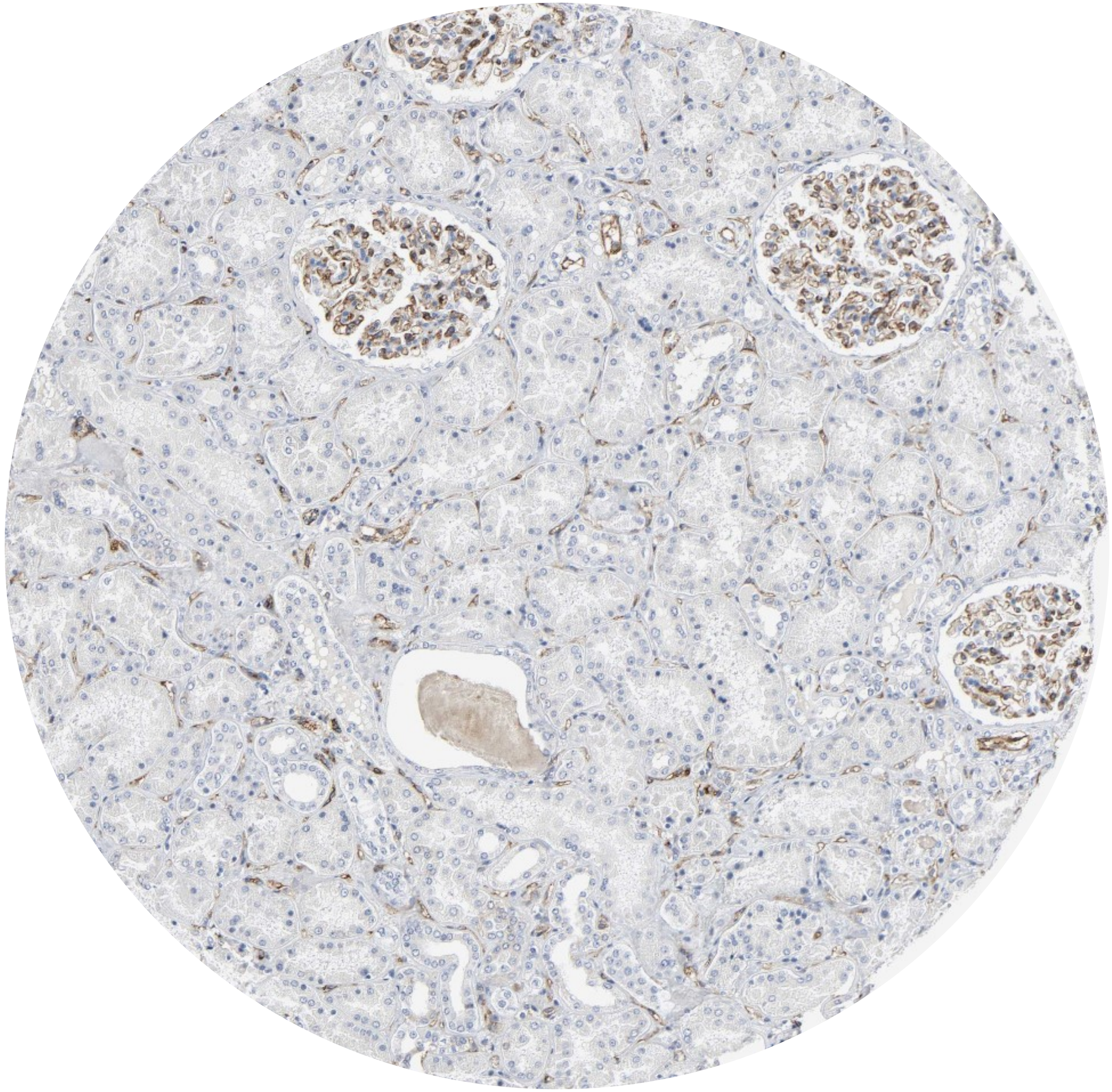

### CD31 (PECAM1)

**PMID 16234507**

**Figure 1** (**A**) CD31 expression in the kidney showing 2 glomeruli with strong positivity. (**B**) CD34 expression pattern in the kidney is similar to CD31 but more interstitial capillaries are stained. (**C**) In contrast to CD31 and CD34, the glomeruli are negative for von Willebrand factor (vWF). (**D**) Fli-1 nuclear immunostaining of EC present in the glomeruli and interstitial capillaries.

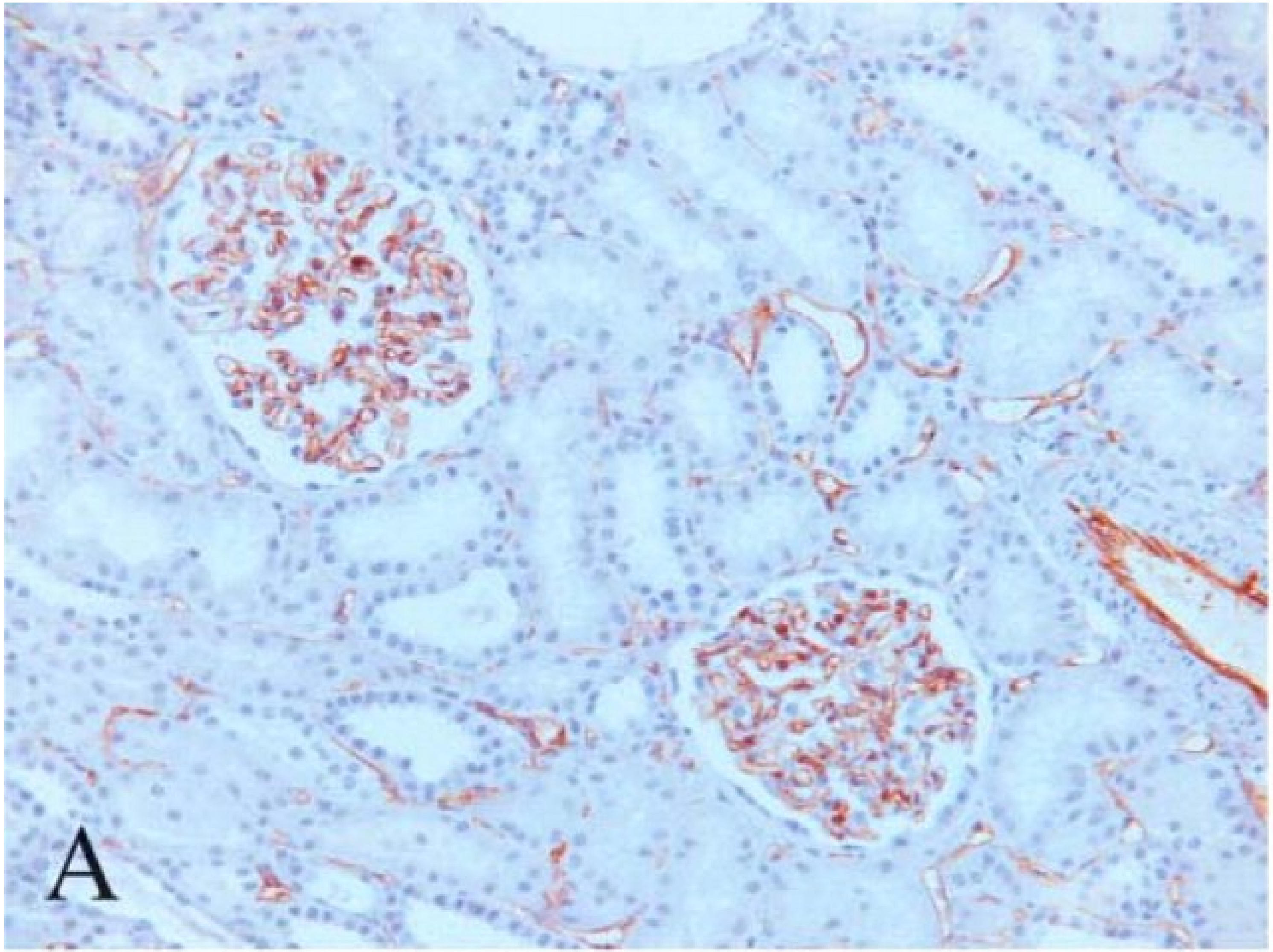

### CD31 (PECAM1)

**PMID 16234507**

**Table 2** Staining of endothelial cells for CD31, CD34, vWF, and Fli-1 in various normal adult tissues

| Tissue | CD31 | CD34 | VWF | Fli-1 (nuclear) |
|---|---|---|---|---|
| Kidney | | | | |
| Glomeruli | +++ | +++ | 0/+ | +++ |
| Capillaries | ++ | +++ | ++ | +++ |
| Venules | +++ | +++ | +++ | +++ |
| Arteries | +++ | +++ | +++ | +++ |
| Lung | | | | |
| Capillaries | +++ | +++ | 0/+ | +++ |
| Arterioles | +++ | +++ | ++ | +++ |
| Venules | +++ | +++ | ++ | +++ |
| Veins | +++ | +++ | +++ | +++ |
| Arteries | +++ | +++ | +++ | +++ |
| Liver | | | | |
| Periportal sinusoids | +++ | +++ | ++ | +++ |
| Centrolobular sinusoids | +++ | 0/+ | ++ | +++ |
| Centrolobular veins | +++ | +++ | ++ | +++ |
| Portal venules | +++ | +++ | ++ | +++ |
| Arterioles | +++ | +++ | ++ | +++ |
| Spleen | | | | |
| Sinusoids | ++ | 0 | ++ | ++[a] |
| Capillaries | 0 | +++ | 0/+ | ++[a] |
| Central arteries | +++ | +++ | ++ | ++[a] |
| Venules | +++ | +++ | ++ | ++[a] |
| Lymph nodes | | | | |
| Capillaries | +++ | +++ | ++ | ++[a] |
| Marginal sinuses | +++ | 0/+ | 0/+ | ++[a] |
| High endothelial venules | +++ | +++ | +++ | ++[a] |

Intensity of immunohistochemical staining: 0, absent; +, low; ++, medium; +++, high.
[a]Most cells present (lymphocytes) are stained.

### CD31 (PECAM1)

https://www.lsbio.com/antibodies/ihc-plus-pecam-1-antibody-cd31-antibody-clone-2f7b2-elisa-ihc-wb-western-ls-b2364/83552

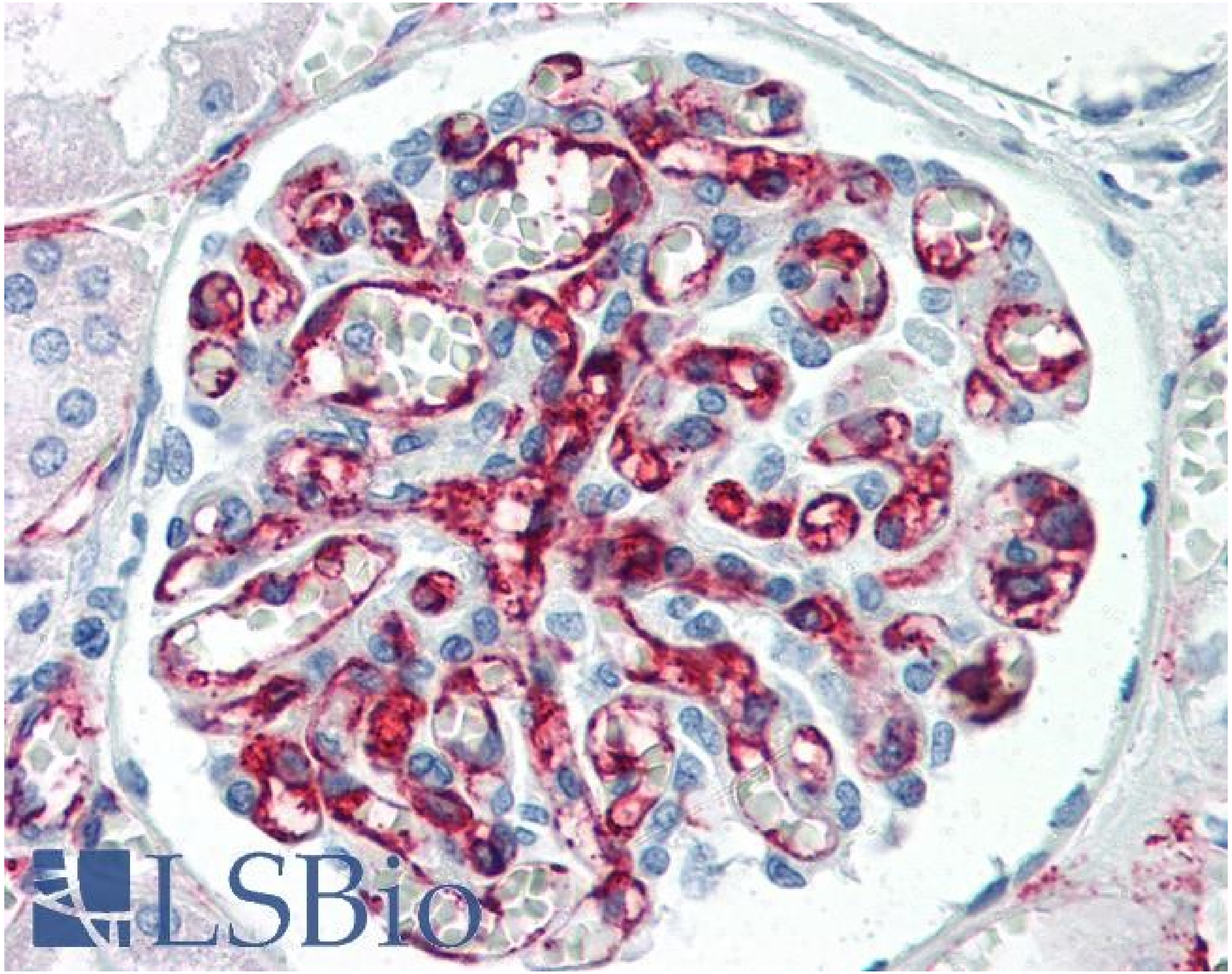

# CD45

**Immunohistochemistry (Formalin/PFA-fixed paraffin-embedded sections) - Anti-CD45 antibody [EP322Y] - (ab214437)**
Unpurified ab40763 showing negative staining in Normal kidney tissue. This data was developed using the same antibody clone in a different buffer formulation containing PBS, BSA, glycerol, and sodium azide (ab40763).

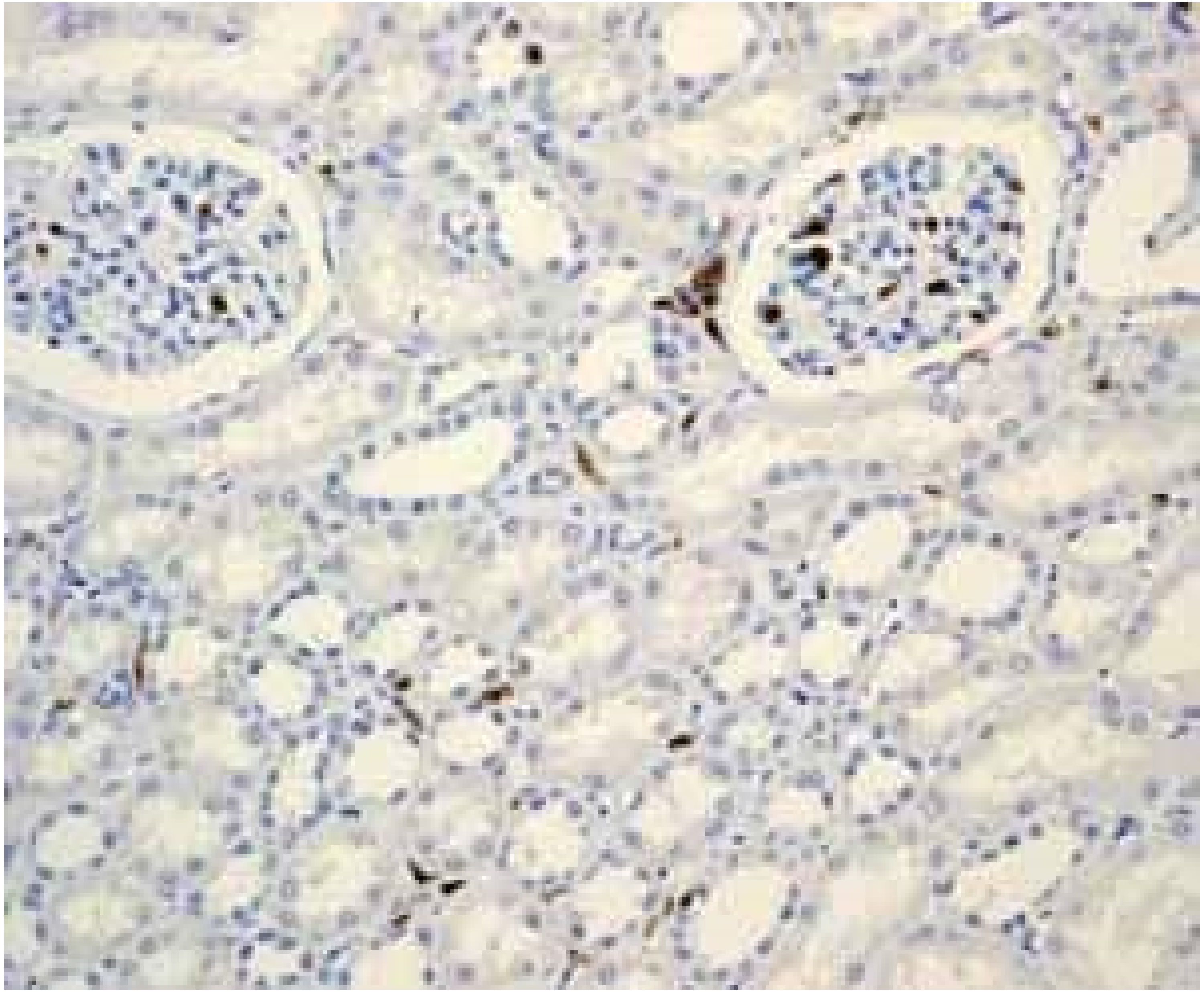

### CD45 (PTPRC)

**PMID: 34039664** – a transmembrane glycoprotein, expressed on almost all hematopoietic cells except for mature erythrocytes, and is an essential regulator of T and B cell antigen receptor-mediated activation

### CD68 (Macrosialin)

**PMID: 18405323**
Highly expressed by blood monocytes and tissue macrophages. Also expressed in lymphocytes, fibroblasts and endothelial cells.

### Collagen IV

#### Human Protein Atlas

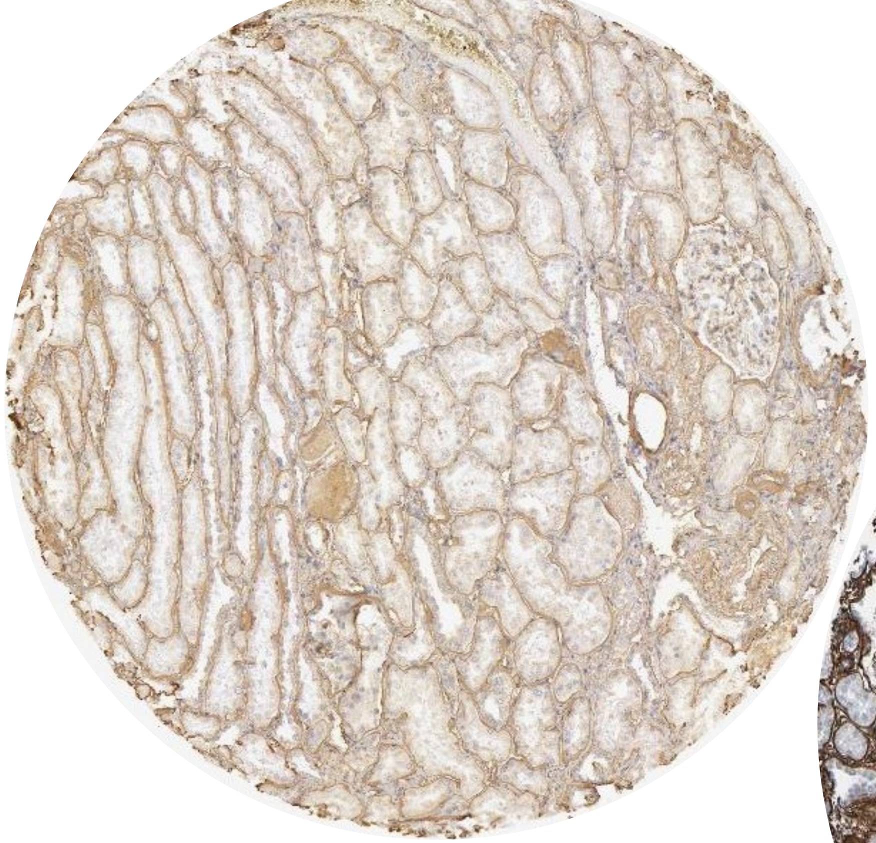

COL4A1

COL4A2

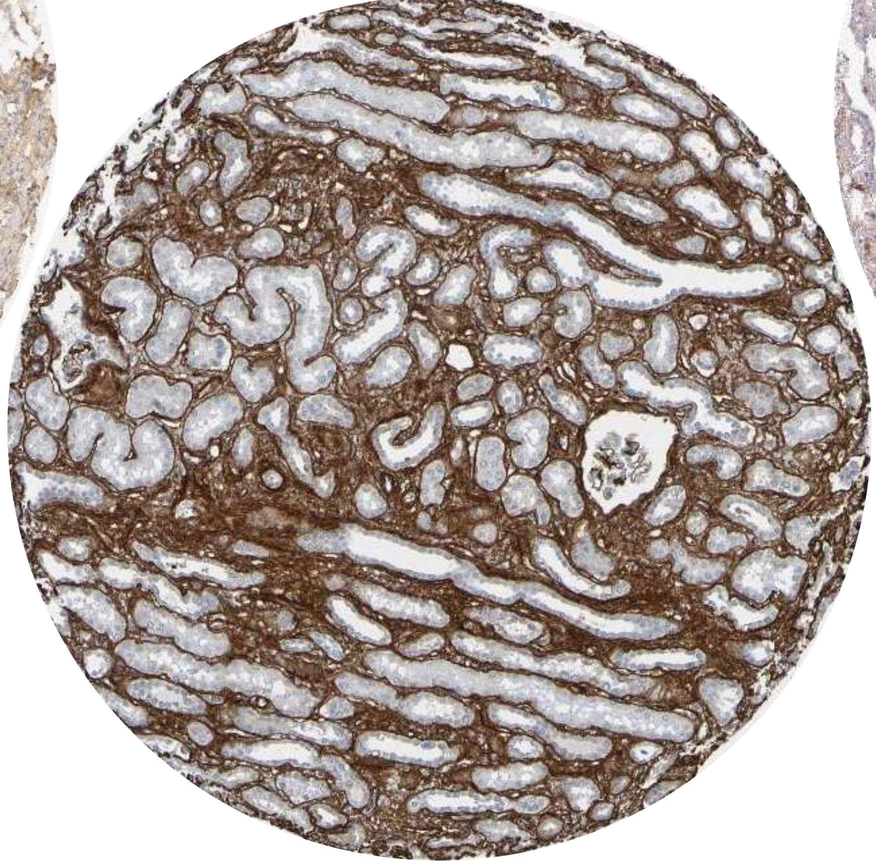

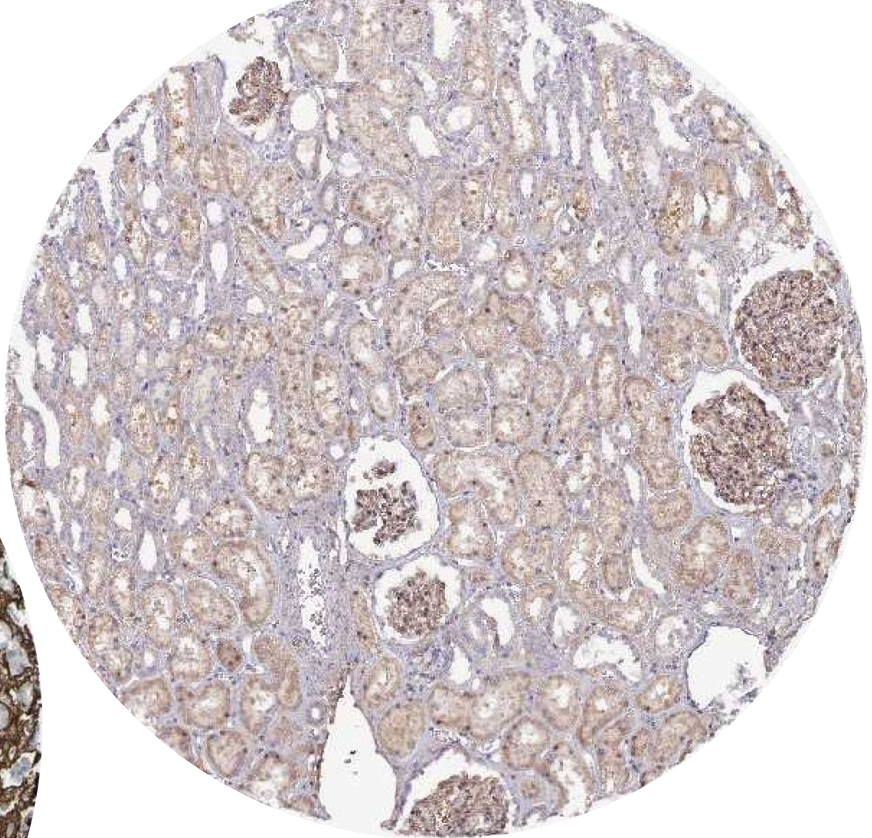

COL4A3

### Collagen IV

#### PMID 10854213

**Figure 2.** Immunohistochemical distribution of type IV collagen α1, α3, and α5 chains in the kidney of controls (a–c) and X-linked AS male patients (d–i).

Controls: a, anti-α1(IV) antibodies strongly stained the mesangial matrix, Bowman's capsule, and tubular basement membranes and reacted faintly with the GBM; b, anti-α3(IV) antibodies stained the GBM; and c, with anti-α5(IV) antibodies, the GBM and also the Bowman's capsule are strongly stained.

AS patients: d, strong and diffuse labeling of the GBM is observed with anti-α1(IV) antibodies; e, no GBM labeling is observed with anti-α3(IV) antibodies but faint staining of two podocytes (arrow) is seen (patient 1); f, absence of GBM staining with anti-α5(IV) antibodies; g–i, after amplification of the α3(IV) signal, strong podocyte labeling is observed (arrow) (patients 1, 2, and 5). Magnification, ×150 (a–e, g and h), ×180 (f), ×240 (i).

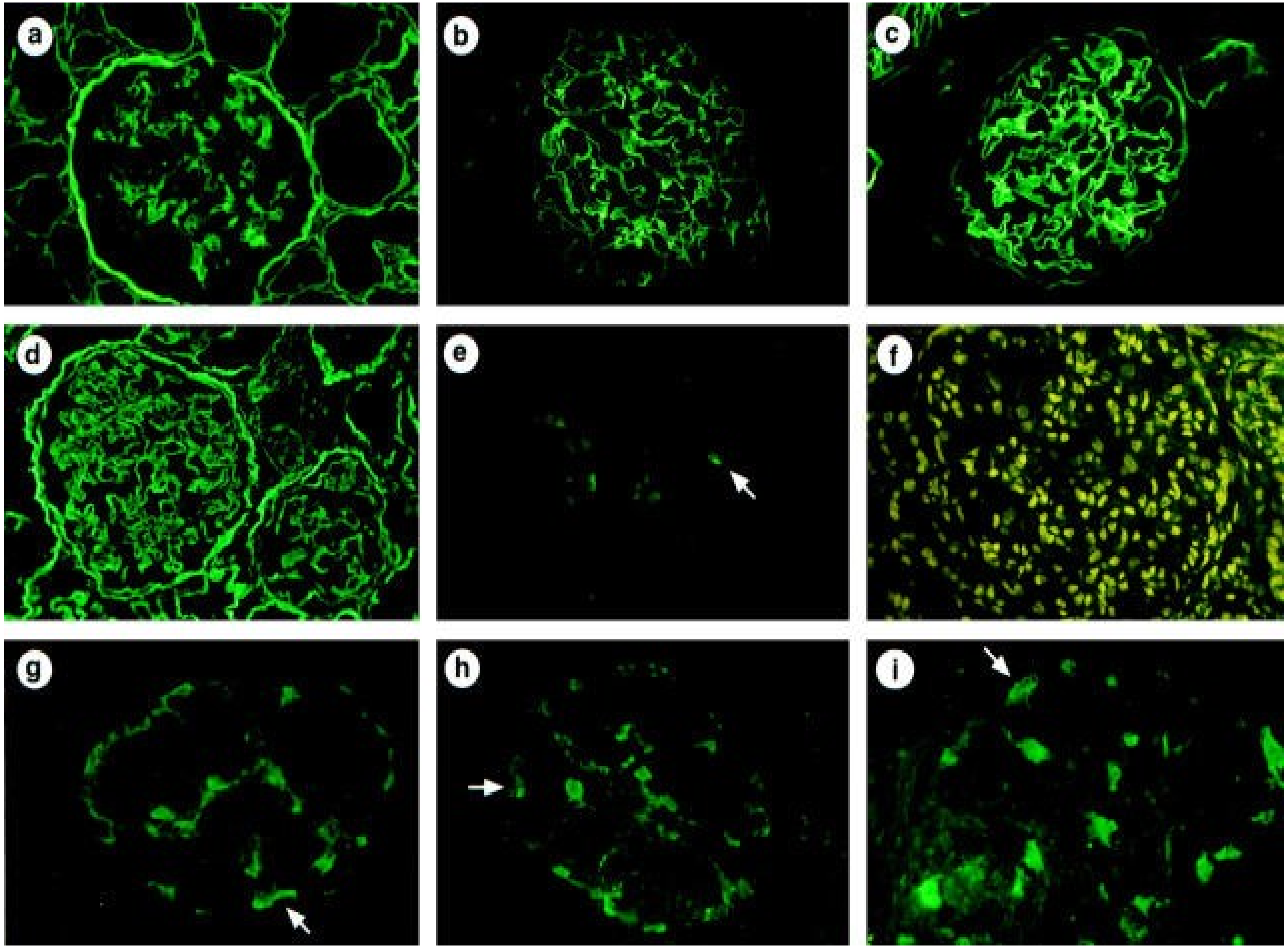

### β-catenin (CTNNB1)

**Human Protein Atlas**

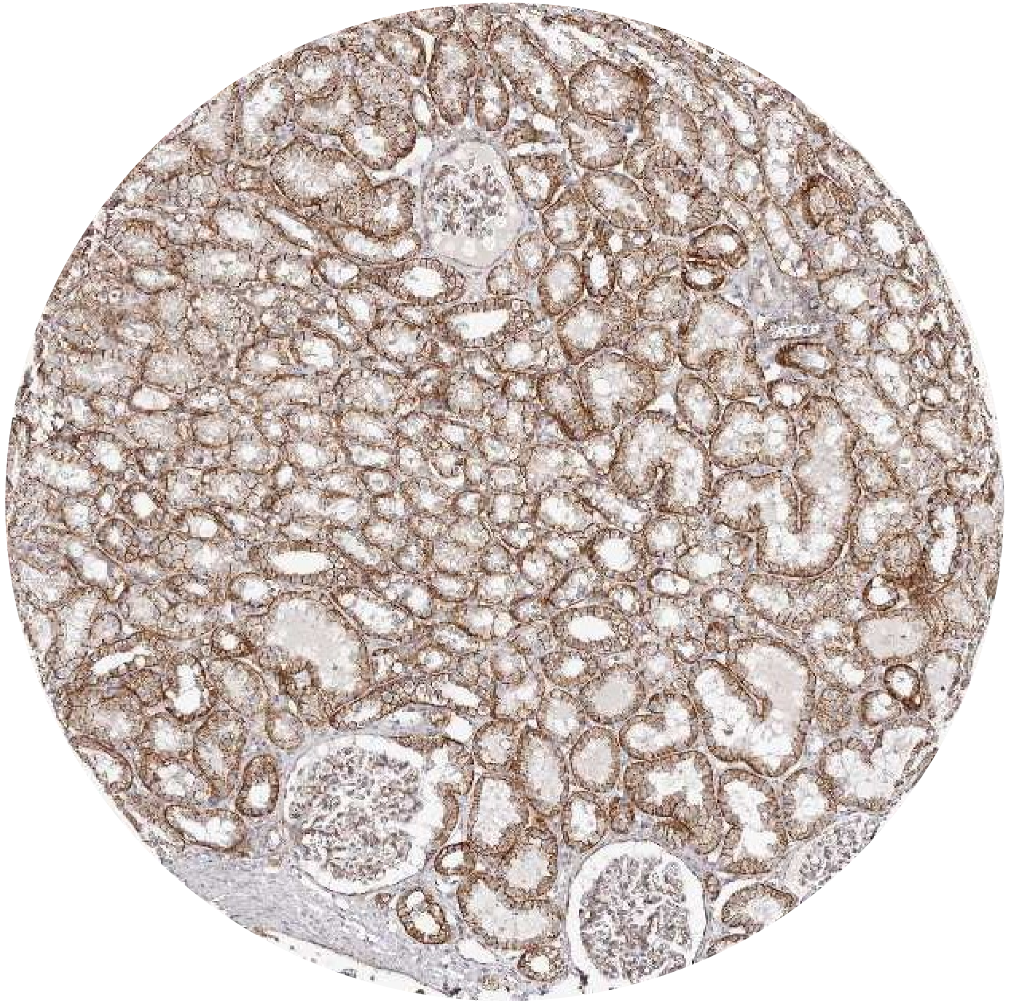

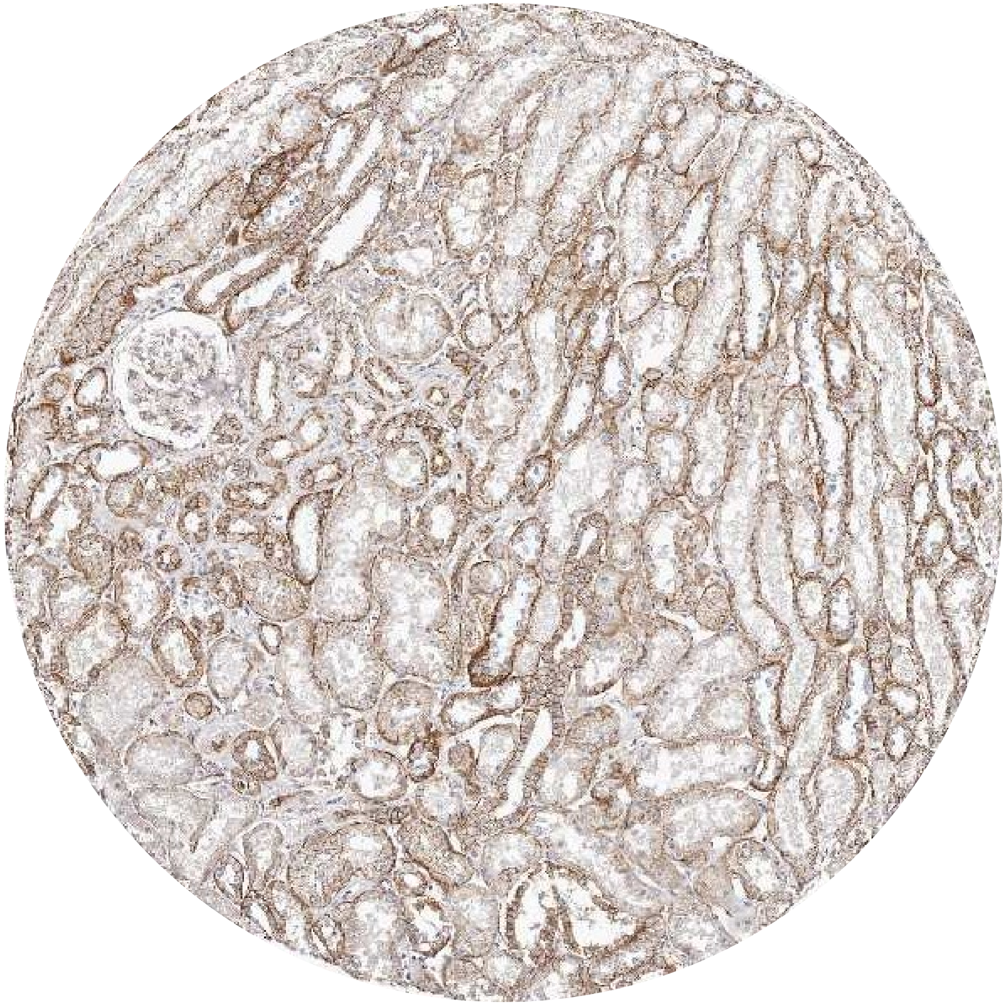

### β-catenin (CTNNB1)

**PMID: 31217358**

β-catenin is expressed in:
- Proximal tubules
- Thin ascending limb
- Thick ascending limb
- Distal convoluted tubules
- Collecting duct

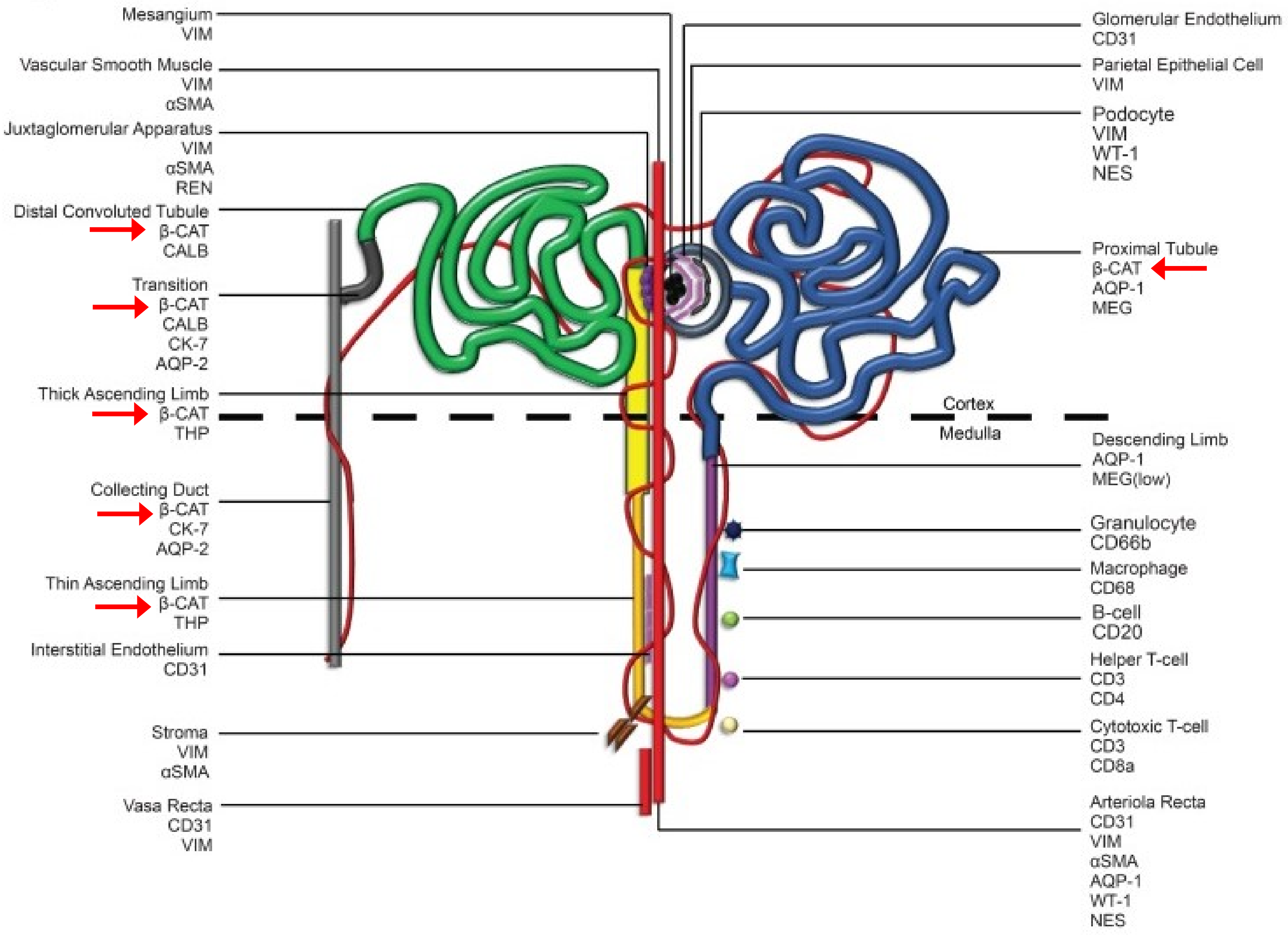

### β-catenin (CTNNB1)

**PMID:**

**Figure 1. A** Representative micrographs of B7-1 RNAscope staining and β-catenin immunochemistry staining in human kidney cortical tissue from CKD patients with lupus nephritis class III (LN), and the healthy control was from paracancerous tissue. Arrows indicate positive staining. Bar = 50 μm.

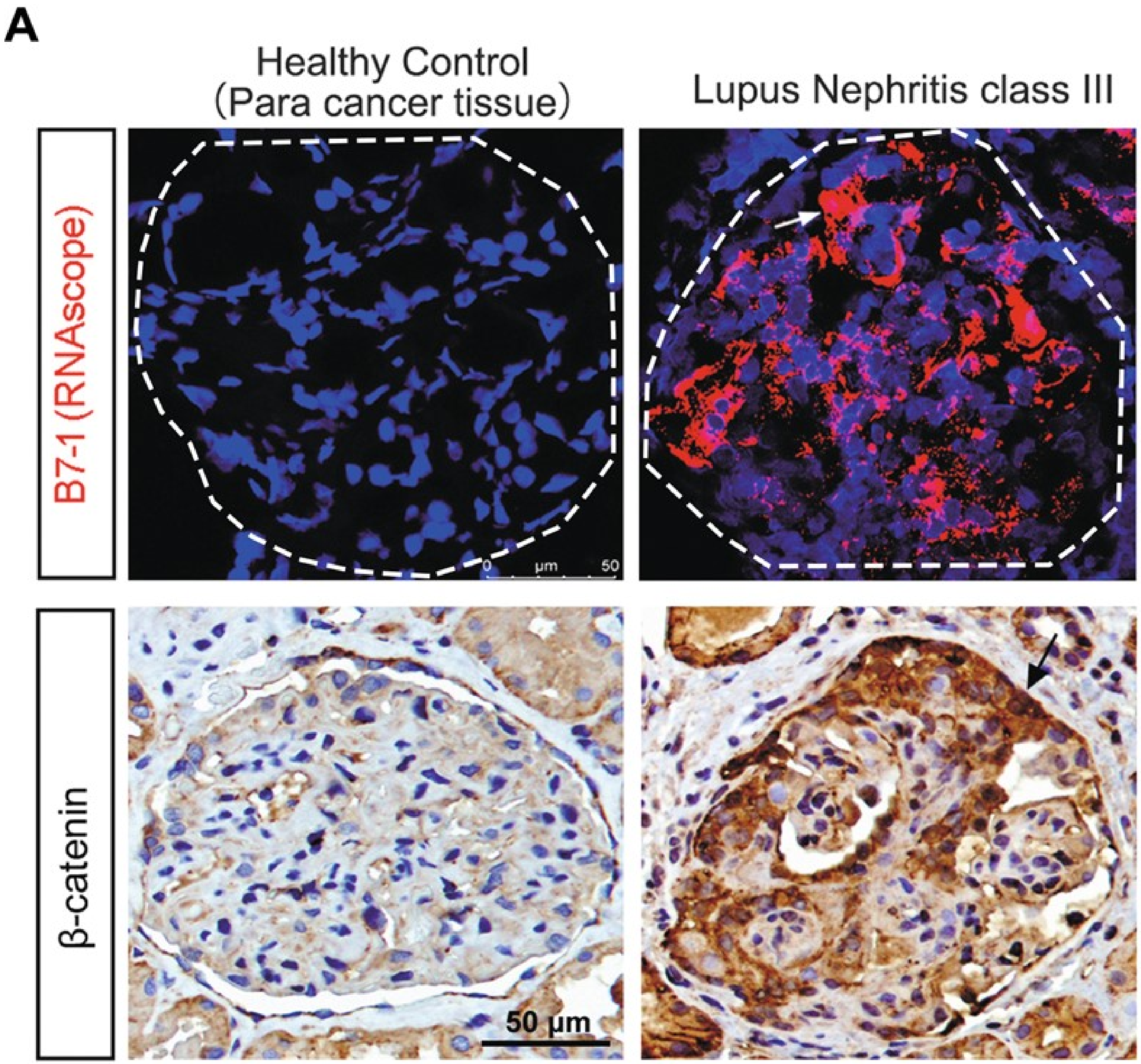


- Biopsy from CKD, healthy control from nephrectomy (malignancy)
- Unclear tissue preservation and staining process.
- Likely one or both monoclonal Abs to β-catenin (BD, 610154 OR Abclonal, A19657)

### β-catenin (CTNNB1)

**PMID: 24465439**

**Figure 1.** The protein levels of β-catenin in LN and control kidney tissues. The IHC results of β-catenin in LN and control kidney tissues(A and B). The IHC results showed that the staining of β-catenin was significantly stronger in lupus nephritis samples compared with controls (the levels of β-catenin were 0.467±0.500 vs. 0.093±0.129, respectively, $p<0.01$.) (C). Quantification of western blotting revealed significantly increased β-catenin in 20 renal biopsy specimens compared with 8 control renal specimens (D) (1.45±0.11 vs. 0.77±0.21, $p<0.05$).

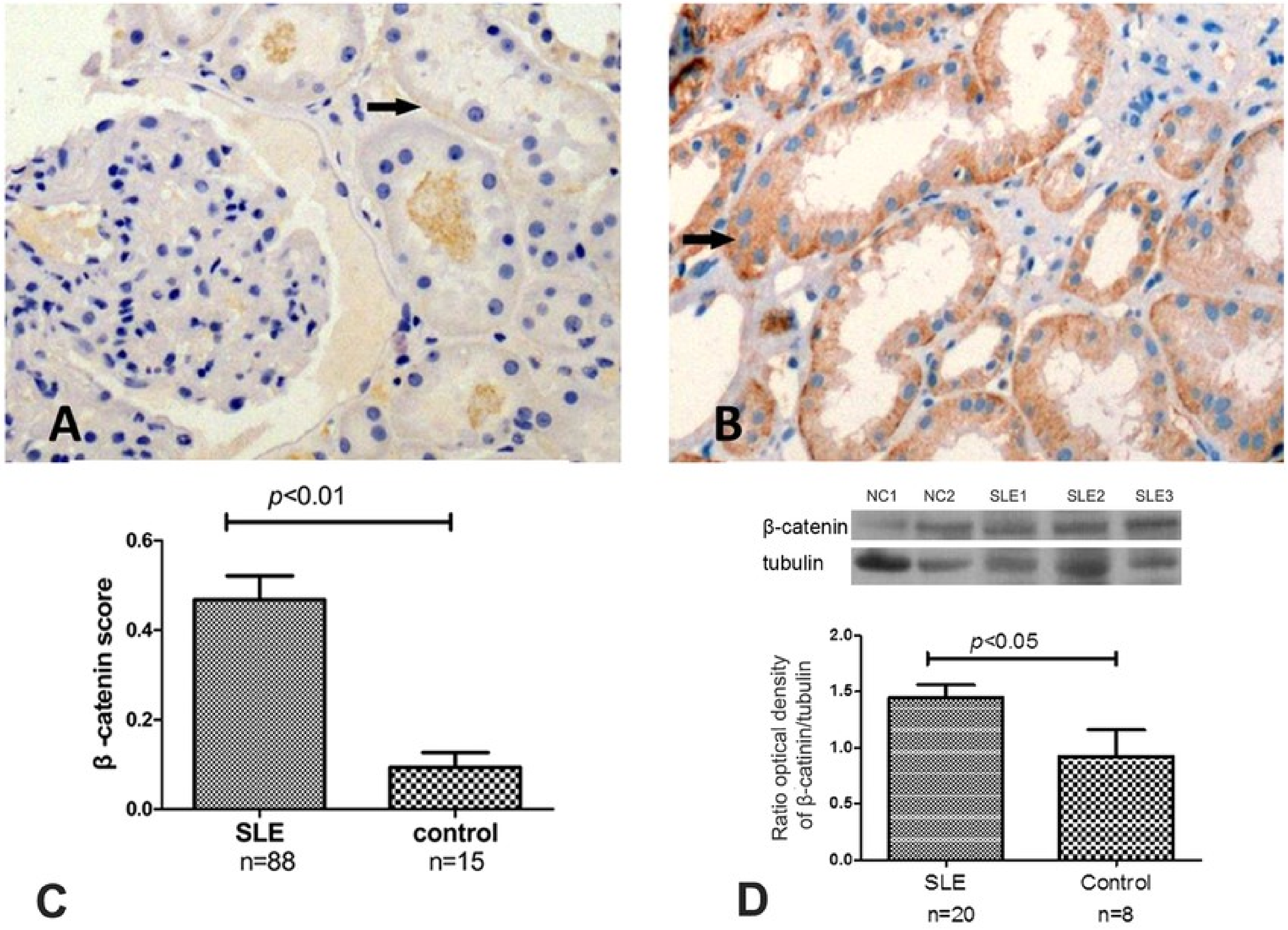


- Tissue from biopsy/nephrectomy (controls)
- FFPE preservation
- Citrate, microwave antigen retrieval
- anti-β-catenin antibody (Millipore) but unspecified mono or polyclonal or the clone

### CD183 (CXCR3)

**PMID: 12782716**

- Isoform 1 (Transmembrane)
  - ➢ Receptor for the C-X-C chemokine CXCL9, CXCL10 and CXCL11 and mediates the proliferation, survival and angiogenic activity of human mesangial cells (HMC) through a heterotrimeric G-protein signaling pathway
  - ➢ Binds to CCL21. Probably promotes cell chemotaxis response

- Isoform 2 (Transmembrane)
  - ➢ Receptor for the C-X-C chemokine CXCL4 and also mediates the inhibitory activities of CXCL9, CXCL10 and CXCL11 on the proliferation, survival and angiogenic activity of human microvascular endothelial cells (HMVEC) through a cAMP-mediated signaling pathway
  - ➢ Does not promote cell chemotaxis response. Interaction with CXCL4 or CXCL10 leads to activation of the p38MAPK pathway and contributes to inhibition of angiogenesis. Overexpression in renal cancer cells down-regulates expression of the anti-apoptotic protein HMOX1 and promotes apoptosis

- Isoform 3 (listed in Uniprot)
  - ➢ Mediates the activity of CXCL11.

**PMID: 31216755**

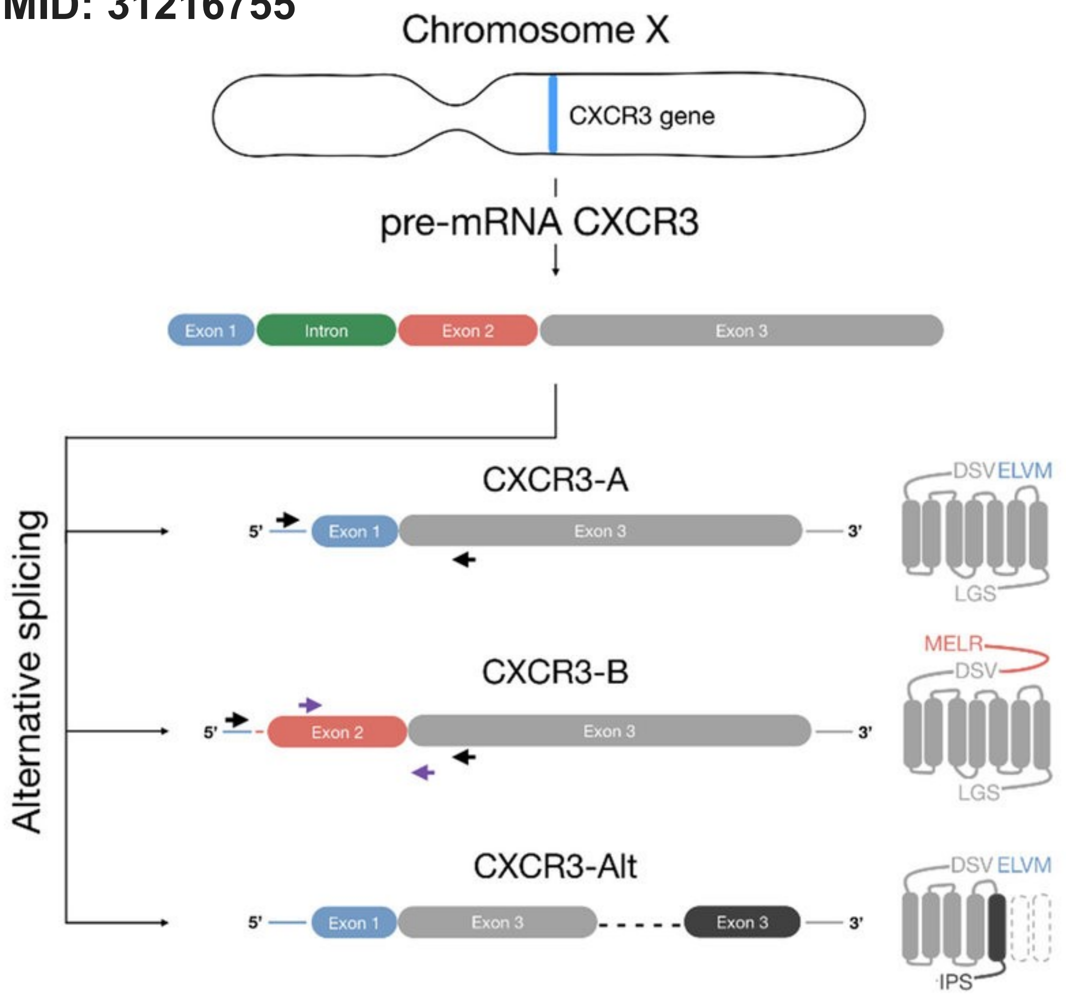

### CD183 (CXCR3)

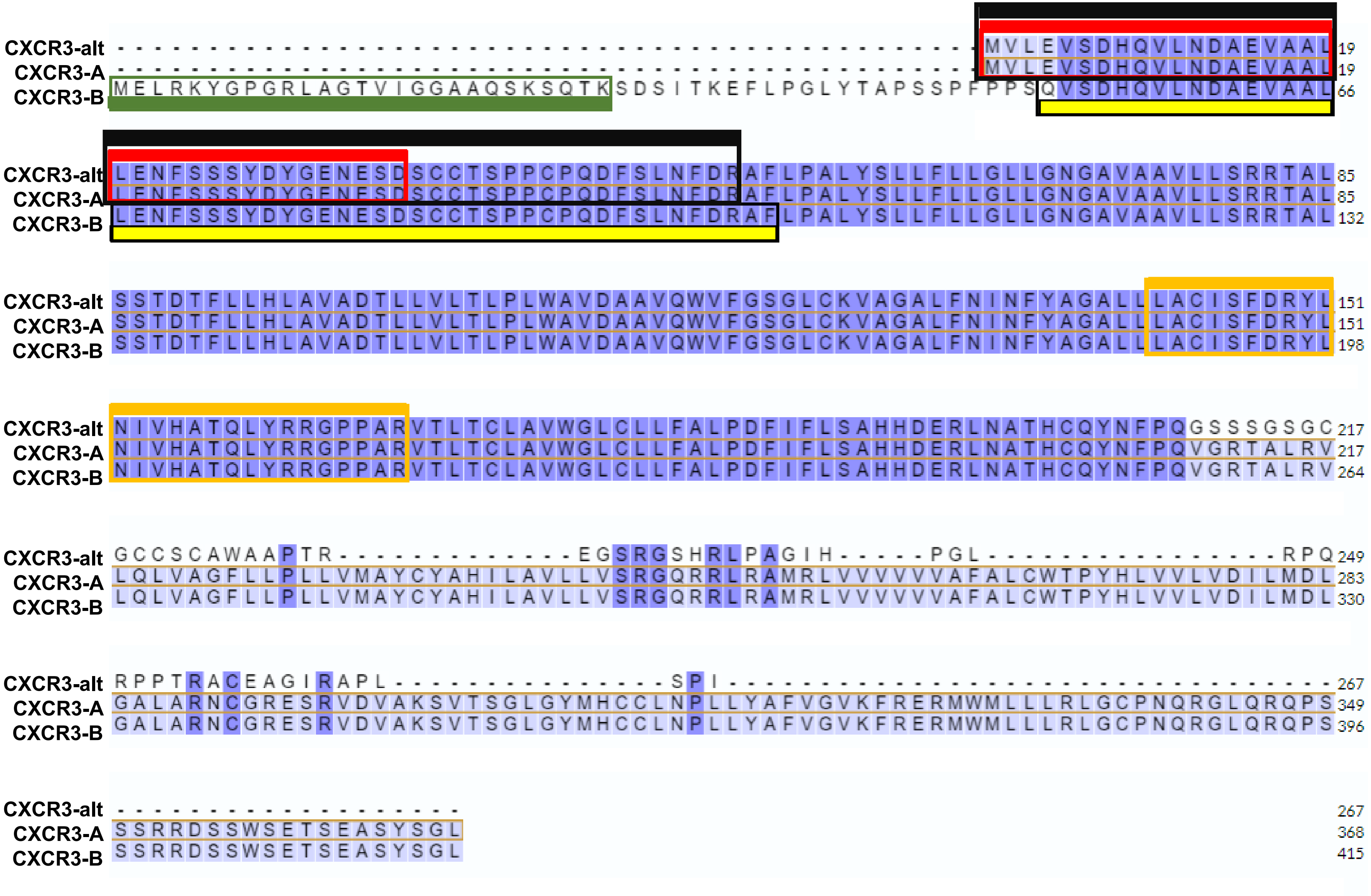


| | Antibody | Kidney Staining |
|---|---|---|
| (red) | 1C6 | None, inflammatory infiltrate |
| (yellow) | HPA | None |
| (green) | PL1 | None |
| (black) | 26756-1-AP | All tubules |
| (orange) | PA5-23679 | Tubules |

**CXCR3-A** (P49682-1)
**CXCR3-B** (P49682-2)
**CXCR3-alt** (P49682-3)

### CD183 (CXCR3)

**MAb: Anti-human CD183 (CXCR3), clone G025H7 (BioLegend)**
Monoclonal mouse anti-human, IgG1k
Antigen: human CXCR3 transfectants

Human peripheral blood lymphocytes were stained with CD3 APC and CXCR3 (clone G025H7) PE (top plot) or mouse IgG1 PE isotype control (bottom plot).

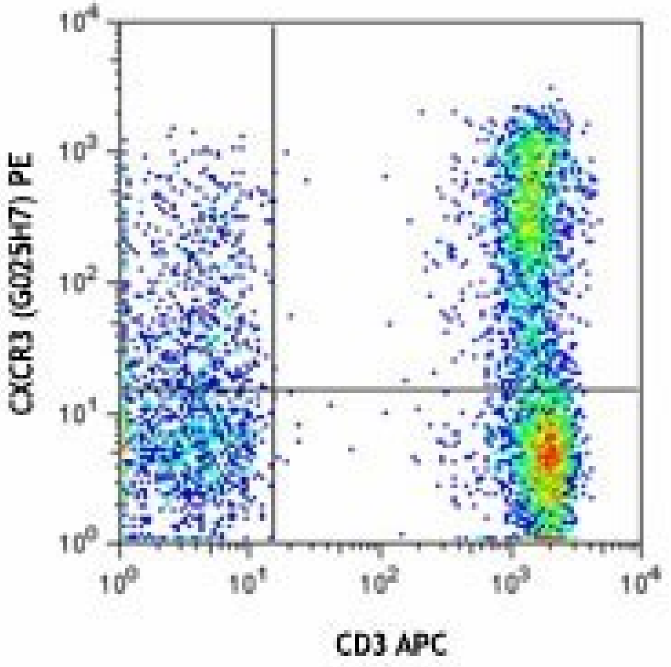


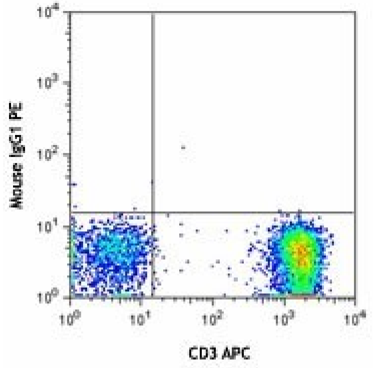

### CD183 (CXCR3)

#### PMID: 12782716

**Fig 1.** Identification, molecular characterization, and tissue distribution of CXCR3-B. (a) Structure of the human CXCR3 gene. Upper case, the exons; lower case, the intron; arrow, the CXCR3-B transcription start; underline, the CXCR3-B translation start site (ATG); light shading, the donor (AG/gt, common for both CXCR3-A and CXCR3-B) and acceptor (ag/GG, CXCR3-B; AG/GT, CXCR3-A) splice sites. The nested primers used for 5′ and 3′ RACE are boxed and the direction is pointed by the arrow. The 182-bp probe (recognizing CXCR3-B) and the 401-bp probe (recognizing both CXCR3-A and CXCR3-B) used for Northern blot analysis are evidenced in yellow and light green, respectively. (b) RACE performed on primary cultures of HMVEC. Lane 1, DNA ladder Marker VIII; lane 2, 5′ RACE product; lane 3, 3′ RACE product; lane 4, DNA ladder 1 Kb plus. (c) Schematic representation illustrating the generation of CXCR3-A and CXCR3-B mRNA through alternative splicing. (d) Multiple tissue Northern blot hybridization with the 182-bp probe (top) or the 401-bp probe (bottom). These sequence data are available from GenBank/EMBL/DDBJ under accession no. AF469635.

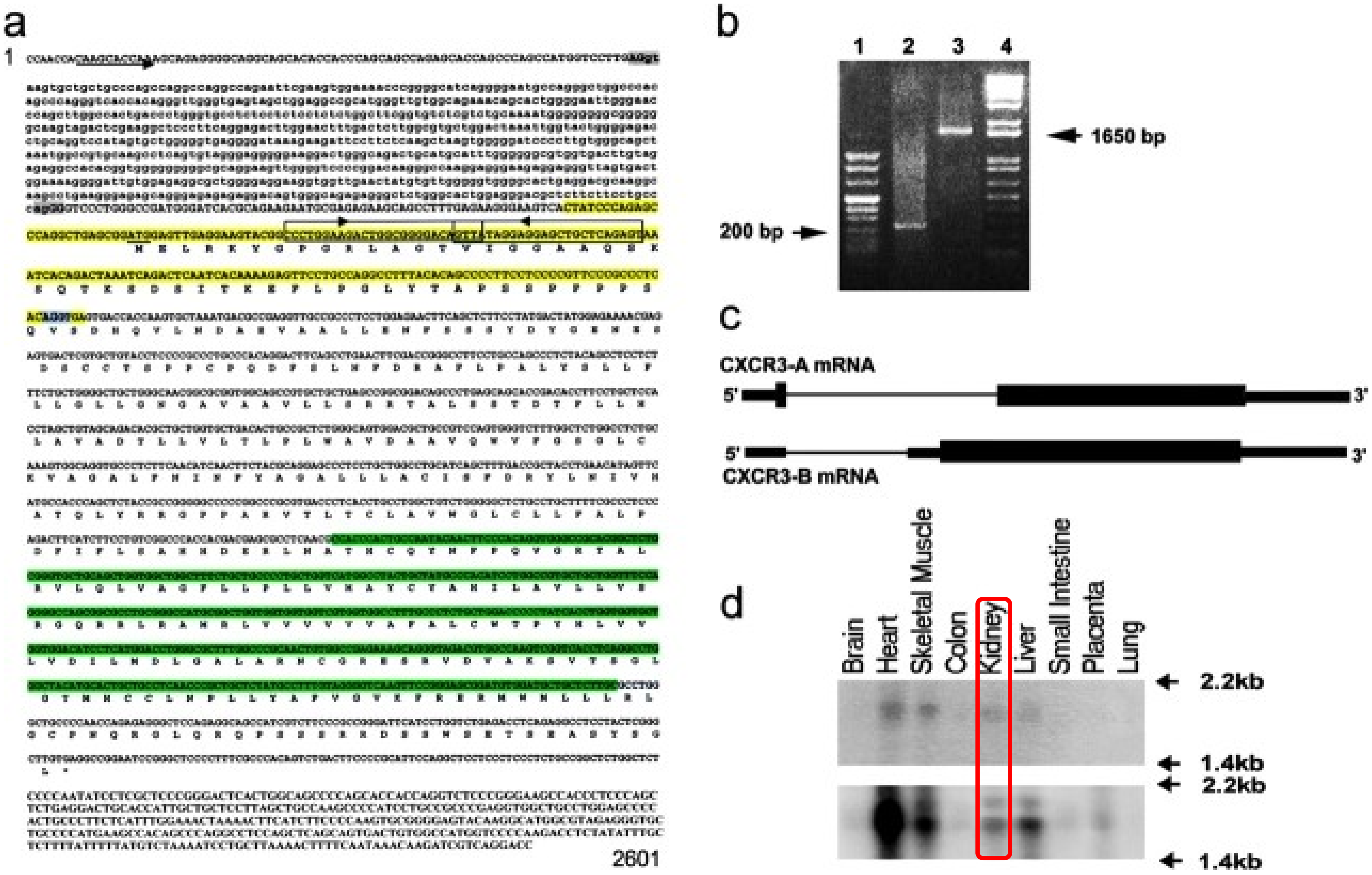

### CD183 (CXCR3)

#### PMID: 12517959

**Figure 1. The CXCR3 receptor is expressed in proximal tubules and IHKE-1 cells. A,** RT-PCR results for the mRNA expression of the human CXCR3 receptor in micro-dissected human proximal tubules (PT) gained from unaltered cortices of patients (with their consent) undergoing tumor nephrectomy and in the immortalized human proximal tubule cell line IHKE-1. In both tissues, a 379-bp fragment of the CXCR3 receptor was found with primers specific for the CXCR3 receptor. Amplified fragments were directly sequenced to confirm the presence of the CXCR3 receptor. In all experiments, a negative control (−) was included where 1) RNA was excluded, and 2) reverse transcriptase was omitted from the reverse transcriptase mixture. Amplification of GAPDH was used to confirm RNA integrity. **B,** Immunohistochemical stains (magnification, ×40) of a rat kidney showing the presence of the CXCR3 receptor in proximal tubules beneath the brush border membrane (upper panel). In negative controls (magnification, ×63; lower panel) the primary Ab was heat-denatured.

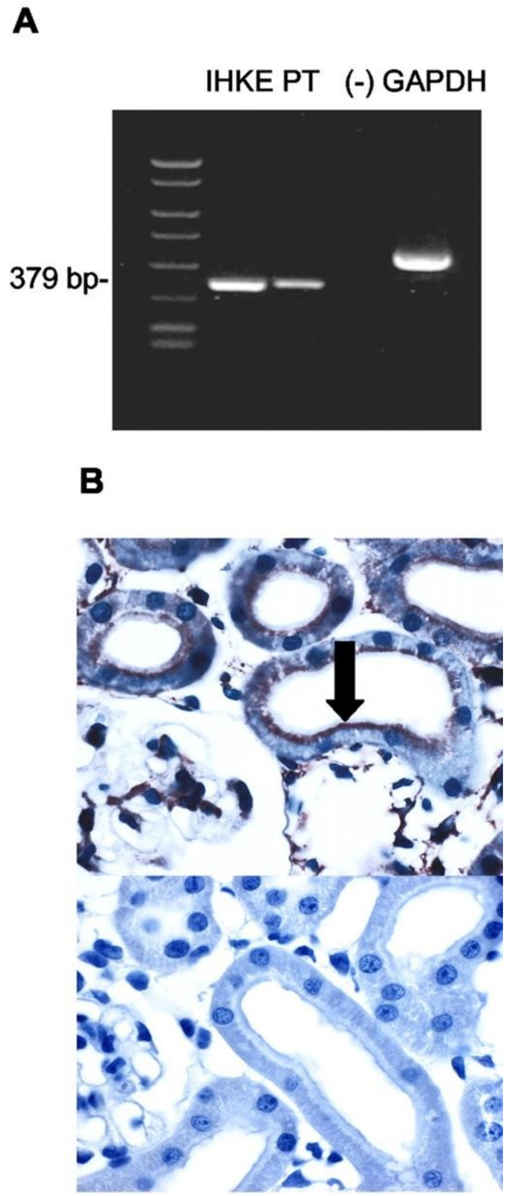


- IHKE-1 are immortalized human proximal tubular cells (10457080).
- The PCR used here identifies at least isoforms A and B:

  5' oligo:
  CCACCCACTGCCAATACAAC

  3' oligo: CGGAACTTGACCCCTACAAA
  Reversed: tttgtaggggtcaagttccg

### CD183 (CXCR3)

#### Human Protein Atlas

- One pAb – not detected
- Atlas Antibodies, rabbit pAb (HPA045942), IgG
    - Medium consistency between Ab staining and RNA expression data
    - No bands on Western
    - Single peak corresponding to interaction with its own antigen on Protein Array

- Validation:
    - IHC- staining mainly consistent with RNA expression across 45 tissues.
    - ICC- no data reported.
    - WB- uncertain: No bands detected in analysis including a standard sample panel
    - Protein array- supported: single peak corresponding to interaction with its own antigen

ICC

N/A

IHC

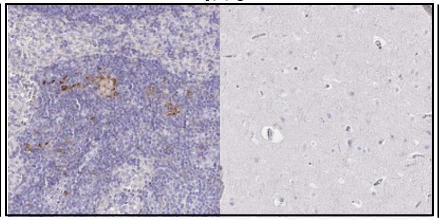

High expression Low expression

WB

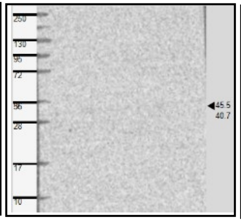

Protein Array

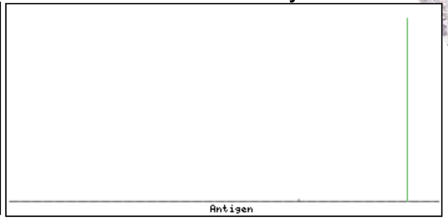


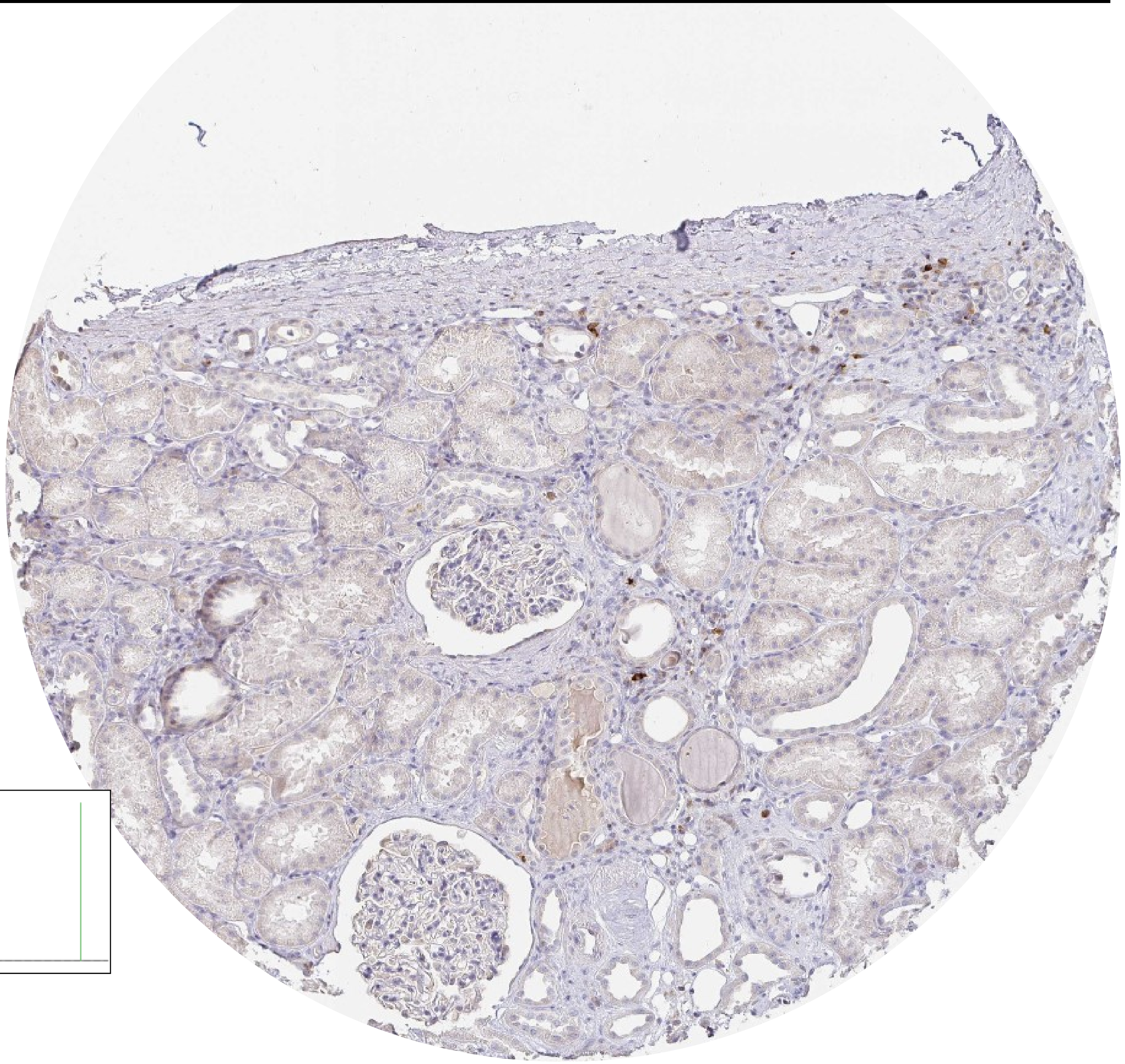

### CD183 (CXCR3)

**PMID: 16421159**

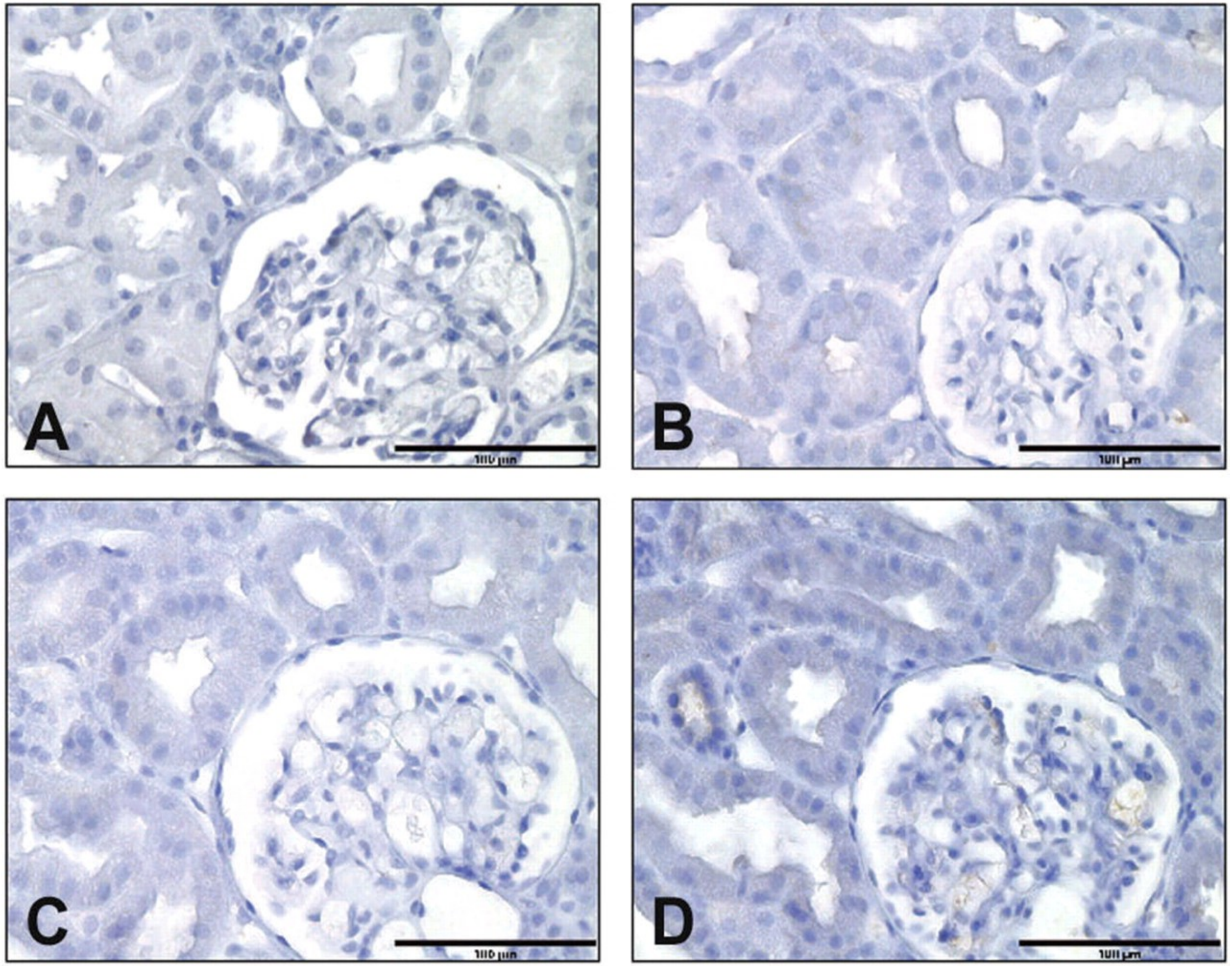


**Fig. 2.** Immunohistochemistry using antibodies against CXCR3 (A), CD3 ( B ), CD4 ( C ) and CD8 ( D ) performed on consecutive sections of a normal renal graft biopsy (original magnification ×400). Note the low number of positive cells

### CD183 (CXCR3)

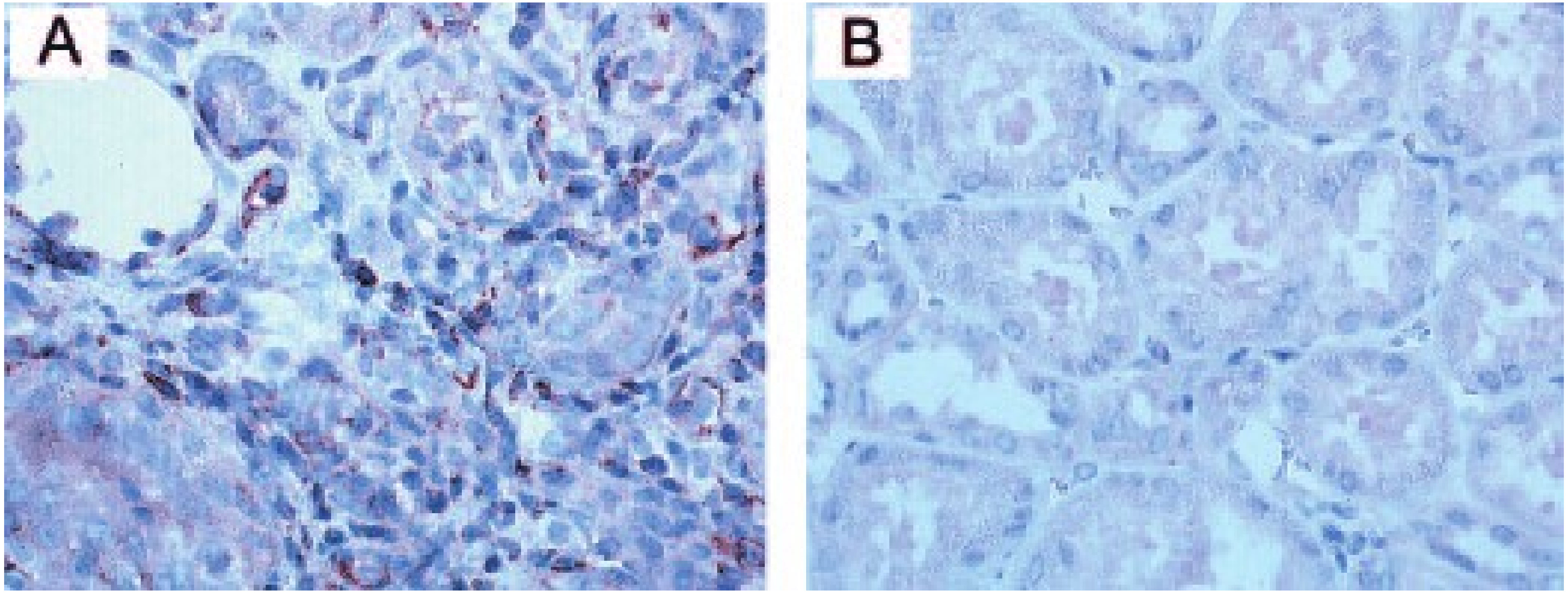


**PMID 15857922**

**Figure 5.** Immunohistochemical determination of CXCR3-positive cells in a renal biopsy of a patient with acute rejection (A) and normal control kidney specimen (B). Out of the series of AR patients, a representative biopsy specimen is shown (A) and compared with a control biopsy specimen (B). Paraffin sections were stained with a polyclonal antibody specific for human CXCR3 and counterstained as described in the Materials and Methods section (PMID 14742268).

### CD183 (CXCR3)

**PMID 11134180**

**Fig 1.** CXCR3 protein expression by a small proportion of microvascular endothelial cells in normal and pathological human tissues. (a) CXCR3 immunostaining (red) of endothelial and smooth muscle cells in a small arteriola of human thymus and in some thymocytes of the subcapsular areas. ×250. (b) CXCR3 immunostaining (red) of endothelial cells in small vessels surrounding thyroid follicles in a biopsy specimen from a patient suffering from Graves' disease. ×250. (c) CXCR3 expression (red) of endothelial and smooth muscle cells in an arteriola of normal liver. × 400. (d) CXCR3 immunostaining (red) of endothelial cells from small vessels and of infiltrating inflammatory cells present in a biopsy specimen from a patient with active cirrhosis. ×250. (e) CXCR3 expression (red) in a normal human kidney. Signal is clearly visible only at level of vascular smooth muscle cells. ×100. (f) CXCR3 immunostaining (red) in the biopsy specimen of a patient suffering from glomerulonephritis. Signal is visible in both vascular smooth muscle cells and endothelial cells. ×100. (g and h) Double-label immunohistochemistry for CXCR3 (red) and vWF (bluish-gray) showing CXCR3 expression by endothelial cells from some microvessels in the normal part of a human kidney (×100) and from a much higher number of microvessels present in its neoplastic counterpart (×100), respectively. Inset: Higher-power magnification of some microvessels in the normal part of the kidney. ×400

49801.111 anti-CXCR3 mAb

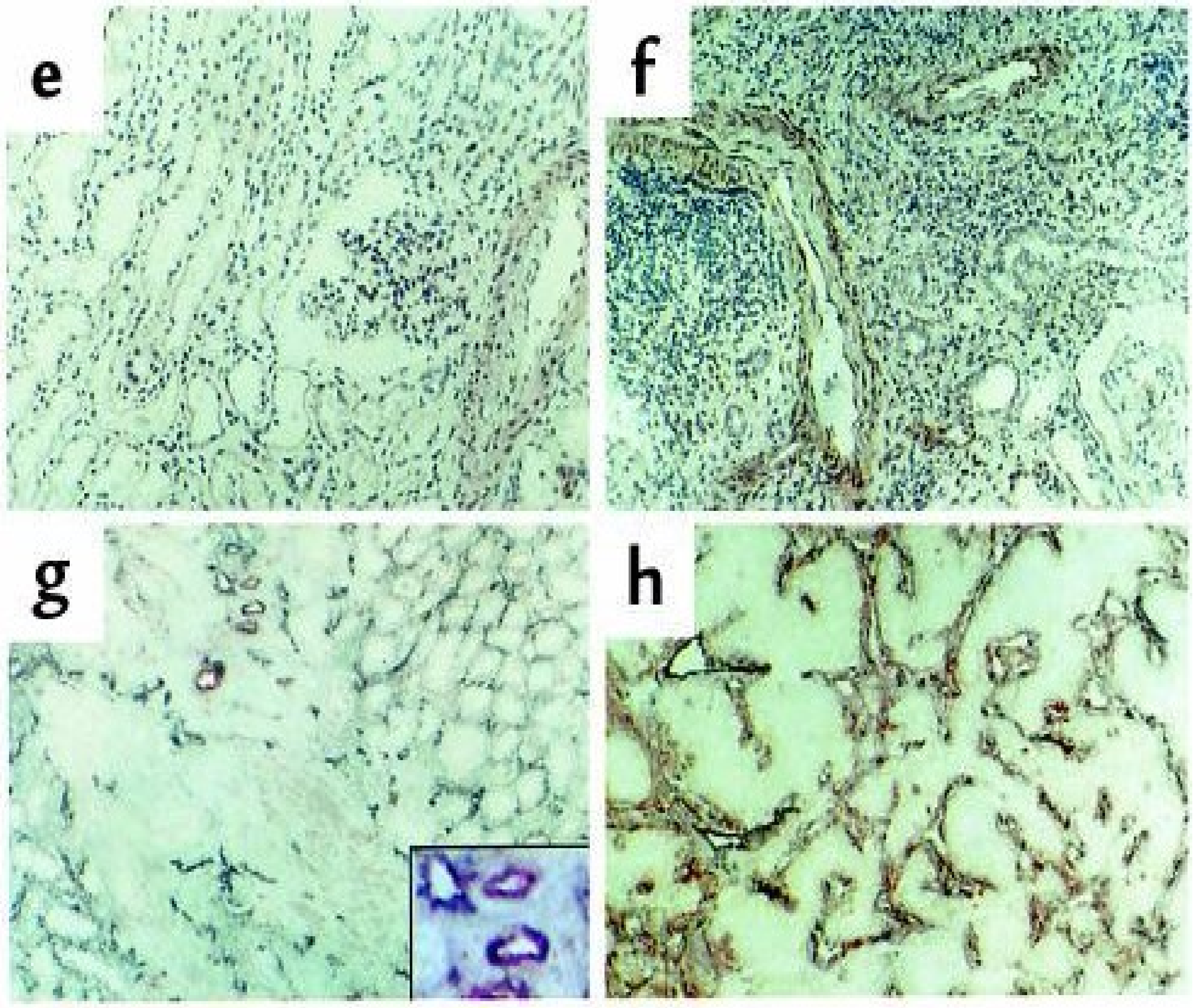

### CD183 (CXCR3)

**PMID: 12782716**

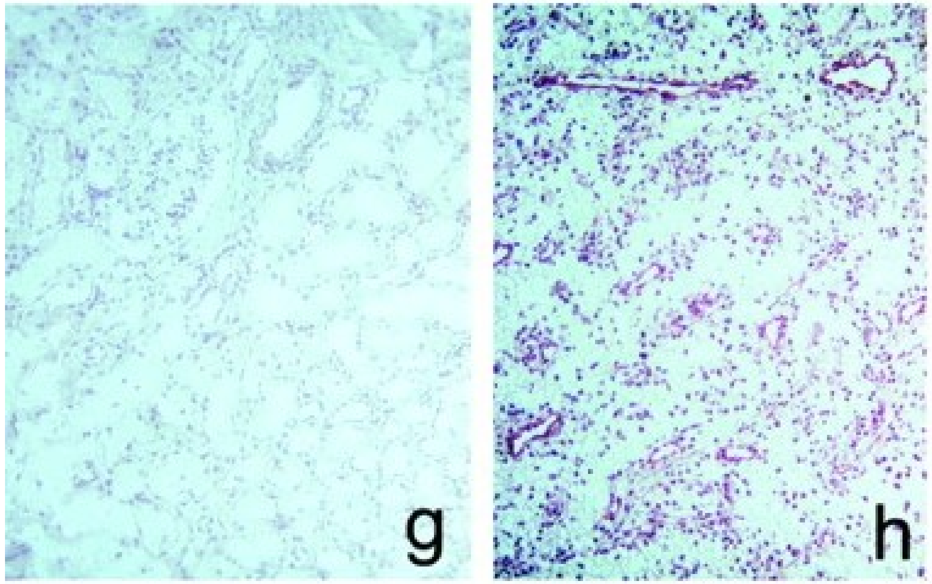


**Fig 6.** Detection of CXCR3-B protein expression by different types of cell cultures and by endothelial cells of human neoplastic tissues by CXCR3-B–specific mAbs. (a) Absence of reactivity in mock transfectants stained with an anti-CXCR3-B mAb. ×100. (b) Intense staining of CXCR3-B transfectants with an anti–CXCR3-B mAb. ×100. (c) Absence of reactivity in CXCR3-A transfectants stained with an anti–CXCR3-B mAb. ×100. (d) Absence of reactivity in primary cultures of HMC stained with an anti–CXCR3-B mAb. ×100. (e) Positive staining of primary cultures of HMVEC as well as the ACHN cell line (f) with an anti–CXCR3-B mAb. ×100. (g) Absence of reactivity in normal human renal tissue stained with PL1 anti–CXCR3-B mAb. ×10. (h) Staining with the same mAb of endothelial cells in a specimen of renal cell carcinoma. (i) Staining of both endothelial and tumor cells with the anti-CXCR3 mAb 49801.111 tested in an adjacent section. (j) Absence of reactivity in an adjacent section stained with an isotype-matched control mAb. (k and l) Reactivity of endothelial cells from the same renal carcinoma specimen before and after adsorption of PL1 mAb with the peptide used for mouse immunization. (m) Reactivity with PL1 mAb of endothelial cells from a group of vessels in a renal cell carcinoma specimen as detected at a higher power magnification. ×250. (n) Staining of an adjacent section with PL2 anti–CXCR3-B mAb. (o) Double label immunohistochemistry for CXCR3-B (red) and vWf (blue-gray), showing costaining (brown). (p) Absence of CXCR3-B reactivity in normal human lung tissue stained with PL1 anti–CXCR3-B mAb. (q) Staining with the same mAb of endothelial cells in a specimen of NSCLC. (r) Staining of both endothelial cells and other cell types with the anti-CXCR3 mAb 49801.111 in an adjacent section

### CD183 (CXCR3)

**PMID 10589690**

**Figure 1. CXCR3 expression in normal kidney and in non-proliferative glomerulopathies**. (A) CXCR3 expression in normal kidney. Section was immuno-stained with anti-CXCR3 monoclonal antibody (mAb), using the avidin-biotin-peroxidase method, and the 3-amino-9-ethylcarbazole substrate (red color). A slight signal is also present over some cells in the glomerulus. (B) High-power magnification of a glomerulus showing slight or no signal for CXCR3. (C) CXCR3 expression in non-proliferative glomerulopathies. Infiltrating cells stain positive for CXCR3. (D) High-power magnification of a glomerulus showing slight or no signal for CXCR3; the arrow points to some mononuclear infiltrating inflammatory cells in the interstitium. (E) A section adjacent to C was immunostained with an isotype control-matched mAb; no signal is visible. (F) **CXCR3 expression in the gut of a patient suffering from Crohn's disease. Inflammatory cells are immunostained with anti-CXCR3 mAb.** Magnification: ɤ 100 in A, C, E, and F; ɤ 250 in B and D.

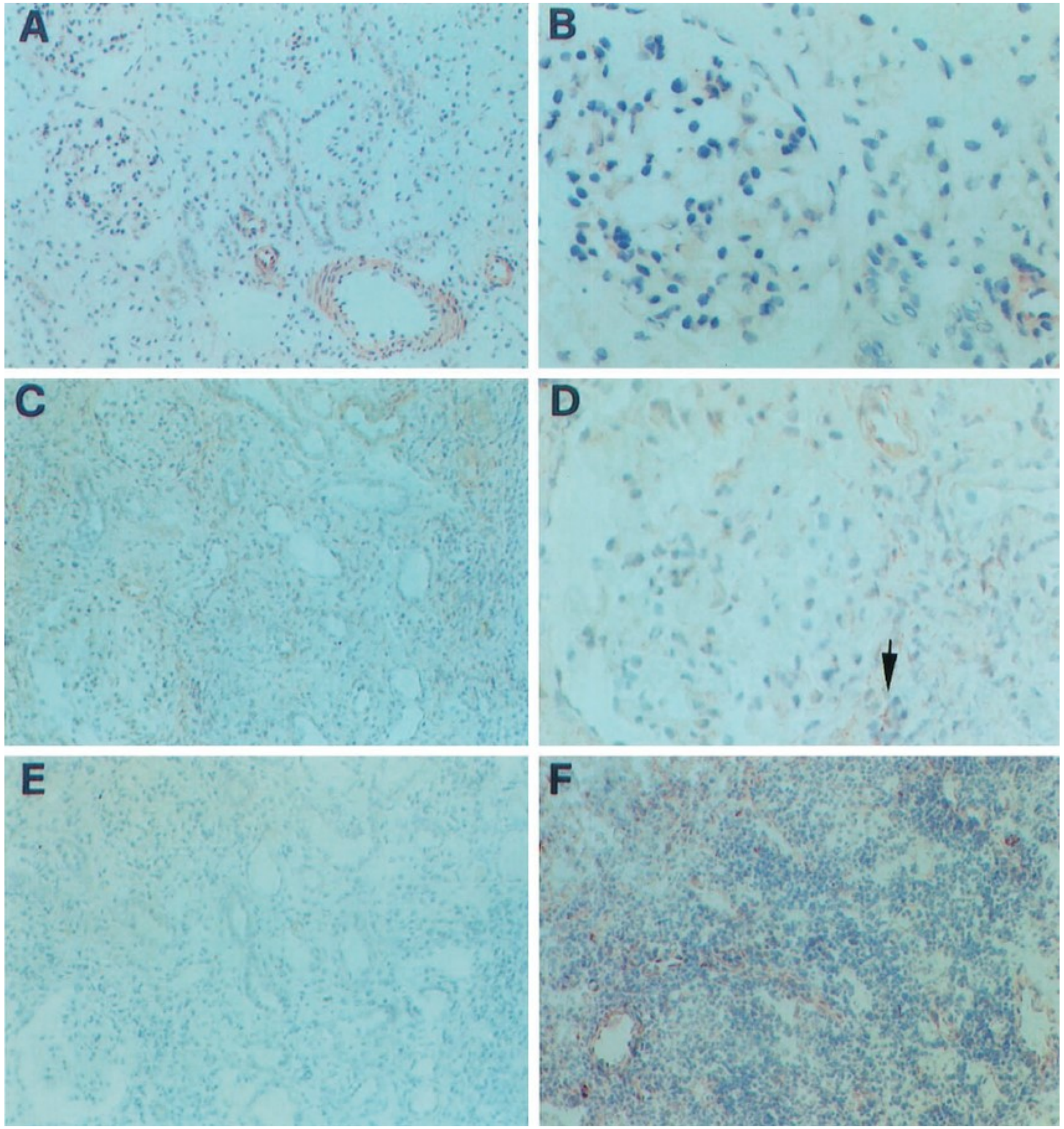

### CD183 (CXCR3)

**PMID: 19116922**

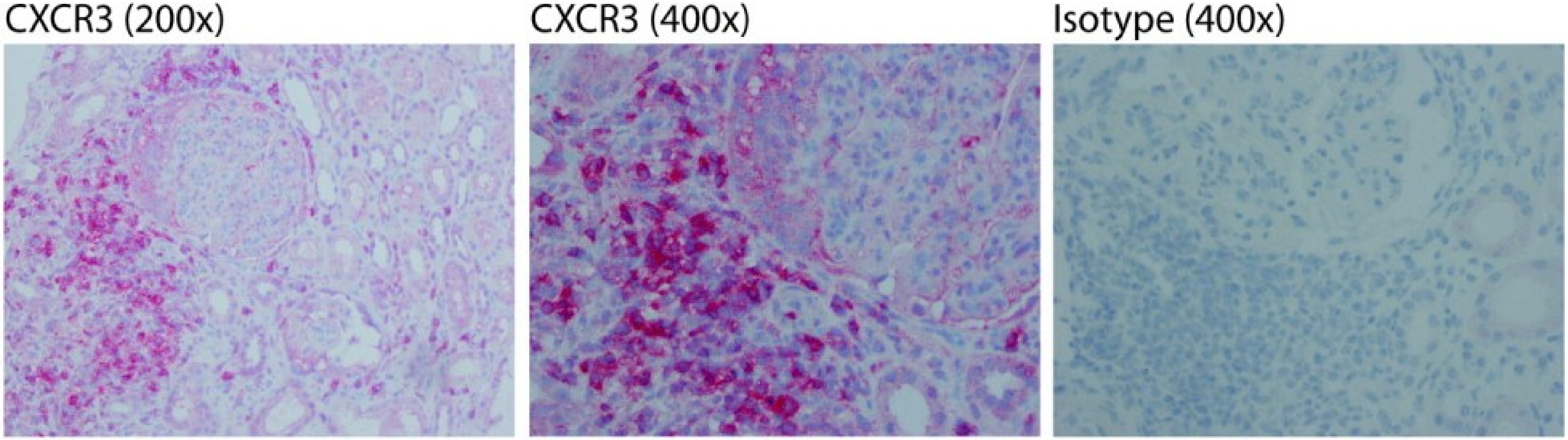


**Fig. 1.** Interstitial CXCR3 and CXCL10 expression in kidney biopsy tissues from patients with lupus nephritis. A, Typical periglomerular accumulation of CXCR3-expressing infiltrating cells in a patient with type IV lupus nephritis. B, Expression of the chemokine CXCL10 in a patient with type IV lupus nephritis. Isotype controls are shown for comparison.

### CD183 (CXCR3)

**PMID: 30273603**

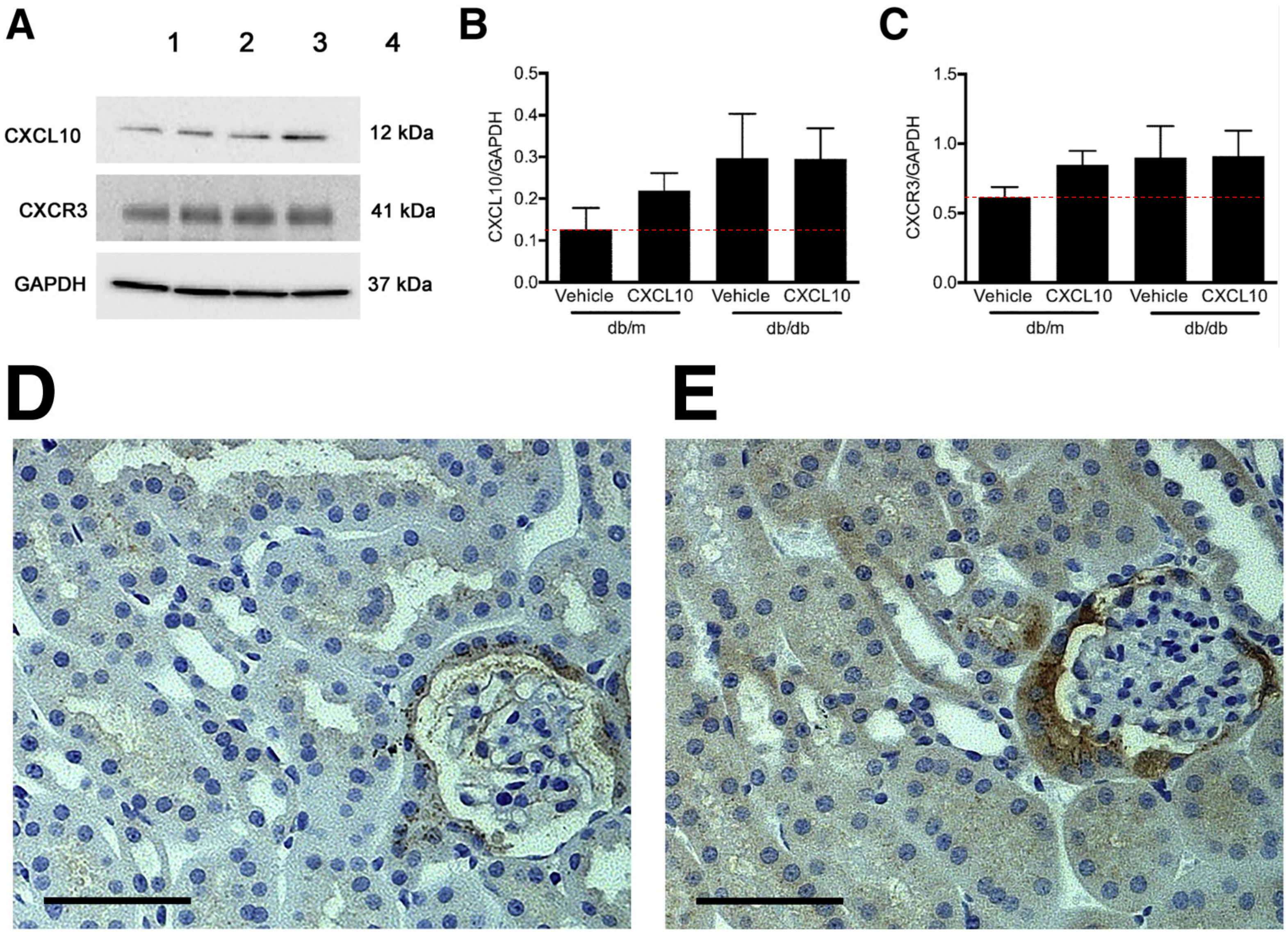


**Fig. 8.** A–C: Expression CXCL10 and CXCR3 in kidney was assessed by immunoblot (A) and quantified relative to glyceraldehyde-3-phosphate dehydrogenase (GAPDH; B and C). No difference among groups was noted for either CXCL10 (B) or CXCR3 (C). D and E: Immunohistochemistry showed expression of CXCR3 predominantly in the parietal epithelium of glomeruli in both nondiabetic db/m and diabetic db/db mice. Data are expressed as means ± SEM. Scale bar = 100 μm. Original magnification, ×160.

polyclonal anti-mouse CXCR3 (NBP2-12239)

### CD183 (CXCR3)

**PMID: 17671396**

**Figure 1.** The renal expression of IP-10 and CXCR3 was upregulated in diseased kidneys. **a** The number of IP-10-positive cells was increased in the renal interstitium after ureteral ligation which was not different from that observed in mice treated with anti-IP-10 antibody. **b** Transcripts of IP-10 were faintly detected in normal kidneys by real-time RT-PCR. In contrast, transcripts of IP-10 in diseased kidneys were upregulated. However, anti-IP-10 antibody treatment did not change the levels of IP-10 transcripts. **c** CXCR3-positive cells were hardly detected in a normal mouse kidney. **d** CXCR3-positive cells, detected in tubular epithelial cells (arrow), were increased on day 7. **e** Number of CXCR3-positive cells in the interstitium. IP-10 blockade increased the number of CXCR3-positive cells on day 7. **f** CXCR3 mRNA expression in diseased kidneys. IP-10 blockade increased the CXCR3 mRNA expression on day 4. Values are mean ± SEM. * $p < 0.05$ as compared with nonspecific IgG-treated mice.

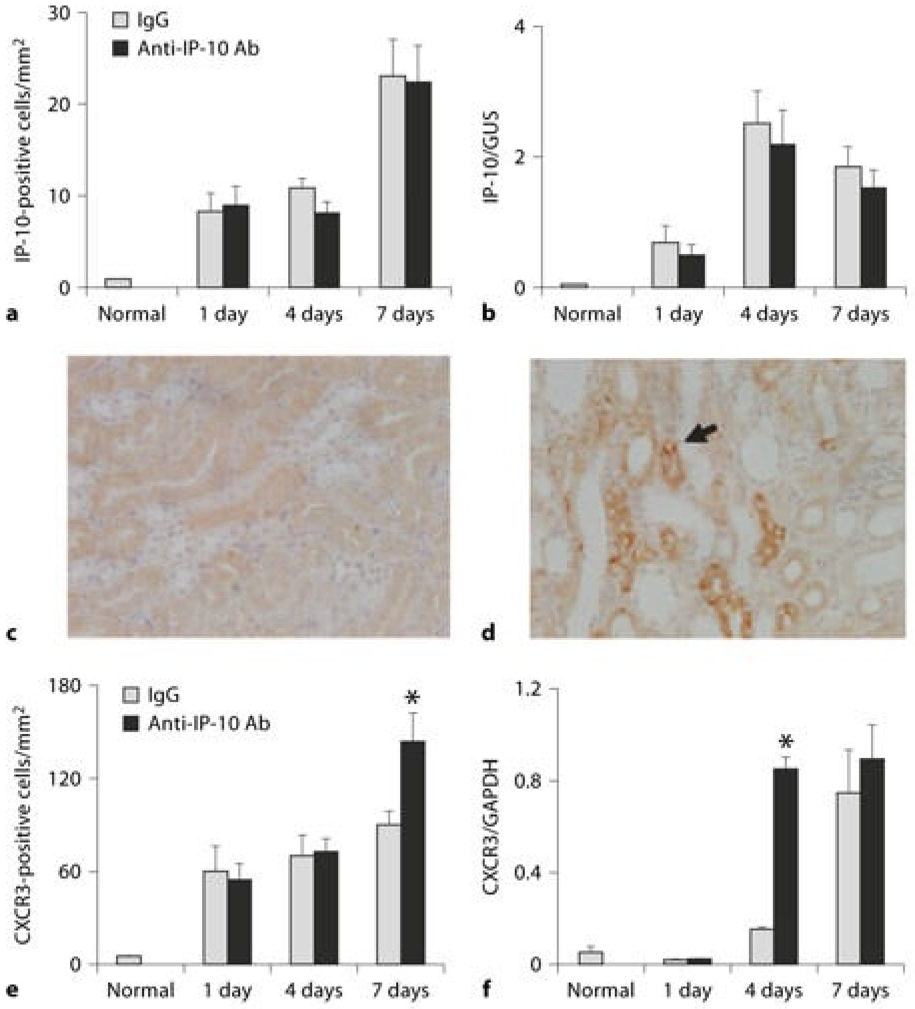

### CD183 (CXCR3)

**Abcam antibody (SP103) used in the current report:**

https://www.abcam.com/products/primary-antibodies/nestin-antibody-sp103-ab105389.html#lb

**Immunohistochemistry (Formalin/PFA-fixed paraffin-embedded sections) - Anti-Nestin antibody [SP103] (ab105389)**

Formalin-fixed, paraffin-embedded human kidney tissue stained for Nestin using ab105389 at 1/100 dilution in immunohistochemical analysis.

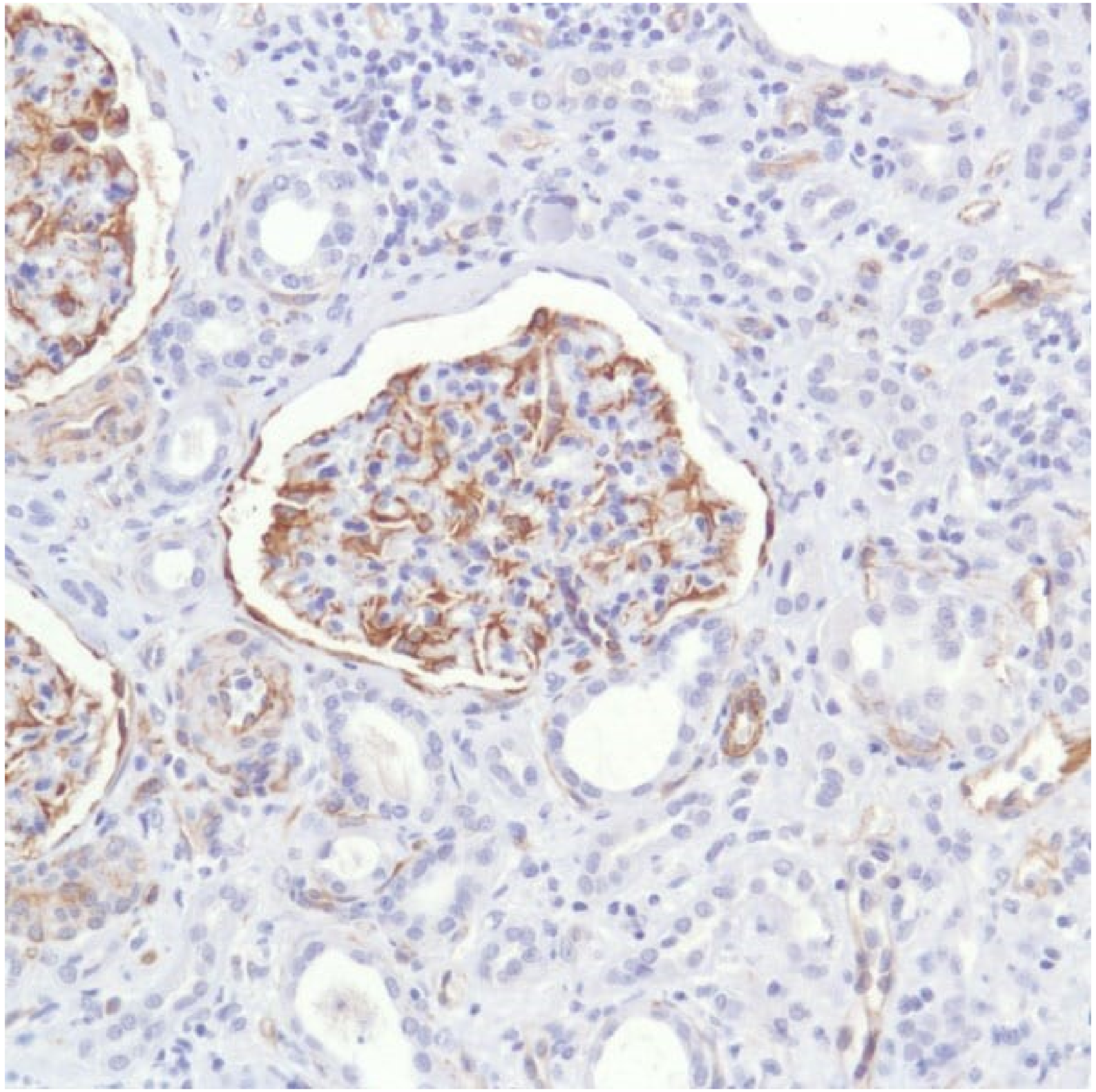

### CD183 (CXCR3)

**Abcam antibody (SP103) used in the current report:**

https://www.abcam.com/products/primary-antibodies/nestin-antibody-sp103-ab105389.html#lb

**Immunohistochemistry (Frozen sections) - Anti-Nestin antibody [SP103] (ab105389)**
IHC image of Nestin staining in a section of frozen normal human kidney* performed on a Leica BOND™ system using the standard protocol. The section was fixed in 10% paraformaldehyde (10 min) prior to staining. The section was incubated with ab105389, 5ugml, for 15 mins at room temperature and detected using an HRP conjugated compact polymer system. DAB was used as the chromogen. The section was then counterstained with haematoxylin and mounted with DPX. The inset secondary-only control image is taken from an identical assay without primary antibody.

For other IHC staining systems (automated and non-automated) customers should optimize variable parameters such as antigen retrieval conditions, primary antibody concentration and antibody incubation times.

*Tissue obtained from the Human Research Tissue Bank, supported by the NIHR Cambridge Biomedical Research Centre

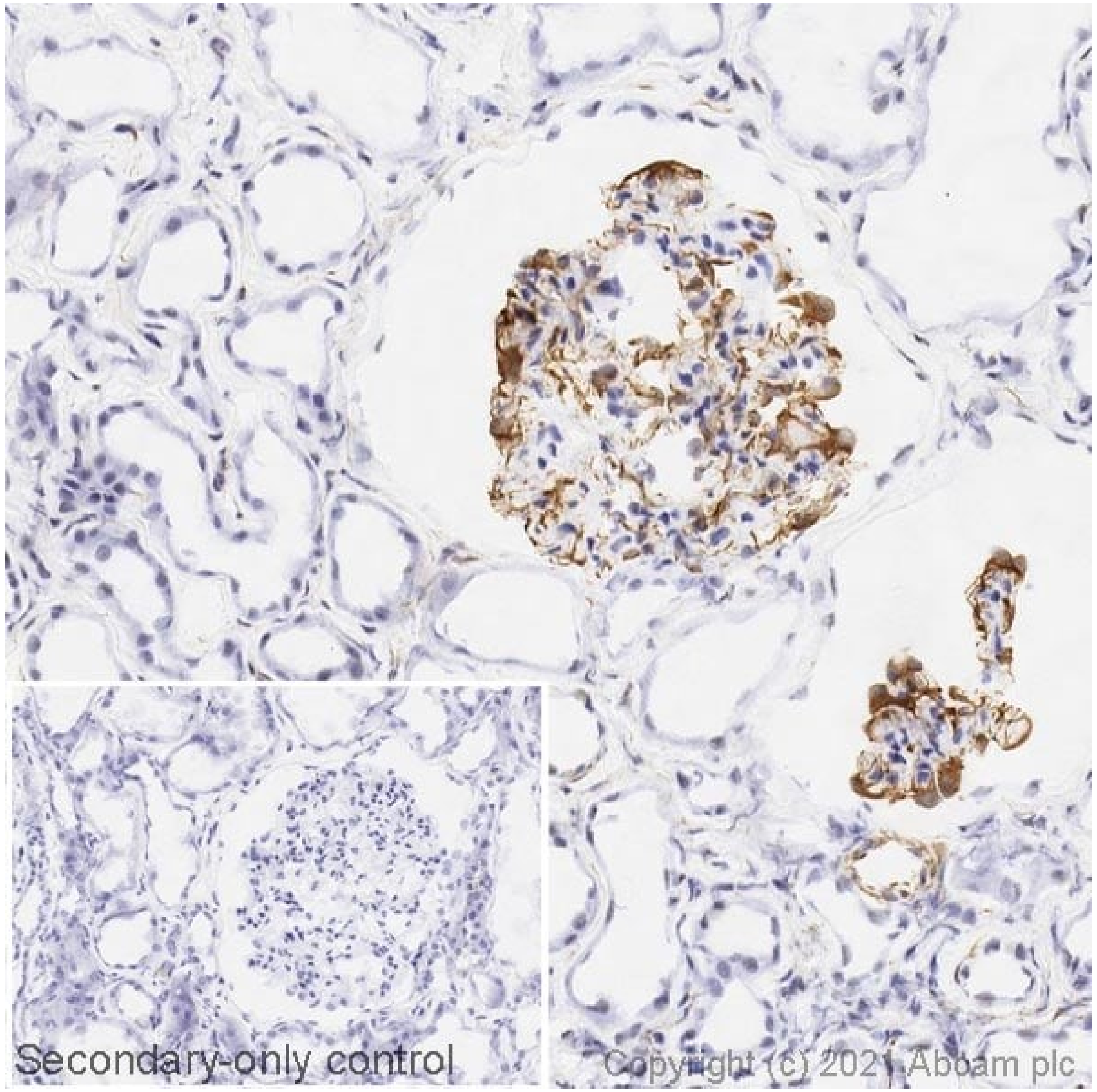

### CD183 (CXCR3)

https://www.thermofisher.com/antibody/product/CXCR3-Antibody-Polyclonal/PA5-23679

Immunohistochemistry analysis in formalin-fixed, paraffin-embedded human kidney tissue using a CXCR3 **polyclonal** antibody (Product # PA5-23679), followed by HRP-conjugated secondary antibody and DAB staining.

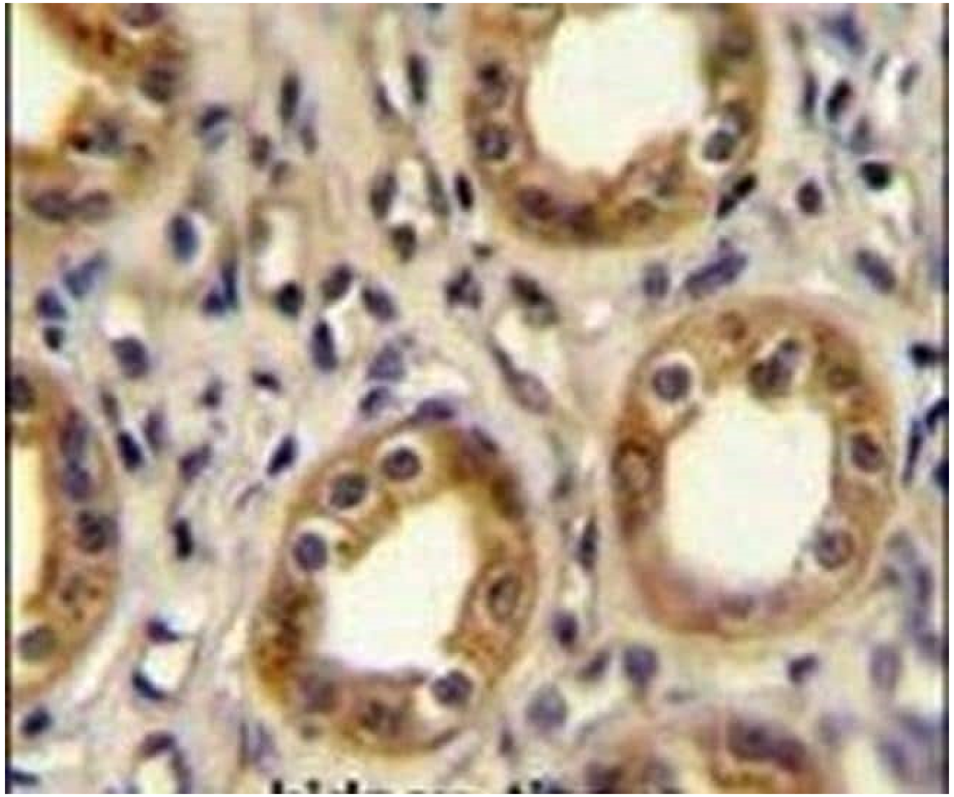

### CD183 (CXCR3)

https://www.thermofisher.com/antibody/primary/query/cxcr3/filter/application/Immunohistochemistry/species/Human/clonality/Polyclonal

CXCR3 Antibody (26756-1-AP) in IHC (P) Immunohistochemistry of paraffin-embedded human kidney tissue slide using 26756-1-AP (CXCR3 Antibody) at dilution of 1:200 (under 10x lens).

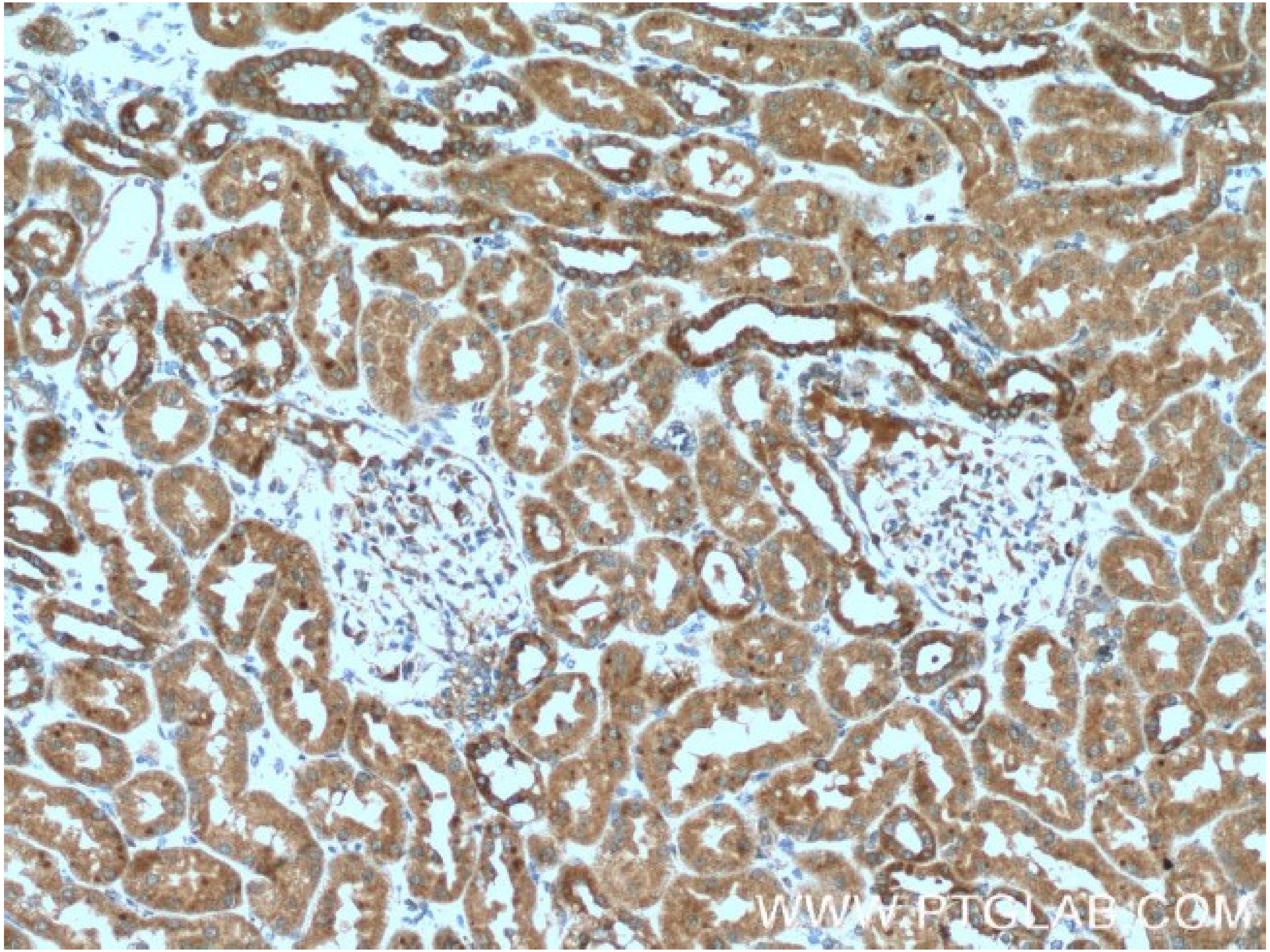

### EpCAM

Human Protein Atlas

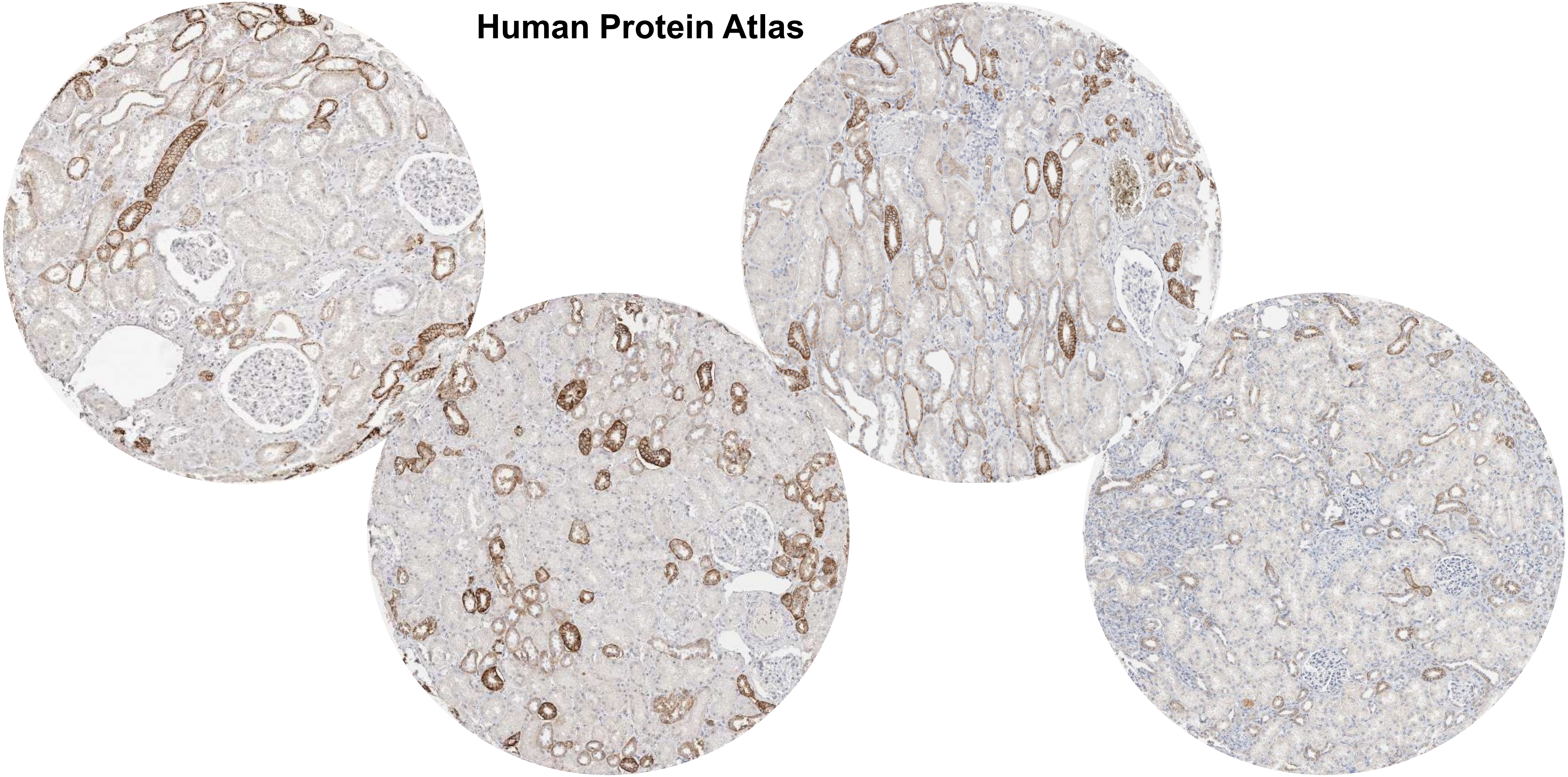

### EpCAM

**PMID 18025791**

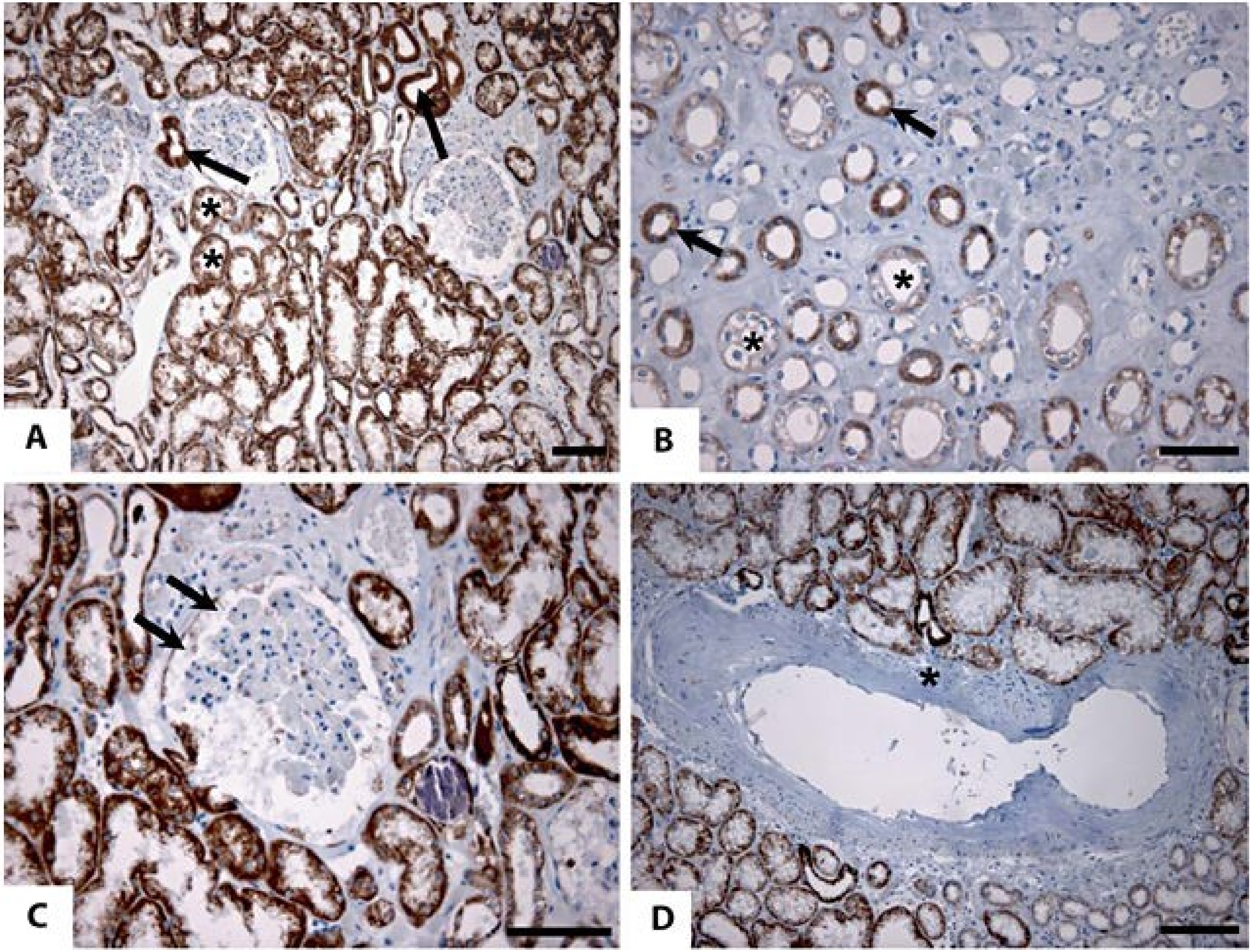


Expression pattern of hEpCAM in the adult human kidney. The **strongest staining for hEpCAM was observed in distal convoluted tubules** (**A**, examples indicated by arrows) **and medullary loops of Henle** (**B**, examples indicated by arrows). The weakest staining for hEpCAM was found in medullar proximal tubules (**A**, examples indicated by asterisks), collecting ducts (**B**, examples indicated by asterisks), and in some of the parietal epithelial cells in the capsule of Bowman (**C**, arrows). Nonepithelial structures, such as blood vessels (**D**, example indicated by asterisk), did not express hEpCAM. Original magnification: **A**, **D** ×100; **B**, **C** ×200. Scale bars: 40 μm

### EpCAM

**PMID 18025791**

**Expression of EpCAM homologues in human, hEpCAM Tg and wt murine adult kidneys**

| Kidney region | Human | hEpCAM Tg mouse | | wt mouse |
|---|---|---|---|---|
| | hEpCAM | hEpCAM | mEpCAM | mEpCAM |
| Capillary tufts | – | – | – | – |
| Mesangial cells | – | – | – | – |
| Bowman's space | – | – | – | – |
| Podocytes | – | – | – | – |
| Capsules of Bowman | + | + | + | + |
| Proximal tubules | | | | |
| Outer zone of cortex | – | – | – | – |
| Medulla, cortex between pyramids | + | + | + | + |
| Distal convoluted tubules | ++ | ++ | ++ | ++ |
| Loops of Henle | ++ | + | + | + |
| Collecting ducts | + | ++ | ++ | ++ |
| Interstitial space | – | – | – | – |

Staining intensity analysis: – = negative (no staining); + = weakly positive (low-level staining); ++ = strongly positive (highest-level staining).

### EpCAM

**PMID 24215118**

A) EpCAM expression in normal renal tissue: negativity in glomeruli and positivity related to the distal tubule system and collecting ducts (EpCAM, 10×). (B) ccRCC with negative EpCAM-immunoreactivity (IRS score 0; EpCAM, 10×). (C) Moderate EpCAM-expression (IRS score 6) in a ccRCC case (EpCAM, 10×). (D) Strong EpCAM expression (IRS score 12) in a pRCC case (EpCAM, 10×).

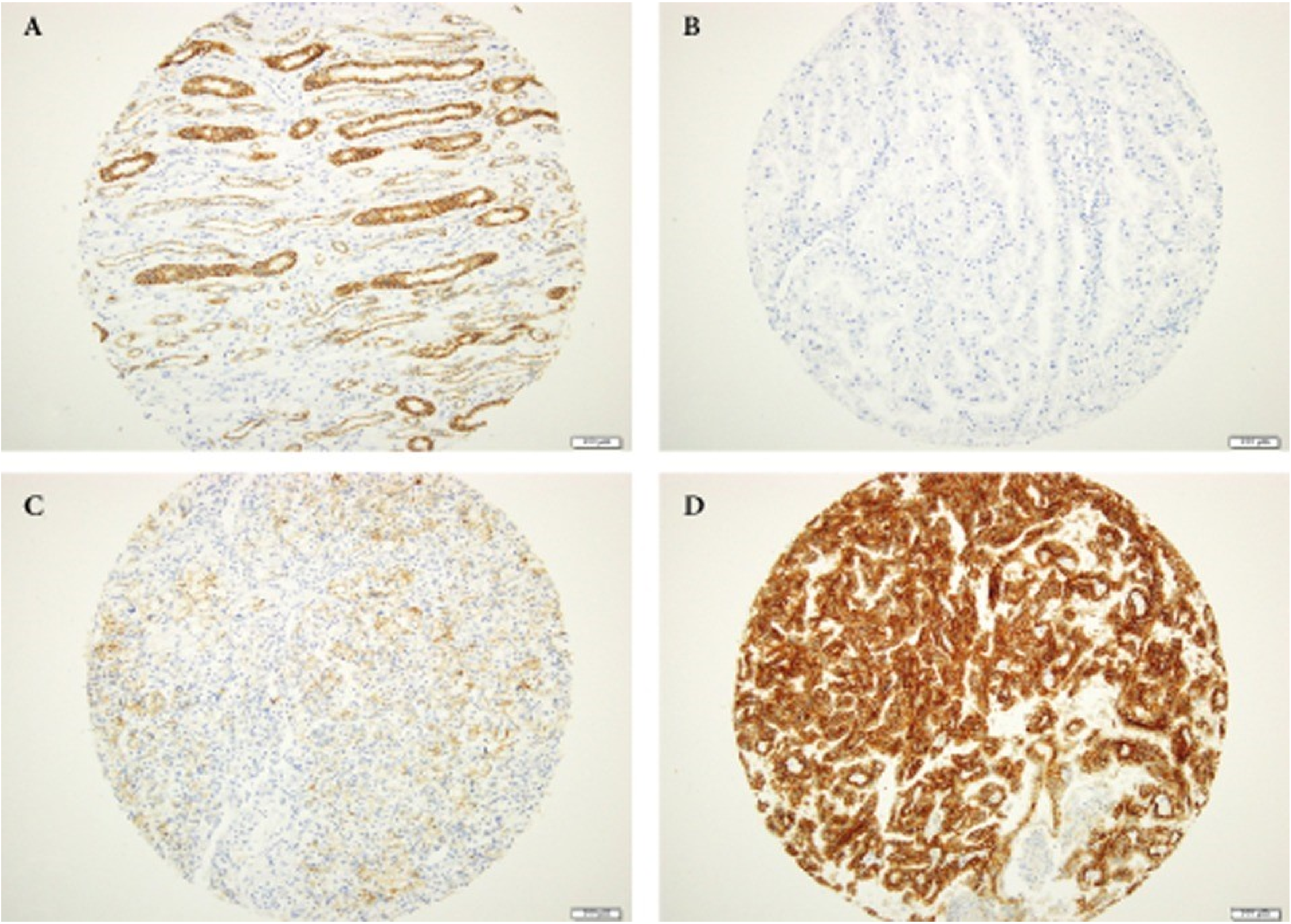

### CD107a (LAMP1)

**Human Protein Atlas**

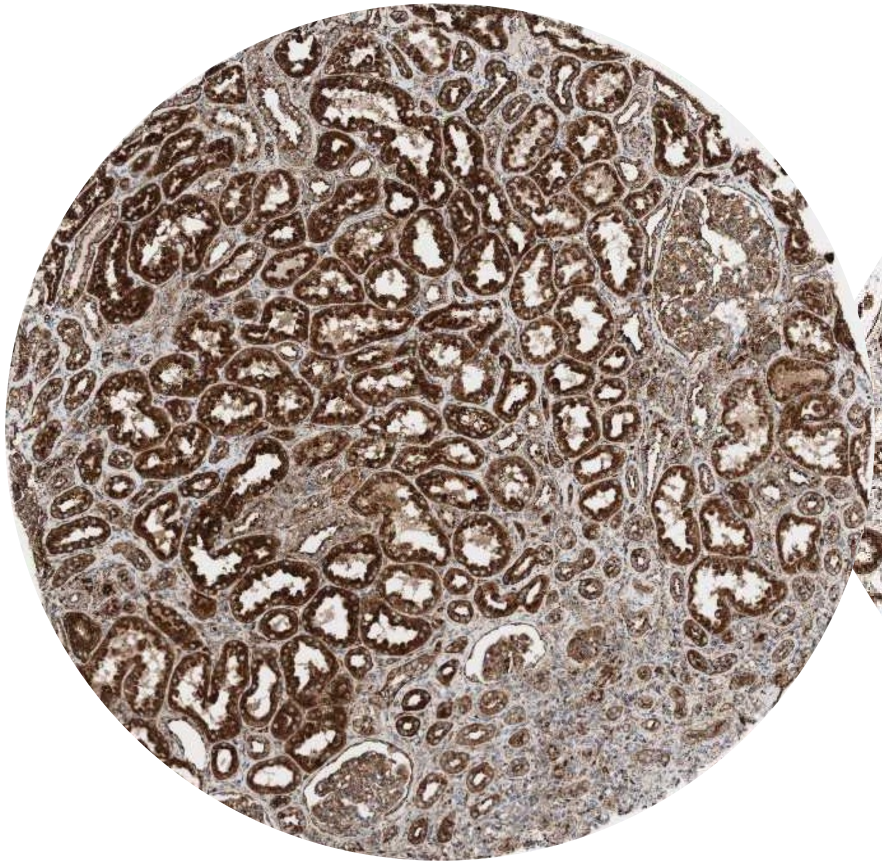

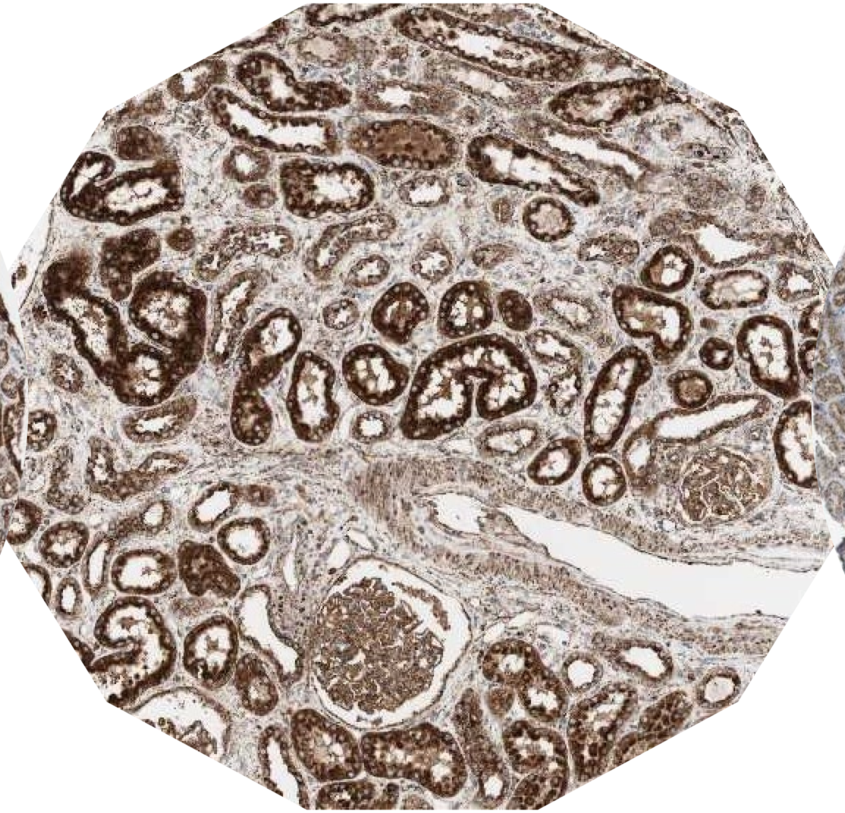

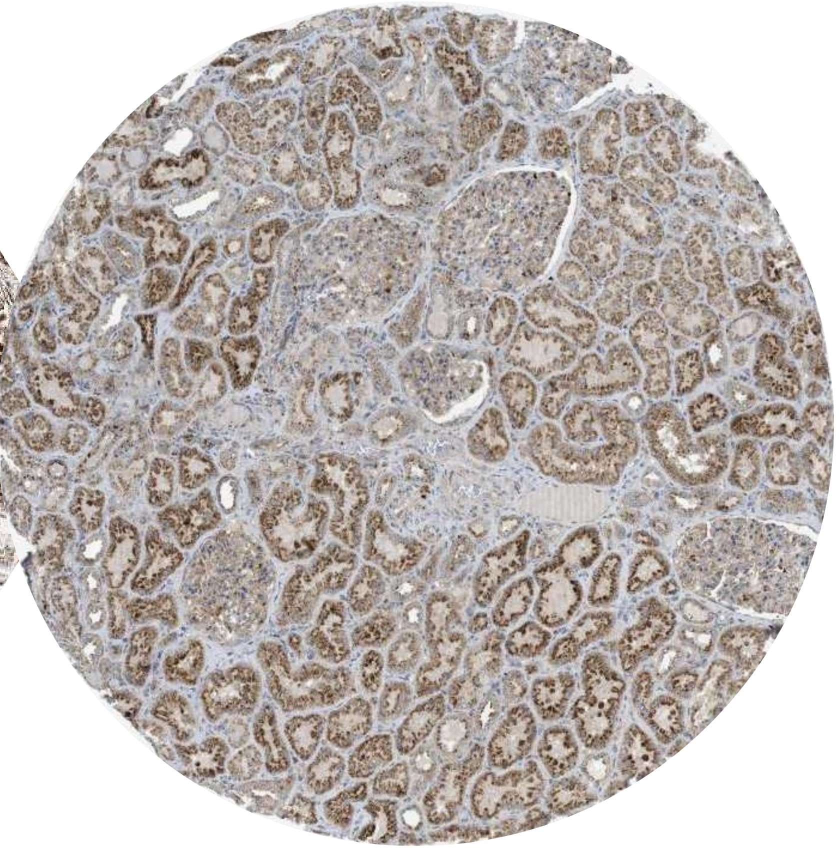

# CD107a

**PMID: 33441424**

**Fig. 7.** Phenotypic characterization of kidneys obtained from subjects with CKD associated genetic-risk variants (A) Representative images of LAMP1 staining in human kidney tissues of subjects with AA, AG and GG alleles, respectively. The A allele is the risk allele. White squares indicate images shown at higher magnification. Scale bar, 10 μm. (B) Gene expression analysis of microdissected kidney tubule samples from risk allele vs. reference allele. Pathway enrichment analysis (DAVID) of top differentially expressed genes. The bar plot shows the significance (log10(p) and log10(odds ratio)) of the enrichment of specific pathways. (C) Relative mRNA expression of C1QL1 in reference and risk genotype kidney tubule samples.

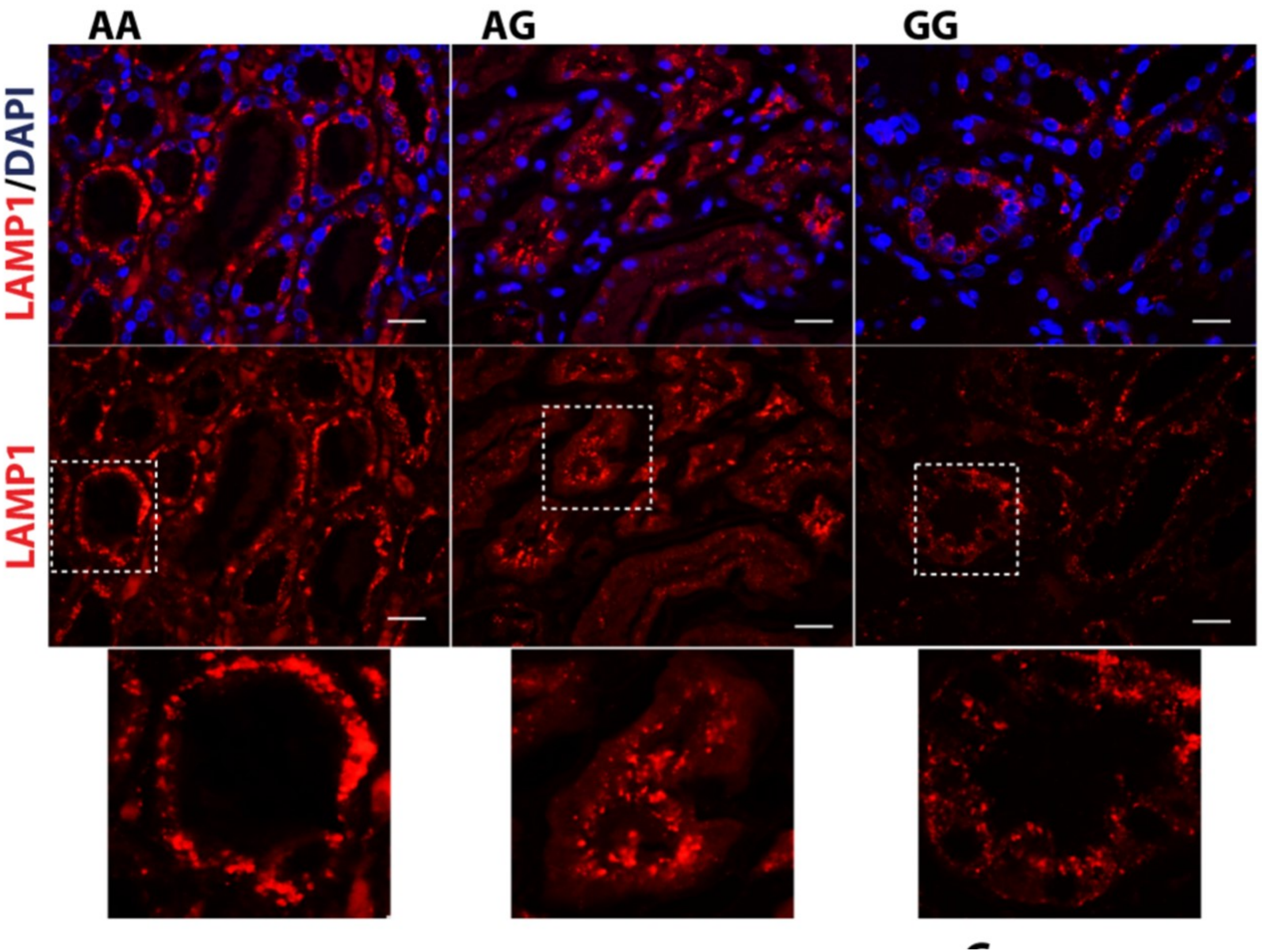


- FFPE human kidney tissue
- Anti-LAMP1 Ab: Abcam rabbit polyclonal ab24170

### CD107a (LAMP1)

https://www.abcam.com/lamp1-antibody-lysosome-marker-ab24170.html

IHC image of LAMP1 staining in a section of formalin-fixed paraffin-embedded normal human kidney* performed on a Leica BOND$^{TM}$ system using the standard protocol B. The section was pre-treated using heat mediated antigen retrieval with sodium citrate buffer (pH6, epitope retrieval solution 1) for 20mins. The section was then incubated with ab24170, 1ug/ml, for 15 mins at room temperature and detected using an HRP conjugated compact polymer system. DAB was used as the chromogen. The section was then counterstained with haematoxylin and mounted with DPX. The inset secondary-only control image is taken from an identical assay without primary antibody.

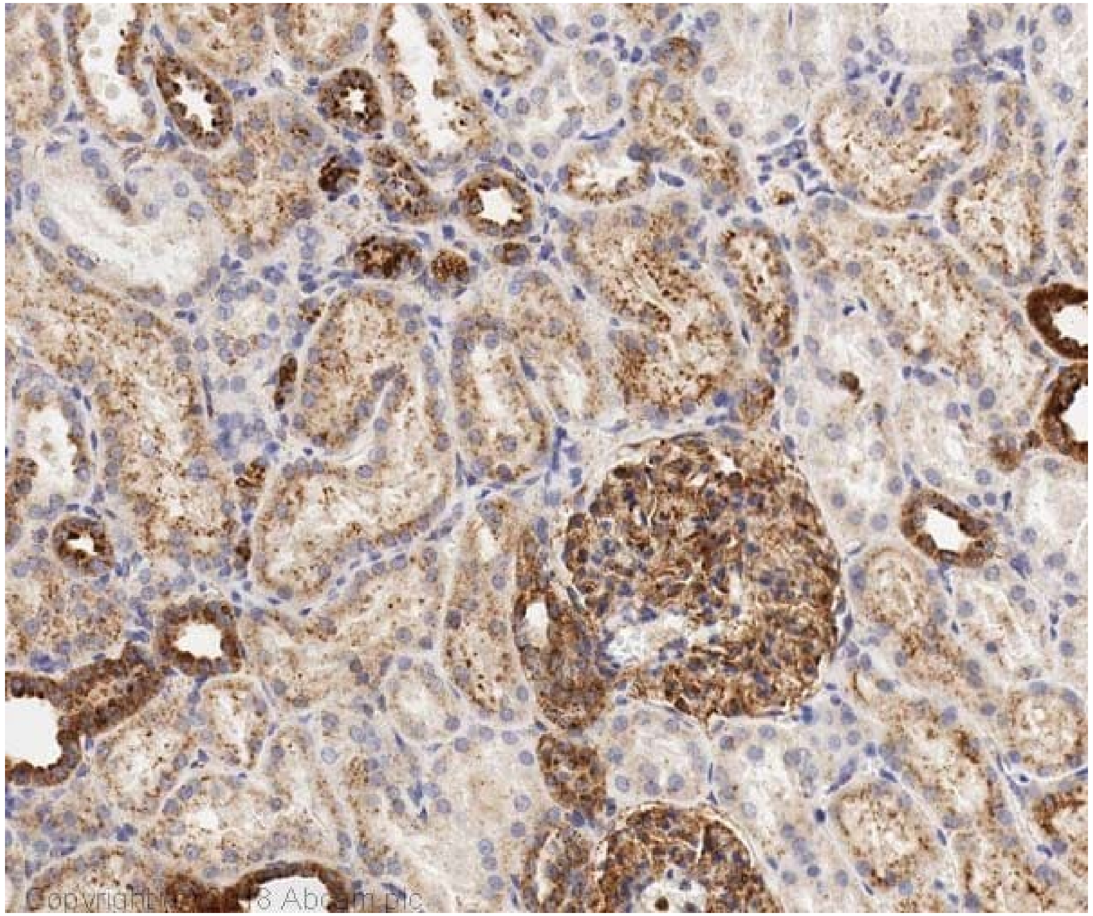


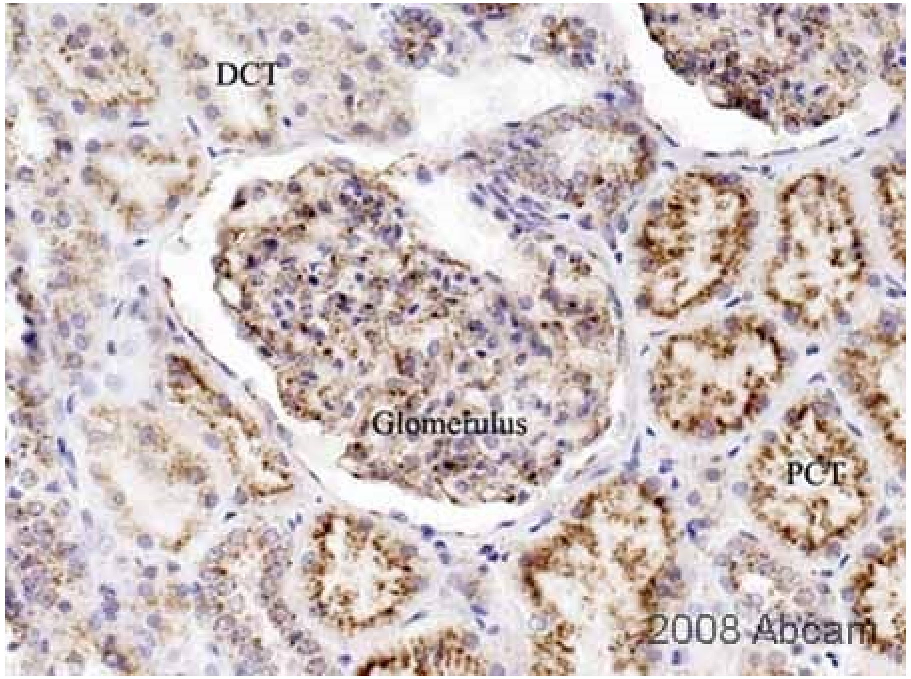

### CD107a (LAMP1)

https://www.novusbio.com/products/lamp-1-cd107a-antibody-5e7_nbp2-52721

Immunohistochemistry-Paraffin: LAMP-1/CD107a Antibody (5E7) [NBP2-52721] - Analysis of FFPE human kidney tissue section using LAMP-1 antibody (clone 5E7) at 1:100 dilution. The antibody generated a specific cytoplasmic staining (with dotted appearance) in all the ductal and tubular epithelial cells.

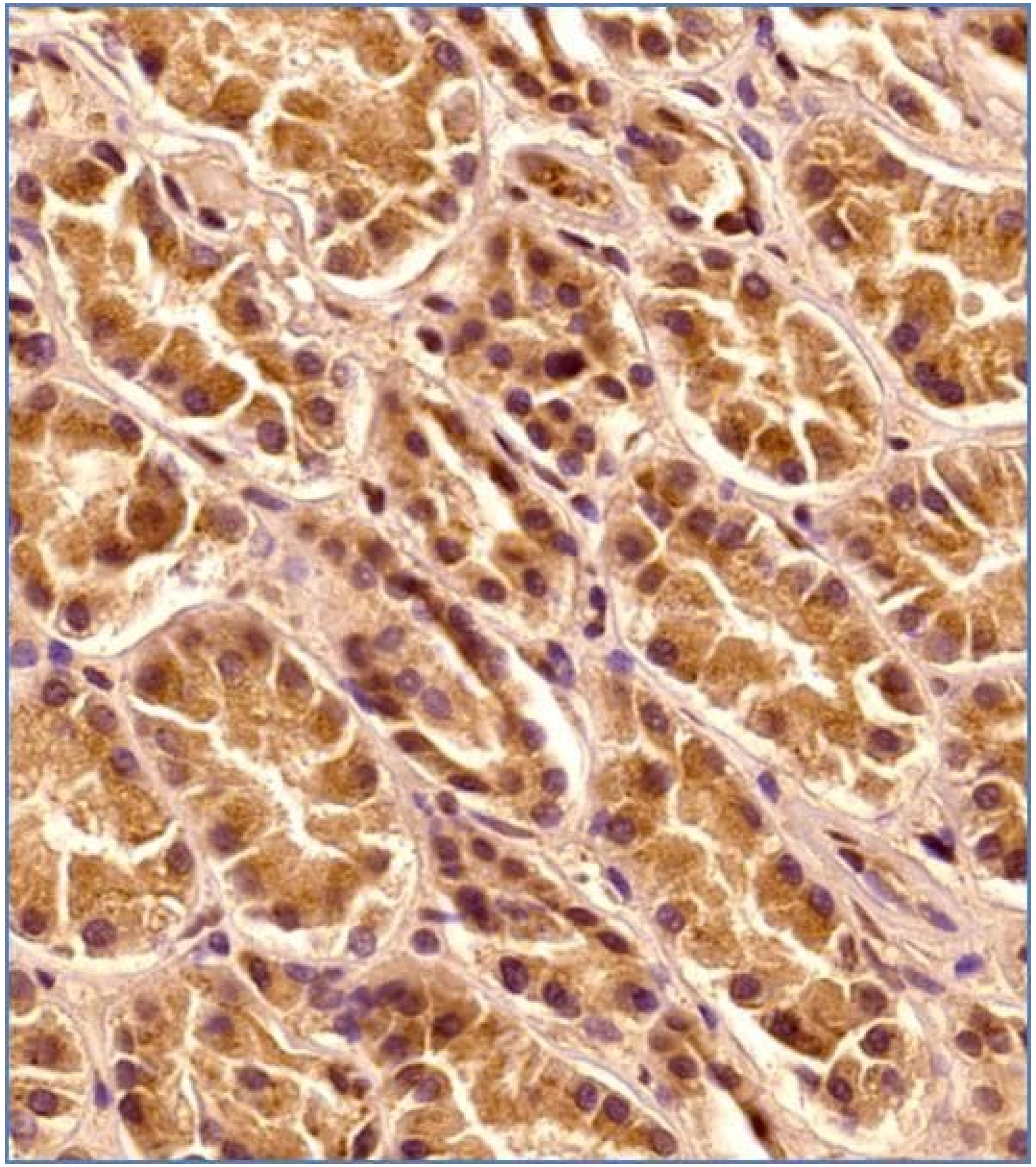

### CD107a (LAMP1)

https://www.novusbio.com/products/lamp-1-cd107a-antibody-5e7_nbp2-52721

Immunohistochemistry-Paraffin: LAMP-1/CD107a Antibody (5E7) [NBP2-52721] - Analysis of FFPE human kidney tissue section using LAMP-1 antibody (clone 5E7) at 1:100 dilution. The antibody generated a specific cytoplasmic staining (with dotted appearance) in all cells, especially the ductal and tubular epithelial cells. Few cells from the glomeruli showed nuclear signal also.

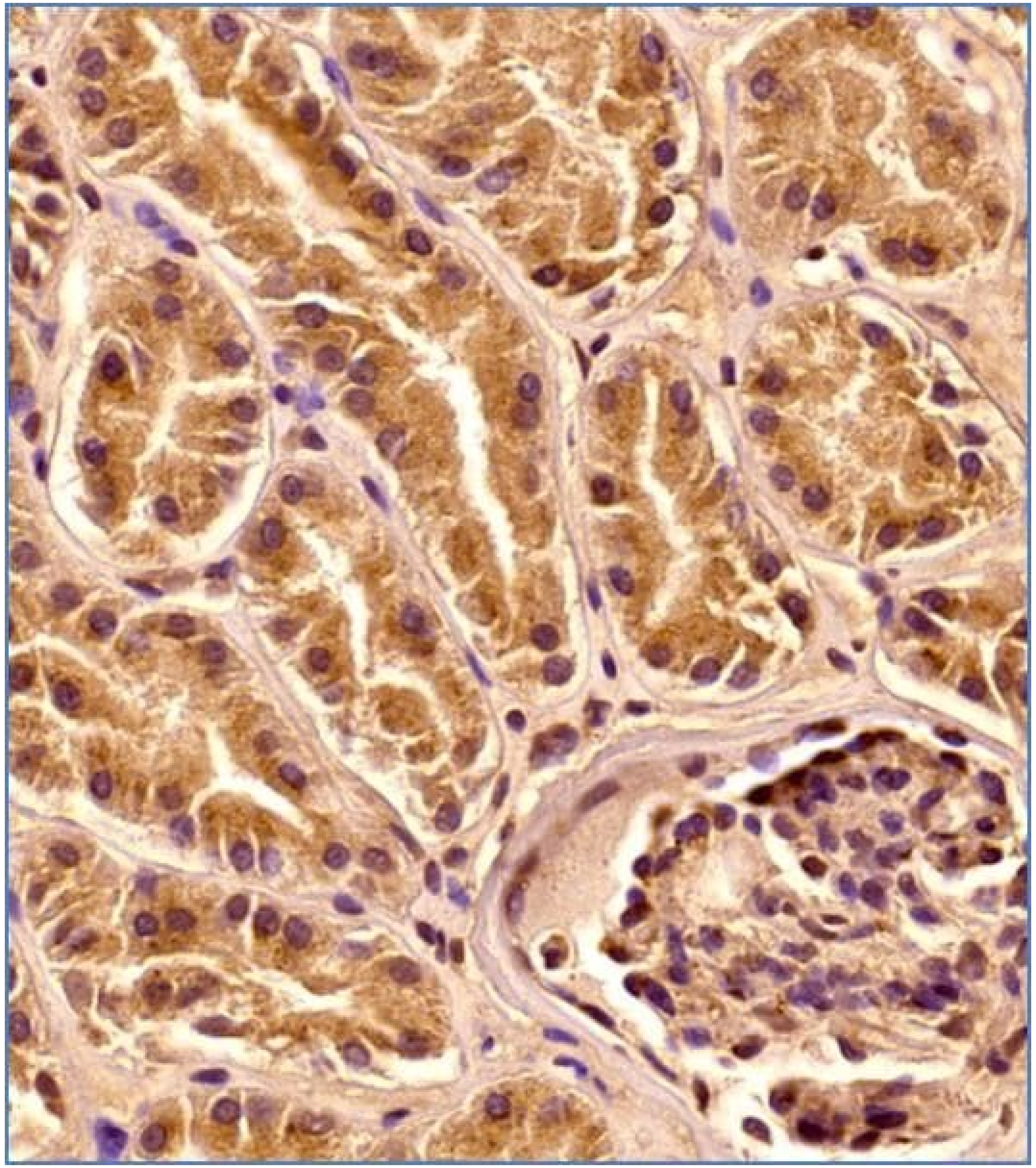

### CD107a (LAMP1)

https://www.novusbio.com/products/lamp-1-cd107a-antibody-cl4489_nbp2-59053

Immunohistochemistry-Paraffin: LAMP-1/CD107a Antibody (CL4489) [NBP2-59053] - Staining of human kidney shows strong granular cytoplasmic positivity in renal tubules and glomeruli.

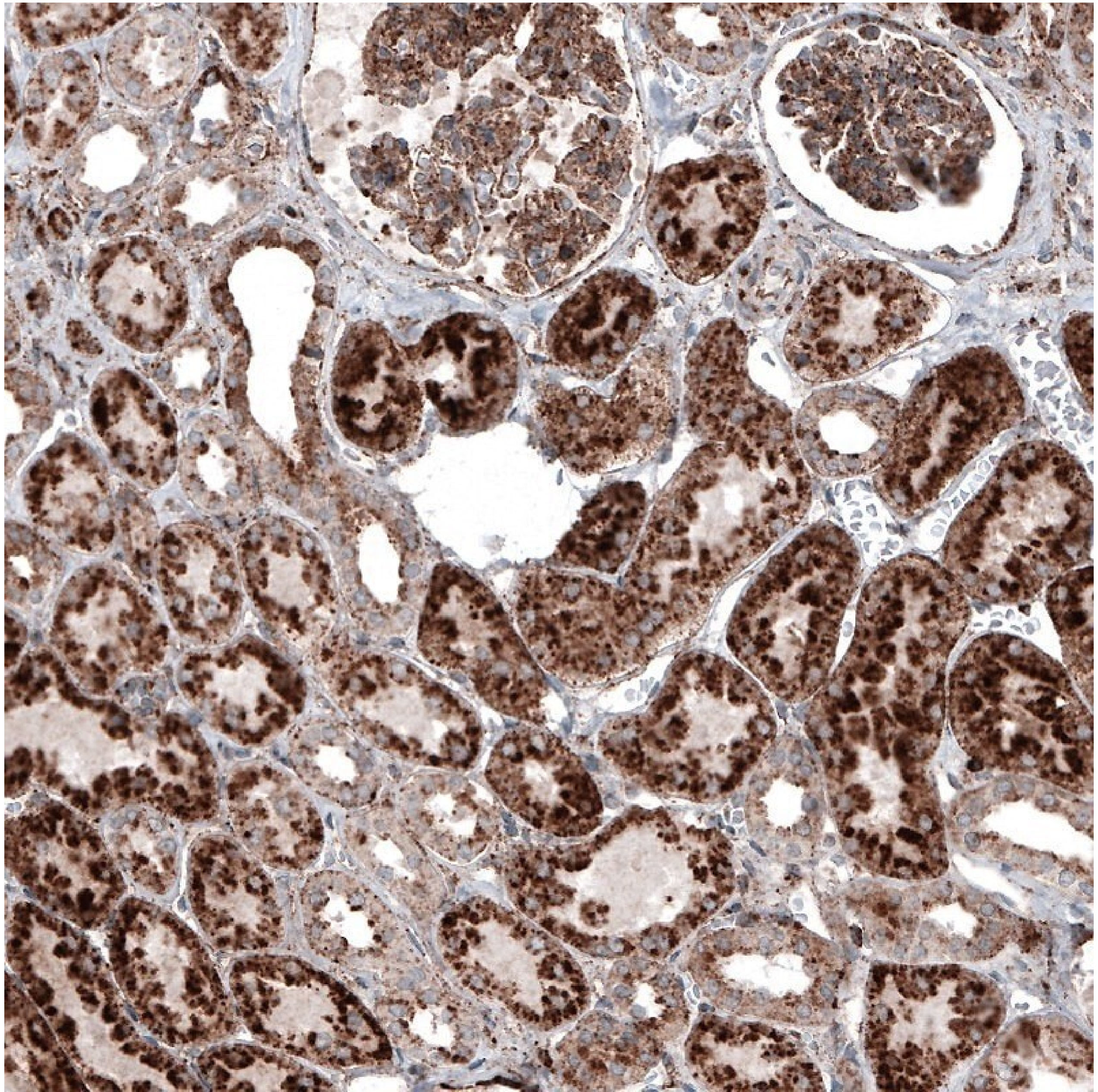

### CD107a (LAMP1)

https://www.thermofisher.com/antibody/product/CD107a-LAMP-1-Antibody-clone-JJ0940-Recombinant-Monoclonal/MA5-32491

Immunohistochemical analysis of CD107a (LAMP-1) of paraffin-embedded Human kidney tissue using a CD107a-LAMP-1 Monoclonal antibody (Product #MA5-32491). Counter stained with hematoxylin.

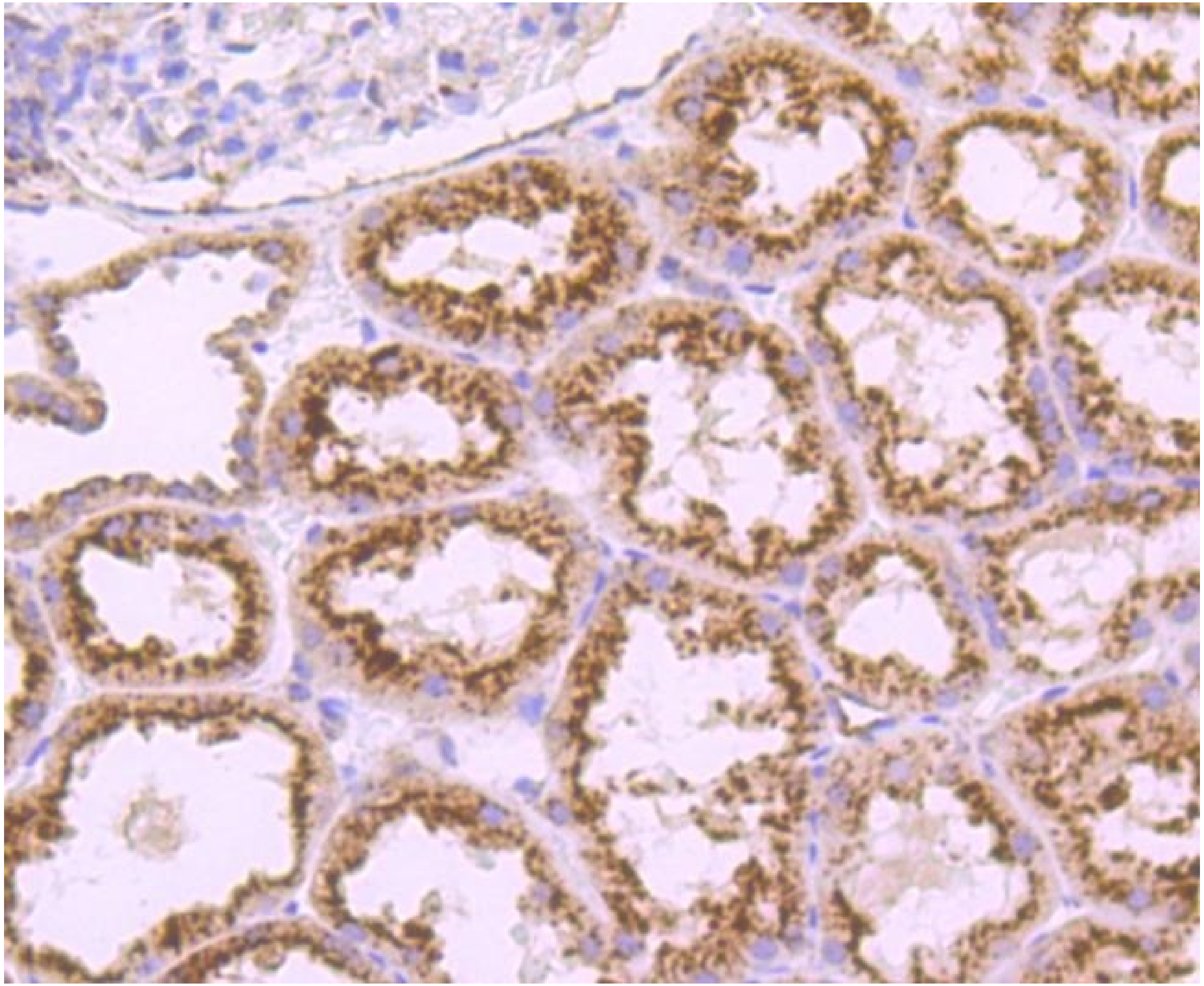

### CD107a (LAMP1)

https://www.lsbio.com/antibodies/ihc-plus-epcam-antibody-clone-vu-1d9-flow-icc-ihc-ip-wb-western-ls-b4511/120351

Anti-EPCAM antibody IHC of human kidney. Immunohistochemistry of formalin-fixed, paraffin-embedded tissue after heat-induced antigen retrieval. Antibody concentration 10 ug/ml.

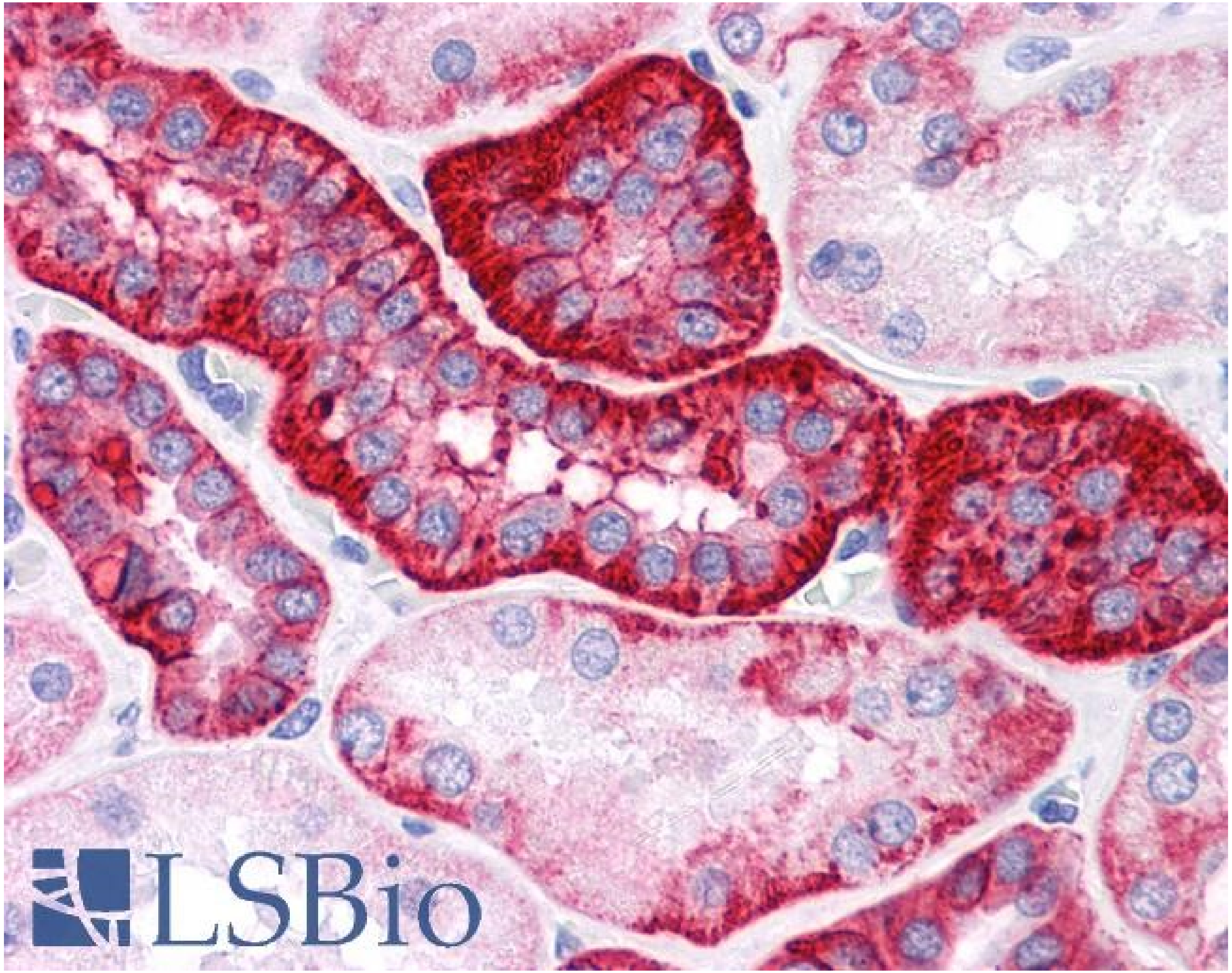

### mtTFA

**Human Protein Atlas**

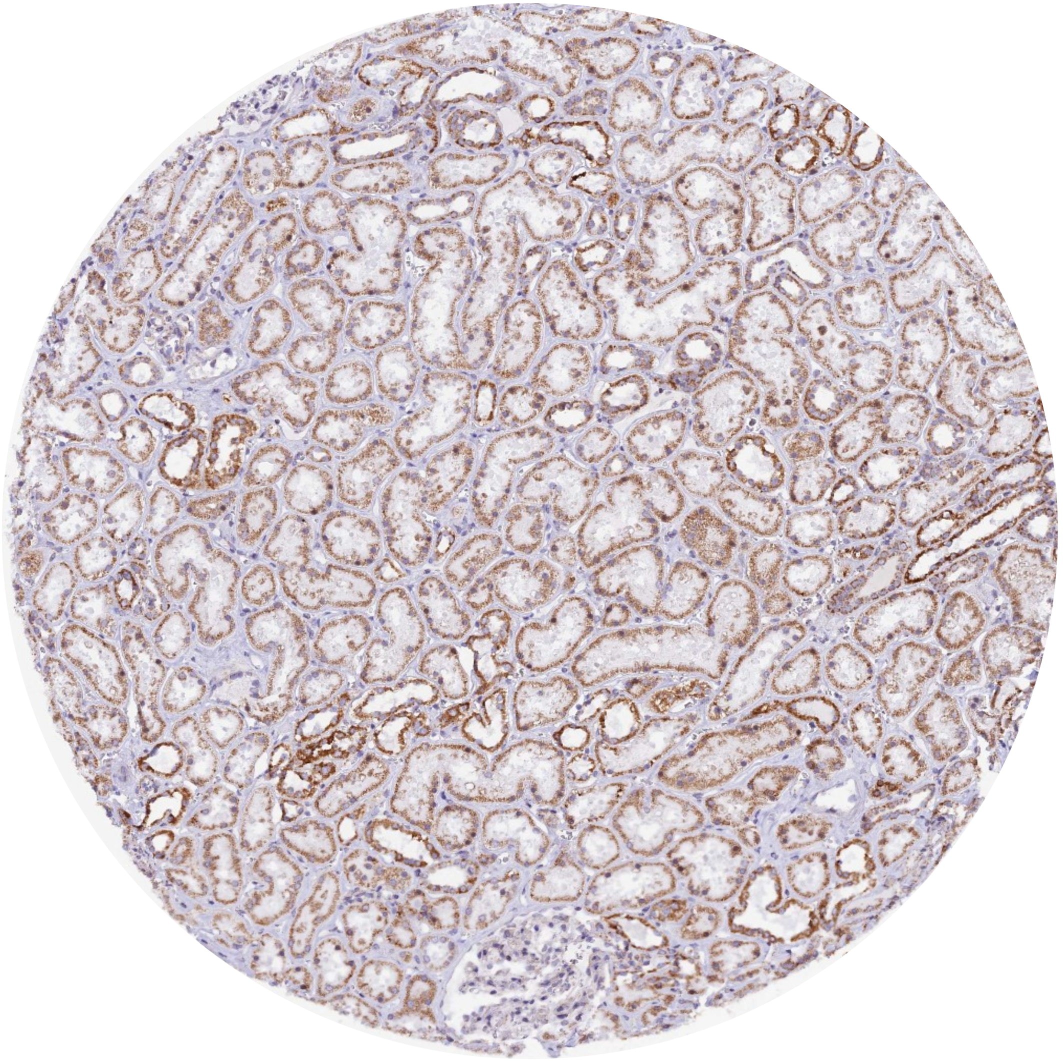

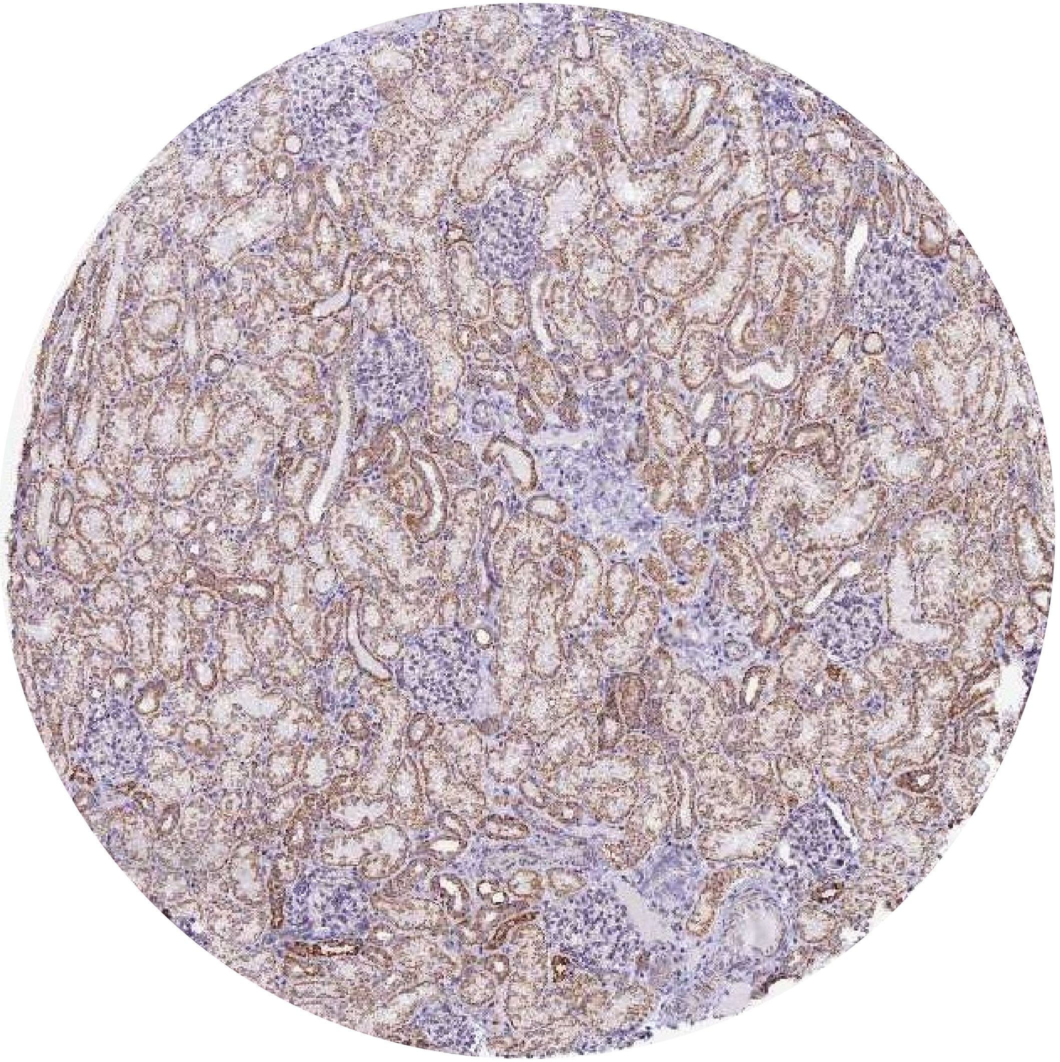

### mtTFA

**PMID: 33159962**

**Fig 7.** Mitochondrial transcription factor A (TFAM) expression in renal cysts from patients with polycystic kidney disease is reduced. (a) Representative images of formalin-fixed paraffin-embedded sections from normal human kidneys and kidneys from polycystic kidney disease (PKD) patients analyzed by immunohistochemistry for TFAM expression, by immunofluorescence (IF) for voltage-dependent anion-selective channel 1 (VDAC) expression, and by RNA fluorescent in situ hybridization for mitochondrially encoded cytochrome c oxidase 1 (MT-CO1) and mitochondrially encoded ATP synthase membrane subunit 6 (MT-ATP6) mRNA expression. Arrows identify cyst lining epithelial cells, number signs depict cyst lumina, and asterisks depict glomeruli. Bar = 100 μm for low-magnification images and 10 μm for high-magnification images.

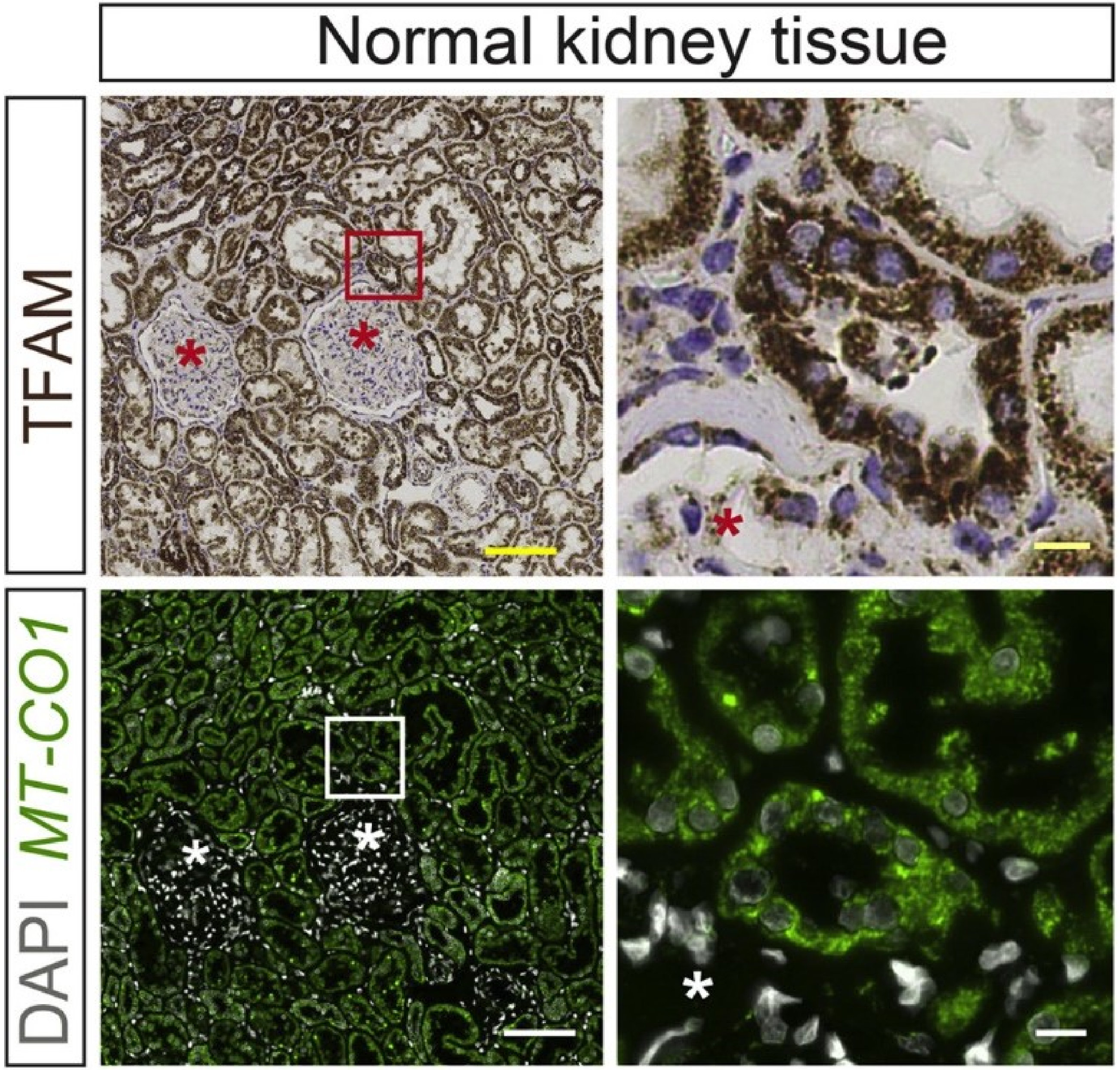

### mtTFA

https://www.sinobiological.com/antibodies/tfam-201038-t46#pid=1

Immunochemical staining of human TFAM in human kidney with rabbit polyclonal antibody at 1:100 dilution, formalin-fixed paraffin embedded sections.

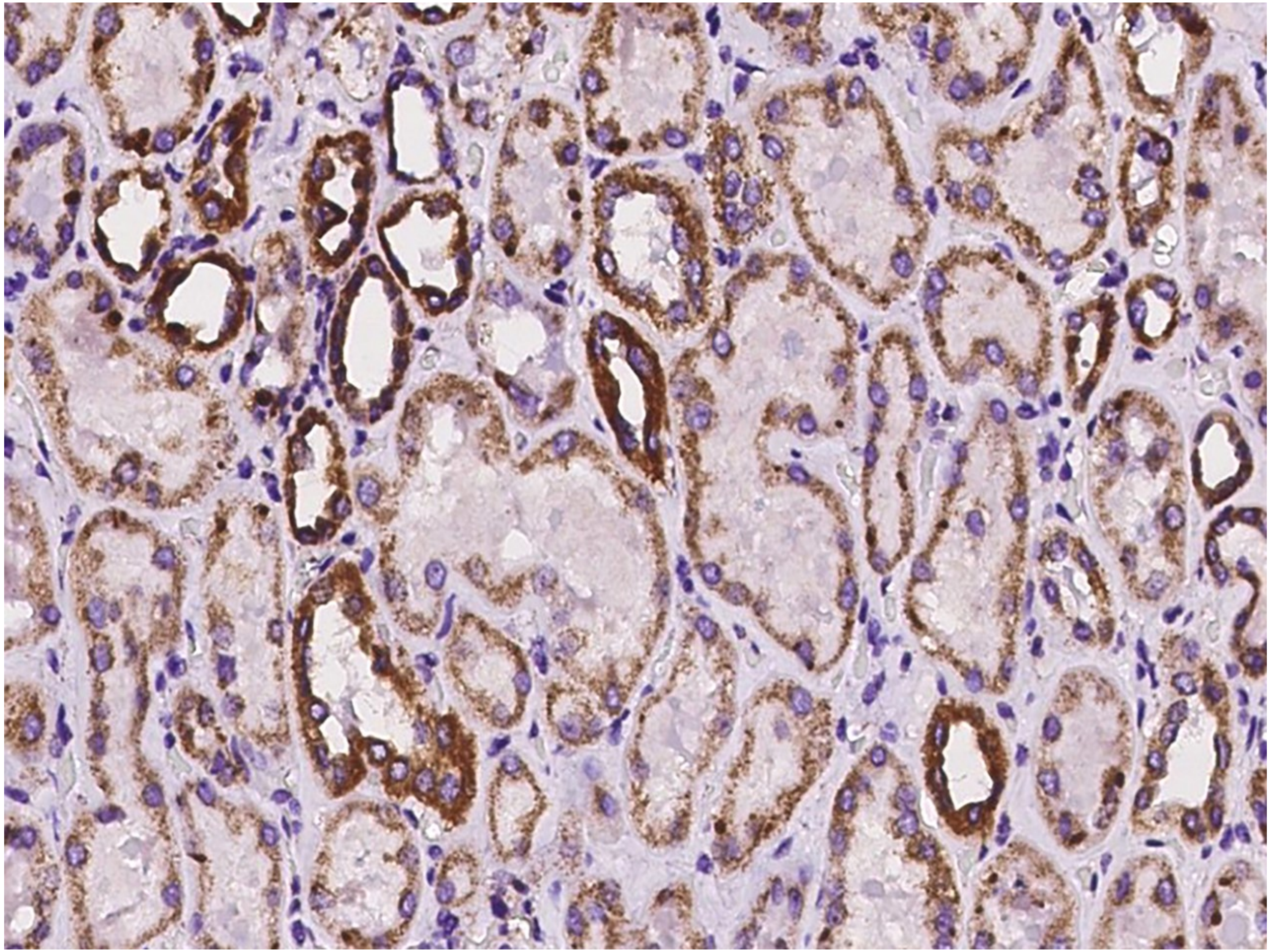

### mtTFA

https://www.biossusa.com/products/bsm-54118r

Paraformaldehyde-fixed and paraffin-embedded Human Kidney tissue incubated TFAM (16A1) Monoclonal Antibody (bsm-54118R) at 1:100, overnight at 4°C, followed by a conjugated secondary antibody and DAB staining. Counterstained with hematoxylin

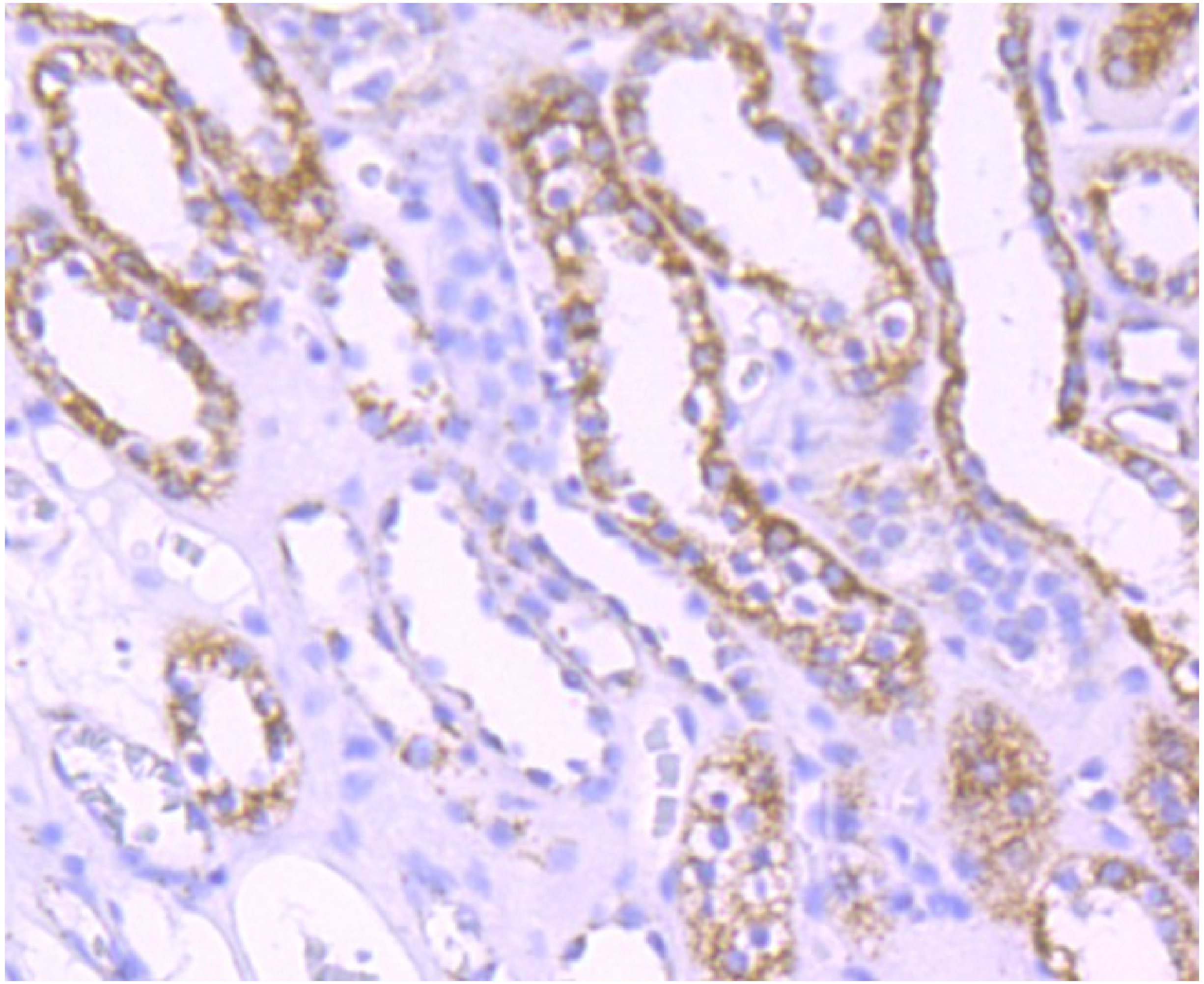

### CD227 (MUC1)

**Human Protein Atlas**

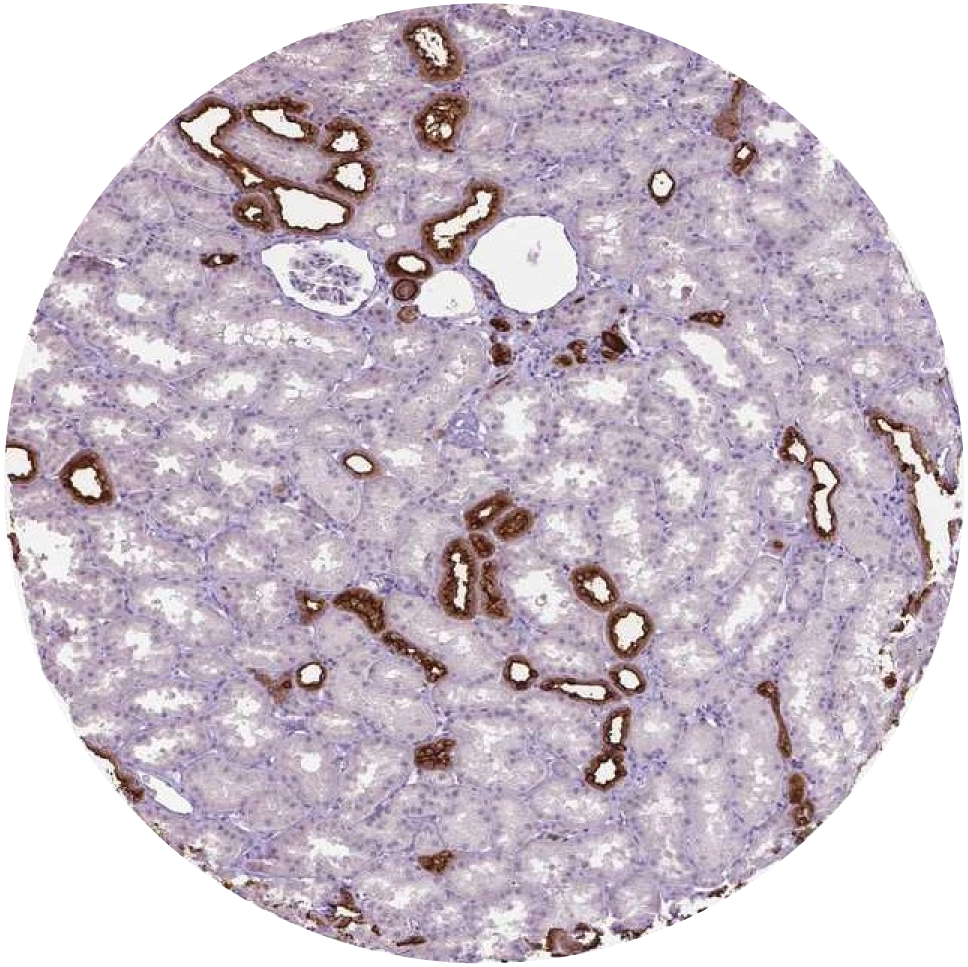

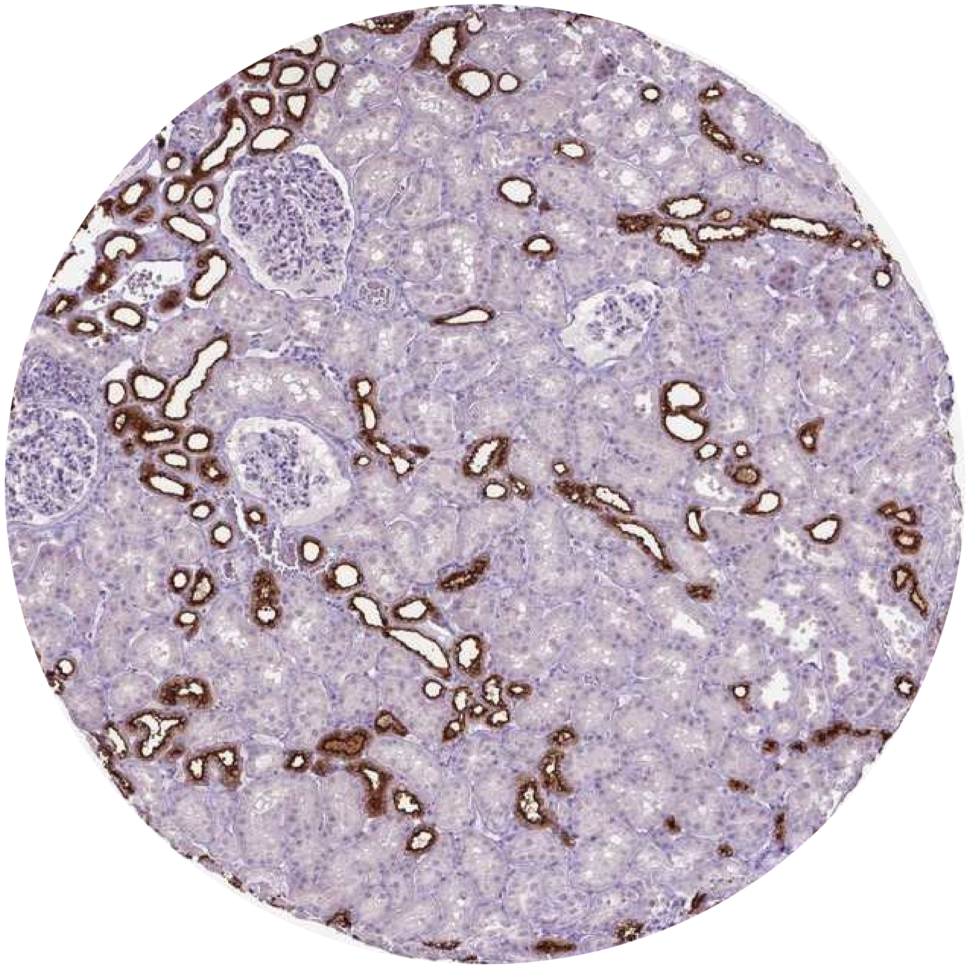

### CD227 (MUC1)

**PMID: 10755601**

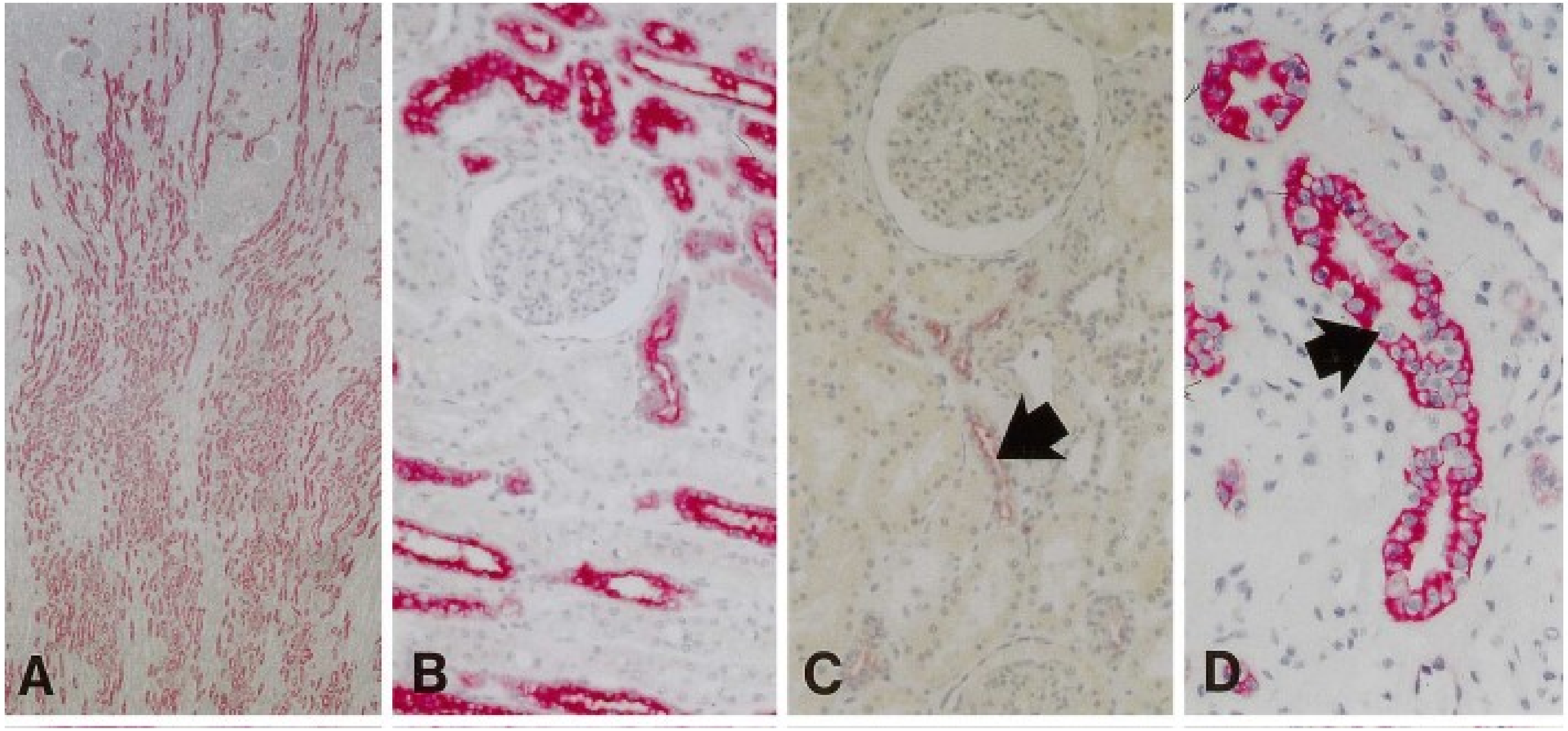


**Fig 2 A, B.** Normal kidney, distal tubules and collecting ducts are stained with A76-A/C7 (anti-MUC1), whereas renal glomeruli and proximal tubules are negative. **C.** Normal kidney, A78-G/A7 (anti-TF) stains the luminal surface of the distal tubules and collecting ducts, but not the renal glomeruli and proximal tubules**. D.** Normal kidney: cytokeratin 19 is detected with A53-B/A2 in distal tubules and collecting ducts. Note that the intercalated cells (arrow) are negative for A53-B/A2. E. Oncocytoma: A76-A/C7 stains the cytoplasm of tumour cells strongly. F. Clear-cell RCC: most tumour cells show staining of entire membrane for A76-A/C7, but some also have a cytoplasmic reaction. G. Chromophobic cell RCCs positive for A76-A/C7. H Clear/granular RCC: note A53-B/A2 staining in the cytoplasm. I, J. Serial sections of a clear-cell RCC, I weakly stained with A78-G/A7 (antiTF; arrows) and J more strongly stained after neuraminidase treatment (arrows). K. Clear-cell RCC: HBTn (anti-Tn) reacts with some tumour cells. L Clear-cell RCC: B72.3 (anti-s-Tn) reacts strongly.

### CD227 (MUC1)

**PMID: 27768738**

**Fig 7.** MUC1-TM and MUC1-ARF expression in normal human kidney and pancreas. Serial sections of paraffin-embedded human pancreatic and renal tissues were immunohistochemically stained with anti-MUC1-TM SEA module antibodies (anti-MUC1-TM [DMB5F3]) and anti-MUC1-ARF antibodies (anti-MUC1-ARF [MPR2G10]) as indicated. Normal kidney is shown in Panels A-i (MUC1-TM) and A-ii (MUC1-ARF); larger fields and higher magnifications are shown in Panels C-i, C-i' (MUC1-TM), and D-i, D-i' (MUC1-ARF)]. Glomerulus, proximal tubule, and distal tubule are designated by G, PT and DT respectively. Filled green arrows designate sites of MUC1-TM-SEA protein at the cell surface whereas filled red arrows designate MUC1-ARF protein; absence of anti-MUC1-ARF immunoreactivity is shown by filled red and white arrows.

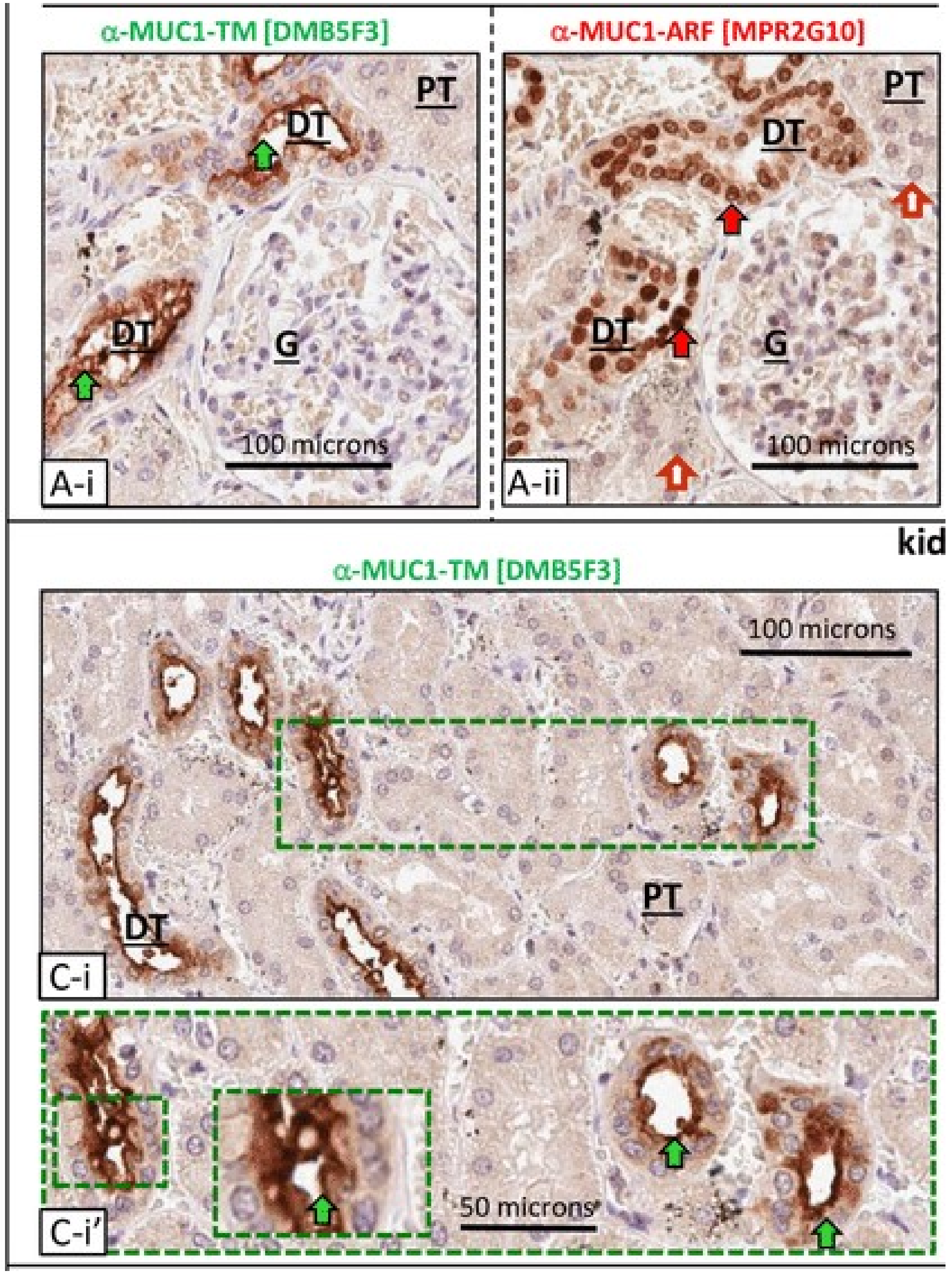

### HSPG2 (Perlecan)

Human Protein Atlas

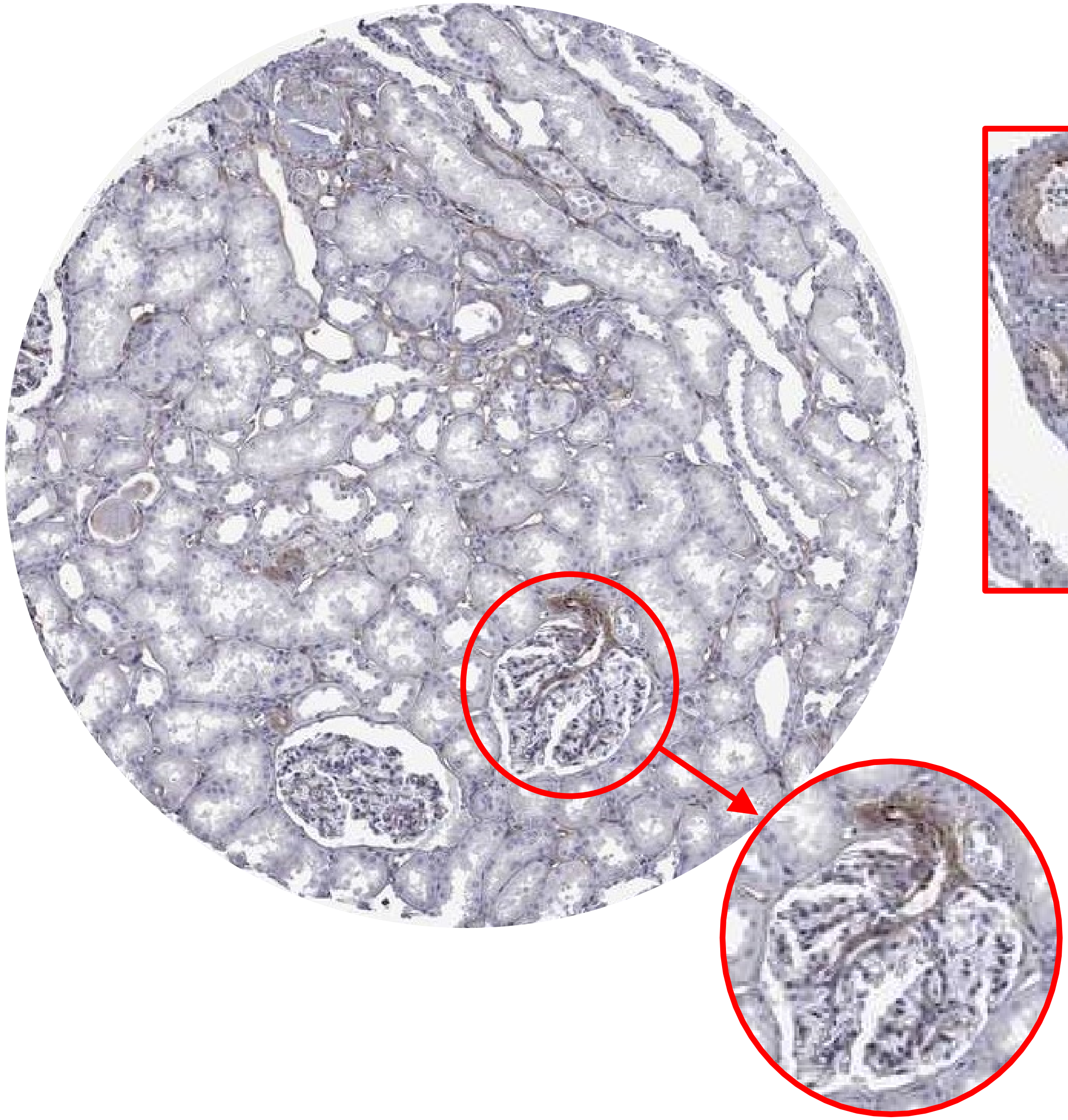

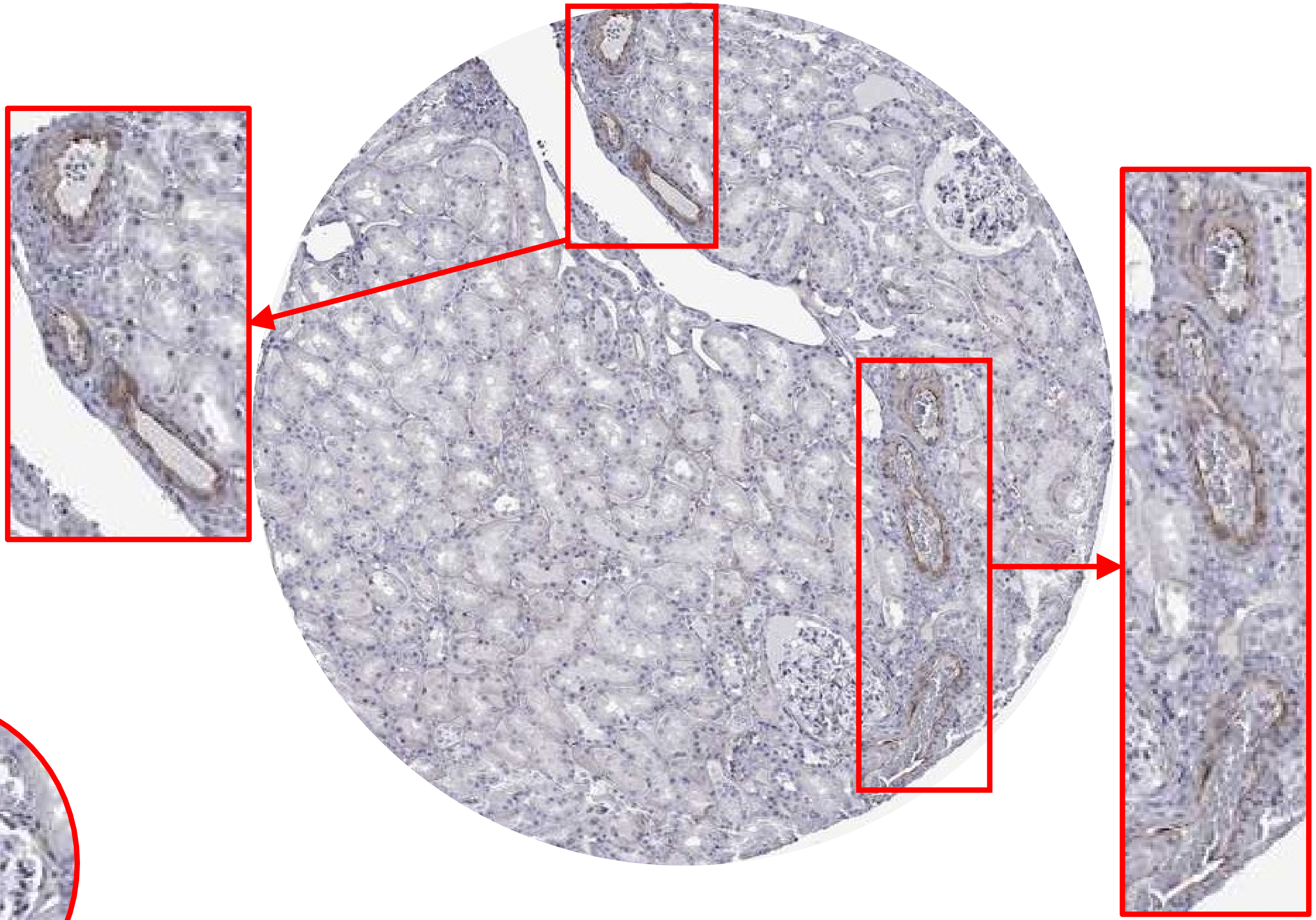

### HSPG2 (Perlecan)

**PMID 18300285**

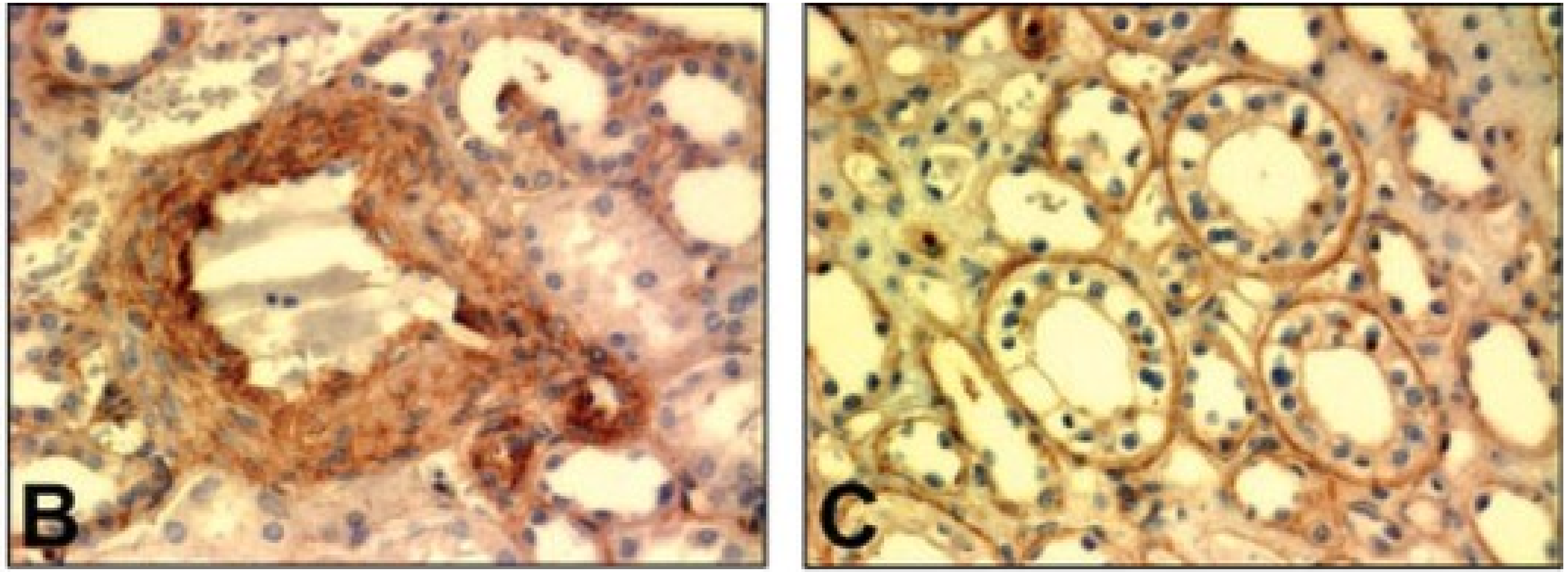


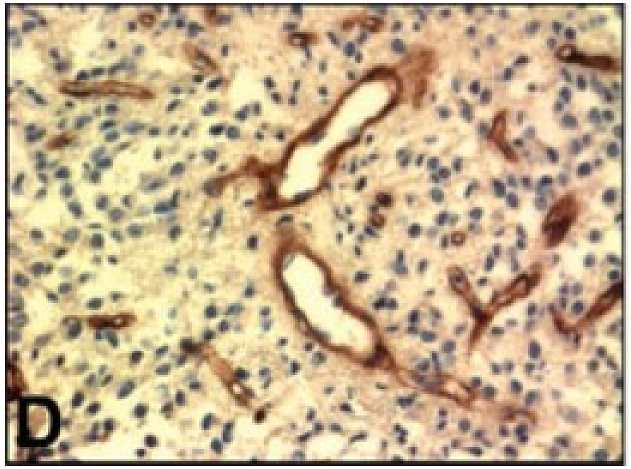


**Fig 1.** Immunolocalization of perlecan protein core in the blood vessels of normal human breast (**A**), renal arteriole (**B**), renal tubules (**C**), astrocytoma (**D**), renal cell carcinoma (**E**), and in the stroma of an ovarian cancer (**F**). The dark brown pigment is indicative of strong perlecan expression and deposition

### HSPG2 (Perlecan)

**PMID 31116062**

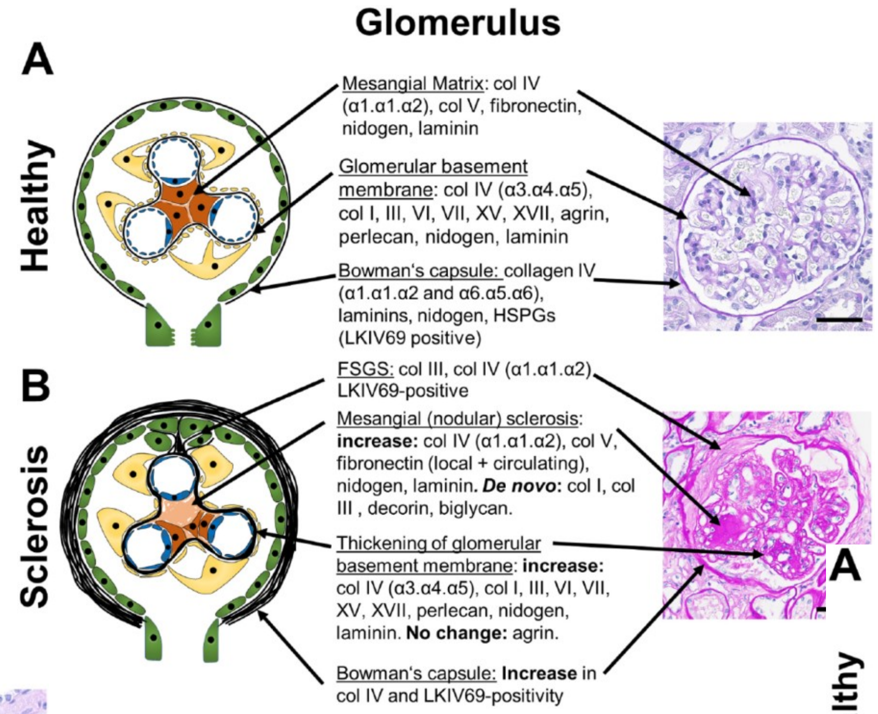


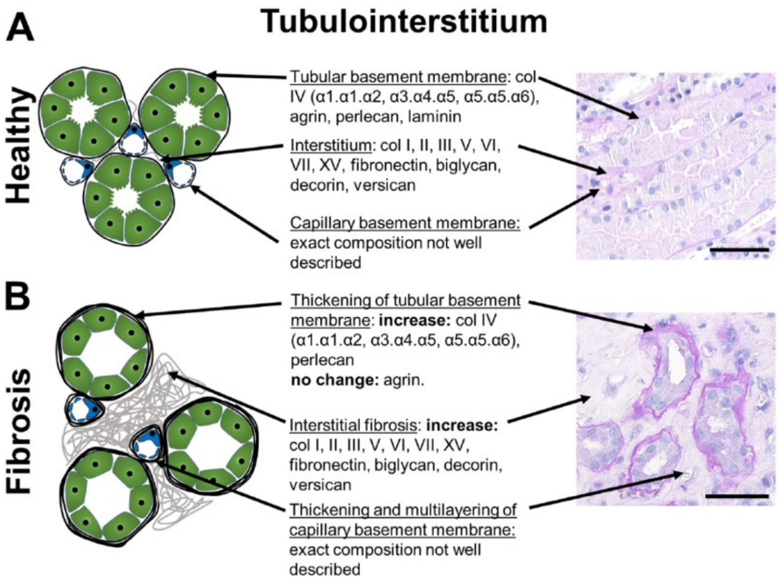


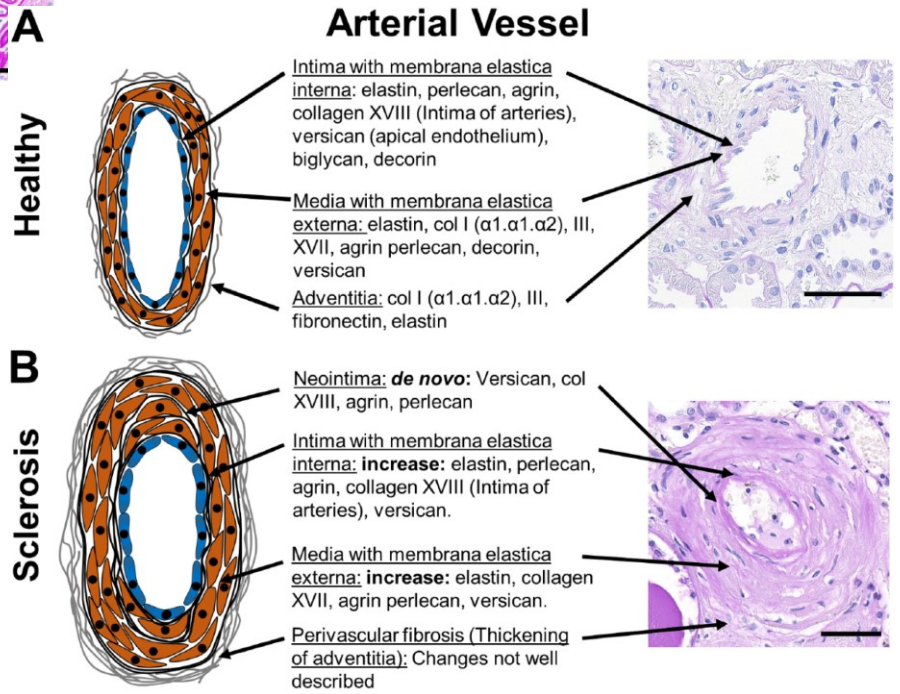

### HSPG2 (Perlecan)

https://www.lsbio.com/antibodies/ihc-plus-heparan-sulfate-proteoglycan-antibody-icc-ihc-ip-wb-western-ls-b5244/139793

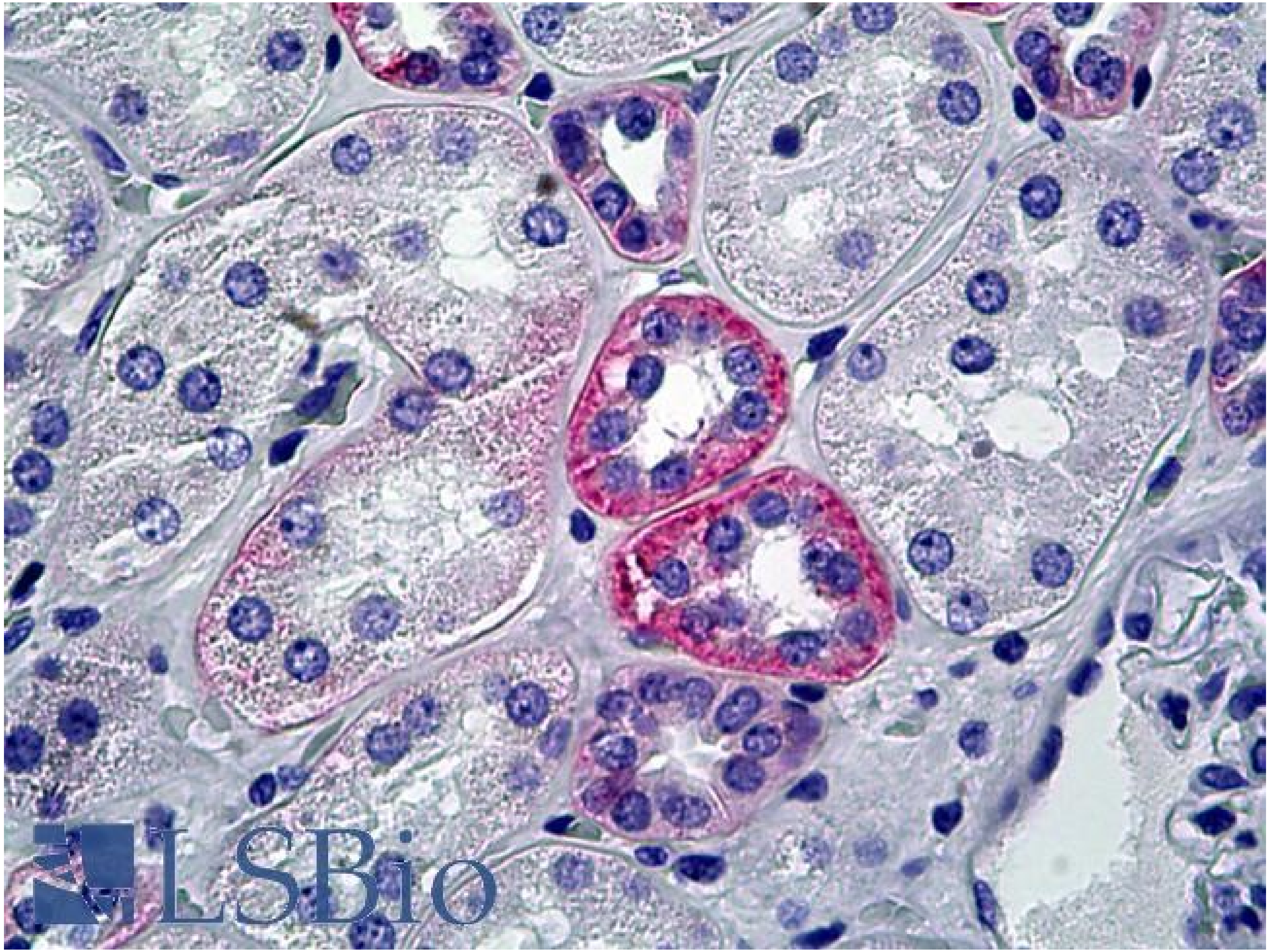


Anti-Heparan Sulfate Proteoglycan antibody IHC of human kidney. Immunohistochemistry of formalin-fixed, paraffin-embedded tissue after heat-induced antigen retrieval. Antibody concentration 10 ug/ml.

### Nestin

#### Human Protein Atlas

HPA007007

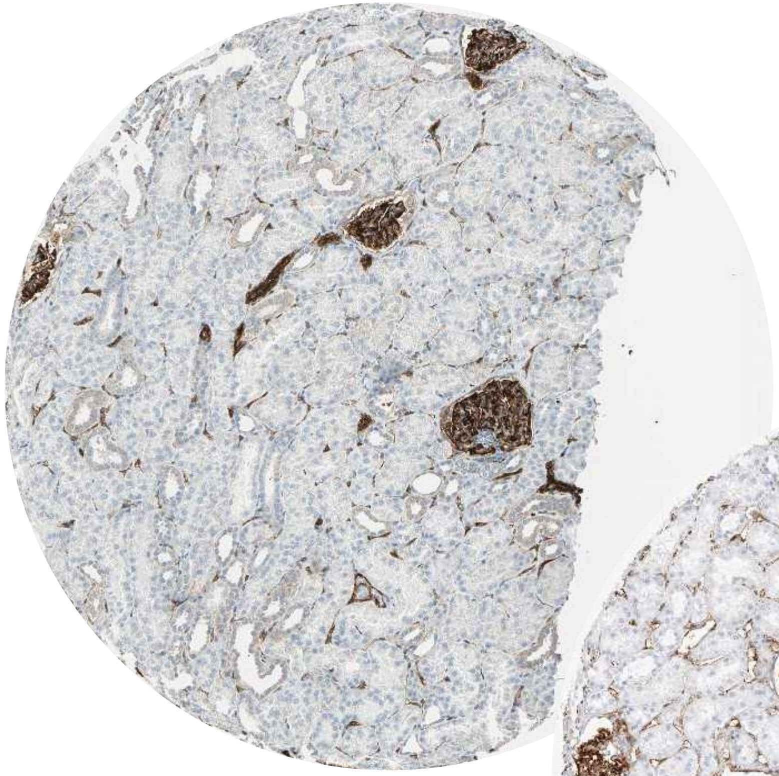

HPA026111

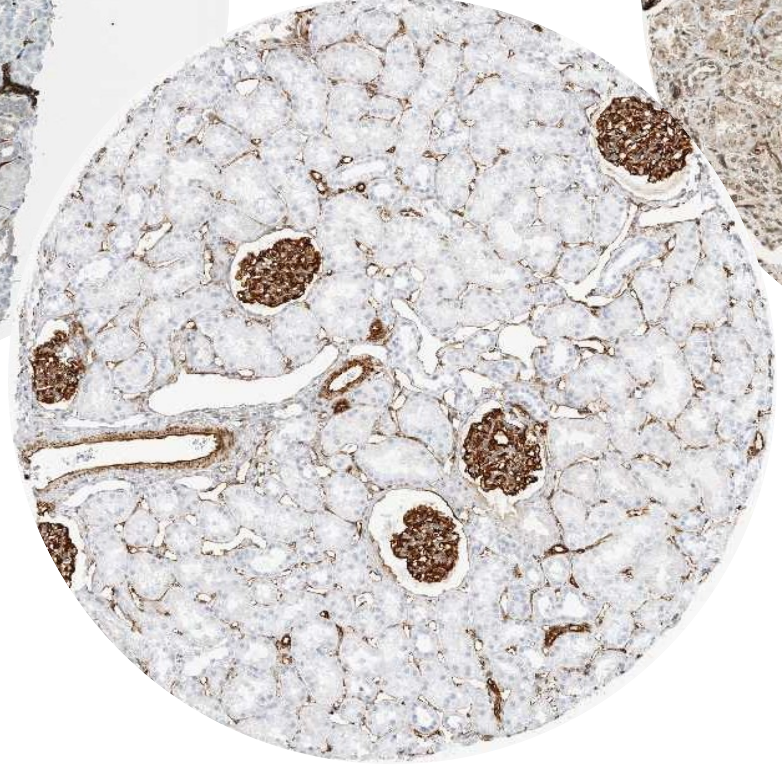

CAB005889

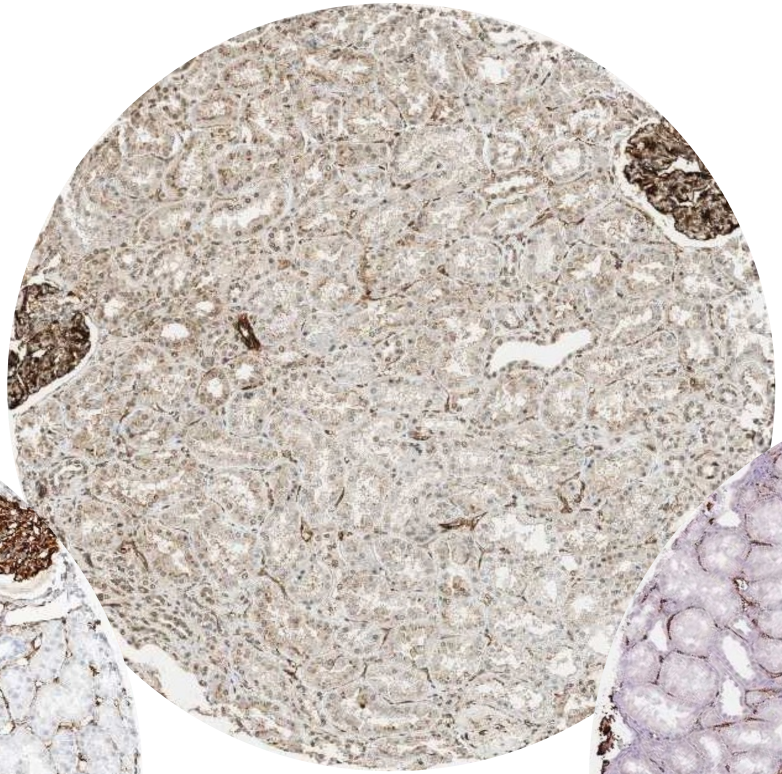

CAB058692

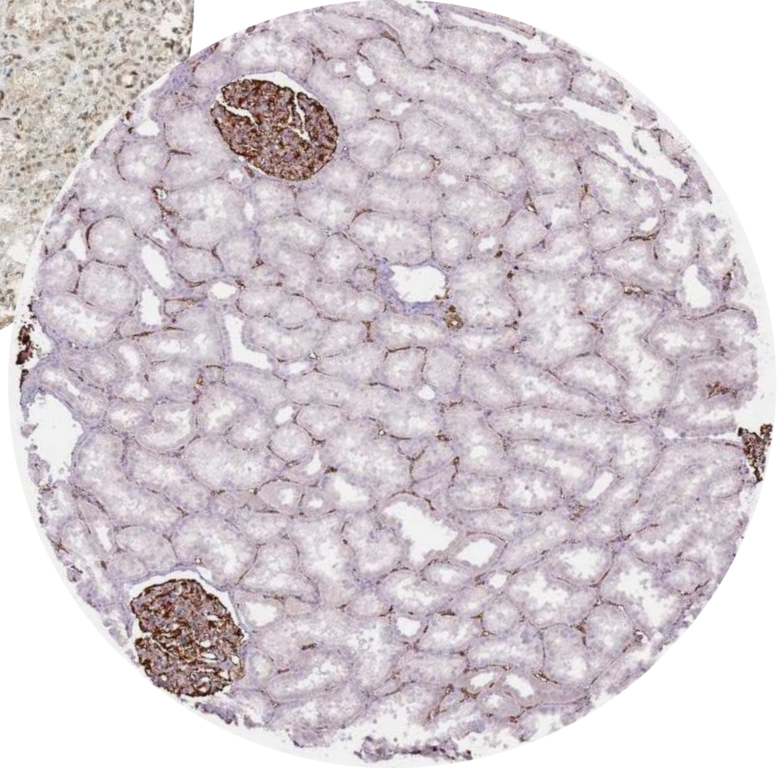

### Nestin

#### Human Protein Atlas – 4/5 antibodies worked for IHC

- Atlas Antibodies, rabbit pAb (HPA007007)
  - Immunogen:DPEGQSQQVGAPGLQAPQGLPEAIEPLVEDDVAPGGDQASPEVMLGSEPAMGESAAGAEPGPGQGVGGLGDPGHLTREEVMEPPLEEESLEAKRVQGLEGPRKDLEEAGGLGTEFSELP
- Validation:
  - ICC- No data.
  - **IHC** (HIER pH 6)- **Enhanced/orthogonal:**
    - Protein distribution across 45 tissues similar between the independent antibodies HPA007007 and HPA026111
    - staining mainly consistent with RNA expression data across 42 tissues.(high and low expression in spleen and pancreas below).
  - **WB- Enhanced:** Downregulation visible in both siRNA lanes
  - **Protein array- Supported:** Pass with single peak corresponding to interaction only with its own antigen.

ICC

N/A

IHC

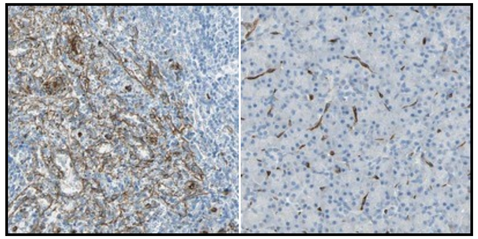

High expression Low expression

WB

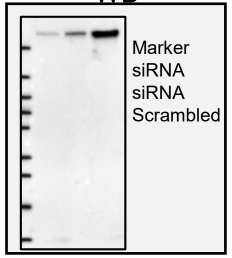


Protein Array

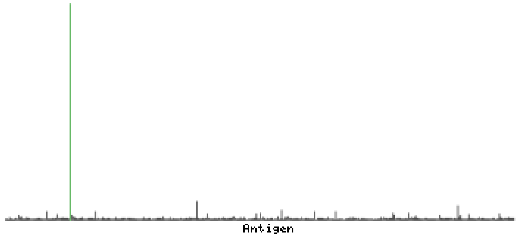


- Atlas Antibodies, rabbit pAb (HPA026111)
  - Immunogen:EPLRSLEDENKEAFRSLEKENQEPLKTLEEEDQSIVRPLETENHKSLRSLEEQDQETLRTLEKETQQRRRSLGEQDQMTLRPPEKVDLEPLKSLDQEIARPLENENQEFLKSLKEESVEAVKSLETEILESLKSAGQENLETLK
- Validation:
  - **ICC- Enhanced:** Antibody staining overlaps with antibody HPA006286.
  - **IHC** (HIER pH 6)- **Enhanced/orthogonal:**
    - Protein distribution across 45 tissues similar between the independent antibodies HPA007007 and HPA026111
    - staining mainly consistent with RNA expression data across 45 tissues.(high and low expression in spleen and pancreas below).
  - WB- Uncertain: Weak band of predicted size but with additional bands of higher intensity in standard panel of samples.
  - **Protein array- Supported:** Pass with single peak corresponding to interaction only with its own antigen.

ICC

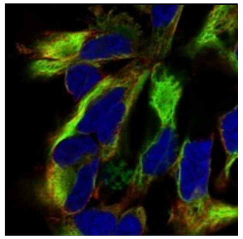

IHC

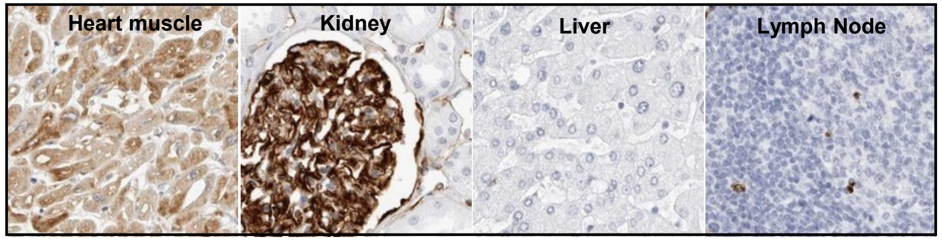


WB

N/A

Protein Array

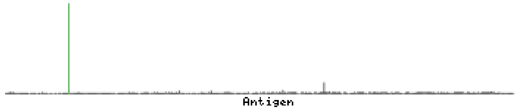

### Nestin

#### Human Protein Atlas – 4/5 antibodies worked for IHC

- Abcam, Rabbit pAb (CAB005889)
  - Immunogen (PMID 29205371): QFLKFTQREGDRESWSSGED

- Validation:
  - **ICC- Supported:** Subcellular location supported by literature. IF staining of human U2OS cell line shows localization to intermediate filaments.
  - **IHC (HIER pH 6)- Enhanced/Orthogonal:** Antibody staining mainly consistent with RNA expression data across 44 tissues (high and low expression in spleen and pancreas below).
  - WB- Uncertain: Weak band of predicted size but with additional bands of higher intensity in standard panel of samples.
  - Protein array- No data.

ICC

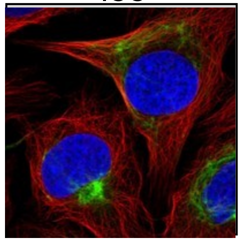

IHC

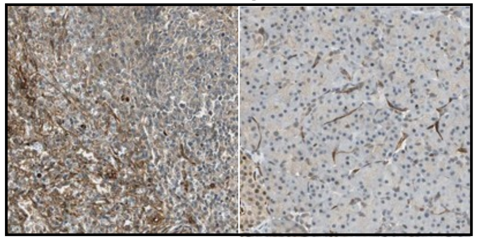

High expression Low expression

WB

N/A

Protein Array

N/A

- Atlas Antibodies, mouse mAb (CAB058692)
  - Immunogen: Binds to an epitope located within the peptide sequence VGGLGDPGHL as determined by overlapping synthetic peptides

- Validation:
  - ICC- No data.
  - **IHC (HIER pH 6)- Enhanced/Orthogonal:** Antibody staining mainly consistent with RNA expression data across 44 tissues (high and low expression in spleen and pancreas below).
  - WB- Uncertain: Single band differing more than +/-20% from predicted size in kDa and not supported by experimental and/or bioinformatic data, in a standard panel of samples.
  - Protein array- No data.

ICC

N/A

IHC

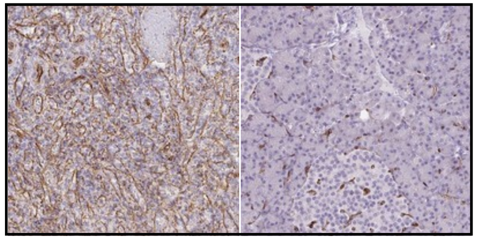

High expression Low expression

WB

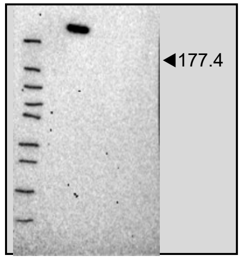


Protein Array

N/A

### Nestin

#### PMID 17929992

**Figure 1. Immunohistochemical localization of nestin in human kidney.** (A) In mature kidney, nestin staining is restricted to the glomeruli and localizes to podocytes. (B) Nestin localizes to both the cell body and processes (arrows) of podocytes.

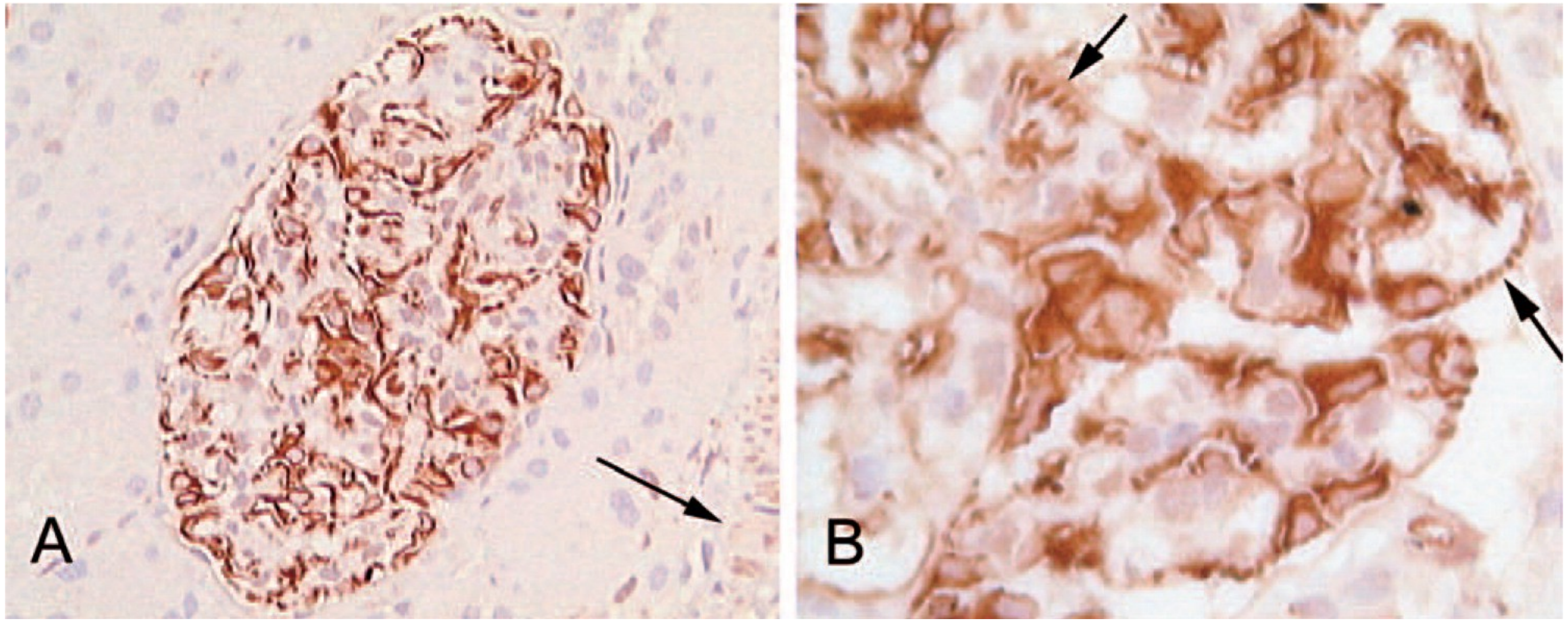

### Nestin

#### PMID 17210924

**Figure 1 Nestin expression in adult kidney.** (A-C) Nestin staining is located within glomeruli. (A) A glomerulus with an arteriole at the vascular pole (arrow). Staining is observed in the endothelial cells of the arteriole. (B,C) A glomerulus at low (B) and higher magnification (C). Staining is restricted to cells with the morphological features of podocytes. (D-L) Confocal microscopy analysis of nestin/vimentin (D-I) and nestin/CD31 (J-L) double-labeling experiments. (D-F) All nestin+ cells (red) costain with vimentin (green). (G-I) The morphology of nestin/vimentin double-labeled cells is typical of podocytes, with long and branched primary processes. (J-L) Endothelial CD31+ cells do not co-stain with nestin. Bars: A-C = 100 μm; D-L = 10 μm.

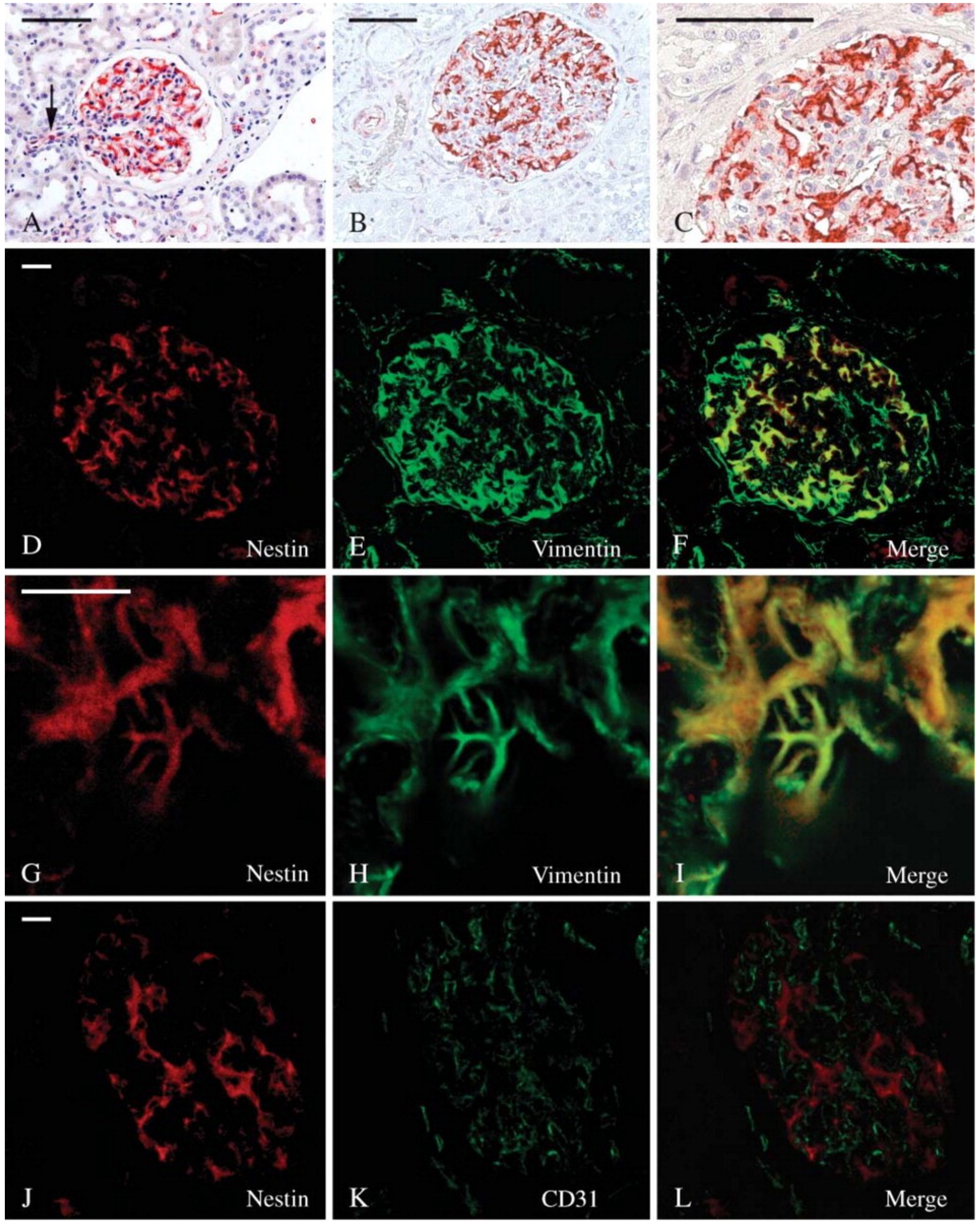

### Nestin

https://www.novusbio.com/products/nestin-antibody-10c2_nb300-266

Immunohistochemistry-Paraffin: Nestin Antibody (10C2) [NB300-266] - Nestin was detected in immersion fixed paraffin-embedded sections of human kidney using anti-human mouse monoclonal antibody (Catalog # NB300-266) at 1:500 dilution overnight at 4C. Tissue was stained using the VisuCyte anti-mouse HRP polymer detection reagent (Catalog # VC001) with DAB chromogen (brown) and counterstained with hematoxylin (blue).Images may not be copied, printed or otherwise disseminated without express written permission of Novus Biologicals a bio-techne brand.

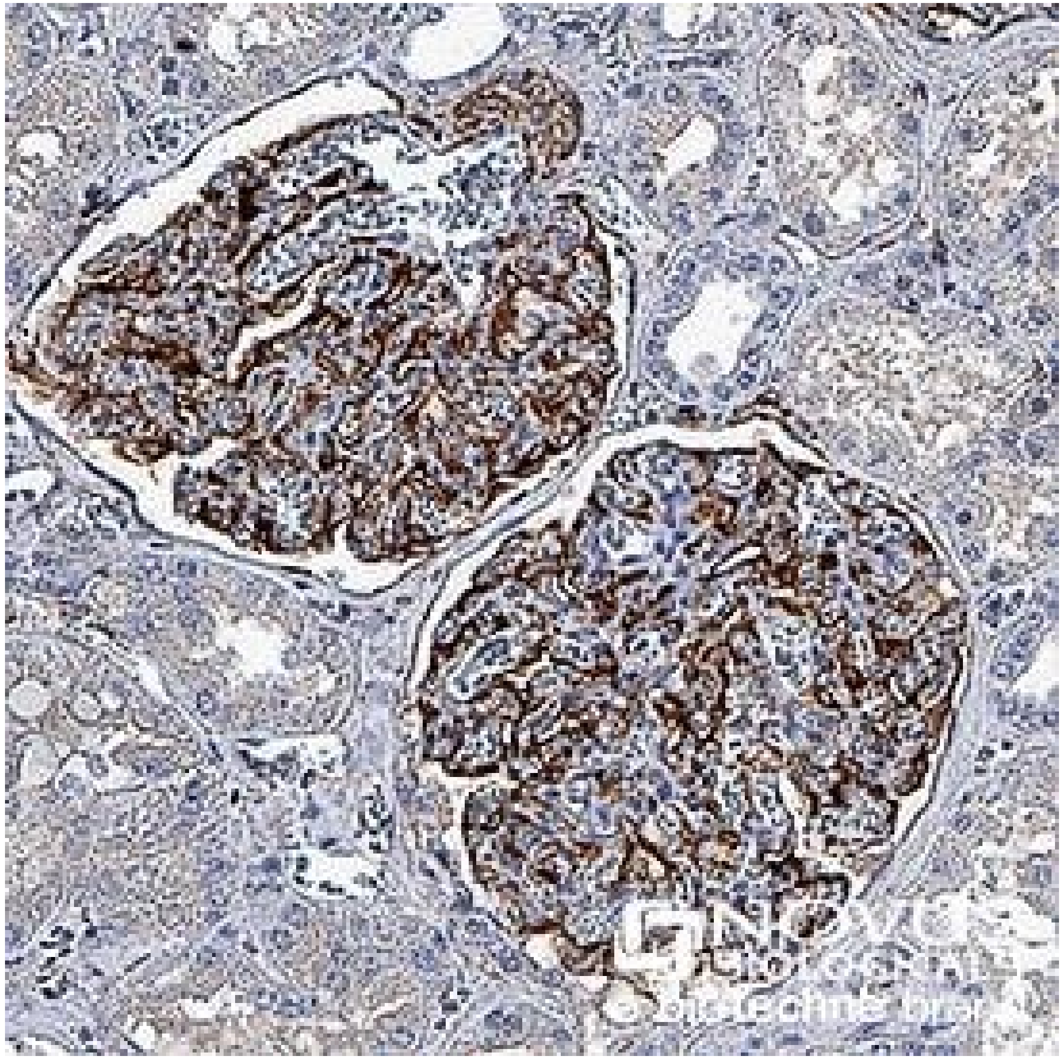

### RORγ (RORC)

#### Human Protein Atlas

- One pAb – pending normal tissue annotation

- Atlas Antibodies rabbit pAb (HPA065620)

- Validation:
  - IHC- Pending annotation.
  - ICC- supported: Immunofluorescent staining of human cell line HeLa shows localization to nuclear bodies.
  - WB- uncertain: Only visualized bands did not correspond to the predicted size in analysis including a standard sample panel
  - Protein array- supported: single peak corresponding to interaction with its own antigen

ICC

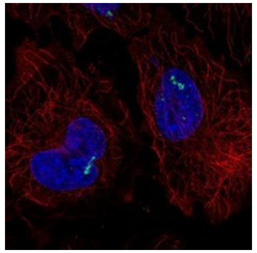

IHC

N/A

WB

N/A

Protein Array

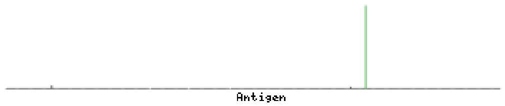

### RORγ (RORC)

**PMID: 27481185**
Isoform 1 (RORγ1) is widely expressed in many tissues, including liver, kidney, lung, muscle, heart, brain and adipose tissue.
Isoform 2 (RORγ2 or RORγt) is primarily expressed in immature thymocytes and some immune cells.

**Fig. 1**. Overview of the ROR family members.

(A) Schematic representation of the domain structure of the three ROR family members (α, β, γ) and their isoforms. The isoforms are generated by alternative promoter usage and/or alternative splicing in the variable A/B domain. The other regions of the proteins are the DNA-binding domain (DBD), the hinge region and the ligand-binding domain (LBD) containing the activation function 2 (AF2). The size of the protein in terms of number of amino-acids is indicated on the right. (B) Organization of the RORc locus. The usage of exons 1γ and 2γ produces the RORγ protein. These are replaced by a unique exon (exon 1γt) to produce the immune-specific isoform RORγt. As a result, the 24 first amino-acids of RORγ are replaced by 3 residues from exon 1γt.

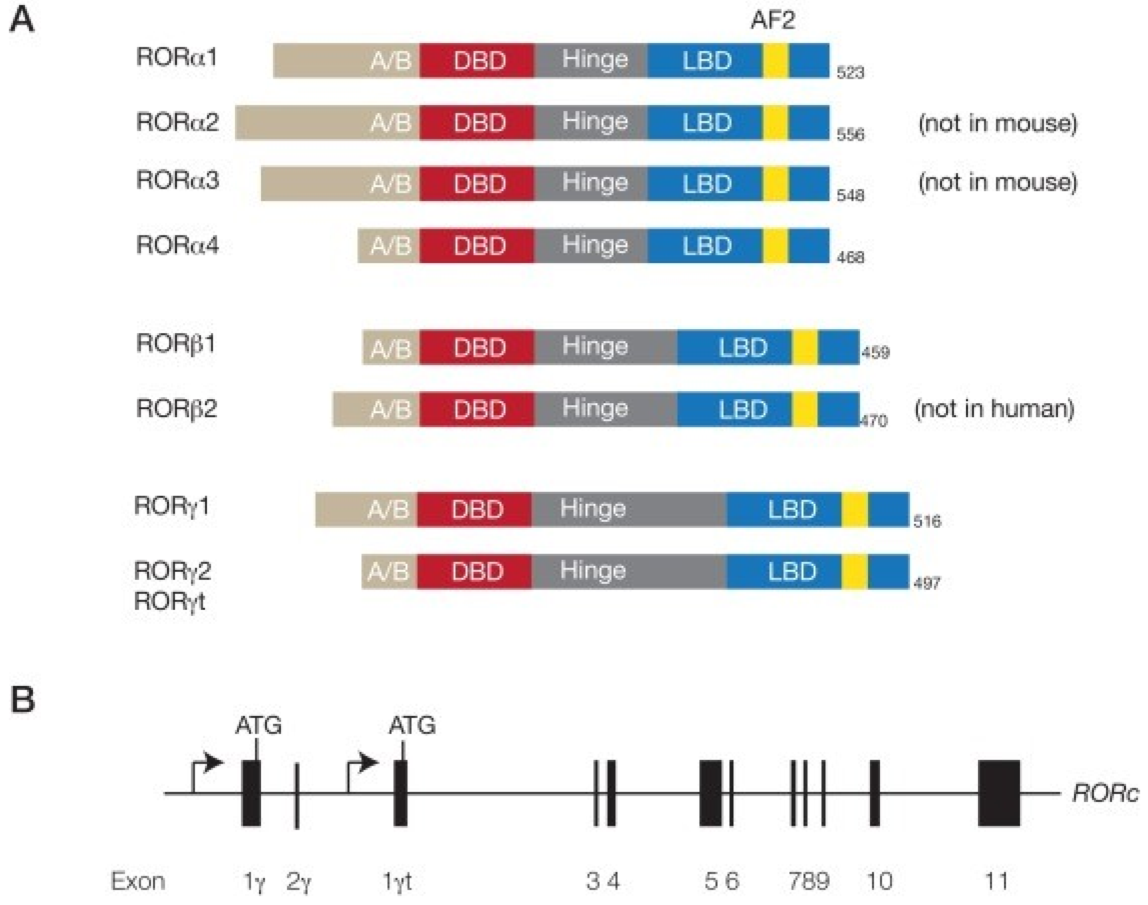

### RORγ (RORC)

#### PMID 25970244

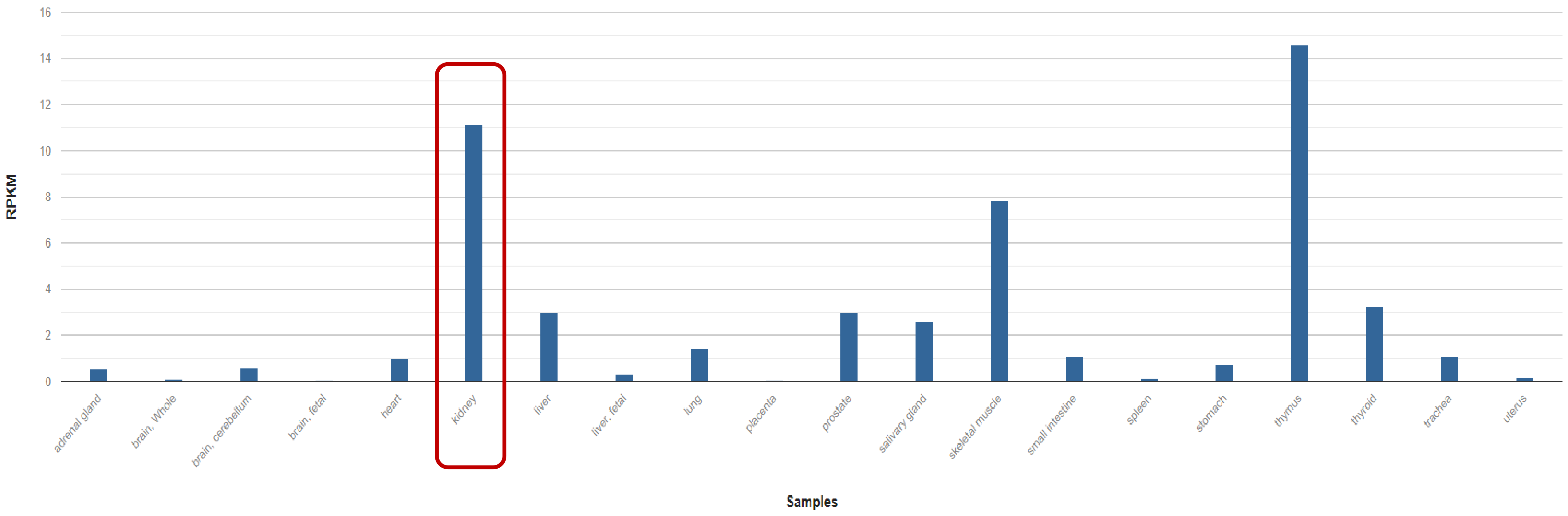

### RORγ (RORC)

**PMID: 9881970**

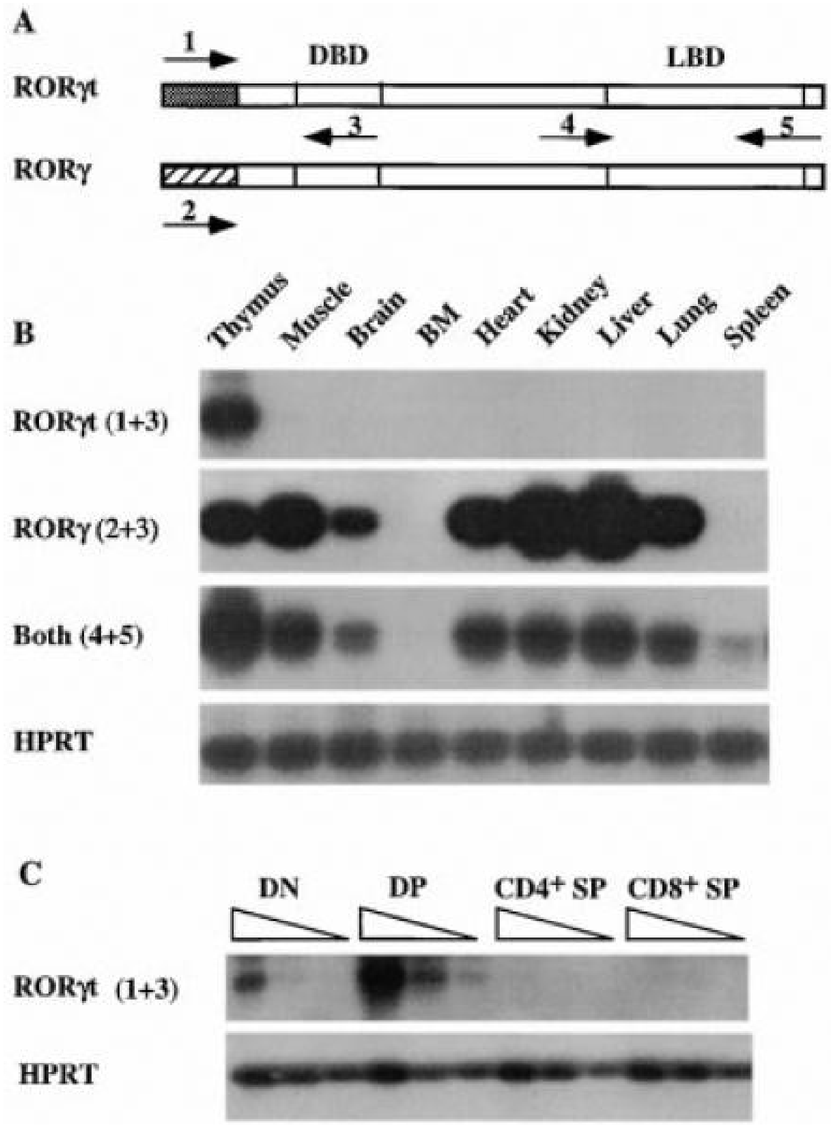


**Figure 2. Differential Expression of RORγt and RORγ in mouse tissues**
(A) Schematic representation of RORγt and RORγ cDNAs and the locations of the primers used in (B) and (C). The shaded region represents the distinct nucleotide sequences. Specific primers are 1 and 2. Common primers are 3, 4, and 5. Diagram is not to scale.

(B) Expression of RORγt and RORγ. RT-PCR products of different tissues were blotted and probed with a full-length RORγt cDNA. HPRT serves as an internal control. Numbers in parentheses indicate the primers shown in (A) used in the PCR.

(C) Expression of RORγt in thymocyte subpopulations. RT reactions of different thymocyte subpopulations were serially diluted at 1:3 and subjected to PCR with primers shown in (A). PCR products were blotted and probed as in (B). HPRT RT-PCR was performed from the same RT samples for cDNA template quantity control.

### RORγ (RORC)

**PMID: 8973331**

At least mRNA is expressed in the kidney, ?two isoforms.

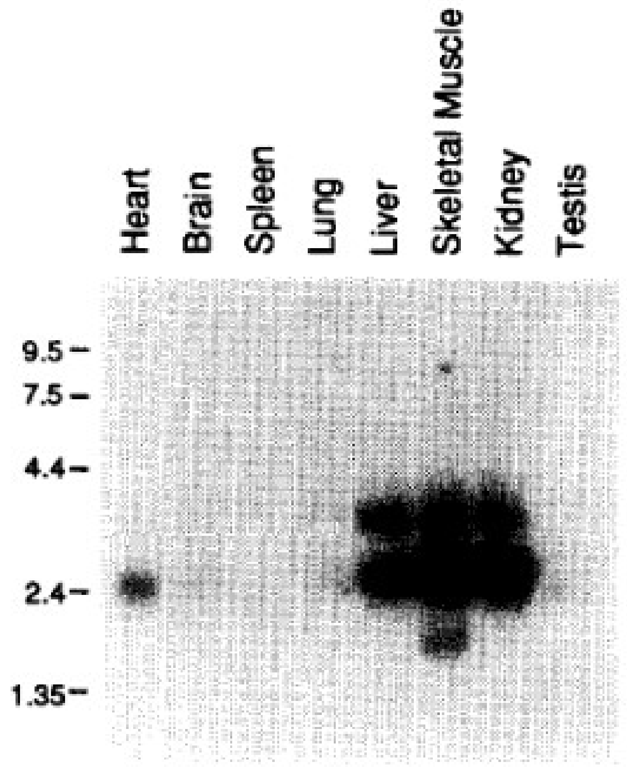


Fig. 2. Tissue-specific expression of mRZR/RORγ. **Methods:** Poly(A)$^+$ RNA prepared from several mouse tissues (2 μg) were analyzed by Northern analysis using a $^{32}$P-labeled RZR/RORγ probe. RNA size markers are indicated on the left side of the blot.

### RORγ (RORC)

**PMID: 19381306-** RORγ exhibits a strong oscillatory expression pattern in kidney (16267379). RORγ is expressed at low levels during the day and at optimum levels at night.

**PMID: 22753030**

**Fig 1.** Oscillatory pattern of expression of RORγ1, RORα1 and RORα4 in mouse BAT, kidney, WAT and small intestines. Liver, BAT, kidney, WAT and small intestines (jejunum) from WT, RORαsg/sg and RORγ−/− mice (n = 4) were isolated every 4 h over a period of 24 h. Subsequently, the expression of RORγ1, RORα1 and RORα4 was analyzed by QRT-PCR. The oscillatory expression patterns in liver were similar to those previously reported (25) using samples from different WT mice. The 24 h expression pattern was double-plotted. The open and solid boxes indicate the 12 h light and dark periods, respectively. Data represent mean ±SD; *P

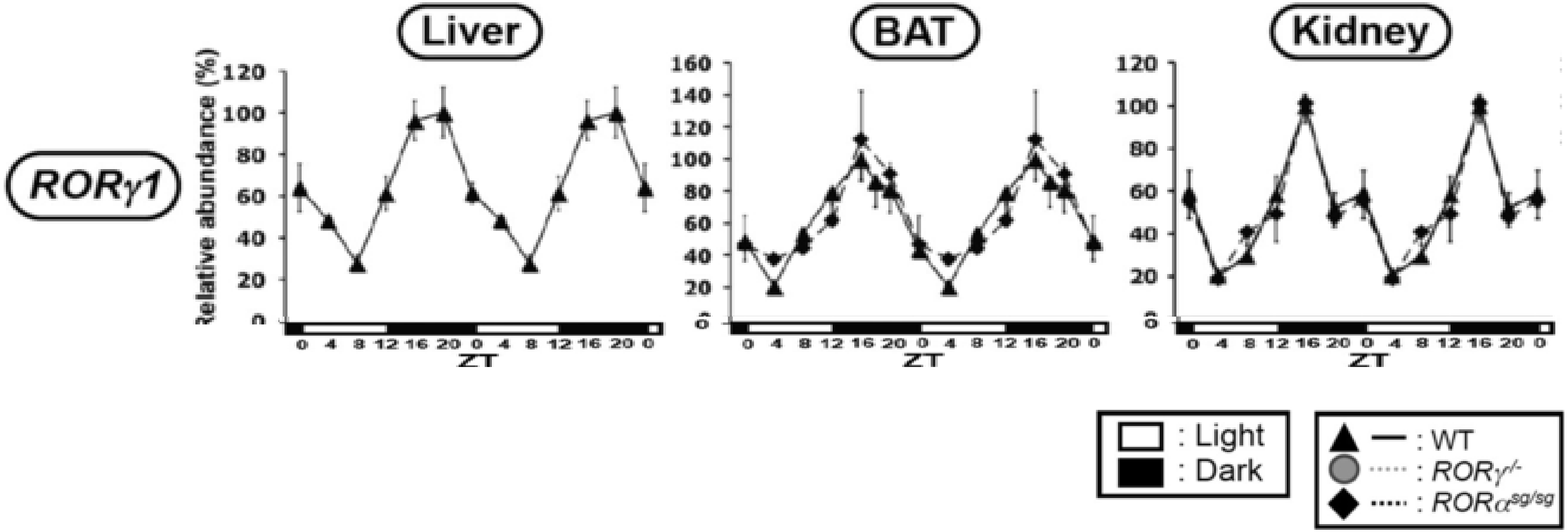

### RORγ (RORC)

**mAb: Anti-ROR gamma T Antibody, clone 6F3.1 (MABF81, Millipore Sigma), IgG2ak**

Antigen: GST-tagged recombinant protein corresponding to human ROR gamma T.

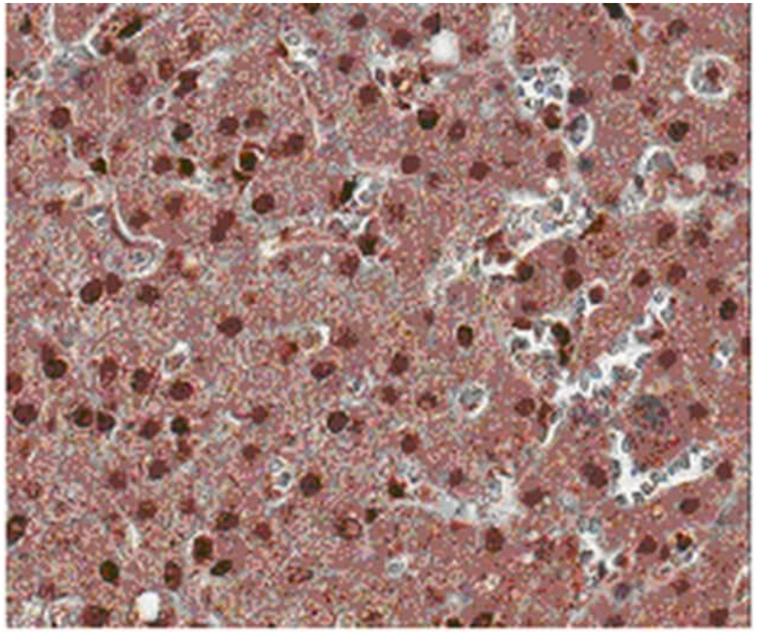

IHC Analysis: Representative lot data. FFPE normal human liver tissue was prepared using HIER. Immunostaining was performed using a 1:50 dilution of MABF81, anti-ROR gamma T, clone 6F3.1. Reactivity was detected using an anti-mouse secondary antibody and HRP-DAB. Positive staining was observed in hepatocytes from normal human liver tissue.

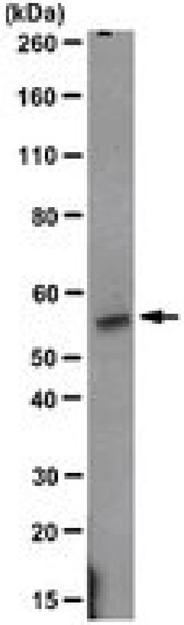


Western blotting Analysis: Representative lot data. T47D cell lysate was probled withi Anti-RORgamma T, clone 6F3.1 (1 ug/ml). Proteins were visualized using a goat anti-mouse IgG secondary antibody conjugated to HRP and a chemiluminescence detection system. Arrow indicates ROR gamma T (~55 kDa).

### RORγ (RORC)

https://www.thermofisher.com/antibody/product/RORG-Antibody-Polyclonal/BS-6217R

Formalin-fixed and paraffin embedded human kidney carcinoma labeled with Rabbit Anti RORG/ROR gamma Polyclonal Antibody, Unconjugated (bs-6217R) at 1:200 followed by conjugation to the secondary antibody and DAB staining.

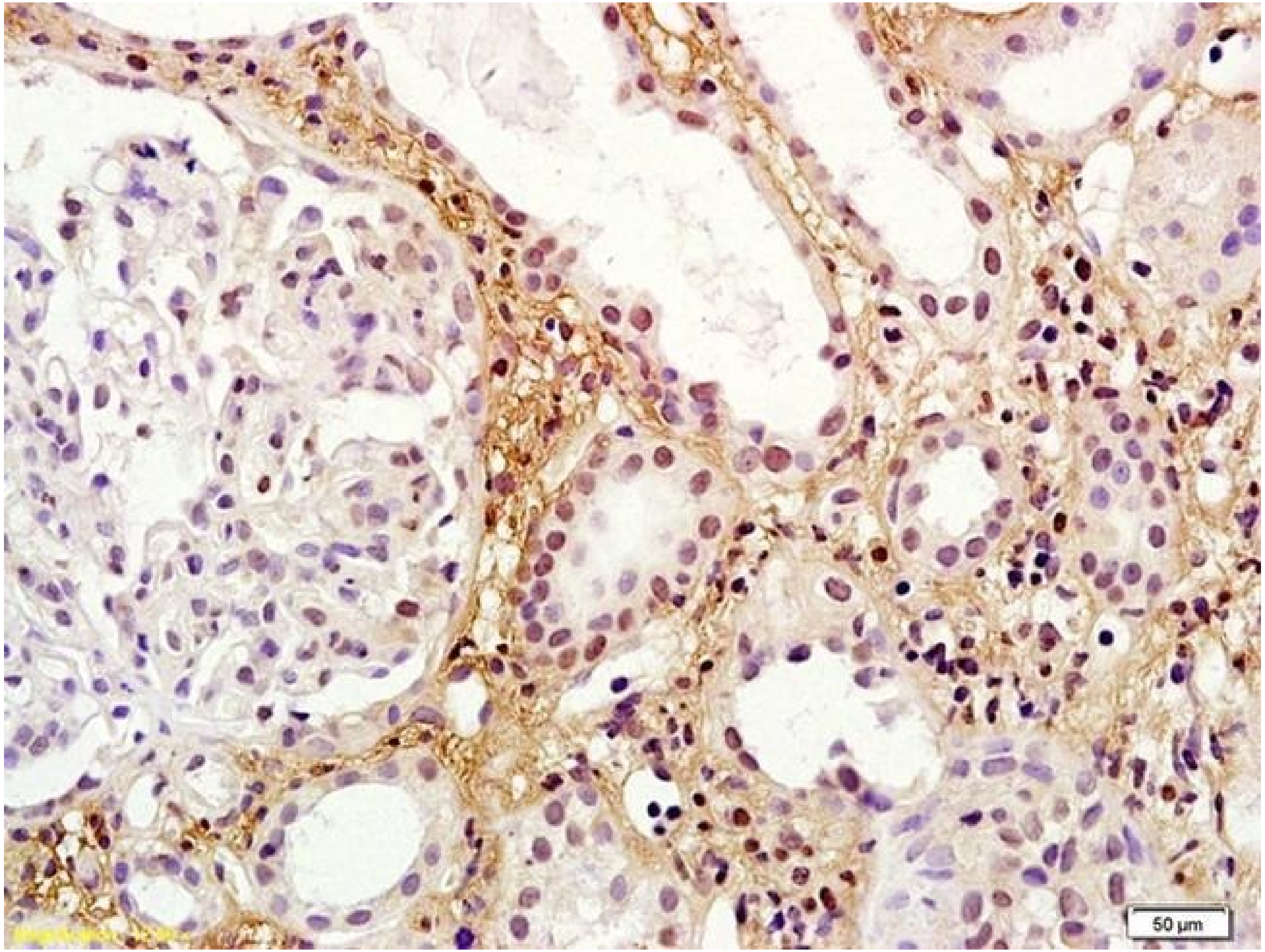

### SPP1 (OPN)

```
3 MRIAVICFCLLGITCAIPVKQADSGSSEEK---------------------------QNAVSSEET 39
4 MRIAVICFCLLGITCAIPVKQADSGSSEEKQLYNKYPDAVATWLNPDPSQKQNLLAPQNAVSSEET 66
1 MRIAVICFCLLGITCAIPVKQADSGSSEEKQLYNKYPDAVATWLNPDPSQKQNLLAPQNAVSSEET 66
5 MRIAVICFCLLGITCAIPVKQADSGSSEEKQLYNKYPDAVATWLNPDPSQKQNLLAPQ-------- 58

3 NDFKQETLPSKSNESHDHMDDMDDEDDDDHVDSQDSIDSNDSDDVDDTDDSHQSDESHHSDESDEL 105
4 NDFKQETLPSKSNESHDHMDDMDDEDD----------------------DSHQSDESHHSDESDEL 110
1 NDFKQETLPSKSNESHDHMDDMDDEDDDDHVDSQDSIDSNDSDDVDDTDDSHQSDESHHSDESDEL 132
5 ------TLPSKSNESHDHMDDMDDEDDDDHVDSQDSIDSNDSDDVDDTDDSHQSDESHHSDESDEL 118

3 VTDFPTDLPATEVFTPVVPTVDTYDGRGDSVVYGLRSKSKKFRRPDIQYPDATDEDITSHMESEEL 171
4 VTDFPTDLPATEVFTPVVPTVDTYDGRGDSVVYGLRSKSKKFRRPDIQYPDATDEDITSHMESEEL 176
1 VTDFPTDLPATEVFTPVVPTVDTYDGRGDSVVYGLRSKSKKFRRPDIQYPDATDEDITSHMESEEL 198
5 VTDFPTDLPATEVFTPVVPTVDTYDGRGDSVVYGLRSKSKKFRRPDIQYPDATDEDITSHMESEEL 184

3 NGAYKAIPVAQDLNAPSDWDSRGKDSYETSQLDDQSAETHSHKQSRLYKRKANDESNEHSDVIDSQ 237
4 NGAYKAIPVAQDLNAPSDWDSRGKDSYETSQLDDQSAETHSHKQSRLYKRKANDESNEHSDVIDSQ 242
1 NGAYKAIPVAQDLNAPSDWDSRGKDSYETSQLDDQSAETHSHKQSRLYKRKANDESNEHSDVIDSQ 264
5 NGAYKAIPVAQDLNAPSDWDSRGKDSYETSQLDDQSAETHSHKQSRLYKRKANDESNEHSDVIDSQ 250

3 ELSKVSREFHSHEFHSHEDMLVVDPKSKEEDKHLKFRISHELDSASSEVN Isoform 3 (OPN-c) 287
4 ELSKVSREFHSHEFHSHEDMLVVDPKSKEEDKHLKFRISHELDSASSEVN Isoform 4 (OPN-d) 292
1 ELSKVSREFHSHEFHSHEDMLVVDPKSKEEDKHLKFRISHELDSASSEVN Isoform 1 (OPN-a, OPN1B) canonical 314
5 ELSKVSREFHSHEFHSHEDMLVVDPKSKEEDKHLKFRISHELDSASSEVN Isoform 5 (OPN-b, OP1A) 300
```

### SPP1 (OPN)

#### Human Protein Atlas – two antibodies, one worked for IHC

- Atlas Antibodies, rabbit pAb (HPA027540)
  - Immunoge: VKQADSGSSEEKQLYNKYPDAVATWLNPDPSQKQNLLAPQNAVSSE ETNDFKQETLPSKSNESHDHMDDMDDEDDDDHVD
- Validation:
  - **ICC- Enhanced:** Overlapping staining with HPA027541.
  - IHC- No data.
  - WB- Uncertain: No bands detected in analysis including a standard sample panel
  - **Protein array- supported:** single peak corresponding to interaction with its own antigen

ICC

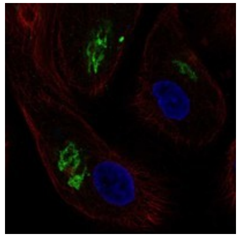

IHC

N/A

WB

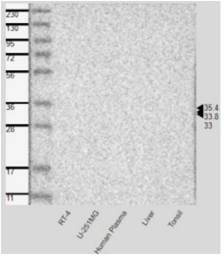


Protein Array

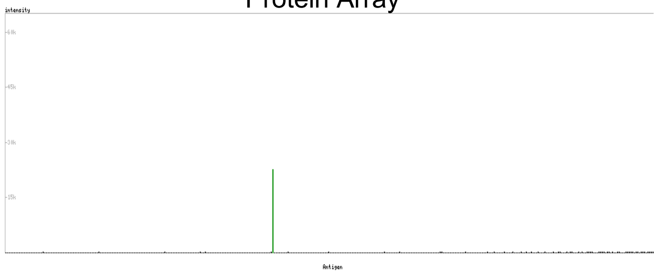

- Atlas Antibodies, rabbit pAb (HPA027541), IgG
  - Immunogen: SQDSIDSNDSDDVDDTDDSHQSDESHHSDESDELVTDFPTDLPATEVFTPVV PTVDTYDGRGDSVVYGLRSKSKKFRRPDIQYPDAT
- Validation:
  - **ICC- Enhanced:** overlapping staining with HPA027540
  - **IHC- Orthogonal:** staining mainly consistent with RNA expression across 45 tissues.
  - **WB- Enhanced** (recombinant expression): Band of expected size with extra band present
  - **Protein array- Approved:** Pass with quality comment- low specificity, binding to 1-2 antigens

ICC

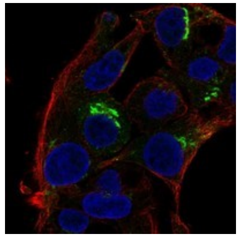

IHC

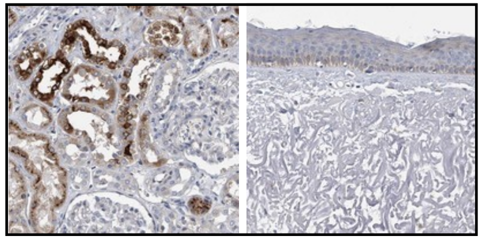

High expression Low expression

WB

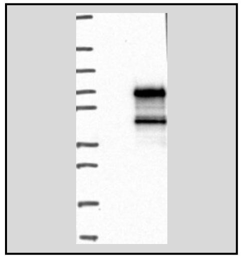

Protein Array

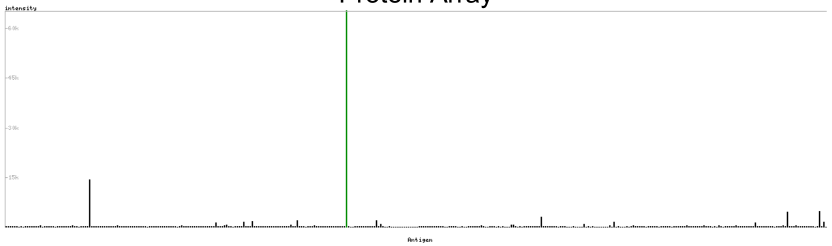

### SPP1 (Osteopontin)

**Human Protein Atlas**

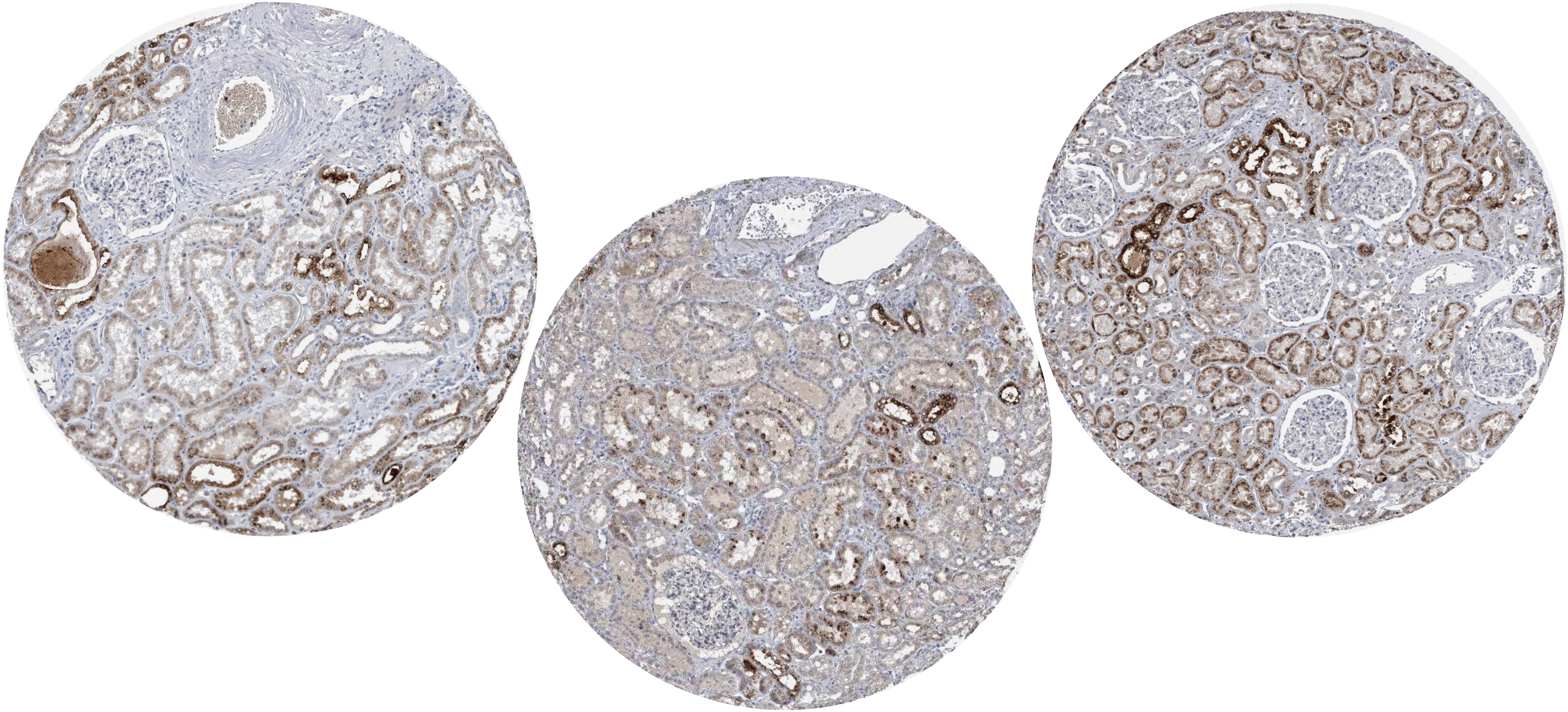

### SPP1 (Osteopontin)

**PMID: 11961008**

**Fig 1.** Osteopontin (OPN) immunostaining of human kidney tissue from implantation biopsies, using a peroxidase-diaminobenzidine procedure. Tissue sections were counterstained with periodic acid-Schiff reagent. Distal tubular cells (DTC) exhibited pronounced apical OPN immunostaining (black arrows); however, some DTC also exhibited OPN signals in perinuclear vesicles (white arrow). Slightly damaged proximal tubular cells (PTC) demonstrated perinuclear OPN staining (arrowheads). Magnification, ɤ 400.

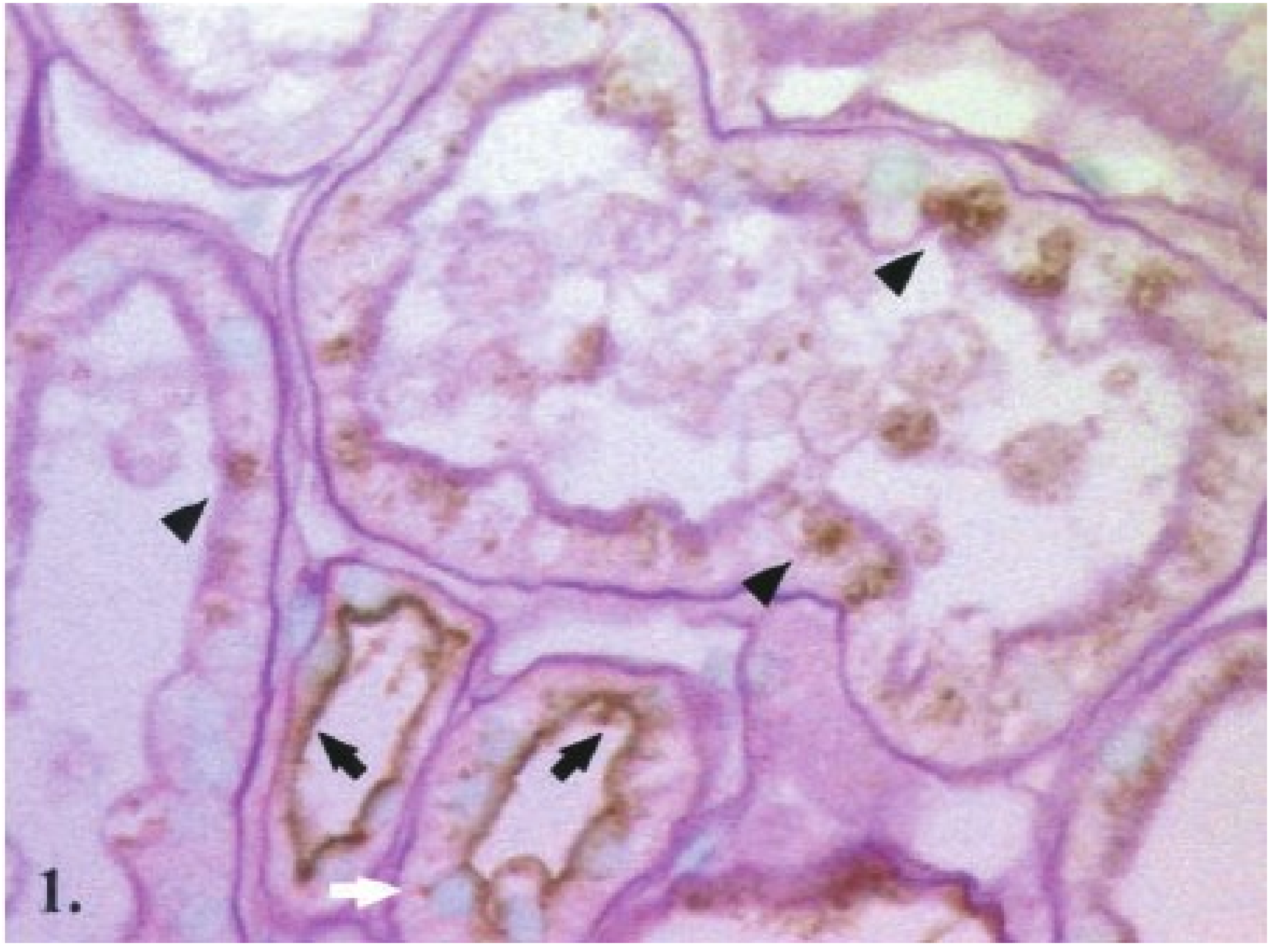

### SPP1 (Osteopontin)

**PMID: 10073594**

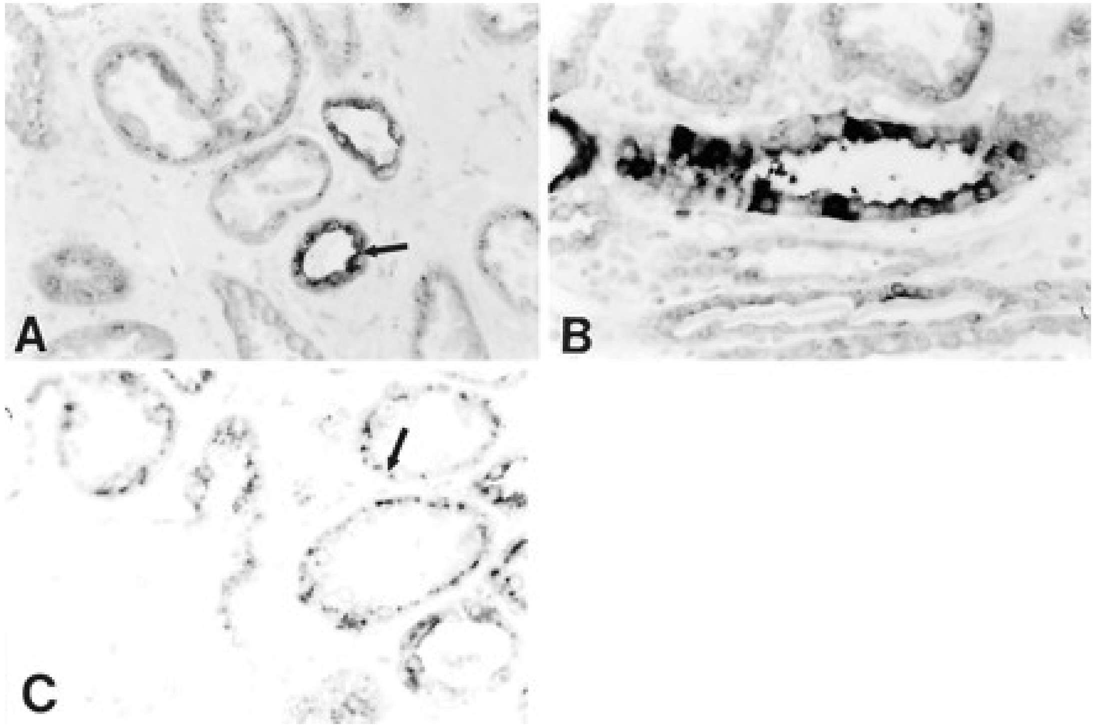


**Figure 4. Immunocytochemical demonstration of osteopontin expression in adult human kidney.** (A) Osteopontin is constitutively expressed in distal tubules of adult human kidney (arrow). (B) Osteopontin is expressed in individual cells of the collecting duct in some kidney sections. (C) Expression of osteopontin is seen in the proximal tubules of some adult kidneys. This expression is usually seen in a very distinct, perinuclear pattern (arrow). Magnification: ×600 in A through C.

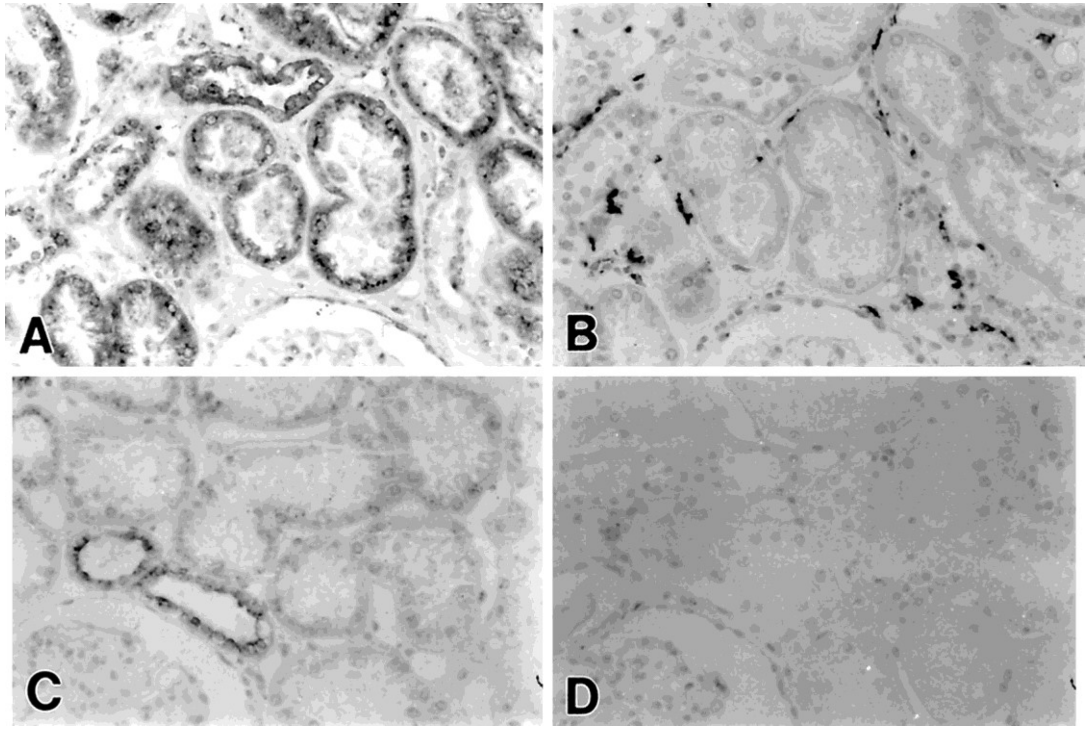


**Figure 7. Comparison of osteopontin expression and the presence of CD68-positive macrophages.** (A) Osteopontin expression in adult human kidney with tubulointerstitial fibrosis. Osteopontin is expressed by nearly all tubules present in the cortex. (B) Staining of serially sectioned tissue from the same specimen as A with anti-CD68 antibody, demonstrating widespread presence of interstitial macrophages. (C) Normal adult kidney with no expression of osteopontin in proximal tubular epithelium. Osteopontin immunoreactivity can be seen in two distal tubular segments directly adjacent to the glomerulus. (D) Serially sectioned tissue from the same specimen as C, stained with anti-CD68 antibody, showing essentially no macrophages present in the interstitium. Magnification: ×400 in A through D.

### SPP1 (Osteopontin)

**PMID 10073594**

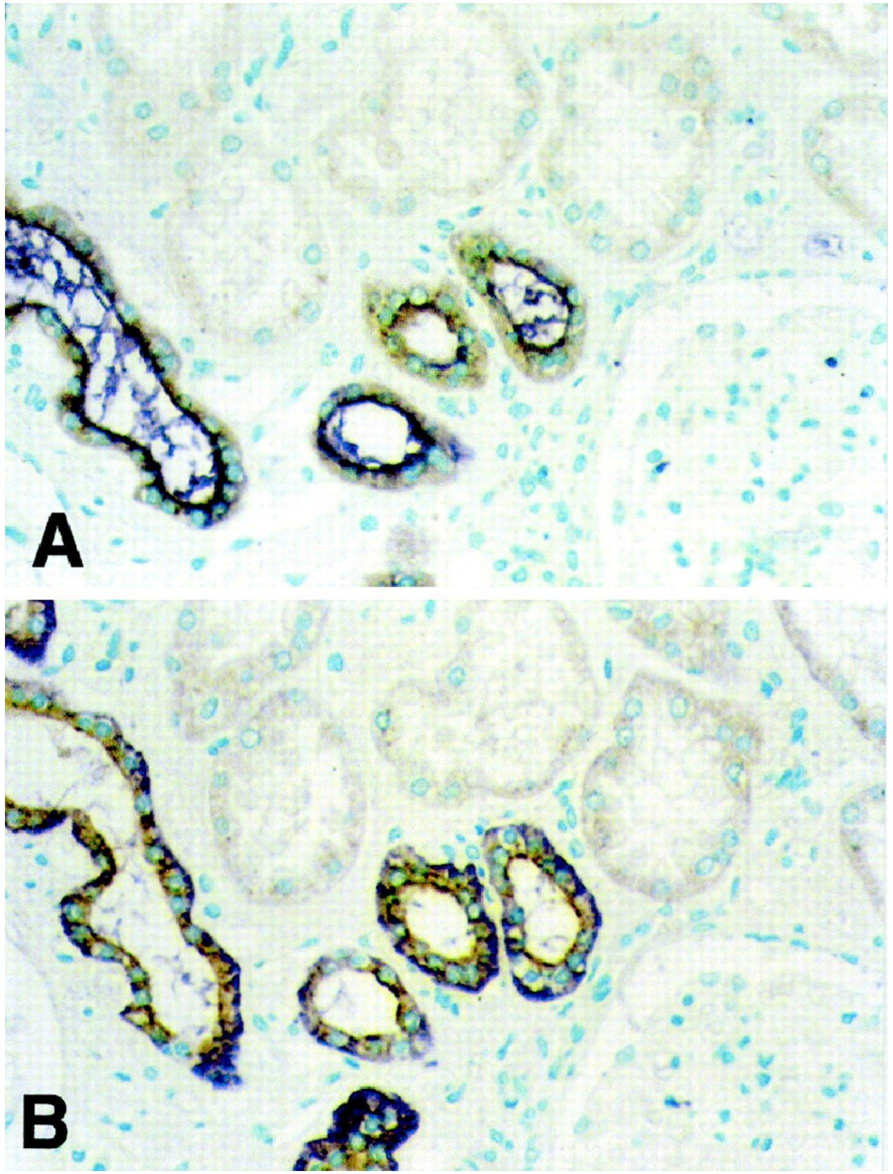


**Figure 9. Double immunohistochemistry for osteopontin and markers of distal tubules.** (A) Distal tubules produce osteopontin as shown by double labeling with antibodies to osteopontin (brown) and epithelial membrane antigen (EMA) (purple). (B) Replicate tissue section of that shown in A, demonstrating double labeling of the same distal tubules with osteopontin (brown) and Na,K-ATPase (purple) antibodies. Magnification: ×400 in A and B.

### SPP1 (Osteopontin)

**PMID 1421573**

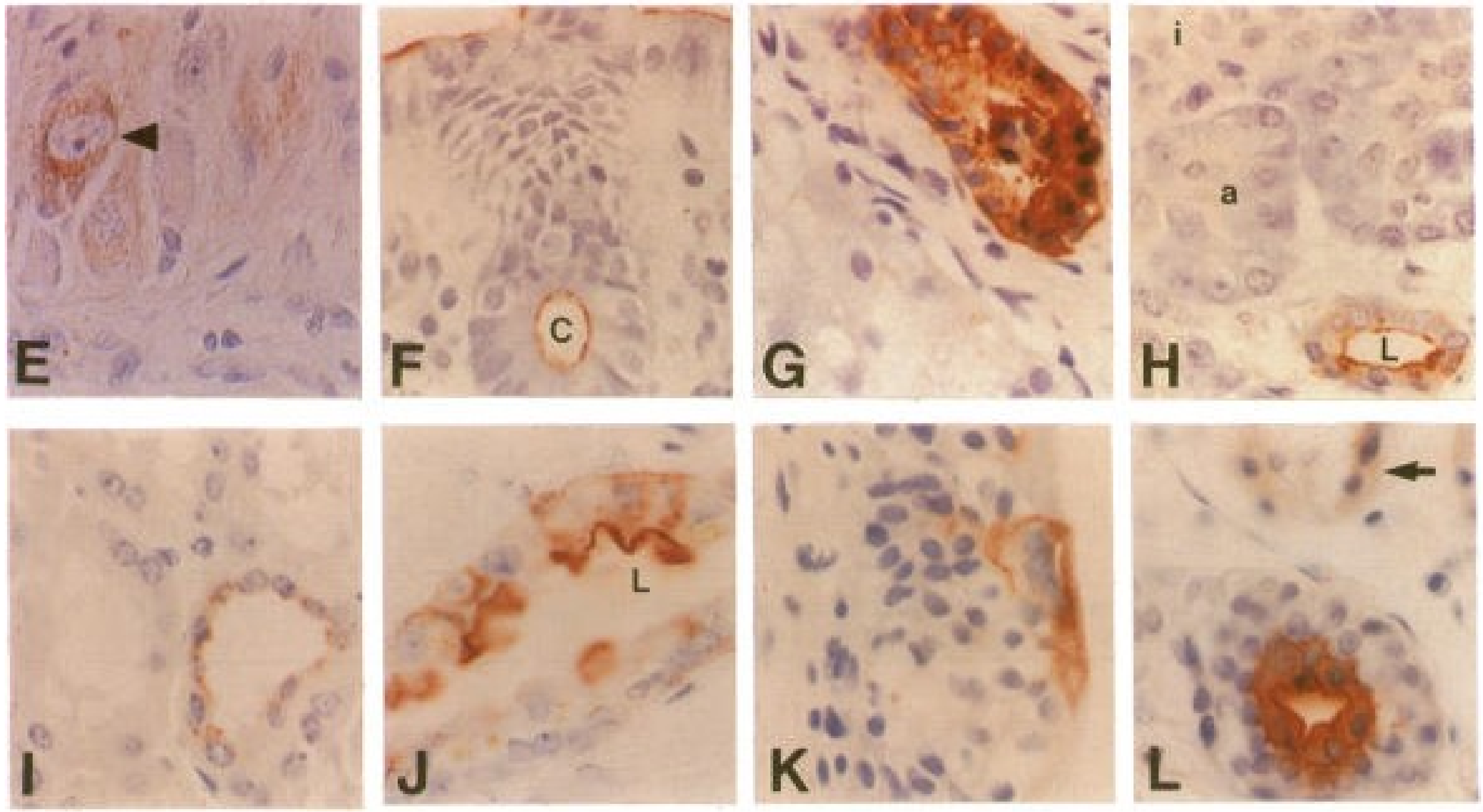


**Figure 1. Immunohistochemical staining of normal adult human tissues for osteopontin.** Positive reactions are denoted by red-colored reaction product. L, lumen. (E) Several ganglion cells within the myenteric plexus of small bowel muscle stain for osteopontin; the most intensely stained ganglion cell is indicated with an arrowhead. (F) Colonic epithelium stains with accentuation both at the luminal surface (top) and within a crypt lumen (c, bottom). (G) Intrahepatic bile duct epithelium stains, whereas hepatocytes (lower left) do not. (H) Pancreatic duct epithelium stains with luminal accentuation (lower right). Acini (a, center) and islets (i, upper left) do not stain. (I) Epithelial cells lining distal tubules of renal cortex stain (right); proximal tubules do not stain (left). (j) Collecting duct epithelium of renal medulla stains with luminal accentuation. (K) Superficial transitional epithelial cells lining the bladder stain. (L) Epithelium lining sweat ducts (bottom) of skin stain with luminal accentuation, whereas the sweat glands stain less intensely (arrow, top).

### SPP1 (Osteopontin)

#### PMID (review) 11703581

**Figure 1. Expression of osteopontin (OPN) in normal adult kidneys.** OPN expression in normal mouse kidneys is primarily restricted to the thick ascending limbs of the loop of Henle and distal convoluted tubules. In normal rat kidneys, OPN is primarily present in the descending thin limbs of the loop of Henle in the outer medulla. In normal human kidneys, OPN is localized primarily to the distal nephrons and is strongly expressed by the thick ascending limbs of the loop of Henle.

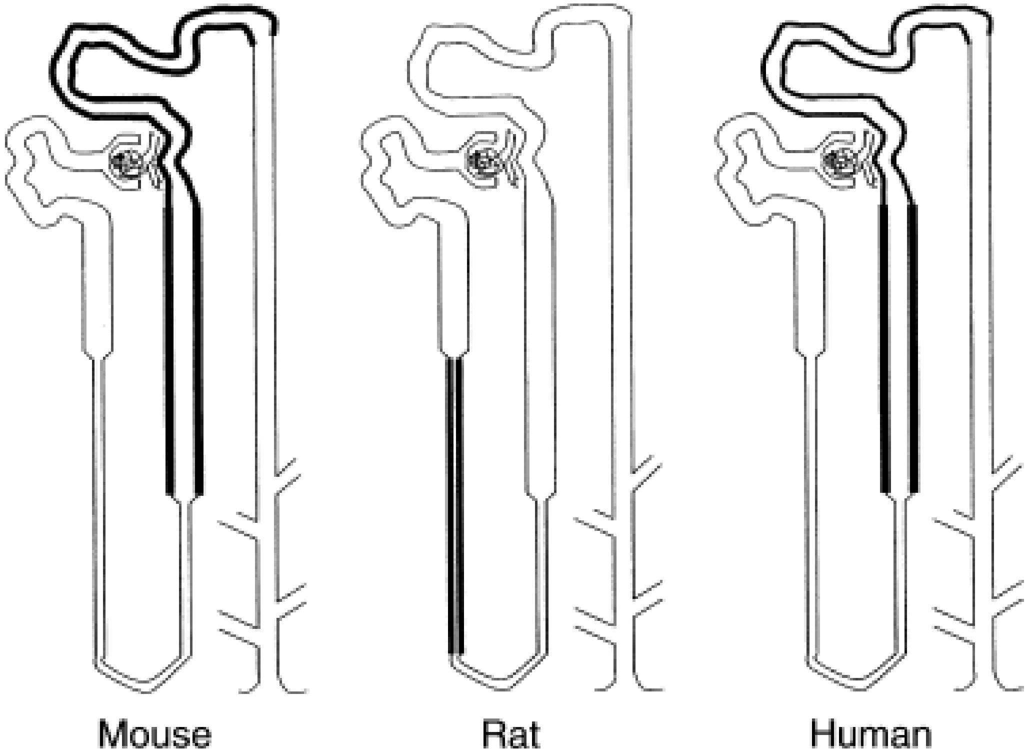

### SPP1 (Osteopontin)

**PMID 15954904**

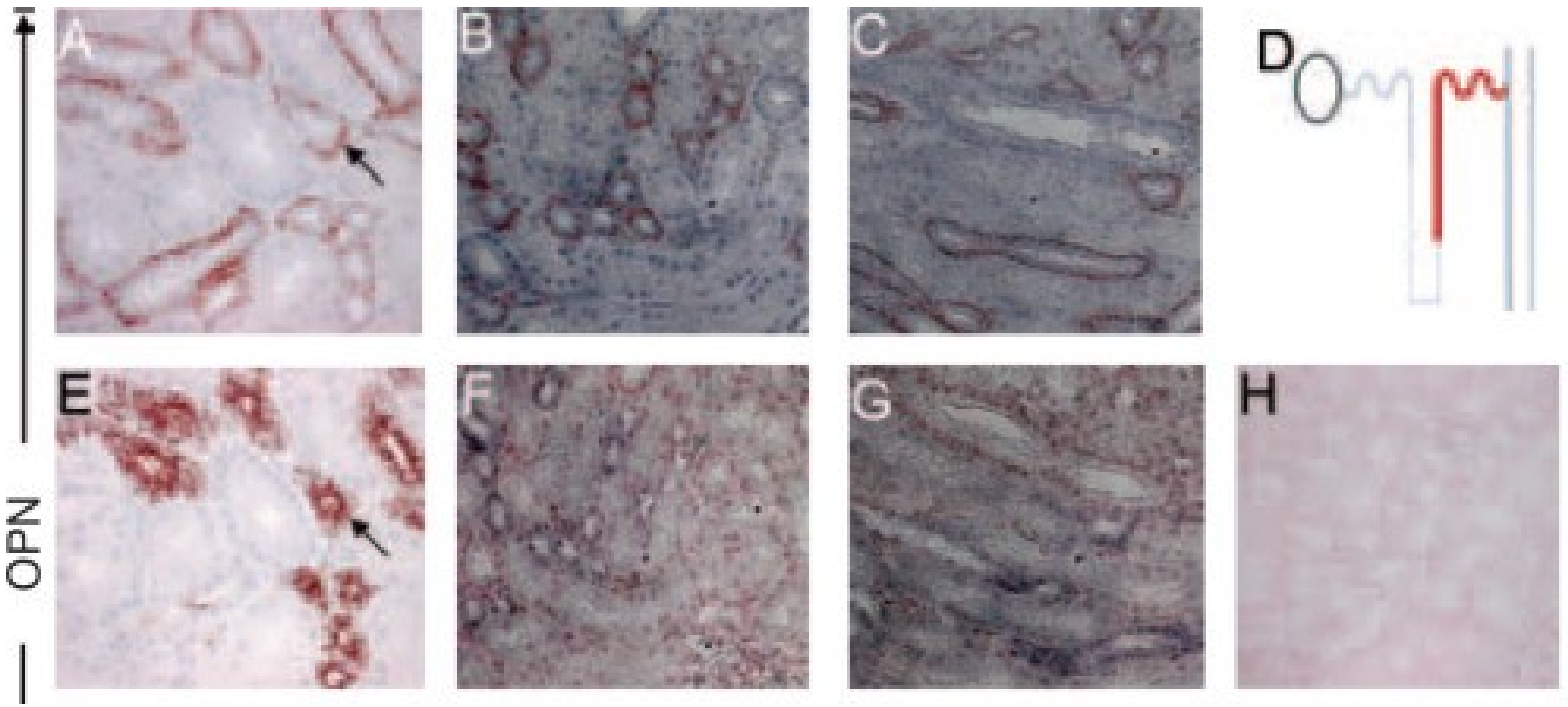


**Figure 2.** Osteopontin (OPN) and matrix metalloproteinase (MMP)-3 localization in monkey and rat kidneys. Immunolocalization (reddish/brown) of OPN (*A* to *C*) to the distal tubules of **monkey** kidney with prominent basal localization (A, arrow). Distal convoluted tubules in the cortical labyrinth (A) and distal straight tubules in the medullary rays (B and C) are positive for osteopontin monoclonal antibody LF-Mb14, and negative for all other segments, including the glomeruli; schematic (*D*) summarizes OPN segmental localization (red). Positive control (*E*, arrow) showing distal tubule localization of Tamm-Horsefall protein (THP) but without basal polarity [contrast with (A)]. Antibody results for OPN were verified by in situ hybridization (*F* and *G*) (purple/blue) on corresponding serial sections.

### SPP1 (Osteopontin)

https://www.scbt.com/p/opn-antibody-akm2a1

OPN Antibody (AKm2A1): sc-21742. Immunoperoxidase staining of formalin fixed, paraffin-embedded human kidney tissue showing cytoplasmic staining of cells in glomeruli and cytoplasmic and faint nuclear staining of cells in tubules. mouse monoclonal IgG1 κ

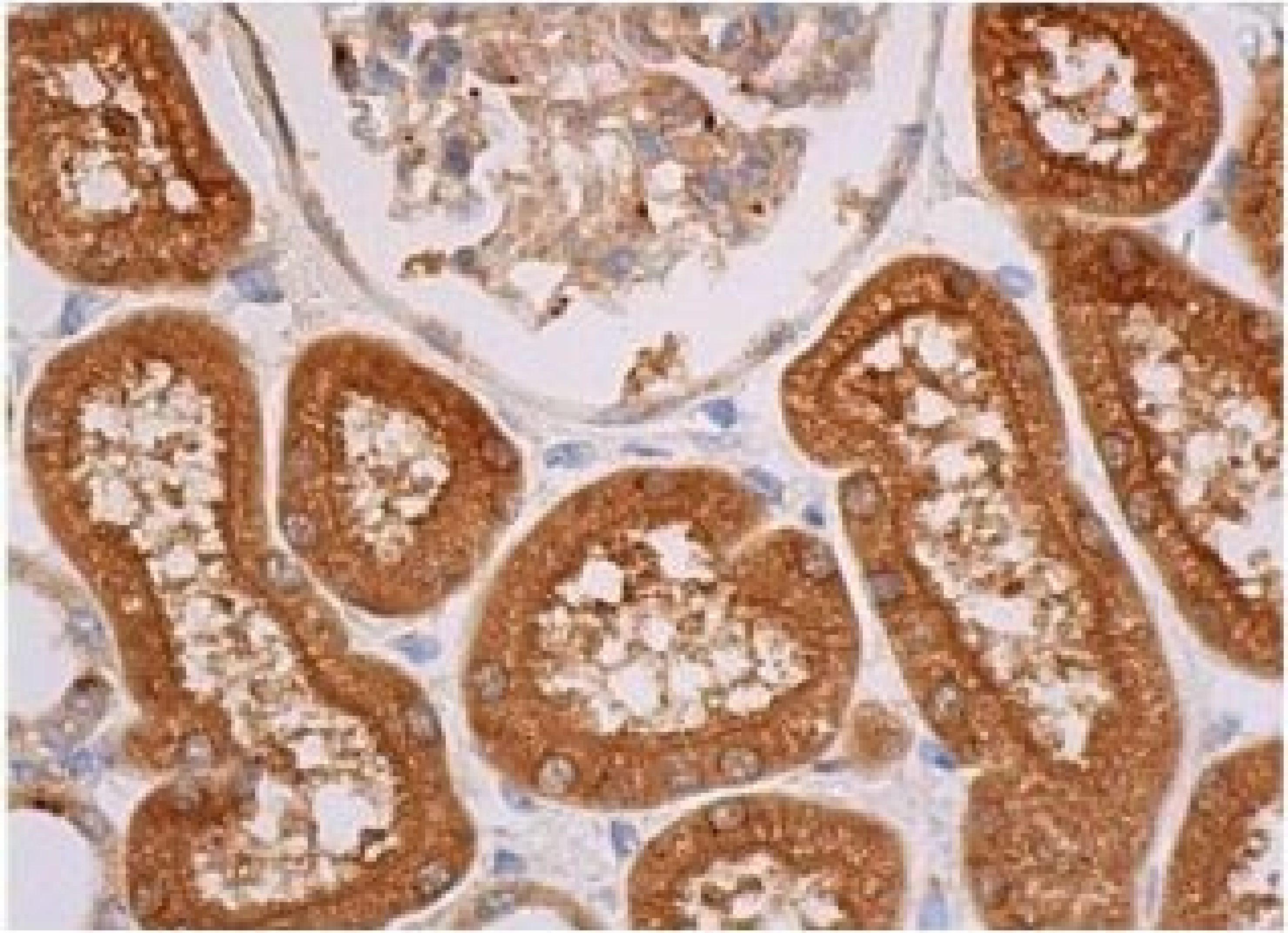

### Thrombomodulin (TM, CD141)

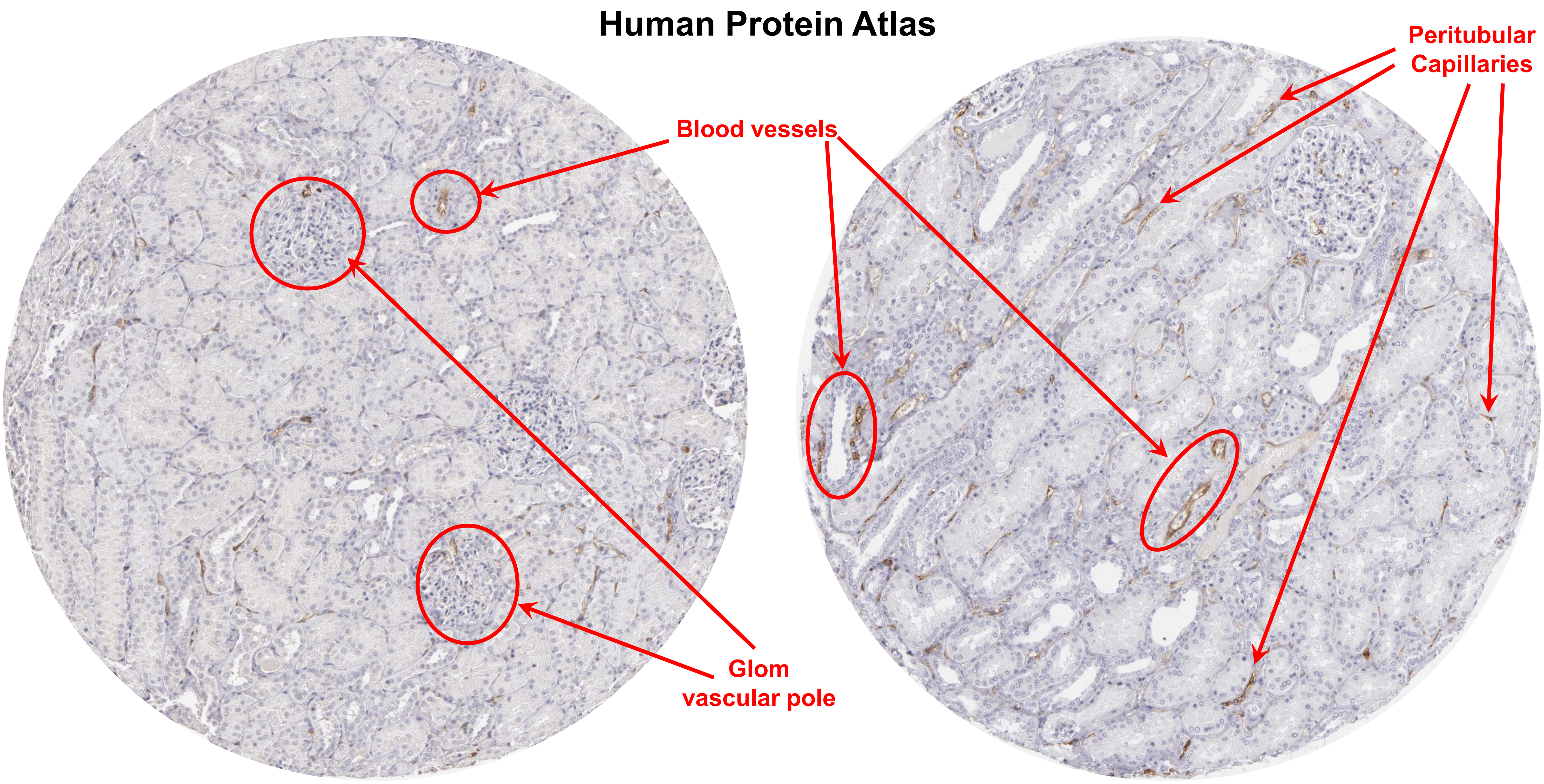

### Thrombomodulin (TM, CD141)

**PMID: 7933663**

**Fig 1.** Normal glomerulus. The intensity of TM expression is weak and its distribution is segmental. TM expression is stronger in the endothelial cells of the peritubular capillaries. (x200).

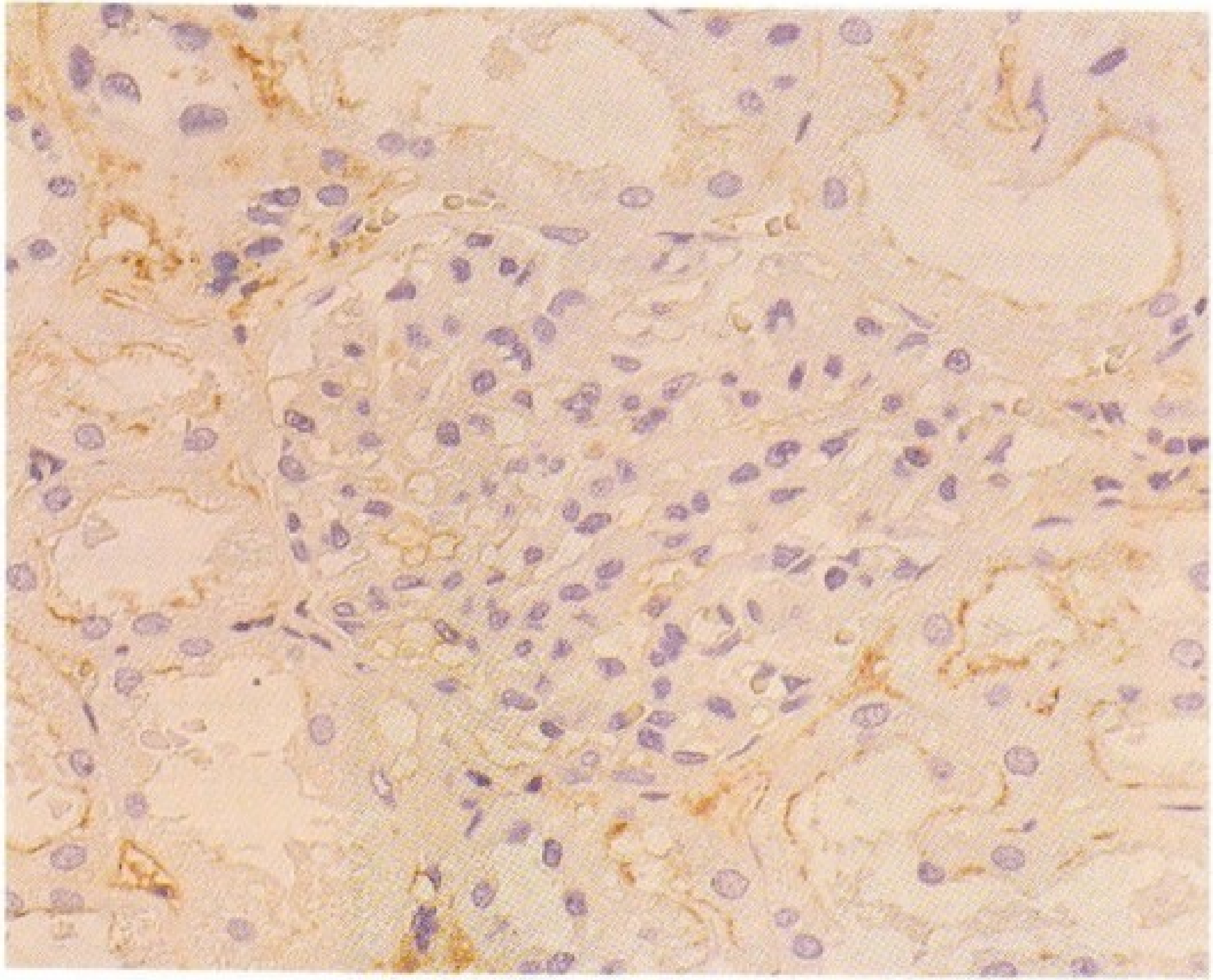

### Thrombomodulin (TM, CD141)

**PMID: 8223719**

**Fig 1.** Normal Eur J Cell Biol. 1993 Aug;61(2):299-313.

### Thrombomodulin (TM, CD141)

**PMID: 33617784**

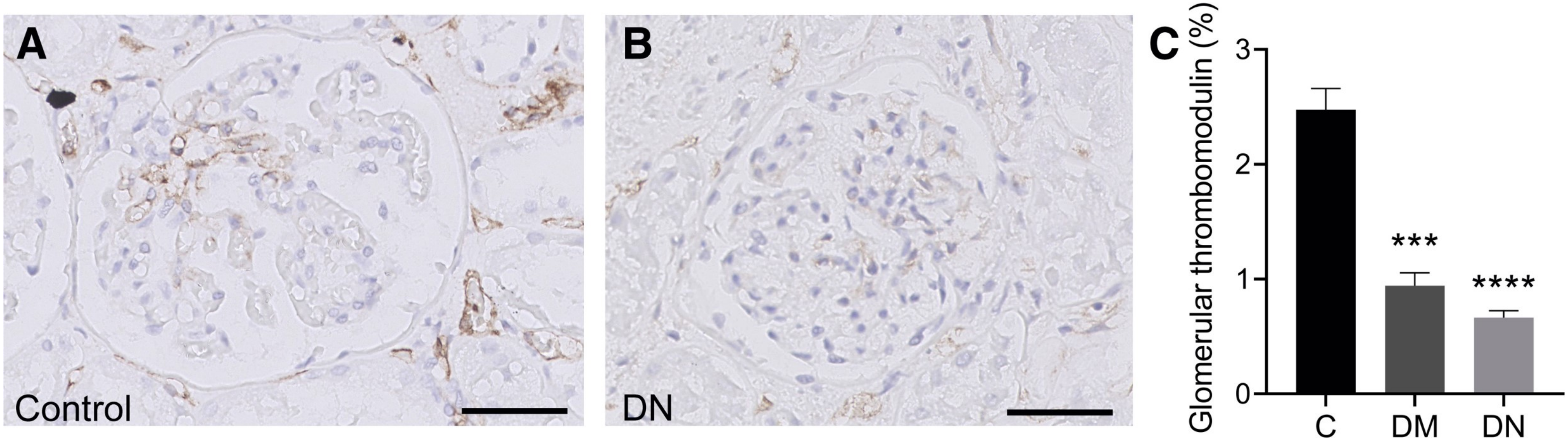


**Figure 1.** Diabetic patients have reduced glomerular thrombomodulin protein. Immunohistochemical staining using an antibody against the extracellular domain of thrombomodulin in the glomeruli of diabetic patients with diabetic nephropathy (DN), diabetic patients without DN (DM), and nondiabetic controls (C). A and B: Representative images of thrombomodulin staining in a nondiabetic control (A) and a patient with DN (B). C: Thrombomodulin-positive glomerular area in diabetic patients with DN, diabetic patients without DN, and nondiabetic controls. n = 90 diabetic patients with DN (C: DN); n = 55 diabetic patients without DN (C: DM); n = 37 nondiabetic controls (C: C). ***$P < 0.001$, ****$P < 0.0001$ versus control (linear mixed model). Scale bars = 50 μm (A and B).

### Thrombomodulin (TM, CD141)

**PMID: 33617784**

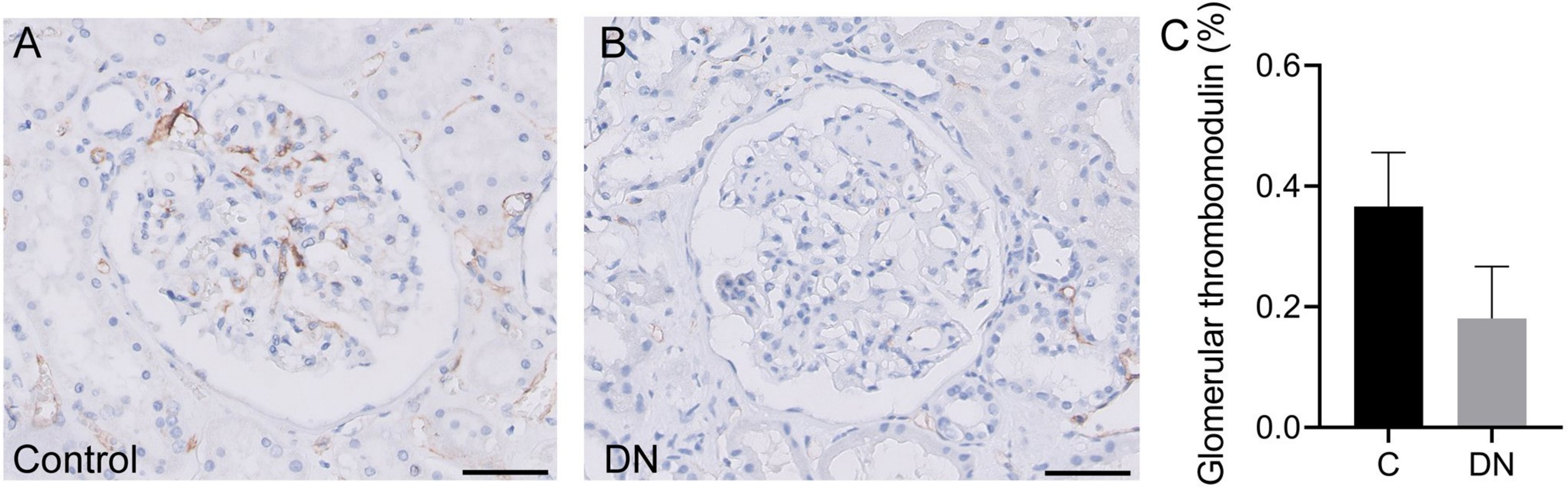


**SupFig S2.** Glomerular thrombomodulin protein levels are reduced in biopsies obtained from diabetic patients with diabetic nephropathy (DN). Biopsy samples were obtained from patients with DN; unaffected renal tissue from nondiabetic patients undergoing tumor nephrectomy was used as a control (C). A and B: Representative images of immunohistochemical staining using an antibody against the extracellular domain of thrombomodulin in sections obtained from a nondiabetic control and a patient with DN. C: Glomerular thrombomodulin measured in patients with DN and nondiabetic controls. n = 9 patients with DN; n = 8 nondiabetic patients undergoing tumor nephrectomy. Scale bars = 50 μm (A and B).

### Thrombomodulin (TM, CD141)

**PMID: 33617784**

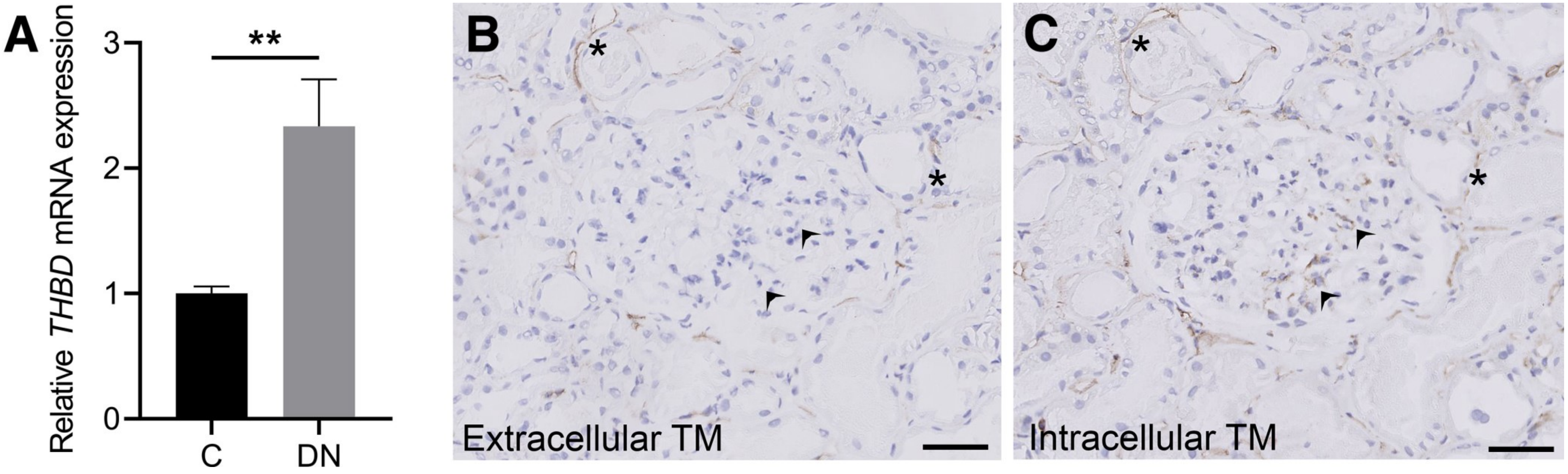


**Figure 2.** Glomerular THBD mRNA is increased in patients with diabetic nephropathy (DN). A: Summary of THBD mRNA measured in microdissected glomeruli obtained from biopsy samples taken from diabetic patients with DN and nondiabetic controls, expressed relative to control. B and C: Representative images of adjacent kidney sections obtained from a patient with DN, stained using an antibody against the extracellular (B) and intracellular (C) domains of thrombomodulin (TM). Note the increased immunoreactivity using the intracellular domain antibody in the same glomerulus (arrowheads), with similar immunoreactivity in the surrounding microvessels (asterisks). n = 24 diabetic patients with DN (A); n = 13 nondiabetic controls (A). $**P < 0.01$ versus control (t-test). Scale bars = 50 μm (B and C).

### Thrombomodulin (TM, CD141)

**PMID: 33617784**

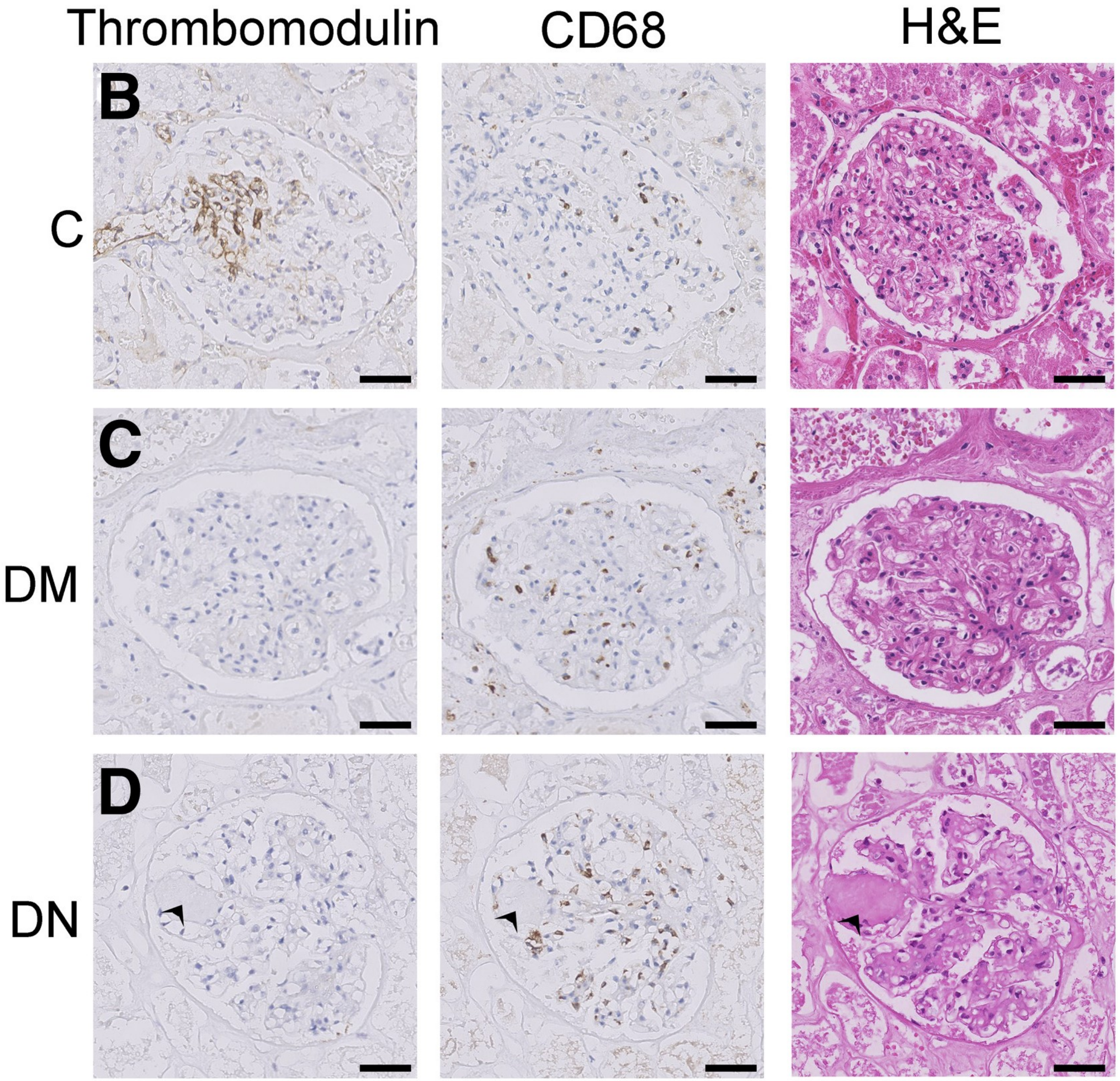


**Figure 3.** Glomerular thrombomodulin protein levels are inversely correlated with the number of glomerular CD68-positive cells. Glomerular thrombomodulin protein levels and the number of CD68-positive cells were measured in renal autopsy samples from diabetic patients with diabetic nephropathy (DN), diabetic patients without DN (DM), and nondiabetic controls (C). A: Linear mixed model regression analysis of the number of CD68-positive cells (on the y axis) on the thrombomodulin-positive area (on the x axis) in the glomeruli of the indicated groups. Each symbol represents an individual glomerulus. B–D: Representative images of three sets of adjacent kidney sections showing thrombomodulin, CD68, and hematoxylin and eosin (H&E) staining in the same glomeruli of a nondiabetic control (B), a diabetic control without DN (C), and a patient with DN (D). Kimmelstiel-Wilson lesion (arrowheads). n = 10 patients per group (A). ****$P < 0.0001$ (linear mixed model). Scale bars = 50 μm (B–D).

### Thrombomodulin (TM, CD141)

**PMID: 33707524**

**Fig 1.** Glomerular thrombomodulin protein is increased in the kidneys of women with pre-eclampsia. (a) Representative example of glomerular thrombomodulin staining in a non-hypertensive pregnant control. (b) Representative example of glomerular thrombomodulin staining in a pre-eclampsia case. (c) Averaged glomerular thrombomodulin protein scores in women with pre-eclampsia (n = 11), pregnant controls (n = 22) and non-pregnant hypertensive controls (n = 11). (d) Distribution of the glomerular thrombomodulin protein scores in the indicated groups; **p < 0.01 and *p < 0.05.

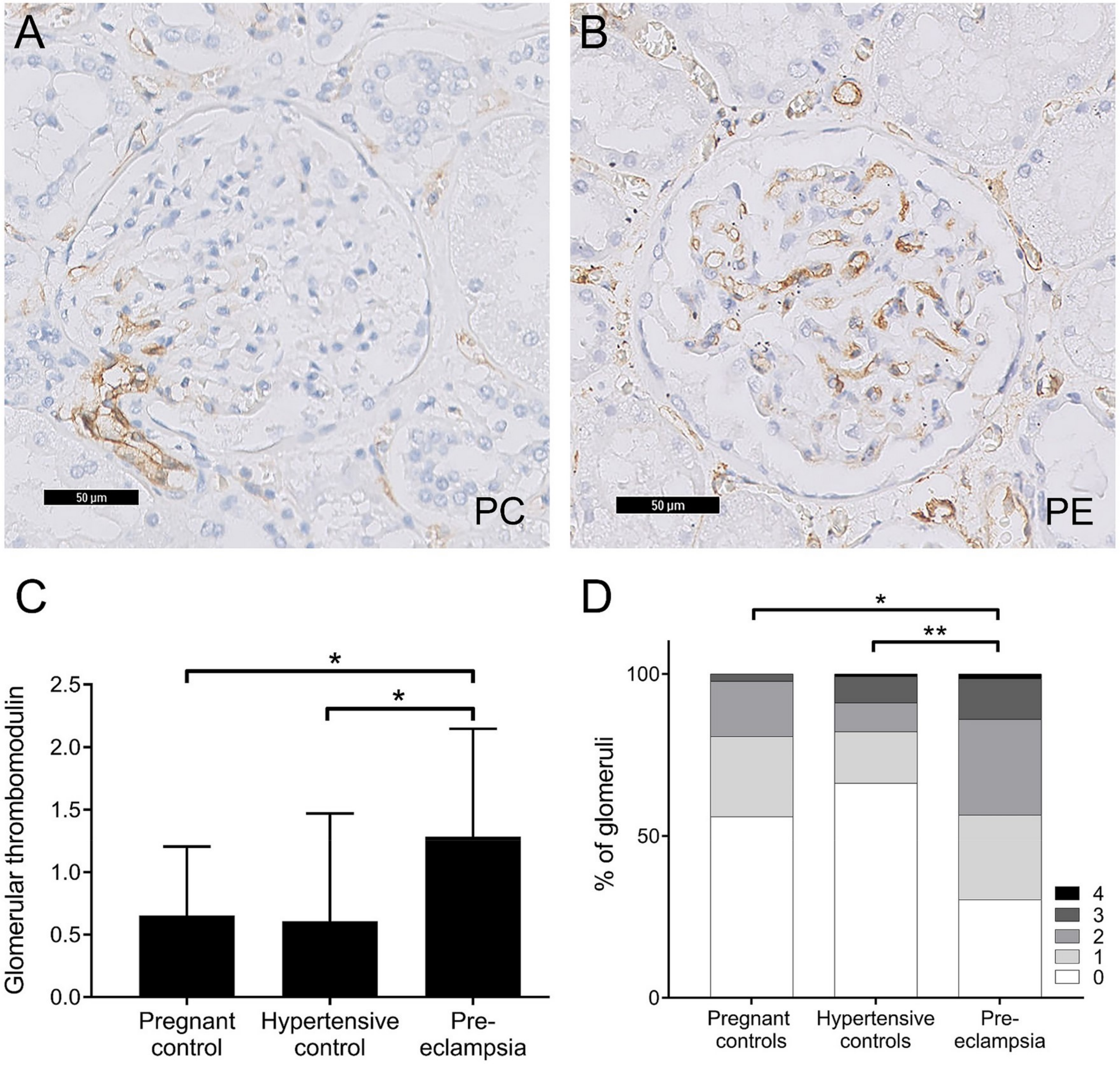

### Thrombomodulin (TM, CD141)

https://www.novusbio.com/products/thrombomodulin-bdca-3-antibody_af3894

Thrombomodulin/BDCA-3 was detected in immersion fixed paraffin-embedded sections of human kidney tissue using Goat Anti-Mouse Thrombomodulin/BDCA-3 Antigen Affinity-purified Polyclonal Antibody (Catalog # AF3894) at 1 µg/mL for 1 hour at room temperature followed by incubation with the Anti-Goat IgG VisUCyte™ HRP Polymer Antibody (Catalog # VC004). Before incubation with the primary antibody, tissue was subjected to heat-induced epitope retrieval using Antigen Retrieval Reagent-Basic (Catalog # CTS013). Tissue was stained using DAB (brown) and counterstained with hematoxylin (blue). Specific staining was localized to cell surface on endothelial cells. View our protocol for IHC Staining with VisUCyte HRP Polymer Detection Reagents.

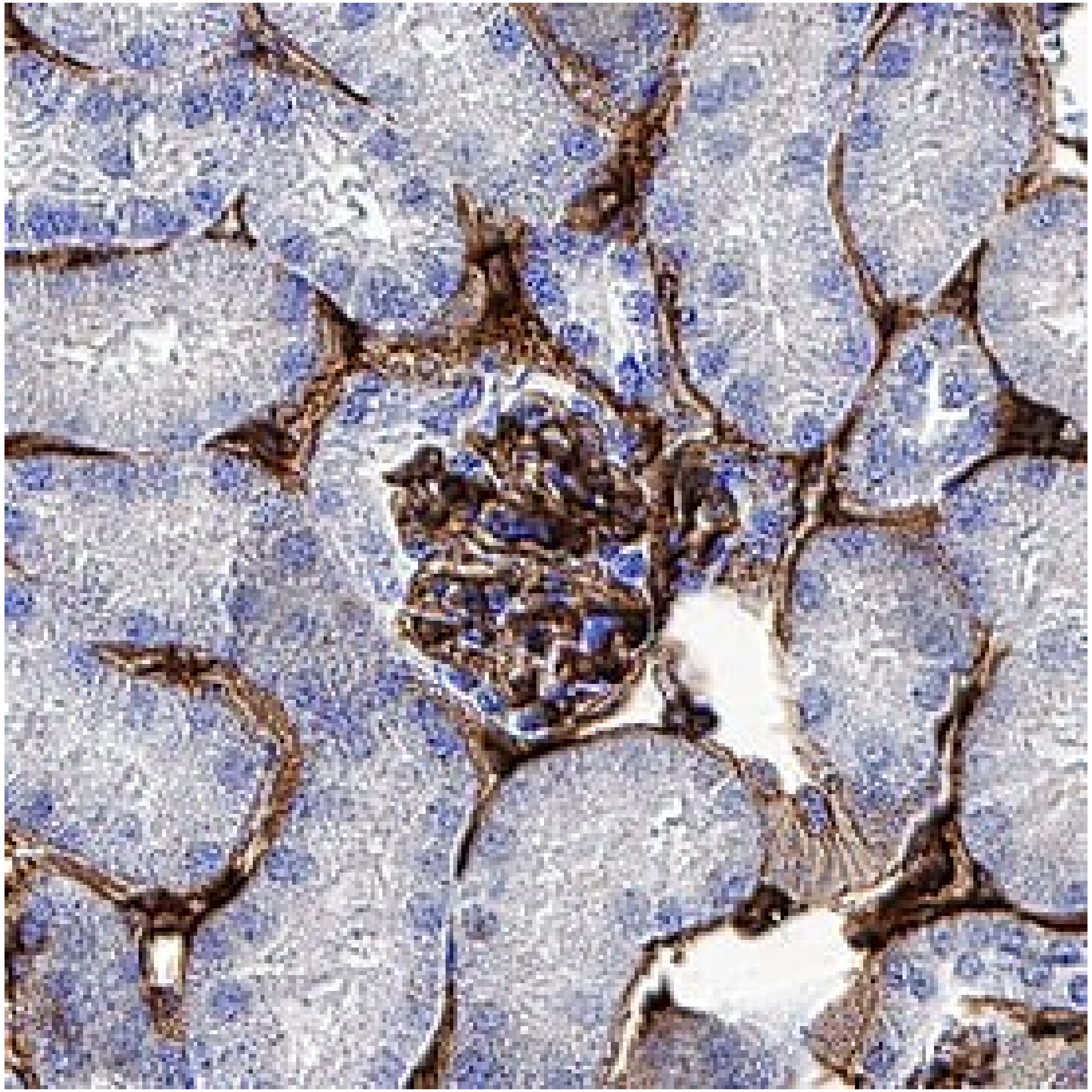

### vWF Human Protein Atlas

Five validated mAbs
No staining reported

**But staining is visible in:**

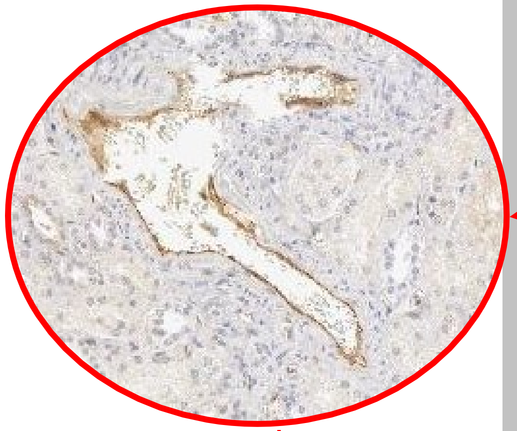

venules

arterioles

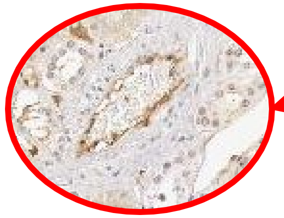

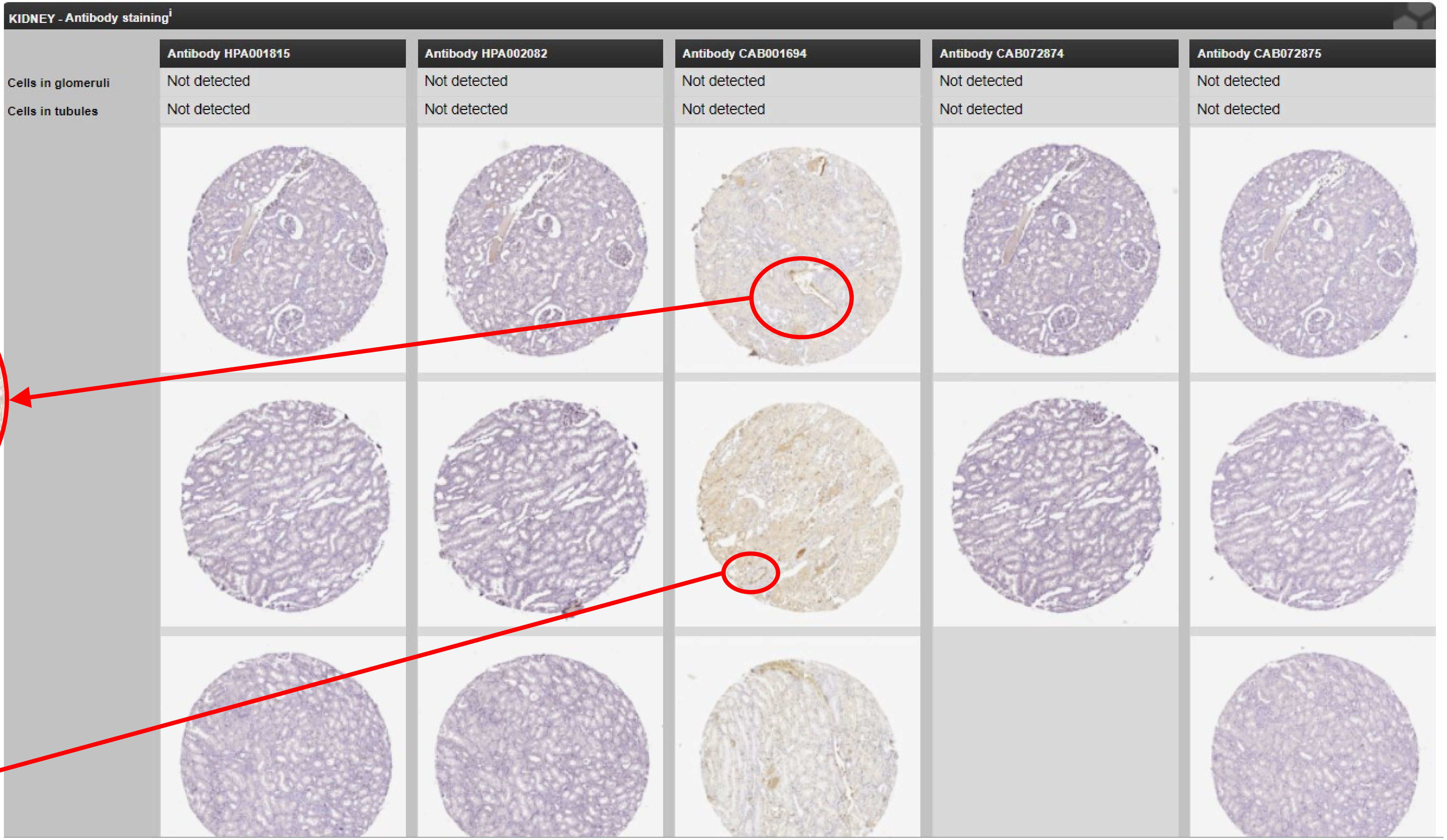

### vWF

**PMID 16234507**

**Figure 1** (**A**) CD31 expression in the kidney showing 2 glomeruli with strong positivity. (**B**) CD34 expression pattern in the kidney is similar to CD31 but more interstitial capillaries are stained. (**C**) In contrast to CD31 and CD34, the glomeruli are negative for von Willebrand factor (vWF). (**D**) Fli-1 nuclear immunostaining of EC present in the glomeruli and interstitial capillaries.

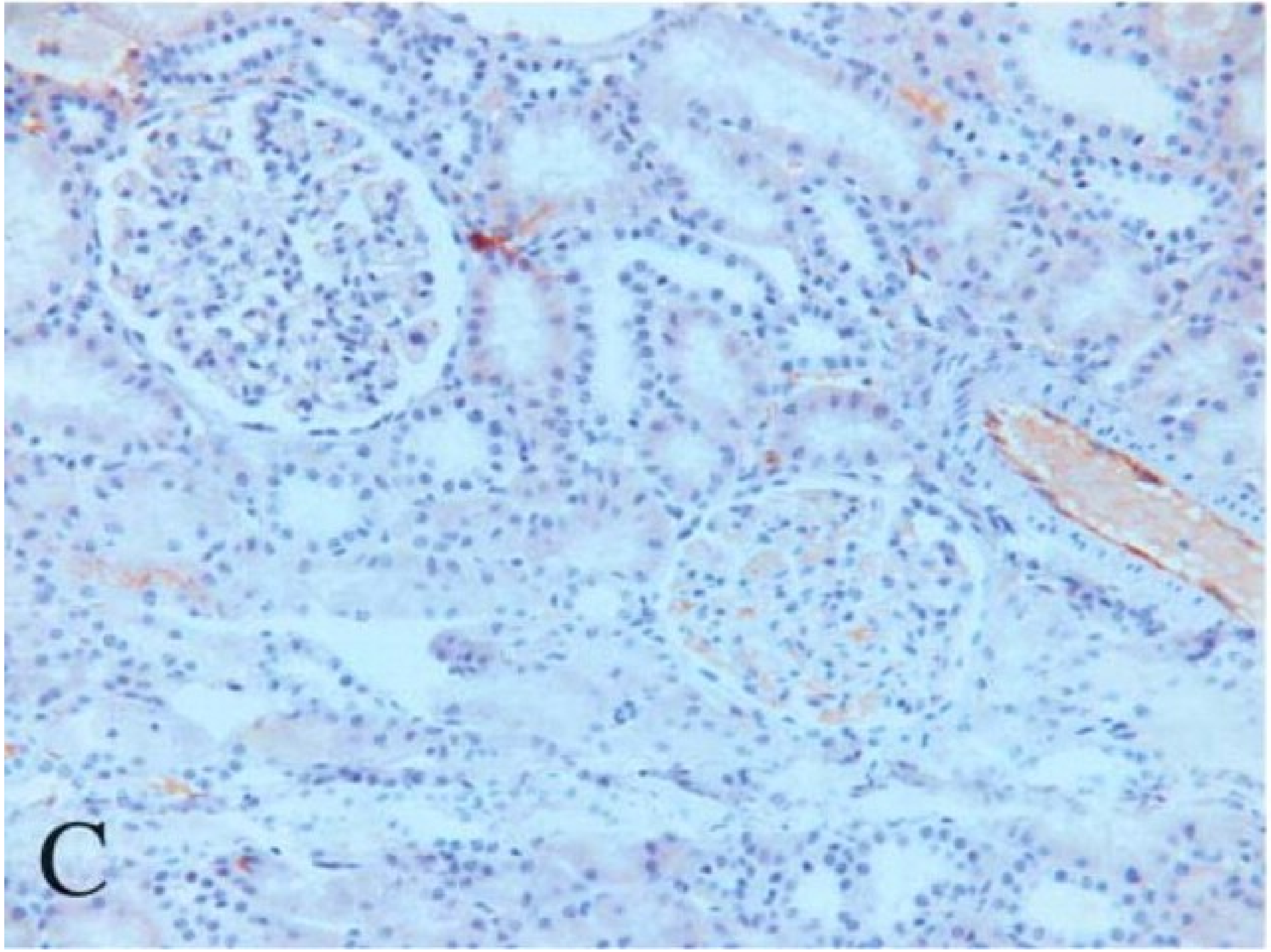


**PMID 16771247**

In control kidneys, vWF seen only in endothelium of blood vessels and glom cap loops, not in mesangium or subendothelial regions in glomeruli.

.

### vWF

**PMID 16234507**

**Table 2** Staining of endothelial cells for CD31, CD34, vWF, and Fli-1 in various normal adult tissues

| Tissue | CD31 | CD34 | VWF | Fli-1 (nuclear) |
|---|---|---|---|---|
| Kidney | | | | |
| Glomeruli | +++ | +++ | 0/+ | +++ |
| Capillaries | ++ | +++ | ++ | +++ |
| Venules | +++ | +++ | +++ | +++ |
| Arteries | +++ | +++ | +++ | +++ |
| Lung | | | | |
| Capillaries | +++ | +++ | 0/+ | +++ |
| Arterioles | +++ | +++ | ++ | +++ |
| Venules | +++ | +++ | ++ | +++ |
| Veins | +++ | +++ | +++ | +++ |
| Arteries | +++ | +++ | +++ | +++ |
| Liver | | | | |
| Periportal sinusoids | +++ | +++ | ++ | +++ |
| Centrolobular sinusoids | +++ | 0/+ | ++ | +++ |
| Centrolobular veins | +++ | +++ | ++ | +++ |
| Portal venules | +++ | +++ | ++ | +++ |
| Arterioles | +++ | +++ | ++ | +++ |
| Spleen | | | | |
| Sinusoids | ++ | 0 | ++ | ++[a] |
| Capillaries | 0 | +++ | 0/+ | ++[a] |
| Central arteries | +++ | +++ | ++ | ++[a] |
| Venules | +++ | +++ | ++ | ++[a] |
| Lymph nodes | | | | |
| Capillaries | +++ | +++ | ++ | ++[a] |
| Marginal sinuses | +++ | 0/+ | 0/+ | ++[a] |
| High endothelial venules | +++ | +++ | +++ | ++[a] |

Intensity of immunohistochemical staining: 0, absent; +, low; ++, medium; +++, high.
[a]Most cells present (lymphocytes) are stained.

### vWF

#### PMID-25163811

**Immunofluorescence images of sections of human kidney showing glomerular expression of NRGN.** Cryosections of human kidney were stained for neurogranin (NRGN) with anti-NRGN antibodies **(C)**, von-Willebrand factor (vWF) with anti-vWF antibodies **(D)** and second antibodies alone **(A** and **B)**. Signals of NRGN (red) and vWF (green) were merged in **(E**

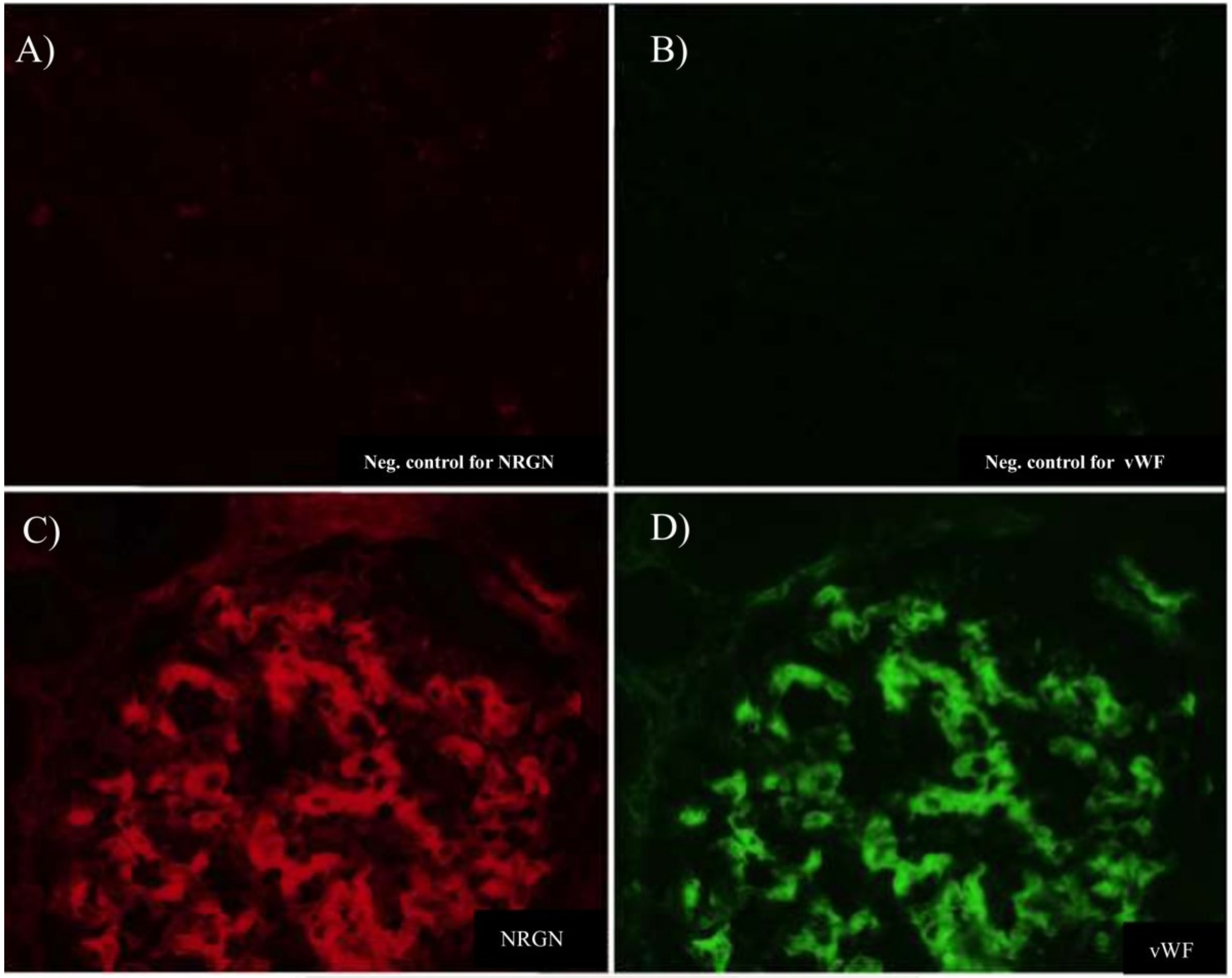

### vWF

biorbyt
https://www.biorbyt.com/vwf-antibody-orb305844.html

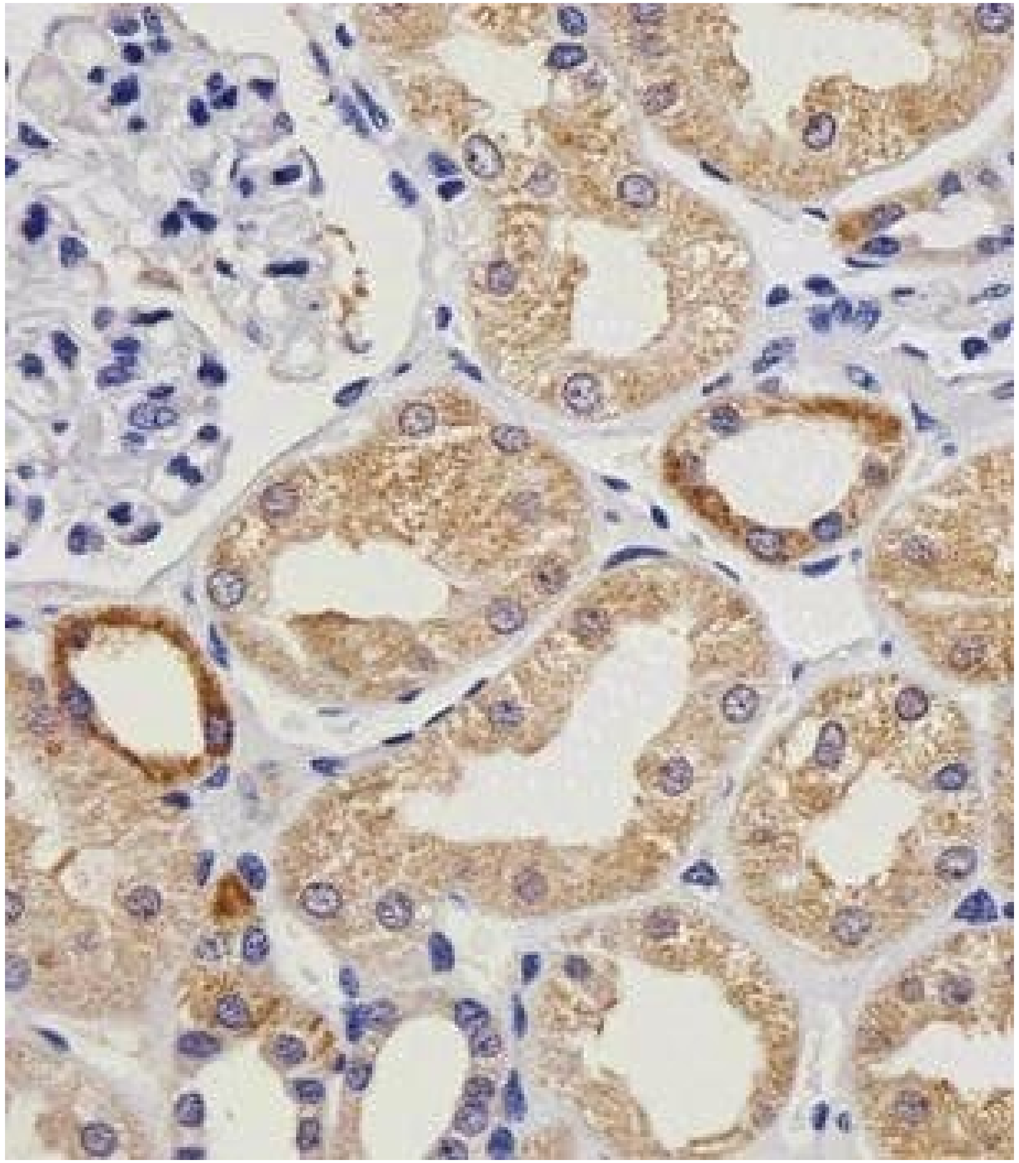

Immunohistochemical staining of paraffin embedded human kidney section tissue using VWF antibody (primary antibody dilution at: 1:25)
