## Supplementary Figure 4 for "Spatial proteomics of human diabetic kidney disease, from health to class III"

### MAP

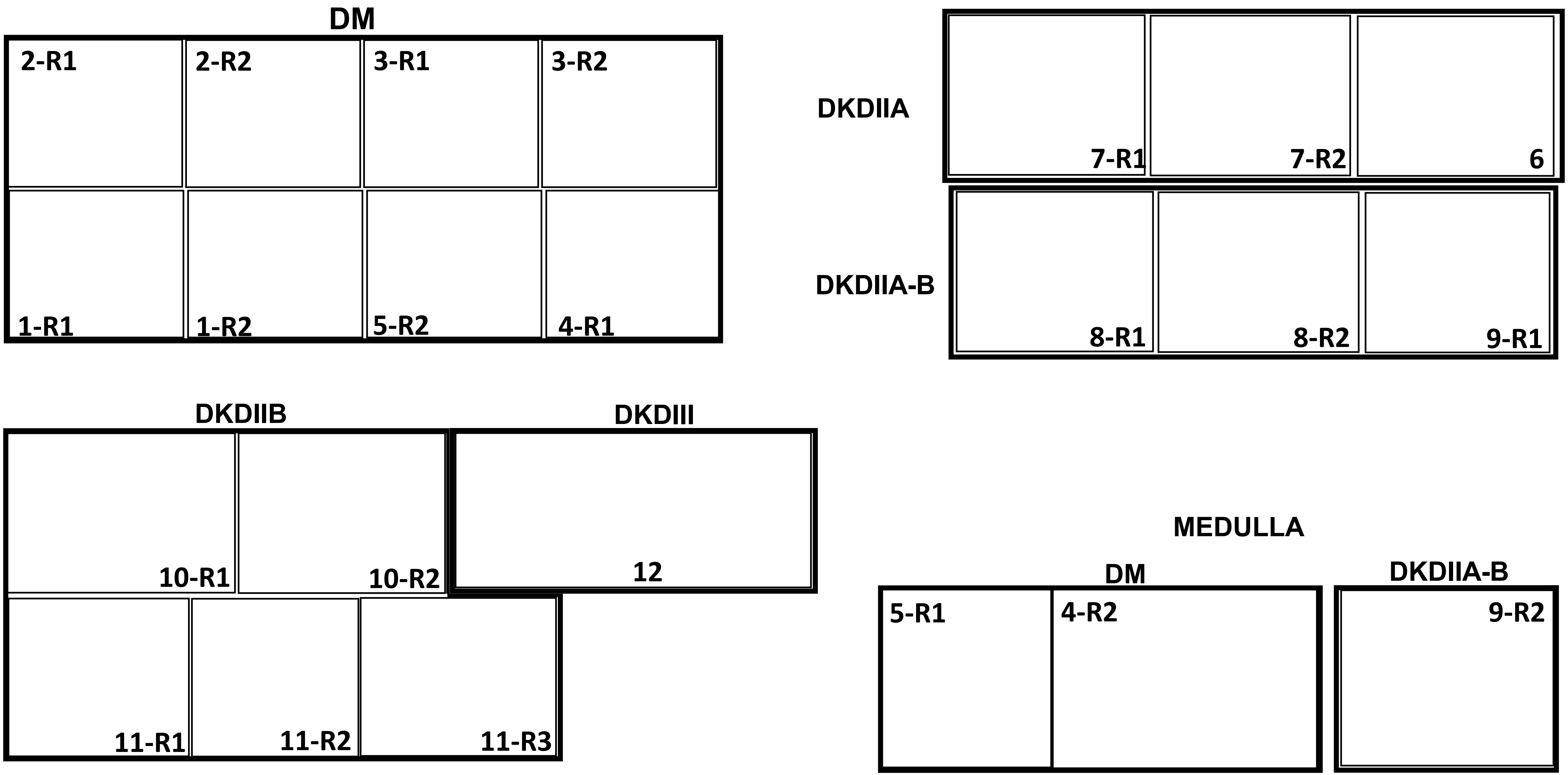
DM
2-R1
2-R2
3-R1
3-R2
1-R1
1-R2
5-R2
4-R1
DKDIIA
7-R1
7-R2
6
DKDIIA-B
8-R1
8-R2
9-R1
DKDIIB
10-R1
10-R2
11-R1
11-R2
11-R3
DKDIII
12
MEDULLA
DM
5-R1
4-R2
DKDIIA-B
9-R2

αSMA (80-500)

DM

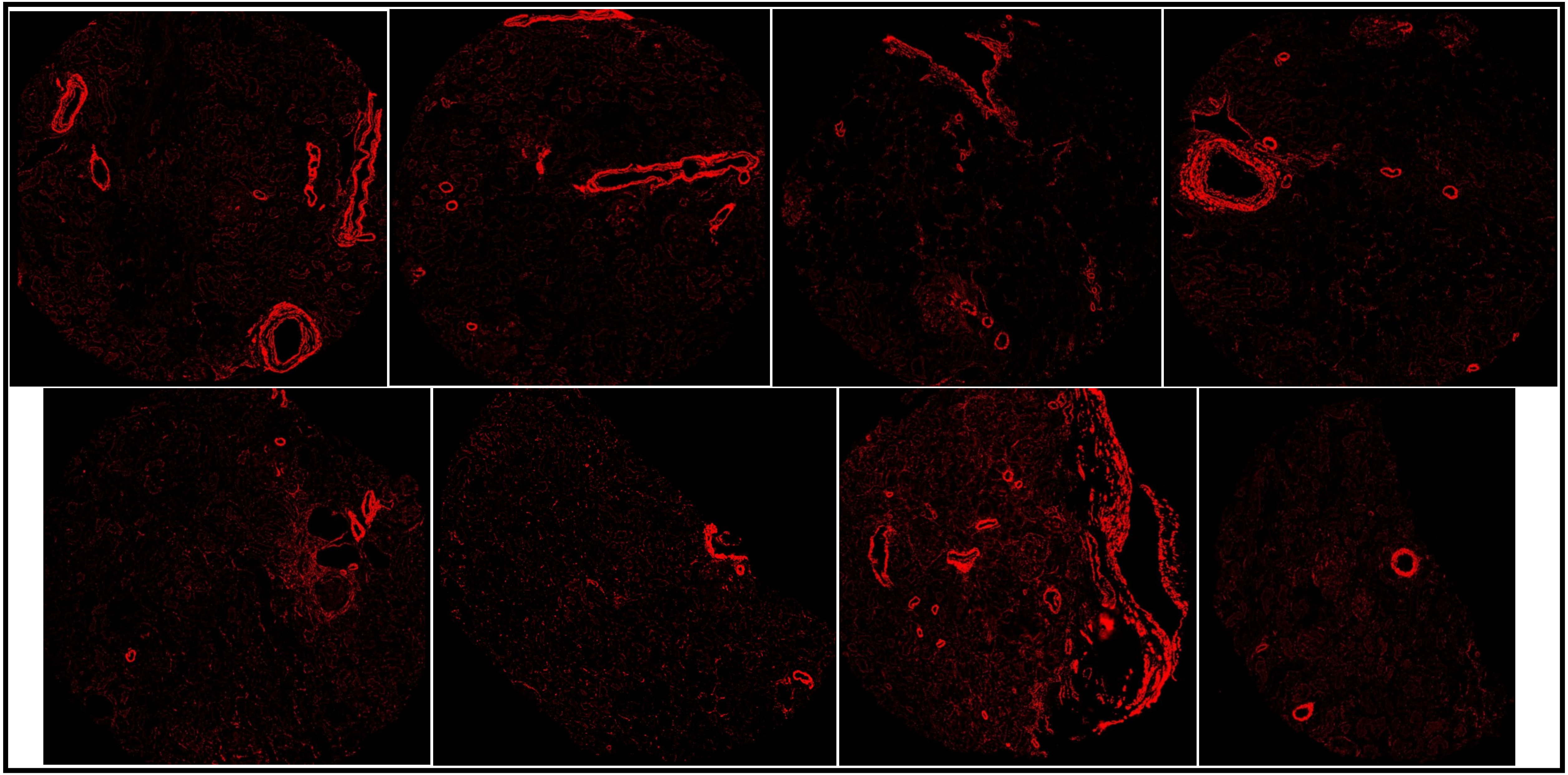

### αSMA (80-500)

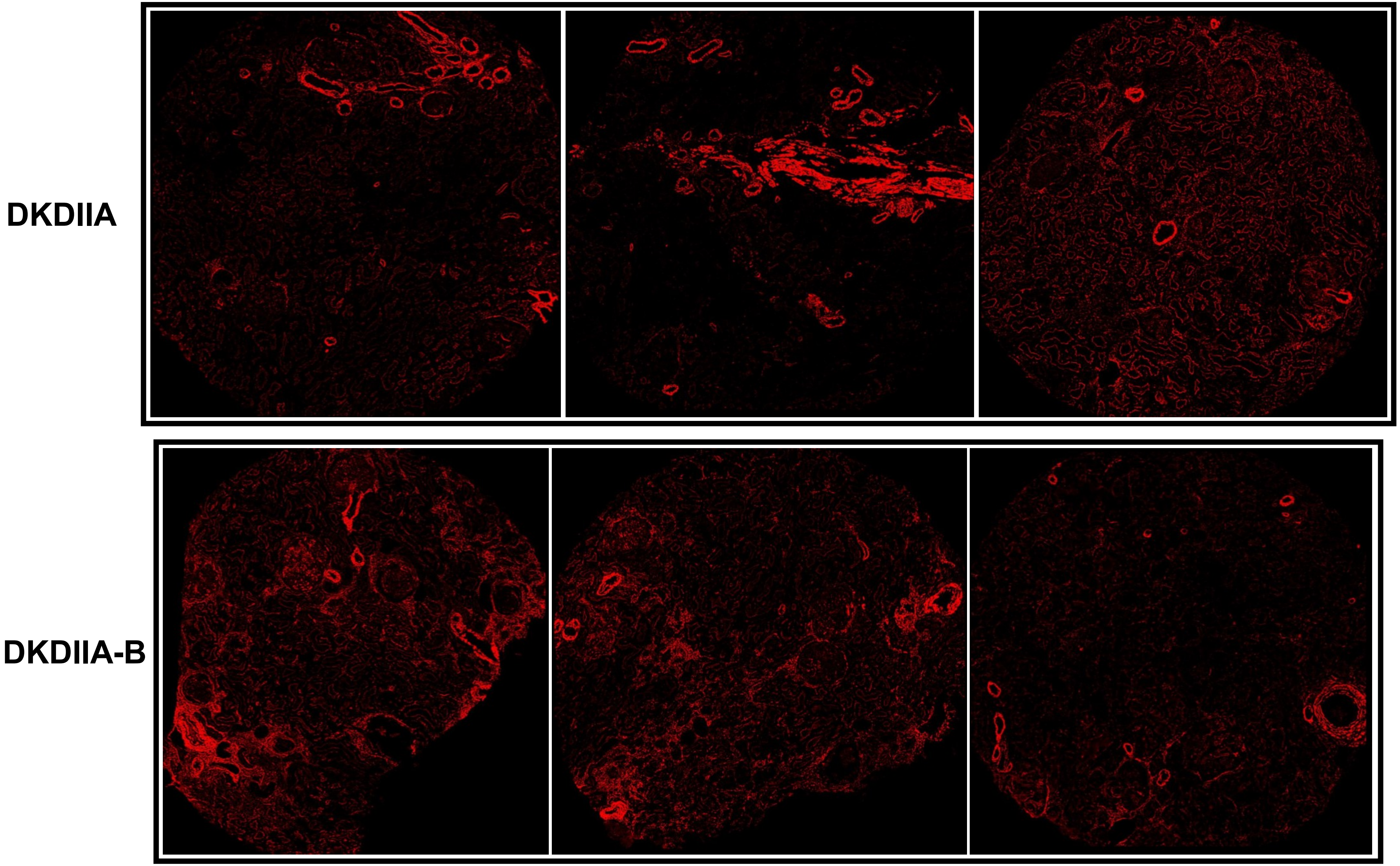

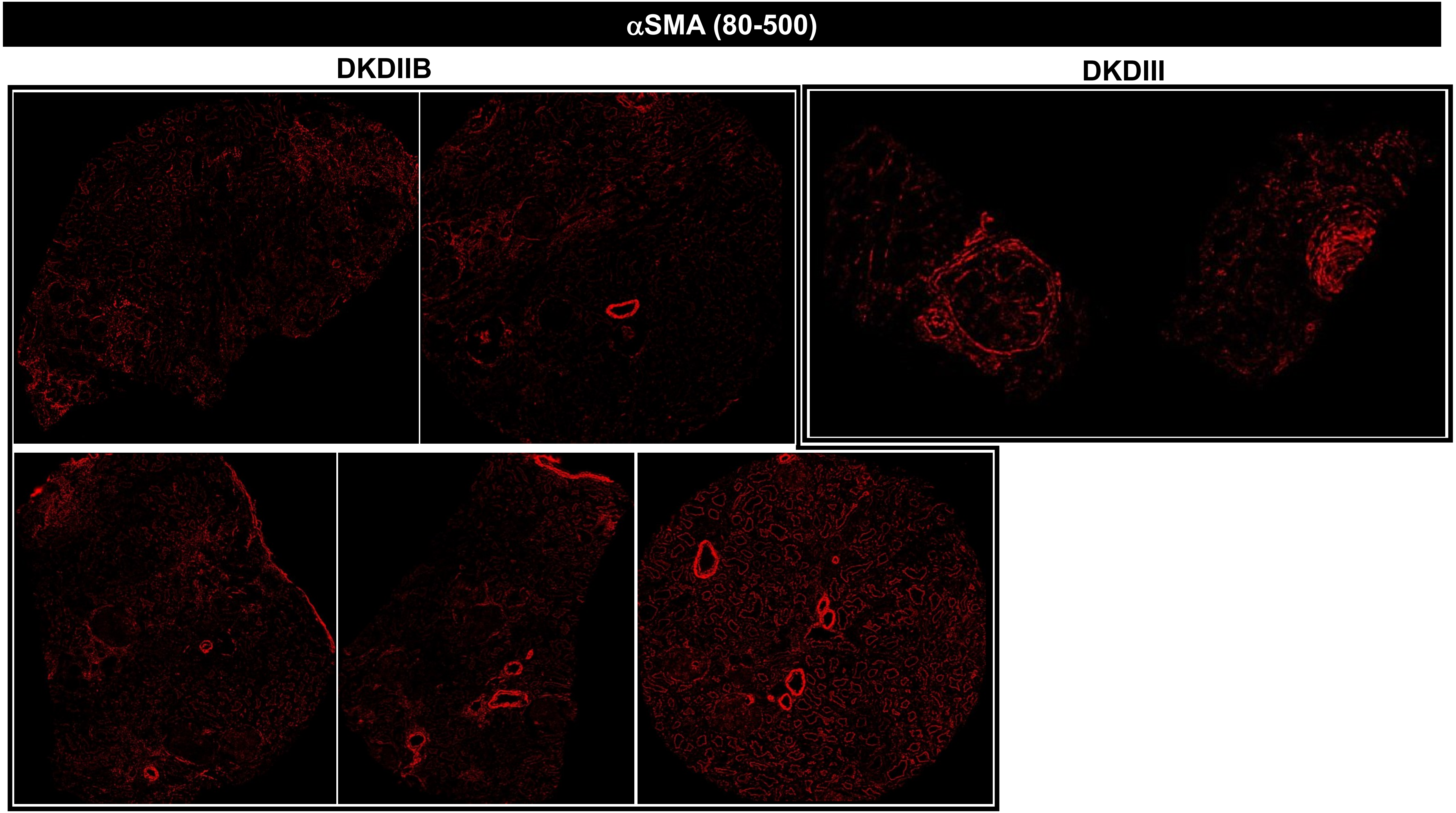
αSMA (80-500)
DKDIIB
DKDIII

#### αSMA (80-500)

### MEDULLA

**DM**

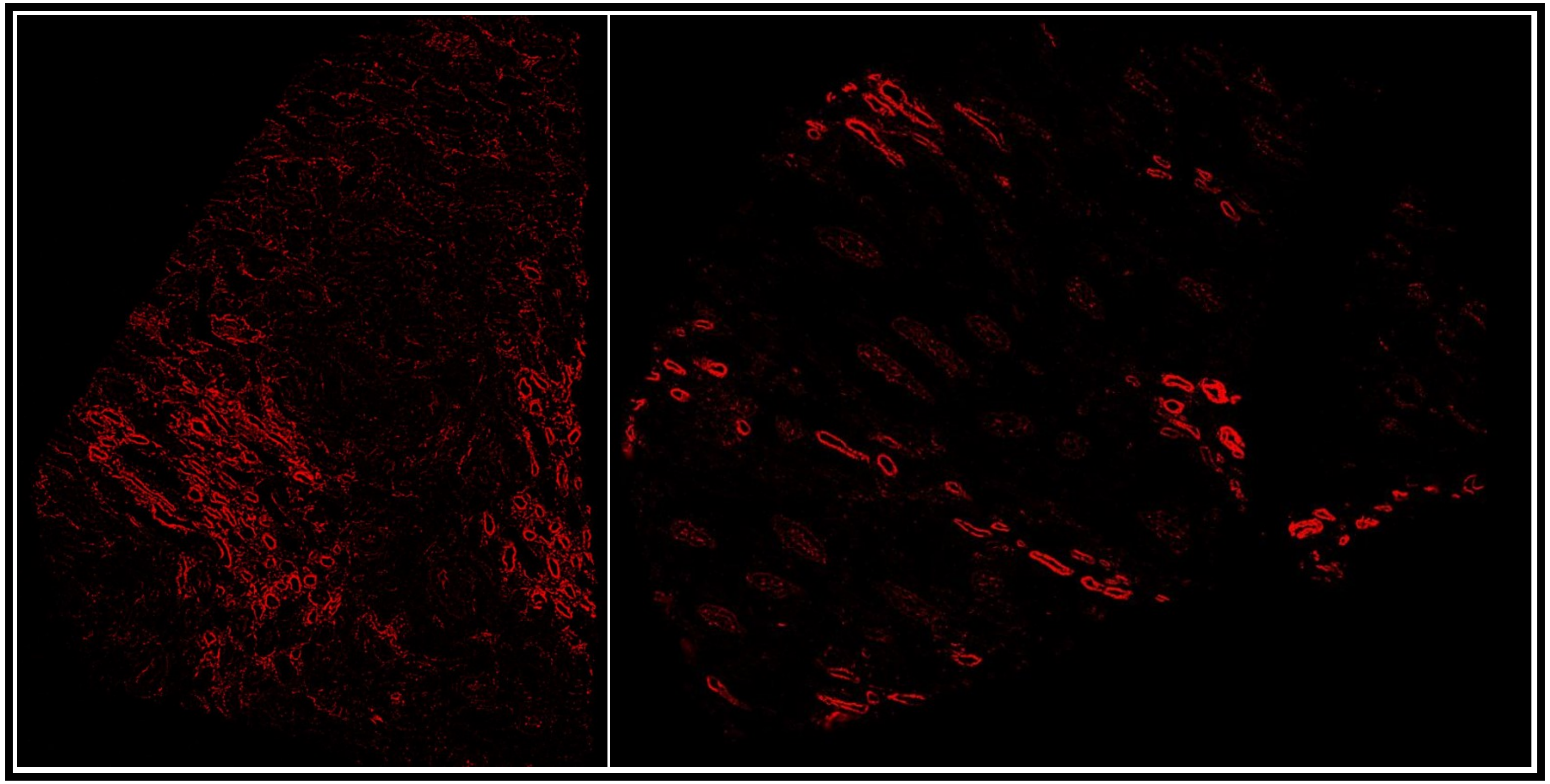

**DKDIIA-B**

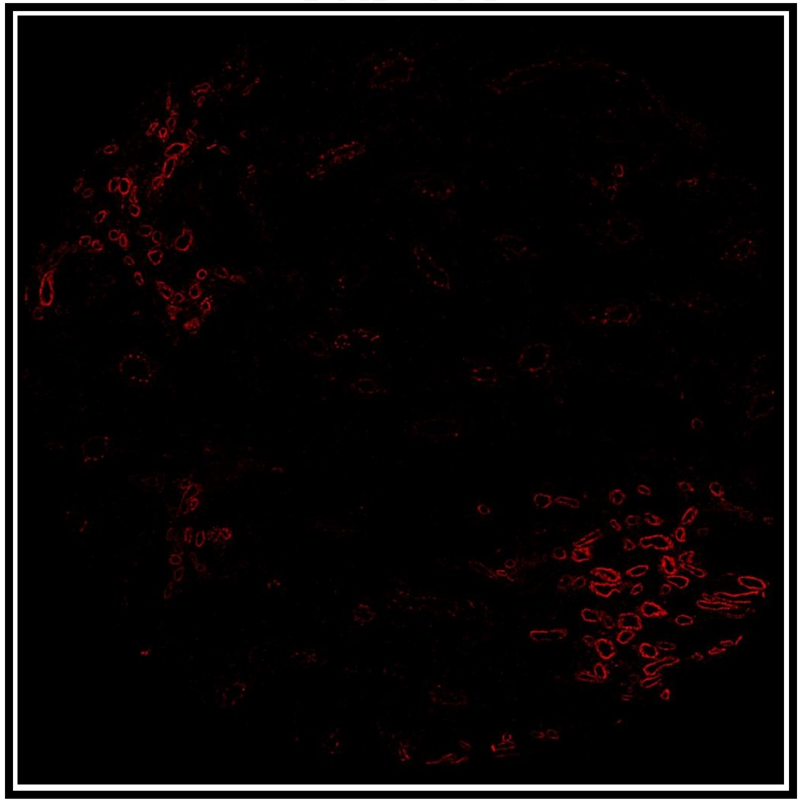

# C1QC (20-1000)

## DM

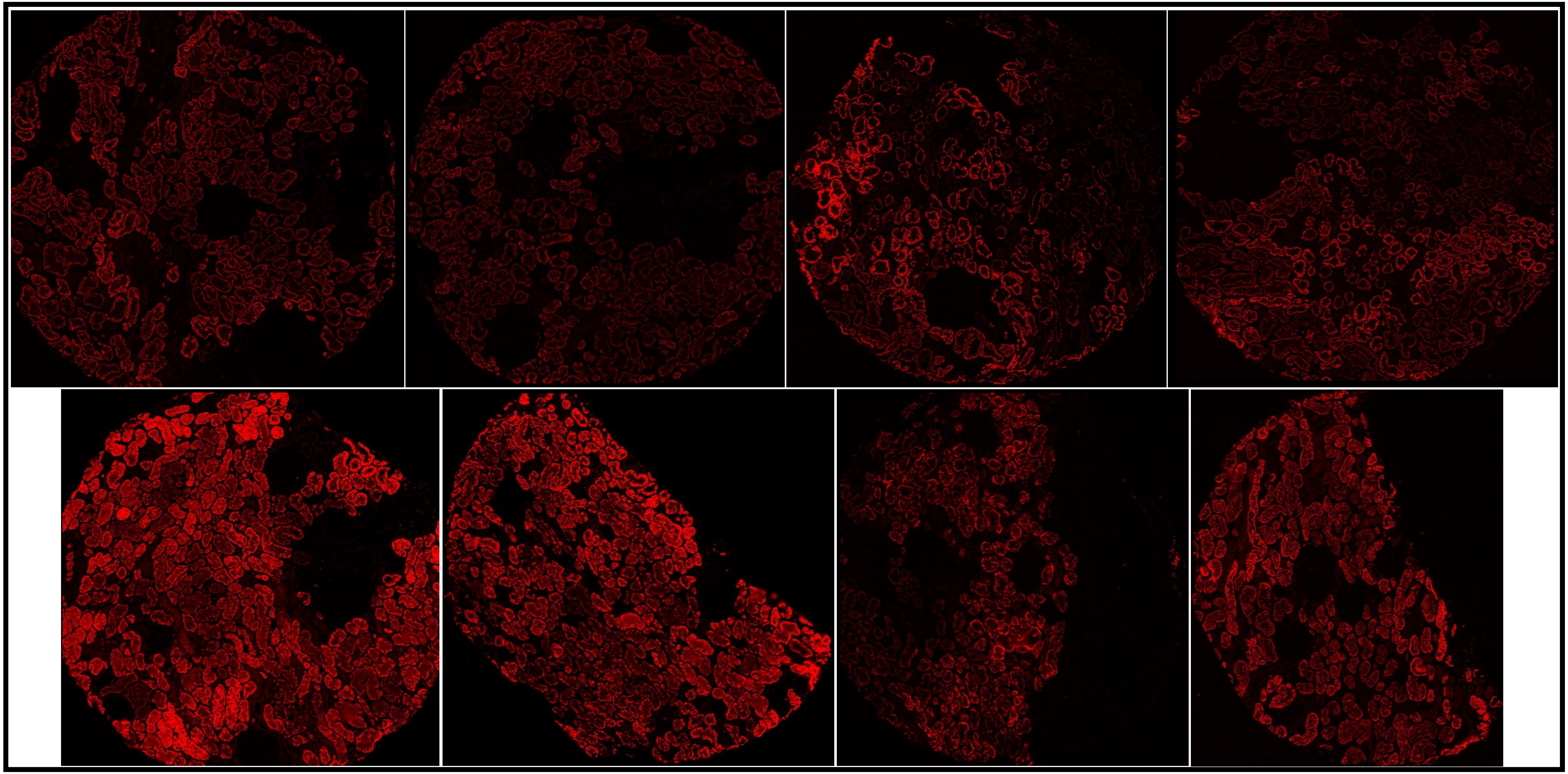

# C1QC (20-1000)

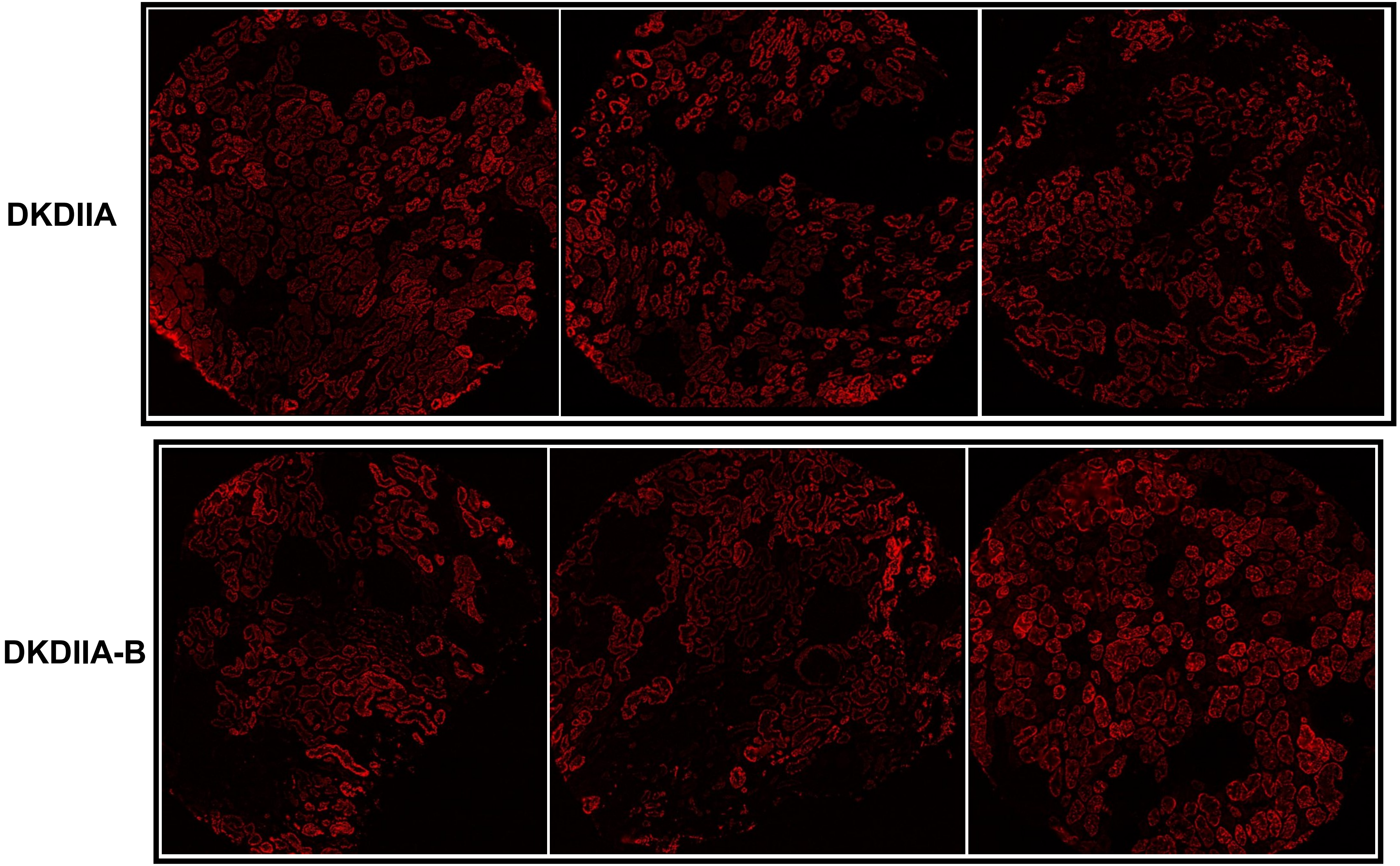

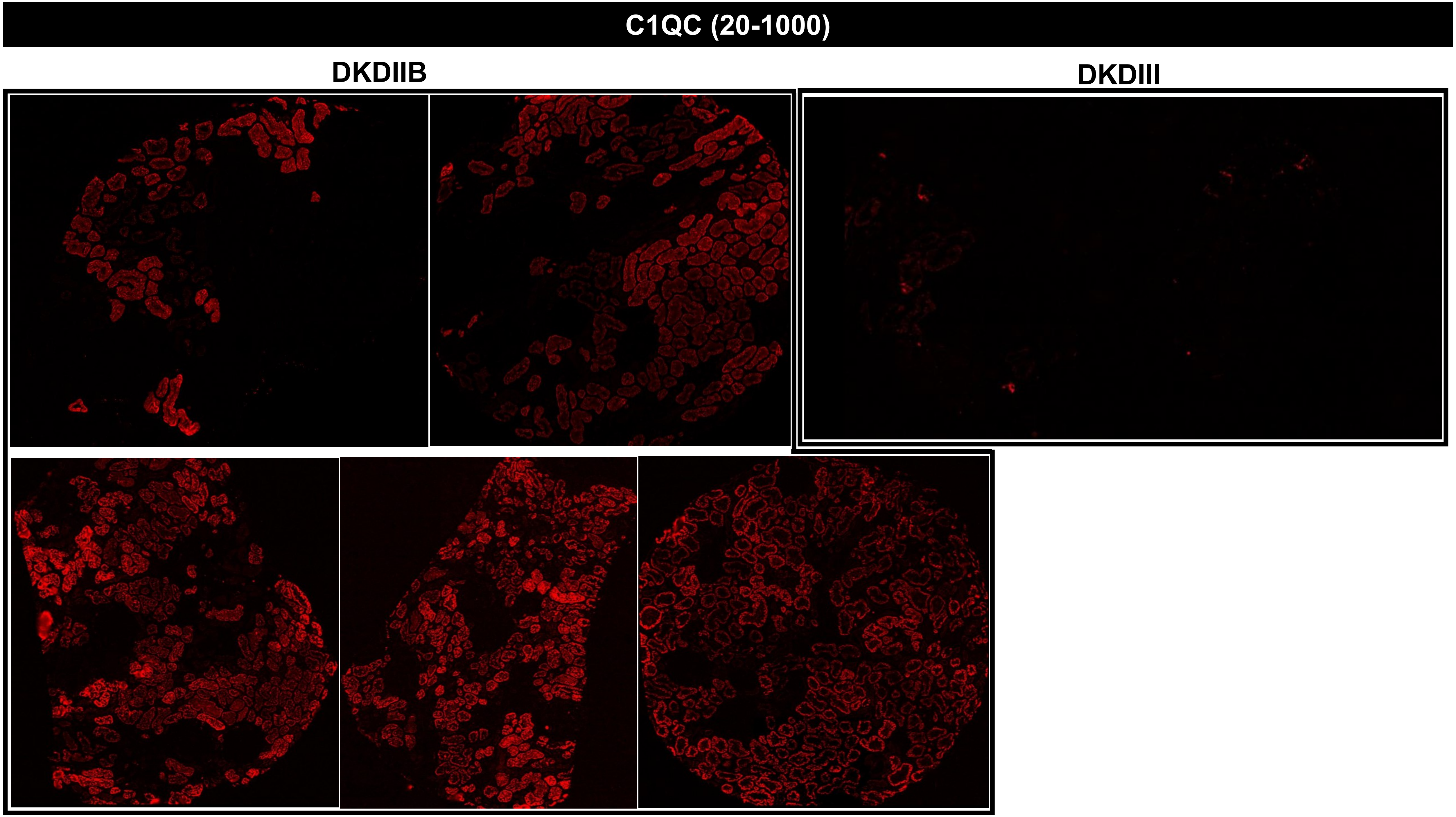
C1QC (20-1000)
DKDIIB
DKDIII

# C1QC (20-1000)

#### MEDULLA

DM

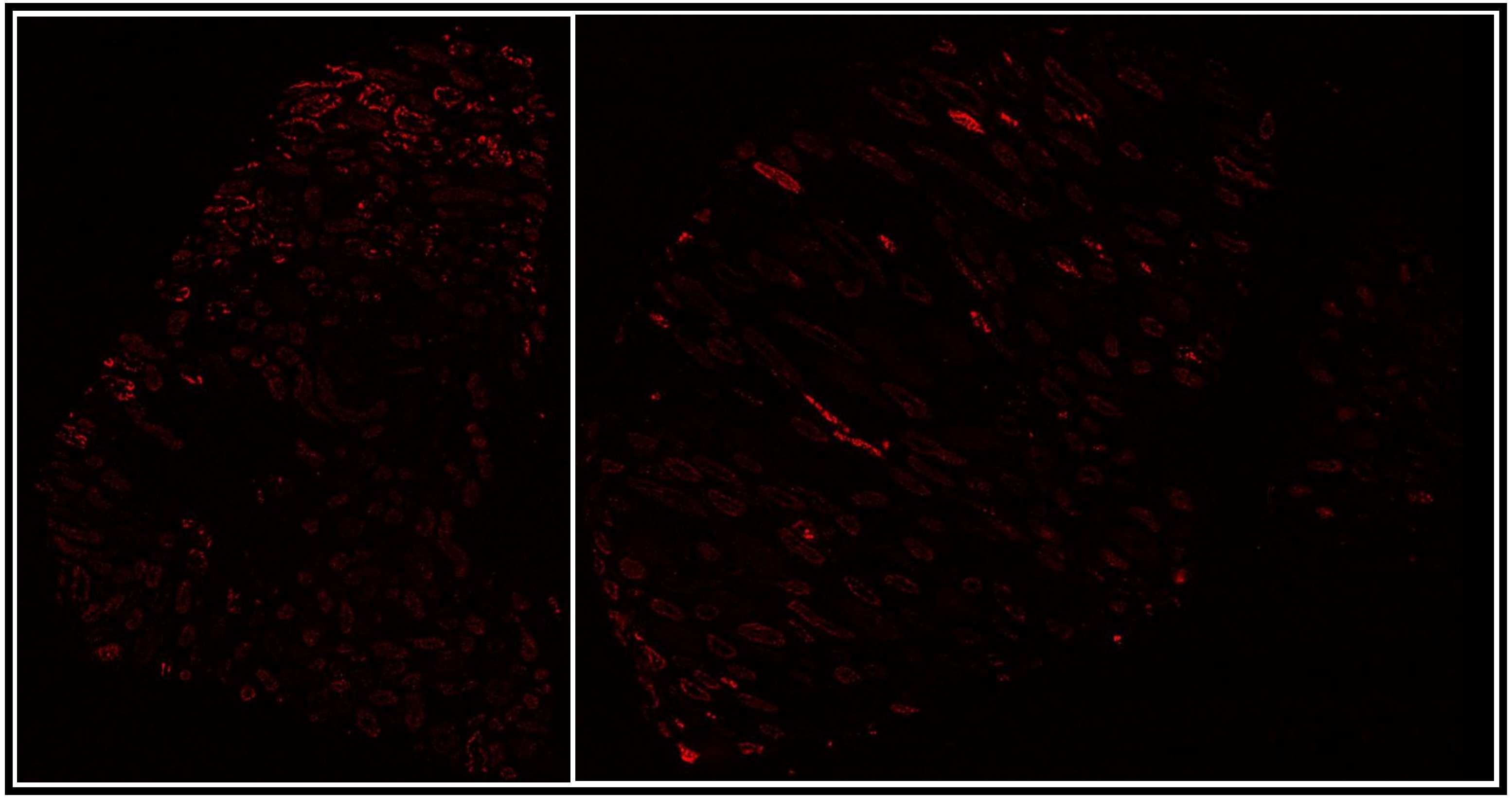

DKDIIA-B

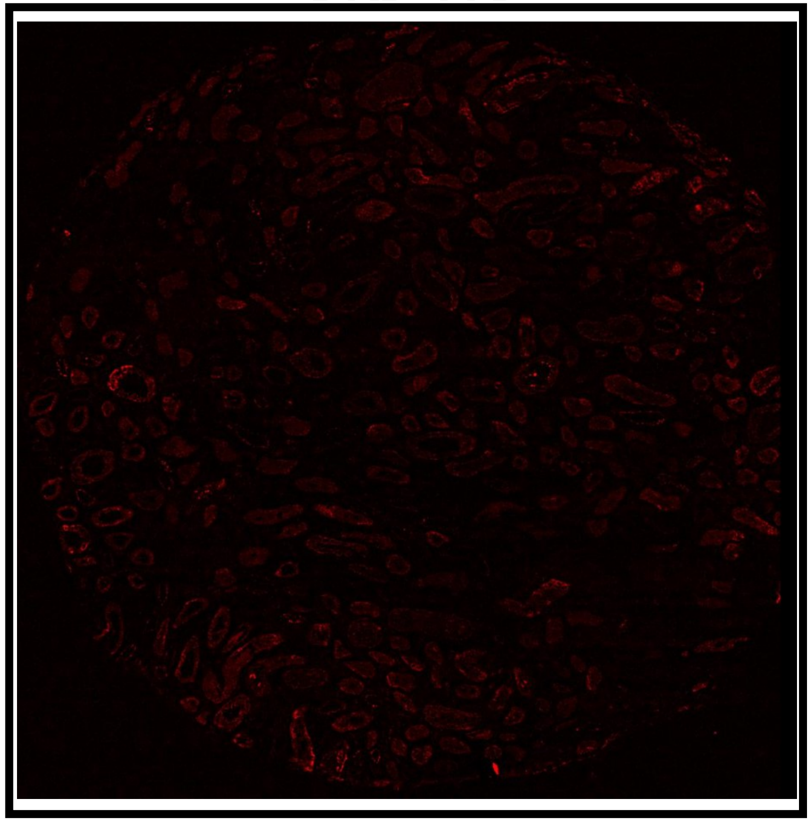

**CCR6 (50-200)**

**DM**

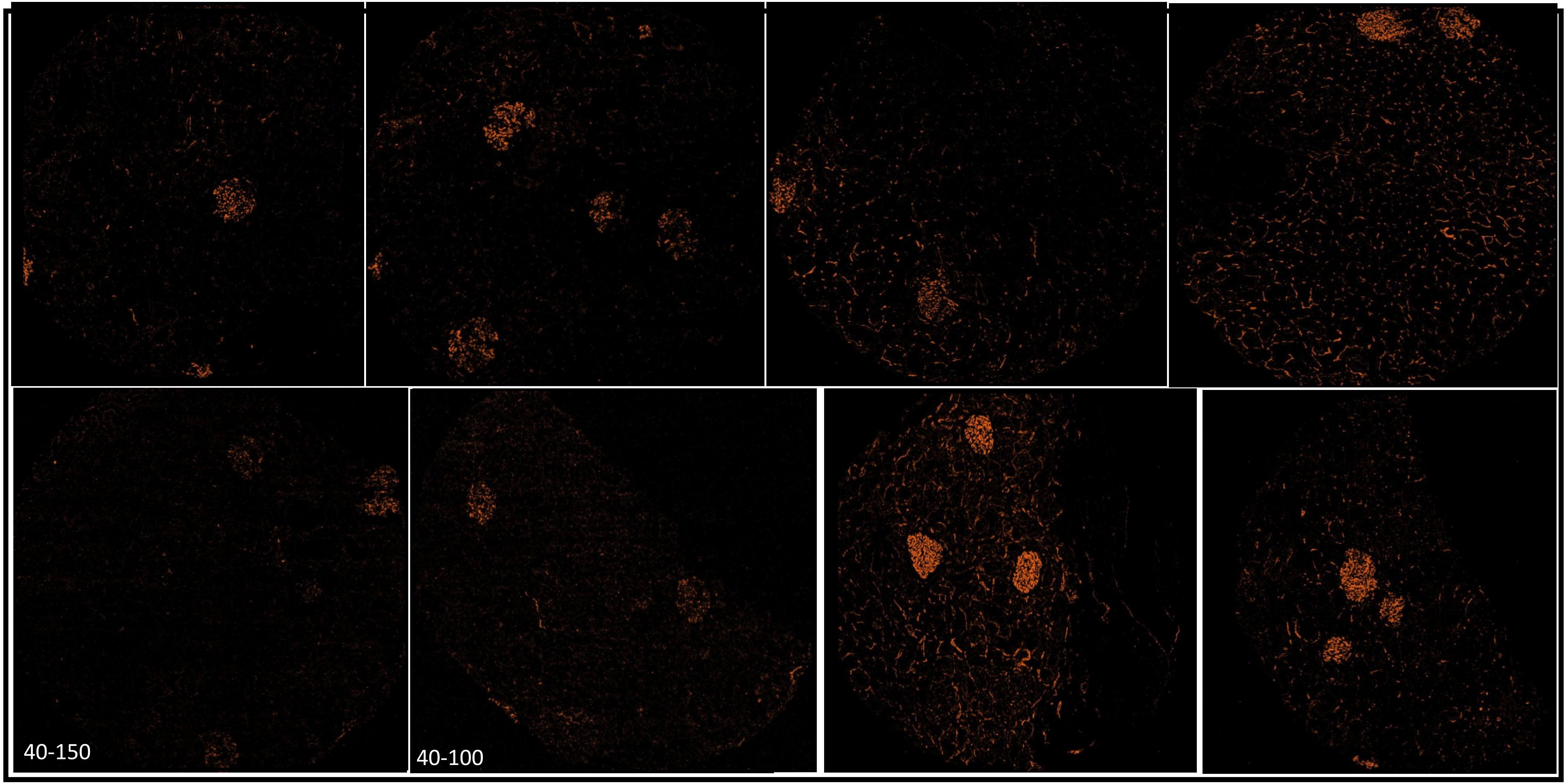

### CCR6 (50-200)

DKDIIA

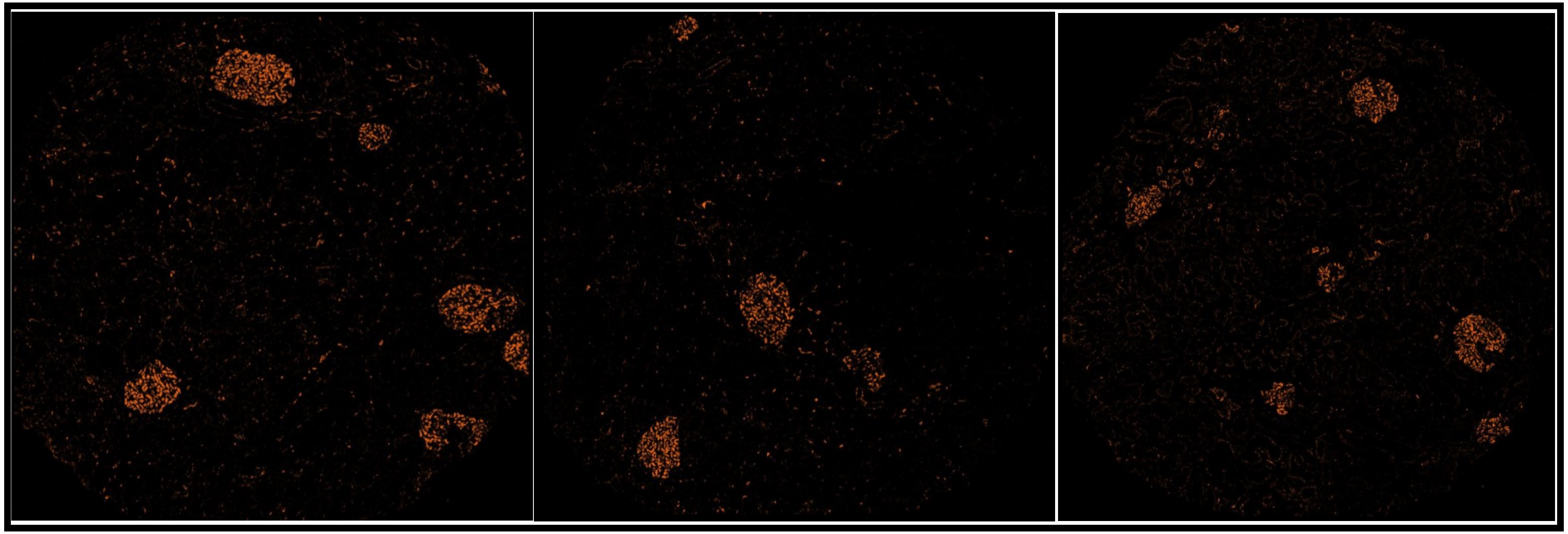

DKDIIA-B

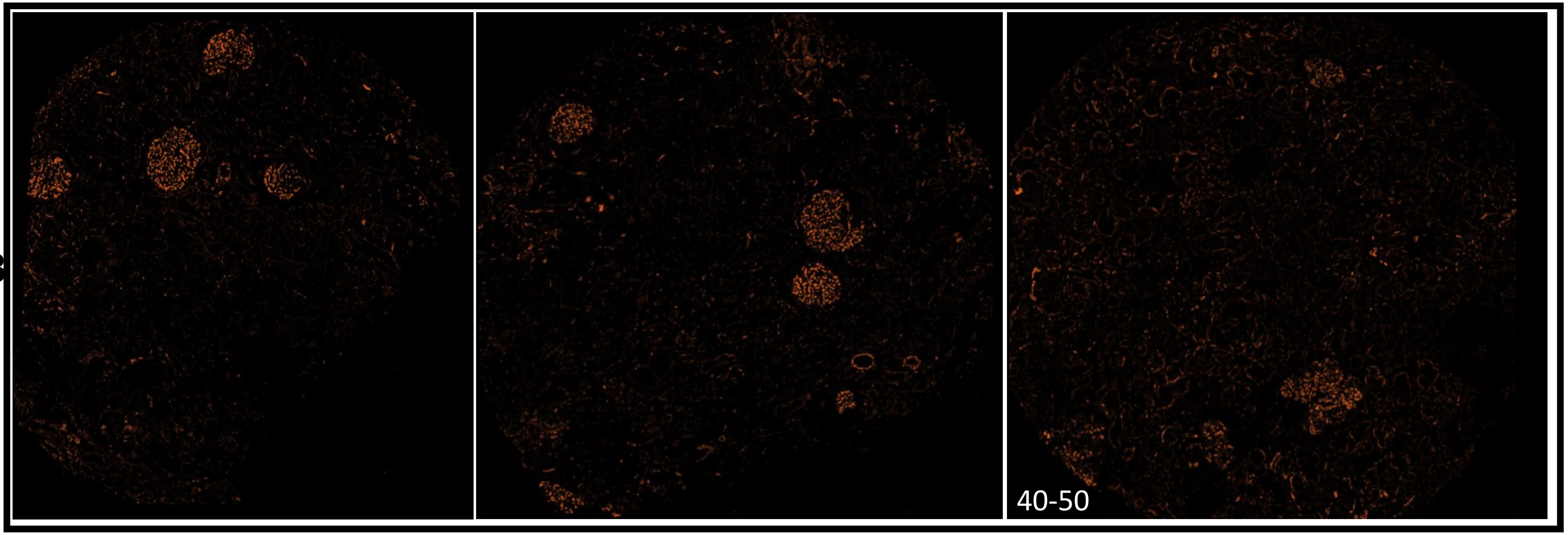

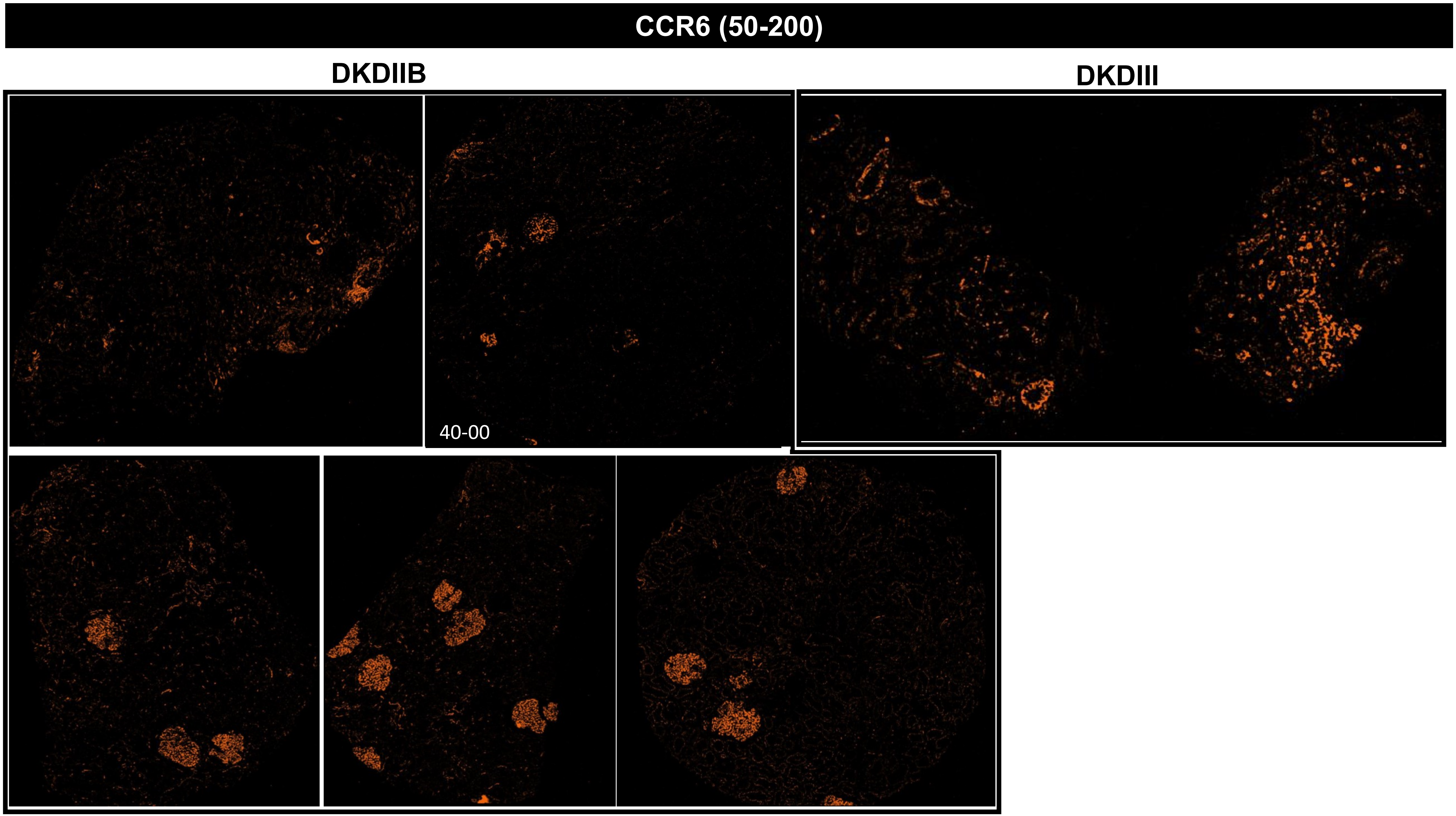
CCR6 (50-200)
DKDIIB
DKDIII
40-00

### CCR6 (50-200)

#### MEDULLA

### DM

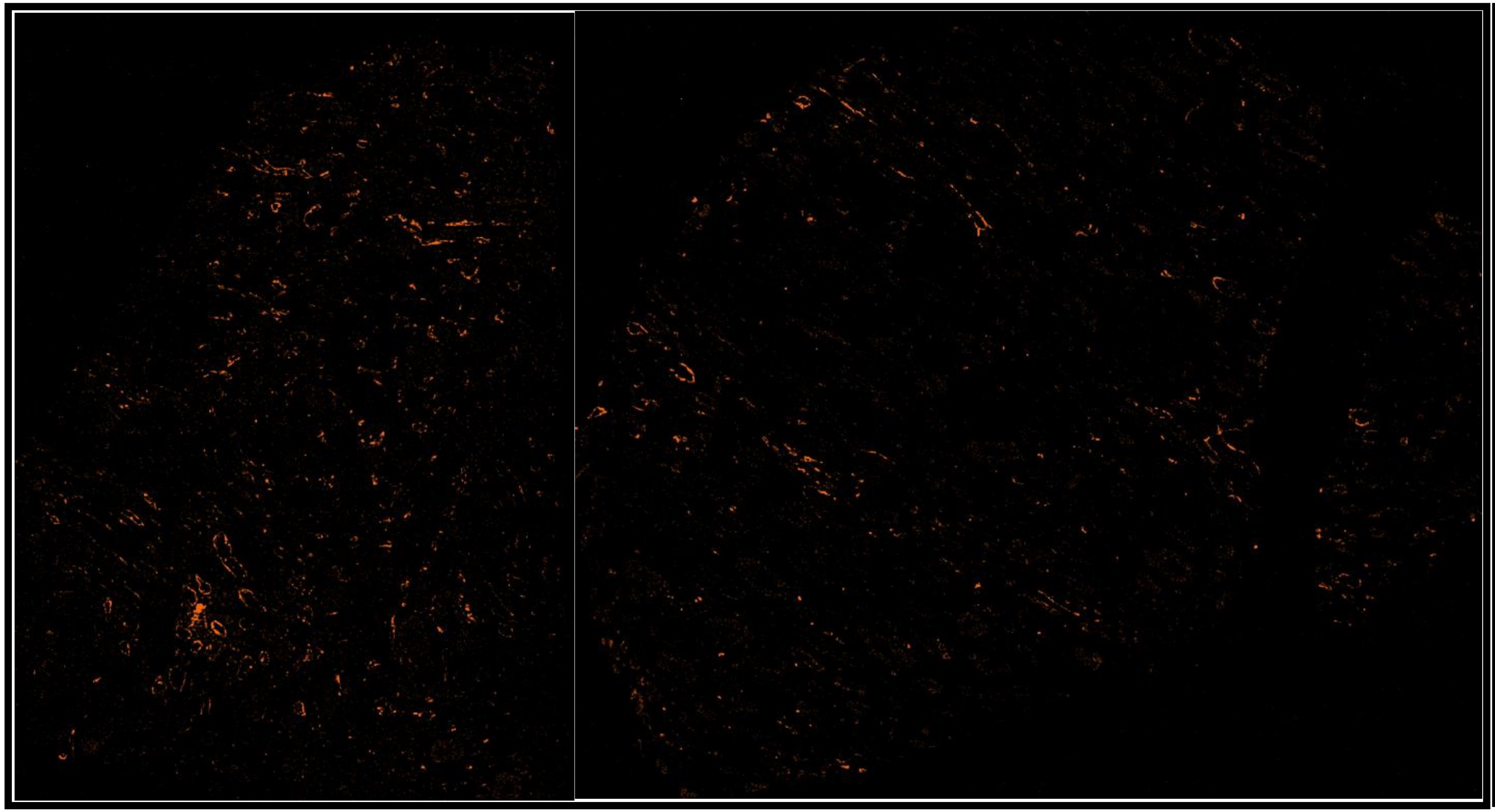

##### DKDIIA-B

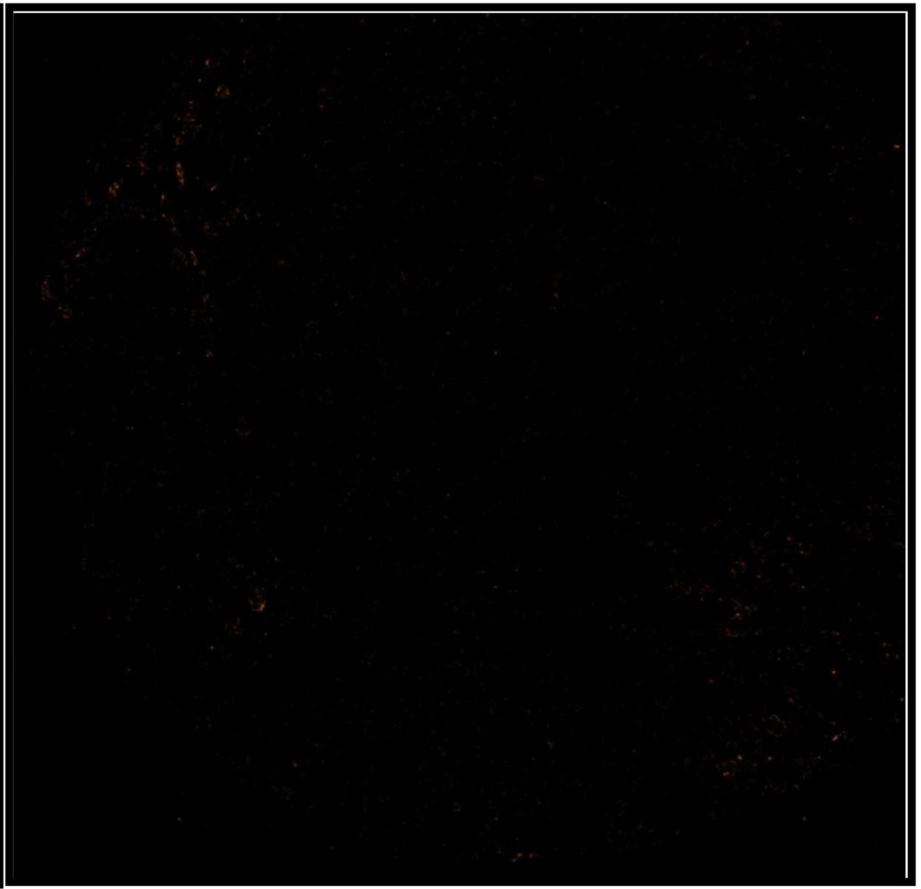

# CD107a (20-1000)

## DM

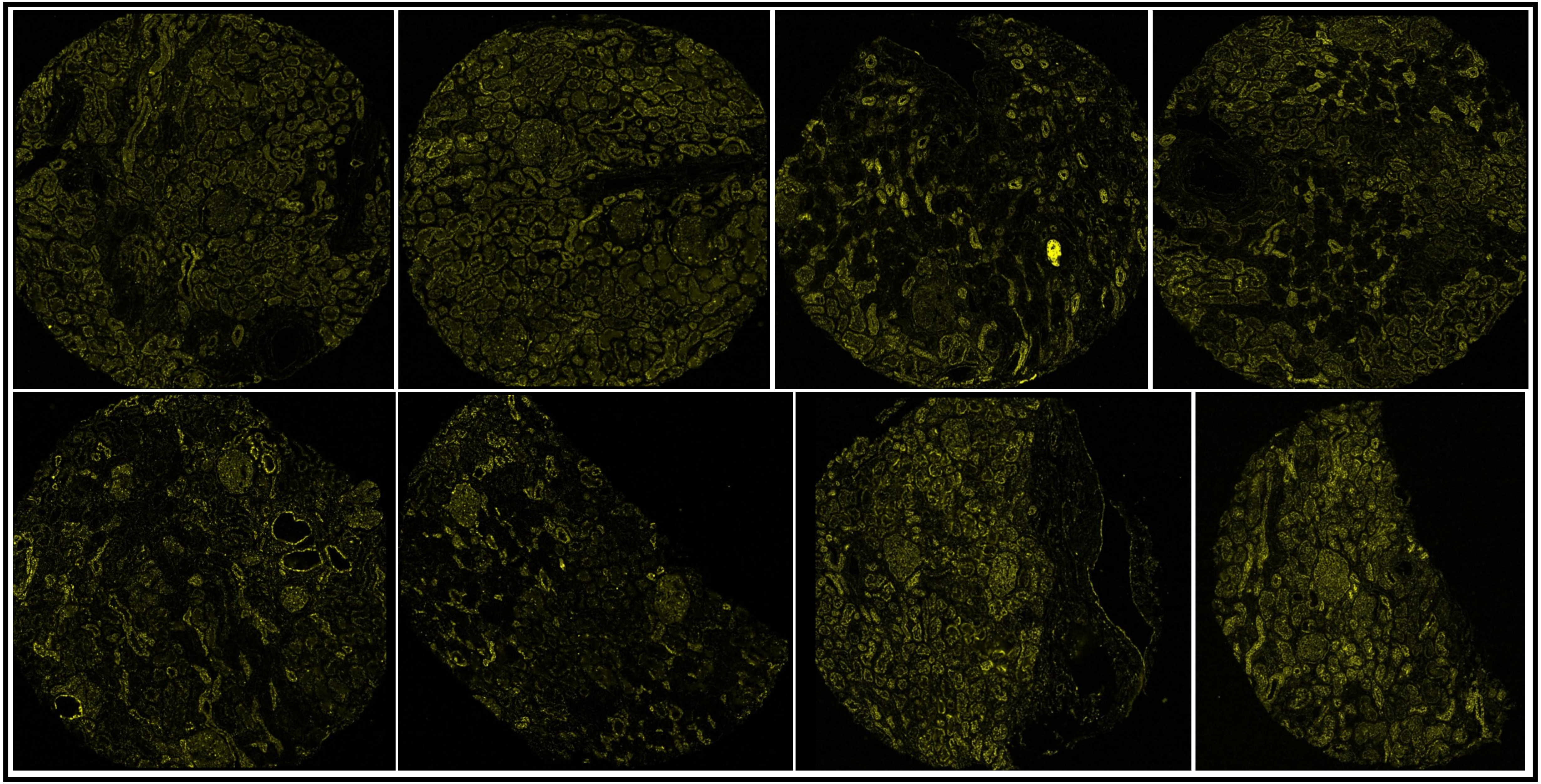

# CD107a (20-1000)

DKDIIA

DKDIIA-B

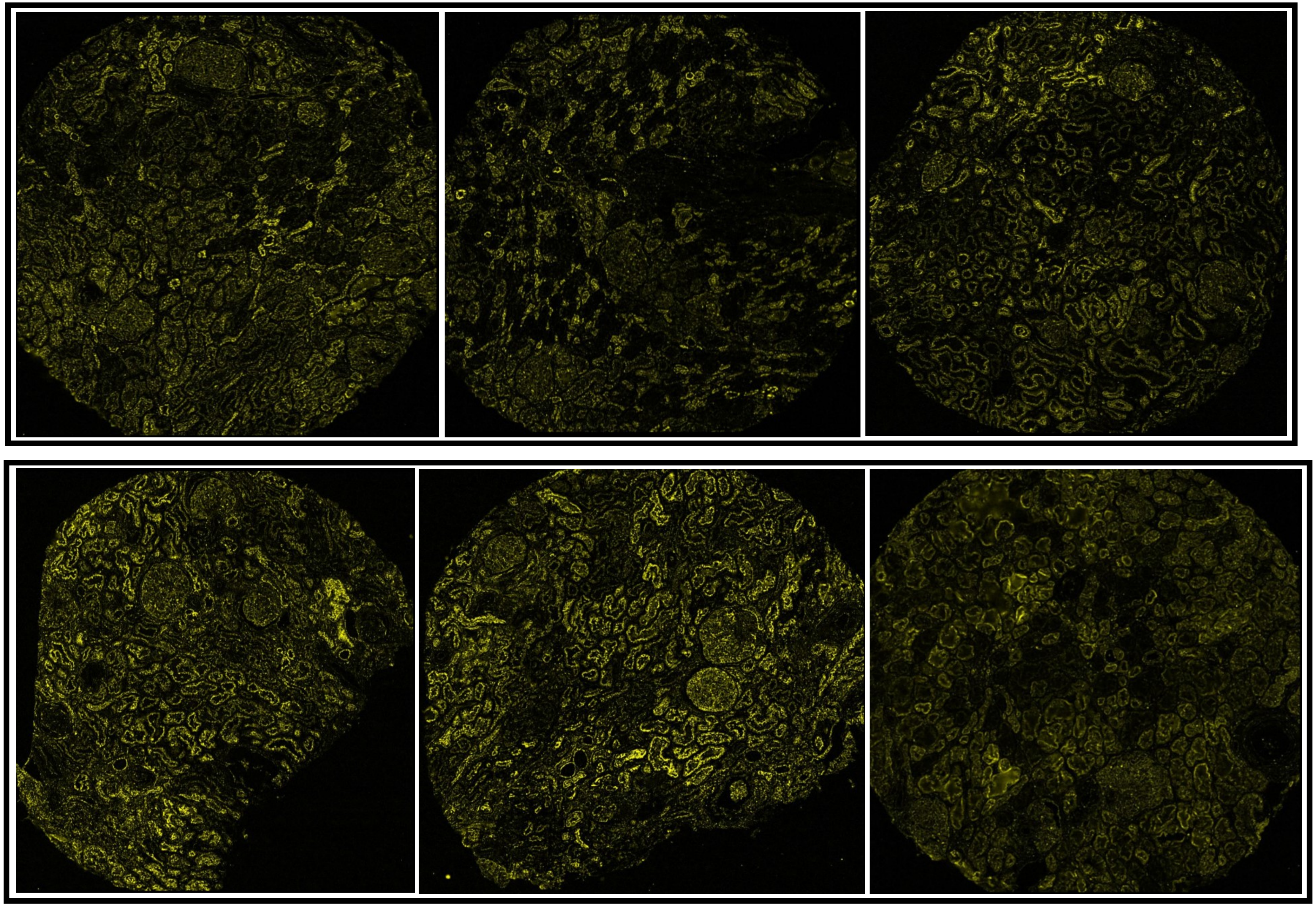

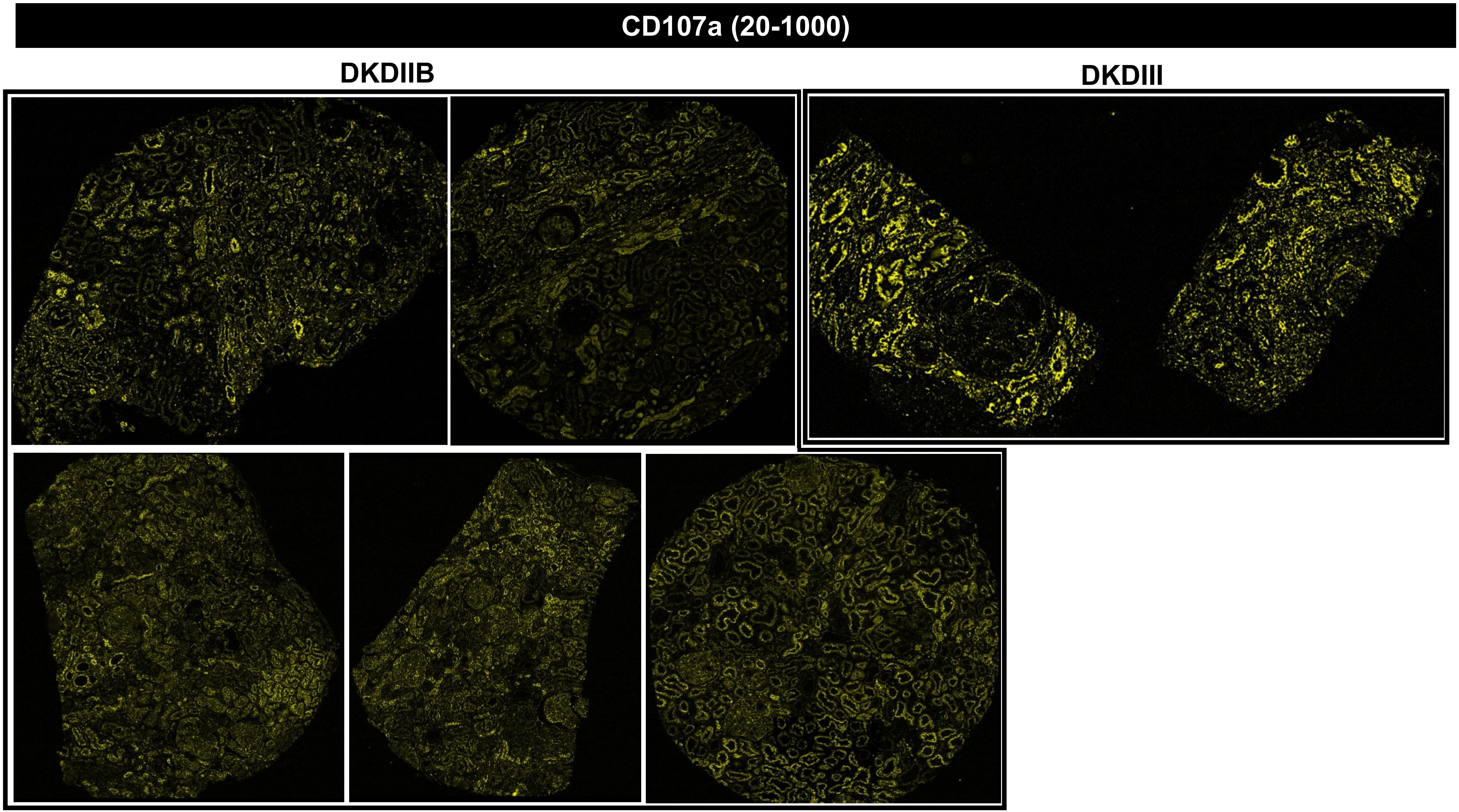
CD107a (20-1000)
DKDIIB
DKDIII

CD107a (20-1000)

### MEDULLA

DM DKDIIA-B

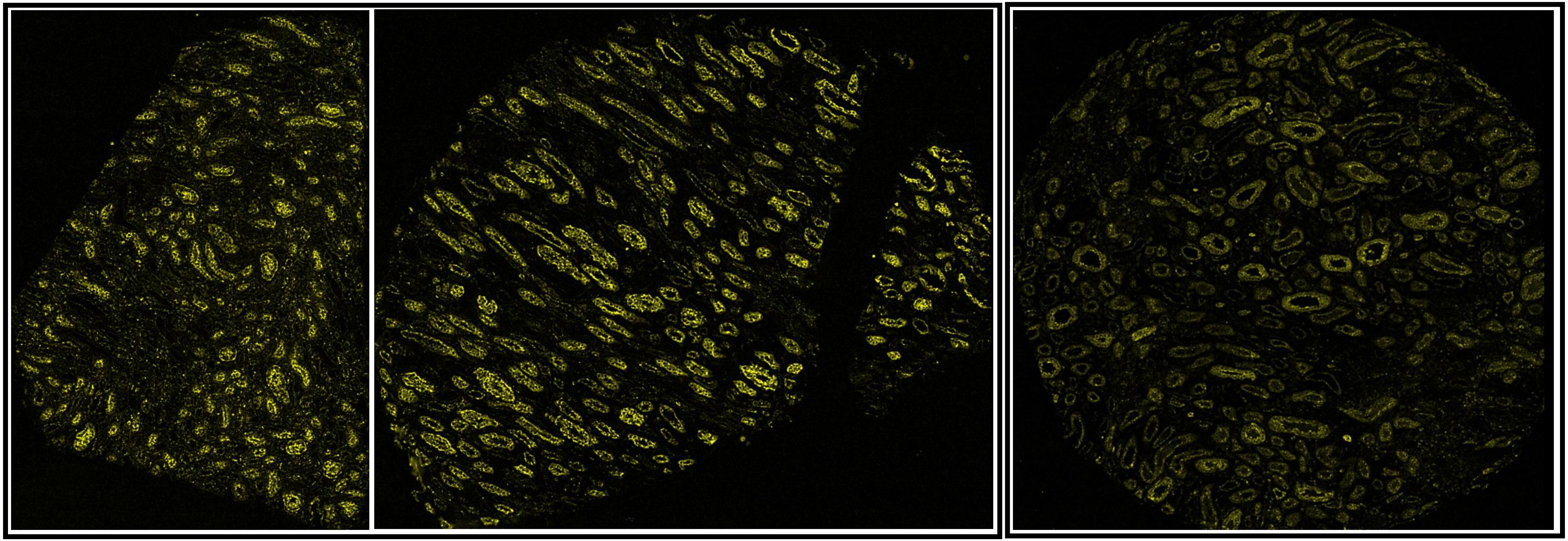

# CD11b (100-500)

## DM

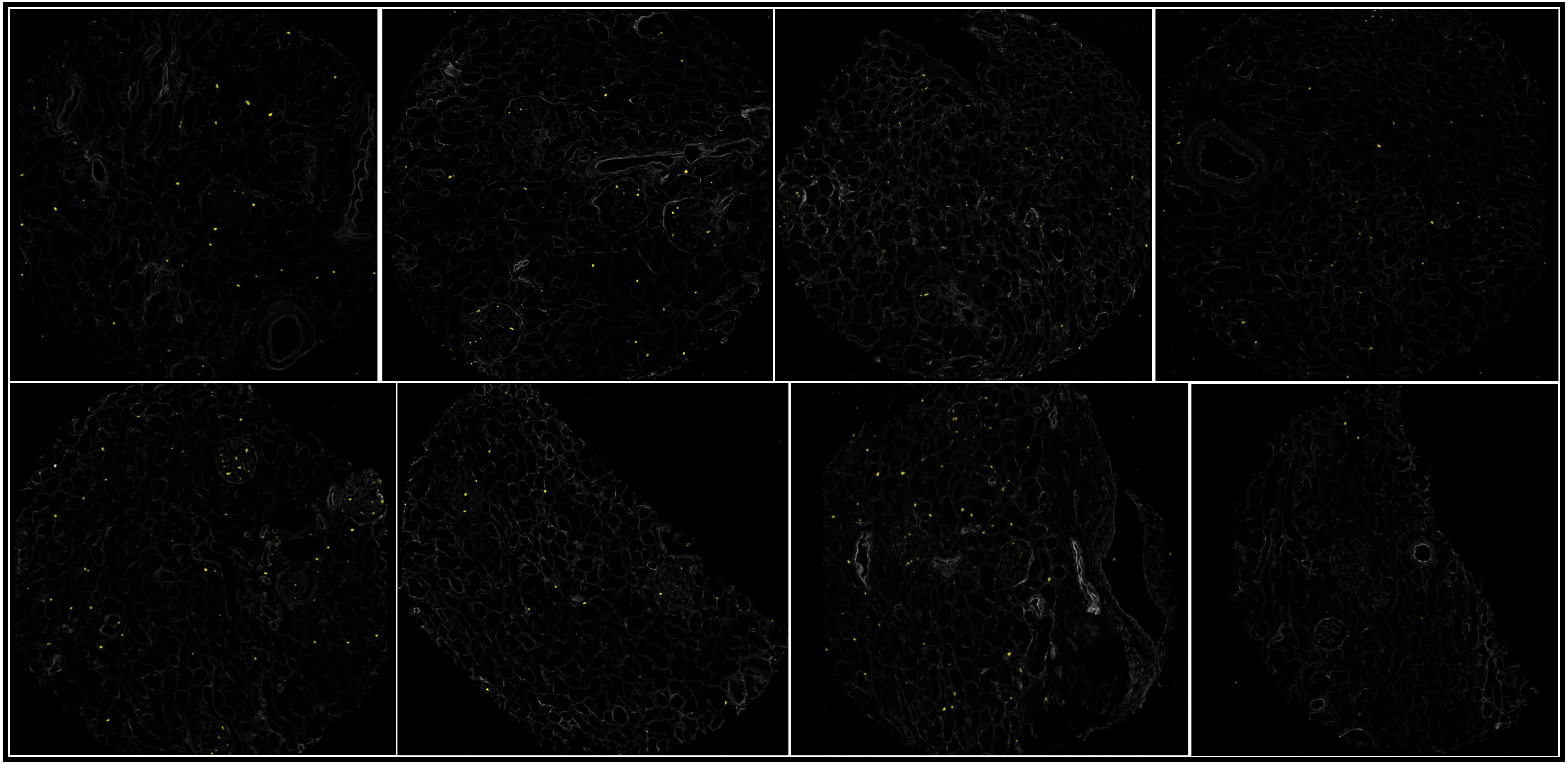

## CD11b (100-500)

DKDIIA

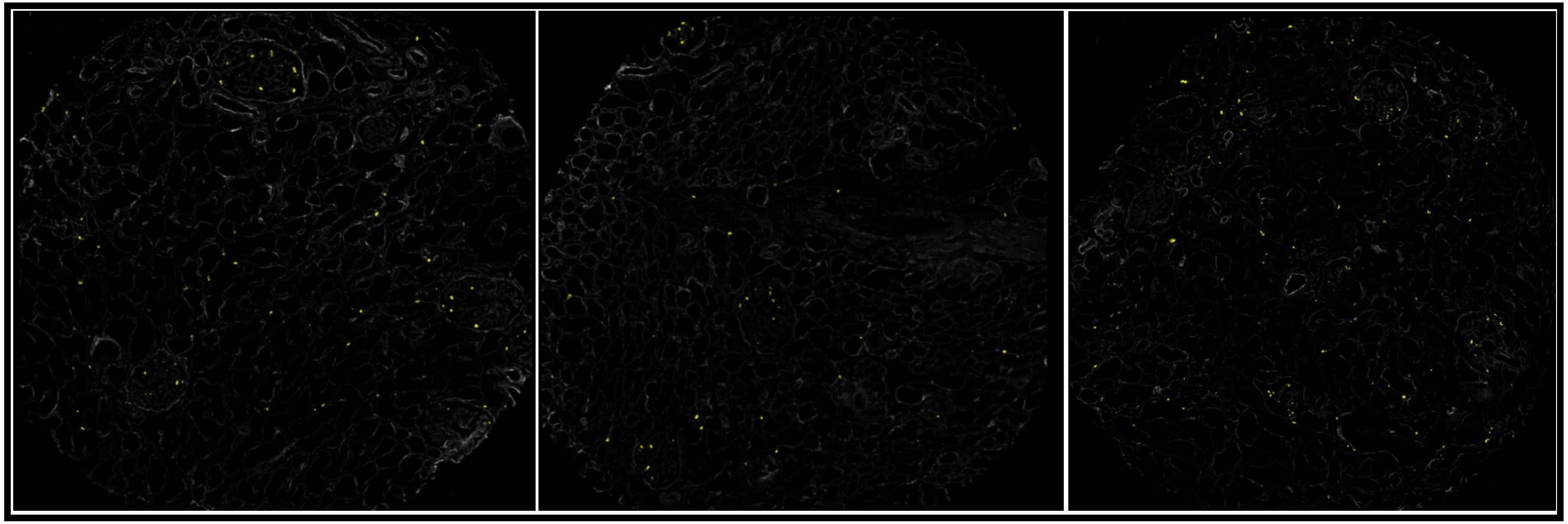

DKDIIA-B

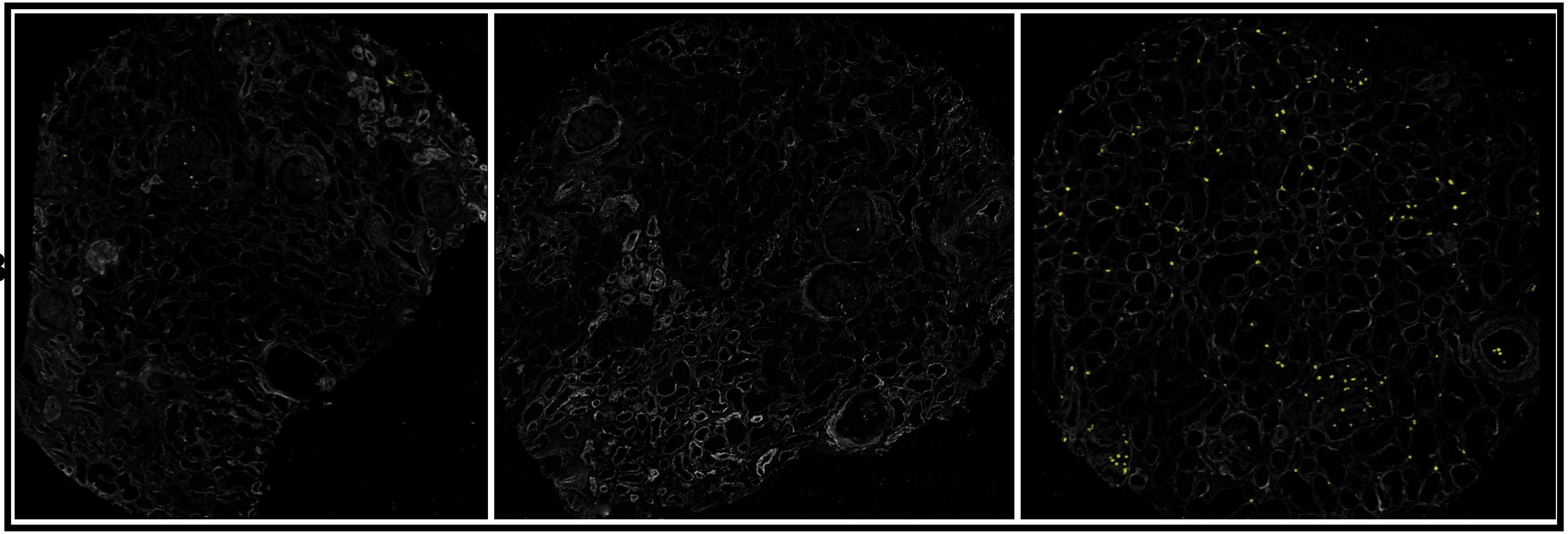

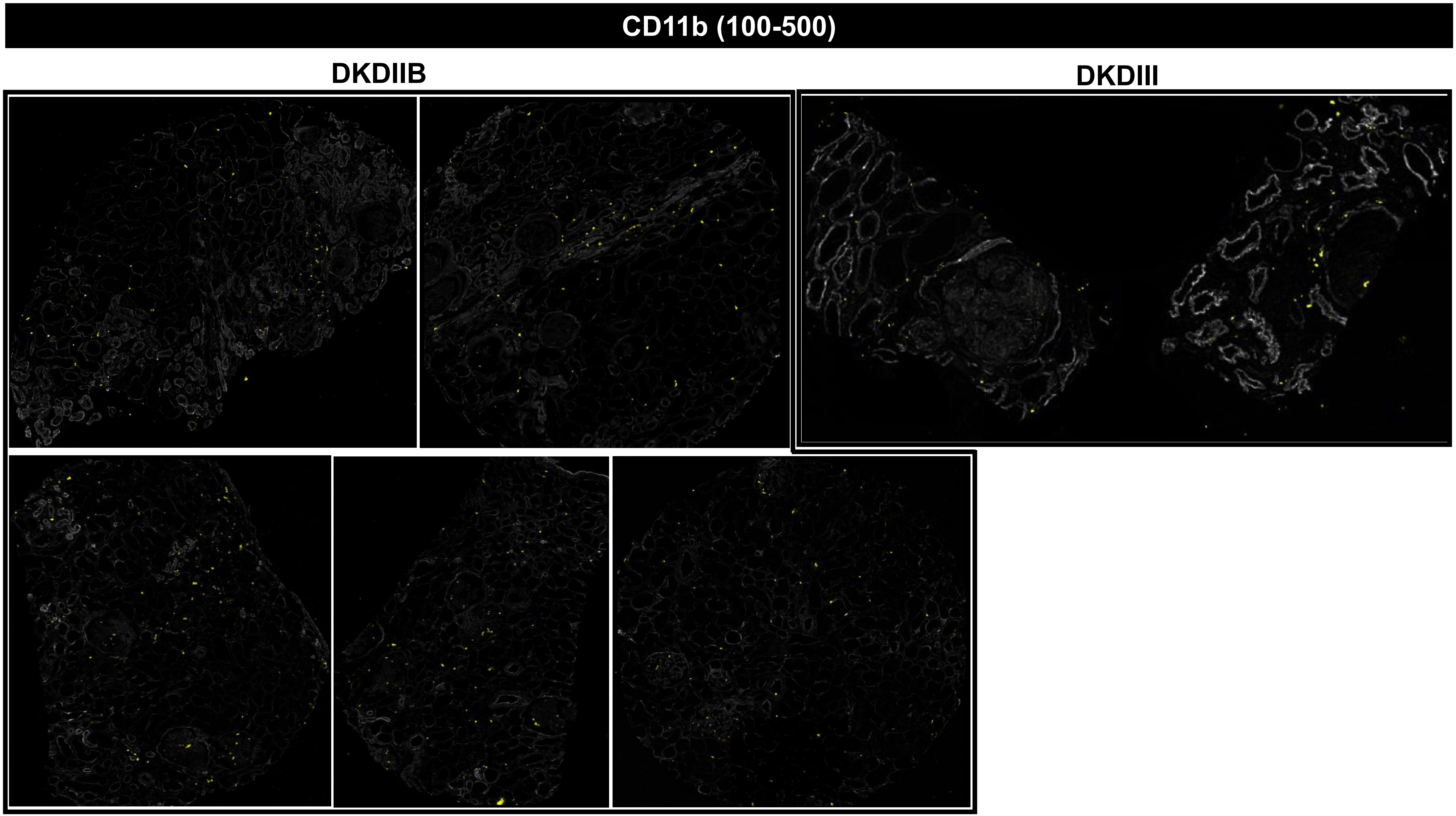
CD11b (100-500)
DKDIIB
DKDIII

# CD11b (100-500)

#### MEDULLA

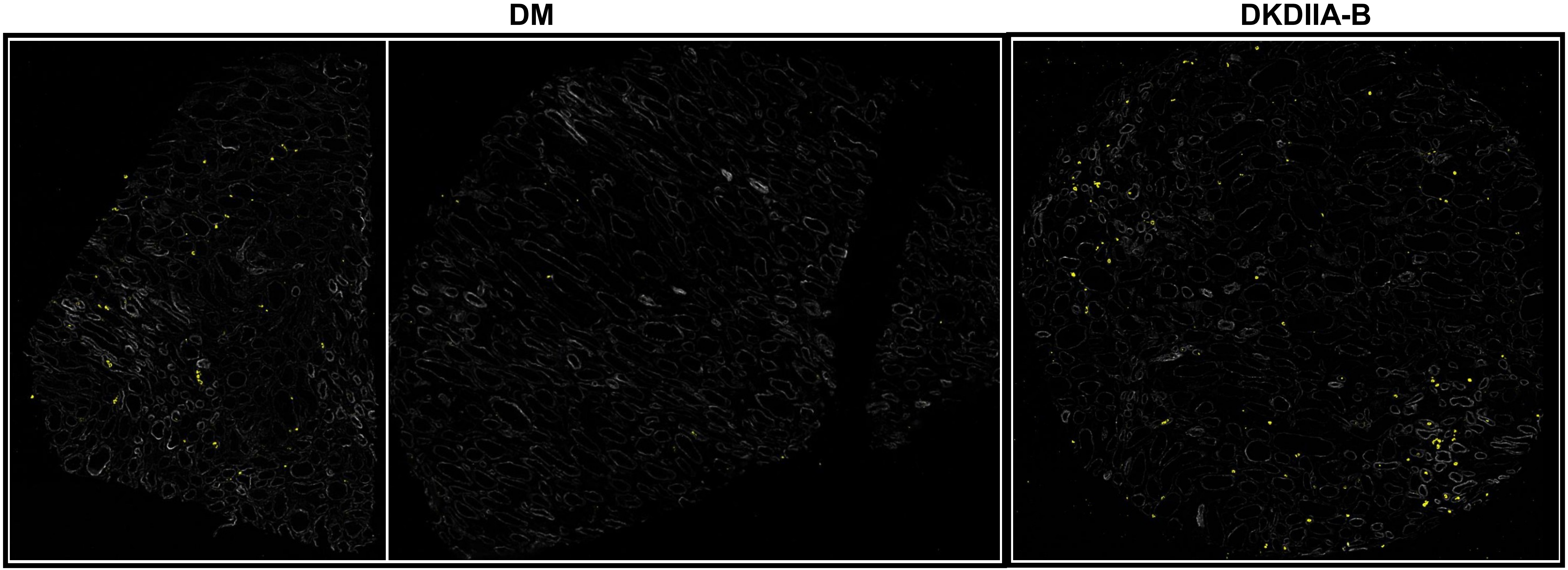

# CD227 (20-2000)

## DM

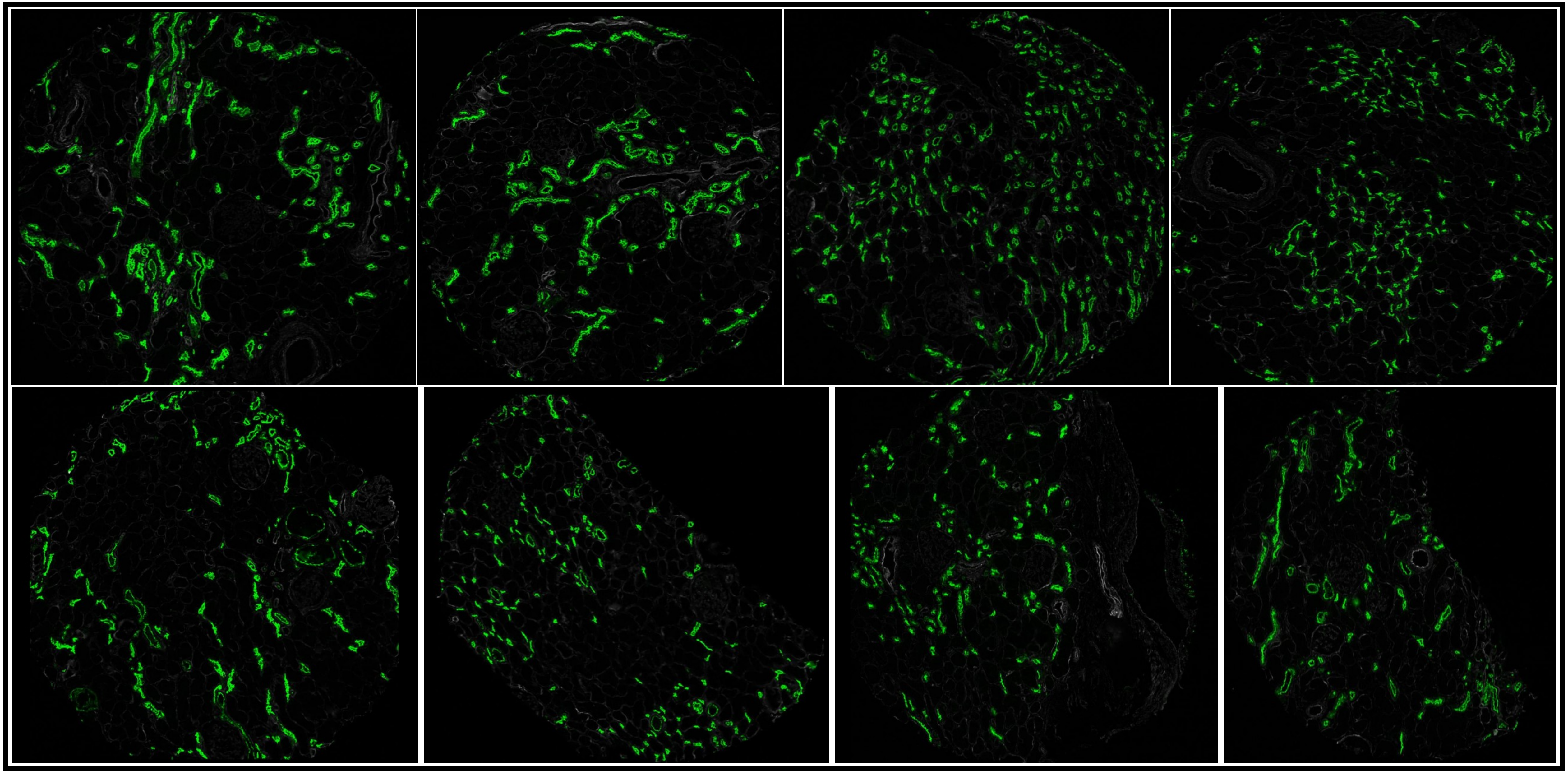

# CD227 (20-2000)

DKDIIA

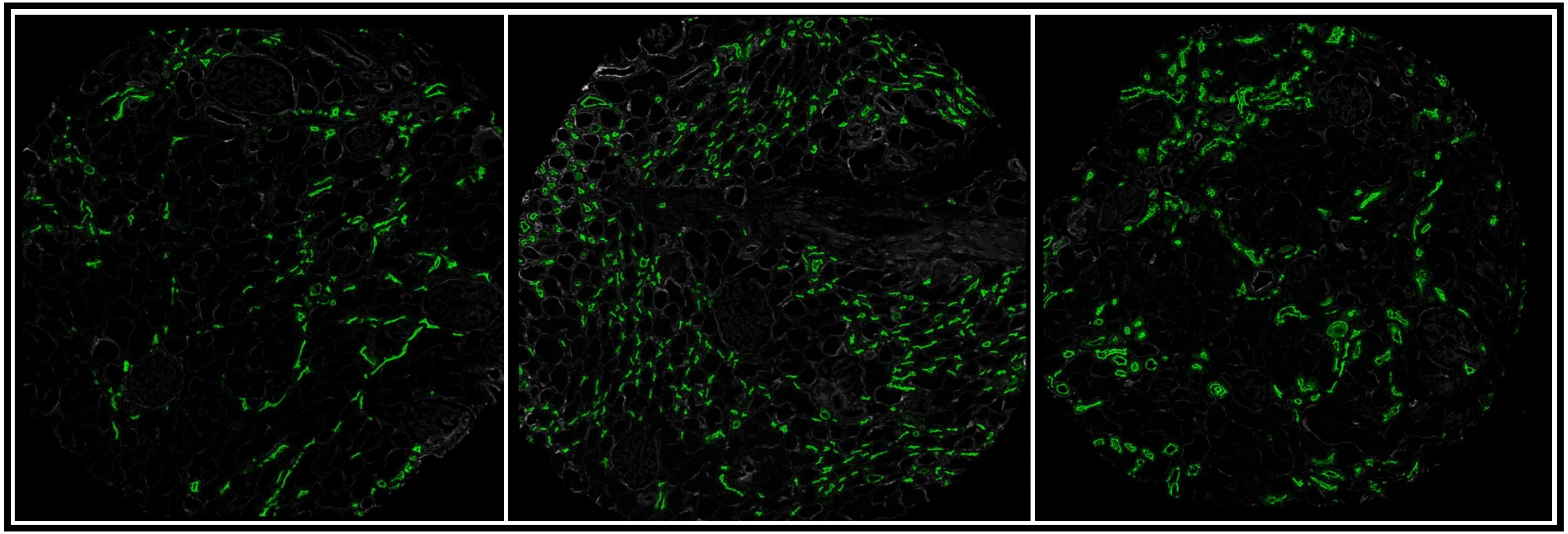

DKDIIA-B

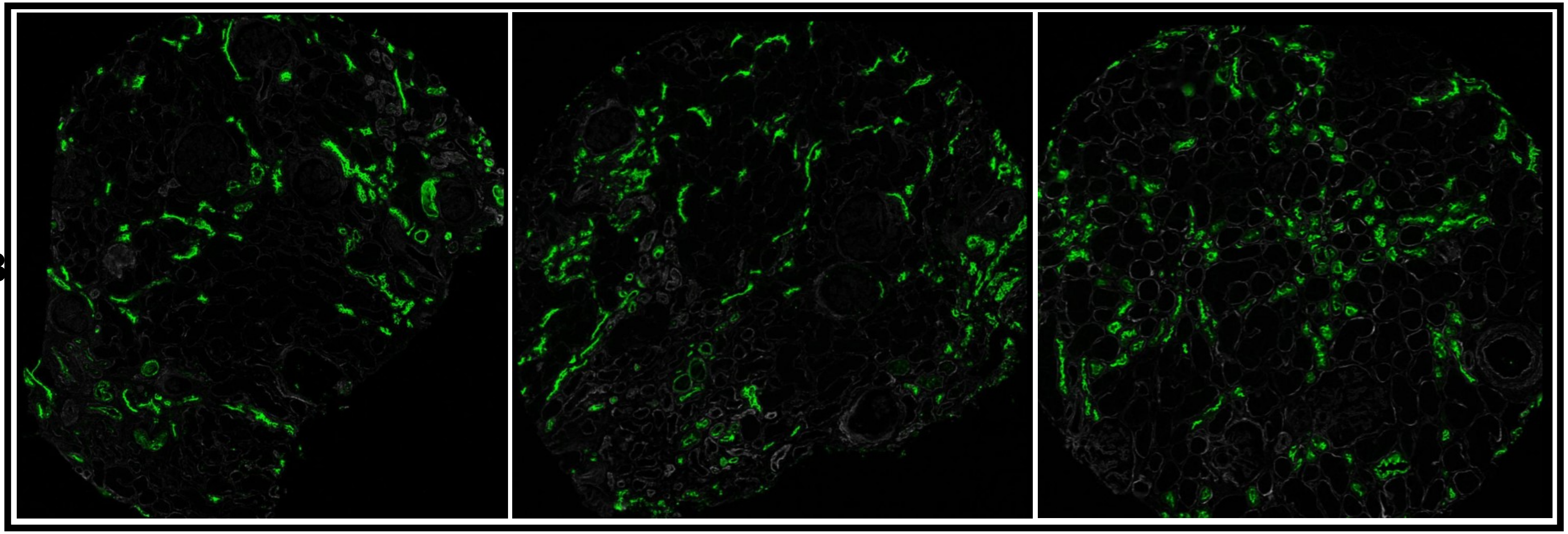

# CD227 (20-2000)

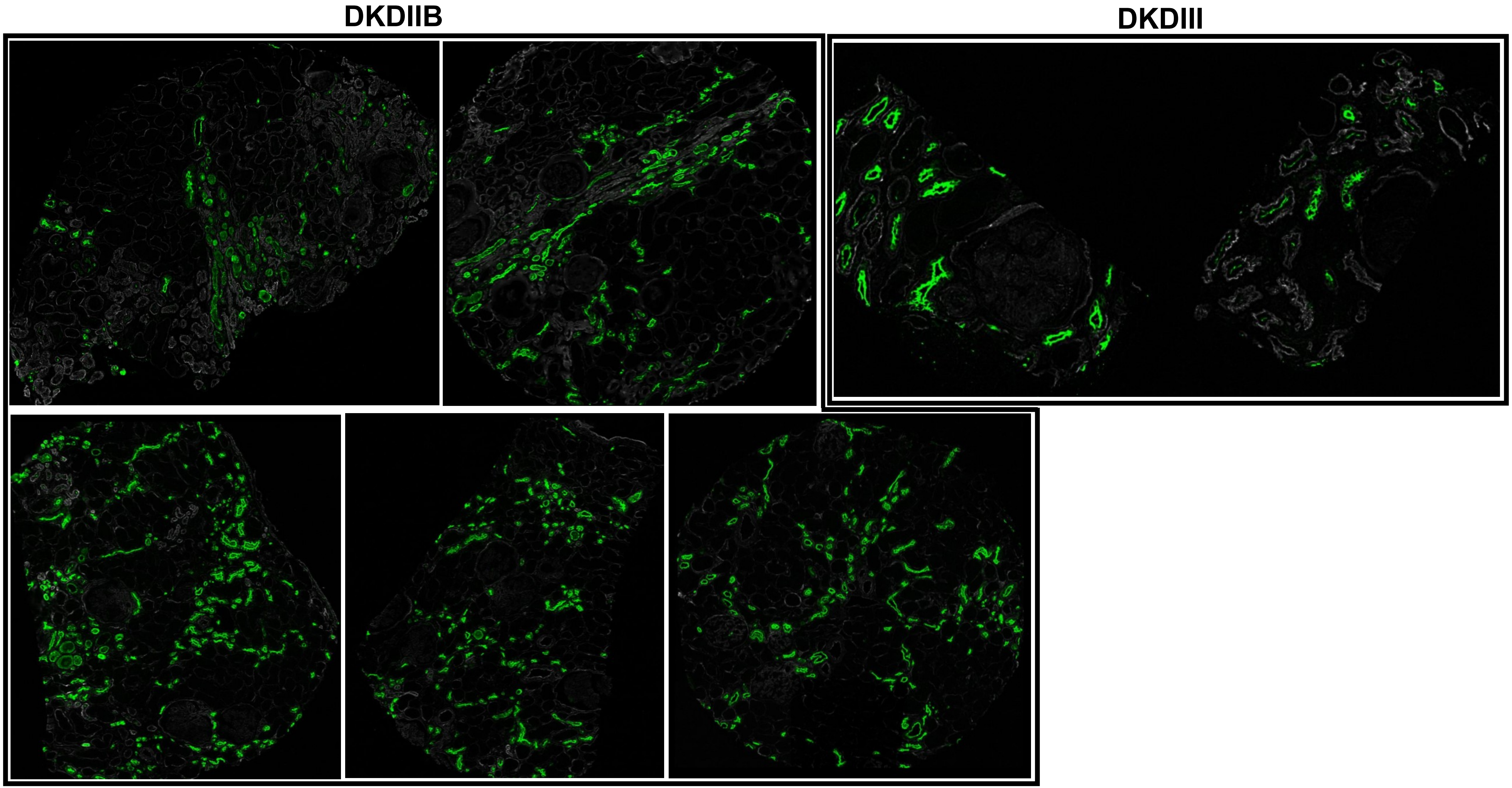

# CD227 (20-2000)

#### MEDULLA

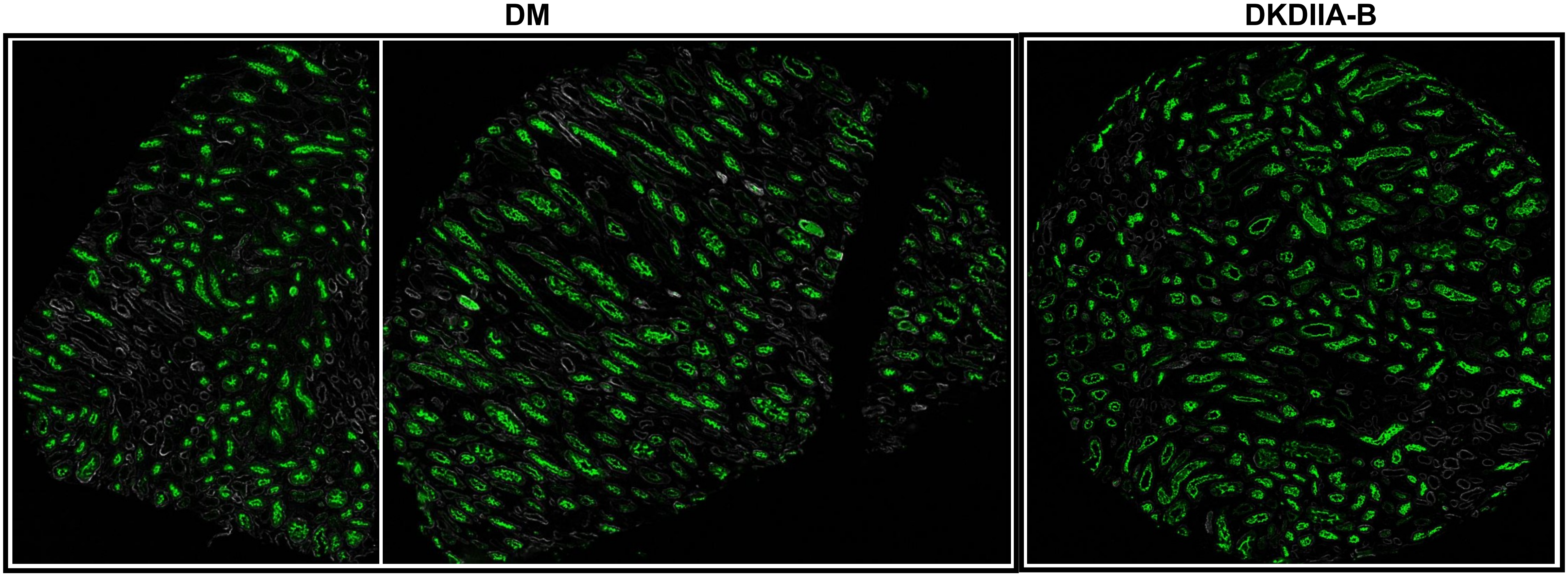

# CD31 (50-100)

## DM

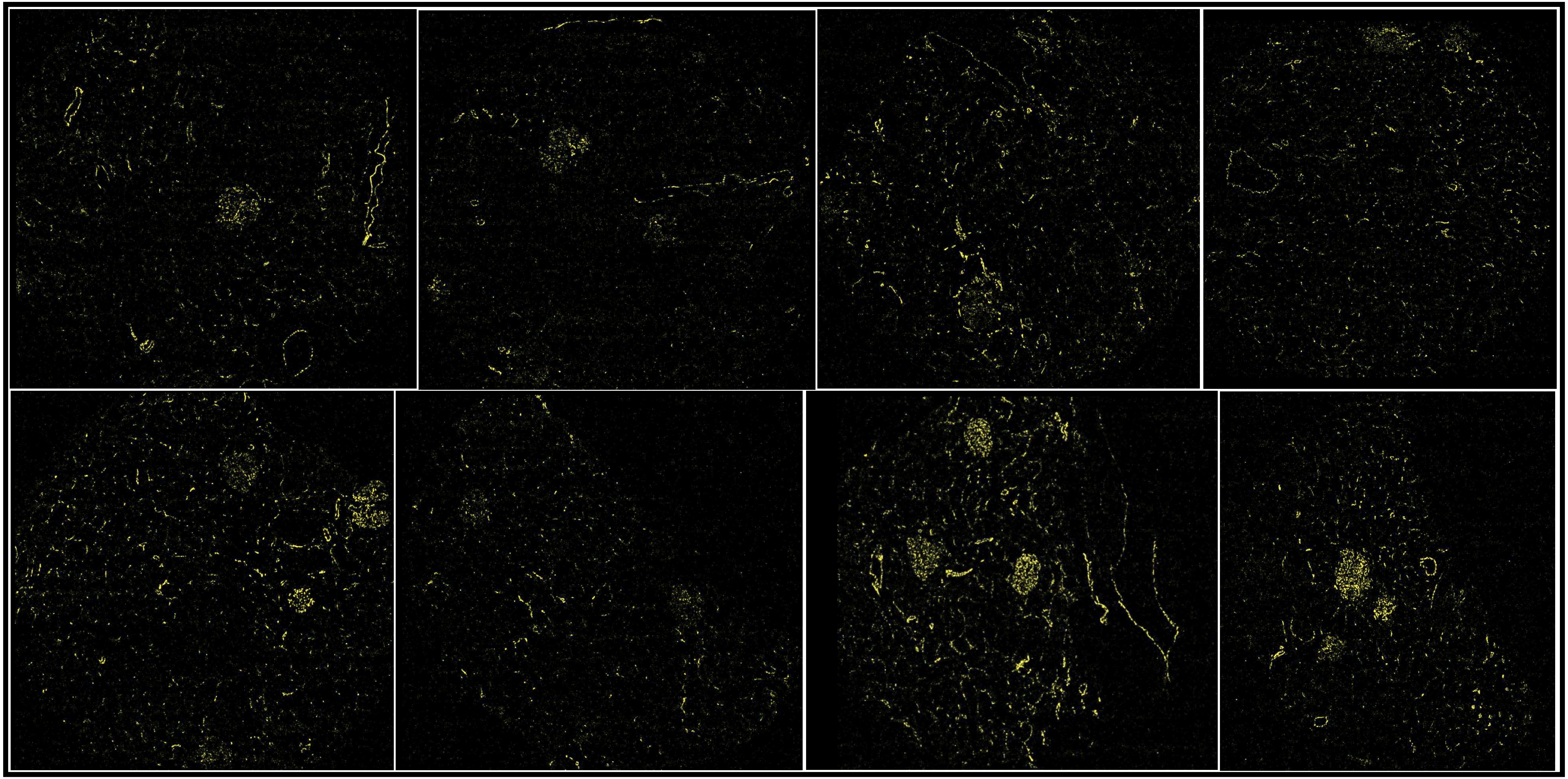

CD31 (50-100)

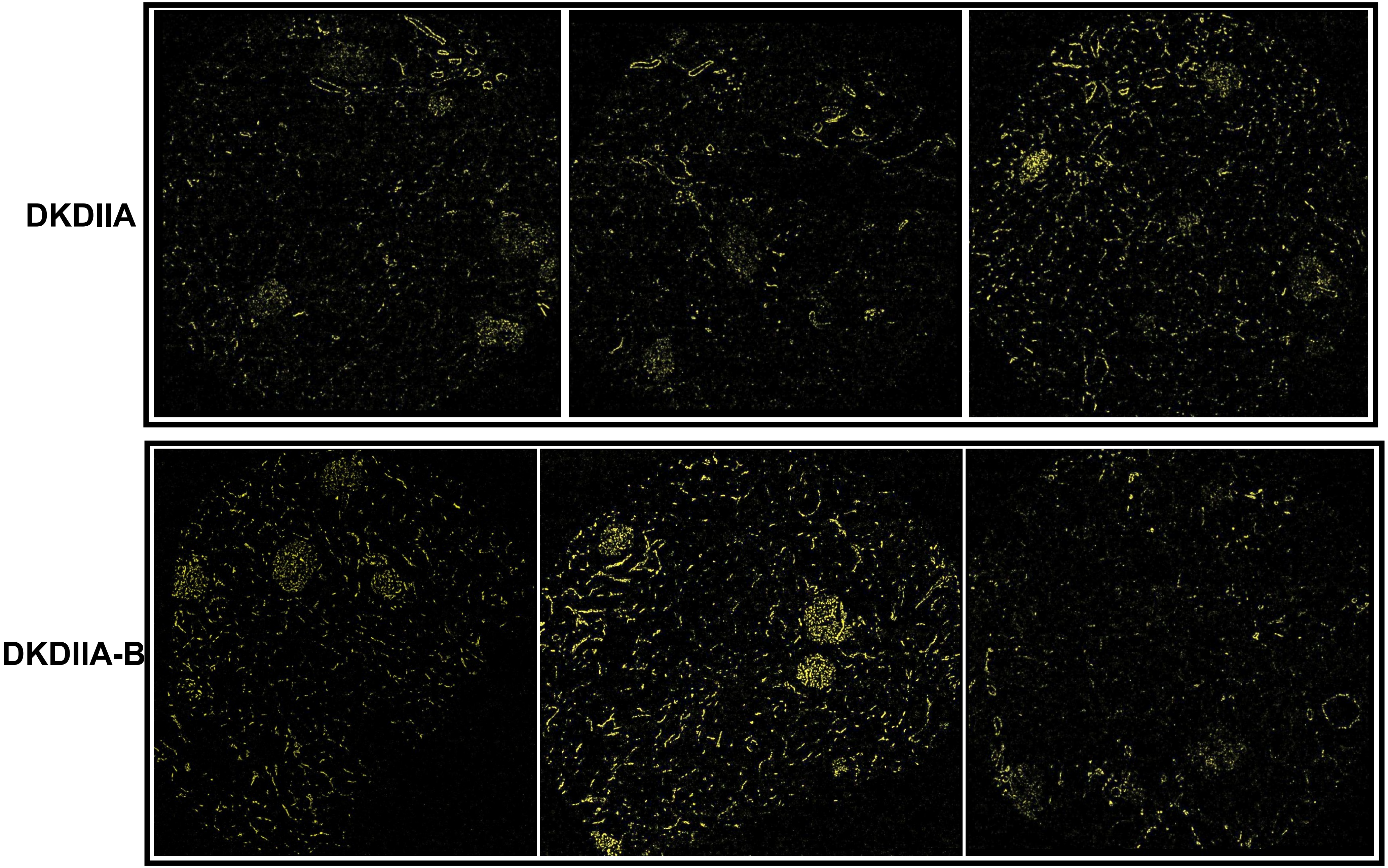

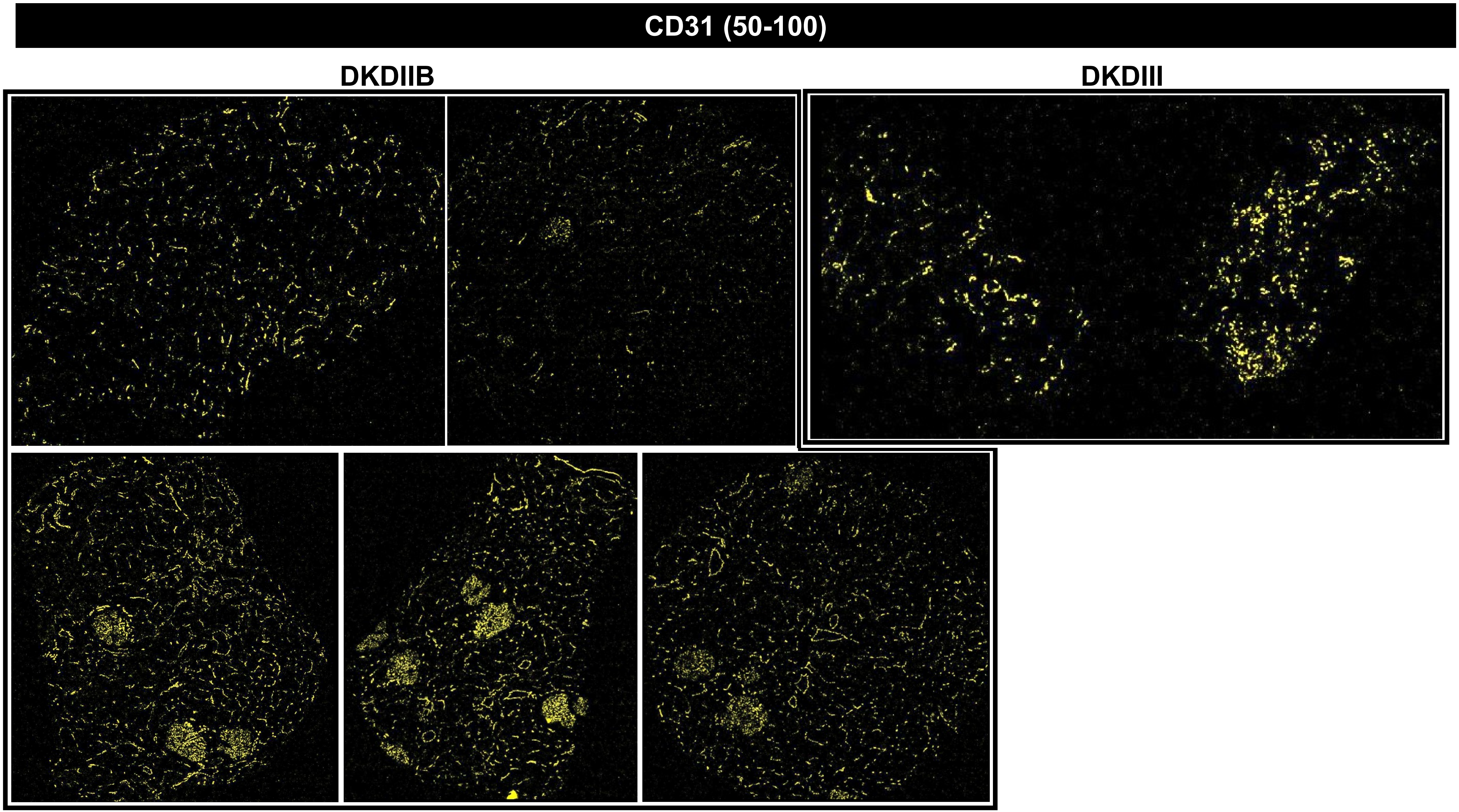
CD31 (50-100)
DKDIIB
DKDIII

CD31 (50-100)

### MEDULLA

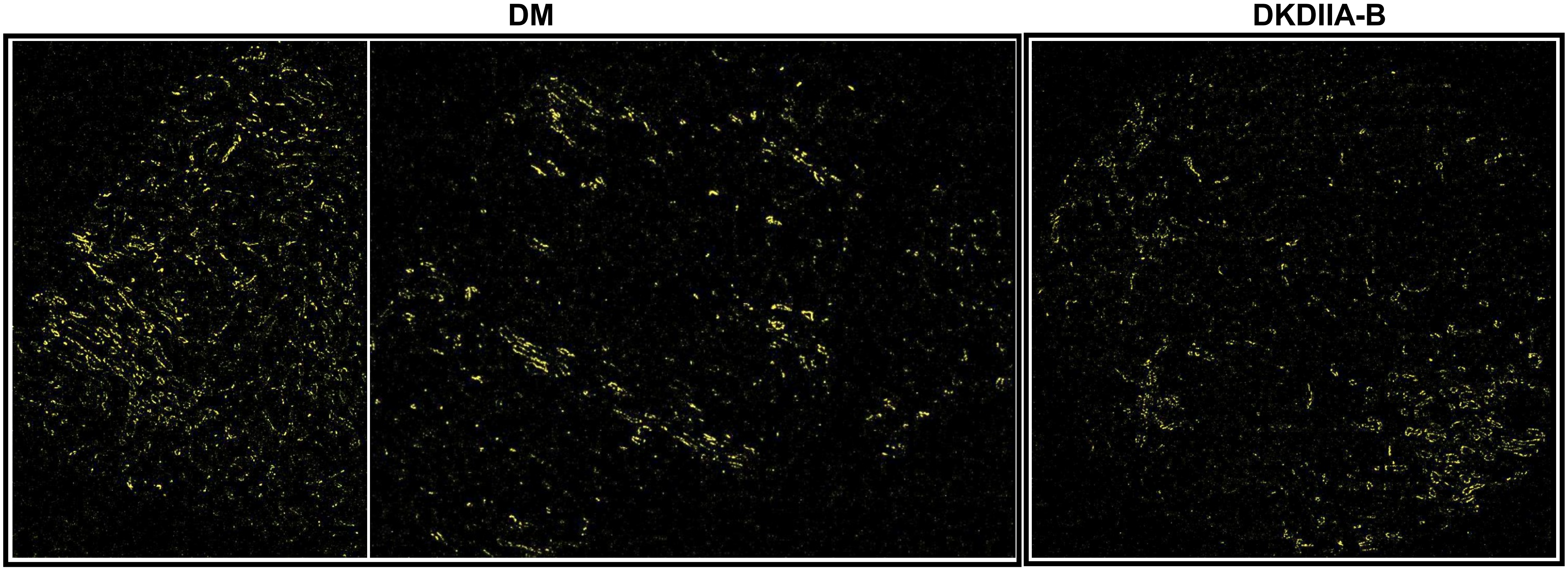

**CD45 (50-1500)**

**DM**

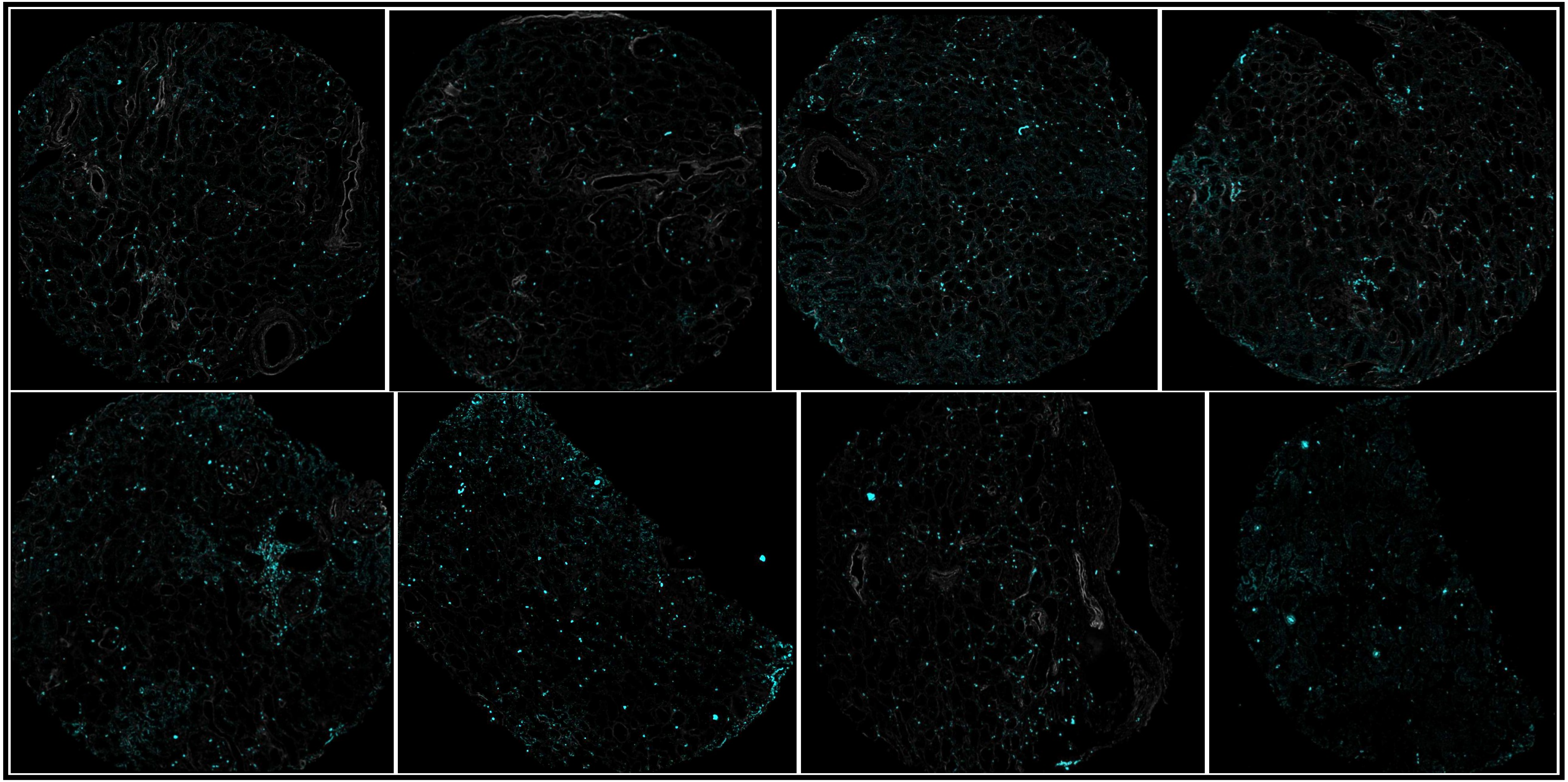

# CD45 (50-1500)

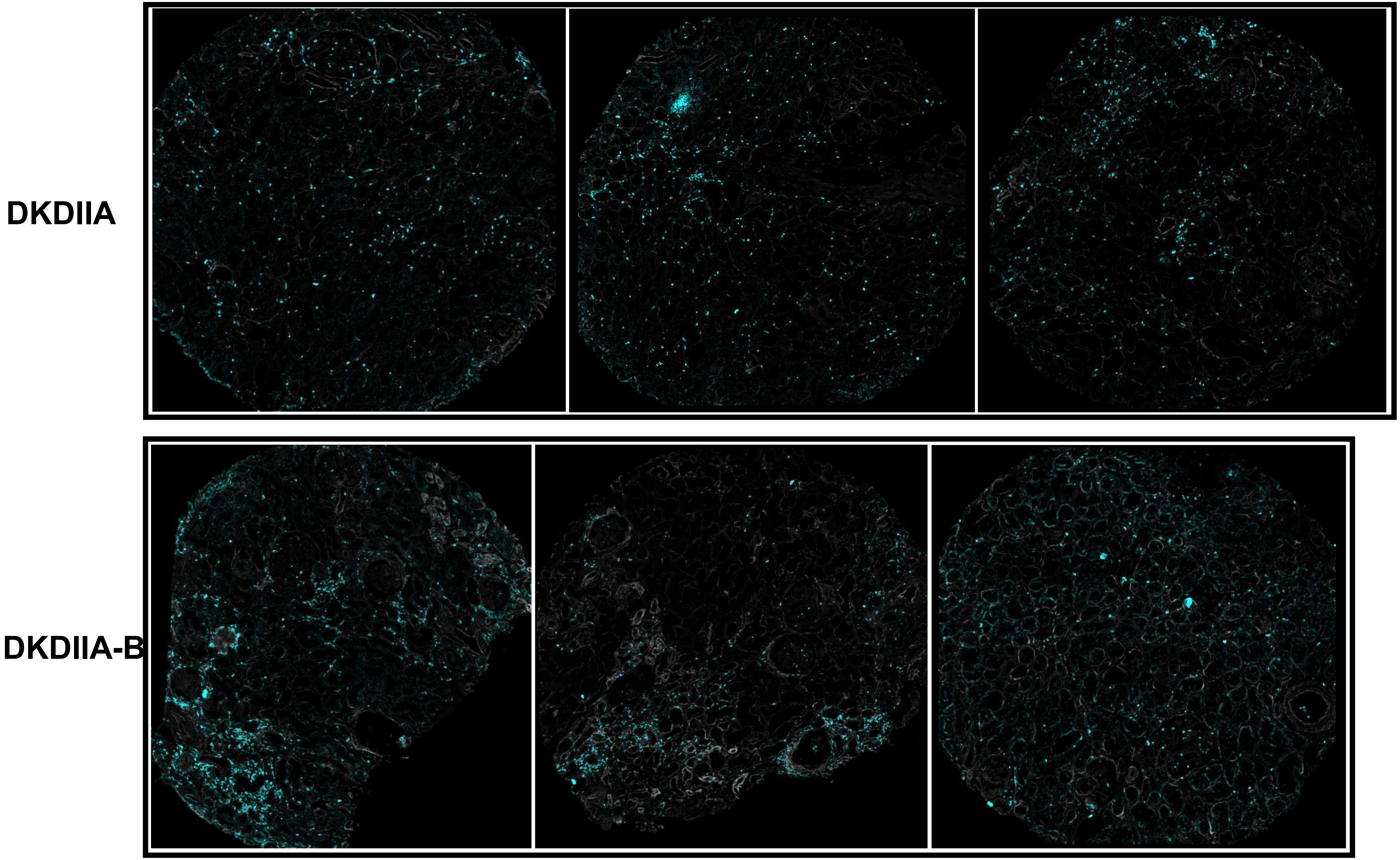

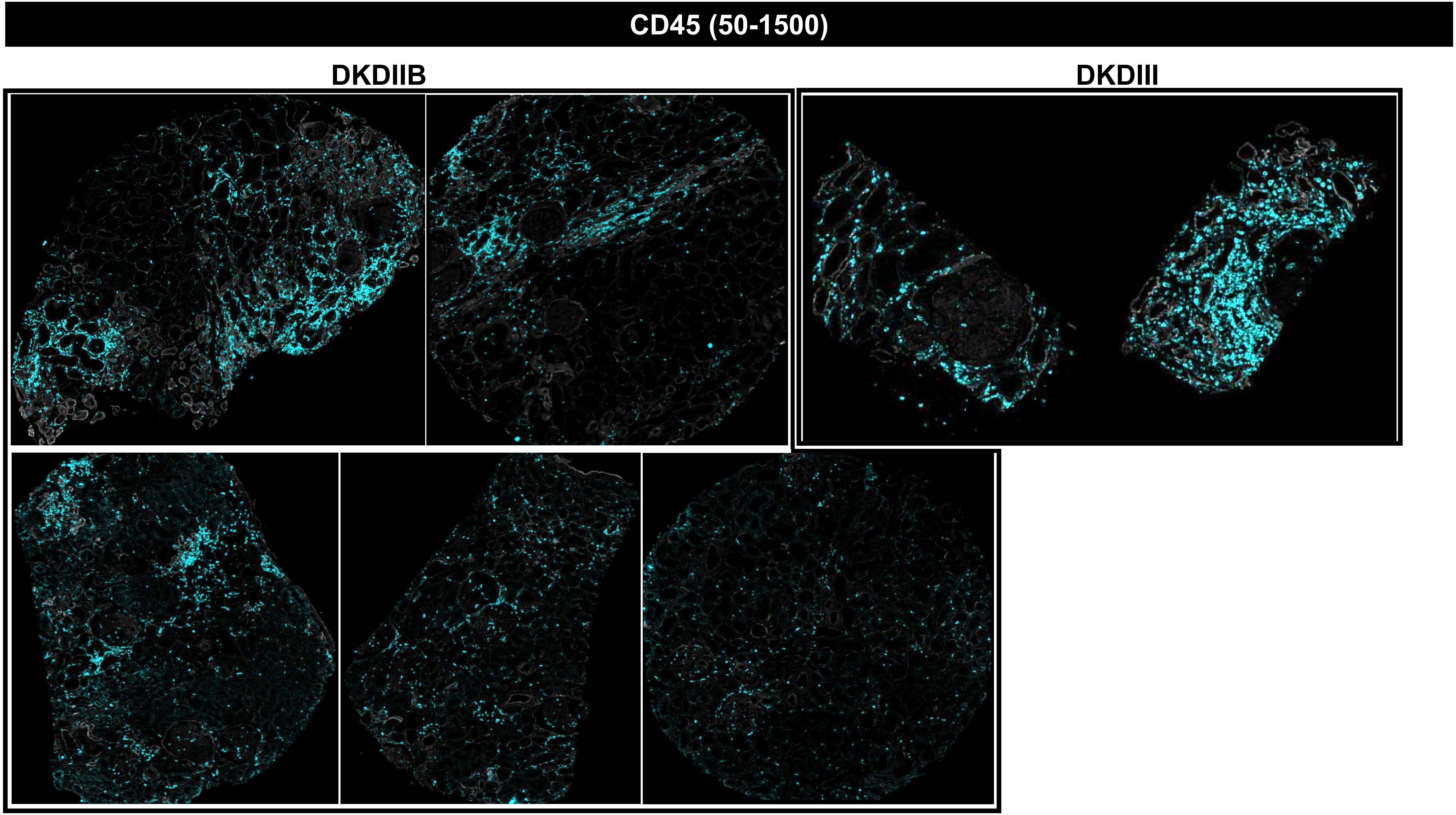
CD45 (50-1500)
DKDIIB
DKDIII

CD45 (50-1500)

### MEDULLA

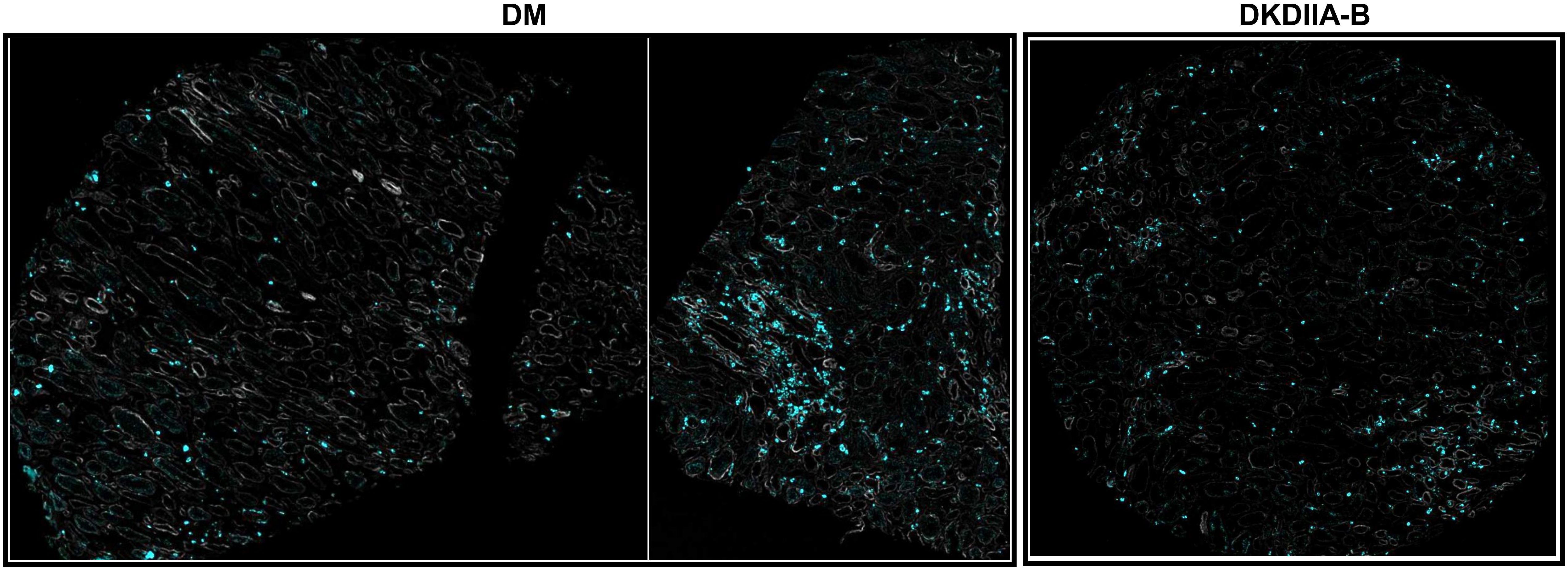

**CD68 (150-300)**

**DM**

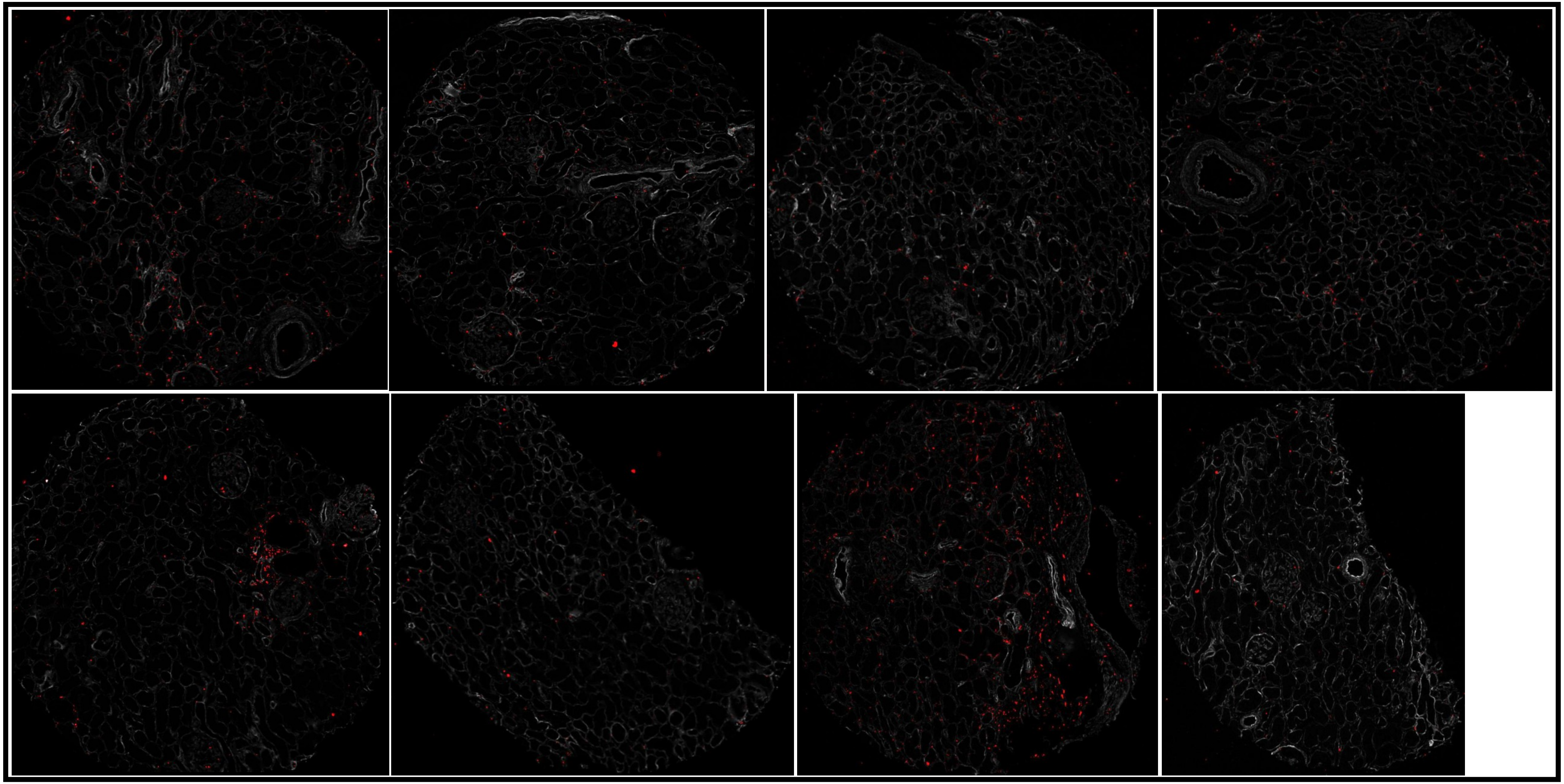

# CD68 (150-300)

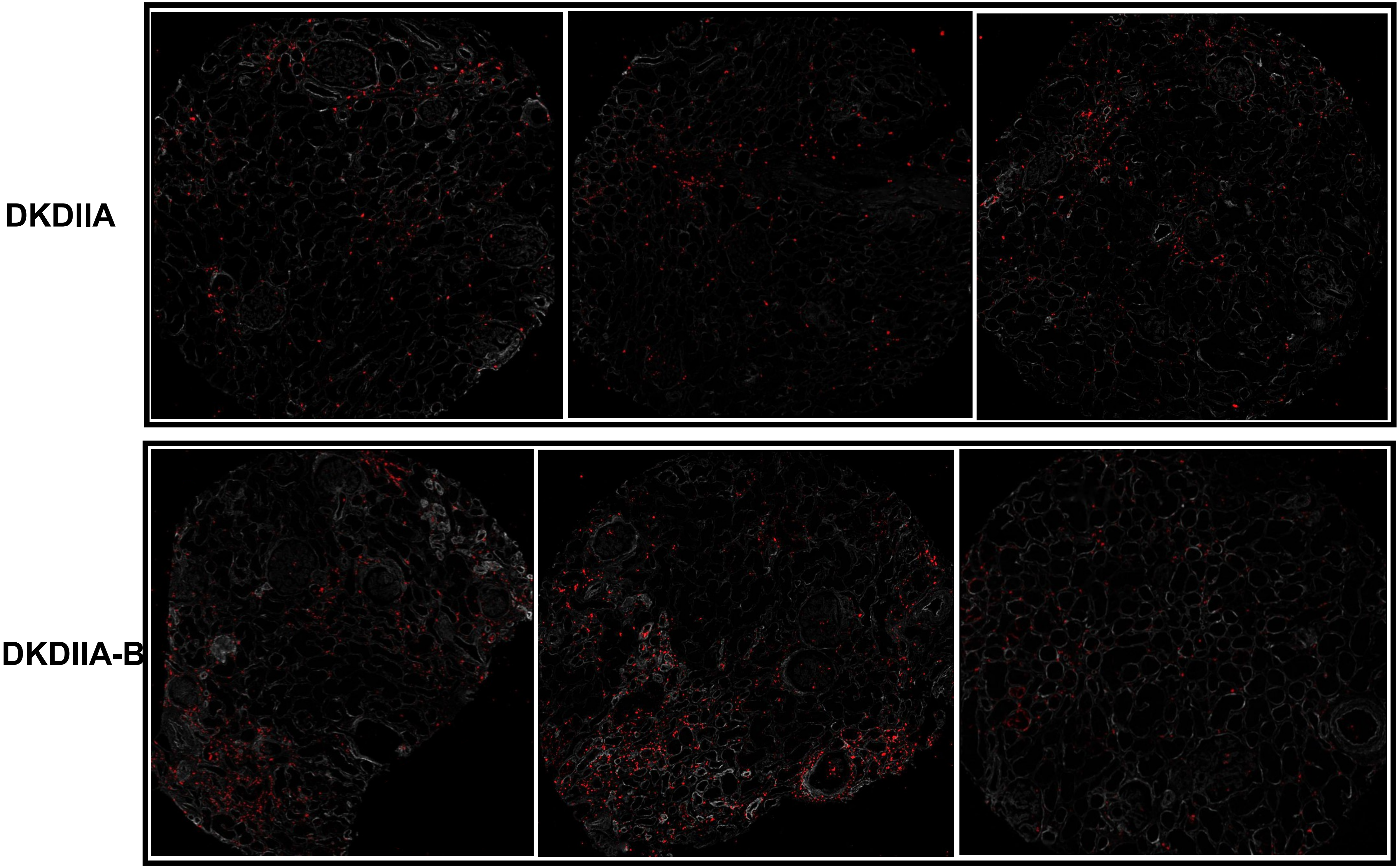

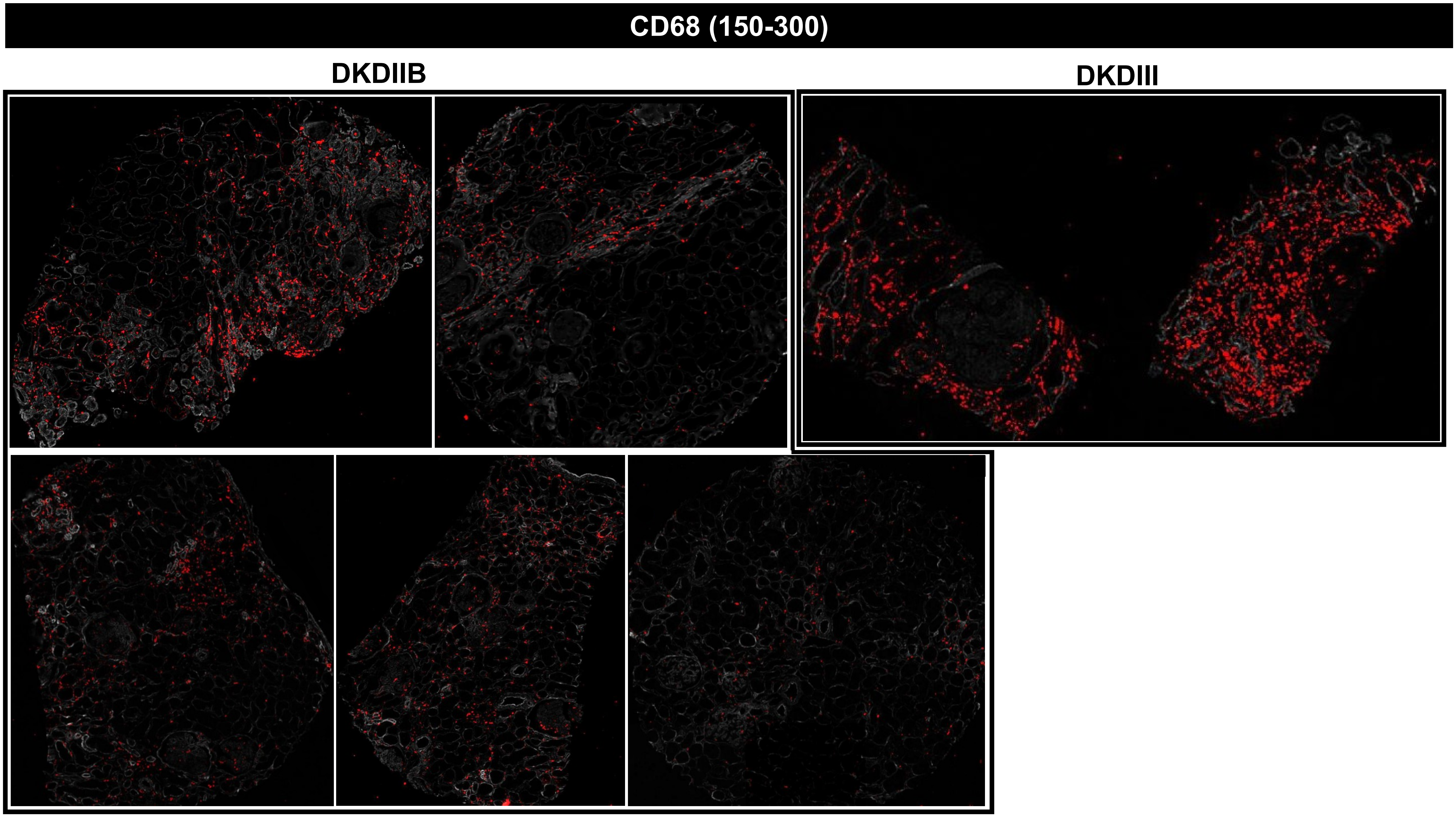
CD68 (150-300)
DKDIIB
DKDIII

CD68 (150-300)

### MEDULLA

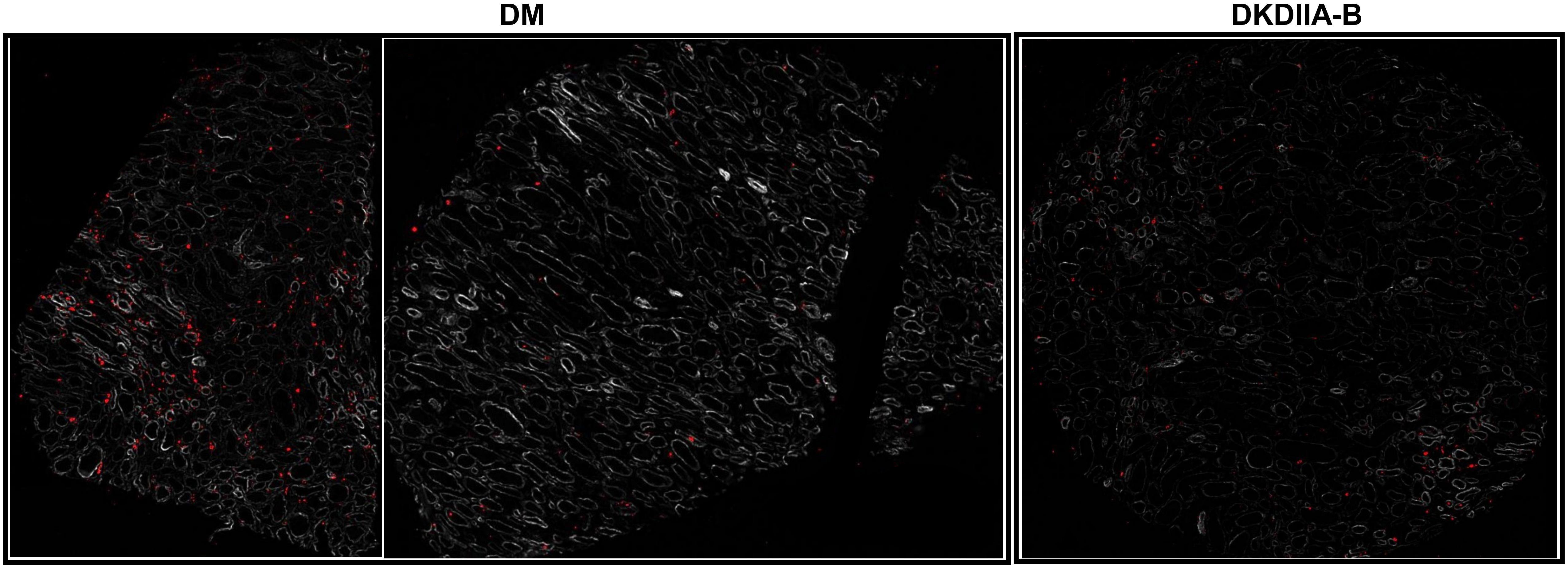

### COL4 (50-2000)

## DM

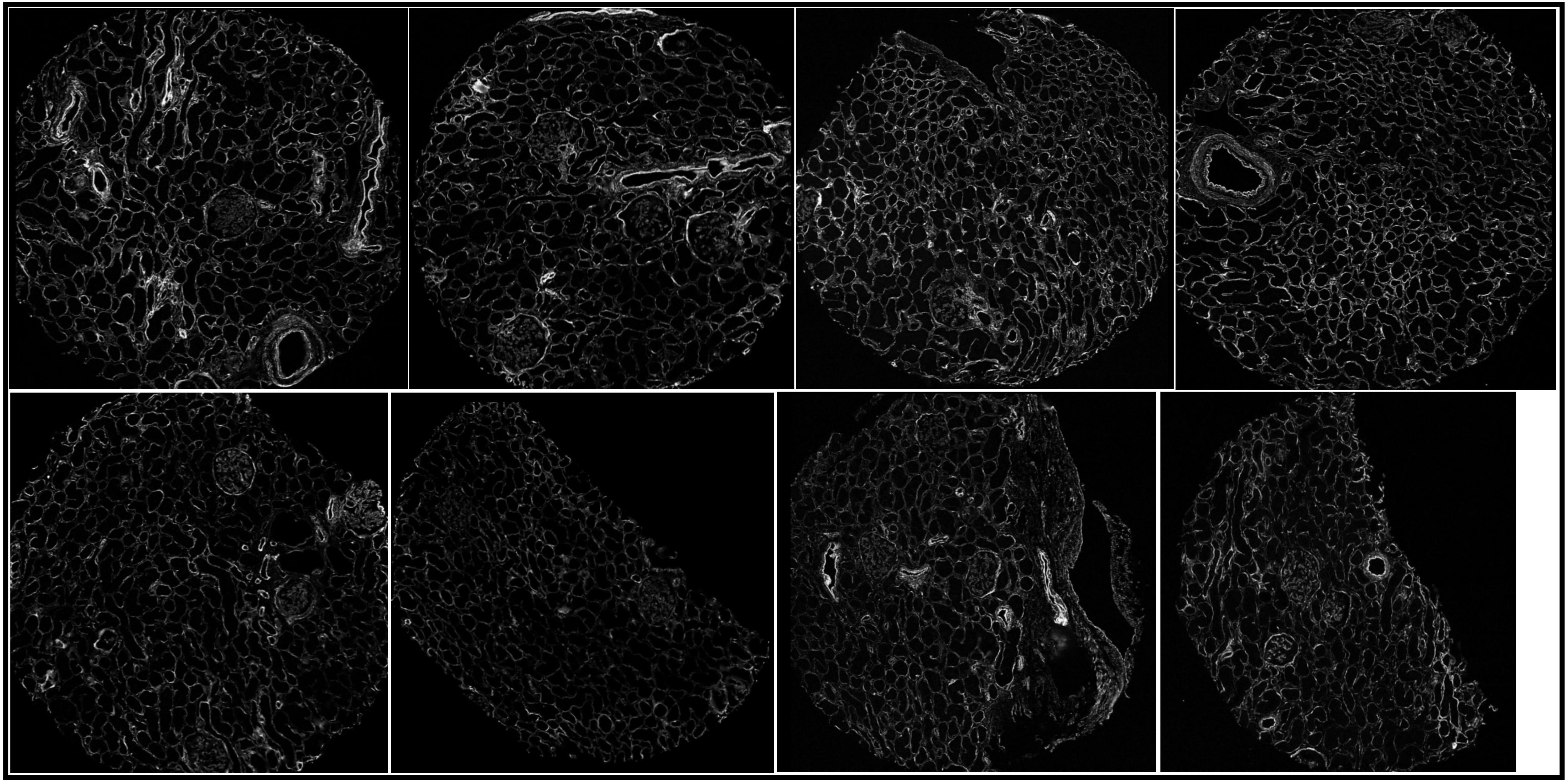

### COL4 (50-2000)

DKDIIA

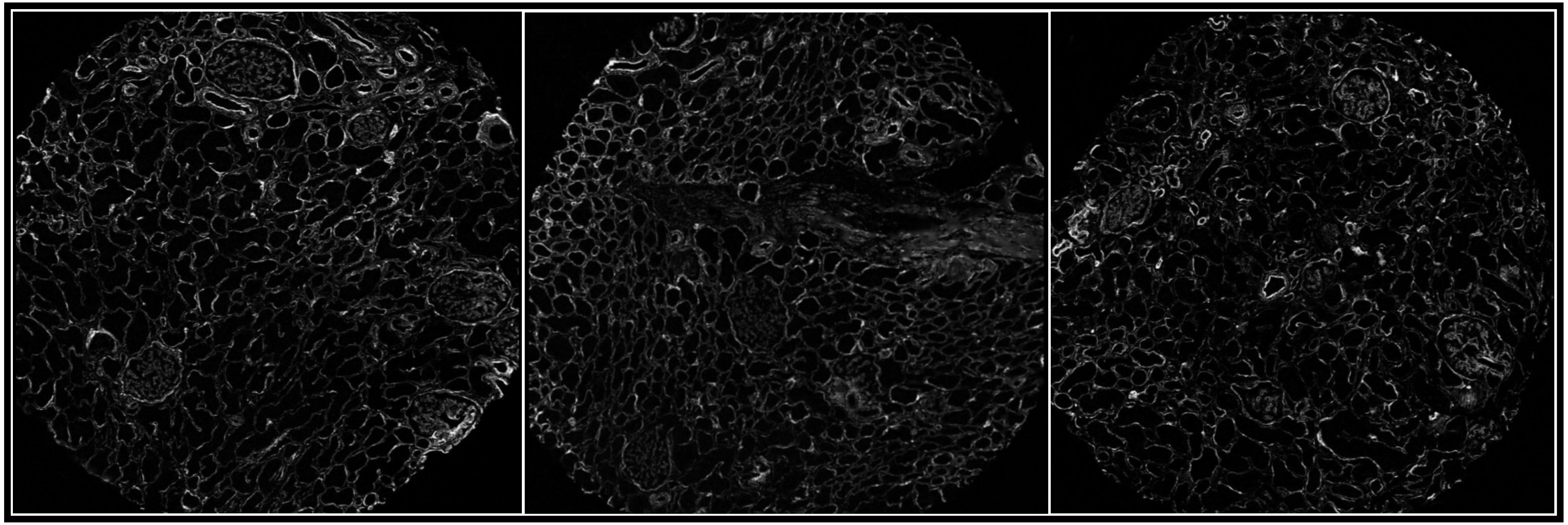

DKDIIA-B

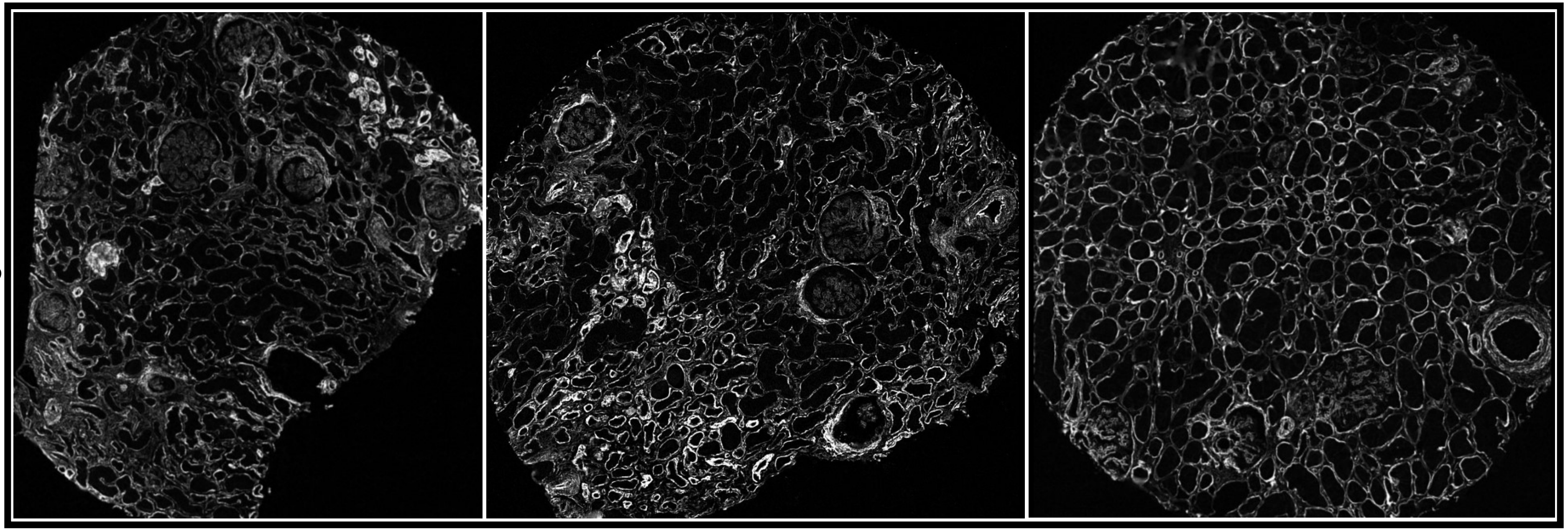

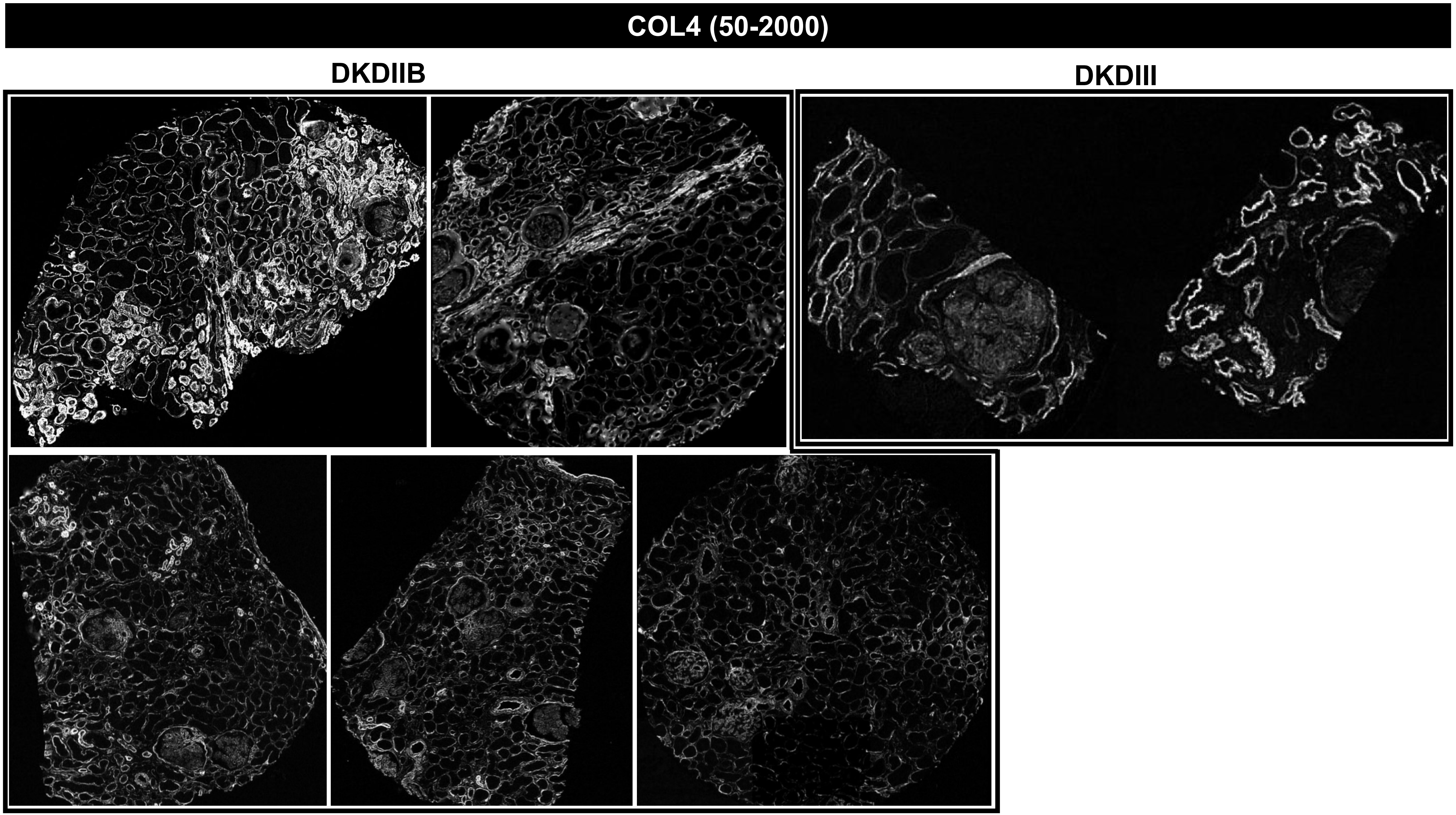

COL4 (50-2000)
DKDIIB
DKDIII

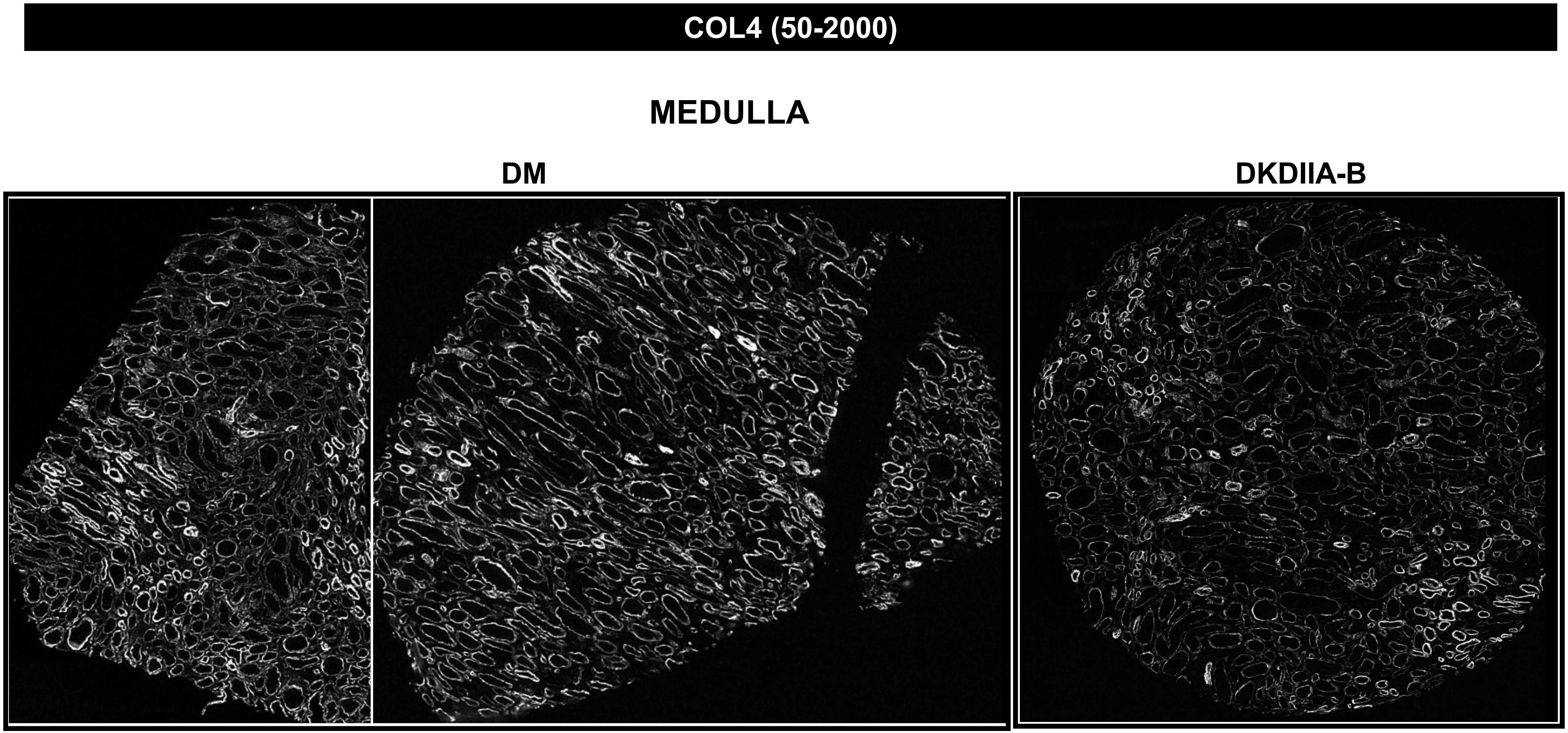
COL4 (50-2000)
MEDULLA
DM
DKDIIA-B

### CTNNB1 (70-500)

## DM

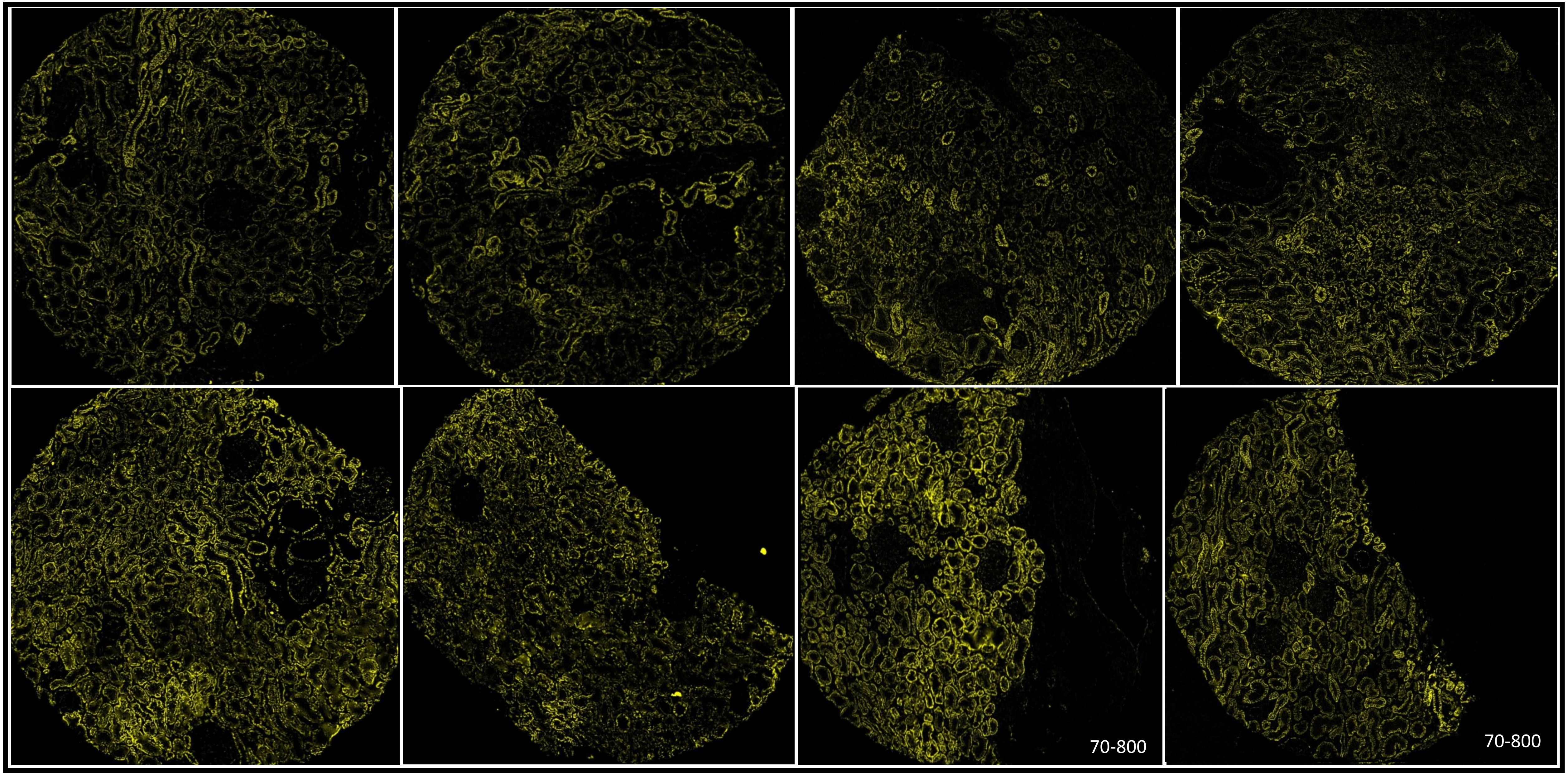

### CTNNB1 (70-500)

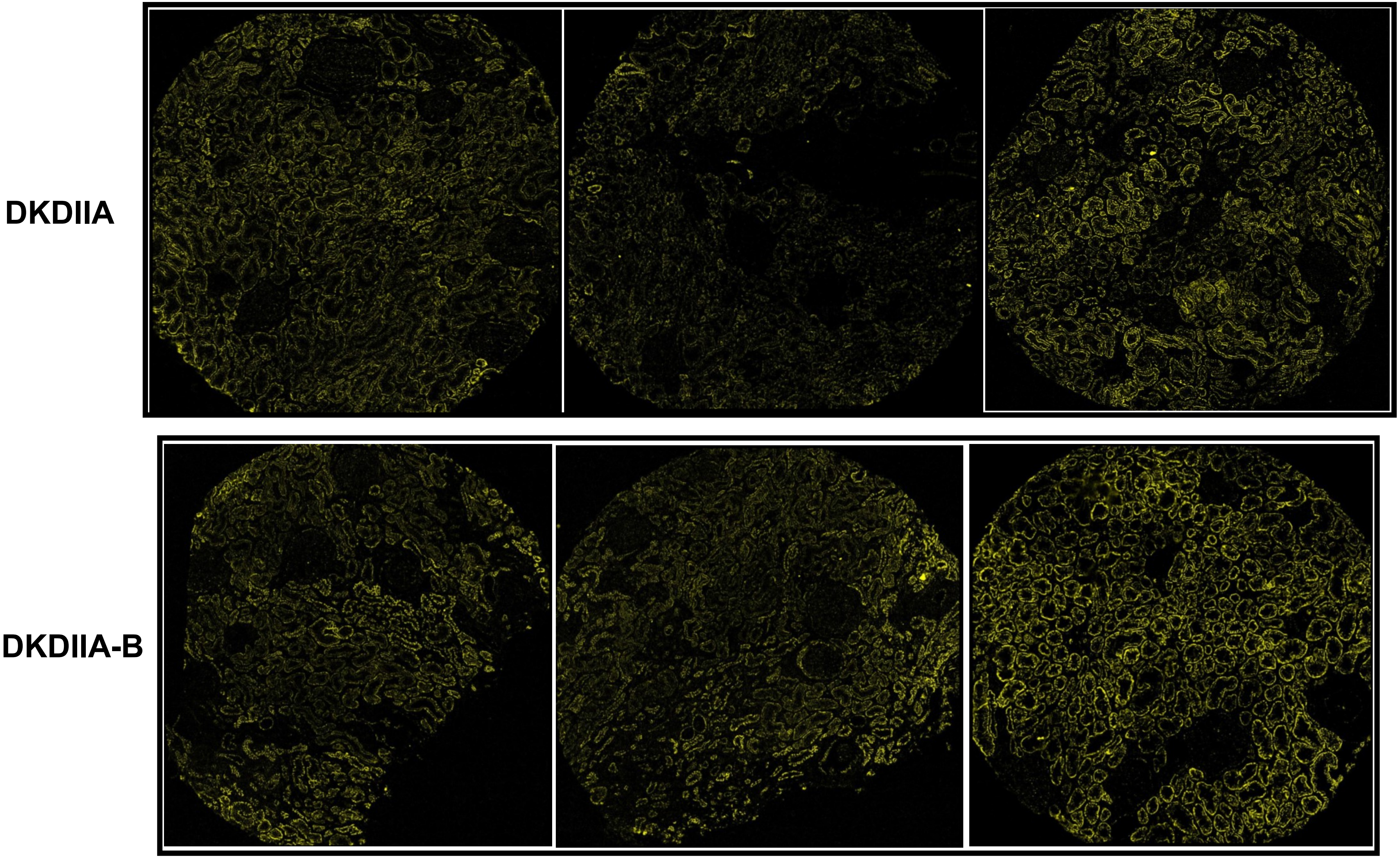

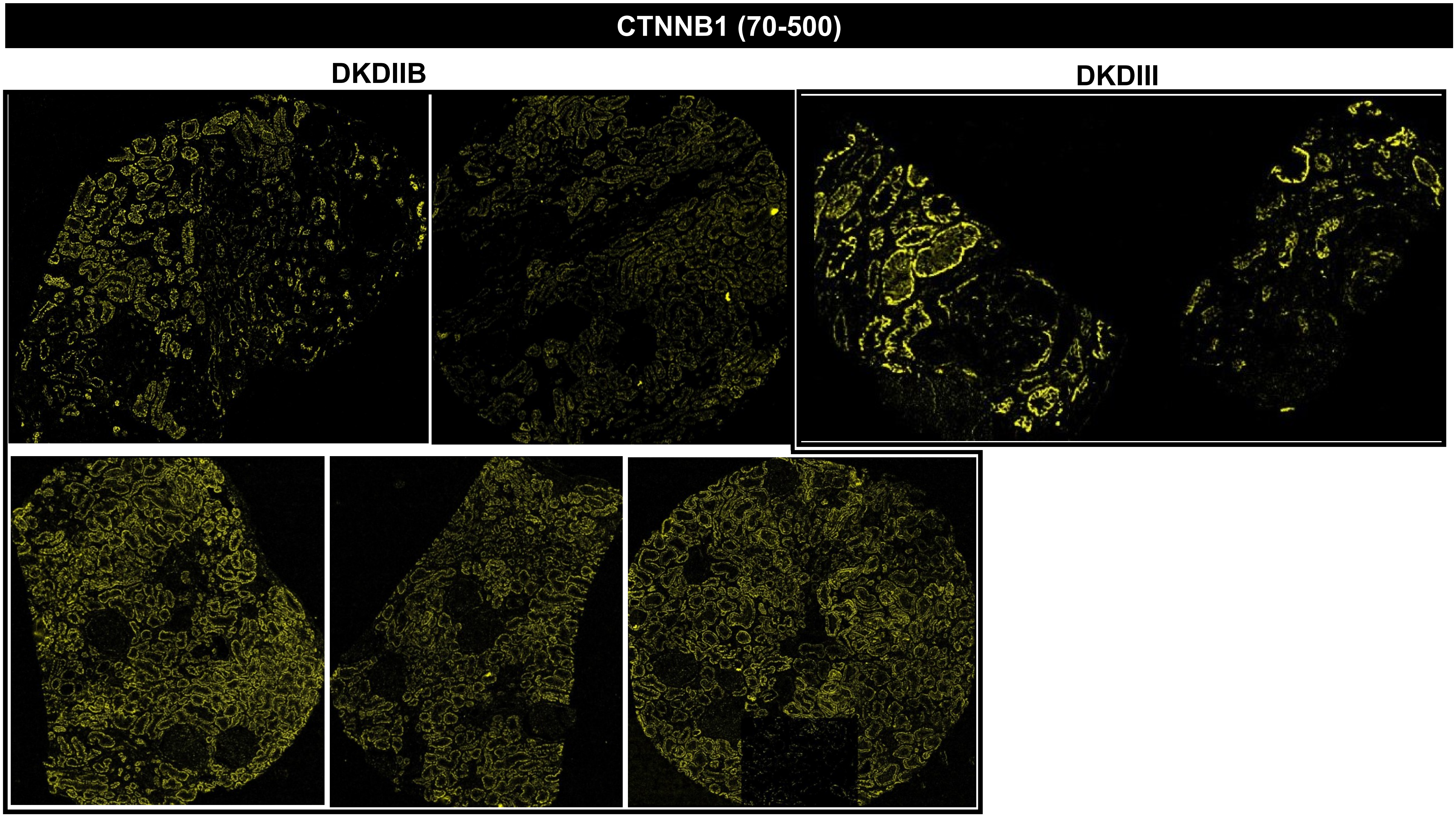
CTNNB1 (70-500)
DKDIIB
DKDIII

### CTNNB1 (70-500)

#### MEDULLA

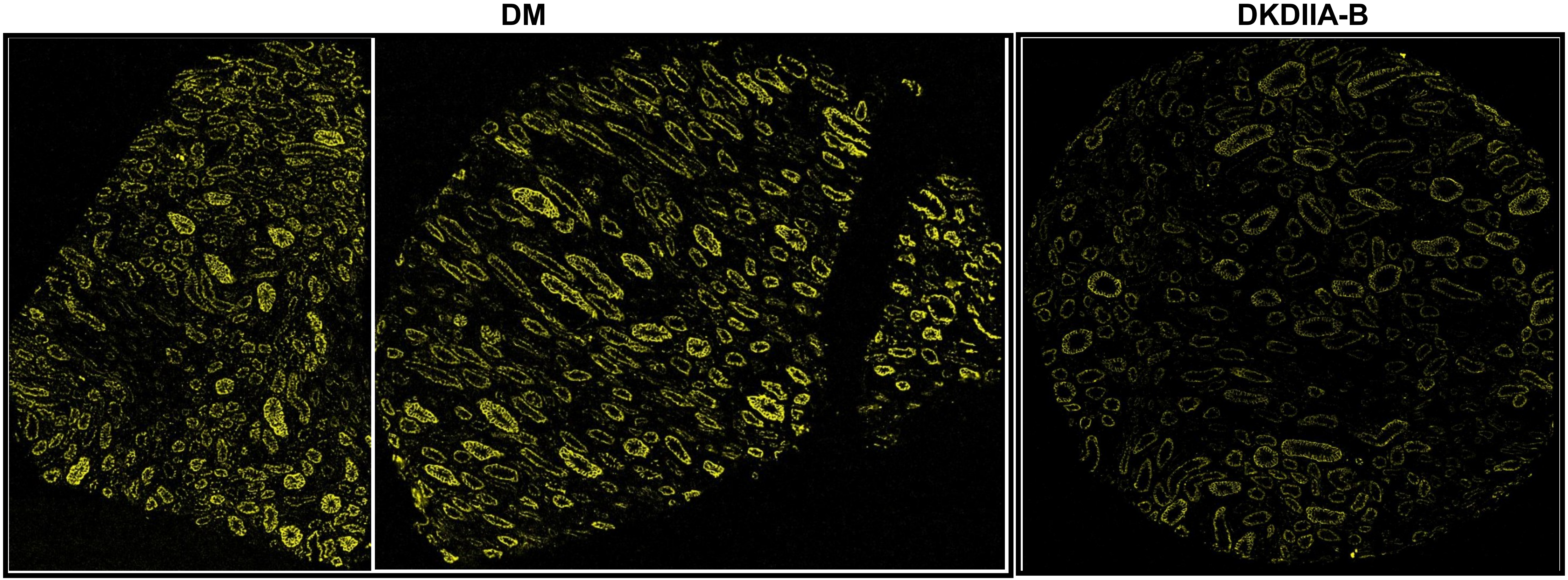

**CD183 or CXCR3 (200-2000)**

**DM**

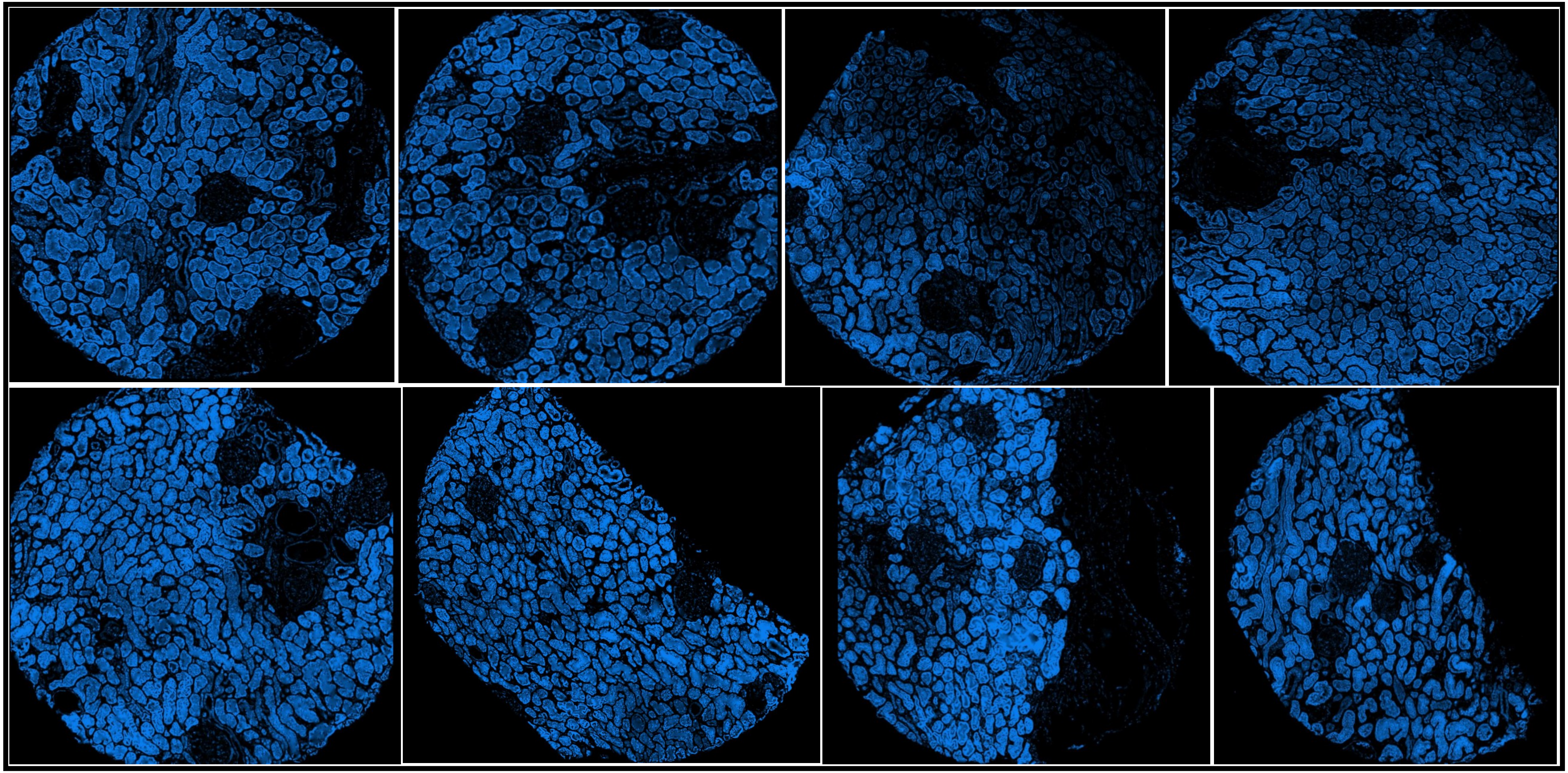

#### CD183 or CXCR3 (200-2000)

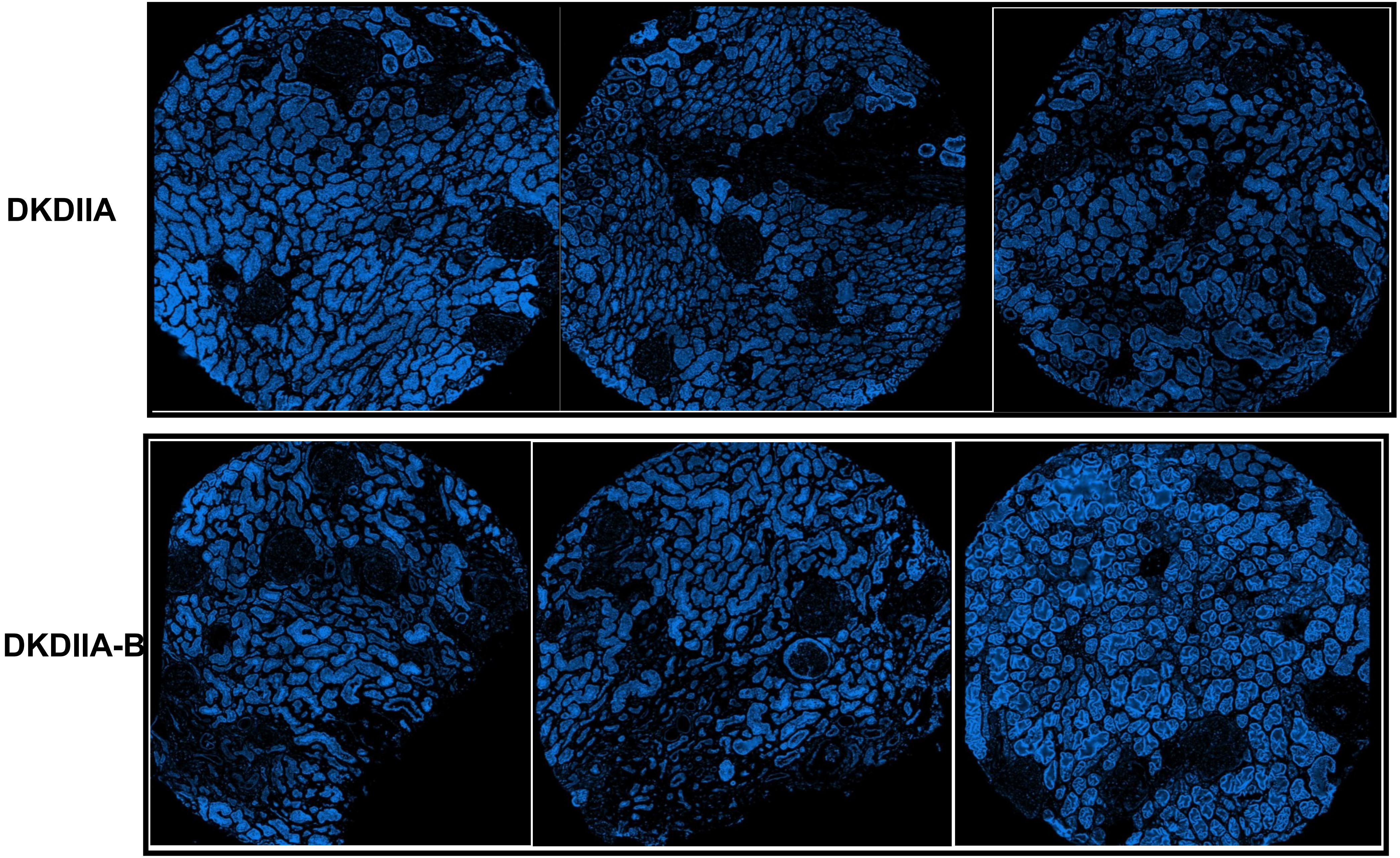

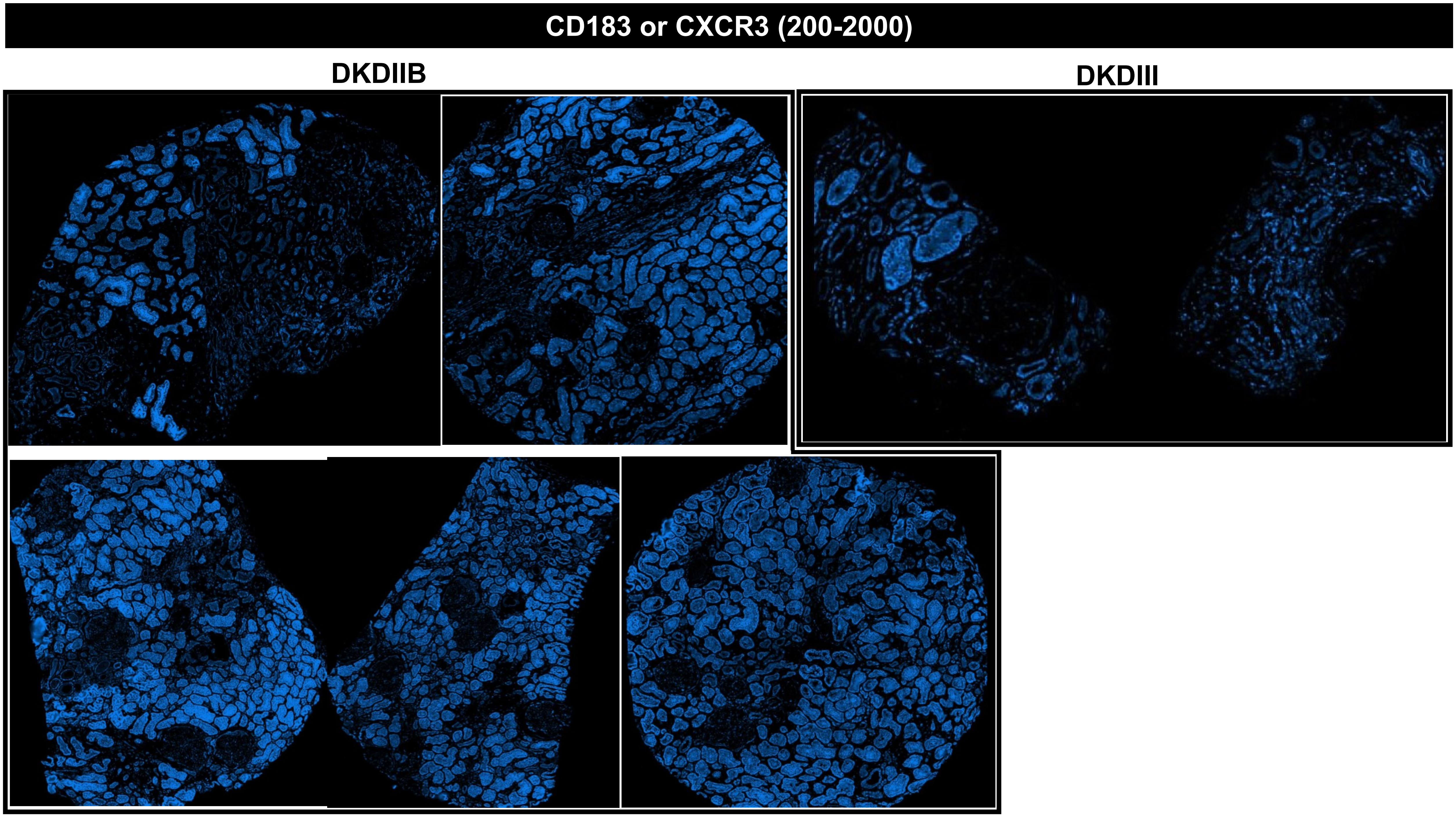
CD183 or CXCR3 (200-2000)
DKDIIB
DKDIII

CD183 or CXCR3 (200-2000)

### MEDULLA

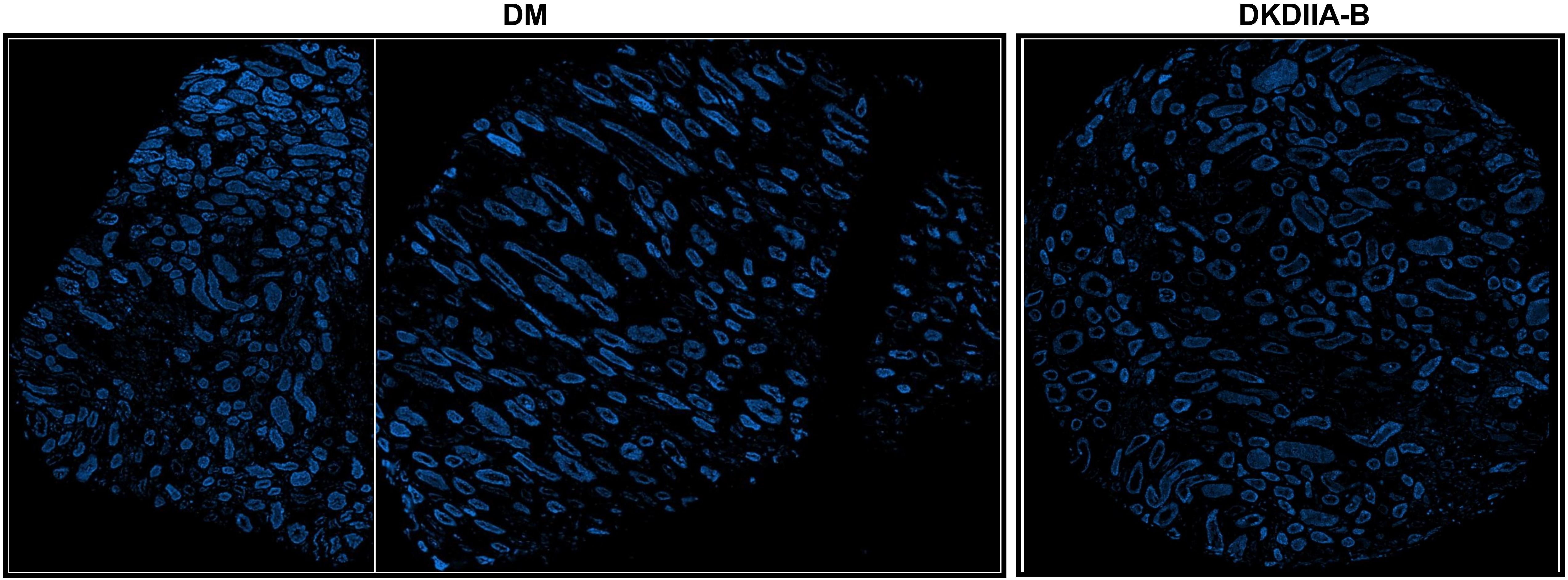

#### EpCAM (35/40-70)

**DM**

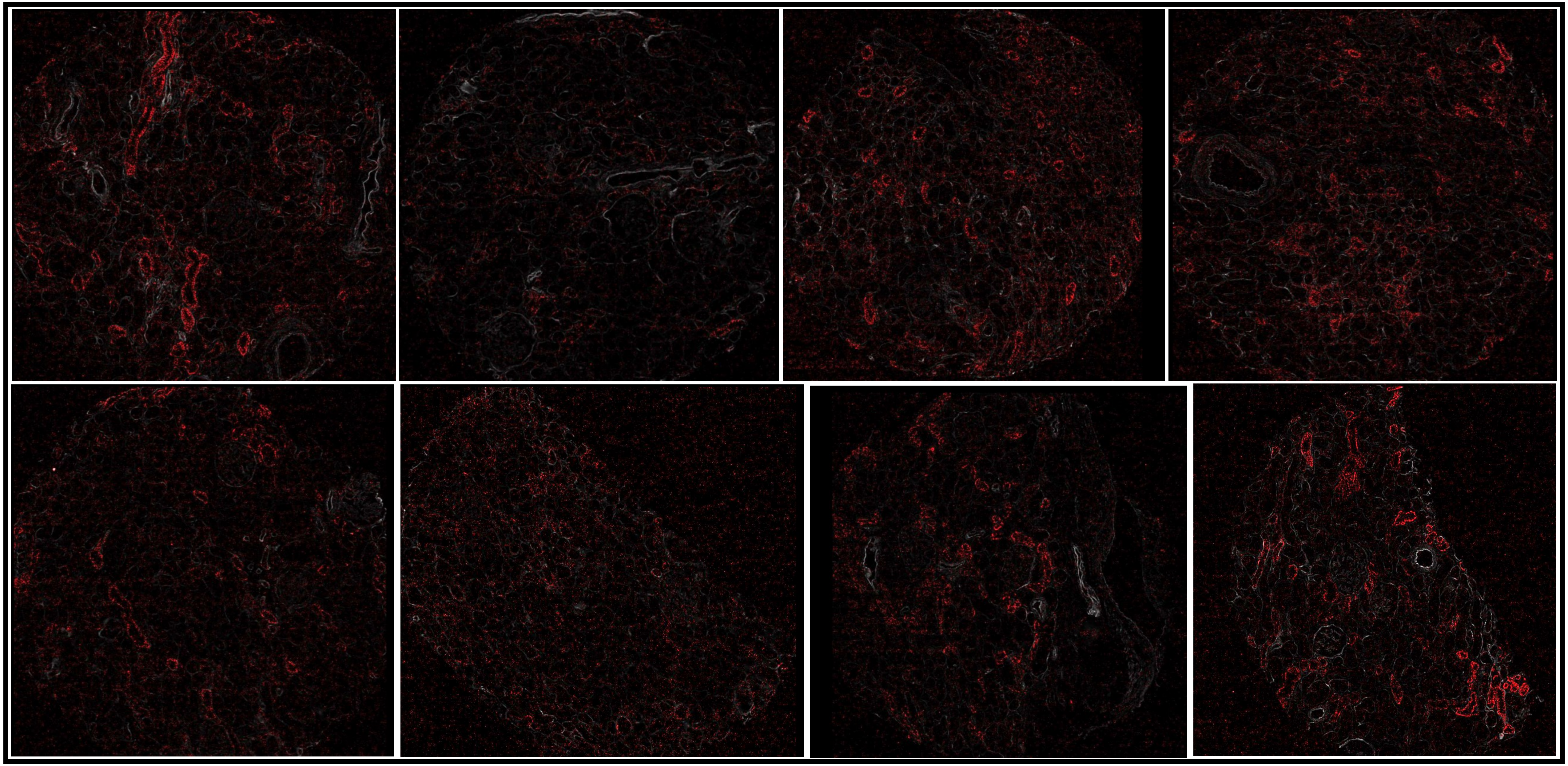

### EpCAM (35/40-70)

DKDIIA

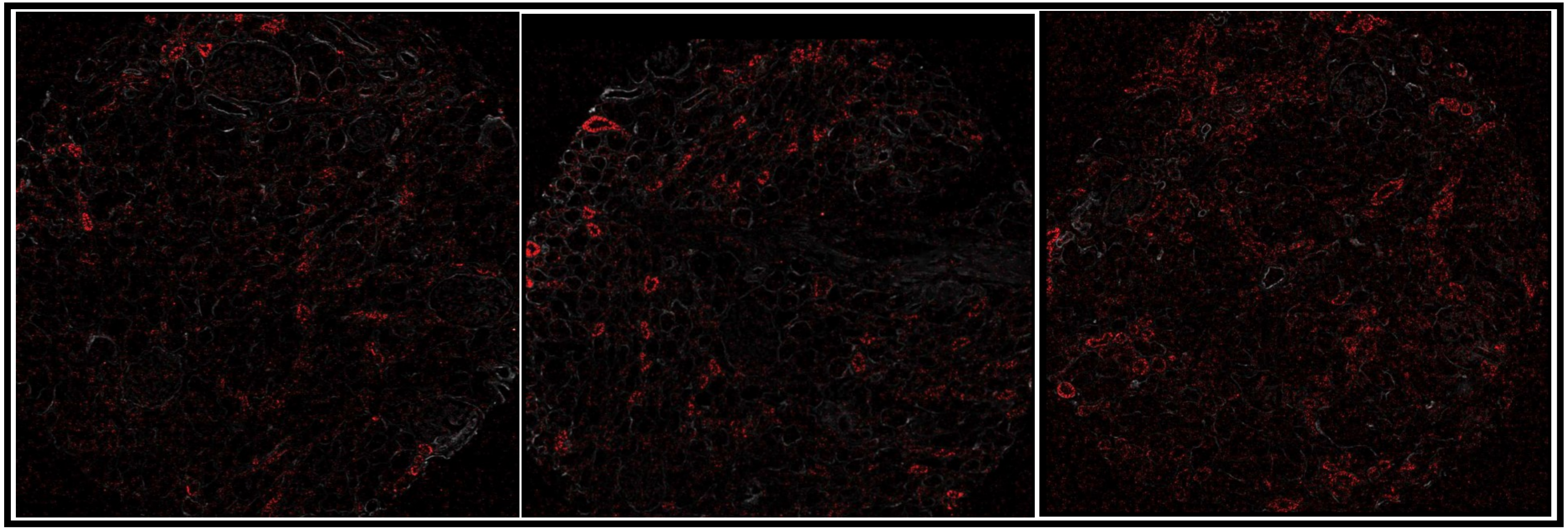

DKDIIA-B

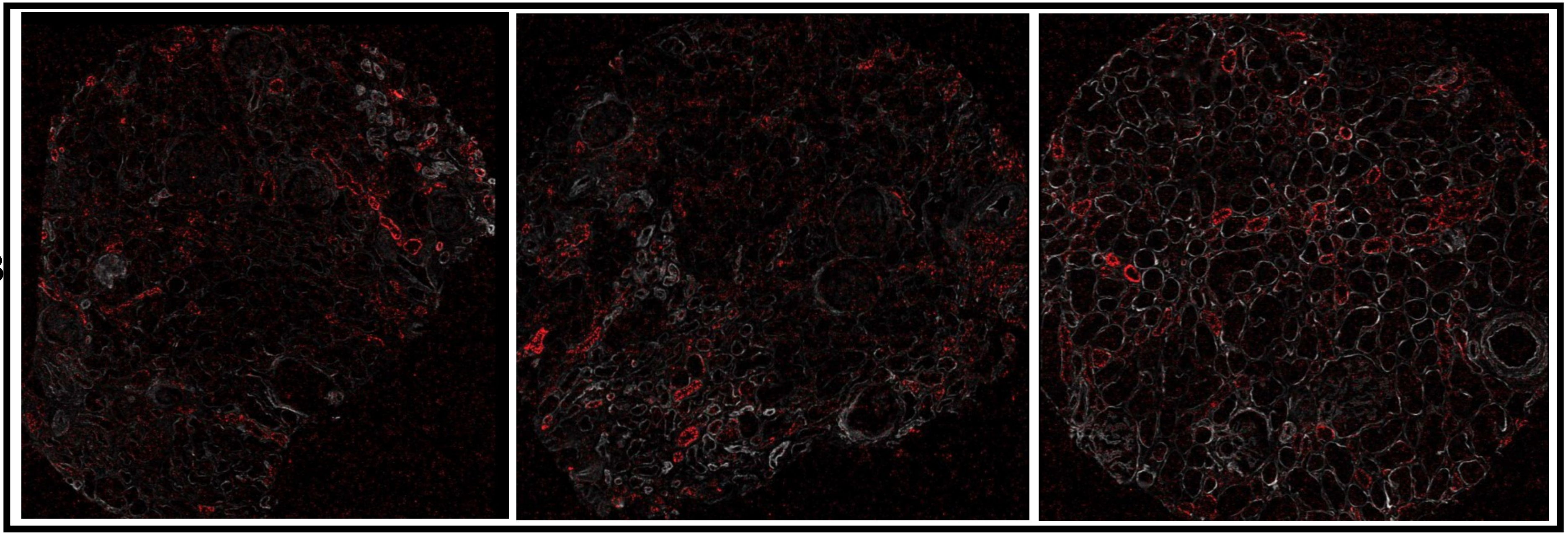

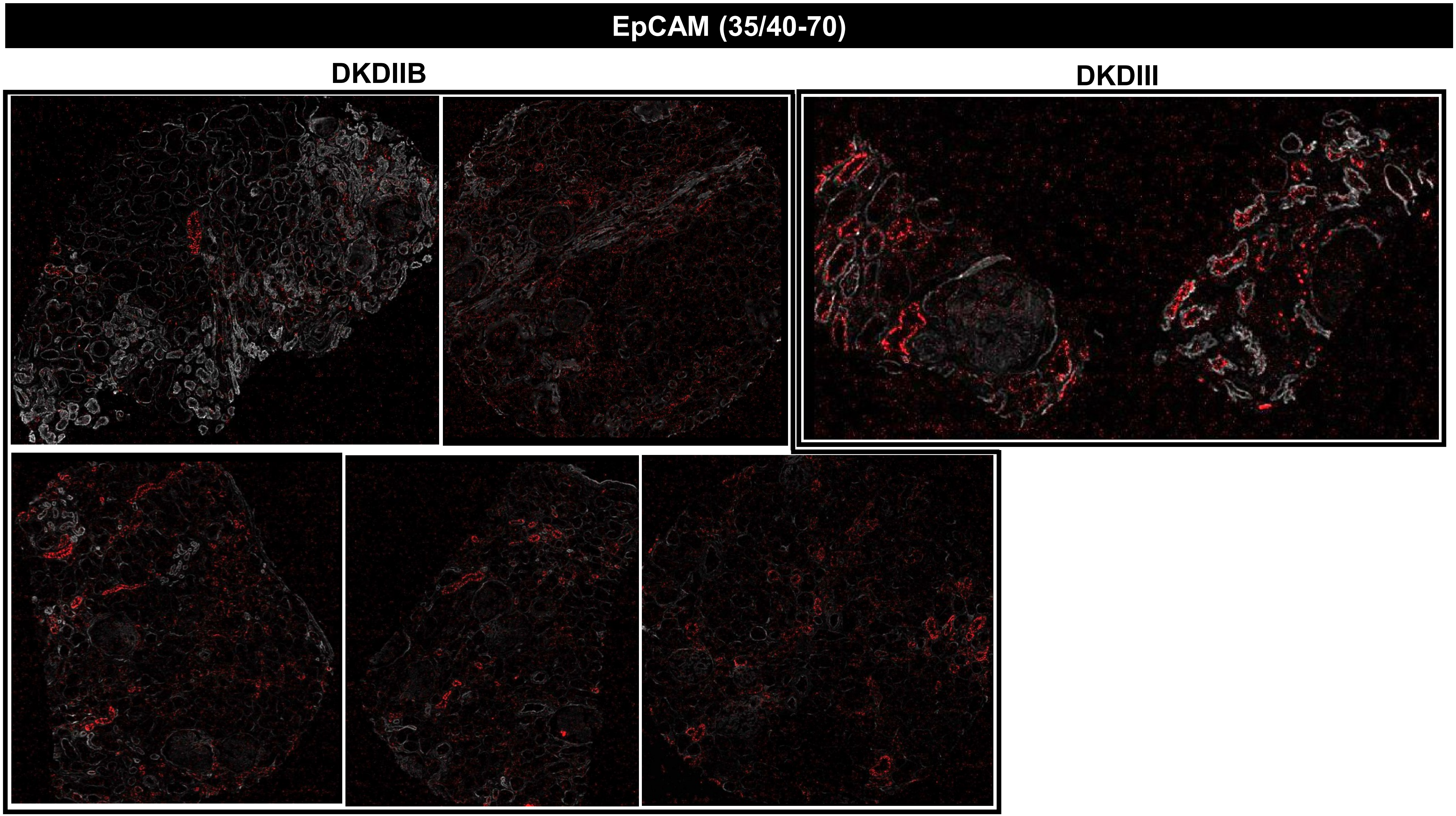
EpCAM (35/40-70)
DKDIIB
DKDIII

**EpCAM (35/40-70)**

### MEDULLA

**DM** **DKDIIA-B**

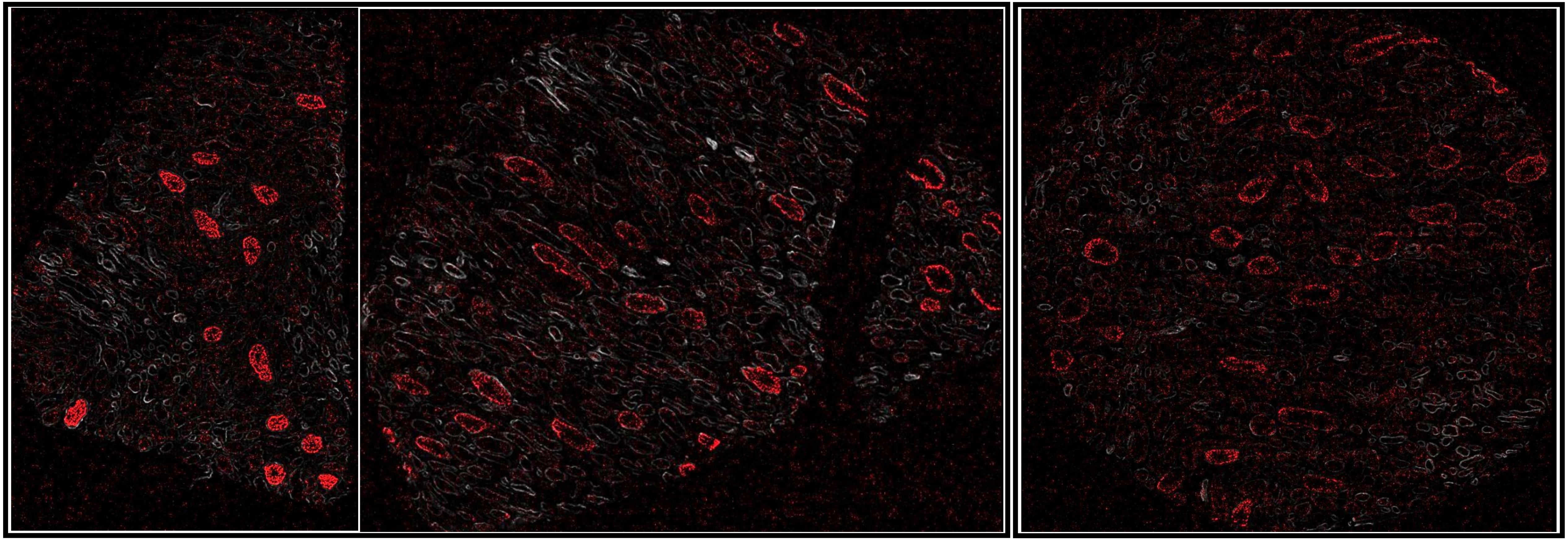

### HSPG (Perlecan, 40-800)

## DM

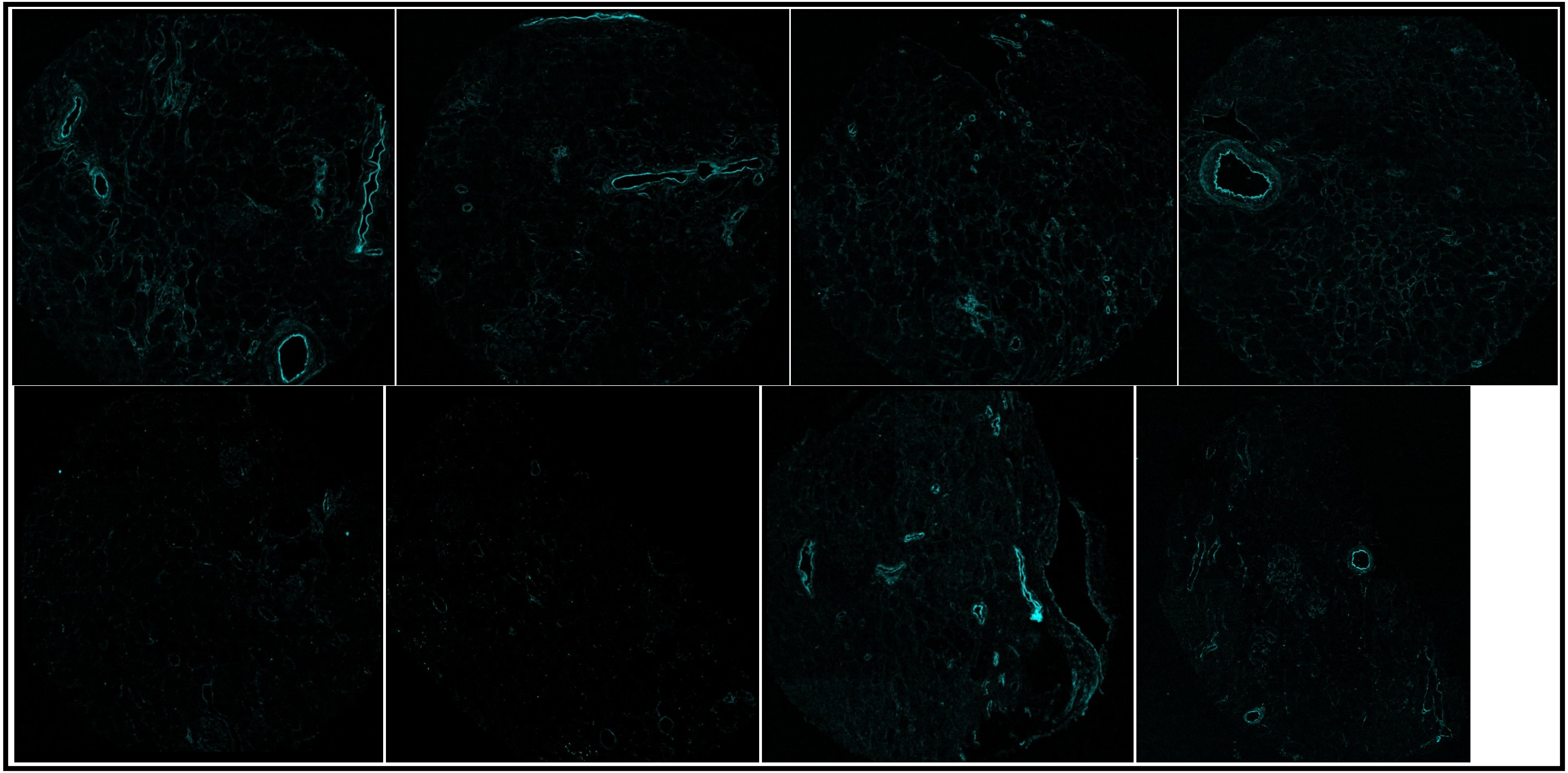

### HSPG (Perlecan, 40-800)

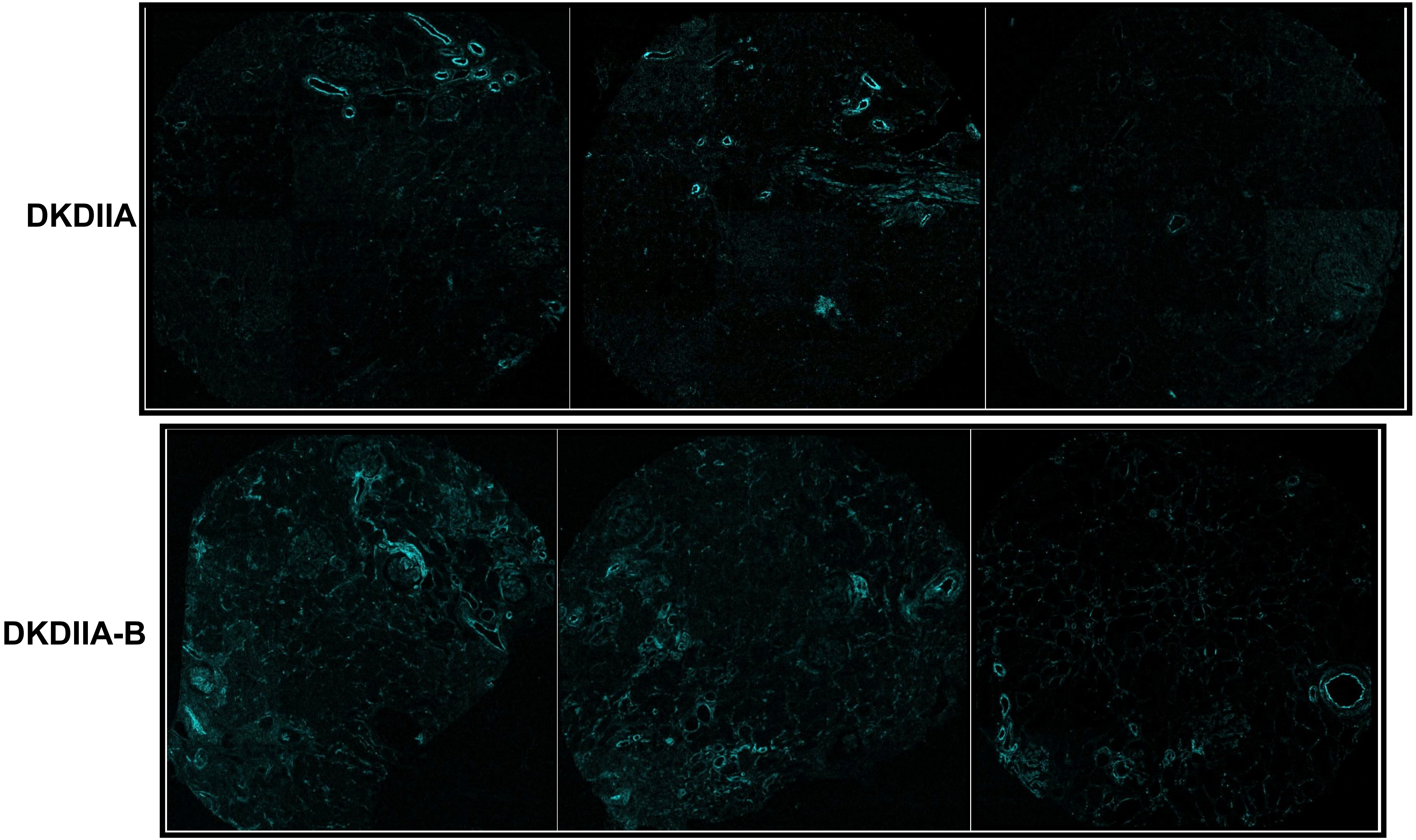

#### HSPG (Perlecan, 40-800)

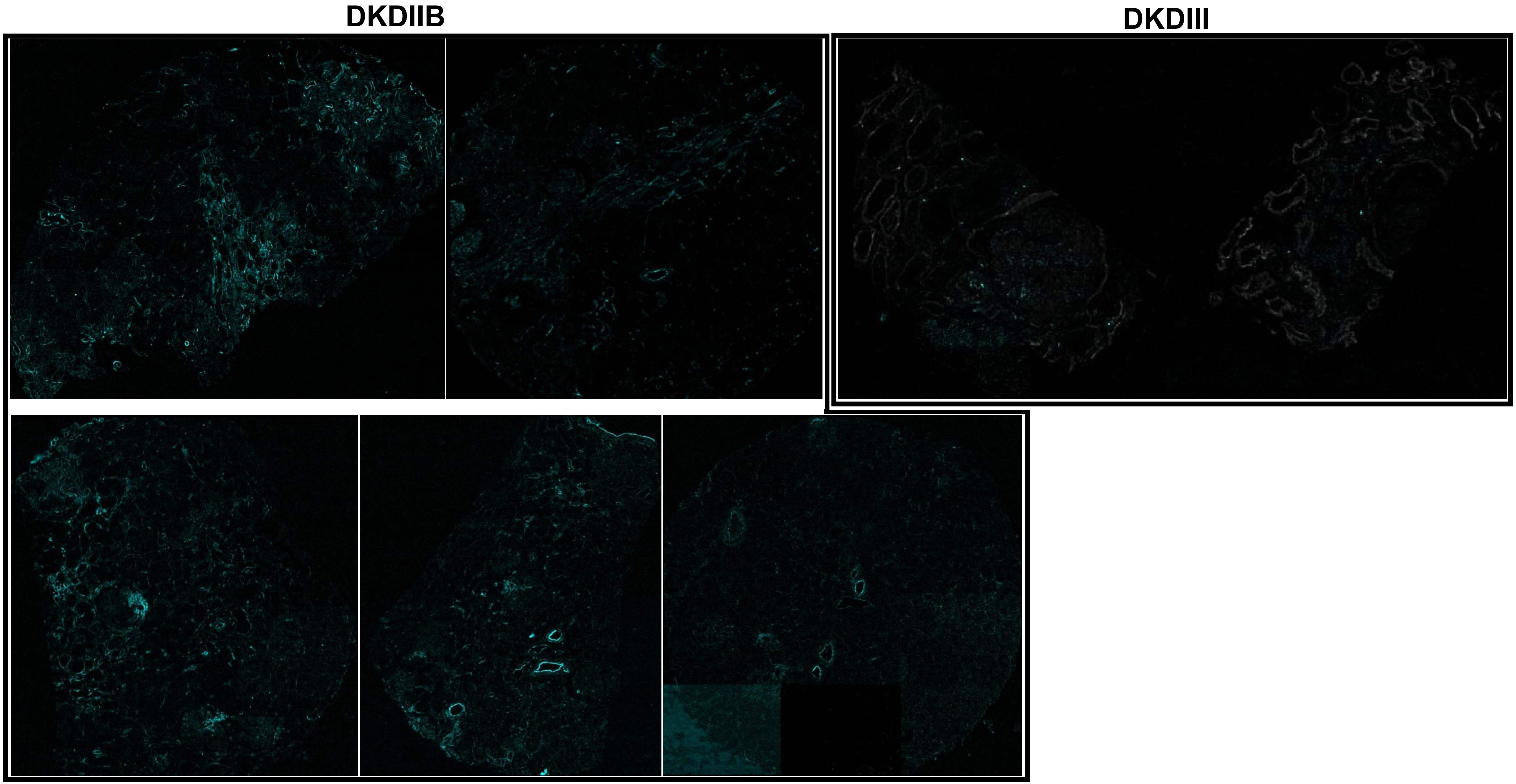

**HSPG (Perlecan, 40-800)**

### MEDULLA

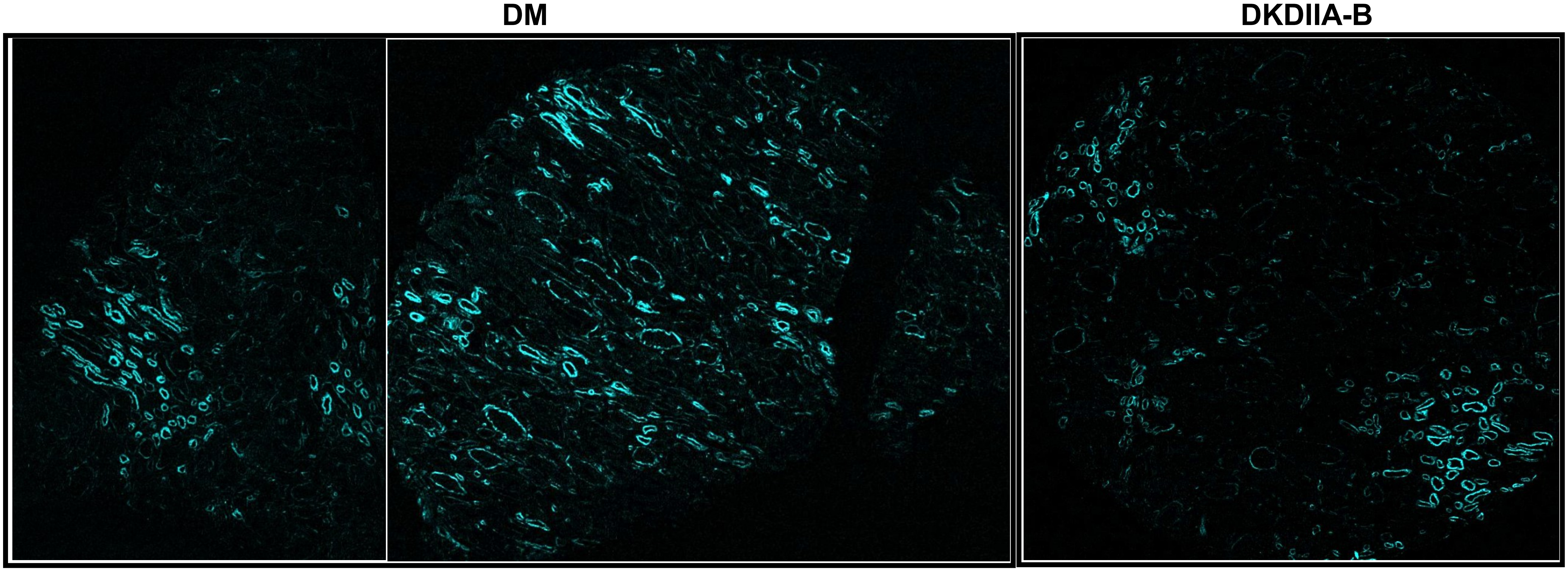

### HSPG (Perlecan, 40-400)

## DM

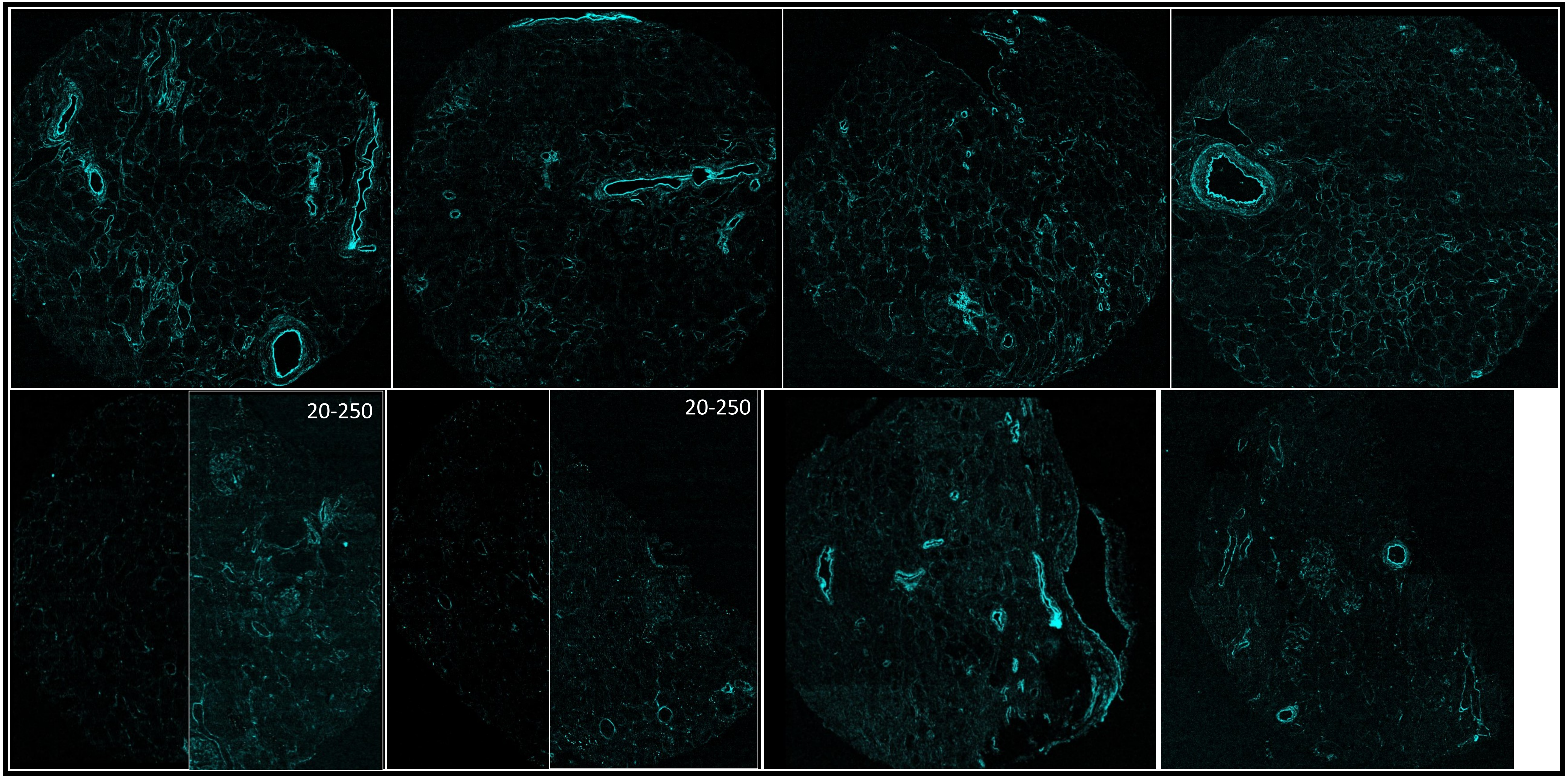

### HSPG (Perlecan, 40-400)

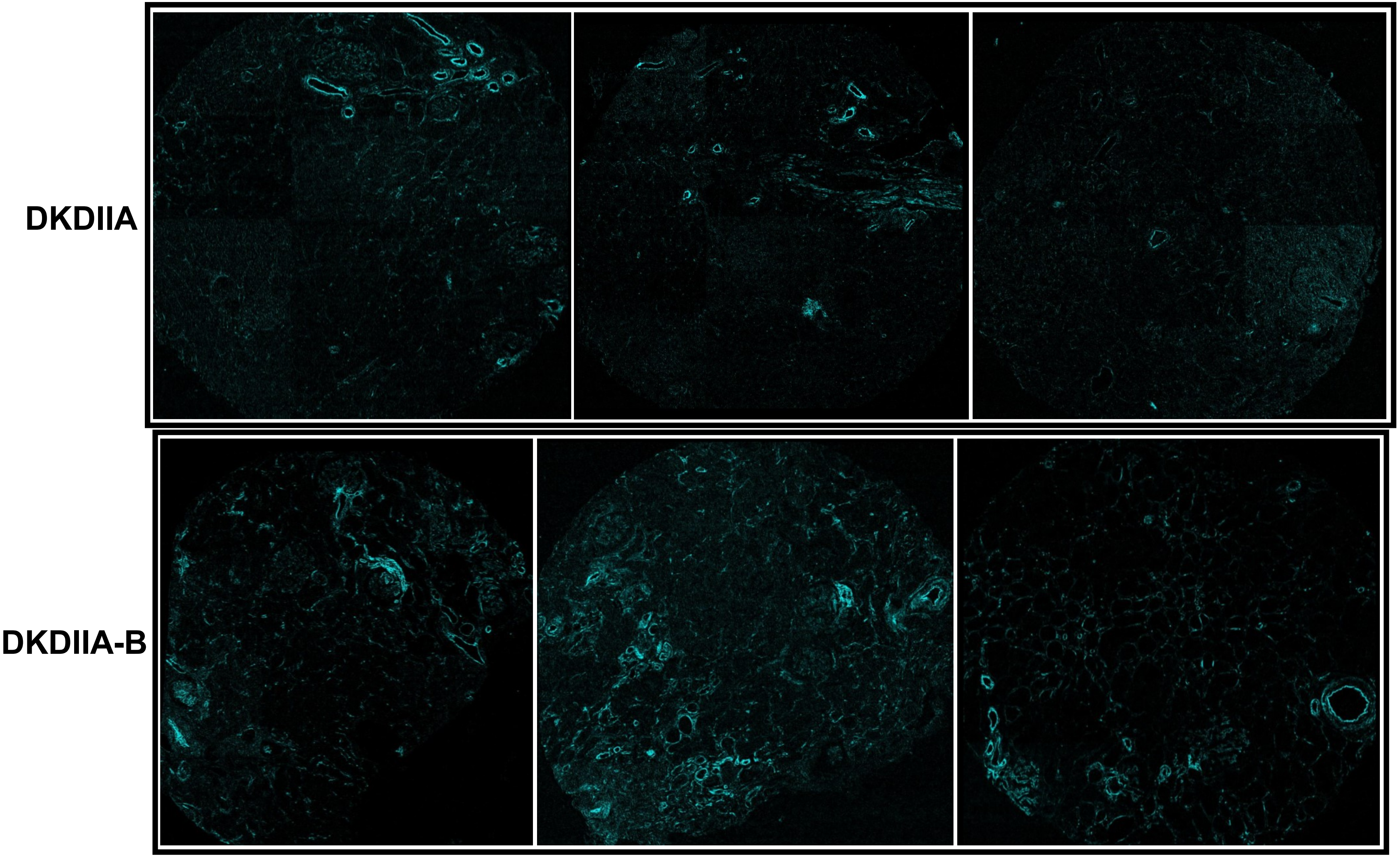

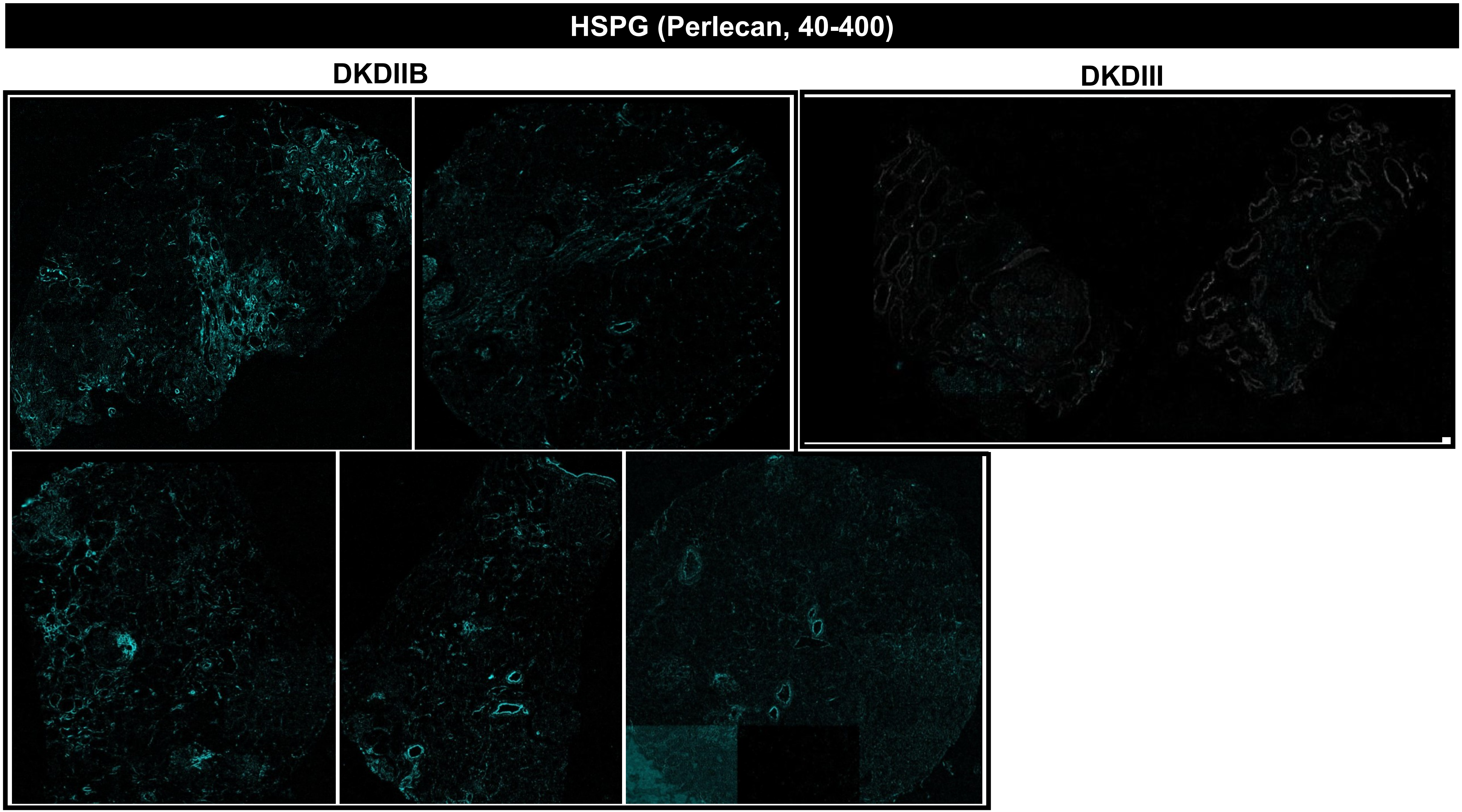
HSPG (Perlecan, 40-400)
DKDIIB
DKDIII

### HSPG (Perlecan, 40-400)

#### MEDULLA

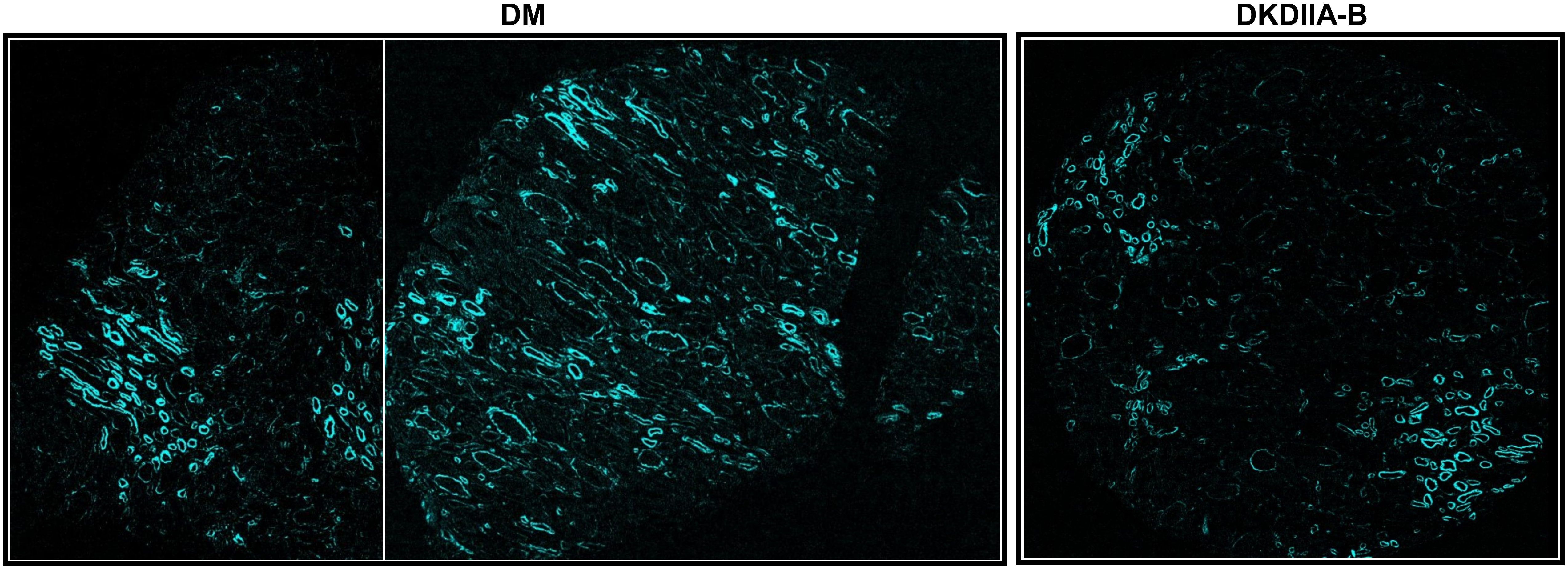

mtTFA (20-1000)

DM

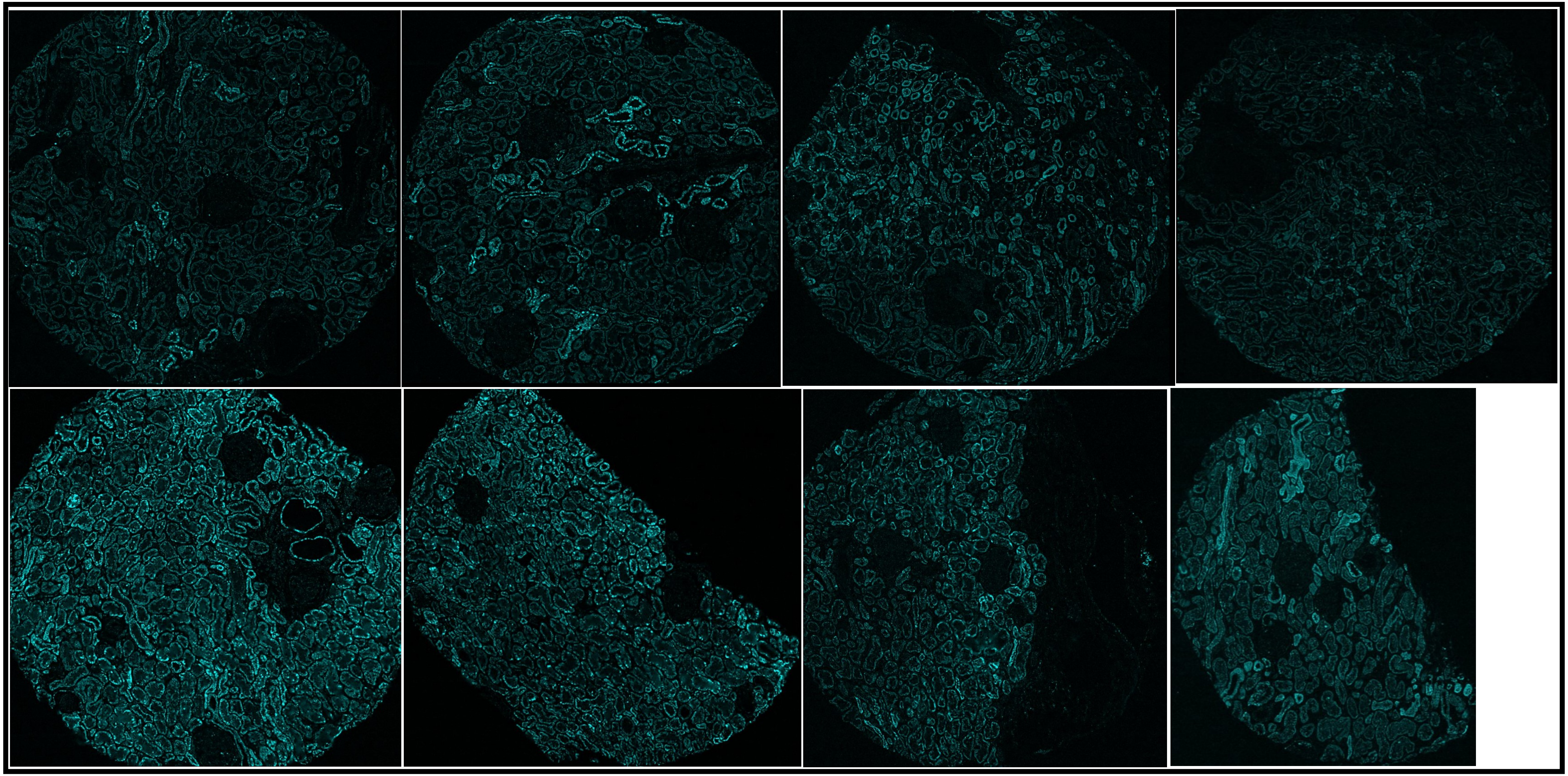

#### mtTFA (20-1000)

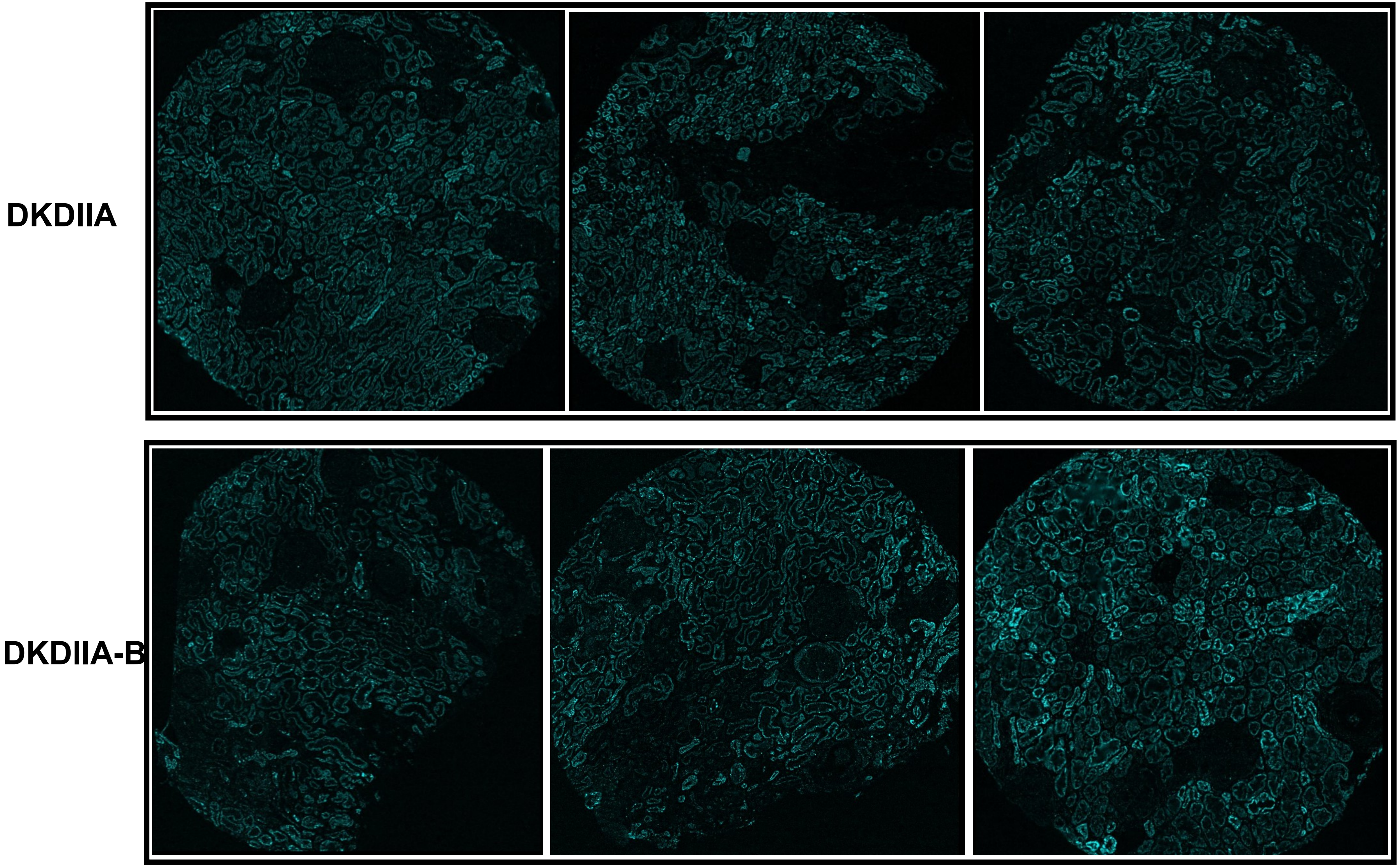

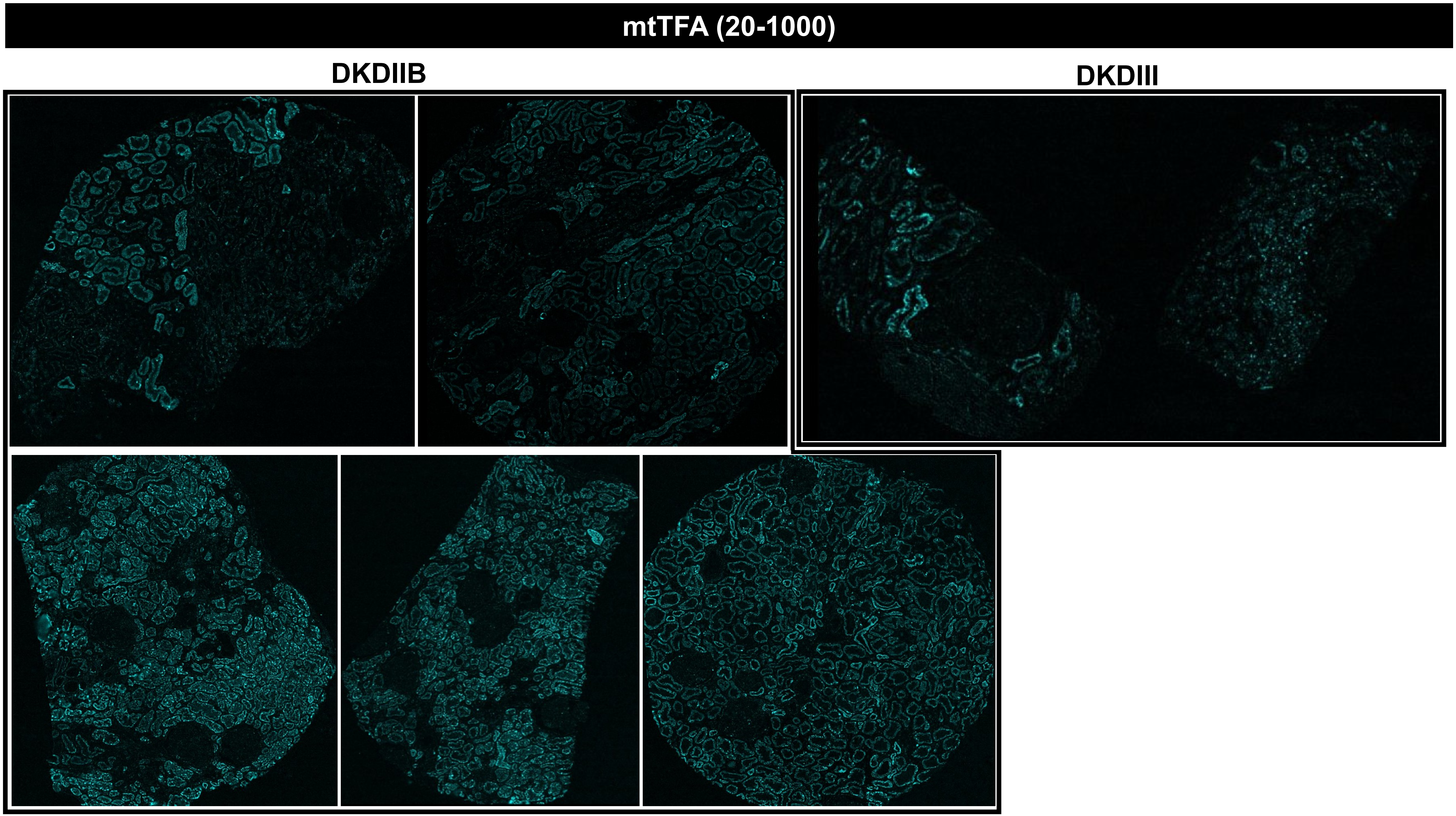
mtTFA (20-1000)
DKDIIB
DKDIII

**mtTFA (20-1000)**

#### MEDULLA

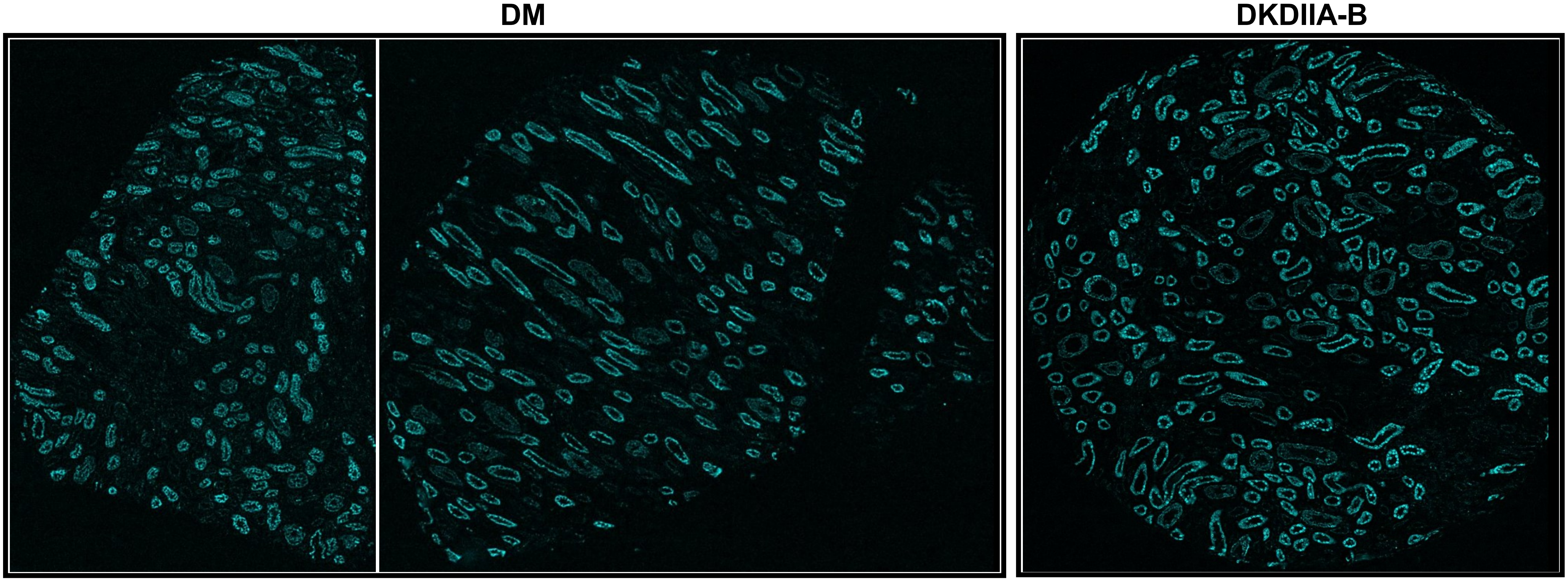

### Nestin (40-200)

## DM

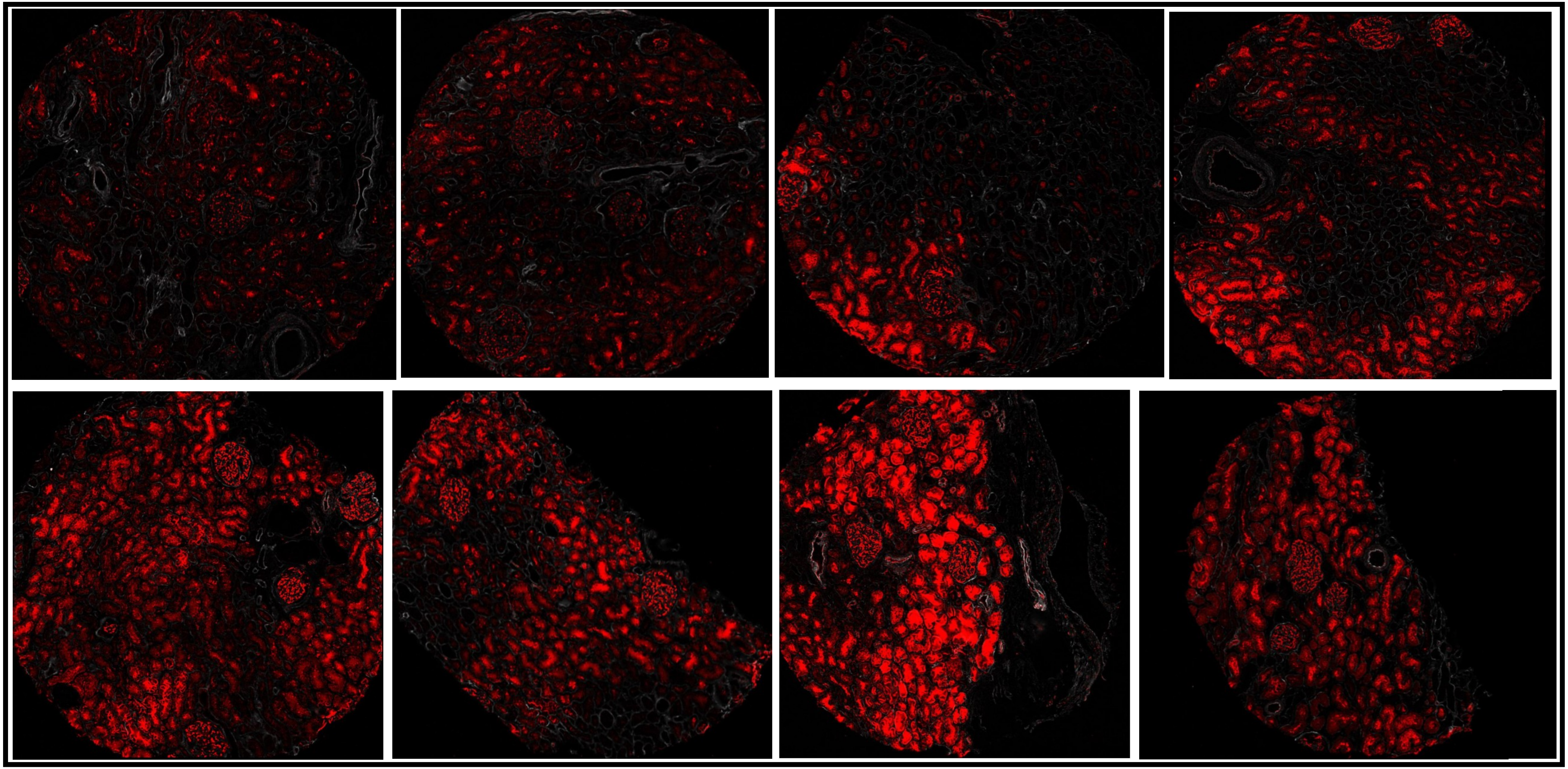

### Nestin (40-200)

DKDIIA

DKDIIA-B

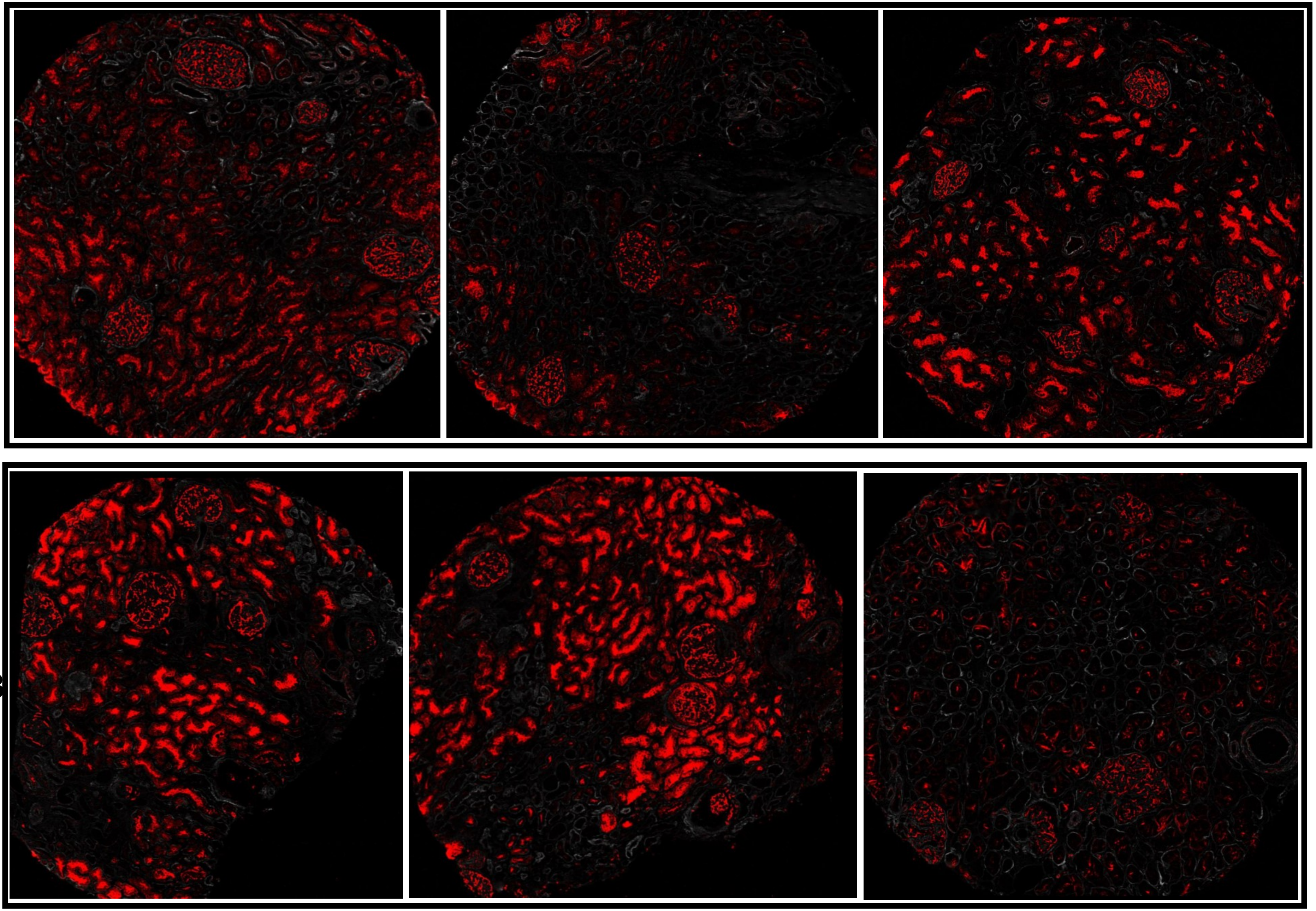

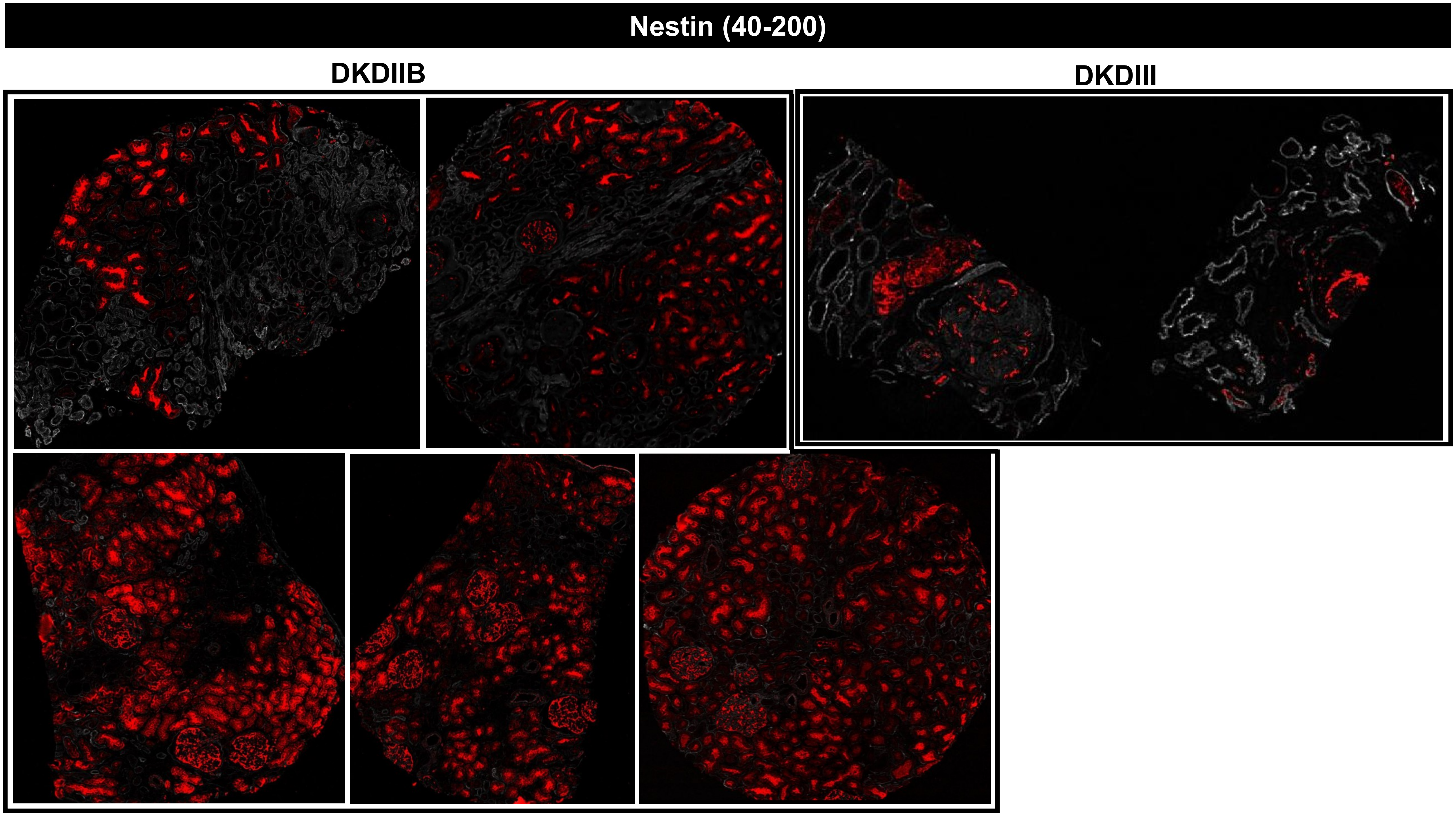
Nestin (40-200)
DKDIIB
DKDIII

Nestin (40-200)

### MEDULLA

DM

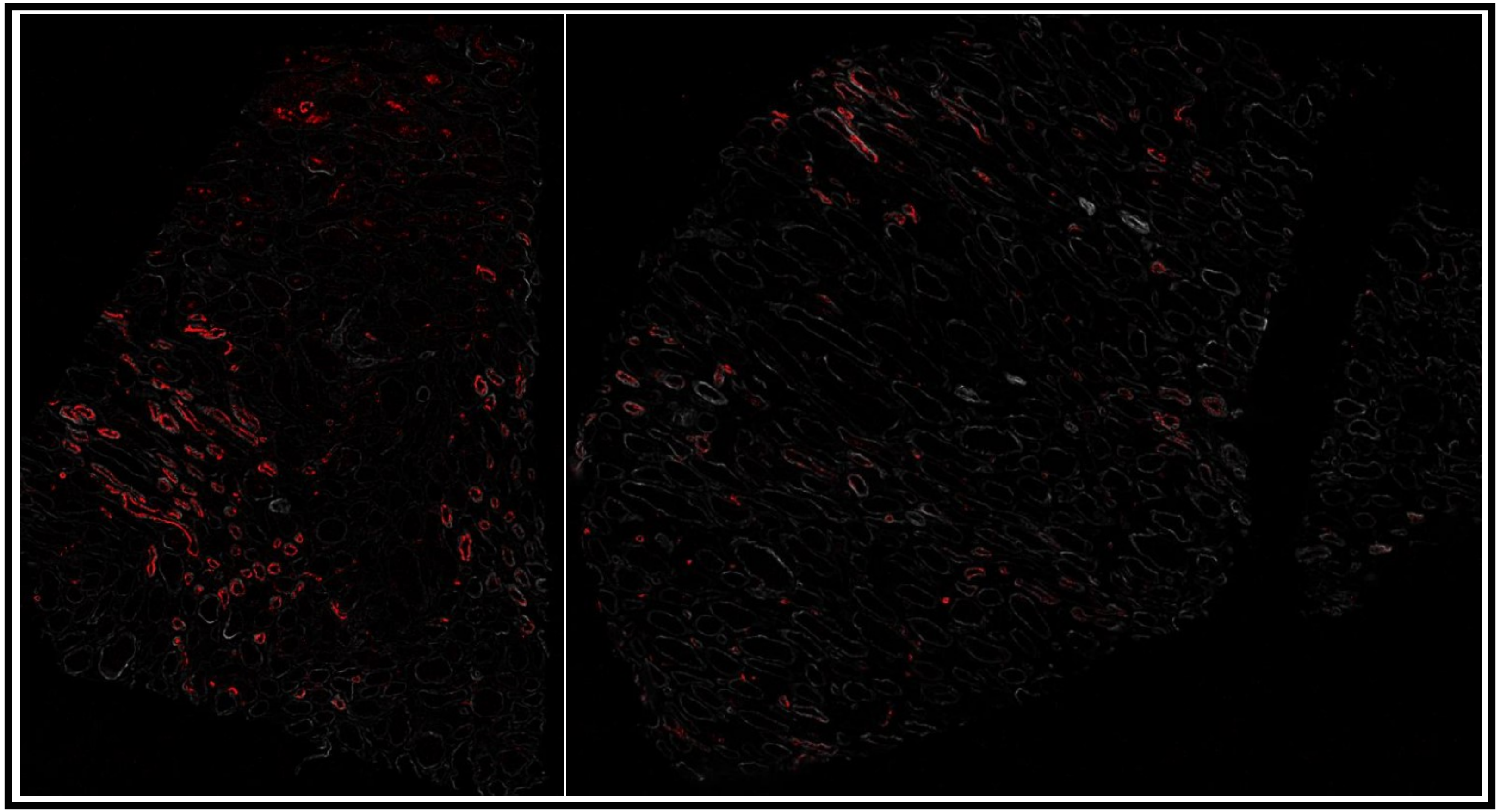

DKDIIA-B

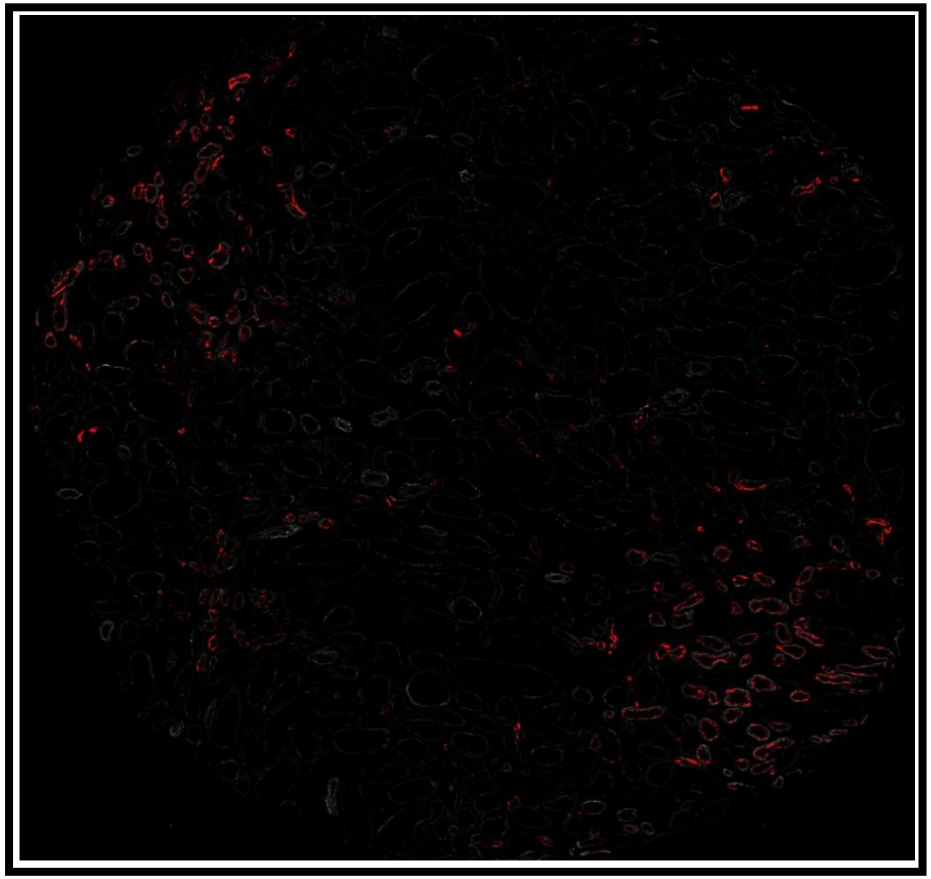

### Nestin (50-500)

## DM

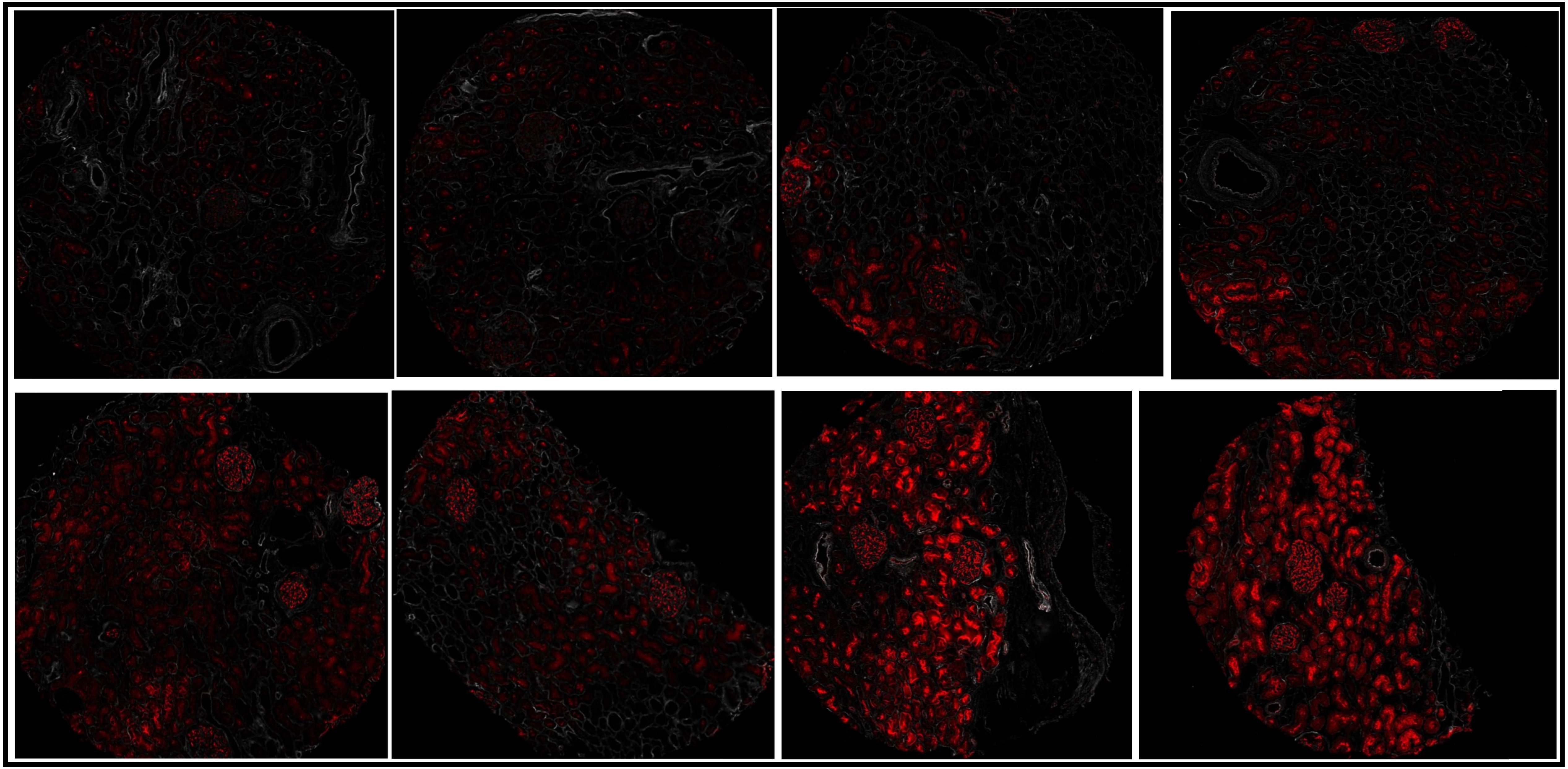

### Nestin (50-500)

DKDIIA

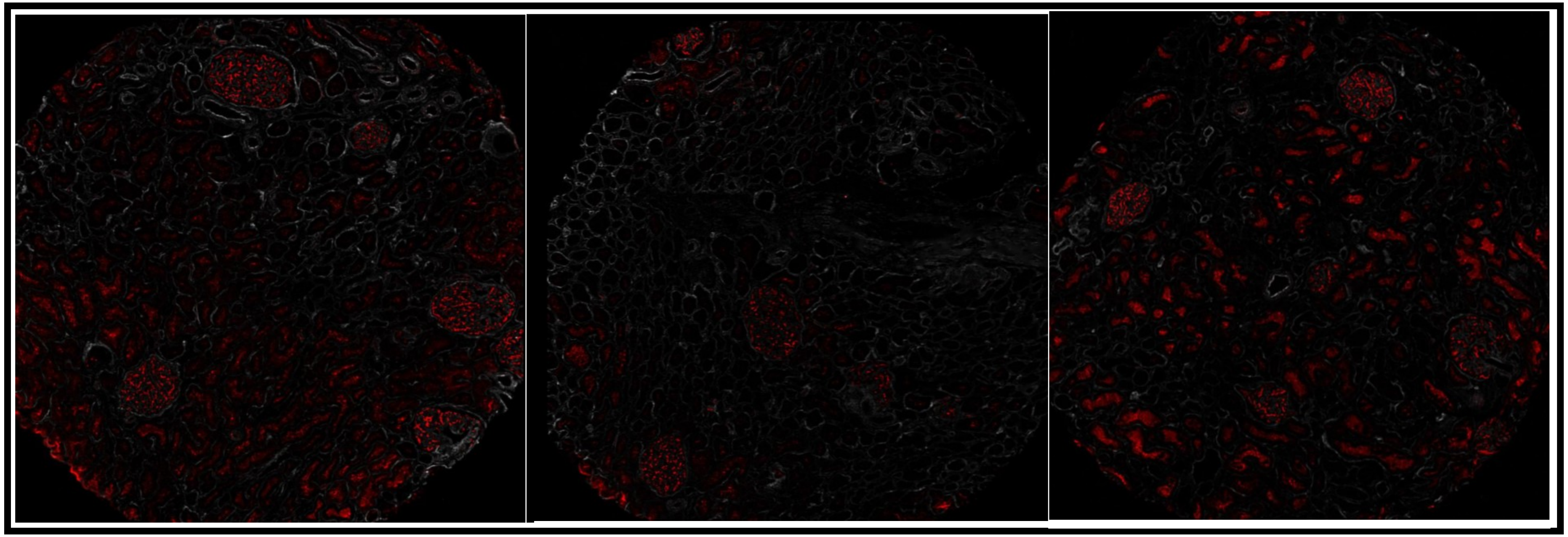

DKDIIA-B

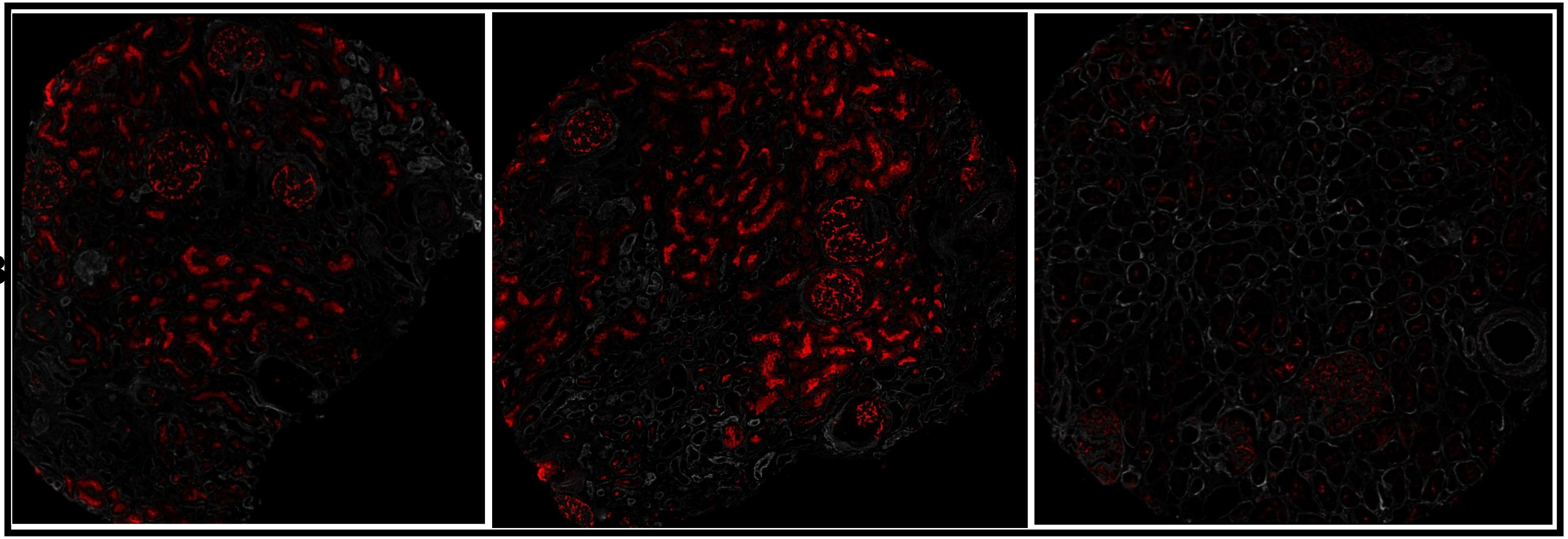

#### Nestin (50-500)

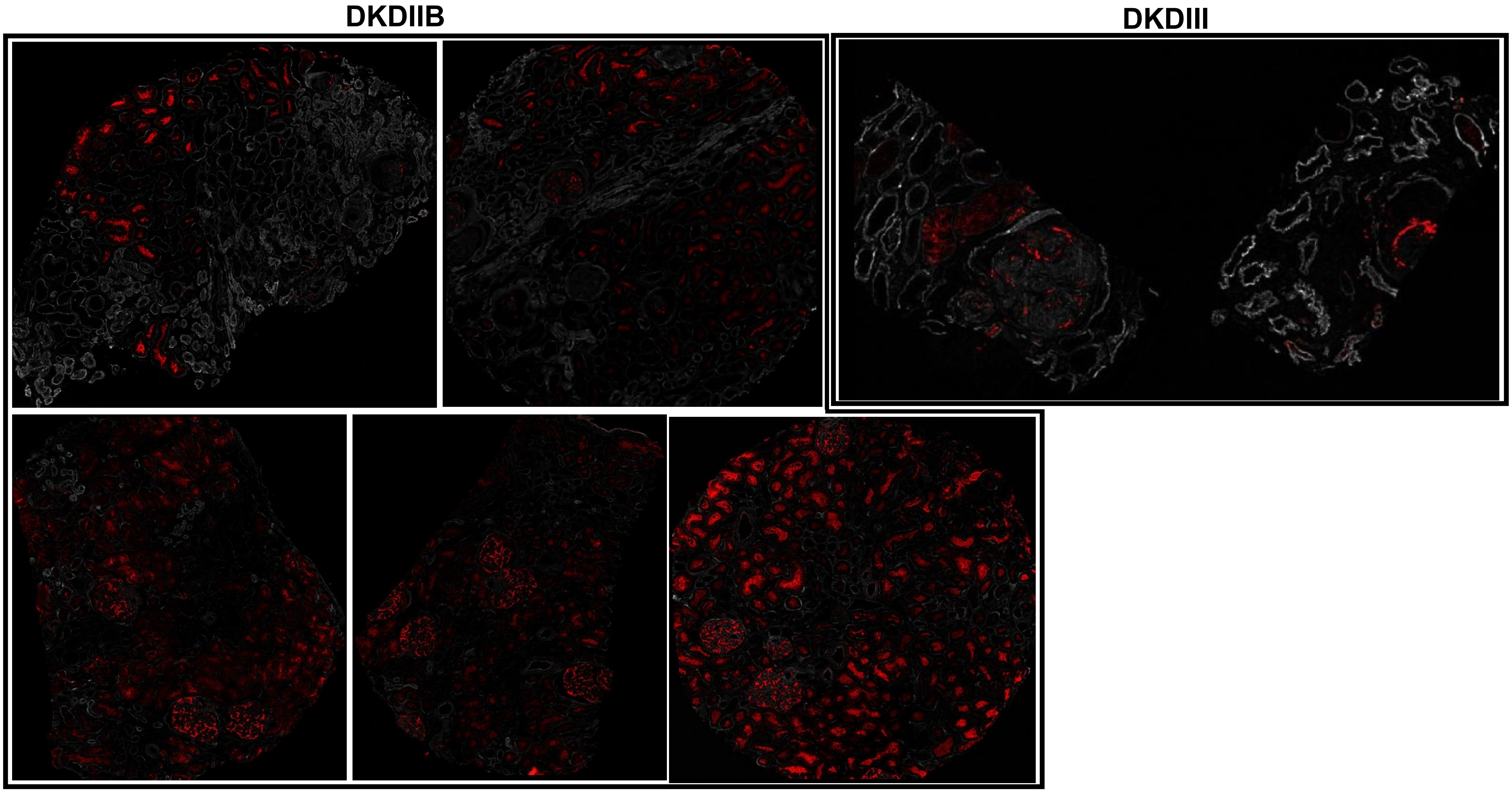

### Nestin (50-500)

#### MEDULLA

DM

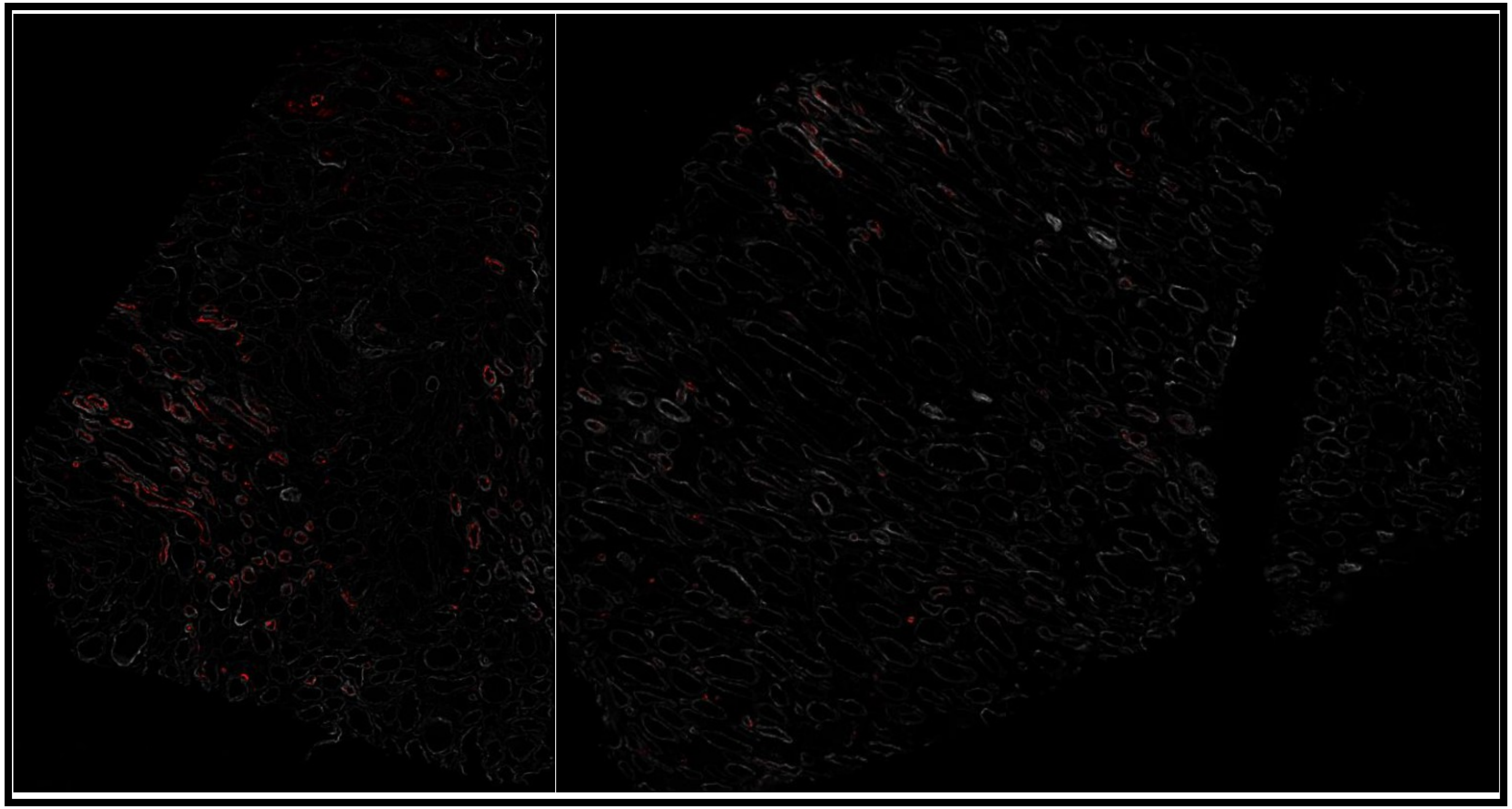

DKDIIA-B

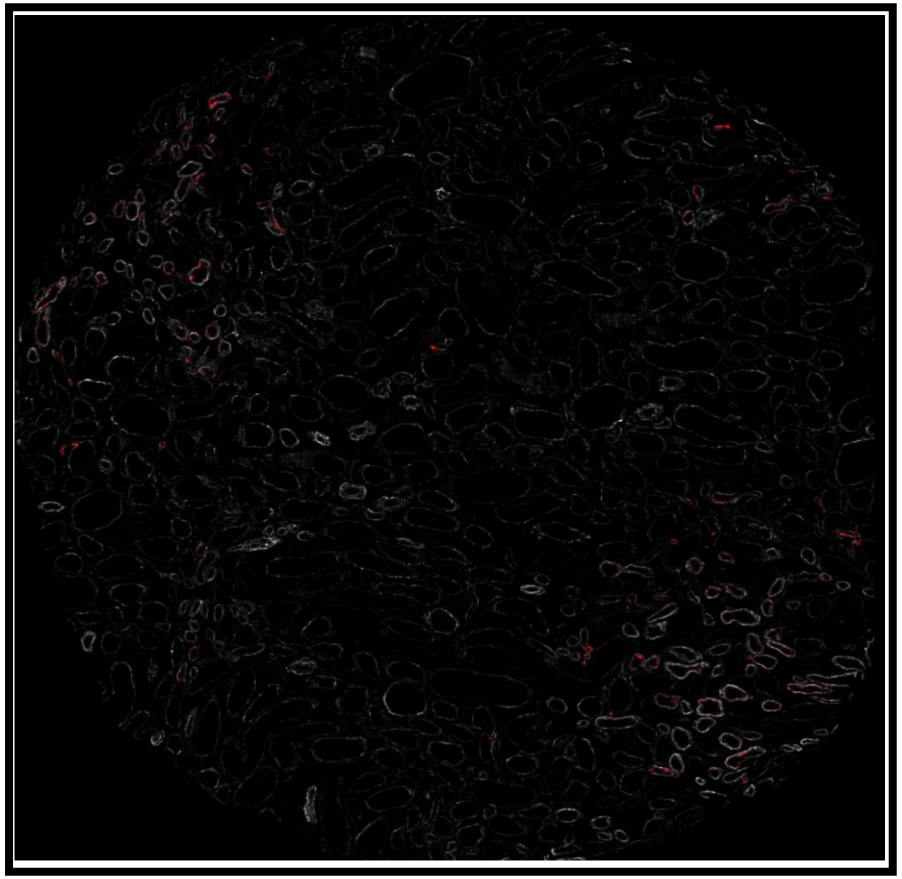

RORγT (500-2000)

DM

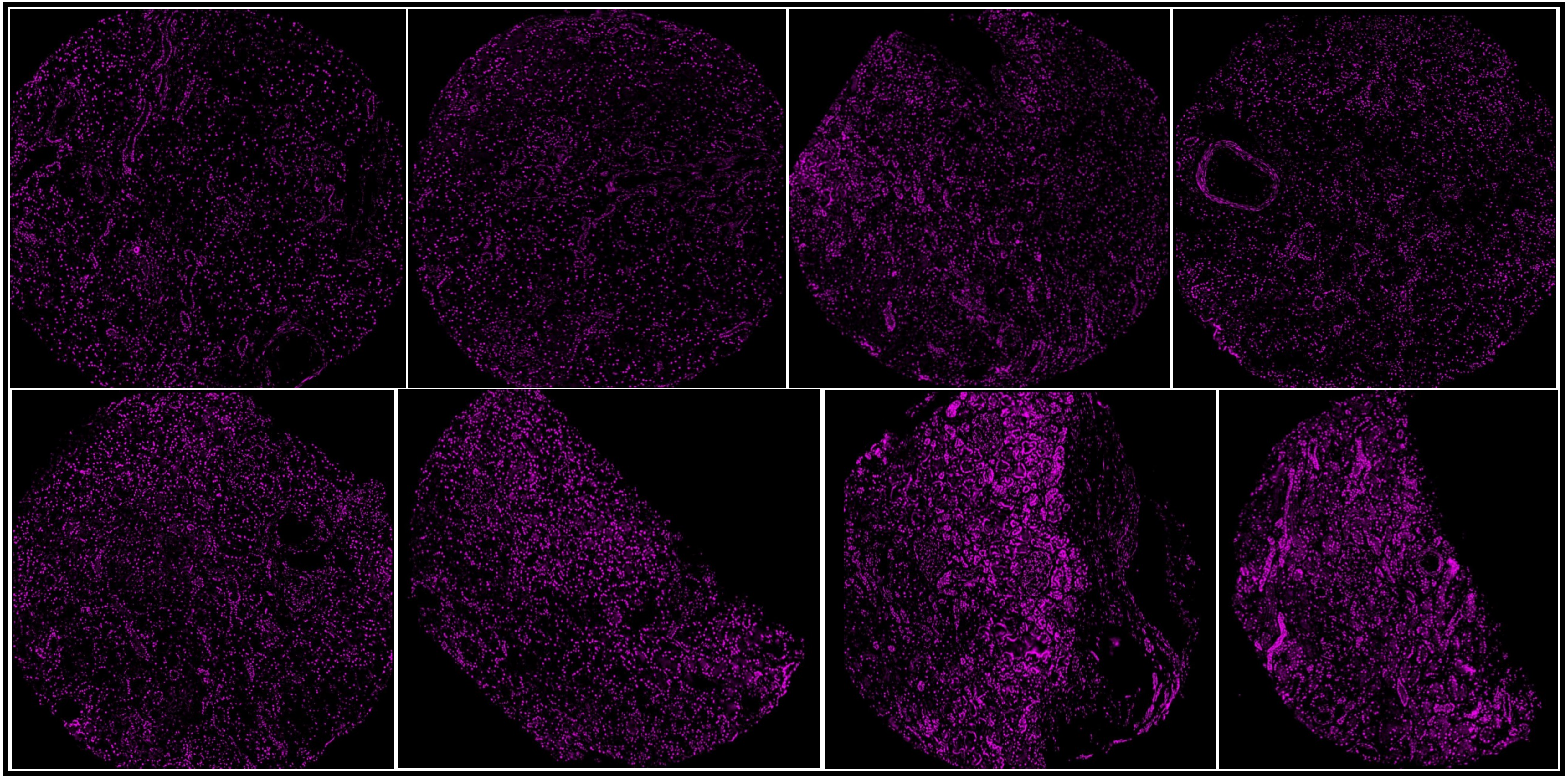

#### RORγT (500-2000)

DKDIIA

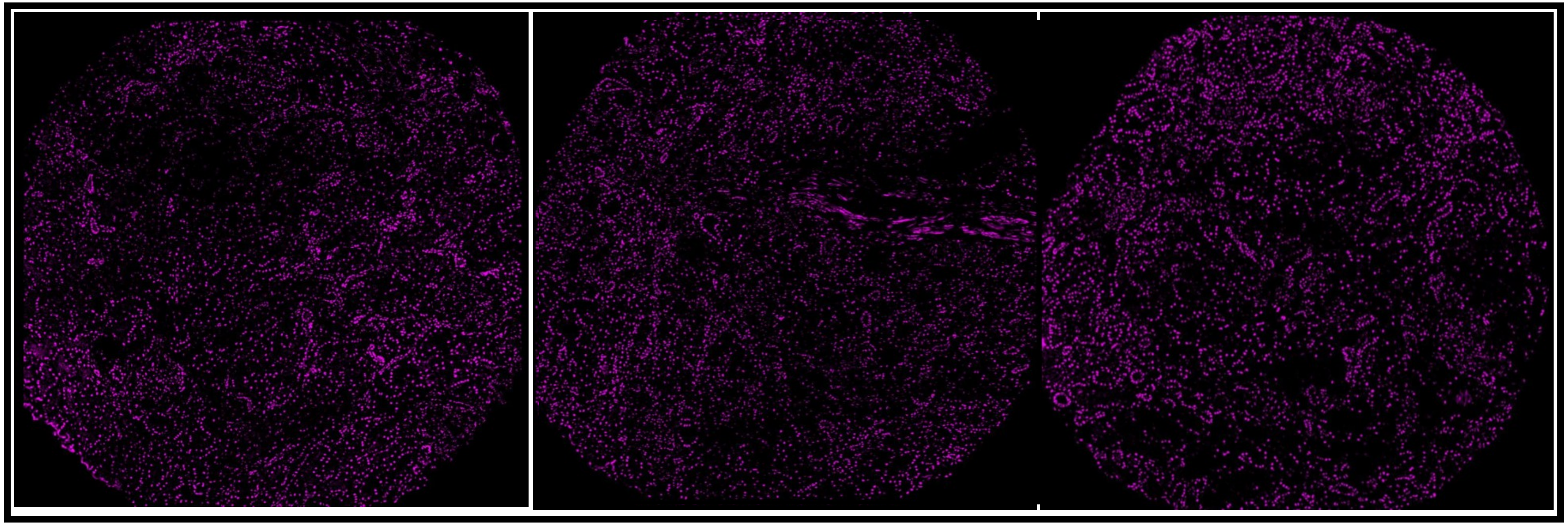

DKDIIA-B

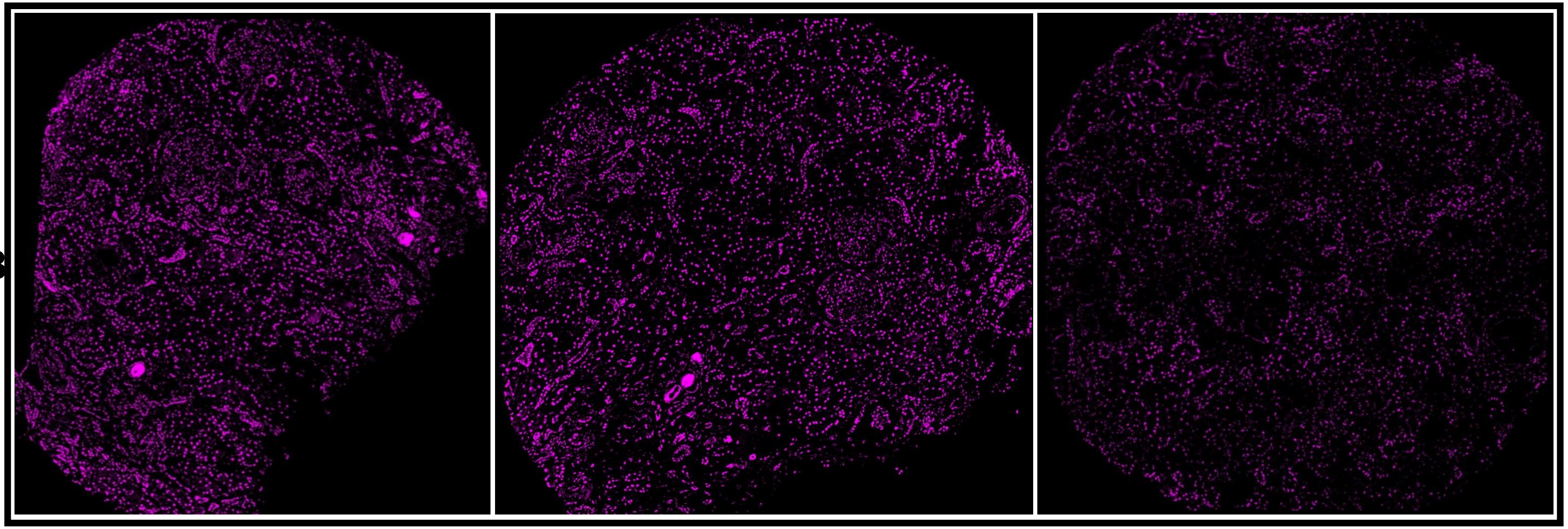

#### RORγT (500-2000)

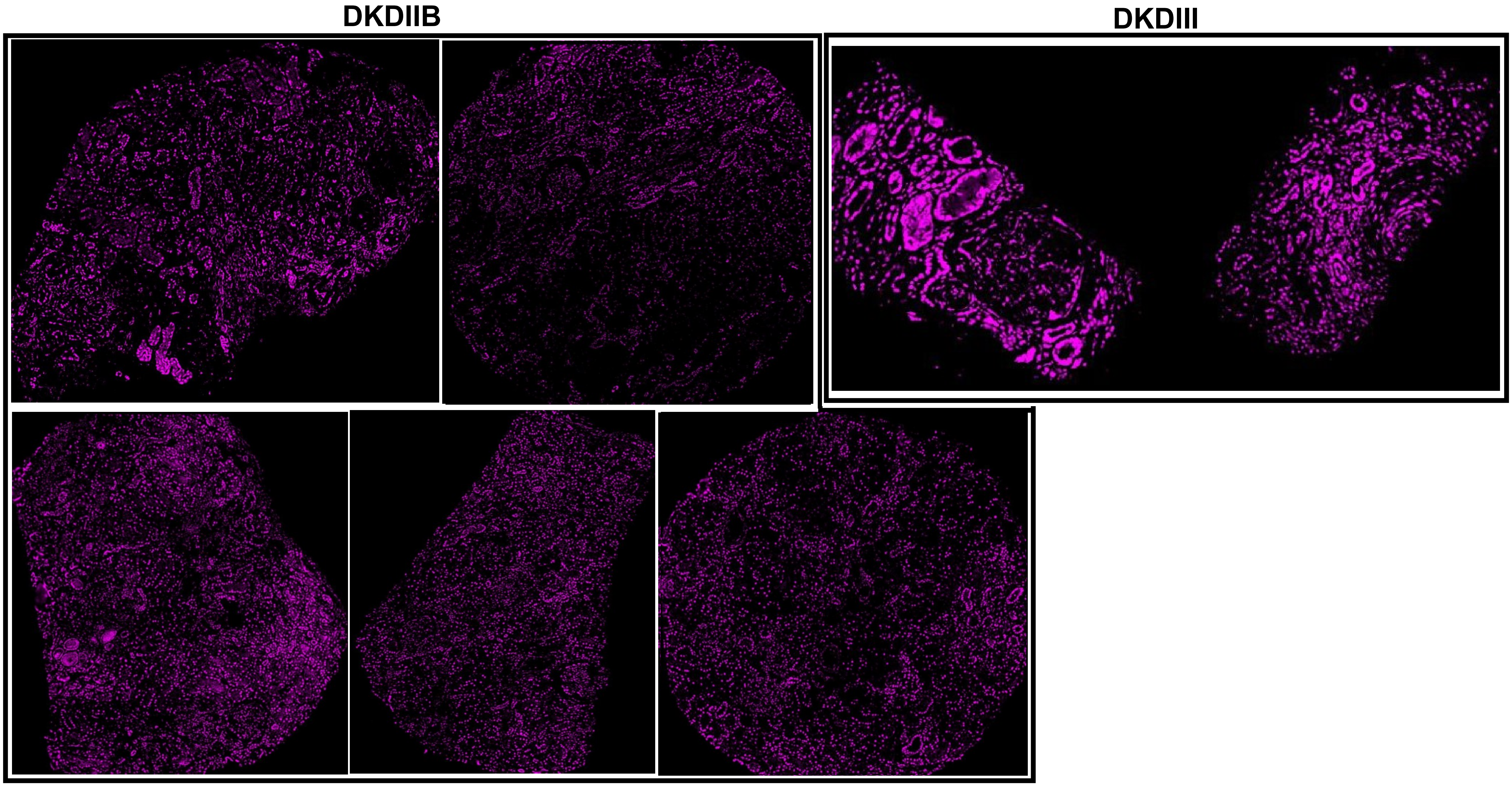

### RORγT (500-2000)

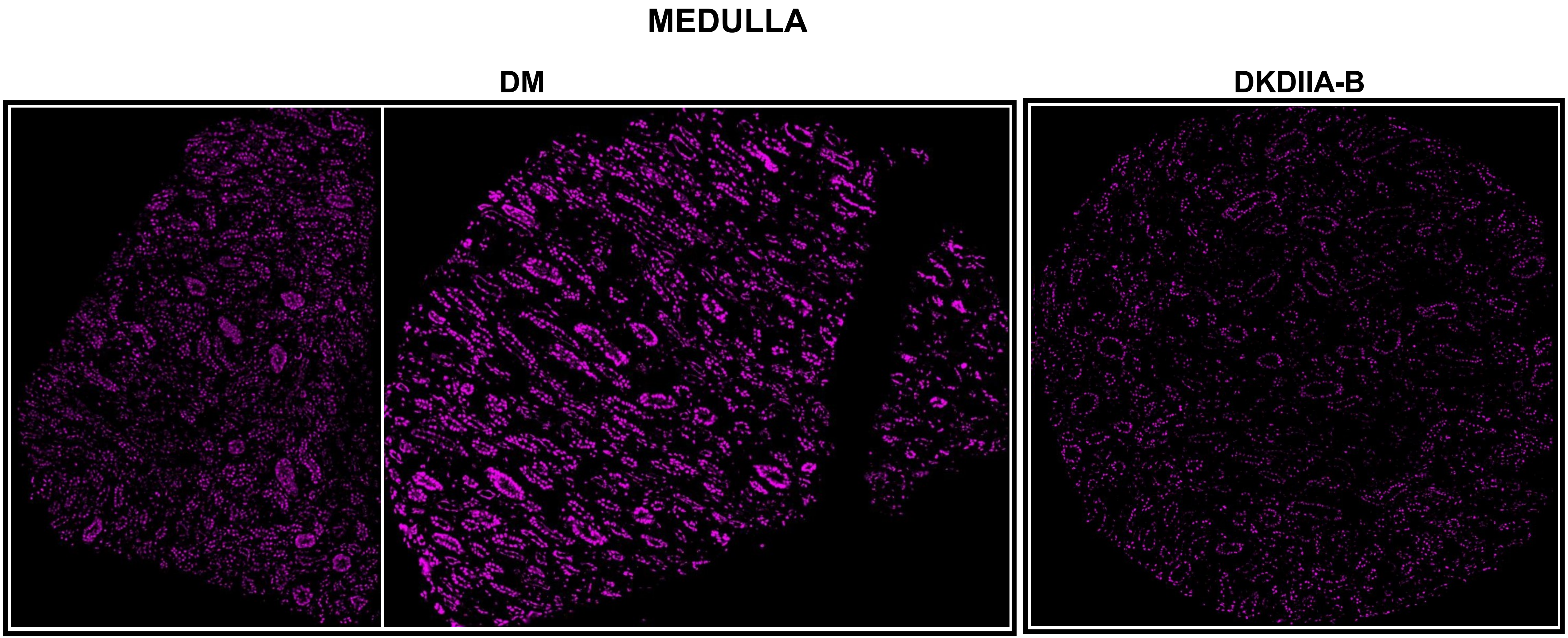

### SPP (50-400)

## DM

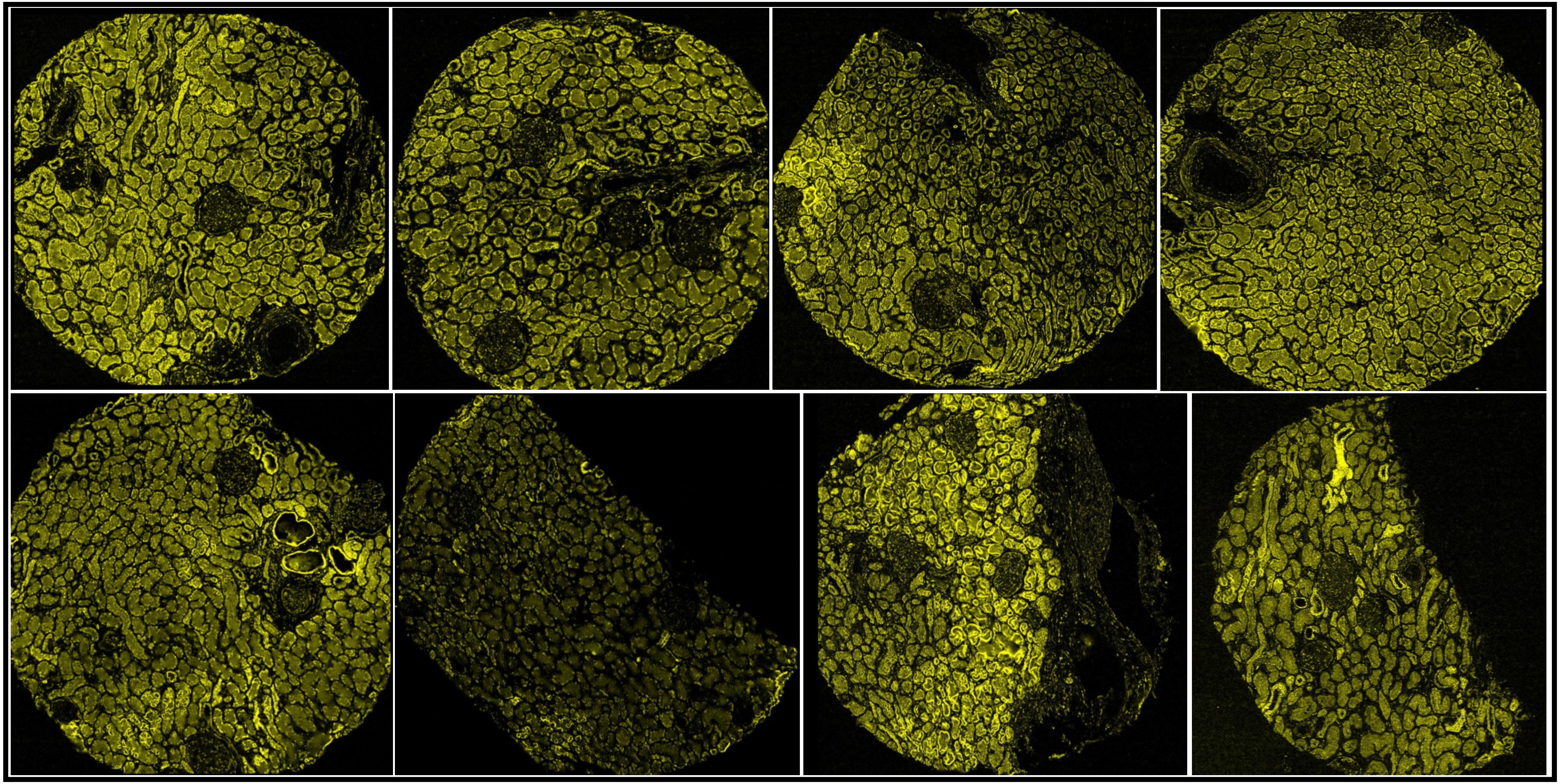

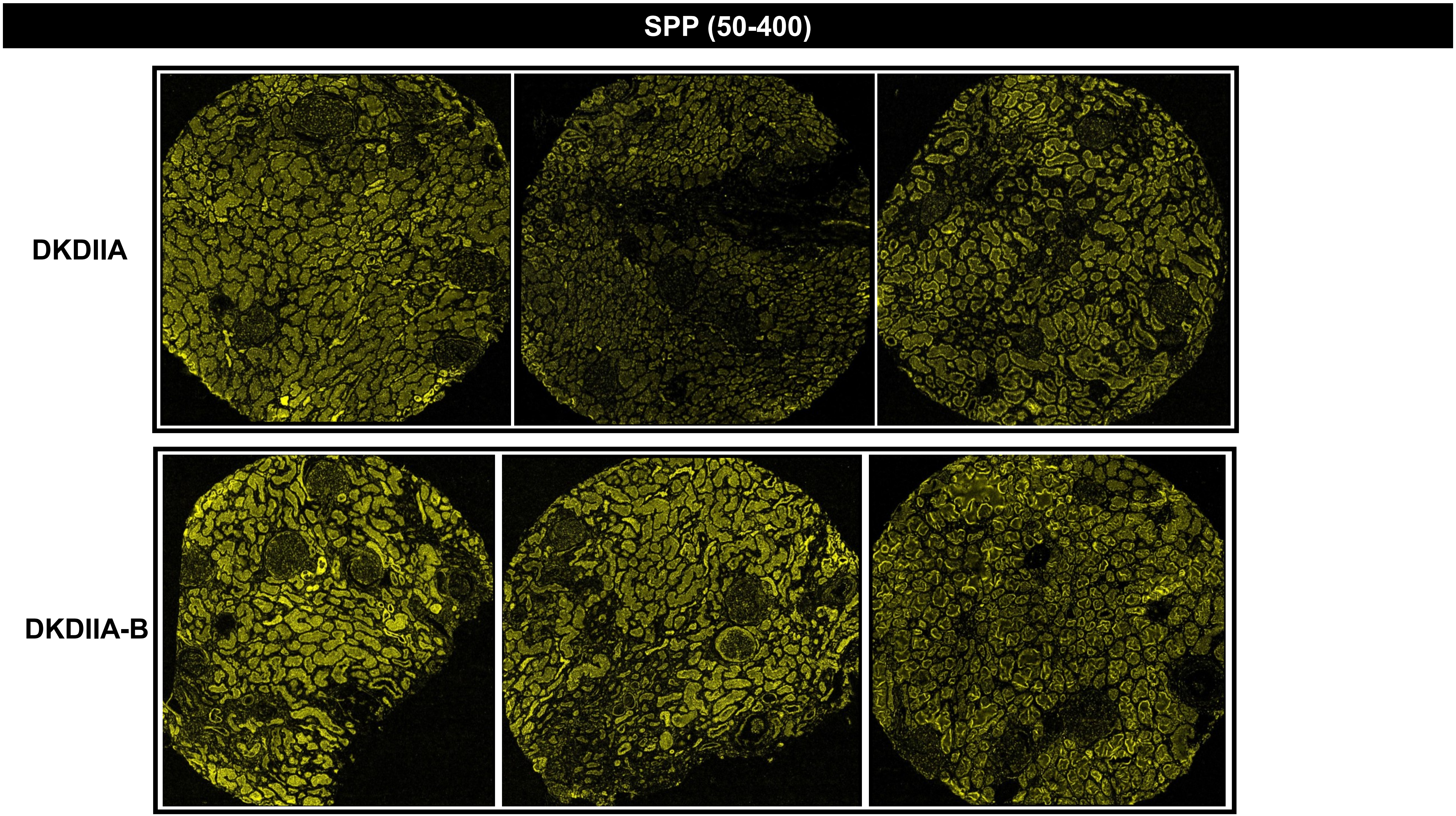
SPP (50-400)
DKDIIA
DKDIIA-B

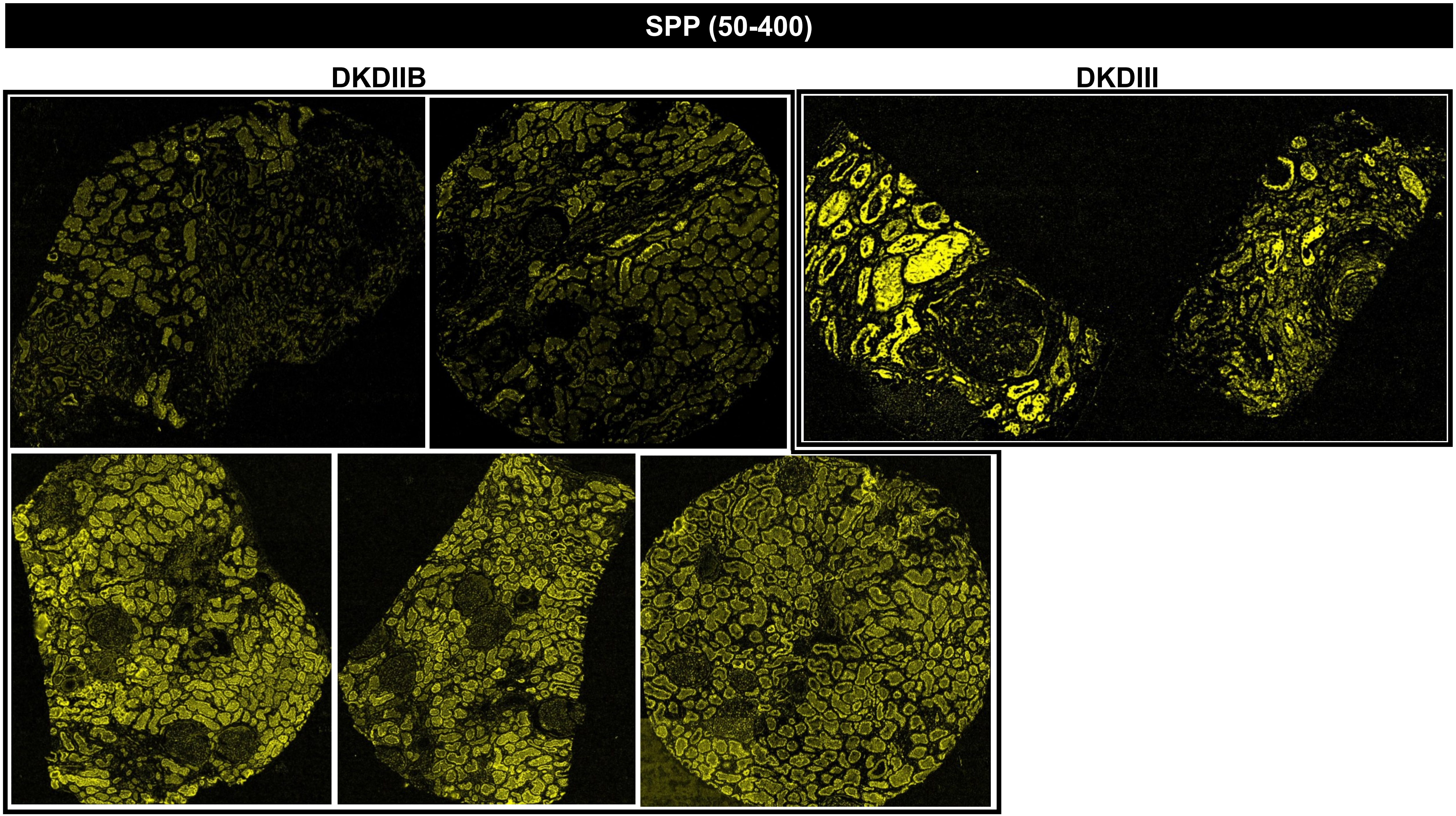
SPP (50-400)
DKDIIB
DKDIII

SPP (50-400)

#### MEDULLA

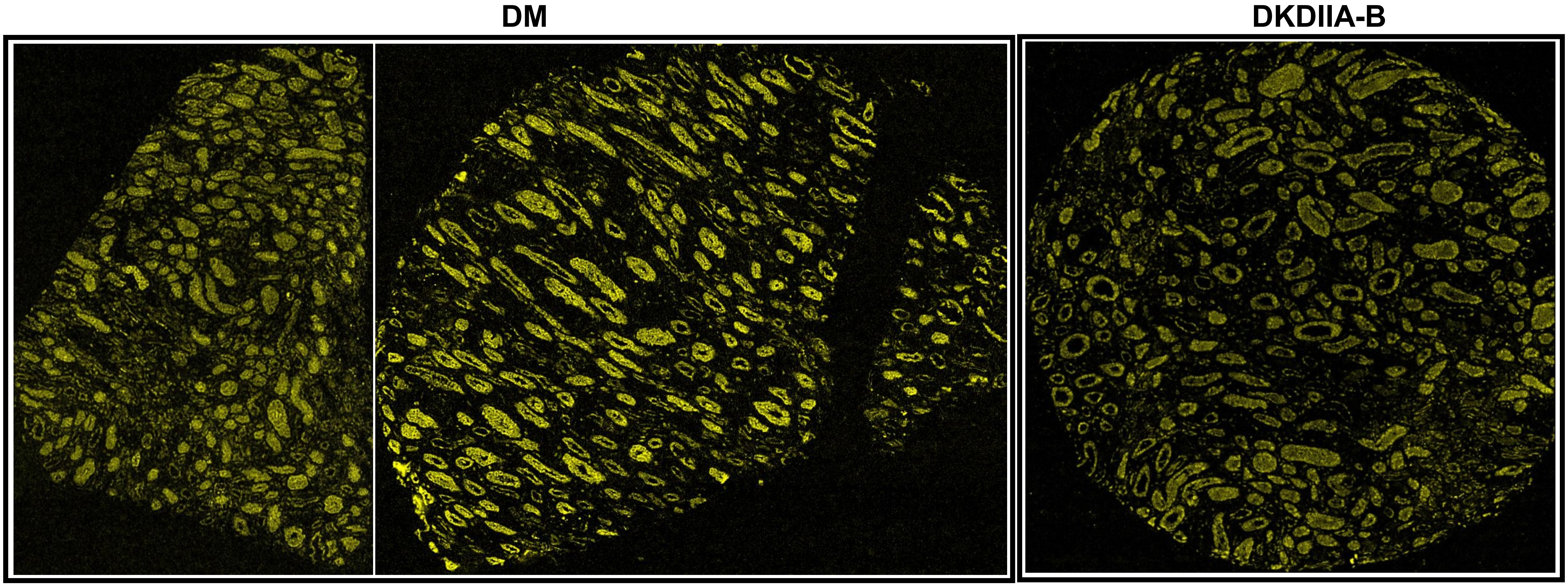

TM (CD141, 40-90)

DM

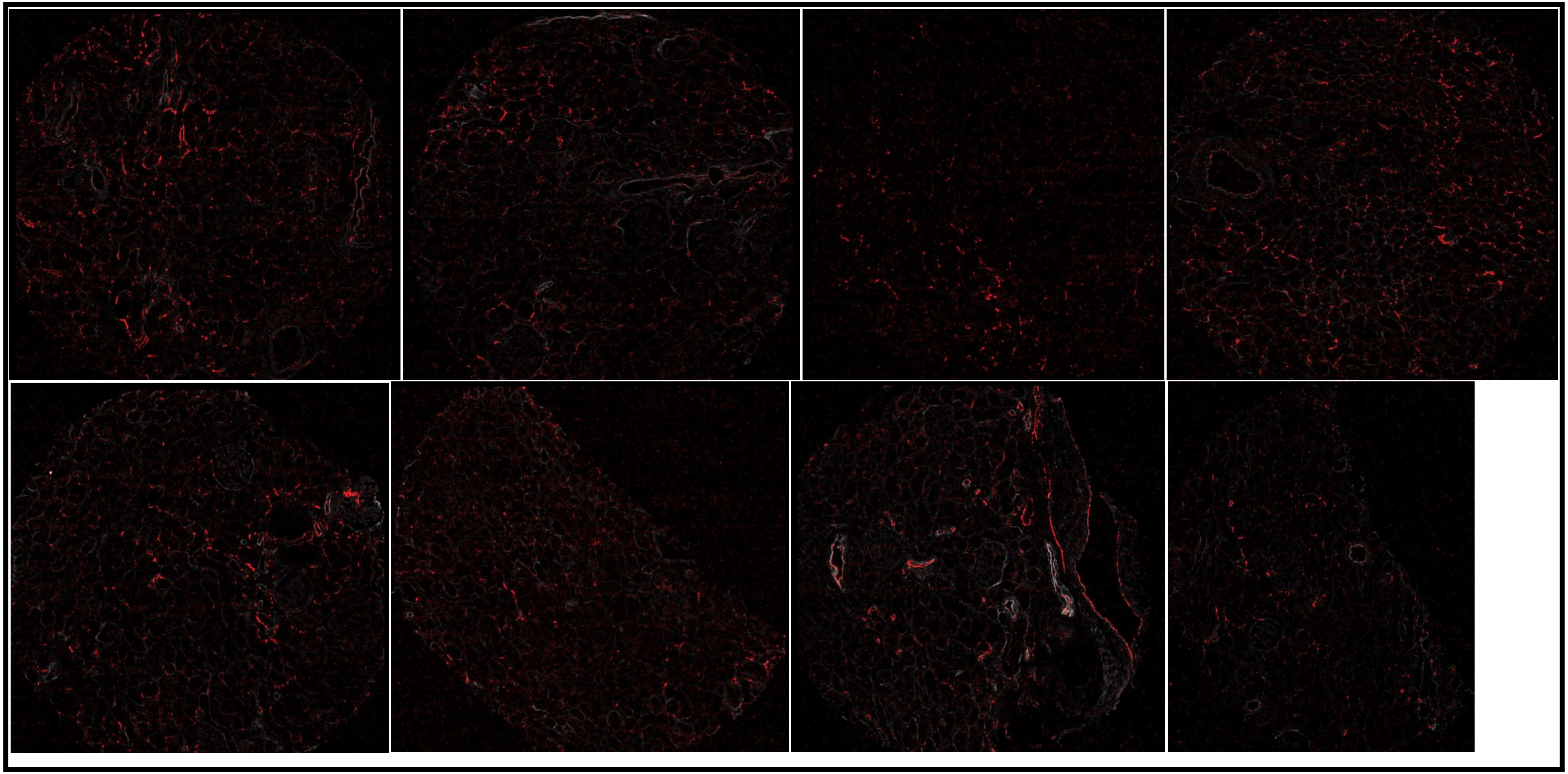

# TM (CD141, 40-90)

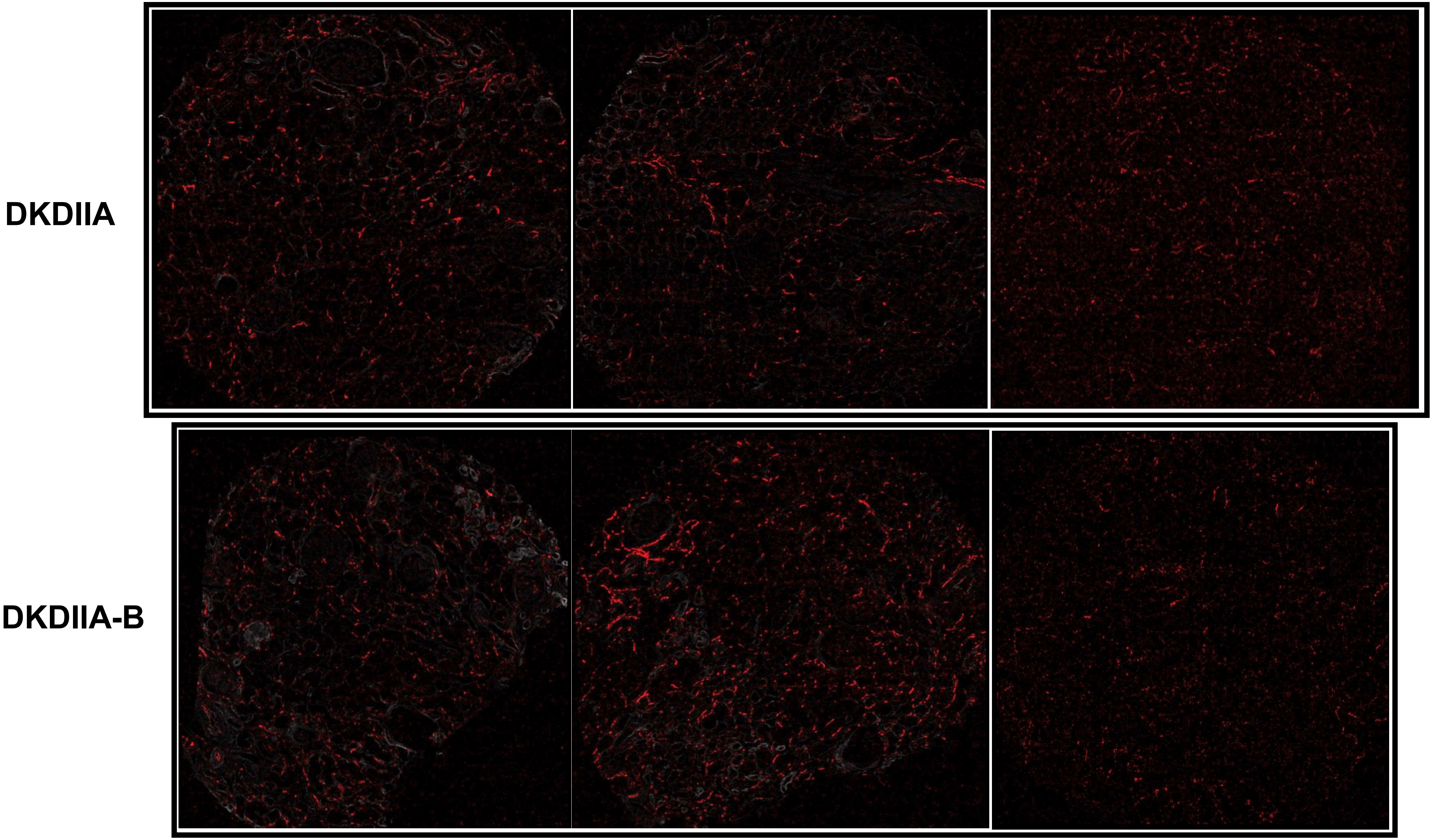

# TM (CD141, 40-90)

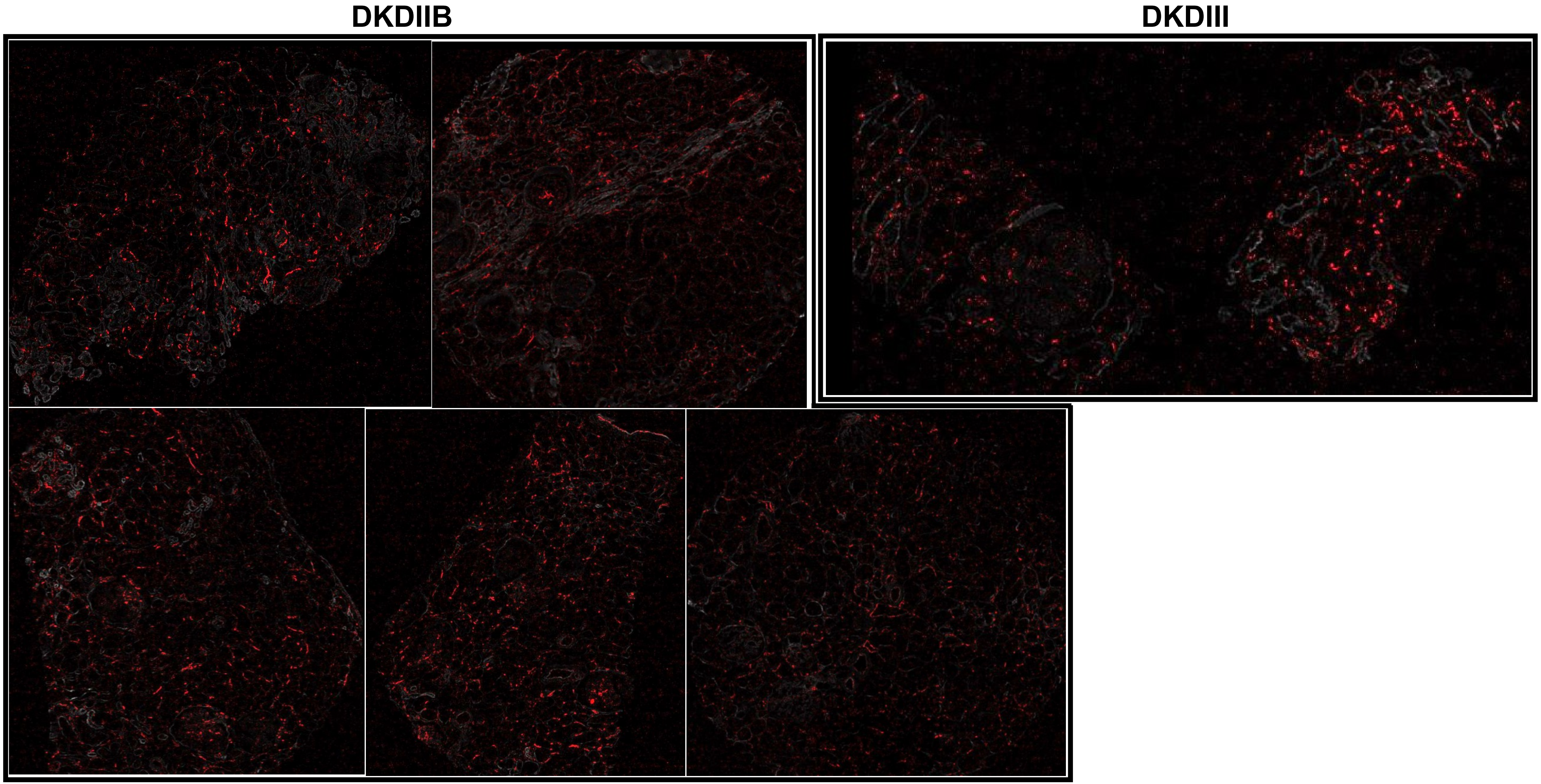

TM (CD141, 40-90)

MEDULLA

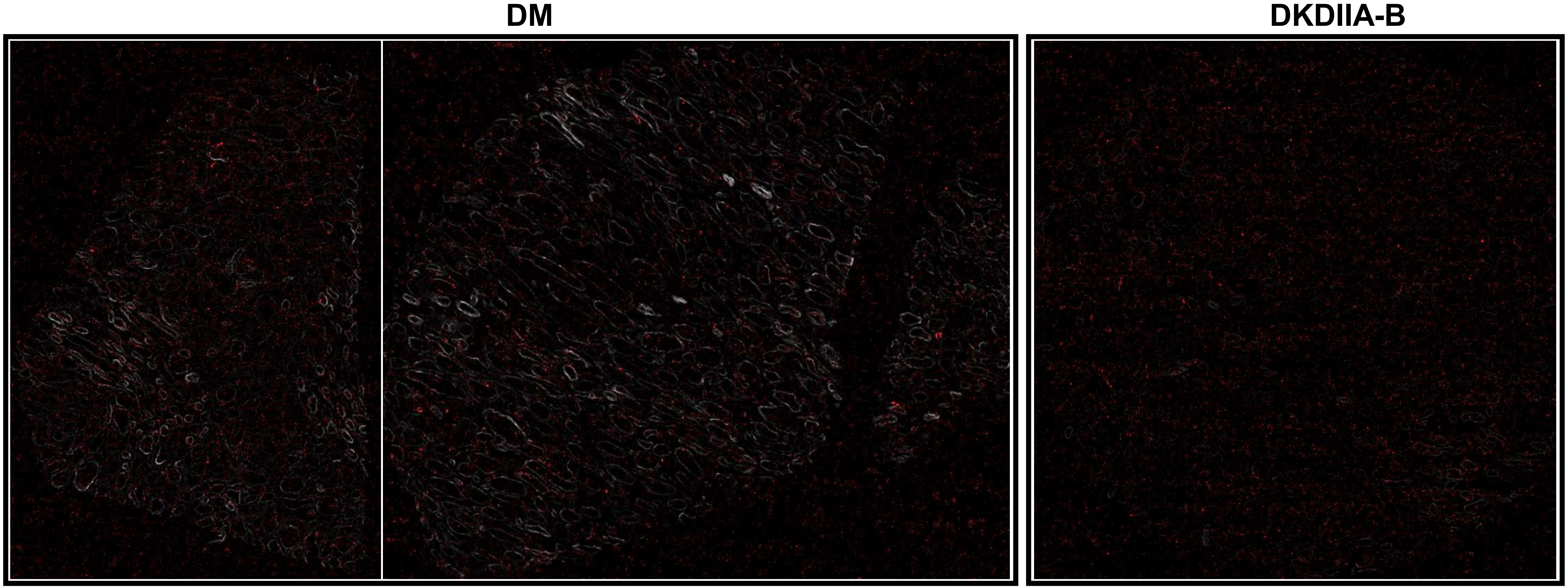

**vWF (200-600)**

**DM**

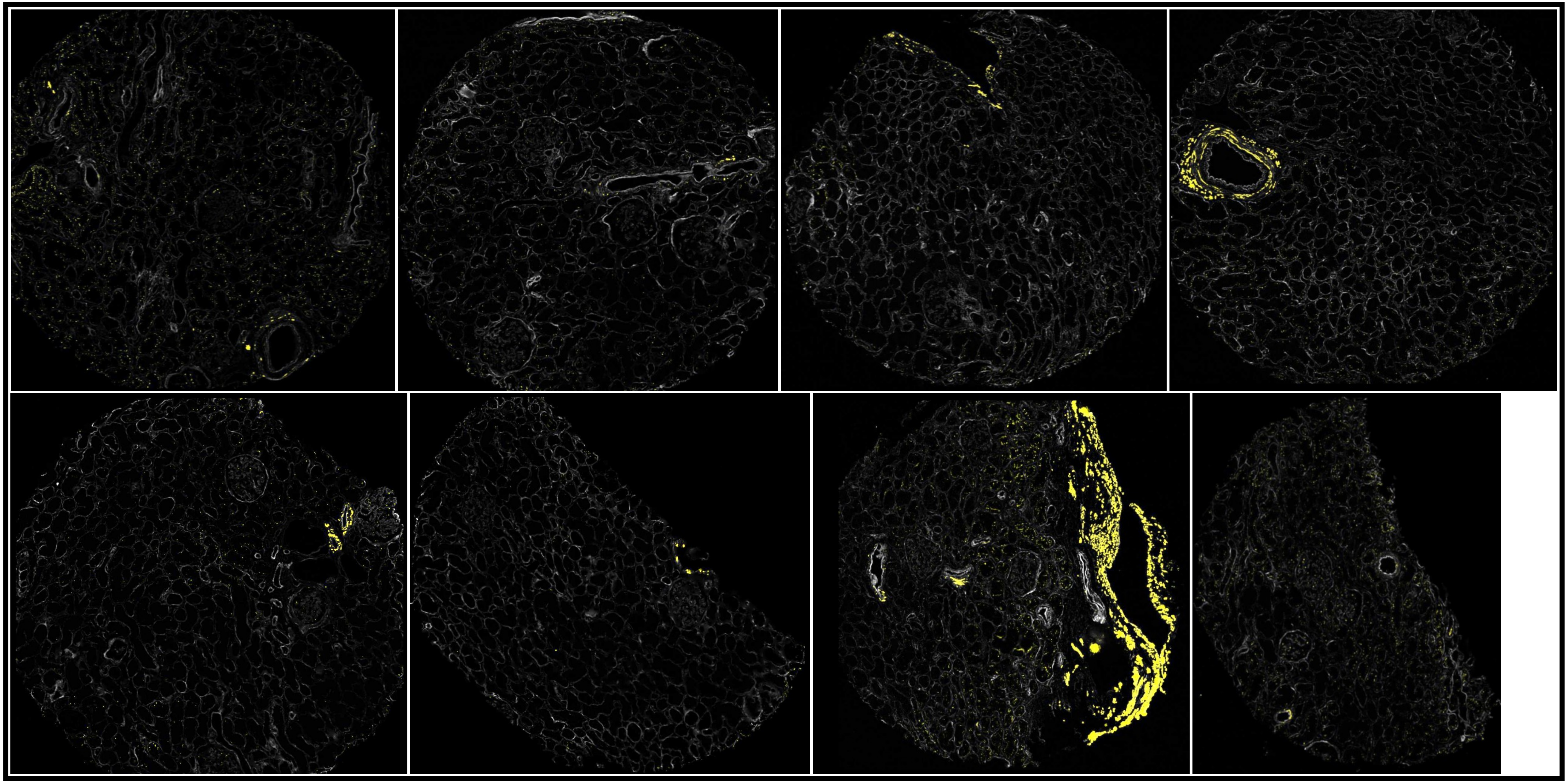

#### vWF (200-600)

DKDIIA

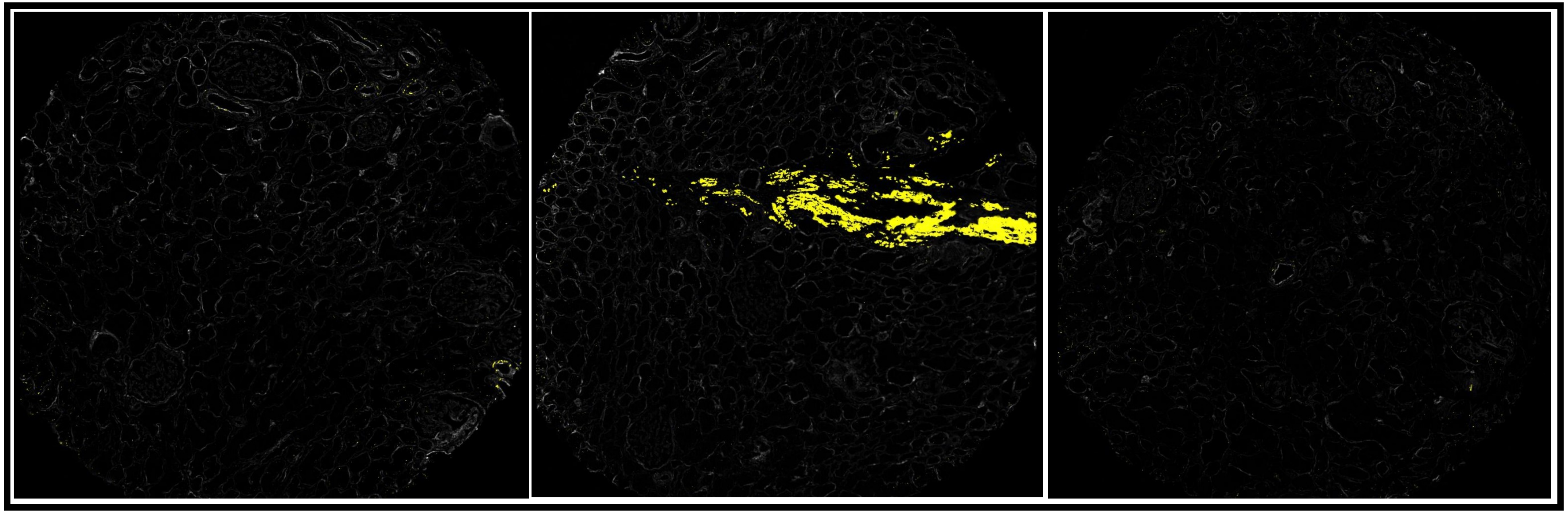

DKDIIA-B

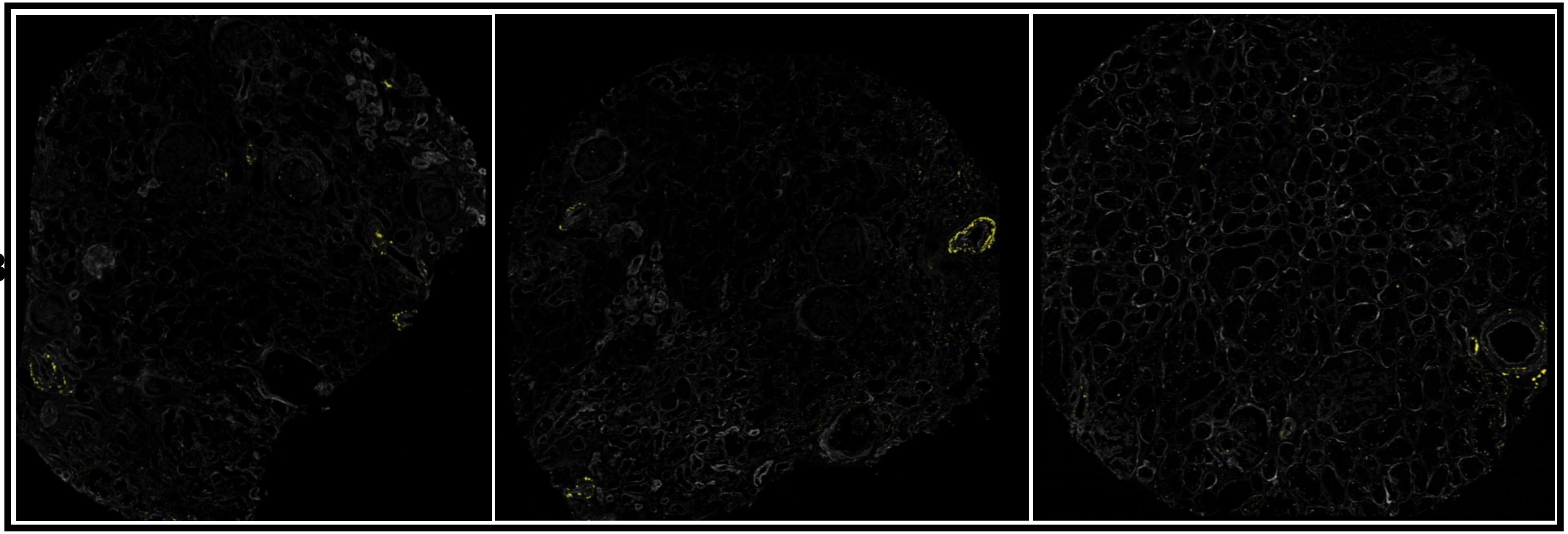

### vWF (200-600)

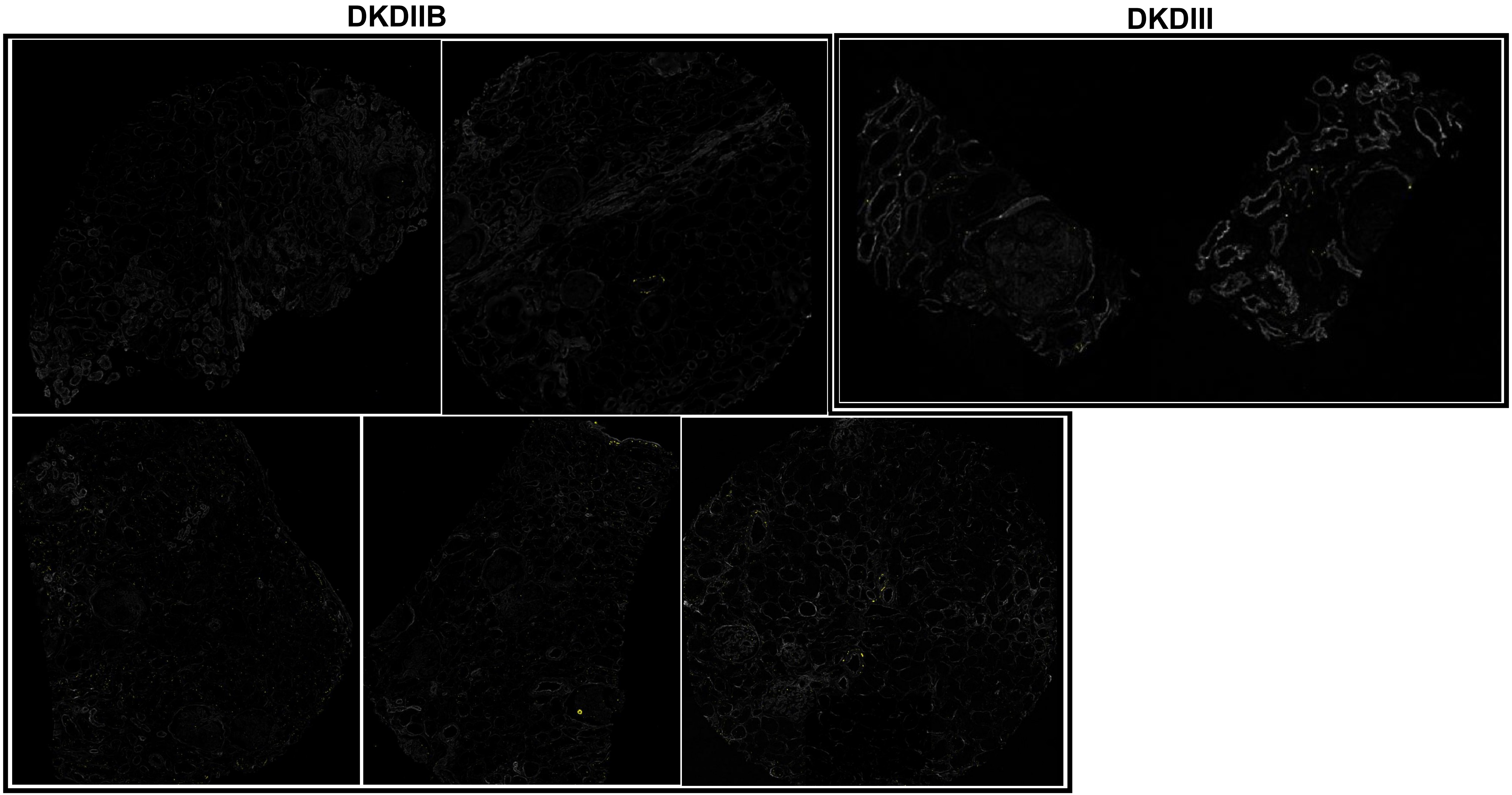

vWF (200-600)

#### MEDULLA

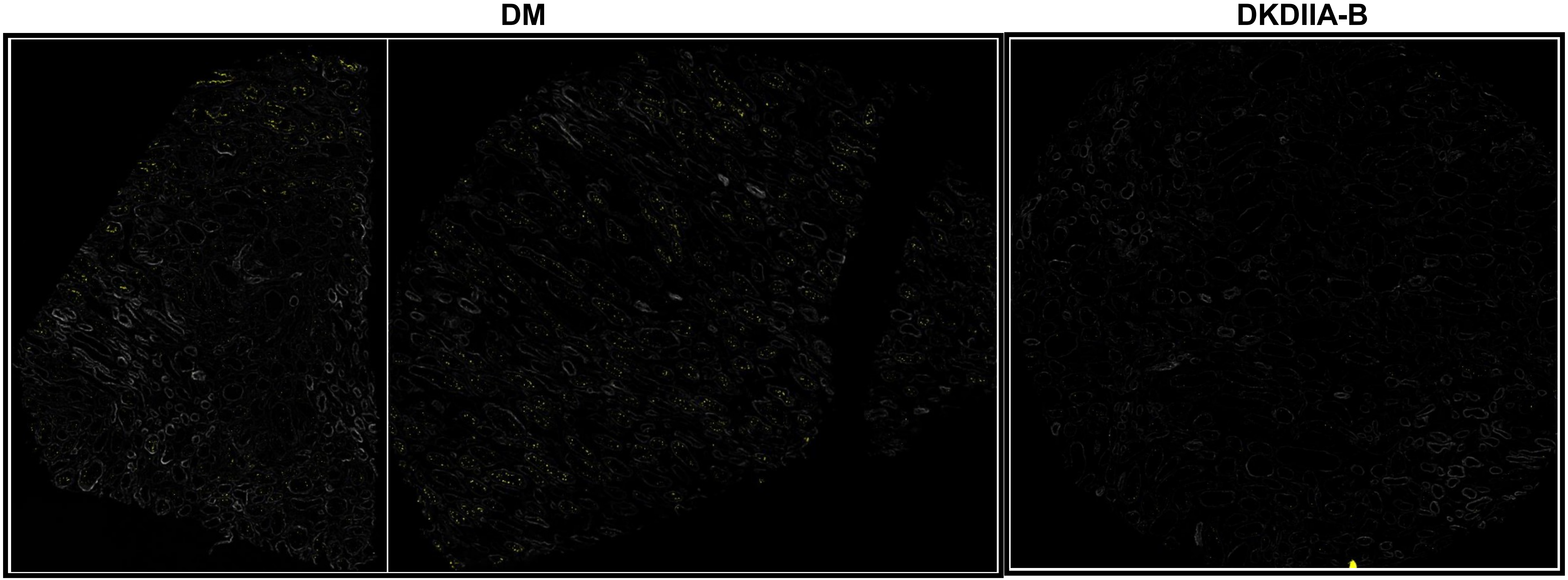

vWF (50-100)

DM

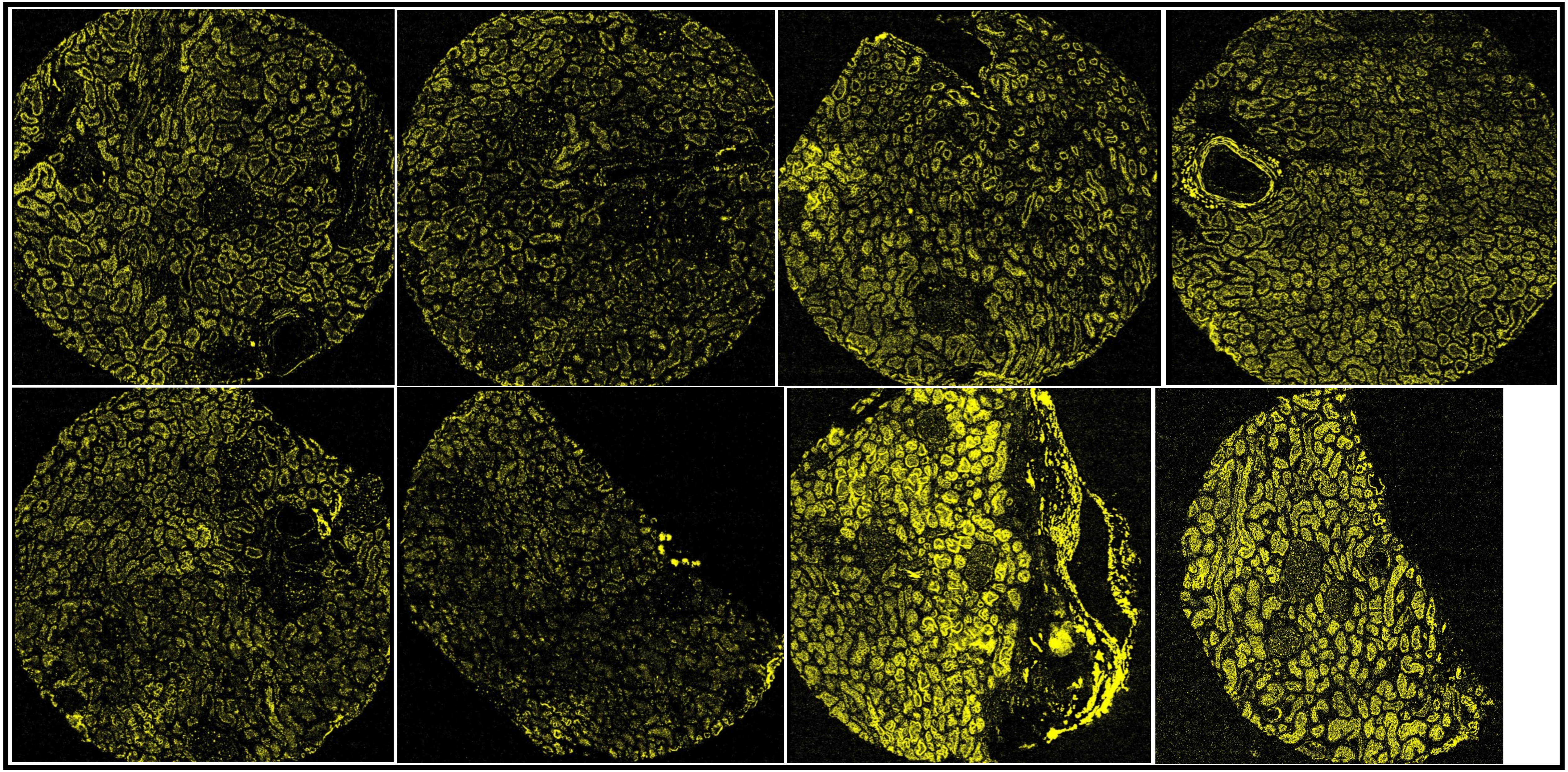

### vWF (50-100)

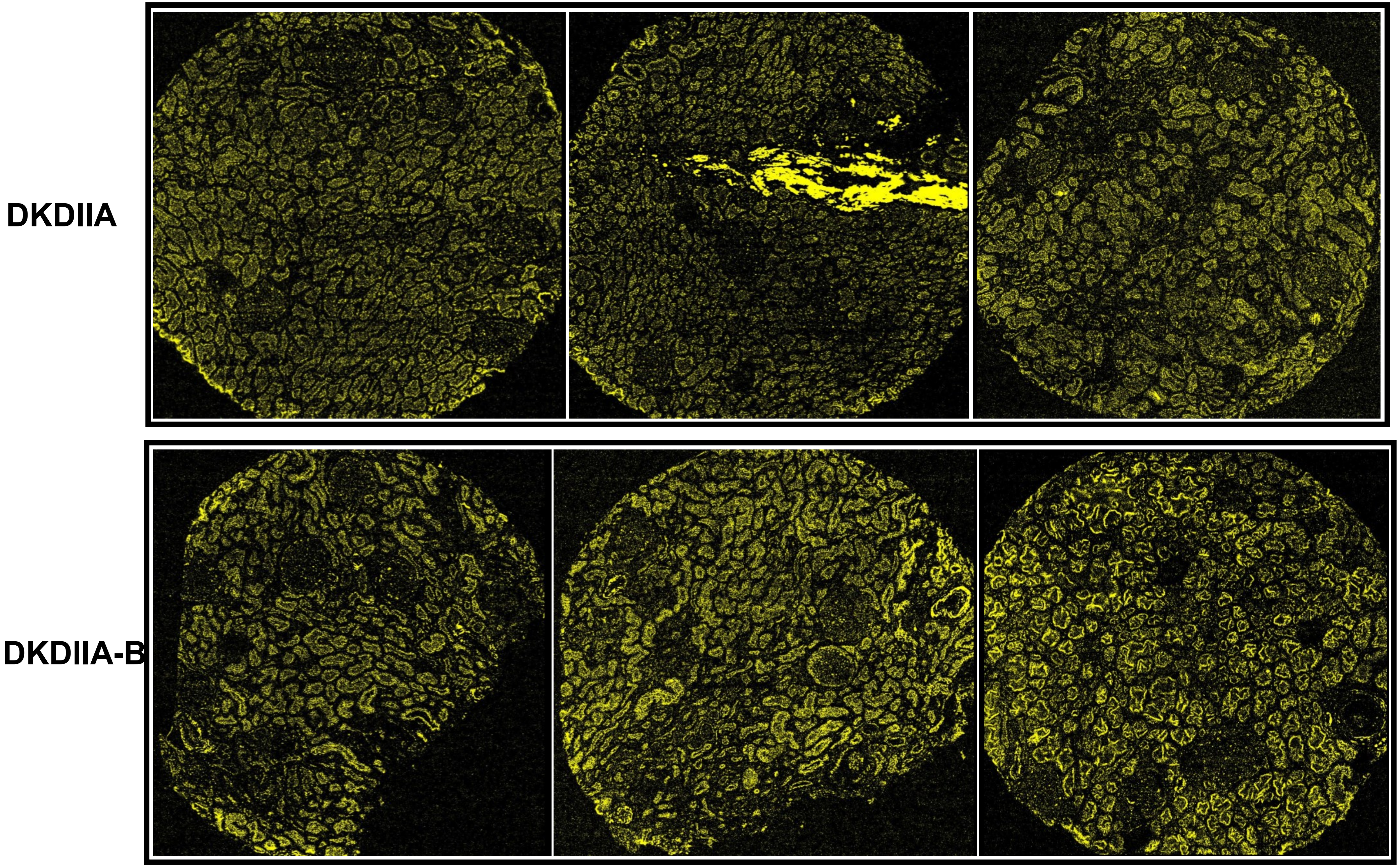

#### vWF (50-100)

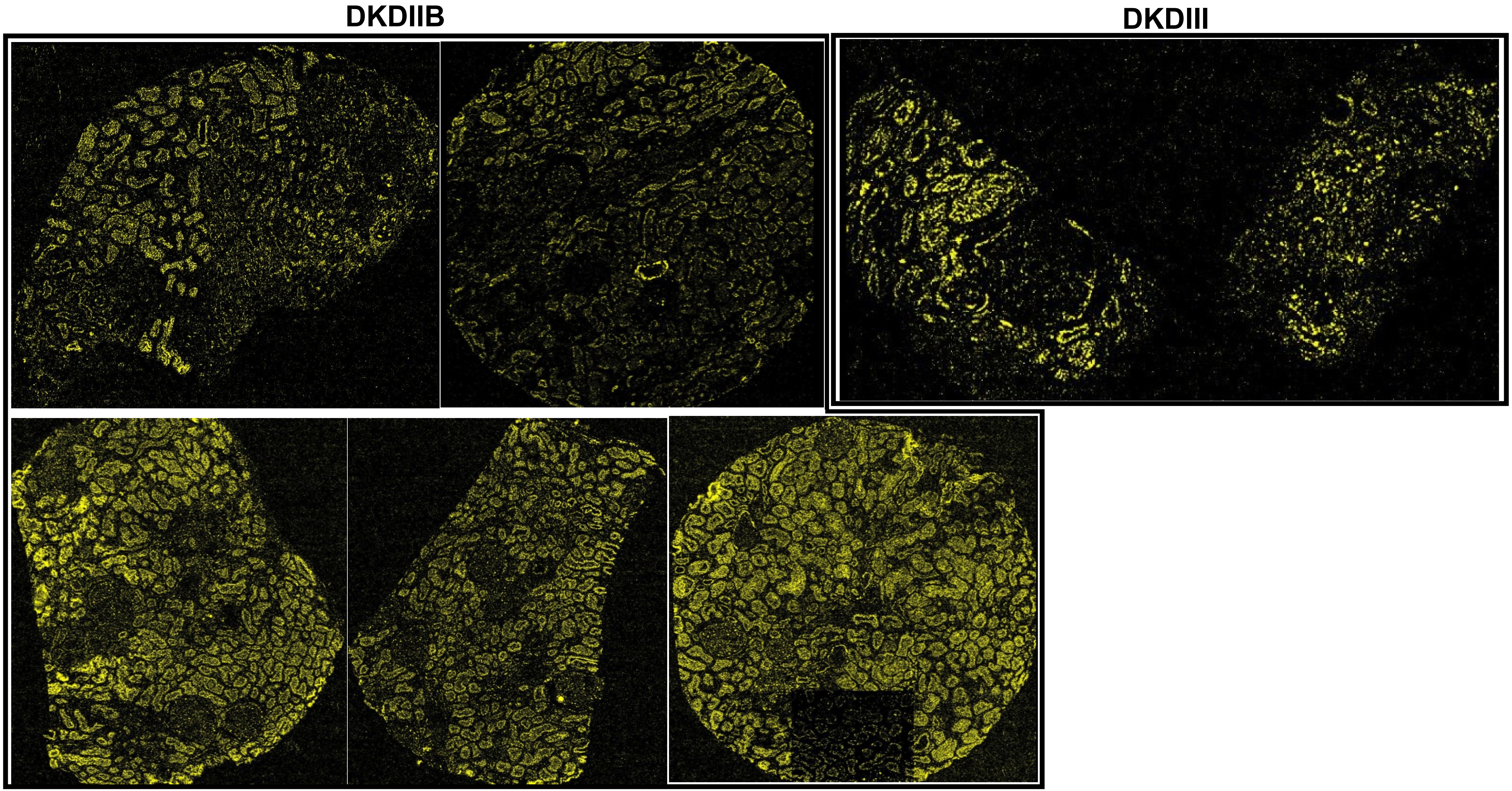

### vWF (50-100)

#### MEDULLA

DM

DKDIIA-B

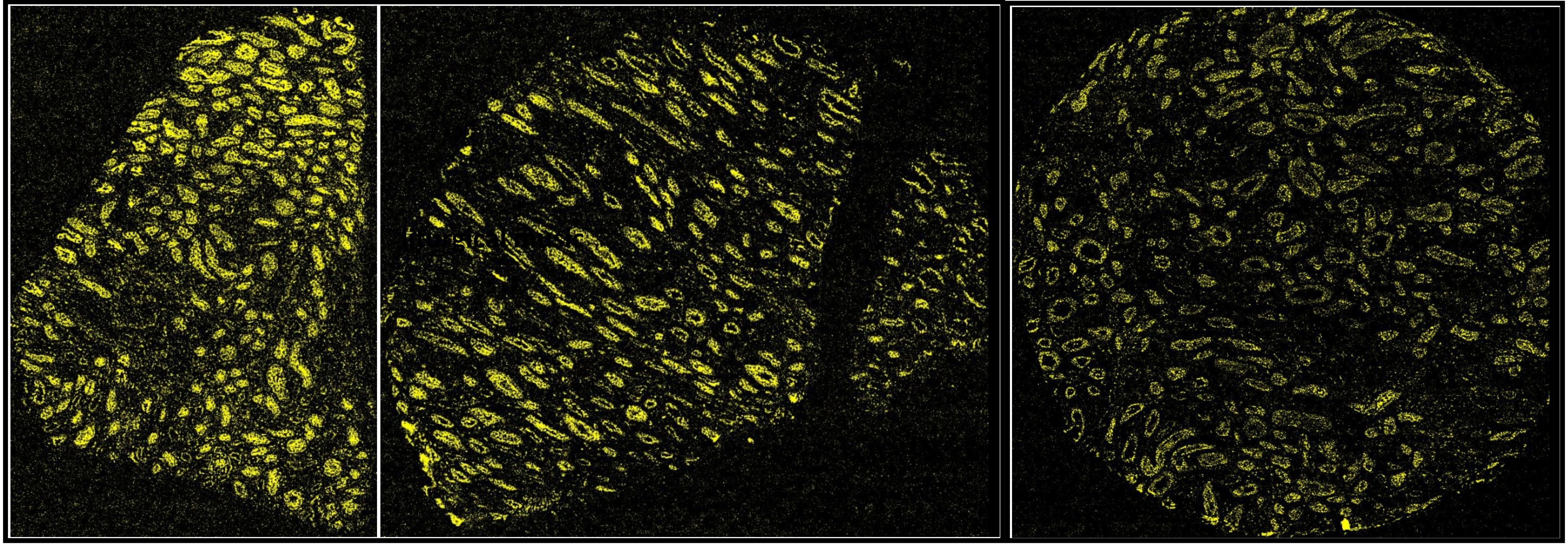

**5 stains** (αSMA, CD7, CD83, CD9, COL4)

**DM**

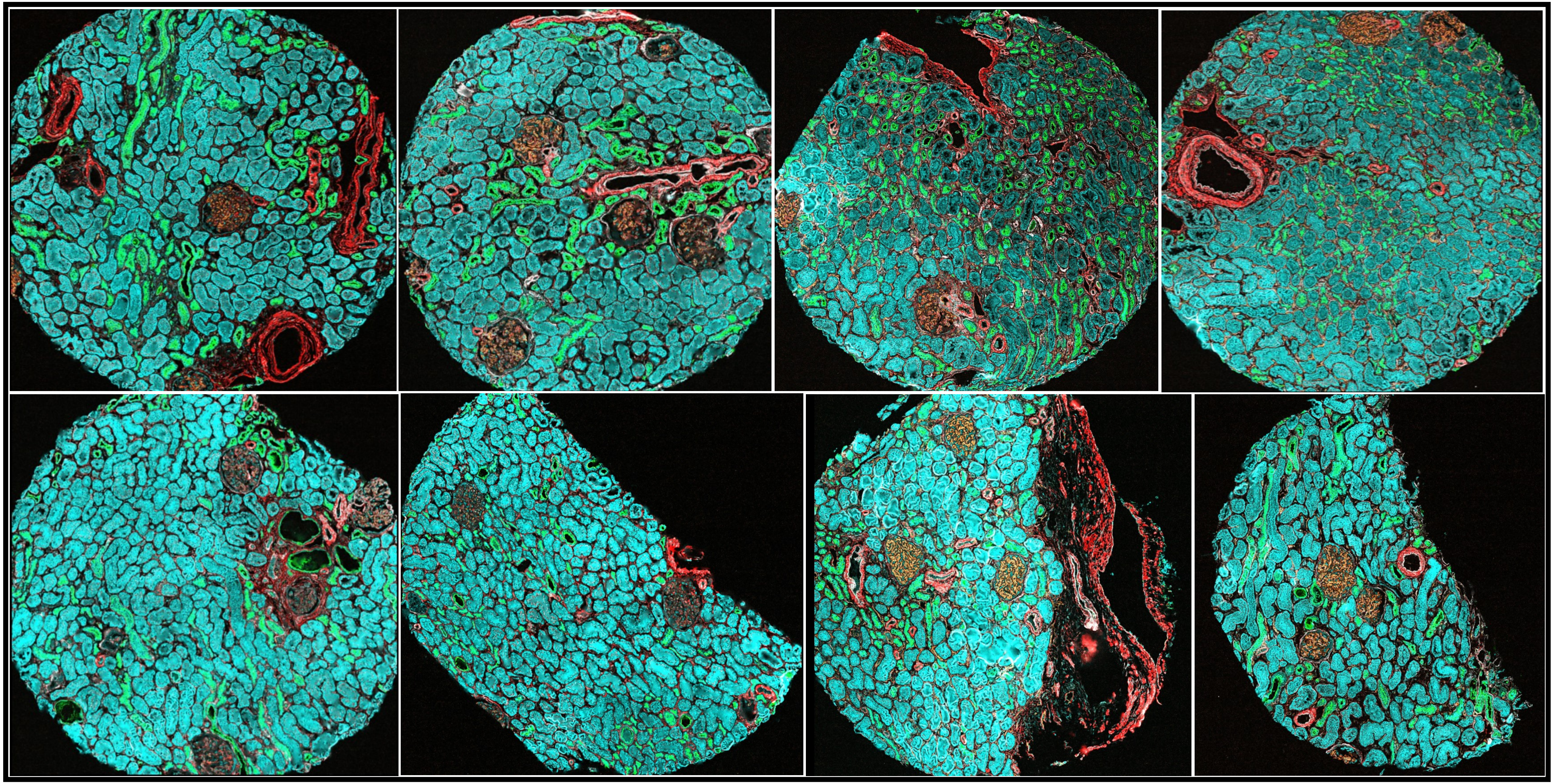

**5 stains** (αSMA, CD7, CD83, CD9, COL4)

DKDIIA

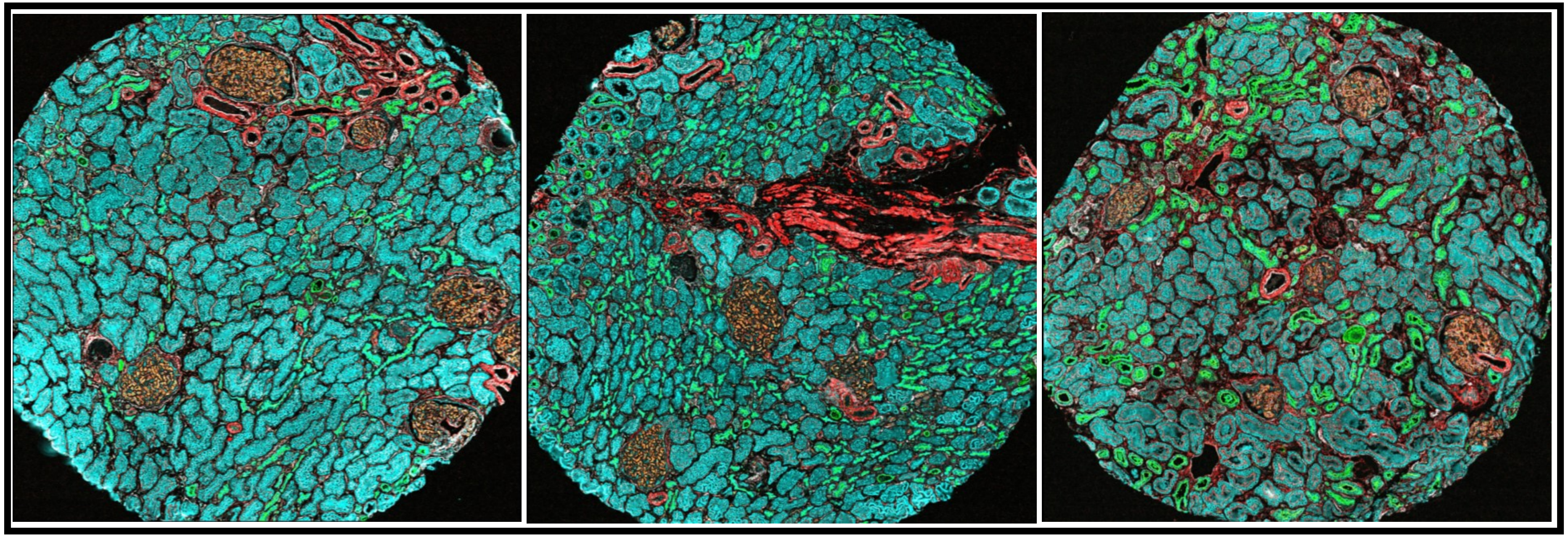

DKDIIA-B

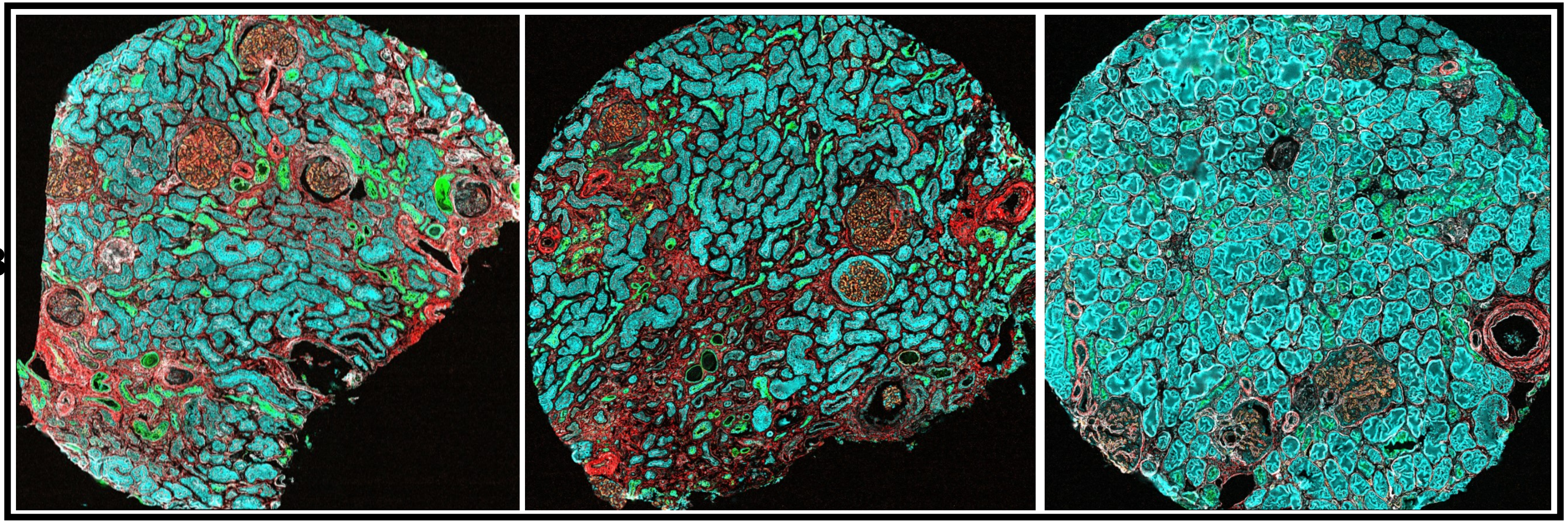

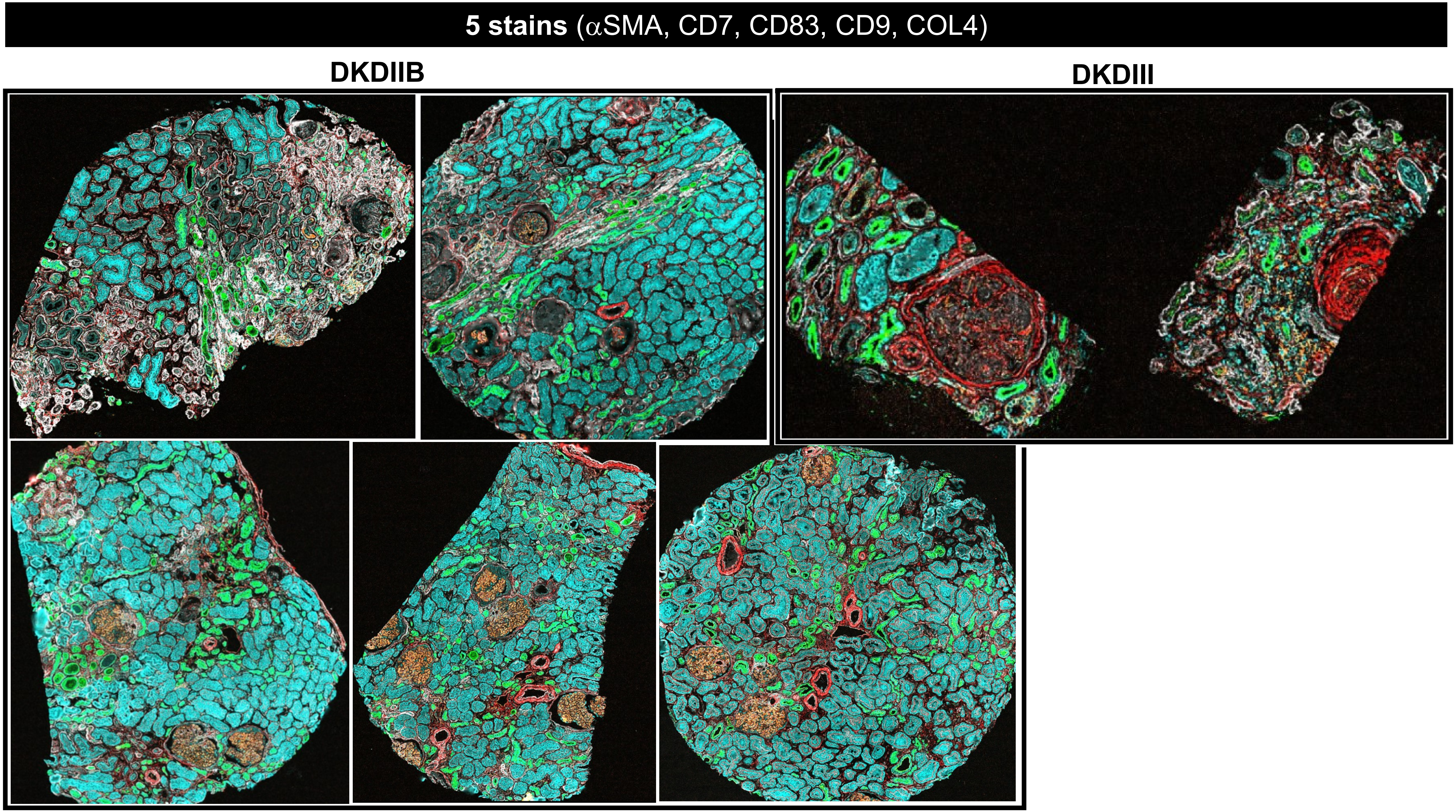
5 stains (αSMA, CD7, CD83, CD9, COL4)
DKDIIB
DKDIII

**5 stains** (αSMA, CD7, CD83, CD9, COL4)

### MEDULLA

**DM**

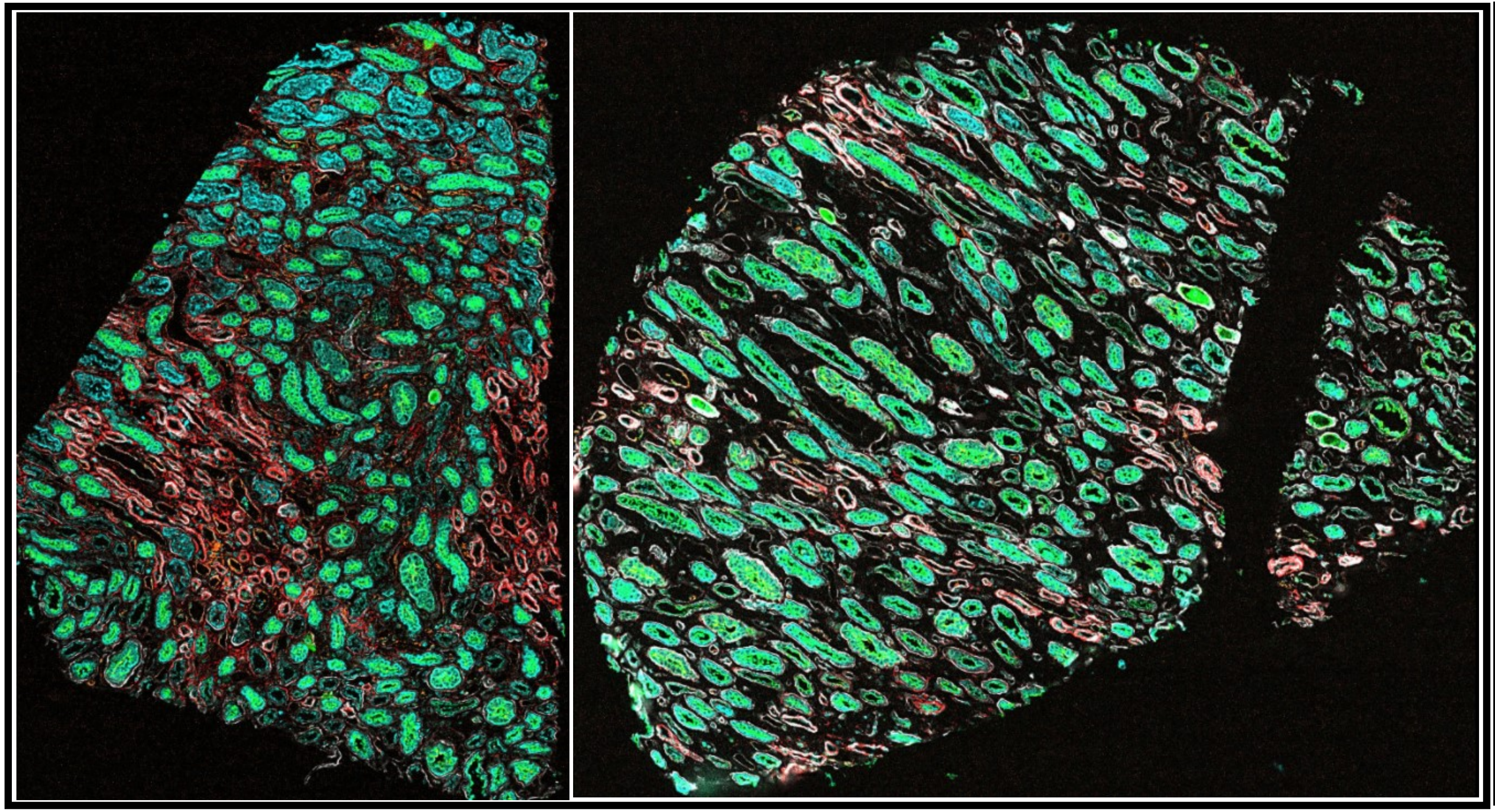

**DKDIIA-B**

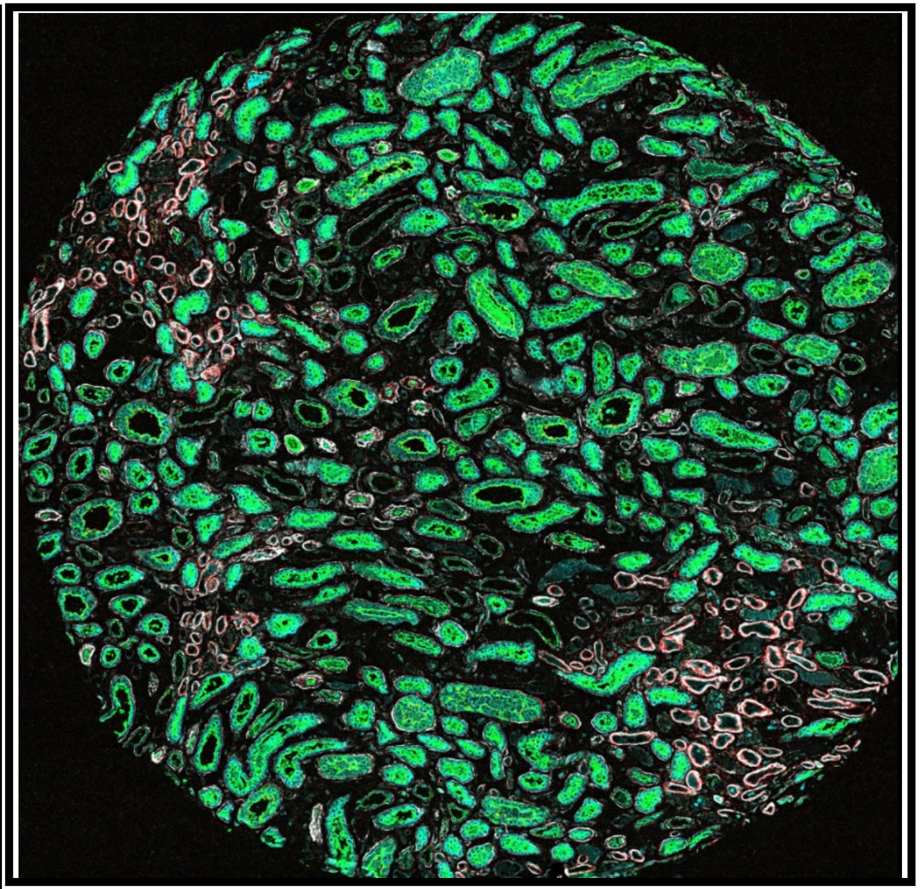
